# Natural and human-mediated drivers of microevolution in Neotropical palms: a historical genomics approach

**DOI:** 10.1101/2022.04.08.486529

**Authors:** Louise Brousseau, Sylvain Santoni, Audrey Weber, Guillaume Odonne

## Abstract

- Microevolution in Amazonia has been far less documented than in temperate regions and still constitutes a major knowledge gap. Moreover, the actual influence of pre-Columbian societies through the Holocene remains unclear, although it could provide interesting insights on the relationship between natural and human-mediated evolutionary processes in driving microevolution.
- Because they are widespread and traditionally managed since pre-Columbian times, Neotropical palms are choice models to investigate the drivers of microevolution in Amazonia. In this study, we carried out a preliminary exploration of the genomic diversity in two pairs of congeneric palm species in French Guiana (*Astrocaryum* spp. and *Oenocarpus* spp.).
- We built upon an original sampling design, taking into account both regional climate variations and local-scale pre-Columbian occupation, and designed a new target capture kit of 20,000 molecular probes scattered across exonic regions of more than 5,000 nuclear genes in Arecoïdeae (“ARECO5000+”). Hundreds of palm libraries were sequenced through Illumina sequencing, providing a unique – high-coverage – genomic dataset in these non-model species.
- We explored patterns of genomic diversity and differentiation within and across populations, bringing state-of-the-art knowledge about the roles of climate adaptation and pre-Columbian domestication. By documenting original cases of “incipient” domestication, these pioneer results open new avenues toward a better understanding of microevolution in Amazonia and of the impact of pre-Columbian societies on present-day biodiversity patterns.

## Introduction

Evolution in Amazonia has been far less documented than in temperate regions and still constitutes a major knowledge gap. Evolutionary studies in Amazonia also largely focused on macroevolution through long-term geological periods, providing interesting insights onto speciation and extinction dynamics in relation to landscape changes associated with the Andean uplift and the formation of the Amazon river (Hoorn et al. 2010), to species dispersal abilities (B. T. Smith et al. 2014) and to biotic interactions (Onstein et al. 2018). On the contrary, less attention has been paid to the microevolution of plants and animals during the Quaternary (Cavers and Dick 2013), and even less during the Holocene. This period, which started about 11,000 years ago, experienced both a global warming subsequent to the last glacial period resulting in the present climate, and the establishment of dense networks small-scale human societies subsequent to the human colonization of South America in late Pleistocene (Llamas et al. 2016; Rothhammer and Dillehay 2009). A better understanding of the drivers of plant and animal evolution during the Holocene could thus bring interesting insights on the interactions between natural and human-mediated evolutionary processes and their role in shaping present-day patterns of biodiversity.

Neotropical palms of the Arecoideae subfamily are choice models to investigate the drivers of microevolution in Amazonia. Dated phylogenies revealed that Arecoideae crown node diversified in mid-Cretaceous in South America, suggesting a Neotropical origin (Baker and Couvreur 2013). Today, Neotropical palms are widespread in Amazonia; Arecaceae standing at the second position of the most abundant plant families in forest tree inventories with some “hyperdominant” species (ter Steege et al. 2013). Neotropical palms also constitute a key resource to human societies since pre-Columbian times, whose multiple uses – including subsistence food and building materials – have been extensively documented at least since the 1850’s (Wallace 1853; N. Smith 2015). Surprisingly, they are also particularly abundant at archaeological sites with evidence of long-term human occupation, and some species (in particular *Astrocaryum* spp. and *Oenocarpus* spp.) are commonly credited as ‘biological indicators’ of ancient settlements (Odonne et al. 2019).

This is notably the case at the so-called “ring ditches” that are abundant in interfluvial terra-firme forests in lowland Amazonia, from Bolivia (John F Carson et al. 2015) to French Guiana (Petitjean Roget 1991). These pre-Columbian earthworks provide valuable evidence of anthropogenic disturbances, with belowground enrichments in pre-Columbian ceramics, organic matter and charcoals characterizing “anthropogenic soils” that are commonly attributed to slash-and-burn agriculture (Bodin et al. 2020; Brancier et al. 2014). Even if the actual magnitude of land use at these sites remain controversial – with hypotheses varying from extensive forest clearance to the exploitation of naturally open savanna (Watling et al. 2017; John Francis Carson et al. 2014) – there is no more doubt that pre-Columbian societies have strongly impacted forest landscapes, thus contrasting with previous depictions of Amazonia as an undisturbed, primary forest (Levis et al. 2017).

These repeated and converging observations feed the long-standing hypothesis that many plant species, including Neotropical palms, may have been “incipiently-” or “semi-” domesticated through pre-Columbian times (Clement 1999; Clement et al. 2010, 2006). Whether pre-Columbian societies have actually shaped patterns of palm diversity through domestication, and how human-mediated processes interacted with natural ones, remain however largely unknown. This knowledge gap is primarily underpinned by a profound lack of observational genomic data in non-model Neotropical palms that we intend to fill in this study.

We carried out a preliminary exploration of the genomic diversity in two pairs of congeneric Neotropical palms in coastal French Guiana: *Astrocaryum paramaca* and *A. sciophilum* on one side, *Oenocarpus bacaba* and *O. bataua* on the other side. We sampled populations following an original sampling design taking into account both regional and local-scale evolutionary processes related to climate variations and pre-Columbian occupation, respectively. At the regional scale, we sampled populations in study sites with contrasted rainfall to catch climate adaptation. At the local scale, we sampled palms both on the top of ring ditches and in the surrounding forests to catch the impact of domestication associated with local anthropogenic disturbances. We designed a new capture kit composed of 20,000 molecular probes to target exonic regions within thousands of nuclear genes, and carried out an extensive targeted-capture experiment and next-generation sequencing (NGS) of hundreds of libraries.

Here, we introduce this new capture kit (“ARECO5000+”) and provide new NGS data on the state and distribution of genomic diversity in Neotropical palms in coastal French Guiana. We investigate intra-specific patterns of diversity and differentiation driven by both neutral evolutionary processes (gene flow and admixture) and selective ones, disentangling the roles of (natural) climate adaptation and of (human-mediated) pre-Columbian domestication that operated at these two spatial scales.

## Materials and Methods

### 1. Palm species

We focused on four palm species that belong to the Arecoideae sub-family and that are widespread in forest landscapes of French Guiana: two species of the genus *Astrocaryum (A. paramaca* and *A. sciophilum*) and two species of the genus *Oenocarpus* (*O. bacaba* and *O. bataua). Oenocarpus bacaba* and *O. bataua* are pan-Amazonian species that are broadly distributed from Colombia to north Brazil. *Oenocarpus bataua* stands at the 7^th^ position amongst the 20 most abundant species in the Amazonian tree flora, while *O. bacaba* stands at the 33^th^ position (ter Steege et al. 2013). The distribution of *A. paramaca* and *A. sciophilum* is restricted to northeast Amazonia (Suriname, French Guiana and Brazil, GBIF Secretariat: GBIF Backbone Taxonomy. https://doi.org/10.15468/39omei). Despite this, *A. sciophilum* stands at the 51^th^ position amongst the most abundant species in forest tree inventories. Moreover, *O. bacaba* and *A. sciophilum* are two biological indicators of pre-Columbian settlements in French Guiana, as their abundance is significantly higher in forest plots with evidence of pre-Columbian occupation than in natural plots (Odonne et al. 2019). Indeed, *Astrocaryum* spp. and *Oenocarpus* spp. are major sources of food and of building and handicraft materials in Amazonia (Sosnowska et al. 2015; Cummings and Read 2016). Their leaves are traditionally used in basketry and to build houses (see for example maloca’s walls and roof, García et al. 2015; Davy 2007; Fadiman 2008; Coomes 2004; Gutierrez 2020; Kahn 1993), while their fruits are used to make oil (*e.g*. ‘seje’ oil) and beverages (*e.g*. so called ‘comou’ and ‘wassaï’ in French Guiana and ‘bacaba wine’ in Northwest Amazonia, Cunha et al. 2019; Castro et al. 2014; Puerari et al. 2015).

### 2. Sampling strategy

Palm populations of the four palms species were sampled in five study sites in the coastal French Guiana with different rainfall ranging from about 2400 to 3500 mm.year^-1^ (**Figure 1**): three archaeological sites with ring ditches (SPA, MC87 and NOU), plus two additional undisturbed sites without evidence of pre-Columbian occupation (LAU and PAR). In the three study sites with ring ditches, we sampled palm populations both at the top of the ring ditches and in the surrounding (undisturbed) forest (< 5 km from the ring ditch), **Table 1**. When the species were present, we collected a piece of leaf for 20 to 100 adult palms per species in each sampling site and local condition (pre-Columbian occupation “+” and “–”).

**Figure 1.**
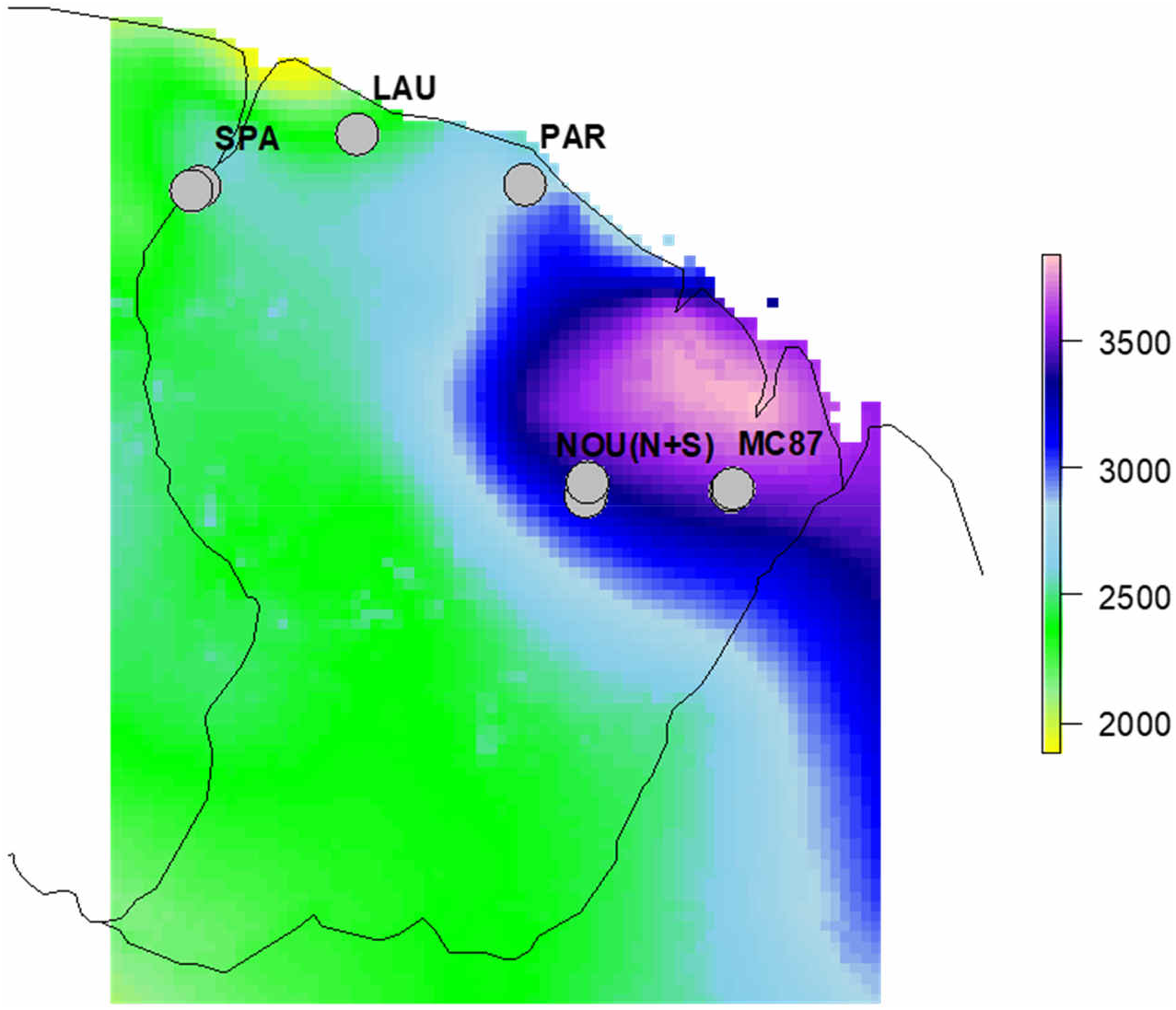
Localization of the study sites and annual precipitation (in mm.year^-1^) in French Guiana. Current precipitations were extracted from WorldClim version 1 using R (libraires “`raster`” and “`sp`”) and are representative of the period 1960-1990 (http://www.worldclim.com/version1).

**Table 1.** Localization of populations in the different sampling sites and local conditions of pre-Columbian occupation. “+” refers the top of the ring ditch with evidence of pre-Columbian occupation, and “–” refers the surrounding sampling forest area without evidence of pre-Columbian occupation. “Nouragues North” and “Nouragues South” indicate the localization of two ring ditches in the north and in the south of the Nouragues Ecological research station, respectively. Coordinates are provided in decimal degrees (WGS84).

| Study site | Site ID | Local condition | Latitude (N) | Longitude (E) | Rainfall (mm.year<sup>-1</sup>) |
| --- | --- | --- | --- | --- | --- |
| Sparouine | SPA | – | 5.272861999 | -54.22715197 | 2529 |
|  | SPA | + | 5.253569037 | -54.253331 | 2491 |
| Laussat | LAU | – | 5.479205009 | -53.59530601 | 2391 |
| Paracou | PAR | – | 5.279138032 | -52.92425998 | 2818 |
| Nouragues South | NOU_S | – | 4.038217971 | -52.68241298 | 3246 |
|  | NOU_S | + | 4.029483022 | -52.681869 | 3246 |
| Nouragues North | NOU_N | – | 4.081238005 | -52.67338903 | 3299 |
|  | NOU_N | + | 4.079981977 | -52.67142003 | 3299 |
| Montagne Couronnée 87 | MC87 | – | 4.053646959 | -52.09517601 | 3517 |
|  | MC87 | + | 4.05699201 | -52.086698 | 3517 |

### 3. Probes design, targeted capture and sequencing

We took advantage of the availability of the chromosome-level reference genome for the oil palm (*Elaeis guineensis*, Arecaceae, Arecoideae) to design a custom of capture kit of 20,000 molecular probes located within 5,209 orthologous nuclear genes scattered across all 16 chromosomes (“ARECO5000+”). Details on probes design is provided in **Supplementary material 1**. Probe sequences are available upon request from the corresponding author. The probes were then synthetized by Daicel Arbor Biosciences (myBaits Custom DNA-Seq©). Molecular experiments were further performed at the AGAP genotyping platform (Montpellier, France). Plant DNA was extracted from 15 mg of dried leaves. Ten to fifteen best-quality DNAs per population were retained for library preparation, targeted-capture and sequencing, resulting in the sampling design described in **Table 2** and totaling 405 libraries (one per palm). Genomic library preparation for multiplexed individuals and targeted-capture by bait hybridization followed published protocols (Rohland and Reich 2012; Mascher et al. 2013) with some modifications. The molecular protocol is provided in **Supplementary material 2**. One genomic library was prepared for each genotype, indexed, and enriched using the myBaits Custom DNA-Seq© kit “ARECO5000+”. 48 barcoded libraries were then equally mixed before sequencing. Libraries of *Oenocarpus bataua* were sequenced (paired-end, 2 × 150bp) on an Illumina MiSeq at the AGAP genotyping platform (Montpellier, France), while libraries of *Astrocaryum* spp. and *Oenocarpus bacaba* were sequenced by the Get-PlaGe core facility (GenoToul platform, INRA Toulouse, France http://www.genotoul.fr) on an Illumina HiSeq3000. Sequence data are available on EMBL-ENA under accession PRJEB51800.

**Table 2.**
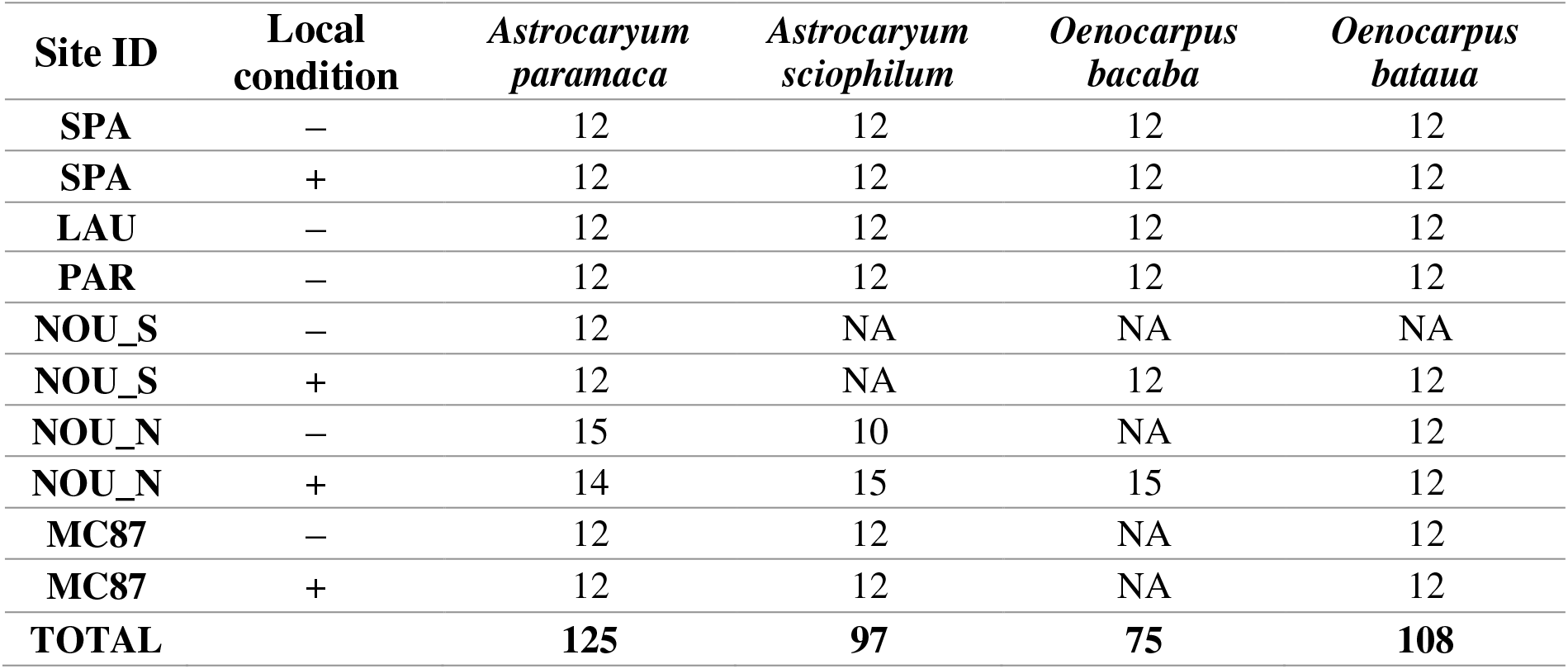
Sampling design for library preparation and sequencing. “NA” indicates that the species was absent at this sites and/or local condition.

### 4. Bioinformatics

#### Reads demultiplexing and pre-processing

Reads were demultiplexed based on their 6 nucleotide barcodes with `fastq-multx` of the `ea-utils-2.7` suit of tools (Aronesty 2013) with options `–b –x –m 0 –d 1`, and split into individual .fastq files (two files per library, corresponding to “R1” and “R2” paired reads). Read sequences that failed to pass the chastity filter (identified “Y” in the “@” line in CASAVA 1.8) were discarded using the command:

~~~
zcat $lib_name.R1.fastq.gz | grep -A 3 ′^@.* [^:]*:N: [^:]*:′|
grep -v “^--$” > $lib_name_pf.R1.fastq.
~~~

Reads were further cleaned using `FASTX-Toolkit (RRID:SCR_005534) v. 0.0.14` and `sickle-1.33`, as follow:

- Trimming with `fastx_trimmer` with option `–Q 33 –f 10 –l 140`, resulting in reads of 140-nucleotide long.
- Masking of individual bases of low quality (phred-score < 25) with `fastq_masker` with options `–q 25 –Q 33`.
- Cutting of 3’-end and 5’-end and discarding of orphan reads with `sickle-1.33` (Joshi and Fass 2011) with options `pe –l 100` (*i.e*. default quality threshold of 20 and length threshold of 100bp).

Once cleaned, reads were quality controlled with `FastQC-0.11.5` (Andrews 2010).

#### Mapping

Cleaned reads of *Astrocaryum* spp. and *Oenocarpus* spp. libraries were separately mapped against *Elaeis guineensis* reference genome (EG5) using `bwa` (H. Li and Durbin 2010, 2009): the reference was indexed with the command `bwa index` and paired-reads were mapped using `bwa mem` with option `–t 10`.

Mapping were post-processed using `samtools-1.9` (H. Li 2011; H. Li et al. 2009). Multiple mapped reads were discarded from .bam alignments using `samtools view` with the command:

~~~
samtools view -h mapping.bam | awk ‘$17 !~ /XA:/ || $1 ~ /^@/’
|    /usr/local/samtools-1.9/bin/samtools     view    -bS   >
mapping_unique.bam
~~~

Alignments were then sorted and indexed using `samtools sort` and `samtools index`, respectively. Mapping statistics were computed using `samtools stats`, `samtools flagstat` and `samtools bedcov`. Sequencing and mapping statistics are provided in **Supplementary materials 3 and 4**.

Following a first mapping against *Elaeis guineensis* reference genome, the reference was updated by editing two genus-specific consensus (one for *Astrocaryum* spp. and one for *Oenocarpus* spp.) to take into account the genomic divergence between *Elaeis guinensis* and the genera under study. This step aimed to avoid detecting polymorphisms caused by inter-genus variation in the following. This was achieved by calling variants using `samtools mpileup` with options `–uf` for each *Astrocaryum* spp. and *Oenocarpus* spp. alignments, respectively. The two consensus references were edited with `bcftools index` (v.1.3) and `bcftools consensus` (Danecek et al. 2021). Then, the libraries of each species (*Astrocaryum paramaca, A. sciophilum, Oenocarpus bacaba* and *O. bataua*) were mapped against the corresponding genus-specific consensus reference following the mapping procedure described above.

#### SNP and genotype calling

Variants were called using `samtools-1.9` and `bcftools-1.3`, as follow:

- Indexing of genus-specific consensus with `samtools faidx`;
- Variant calling with `samtools mpileup` with options `–g –t DP` followed by `bcftools call` with options `–vc`.

The resulting .vcf file was filtered with `grep` to discard monomorphic sites, indels, and variants detected on unassembled scaffolds or chloroplast to retain nuclear single-nucleotide polymorphisms (SNPs) only. SNPs were post-filtered with `R version 3.5.1` (R Core Team 2021) to retain SNPs and genotypes with a quality threshold of 20, and a per-library coverage threshold of 6X. SNPs and genotypes that did not meet the criteria were considered as being missing (“NA”), and SNPs that became monomorphic or contained more than 20% of missing values were discarded. This resulted in 178,358 and 23,252 high-quality SNPs in *Astrocaryum* spp. and *Oenocarpus* spp., respectively. Last, palm samples with more than 20% of missing values were discarded. The list of samples retained for genetic analyses is provided in **Supplementary material 5**.

### 5. Population genetics

Genome-wide patterns of genetic structure and diversity were computed on a random subset of 5,000 SNPs in `R version 3.5.1` (“`sample`”command with argument `replace = F`). Genotype tables were converted from tabular (data frame) to `genind` objects using the function `df2genind` (package `adegenet`).

#### Genetic differentiation

Pairwise genetic differentiation (*F_ST_*) was estimated both between congeneric species and within each species. Weir and Cockerham’s *F_ST_* were estimated using the function `genet.dist` (package `hierfstat`) with argument: `method=“WC84”`.

#### Individual ancestry

Genome-wide genetic structure was assessed for each species by estimating individual ancestry (Frichot et al. 2014) with the function `snmf` (package `LEA`) with arguments: `ploidy = 2, entropy = TRUE and CPU = 8`. The genetic structure was explored from `K=1` to `K=10` ancestral populations (*A. paramaca*), to `K=8` (*A. sciophilum*), to `K=6` (*O. bacaba*) or to `K=9` (*O. bataua*). The most probable number of clusters was retained based on cross-entropy criterion (lower values indicating better fit to the predictions) and each palm sample was assigned to a given group with an ancestry threshold of 0.5.

#### Isolation-by-distance

Isolation-by-distance (i.e. the correlation between genetic and geographic matrices across populations within each species) was tested through Mantel tests. Genetic distances were estimated with the function `dist.genpop` (package `adegenet`) with argument `method = 3` (*i.e*. coancestry coefficient or Reynolds’ distance, Jombart 2008; Jombart and Ahmed 2011). Geographic distances (euclidean) were computed from geographic coordinates using the function `dist` with default arguments. Mantel tests between geographic and genetic distance matrices were computed with the function `mantel.randtest` (package `ade4`, Mantel 1967; Thioulouse et al. 1997).

#### Admixture tests

Admixture between populations was evaluated through *f_3_* admixture tests for all combinations of three populations (Patterson et al. 2012; Peter 2016; Reich et al. 2009): *f_3_*(*P_X_* ;*P_A_,P_B_*), where *P_X_* is the target population and *P_A_* and *P_B_* are two source populations. Admixture tests were encoded with `R version 3.5.1`. *f_3_* statistic was estimated following the equation by Reich et al. (2009):

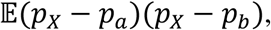

where *p_x_, p_a_* and *p_b_* are the allele frequencies in the populations *P_X_*, *P_A_* and *P_B_*, respectively, and 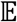 the expected value across all SNPs. Negative values indicate admixture between A and B. The significance of *f_3_* was assessed by bootstrap (function `sample` with argument `replace = T`) through 500 iterations. Z-scores were converted into p-values (left-tailed) with the function `pnorm`.

#### Intra-population diversity

Observed and expected heterozygosity within each species and population (*Ho* and *He*, respectively) were estimated with `adegenet` from `genind` objects. `Genind` objects containing multiple populations were split into separate populations using the function `seppop` and summary statistics were exported with the function `summary`. Whether *Ho* was higher than *He* was assessed for each population through non-parametric Wilcoxon tests (because of non-homogeneity of variances between *Ho* and *He*) with the function `wilcox.test`. Inbreeding coefficients (*Fis*) were estimated following the formula: 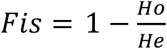. Hardy Weinberg equilibrium was tested for each SNP through Chi-squared tests with the function `hw.test` (package `pegas`), and the proportion of SNPs showing significant departure from HW equilibrium at a 1% threshold was assessed.

#### Selection test

We carried out a genome-wide scan for selective differentiation within each species using `BayPass v.2.1`. Genotype data were converted into allele counts and `BayPass` core model was run a first time using observed SNP counts as input. The number of populations in each dataset was specified with option `–npop` and MCMC parameters were specified with arguments: `–nthreads 8 –thin 15 –burnin 2500 –npilot 15 – pilotlength 5000`. The estimated covariance matrix (*Ω*), which provides an accurate estimate of genome-wide allele frequency covariance across populations, was converted into a correlation matrix in `R version 3.5.1` using the function `cov2cor`. The singular values of the covariance matrix were decomposed using `svd` and left singular vectors (U) were extracted to plot the genome-wide structure across populations. Pseudo-observed data (POD) of 10,000 SNPs (for *Oenocarpus* spp.) or 100,000 SNPs (for *Astrocaryum* spp.) were simulated with the `simulate.baypass` function (`BayPass` miscellaneous R function), specifying the total number of allele counts for each SNP (*i.e*. sample size) and the estimated hyperparameters *a_π_* and *b_π_* (*i.e*. posterior mean and standard deviation of the Beta distribution corresponding to the mean reference allele frequency, ~ancestral reference allele frequency) as input. These POD correspond to a simulated set of SNP counts that take into account the genome-wide (neutral) genetic structure empirically inferred from observed SNP counts. The core model was run again using POD as input under the same parameters to calibrate SNP-specific statistics (standardized allele frequencies *α^std^, XtX* and derived statistics) under neutral hypothesis. `BayPass` outputs were post-processed with `R version 3.5.1`. The robustness of the core model was assessed by evaluating the FMD distance (Förstner and Moonen 2003) between covariance matrices estimated on observed data (*Ω^OBS^*) and on simulated data (*Ω^POD^*) using `fmd.dist` function (`BayPass` miscellaneous R function). SNP-specific departures from genome-wide (*i.e*. neutral) differentiation were detected based on the joint distribution of:

- SNP-specific *XtX_i_* (Günther and Coop 2013) estimated by `BayPass` (which describes the differentiation across all populations for each SNP *i* explicitly corrected for genome-wide covariance, whatever their regional and local environmental conditions), and
- two additional statistics (*δ_i_* and Δ_*i*_) derived from estimated standardized allele frequencies (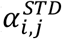, which describes allele frequencies for each SNP *i* within each population *j* explicitly corrected for the genome-wide covariance matrix) and designed to quantify variations in *α^STD^* driven by climate adaptation and pre-Columbian domestication, respectively.

Because *α^std^* are unbounded, *α^std^* were rescaled between 0 and 1 through min-max normalization:

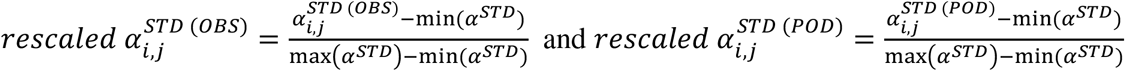

where:

- 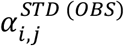 and 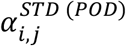 are the standardized allele frequency at SNP *i* in population *j* estimated from observed and simulated data respectively ;
- min(*α^STD^*) = min(*α*^*STD*(*OBS*)^, *α*^*STD* (*POD*)^);
- max(*α^STD^*) = max(*α*^*STD*(*OBS*)^, *α*^*STD*(*POD*)^).

The statistic *δ* quantifies clines in *α^STD^* along the regional precipitation gradient:

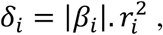

where |*β_i_*| is the absolute regression coefficient and 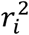 is the r-squared of the linear model between allele frequency at SNP *i* and precipitation: 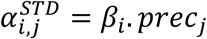, with *prec_j_* the precipitation in population *j* (scaled between 0 and 1). Linear models were fitted for each SNP with the function `lm`.

The statistic Δ quantifies absolute differences in *α^STD^* between contrasted local conditions of pre-Columbian occupation (+ *versus* −) within and across study sites (Δ_*i*_ and Δ*_i,s_*, respectively):

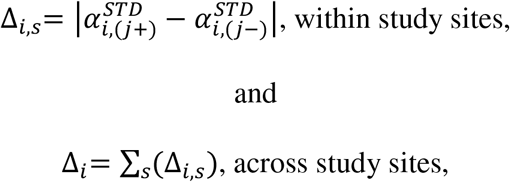

where *j*+ and *j*− refer to populations located on the ring ditch and in the surrounding natural forest for each study site *s* (*i.e*. single-site contrast), respectively, and Δ_*i,s*_ is the sum across study sites *s* (*i.e*. multi-site contrast).

*δ* and Δ statistics were separately computed on observed data 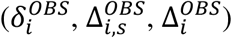 and on POD 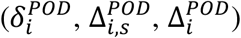. A SNP was retained as being shaped by climate adaptation if it was both over-differentiated across all populations whatever their local environment based on *XtX* at a 95% POD threshold (*i.e*. condition 1), and if it displayed clinal allele frequency variation with regional precipitation based on δ at a 95% POD threshold (*i.e*. condition 2). Similarly, a SNP was retained as being shaped by domestication if both condition 1 was met, and if it displayed local differences in allele frequencies in relation to pre-Columbian occupation based on Δ at a 95% POD threshold (*i.e*. condition 3).

| Climate adaptation SNP | Domestication SNP |
| --- | --- |
| Condition 1: $XtX_i^{OBS} > q_{(95)}(XtX_i^{POD})$ | |
| Condition 2: $\delta_i^{OBS} > q_{(95)}(\delta_i^{POD})$ | Condition 3: $\Delta_i^{OBS} > q_{(95)}(\Delta_i^{POD})$ |

where:

- 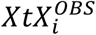 is the observed SNP-specific differentiation, and 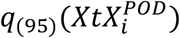 is the 95-quantile of the distribution (across all SNPs) of expected SNP-specific differentiation (estimated from POD);
- 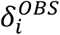 is the observed SNP-specific cline in allele frequency along the regional precipitation gradient, and 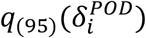 is the 95-quantile of the distribution (across all SNPs) of expected SNP-specific clines in allele frequency along the regional precipitation gradient (estimated from POD);
- 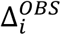 is the observed SNP-specific difference in allele frequency between local conditions of pre-Columbian occupation (+ *versus* −), and 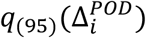 is the 95-quantile of the distribution (across all SNPs) of expected SNP-specific differences in allele frequency between local conditions of pre-Columbian occupation (estimated from POD).

#### Outlier blastn

Last, we retrieved the identity of genes containing precipitation and domestication SNPs. For each species, we extracted the sequences of the genomic regions surrounding each outlier SNP (+/− 200 bp) from consensus genomic sequences using `bedtools getfasta (bedtools-2.27.1)`. Each SNP-surrounding region was then blasted against the NCBI nucleotide database using `blastn` (`NCBI BLAST+ v. 2.10.1+`), limiting blast search by taxonomy (*Elaeis guineensis* taxid = 51953 and *Phoenix dactylifera* taxid = 42345) and returning the first good hit to optimize computation time.

## Results

### 1. Genome-wide genetic structure and differentiation

#### Genetic differentiation

At the inter-specific level, pairwise *F_ST_* across populations ranged between 0.30 and 0.55 in *Astrocaryum* spp. and between 0.23 and 0.30 in *Oenocarpus* spp. Intra-specific *F_ST_* ranged between 0.003 and 0.4 in *A. paramaca*, between 0.01 and 0.11 in *A. sciophilum*, between 0 and 0.06 in *O. bacaba* and between 0 and 0.01 in *O. bataua*, **Figure 2** and **Supplementary material 6 (Tables S5.1 & S5.2)**. In *A. paramaca*, the population originating from the undisturbed area close to the ring ditch in MC87 (i.e “MC87–”) showed a surprisingly high differentiation – ranging between 0.39 and 0.40 – with all other population of this species, while the differentiation between all other populations ranged between 0 and 0.28.

**Figure 2.**
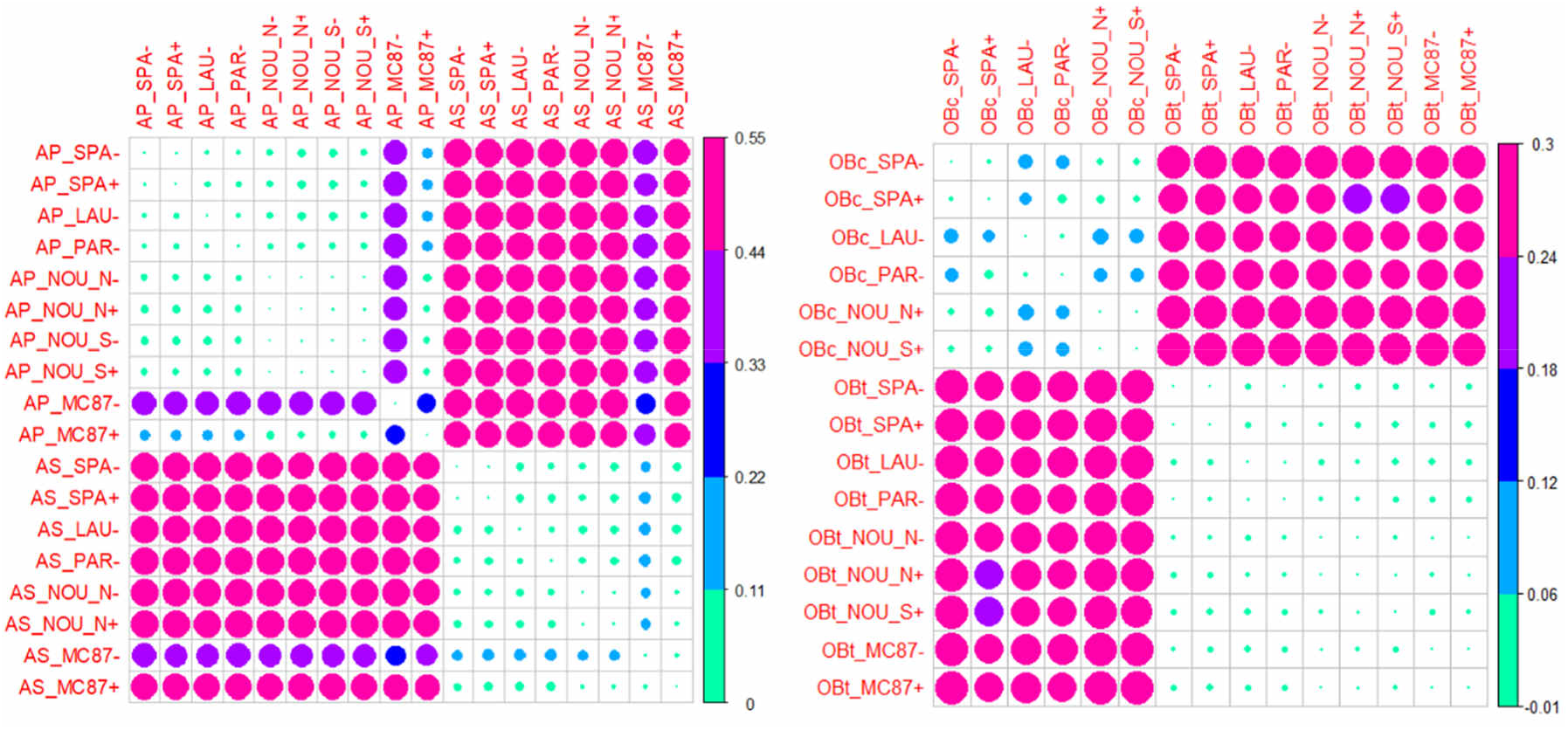
Correlation plots showing patterns of genome-wide genetic differentiation across populations of *Astrocaryum* spp. (left) and *Oenocarpus* spp. (right). “AP” refers to *A. paramaca*, “AS” to *A. sciophilum*, “OBc” to *O. bacaba* and “OBt” to *O. bataua*.

#### Individual ancestry

Cross-entropy criterion of individual ancestry was lower for K=3 and K=4 in *A. paramaca* and *A. sciophilum*, respectively, and for K=4 and K=2 in *O. bacaba* and *O.bataua*, respectively, thus indicating a genome-wide genetic structuring at a grain lower than the species level (**Supplementary material 6, Figures S6.1 & S6.2**). Within each species, the genetic structure was consistent with the west-east regional location of the palm populations, **Figure 3**.

**Figure 3.**
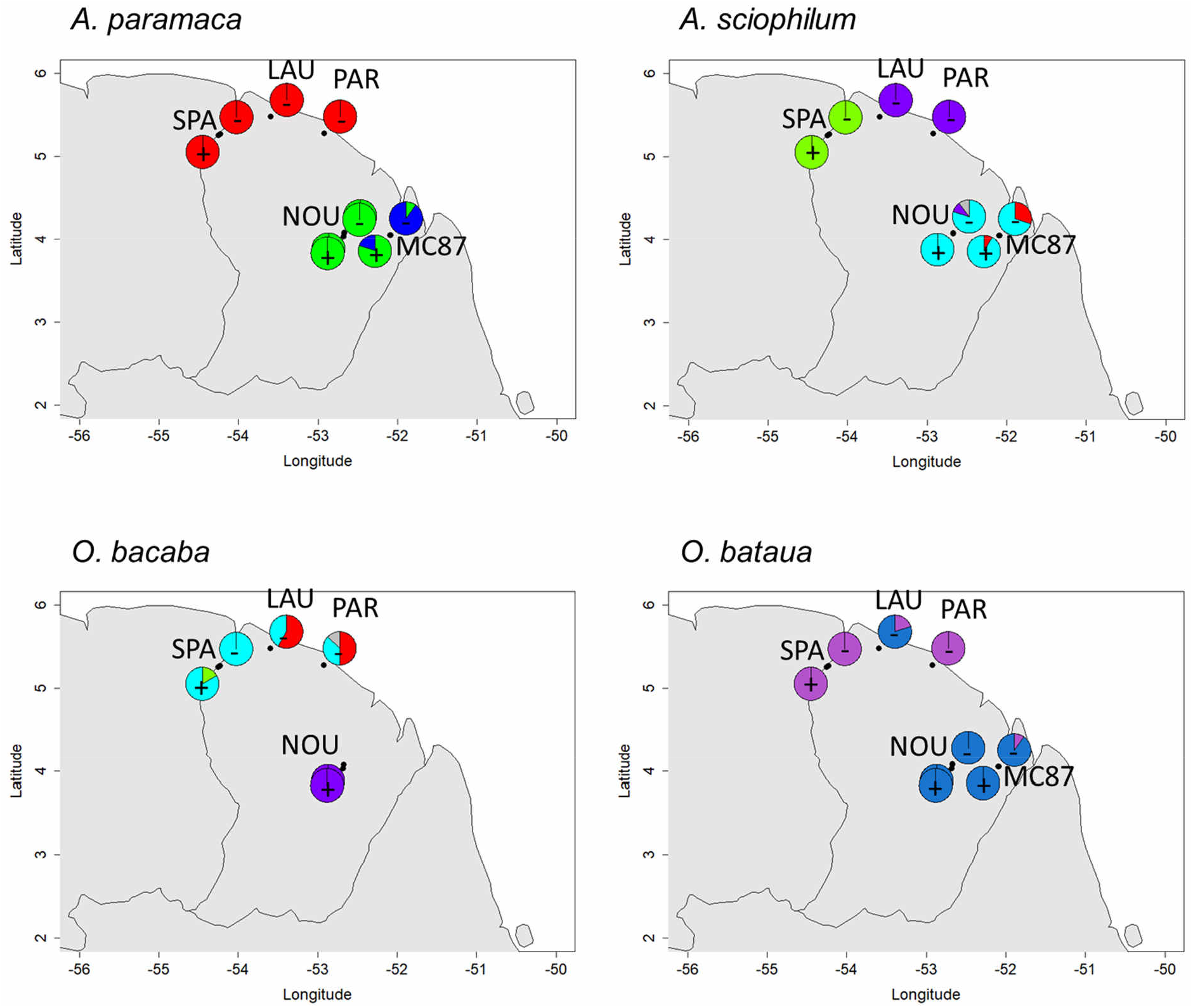
Geographic genetic structure. The pies indicate the proportion of individuals assigned to genetic each cluster within each study site and local condition of pre-Columbian occupation. Grey slices indicate ambiguous individuals that cannot be assigned to a given group with an ancestry coefficient > 0.5.

In *A. paramaca*, we detected one cluster in the west (in red) including all individuals from the sites of Sparouine (SPA), Laussat (LAU) and Paracou (PAR), and two genetic clusters in the east (sites of NOU and MC87), with all individuals associated to the blue group originating from MC87.

In *A. sciophilum*, we detected two genetic clusters in the west: one corresponding to the study site SPA (in green) and one (in purple) corresponding to the study sites LAU and PAR. In the east (NOU and MC87), most individuals were assigned to the light blue group. Few individuals originating from MC87 were assigned to a fourth group (in red). In addition, one individual was assigned to the purple group and one ambiguous individual was not assigned to any group.

In *O. bacaba*, we detected three genetic groups in the west, and one in the east. Palms from SPA were in majority assigned to the light blue group, while the populations from LAU and PAR included palms assigned to both light blue and red groups. The third group (green) corresponded to two individuals originating from the ring ditch (+) of SPA. The palms from NOU were all assigned to the fourth group (purple).

In *O. bataua*, we detected two groups with one group (purple) including palms from SPA and PAR, and another group (blue) corresponding to palms from NOU and MC87 except one palm in the latter that clustered with the purple group with an ancestry coefficient slightly above 0.5. In LAU, quite all palms clustered with the blue group except two palms clustered with the purple group.

#### Isolation-by-distance

Patterns of isolation by distance were significant in *A. sciophilum* (p-value = 0.001) and *O.bataua* (p-value = 0.002), but not in *A. paramaca* (p-value = 0.20) nor in *O. bacaba* (p-value = 0.14). In *A. paramaca*, however, IBD became significant (p-value = 0.003) after exclusion of populations from MC87 (MC87 + and MC87–), **Supplementary material 6 (Figure S6.3)**.

#### Admixture

Significantly negative *f_3_* statistics were detected at a threshold of 5% for four target populations of *Astrocaryum* spp. (*A. paramaca* from MC87 +, *A. sciophilum* from MC87 + and MC87 – and *A. paramaca* from NOU_N–), and for one target population *of Oenocarpus* spp. (*O.bacaba* in SPA +), **Supplementary material 6 (Figure S6.4)**. Populations of *A. paramaca* from MC87 + appeared to be admixed between populations of *A. paramaca* from NOU (N and S, + and –) and populations of *A. paramaca* from MC87 –, while *A. paramaca* from NOU_N– appeared to be admixed between *A. paramaca* from NOU_S – and *A. sciophilum* from NOU_N (+ and −), LAU – and/or PAR, thus suggesting patterns of inter-specific hybridization between *A. paramaca* and *A. sciophilum* in the study site of NOU. Similar patterns of inter-specific admixture also appeared in *A. sciophilum* from MC87 (+ and −), revealing admixture between *A. sciophilum* from NOU_N (+ and −) and *A. paramaca* from MC87 –. *Oenocarpus bacaba* from SPA + also appeared to be admixed between *O. bacaba* from SPA – and *O. bataua* from different sites.

#### Intra-population diversity

In *Astrocaryum* spp., observed heterozygosity (*Ho*) ranged between 0.136 (*A. sciophilum* in MC87 +) and 0.163 (*A. paramaca* in NOU_S –). In *Oenocarpus* spp., *Ho* ranged between 0.351 (*O. bacaba* SPA – and LAU – ; *O.bataua* NOU_S +, SPA – and PAR –) and 0.372 (*O. bacaba* SPA+). *Fis* was negative, indicating outcrossing. *Fis* was slightly higher in *A. paramaca* than in *A. sciophilum* although non-significant (Wilcoxon p-value = 0.1), and higher in *O. bacaba* than in *O. bataua* (Wilcoxon p-value = 0.0004), **Table 3** and **Supplementary material 6 (Figure S6.5)**. In *Astrocaryum* spp., *Fis* was higher in the populations from MC 87 where it became non-significantly negative. *Fis* also tend to be slightly higher in populations on ring ditches (+) than in surrounding natural populations (−) in *Astrocaryum* spp. and *O. bataua* while the trend was opposite in *O. bacaba*, **Table 3** and **Supplementary material 6 (Figure S6.5)**. A large proportion of SNPs shew significant departure from HW equilibrium with large variations across populations ranging from 7% to 12% in *A. paramaca*, from 0 % to 14% in *A. sciophilum*, from 18% to 27% in *O. bacaba* and from 0 to 25% in *O.bataua*.

**Table 3.** Intra-population diversity statistics (*He*, *Ho* and *Fis*) in *Astrocaryum* spp. with Wilcoxon tests for differences between *Ho* and *He* (** : p-value <0.01 ; * : p-value<0.05).

|  | *Ho* | *He* | *Fis* |
| --- | --- | --- | --- |
| ***A. paramaca*** |  |  |  |
| SPA – | 0.156 (sd=0.292) | 0.105 | -0.333 (sd=0.381) ** |
| SPA + | 0.156 (sd=0.294) | 0.105 | -0.337 (sd=0.396) ** |
| LAU – | 0.160 (sd=0.301) | 0.107 | -0.365 (sd=0.401) ** |
| PAR – | 0.157 (sd=0.294) | 0.106 | -0.345 (sd=0.389) ** |
| NOU_N – | 0.165 (sd=0.290) | 0.114 | -0.273 (sd=0.362) ** |
| NOU_N + | 0.162 (sd=0.289) | 0.112 | -0.288 (sd=0.372) ** |
| NOU_S – | 0.163 (sd=0.296) | 0.110 | -0.340 (sd=0.371) ** |
| NOU_S + | 0.162 (sd=0.292) | 0.111 | -0.312 (sd=0.371) ** |
| MC87 – | 0.140 (sd=0.285) | 0.104 | -0.141 (sd=0.587) |
| MC87 + | 0.154 (sd=0.278) | 0.128 | -0.101 (sd=0.556) |
| ***A. sciophilum*** |  |  |  |
| SPA – | 0.143 (sd=0.303) | 0.089 | -0.496 (sd=0.408) ** |
| SPA + | 0.139 (sd=0.289) | 0.092 | -0.384 (sd=0.407) * |
| LAU – | 0.137 (sd=0.287) | 0.090 | -0.403 (sd=0.400) * |
| PAR – | 0.137 (sd=0.285) | 0.091 | -0.363 (sd=0.402) * |
| NOU_N – | 0.144 (sd=0.292) | 0.094 | -0.383 (sd=0.389) ** |
| NOU_N + | 0.137 (sd=0.285) | 0.091 | -0.366 (sd=0.405) * |
| MC87 – | 0.137 (sd=0.257) | 0.142 | 0.030 (sd=0.614) * |
| MC87 + | 0.136 (sd=0.272) | 0.110 | -0.004 (sd=0.625) |

**Table 4.** Intra-population diversity statistics (*He*, *Ho* and *Fis*) in *Oenocarpus* spp. with Wilcoxon tests for differences between *Ho* and *He* (** : p-value <0.01 ; * : p-value<0.05).

|  | *Ho* | *He* | *Fis* |
| --- | --- | --- | --- |
| ***O. bacaba*** |  |  |  |
| SPA – | 0.352 (sd=0.415) | 0.205 | -0.575 (sd=0.389) ** |
| SPA + | 0.373 (sd=0.393) | 0.228 | -0.438 (sd=0.408) ** |
| LAU – | 0.351 (sd=0.381) | 0.226 | -0.427 (sd=0.436) ** |
| PAR – | 0.356 (sd=0.385) | 0.228 | -0.453 (sd=0.442) ** |
| NOU_N + | 0.353 (sd=0.411) | 0.207 | -0.535 (sd=0.400) ** |
| NOU_S + | 0.351 (sd=0.413) | 0.206 | -0.554 (sd=0.398) ** |
| ***O. bataua*** |  |  |  |
| SPA – | 0.351 (sd=0.428) | 0.198 | -0.659 (sd=0.367) ** |
| SPA + | 0.353 (sd=0.427) | 0.200 | -0.639 (sd=0.376) ** |
| LAU – | 0.352 (sd=0.430) | 0.198 | -0.700 (sd=0.347) ** |
| PAR – | 0.352 (sd=0.427) | 0.200 | -0.686 (sd=0.353) ** |
| NOU_N – | 0.358 (sd=0.427) | 0.203 | -0.644 (sd=0.375) ** |
| NOU_N + | 0.361 (sd=0.423) | 0.206 | -0.614 (sd=0.384) ** |
| NOU_S + | 0.357 (sd=0.424) | 0.204 | -0.631 (sd=0.378) ** |
| MC87 – | 0.355 (sd=0.421) | 0.204 | -0.617 (sd=0.376) ** |
| MC87 + | 0.353 (sd=0.422) | 0.202 | -0.642 (sd=0.365) ** |

**Table 5.**
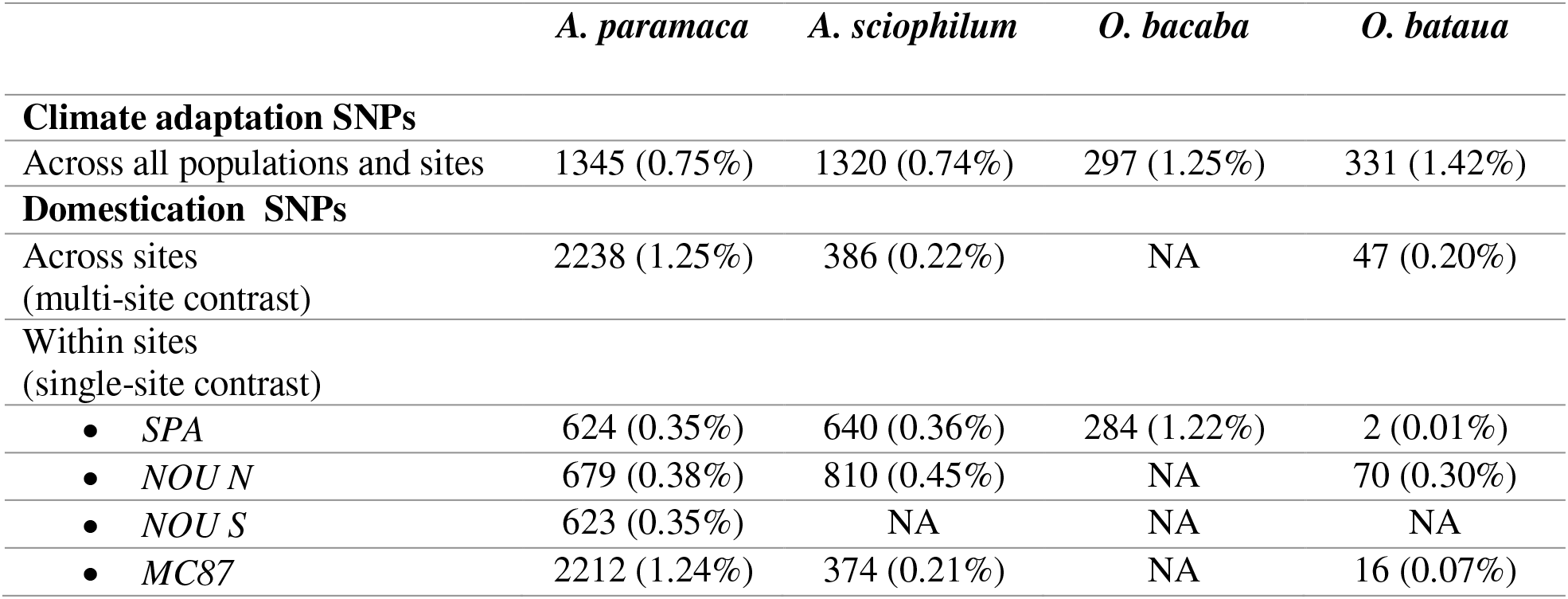
Number of over-differentiated SNPs detected in the four species in relation to climate adaptation and domestication (out of 178,358 SNPs in *Astrocaryum* spp. and 23,252 SNPs in *Oenocarpus* spp.). Because *O. bacaba* was only present on the ring ditches in NOU_N and NOU_S and was absent from MC87, the detection of domestication SNP was based on a single contrast in SPA.

|  | *A. paramaca* | *A. sciophilum* | *O. bacaba* | *O. bataua* |
| --- | --- | --- | --- | --- |
| **Climate adaptation SNPs** |  |  |  |  |
| Across all populations and sites | 1345 (0.75%) | 1320 (0.74%) | 297 (1.25%) | 331 (1.42%) |
| **Domestication SNPs** |  |  |  |  |
| Across sites<br>(multi-site contrast) | 2238 (1.25%) | 386 (0.22%) | NA | 47 (0.20%) |
| Within sites<br>(single-site contrast) |  |  |  |  |
| • SPA | 624 (0.35%) | 640 (0.36%) | 284 (1.22%) | 2 (0.01%) |
| • NOU N | 679 (0.38%) | 810 (0.45%) | NA | 70 (0.30%) |
| • NOU S | 623 (0.35%) | NA | NA | NA |
| • MC87 | 2212 (1.24%) | 374 (0.21%) | NA | 16 (0.07%) |

### 2. SNP-specific differentiation

`BayPass` core model provided accurate estimates of genome-wide genetic structure, SNP-specific differentiation (*XtX_i_*) and derived statistics designed to disentangle footprints of divergent selection caused by precipitation (*δ_i_*) at the regional scale, and pre-Columbian occupation (*i.e*. domestication) at the local scale (*Δ_i_*). Covariance matrices estimated on observed data (Ω^OBS^) were close to covariance matrices estimated on pseudo-observed data (Ω^POD^), with 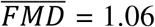 in *A. paramaca*, 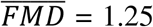 in *A. sciophilum*, 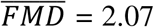 in *O. bacaba* and 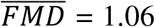 in *O. bataua*, **Supplementary material 6 (Figures S6.6 to S6.9)**. The genome-wide genetic structure estimated by `BayPass` through covariance matrices was also consistent with the genetic structure estimated by `snmf` through individual ancestry, confirming the robustness of `BayPass` to empirically infer the genomic structure across populations **Supplementary material 6 (Figures S6.6 to S6.10)**. We further detected SNPs displaying patterns of differentiation related to precipitation and/or pre-Columbian occupation based on the joint distribution of *XtX_i_* and derived statistics (*δ_i_* and *Δ_i_*), **Figure 4** and **Supplementary material 6 (Figure S6.11)**.

**Figure 4.**
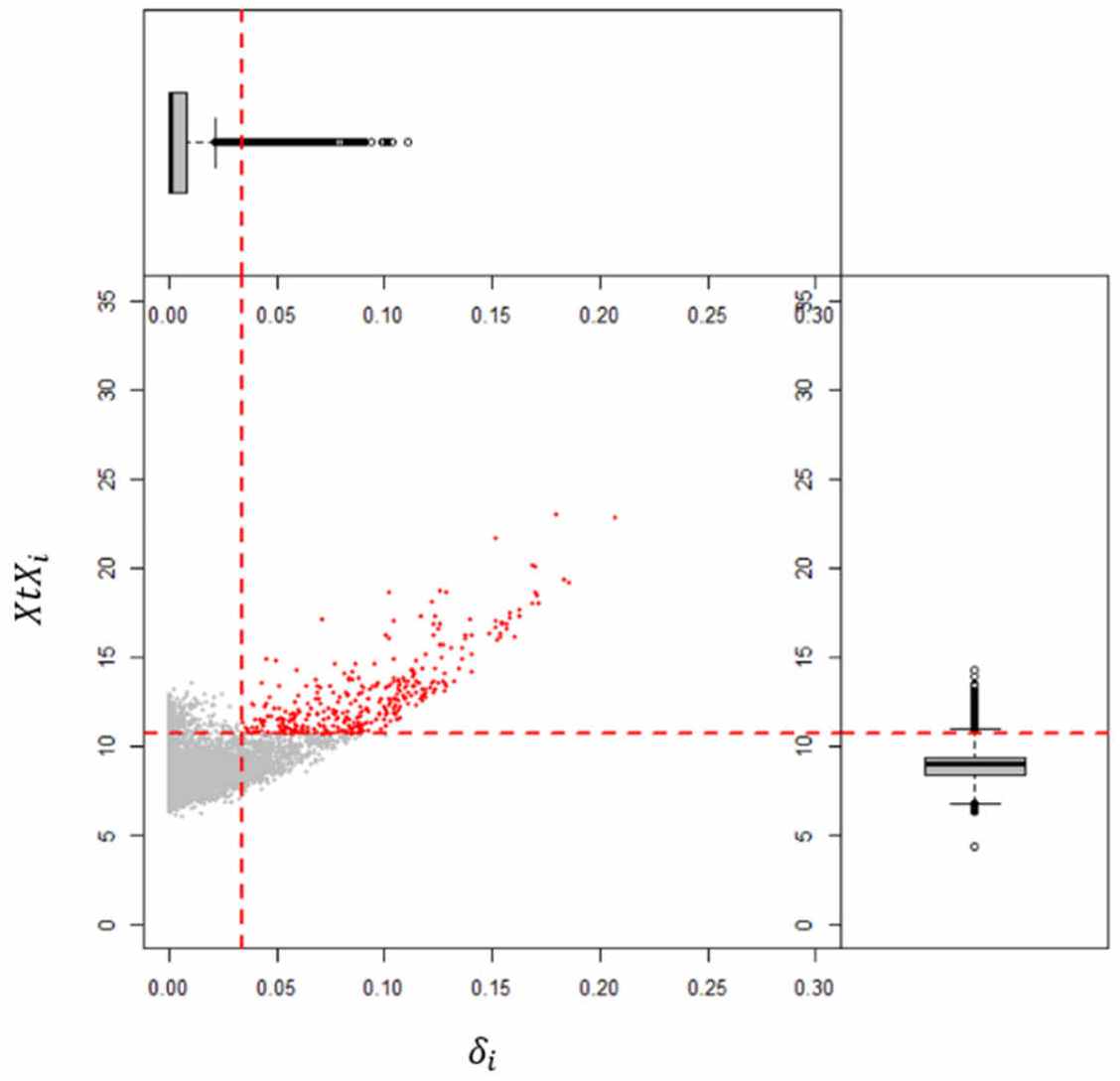
Example of SNP detection in relation to precipitation in *Oenocarpus bataua* based on the joint distribution of *XtX_i_* and *δ_i_*.

Between 0.74% (*A. sciophilum*) and 1.42% (*O. bataua*) climate adaptation SNPs were detected and between 0.20% (*O. bataua*) and 1.25% (*A. paramaca*) domestication SNPs, across all study sites. Moreover, the proportion of domestication SNPs was higher when considering all sites together (i.e. multi-site contrast) than within each site (single-site contrast), with many SNPs detected in one or two study sites only. Nevertheless, many domestication SNPs were shared between study sites: in *A. paramaca*, 1,252 domestication SNPs were detected in two sites, 285 were detected in three sites, and 28 in four sites ; in *A. sciophilum*, 564 SNPs were detected in two sites and 105 in three sites ; in *O. bataua*, only 10 SNPs were detected in two sites. The number of shared SNPs between pairs of study sites is provided in **Supplementary material 6 (Tables S6.3 to S6.6).**

The median of SNP-specific differentiation (*XtXi*) for neutral SNPs was 9.8 (Q90 = [8.6 ; 12.0]) in *A. paramaca*, 8.0 ([6.7 ; 9.9]) in *A. sciophilum*, 5.6 ([4.5 ; 8.1]) in *O. bacaba* and 8.7 ([7.6 ; 11.0]) in *O. bataua. XtXi* ranged between 9.6 ([8.2 ; 14.6]) in *O. bacaba* and 13.9 ([12.5 ; 18.1]) in *A. paramaca* for climate adaptation SNP, and between 10.48 ([8.2 ; 13.8]) in *O. bacaba* and 13.1 ([12.5 ; 16.0]) in *A. paramaca* for domestication SNPs, without differences in the extent of genetic differentiation between climate adaptation and domestication SNPs, **Figure 6**.

**Figure 6.**
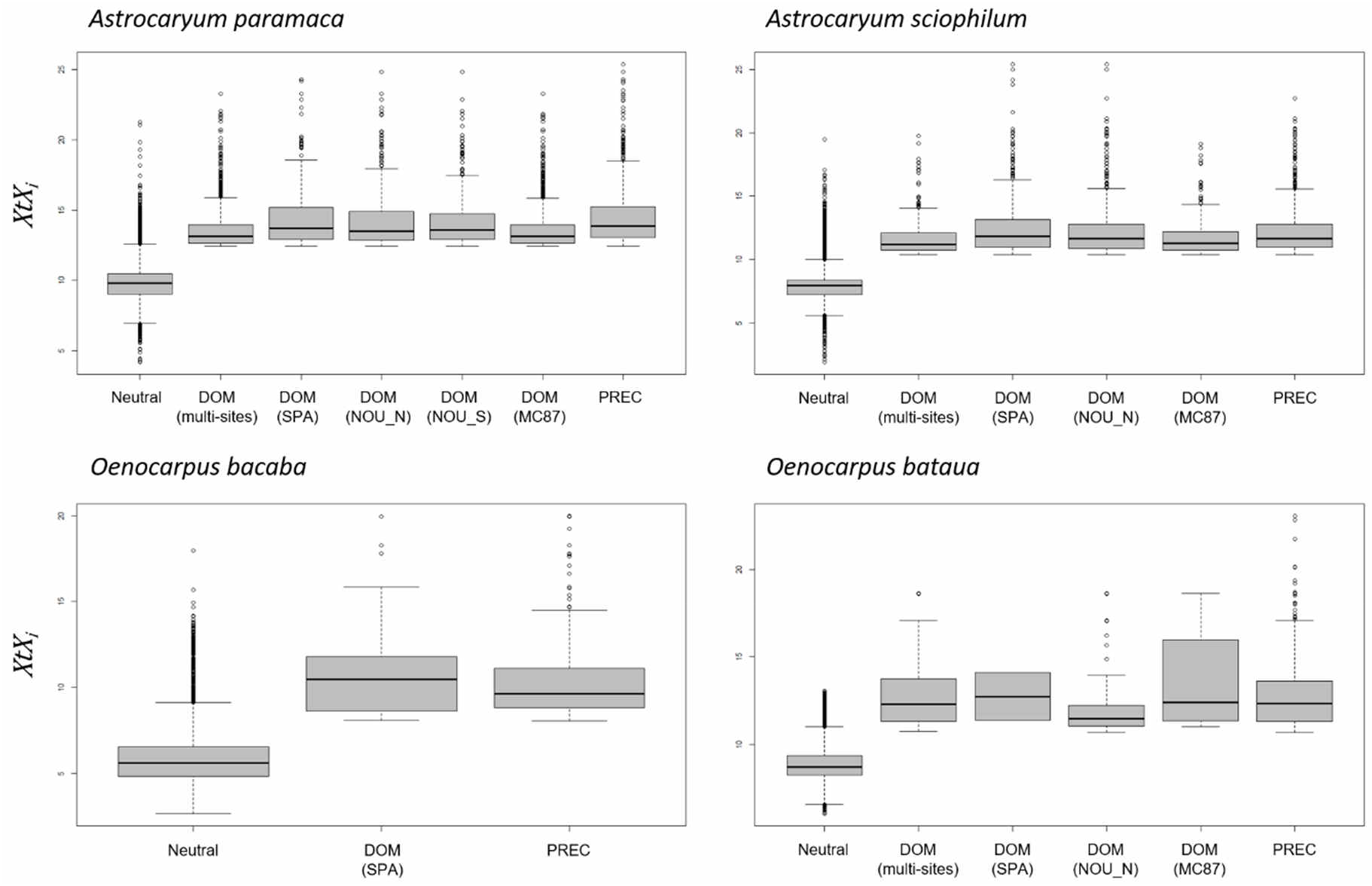
Distribution of SNP-specific differentiation (*XtX_i_*) in neutral SNPs, domestication (DOM) and climate adaptation SNPs (PREC).

Interestingly, several SNPs shew variations in allele frequency in relation to both precipitation and pre-Columbian occupation (**Supplementary material 6**, **Tables S6.3 to S6.6)**), allowing to distinguish four groups of SNPs within each species: (i) neutral SNPs, (ii) climate adaptation SNPs showing clinal variations in allele frequency in relation to regional precipitation, (iii) domestication SNPs showing contrasted allele frequency in relation to local pre-Columbian occupation, and (iv) climate and domestication SNPs showing allele frequency variations in relation to both, **Figure 7**.

**Figure 7.**
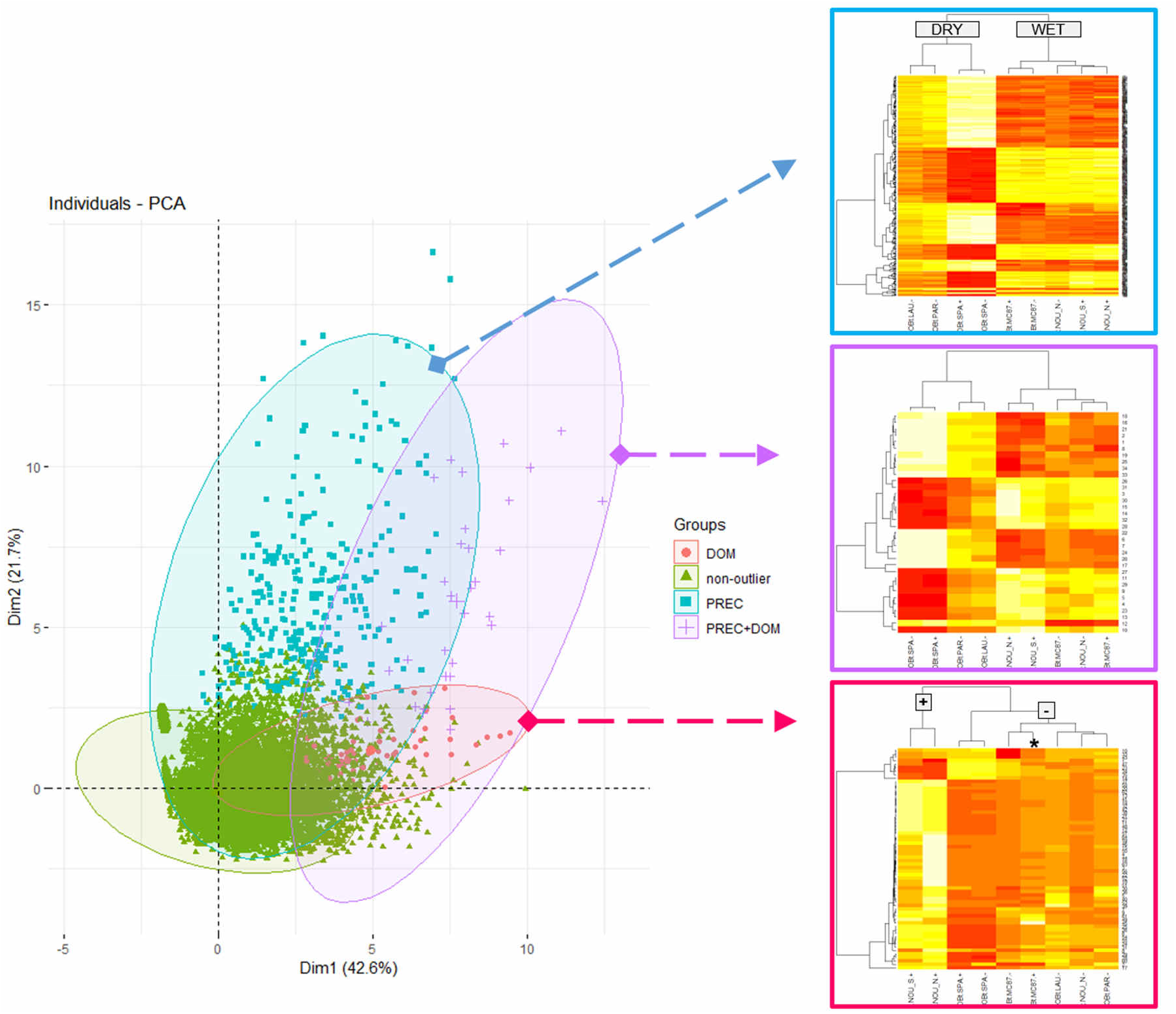
Principal Component Analysis representing the multivariate projection of SNPs based on SNP-specific estimates of *XtX_i_, δ_i_, Δ_i_* and *Δ_i,s_* in *Oenocarpus bataua*. Green: neutral SNPs (*i.e*. non-outlier); blue: climate adaptation SNPs (“PREC”) ; pink: domestication SNPs (“DOM”) ; violin: climate and domestication SNPs (“PREC+DOM”). Heatmaps on the right show standardized allele frequencies in the different populations and for the different SNPs of each group.

## Discussion

This study opens a new research framework to document microevolution in non-model Amazonian palms. We provide original observational genomic data describing the state and distribution of genomic diversity in French Guiana, and we bring new knowledge about the microevolutionary processes at play, disentangling neutral and selective processes on one side, and natural and human-mediated ones on the other.

First of all, we must recall that all the four species were not present in all study sites and local conditions of pre-Columbian occupation. This was particularly the case for *Oenocarpus* spp. that were more difficult to find and to collect in natural forest areas than on ring ditches were these species are frequent. As a result, we did not collect any *Oenocarpus* spp. in NOU_S –, while *O. bacaba* was also absent from NOU_N – and MC87 (+ and −). This is consistent with previous observations at the community level by Odonne et al. 2019 who reported an over-abundance of palms in sites with strong evidence of pre-Columbian occupation compared to natural forest landscapes. This constrained our study, as we had to deal with incomplete sampling designs. Nevertheless, the amount of population-scale NGS data acquired was largely sufficient to explore microevolutionary processes in the studied palm species, and to disentangle natural and human-mediated ones.

### Neutral microevolution driven by gene flow and admixture

Patterns of genome-wide diversity revealed a west-east genetic structuring at the regional scale, with similar trends between the four species. Genome-wide structure and differentiation are commonly driven by neutral evolutionary processes, in particular gene flow. This is comforted by significant isolation-by-distance in three species, indicating that regional patterns of genomic diversity are primarily driven by gene flow, which gradually decreases with the geographic distance between populations. We also detected intriguing patterns of genome-wide genetic structure in *Astrocaryum* spp. in the study site of MC87. In particular, the natural population (–) of *A. paramaca* revealed high levels of differentiation (about 40%) with populations from other study sites, while the population on the ring ditch was closer to populations from other study sites than to the population in the surrounding natural forest. A similar pattern was also observed in the congeneric species *A. sciophilum*, although to a lesser extent, suggesting a site-specific process probably involving inter-specific hybridization. This hypothesis was corroborated by admixture tests, which suggested inter-specific admixture between *A. paramaca* and *A. sciophilum* in the study site of MC87. Similarly, the genetic structure of *Oenocarpus bataua* revealed a site-specific structure in SPA+ corroborated by admixture tests suggesting inter-specific hybridization between *O. bacaba* and *O. bataua*. These site-specific patterns of inter-specific hybridization were neither expected nor anticipated when designing this study and carrying out the sampling. Most interestingly, *Astrocaryum* putative inter-specific hybrids were identified either as *A. paramaca* or as *A. sciophilum*, suggesting that they alternatively exhibit the morphological characteristics of one or the other species. The two putative *Oenocarpus* hybrids, on their side, were both identified as *Oenocarpus bacaba*. While there is no other species of *Oenocarpus* known in French Guiana, there is at least fifteen different species of *Astrocaryum* recorded on *GBIF* (GBIF Secretariat: GBIF Backbone Taxonomy. https://doi.org/10.15468/39omei Accessed via https://www.gbif.org/species/2738048 [30 March 2022]), raising the possibility of complex inter-specific hybridization with possible cryptic species in *Astrocaryum*. Even though we cannot conclude about the exact species involved in the pattern observed here, this preliminary observation would deserve in-depth investigation of inter-specific hybridization and introgression in the *Astrocaryum* complex.

Independently from site-specific hybridization patterns, estimates of intra-population genetic diversity revealed high heterozygosity (*H_o_*) and significantly negative *F_is_*, indicating the absence of inbreeding. These estimates are in line with previous knowledge about the breeding system in Arecoideae that are known to be monoecious and predominantly outcrossed due to protandry and/or genetic incompatibility (Henderson 1986; Eguiarte et al. 1992; Consiglio and Bourne 2001; Picanço-Rodrigues et al. 2015; Gaiotto et al. 2003; Ottewell et al. 2012). The genetic structure and diversity of populations on ring ditches was further close to populations in the surrounding natural forests, indicating the absence of pre-Columbian impacts on population genome-wide diversity and divergence. This result is not surprising, considering the geographic proximity between “+” and “−” populations within study sites (< 5 km), which are probably connected by extensive gene flow. The absence of genome-wide differentiation, however, does not preclude the existence of SNP-specific selective processes driven by adaptation and/or domestication.

### Climate adaptation and pre-Columbian domestication

Adaptation and domestication are acknowledged as important drivers of microevolution worldwide, but they have been poorly documented in lowland Amazonia. While Central and South America are known to be important centers of plant domestication, most studies have focused on major crops such as maize (Kistler et al. 2018), papaya (Chávez-Pesqueira and Núñez-Farfán 2017), manioc (Mühlen et al. 2019; Rival and McKey 2008; Alves-Pereira et al. 2020) or common bean (Rendón-Anaya et al. 2017), which provide the most obvious cases of domestication. On the opposite, studies on minor crops are scarce although they are receiving increasing attention since the last ten years (*e.g*. Brazil nut, Shepard and Ramirez 2011 ; piquiá, Alves et al. 2016 ; peach palm, Clement et al. 2017 ; treegourd, Moreira et al. 2017). For its part, the lack of interest in adaptation can be explained by the popularity of the neutral theory under which the distribution of alleles within species (Kimura 1983) or species within communities (Hubbell 2001) is assumed to be independent from fitness differences between genotypes or species, being thus driven by stochastic mutation, demography and dispersal. In hyperdiverse tropical rainforests, neutral models are powerful to predict global patterns of diversity, as illustrated in the above section, and very useful to overcome the complexity of niche differentiation modelling. Neutral and niche models are however highly complementary (Chave 2004), and the neutral theory provides an elegant neutral hypothesis to test for niche differentiation and adaptation (Leigh 2007). This is precisely the theoretical assumption of most population genetics selection tests, which allows detecting gene- or SNP-specific departures from the genome-wide, supposedly neutral, background of diversity and differentiation (Beaumont and Nichols 1996).

By detecting over-differentiated SNPs in relation to precipitation and pre-Colombian occupation, our results bring new evidence of the joint influence of climate adaptation and ancient domestication in shaping patterns of genomic diversity across multiple spatial scales. As expected, divergent selection related to adaptation and domestication affected a very small fraction of the studied SNPs (always below 1.5%). Both the proportion of SNPs and the extent of differentiation (*XtXi*) were in the same order of magnitude between climate adaptation and domestication SNPs, in spite of the different geographical scales involved (*i.e*. hundreds of kilometers between study sites with different rainfall *versus* hundreds of meters between local populations in contrasting conditions of pre-Columbian occupation). Regarding climate adaptation, the present results are in line with a previous study in the hyperdominant tree *Eperua falcata* which provided another evidence of climate adaptation along the sharp rainfall gradient in coastal French Guiana (Brousseau et al. 2021).

Climate and domestication SNPs were located within or in the neighborhood (< 200bp) of many genes whose exhaustive list is provided in **Supplementary material 7**. Although documenting the functions of these genes in *Astrocaryum* spp. and *Oenocarpus* spp. is beyond the scope of this study, we can nevertheless pick a few examples to illustrate the variety of genes involved. For example, we detected climate adaptation SNPs in several genes of major importance for plant development and response to stresses, including ABC transporters (Yazaki 2006; Do et al. 2018), leaf rust disease resistance loci (Lim et al. 2015), Apetala2-like ethylene responsive transcription factor (Licausi et al. 2013), aquaporins (G. Li et al. 2014; Maurel et al. 2015), and several peroxidases (Blokhina et al. 2003; Yoshida et al. 2003). Several domestication SNPs were detected in genes involved in lipid and sugar metabolism that could be of importance for fruit development, including phospholipase D (Sun et al. 2011; Mao et al. 2007; Pinhero et al. 2003), acyl-transferases, fructose-bisphosphate aldolase (Feng et al. 2020; Yin et al. 2012), phospholipid-transporting ATPases, and UDP-glucose 6-dehydrogenase 4.

Interestingly, many SNPs showed patterns of differentiation in relation to both climate adaptation and domestication, indicating that these two processes are closely intertwined. In other words, pre-Columbian societies had either domesticated “*in situ*” locally pre-adapted populations or had to deal with adaptation during domestication. The absence of genome-wide differentiation between palm populations on ring ditches and surrounding natural populations tends to suggest that palm populations were collected in the forest near ring ditches. Moreover, extensive gene flow between wild and domesticated populations had probably continuously homogenized populations, thus preventing genome-wide differentiation and the fixation of the so-called “domestication syndrome” in domesticated populations (see next section). Instead, SNP-specific patterns of domestication observed here provide a new evidence of the subtle influence of pre-Columbian societies and depicts what is commonly referred to as “incipient” domestication (Lins Neto et al. 2014; Casas et al. 2016; Clement 1999). Gene flow since land abandonment may also have partly erased patterns of differentiation caused by domestication (*i.e*. de-domestication, Qiu et al. 2017). Moreover, many SNPs were detected in one study site only, supporting that pre-Columbian impacts were partly site-specific. Nevertheless, many SNPs were also detected in more than one study site, indicating some common trends in the different study sites, probably caused by similar management practices. Whether observed domestication patterns result from “conscious” domestication through direct selection for desired traits or “unconscious” domestication through indirect local adaptation to anthropogenic landscapes (Gepts 2010; Engelbrecht 1916; Darlington 1973) cannot however be solved yet, and would require in-depth investigation at the phenotypic level to assess which traits have been selected for or against in palm populations established on ring ditches.

### Moving beyond the domestication syndrome paradigm

The domestication of plants is a long-term process, with multiple possible outcomes. However, most studies have focused on major crops exhibiting clear phenotypic differences with wild populations, often underpinned by a sharp – genome-wide – genetic differentiation. Studies on major crops led to the elaboration of an earlier model of domestication through which the domestication is initiated by the isolation of a small population with reduced diversity. Domesticated populations are then expected to accumulate phenotypic differences from their wild ancestors under the action of strong selection (*i.e*. artificial selection for desired traits and natural selection exerted by new environmental conditions) reinforced by inbreeding, both resulting in the fast decrease in genetic diversity and in the fixation of phenotypic differences in domesticated populations: the so-called “domestication syndrome” (Hammer 1984; Harlan et al. 1973; Allaby 2014). The domestication syndrome has been extensively documented, and quite over-emphasized (Kantar et al. 2017), leading to a narrow view of the domestication process and to a misleading dichotomous distinction between domesticated varieties with a clear domestication syndrome, and other (supposedly wild) populations. The domestication process is, however, gradual through time and variable across populations (Meyer et al. 2012; O. Smith et al. 2019; Kantar et al. 2017), and it is not necessarily expected to result in a sharp – genome-wide – differentiation, in particular in cases of domestication “*in situ”* with extensive and continuous gene flow with wild populations (Meyer and Purugganan 2013). In a broader sense, the domestication process is better depicted as the modification of the evolutionary trajectory of a population through human-mediated evolutionary processes (including human-mediated dispersal, artificial selection for useful traits and/or indirect adaptation to anthropogenic landscapes) that can outcome – in most extreme cases – in the fixation of the domestication syndrome.

Because lowland Amazonia offers a great diversity of domestication histories and cases, from “incipiently-” or “semi-” domesticated populations to stable varieties (Lins Neto et al. 2014; Clement 1999), it offers an ideal study system to reconsider the domestication process beyond the domestication syndrome paradigm. Integrating the diversity of evolutionary histories underlying domestication further opens a new perspective toward a better understanding of microevolution in Amazonia, and the present study argues in favor of a paradigm shift to better integrate “incipiently-” and “semi-” domesticated minor crops in domestication studies.

## Supplementary materials

**Supplementary material 1 (S1)**. ARECO5000+ Custom probe kit for targeted capture in Arecoideae.

**Supplementary material 2 (S2)**. Molecular methods – library preparation, capture, sequencing and adapters.

**Supplementary material 3 (S3).** Sequencing and bioinformatics (405 libraries) – supplementary figures, tables and information.

**Supplementary material 4 (S4).** Fastq quality check.

**Supplementary material 5 (S5).** List of palm samples retained for genetic analyses.

**Supplementary material 6 (S6).** Population genetics – supplementary figures and tables.

**Supplementary material 7 (S7).** Genes containing climate adaptation and domestication SNPs.

## Ethics statement

This study was carried out in compliance with the Nagoya protocol and French regulations on access to genetic resources with no direct commercial purposes. Palm samples were collected for fundamental research following declaration to the French Ministry of the Ecological Transition registered under the Internationally Recognized Certificate of Compliance ref. ABSCH-IRCC-FR-245923-1 of 23 May 2018. The authors declare no commercial use of the data generated and that no monetary benefit was derived from this study. Genomic data are made available according to shared values and principles for Open Science.

## Data and codes availability

Sequencing data (i.e. paired-end reads) are available on the European Nucleotide Archive under accession PRJEB51800. Bioinformatics codes are available on GitLab under Project ID 34990830 (https://gitlab.com/LouiseBrousseau/palm-capture-bioinformatics). Data and code remains under embargo until formal acceptance of the present article.

## Authors contribution

LB and GO imagined the study and carried out palm sampling. LB designed the capture kit, implemented the bioinformatics pipeline, and carried out bioinformatics and population genetics. SS and AW carried out the molecular experiments (gDNA extraction and purification, library preparation, targeted capture and sequencing of *O. bataua* librairies). All authors wrote and approved the article.

## Acknowledgements

This study has been conducted as part of the research program *PalmOmix* coordinated by L. Brousseau. *PalmOmix* (ID 1702-01) was publicly funded by the French National Research Agency (Agence Nationale de la Recherche, ANR) under the *“Investissement d’avenir”*program with reference ANR-10-LABX-001-01 LabEx Agro and coordinated by Agropolis Foundation under the frame of I-SITE MUSE (ANR-16-IDEX-0006). Sampling in the Nouragues research Station has been supported by the LongTIme project coordinated by G. Odonne and J-F. Molino and funded by an *“Investissement d’avenir”* grant from the French National Research Agency (Agence Nationale de la Recherche, ANR) with reference CEBA: ANR-10-LABX-25-01.

We are grateful to the Nouragues Ecological Research Station (CNRS) and the Réserve Naturelle des Nouragues, to the Paracou Research Station (CIRAD) and to the National Forests Office (ONF) in French Guiana for access to the study sites. We are also grateful to GenoToul (“GeT” and bioinformatics) facilities for next-generation sequencing of *Astrocaryum* spp. and *O. bacaba* libraries, and for access to computing facilities to carry out bioinformatics analyses. We thank Thierry Joët, Stéphane Dussert and James Tregear for their help in pre-filtering candidate proteins (“bottom-up approach”) to design the custom capture kit. We also thank T. Joët for his help in palm sampling and Anaïs Prud’homme who explored *O. bataua* NGS data as part of her internship at UMR DIADE under the supervision of L. Brousseau.

