## Supplementary material 1 for "Natural and human-mediated drivers of microevolution in Neotropical palms: a historical genomics approach"

### Supplementary Material 1 (S1)

### ARECO5000+ Custom probe kit for targeted capture in Arecoideae

Louise Brousseau

**Foreword.**

This technical note briefly introduces a new custom capture kit of 20,000 molecular probes for targeted-enrichment of 5,209 genes in Arecaceae as part of the article by Brousseau *et al.* entitled "Natural and human-mediated drivers of microevolution in Neotropical palms: a historical genomics approach". This custom capture kit has been designed by L. Brousseau and experimentally synthesized by Daicel Arbor Biosciences (myBaits Custom DNA-Seq©). This supplementary material describes the computing procedure implemented to design the probes (nucleotide sequences) and some useful technical information about their validation and success rates. This kit was preliminary intended to target orthologous nuclear genes across Arecoideae, but recent tests on more distant genera (*Mauritia sp.*, *Phoenix sp.*) tend to indicate that it could be used in any Arecaceae species, although this is beyond the scope of the present study. Sequences of the molecular probes are available upon request from Louise Brousseau.

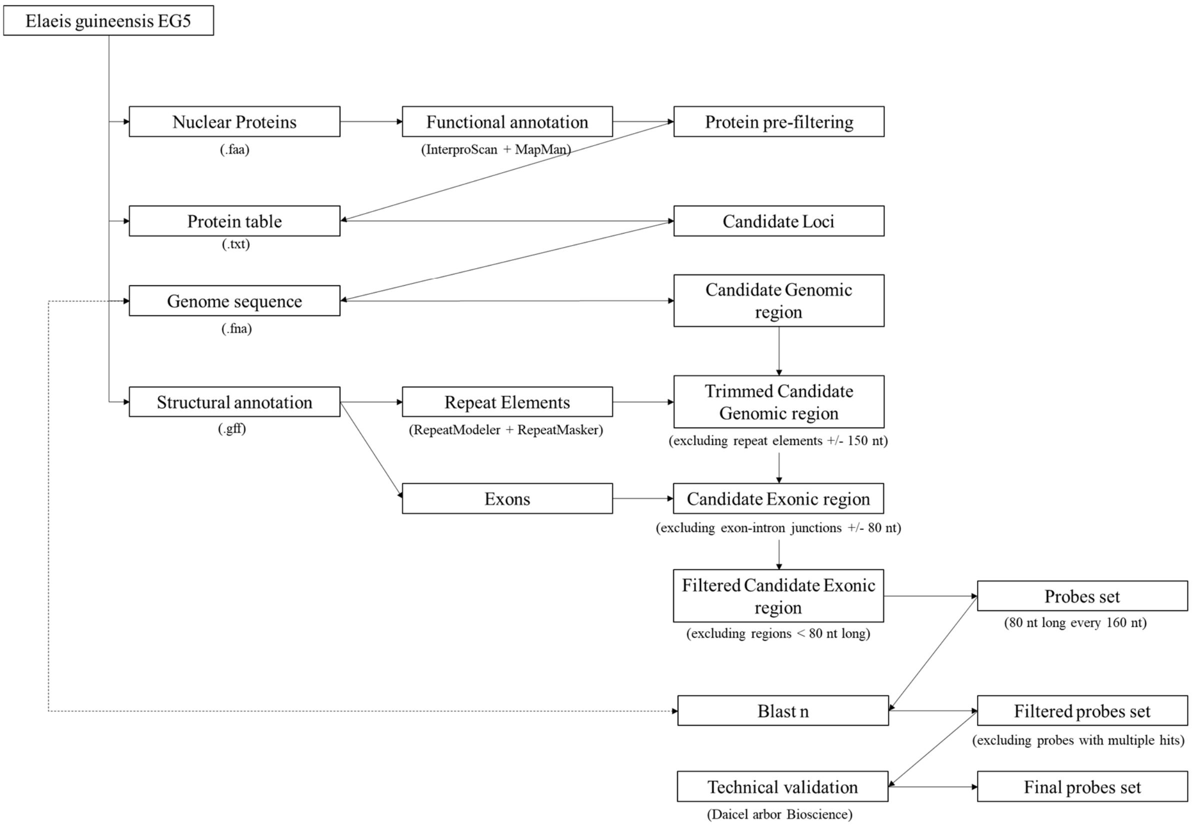


**Fig. S1.1.** Flowchart of probes design.

#### 1. Oil palm reference genome and file download

This capture kit has been designed thanks to the chromosome-level reference genome of *Elaeis guineensis* (oil palm) that has been made available on NCBI as part of the research article by Singh *et al.* 2013[1] under accession GCF_000442705.1 (EG5 assembly[2]) and annotation release 101[3]. Four files were downloaded from NCBI assembly main page[2] and used as inputs in the following steps:

- "GCF_000442705.1_EG5_genomic.fna" containing chromosome (nucleotide) sequences
- "GCF_000442705.1_EG5_protein.faa" containing protein (amino acid) sequences ;
- "GCF_000442705.1_EG5_genomic.gff" containing the structural and functional annotation (including exons and introns);
- "Protein_table.txt" containing the functional annotation in a tabular format ;

#### 2. Functional annotation

Proteins encoded by genes located on chloroplast or unplaced scaffolds were first filtered out from the "Protein_table.txt" file to retain proteins located on nuclear chromosomes only, and the protein sequences file (.faa) was updated accordingly. Nuclear protein sequences were then annotated using `InterProScan 5.21-60.0` with option "`-go terms`" to get Gene Ontology (GO) terms on one side, and with Mercator v.3.6[4] to get Mapman terms on the other.

#### 3. Protein pre-filtering

A large set of candidate proteins for adaptation and domestication (and corresponding genes) was then pre-filtered by combining a "top-down" with a "bottom-up" approach. The bottom-up approach consisted in the manual selection of genes found in the literature and documented as being involved in metabolic pathways of interest, such as oil metabolism, isoprenoid metabolism, carbon metabolism or reproduction. The top-down approach consisted in the automated selection of genes based on their functional annotation, by selecting genes with specific Gene Ontology or MapMan terms of interest (*e.g.* "cellular carbohydrate metabolic

---

[1] Singh, R. *et al.* (2013). Oil palm genome sequence reveals divergence of interfertile species in Old and New worlds. Nature, 500(7462), 335–339. doi: 10.1038/nature12309

[2] https://www.ncbi.nlm.nih.gov/genome/?term=Elaeis+guineensis

[3] https://www.ncbi.nlm.nih.gov/genome/annotation_euk/Elaeis_guineensis/101/

[4] https://plabipd.de/portal/mercator-sequence-annotation

process", "cellular lipid metabolic process", "lipid metabolic process", "developmental growth", "response to stress", "secondary metabolism", etc.) completed by a search for regular expressions (*e.g.* "lipid", "embryogenesis", "seed dormancy process", "seed germination", "reproduction" "reproductive process", "pollination", "nitrate", "light", "temperature", "reactive oxygene species", etc.). This step resulted in the pre-filtering of 11,010 proteins out of the 32,455 nuclear proteins.

**4. Structural annotation of repeat elements**

A custom (species-specific) database of repeat elements was created using `RepeatModeler v.1.0.11`: the database was first created using the utility `BuildDatabae` with option "`-engine ncbi`" and the utility `RepeatModeler` with options "`-engine ncbi and -pa 10`". Chromosome sequences (.fna) were then screened for interspersed repeats and low complexity DNA sequences using `RepeatMasker` with options "`-s -xsmall -pa 10 -gff -engine ncbi`". The localization of every repeat element was extracted from the .gff file produced by `RepeatMasker`.

**5. Trimming of candidate genomic regions, avoiding repeat elements and retaining exons**

The start and end positions of the loci encoding candidate proteins pre-filtered in step 3 were extracted from the file "Protein_table.txt", providing a set of candidate loci. Then, the corresponding genomic regions that intersected a repeat element +/- 150 nucleotides were trimmed to avoid designing probes close to (*i.e.* less than 150 nt from) a repeat element. The start and end positions of exons was then extracted from the "GCF_000442705.1_EG5_genomic.gff" file. Then, only the candidate genomic regions that intersected an exon were retained, trimming exonic regions less than 80 nucleotides from a junction exon-intron. Last, the remaining candidate exonic region less than 80 nucleotides long were filtered out.

**6. Probes design**

Molecular probes 80-nucleotide length were then designed every 160 nucleotides in the remaining candidate exonic regions, resulting in 26,971 probes. This preliminary set of probes was then blasted against the entire reference assembly (including chloroplast or unplaced scaffolds) with blastn to filter out probes similar to multiple regions of the genome that would result in aspecific hybridization. All the probes with multiple hits (with an alignement length of 80 nt and a similarity higher than 80%) were thus excluded from the set of probes, resulting in 20,849 probes. This set of probes was further submitted to technical validation by Daicel Arbor Bioscience to retain the 20,000 best probes located within 5,209 genes. The 20,000 probes of the custom capture are scattered across all chromosomes, as shown in **Fig. 2**.

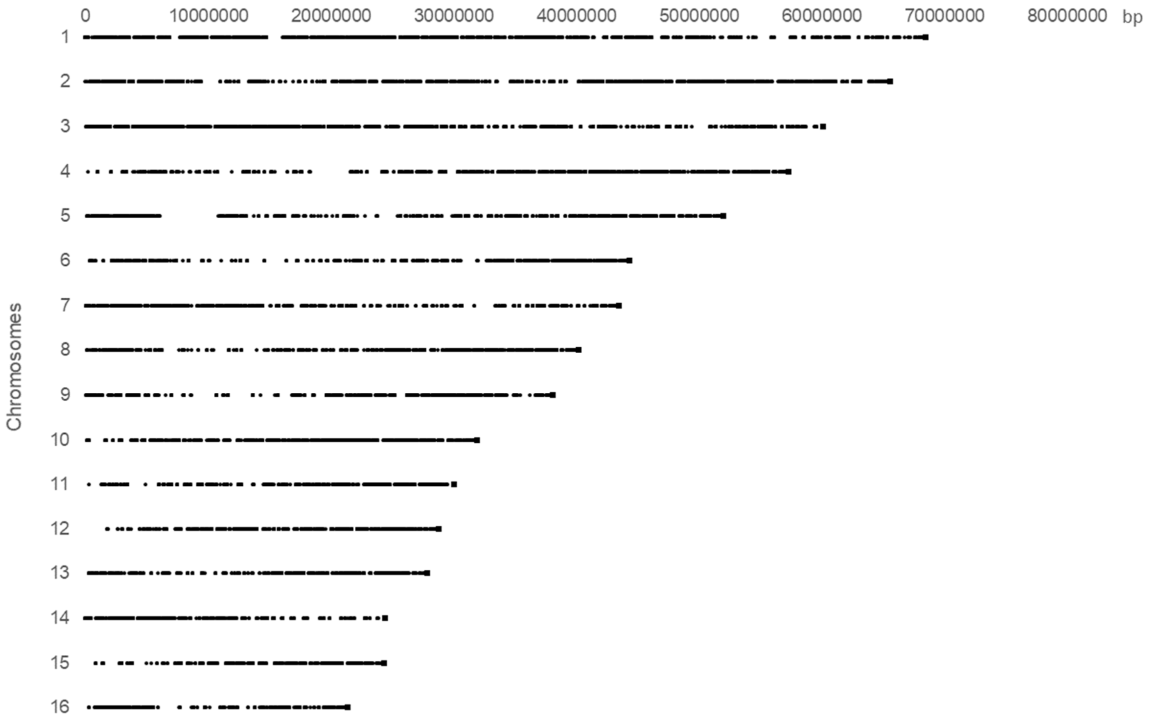


**Fig. S1.2.** Distribution of probes across the 16 chromosomes of *Elaeis guineensis*. The squares indicate chromosome ends.

#### 7. Experimental validation

The final set of probes was synthesized by Daicel Arbor Biosciences (myBaits Custom DNA-Seq©) and experimentally evaluated on 24 libraries of the species (18 libraries of *Astrocaryum spp.*, and 6 libraries of *Oenocarpus bacaba*) on one MiSeq run, using the plant materials collected. The experimental validation of the capture kit revealed success rates above 90% indicating that the probes successfully hybridized with the targeted genomic regions, and a coverage on-target largely above the coverage off-target, indicating a high specificity of the probes design to the targeted regions with marginal enrichment of aspecific (i.e. non-targeted) regions. Sequencing and mapping statistics of the entire dataset (405 libraries) are provided in Supplementary materials 3 and 4.

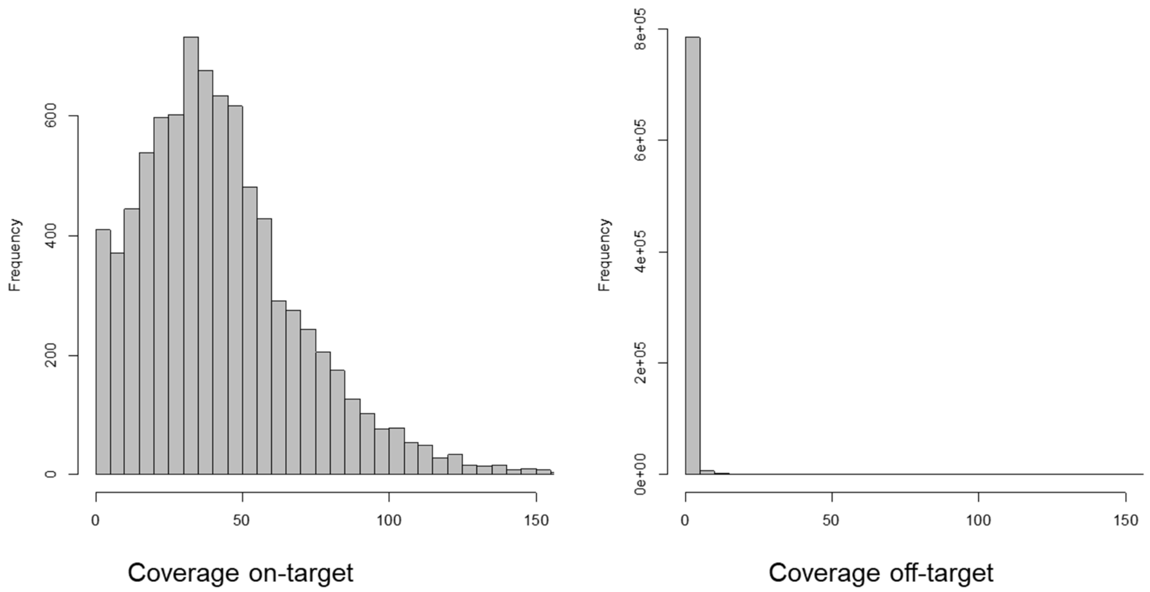


**Table S1.1**. Summary statistics of the libraries sequenced during the experimental evaluation of the custom capture kit. "On-target" refers to the targeted genomic regions where the probes were designed; "off-target" refers to aspecific genomic regions. Notice that the number of "contiguous on-target regions" is lower than the number of probes design, because the reads of regions targeted by adjacent probes are often overlapping due to the proximity of many probes (designed every 160 nt within candidate exonic regions).

| | ***All libs* (x24 libs)** | ***Astrocaryum spp.* (x18 libs)** | ***Oenocarpus bacaba* (x6 libs)** |
|---|---|---|---|
| Sequence duplication | No | No | No |
| N cleaned reads | 7'751'945 | 6'434'984 | 1'316'961 |
| N mapped reads | 7'409'422 | 6'323'710 | 1'085'751 |
| Prop. Mapped reads | 95.58% | 98.27% | 82.45% |
| Insert size (mean) | 227.7 | 225.9 | 238.0 |
| Insert size (sd) | 99.6 | 98.0 | 107.6 |
| N probes mapped | **19'702** | **19'536** | **18'553** |
| N probes unmapped | 298 | 464 | 1'447 |
| N contiguous genomic regions mapped (on- & off-targets) | 805'290 | 710'661 | 166'089 |
| N contiguous on-target regions mapped | 8'419 | 8'439 | 8'988 |
| N contiguous off-target regions mapped | 796'871 | 702'222 | 157'101 |
| Coverage on-target (including probes' flanking regions) | 44.55X | 39.25X | 10.04X |
| Coverage off-target | 1.40X | 1.39X | 1.29X |
