## Supplementary material 2 for "Natural and human-mediated drivers of microevolution in Neotropical palms: a historical genomics approach"

### Molecular methods

This supplementary Material provides the molecular protocol for library preparation and capture

- Plant DNA purification
- Target preparation, construction of barcoded genomic libraries
- PCR post-capture
- Sequencing
- **Table S2.1.** Capture adapters

#### 1. Plant DNA purification

DNA was extracted from 15 mg of piece of leaf, dried in Silicagel, with the Chemagic DNA Plant Kit (Perkin Elmer Chemagen, Baesweller, DE, Part # CMG-194), according to manufacturer's instructions. The protocol was adapted to the use of the KingFisher Flex™ (Thermo Fisher Scientific, Waltham, MA, USA) automated DNA purification workstation.

#### 2. Target preparation, construction of barcoded genomic libraries

Genomic library preparation for multiplexed individuals and enrichment step by capture follow published protocols of Rohland and Reich 2012 and Mascher et al. 2013 respectively with some modifications.

1: For each individual, 1 µg of total DNA (in 100 µL of water) were sheared using a Bioruptor Pico (Diagenode, Seraing, BE) sonication device in 500 µl microtubes to a targeted 300 bp DNA fragment size using parameters of the 300pb standard protocol for DNA shearing[1].

[1] https://www.diagenode.com/files/protocols/Standard_protocols_for_DNAShearing.pdf

2: 400 ng of fragmented DNA (in 40 µl of water) were blunted and 5' phosphorylated using the Thermo Scientific Fast DNA End Repair Kit (Thermo Fischer Scientific, Waltham, MA, USA, Part # K0771). A clean-up step was performed with 1x volume of Agencourt AMPure XP magnetic beads (Beckman Coulter Life Sciences, Indianapolis, IN, USA, Part # A63881).
Elution volume = 20 µL.

3: Fragmented and repaired DNA were individually controlled (sizing and estimation of the concentration) by electrophoresis on a AATI Fragment Analyzer™ (Advanced Analytical Technologies, Ankeny, IA, USA) device with the DNF-474 High Sensitivity Fragment Analysis Kit.

4: 50 ng of fragmented DNA were ligated with 4 pmol of PE-P5 and MPE-P7 adapters. Each PE-P5 and each MPE-P7 adapter carried the same specific hexamer barcode (modified protocol from Rohland and Reich 2012, see **Table S2.1 "Capture adapters"**). Reactions were conducted in 15 µl final volume with 1 unit of T4 DNA ligase for 1 hour at 22 °C followed by a heat inactivation step at 65°C for 10 minutes.

5: 48 samples (corresponding to 48 hexamer barcodes on the PE-P5 and PE-P7 adapter) were pooled. A clean-up step was performed with 1.8x volume of Agencourt AMPure XP magnetic beads.
Elution volume = 94 µL.

6: A nick fill-in step was performed using 64 units of Bst DNA polymerase (New England Biolabs, Ipswich, MA, USA, Part # M0275), 1x ThermoPol® reaction buffer, 250 µM dNTP in 120 µl final volume and incubated for 15 minutes at 37°C. A clean-up step was performed with 1.8x volume of Agencourt AMPure XP magnetic beads.
Elution volume = 40 µL.

7: For each pool of 48 samples, a pre-hybridization PCR was performed using the Phusion® High-Fidelity PCR Master Mix (Thermo Fischer Scientific, Part # 1040-2678) with 200 nM PreHyb-PE_F (CTTTCCCTACACGACGCTCTTC) and 200 nM PreHyb-MPE_R (TGACTGGAGTTCAGACGTGTG) primers in a final volume of 100µl.

Thermocycling parameters: 3 minutes at 98°C, followed by 12 cycles of 80 seconds at 98°C; 45 seconds at 55°C and 60 seconds at 68°C, with a final elongation of 10 minutes at 72°C. A clean-up step was performed with 1.8x volume Agencourt AMPure XP magnetic beads. Elution volume = 20 µL.

#### 3. Enrichment, capture by hybridization

The protocol used is based on Mascher et al. 2013, the User Manual of the myBaits Sequence Enrichment for Targeted Sequencing kit[2] and on the Roche NimbleGen SeqCap EZ Library SR User's Guide[3].

##### 3.1. First round of hybridization of the barcoded libraries to biotinylated RNA probes (custom kit of probes "ARECO5000+", myBaits Custom DNA-Seq©).

8: Prior to hybridization, 10 µl of Roche Diagnostics (Indianapolis, IN, USA) proprietary SeqCap EZ Developer Reagent (Roche, Part # 06684335001) were added to a 1.5-ml tube containing 0.5 µg of the 48 barcoded samples genomic library.
Next, blocking oligos were added:

- 1 µl (100 pmol/µl solution) of the P5 adapter blocking oligo , 5'-AGATCGGAAGAGCGTCGTGTAGGGAAAG
- and 1 µl (100 pmol/µl solution) of the MP7 adapter blocking oligo, 5'-AGATCGGAAGAGCACACGTCTGAACTCCAGTCA,

These oligos were designed to block the truncated segment of TruSeq DNA library adapters during the sequence capture.
The mixture was dried down in a SpeedVac at 43°C during 20 to 30 min.

9: 7.5 µl of 2 × Sequence Capture Hybridization Buffer (tube 5, SeqCap EZ Hybridization and Wash Kit, Roche, Part # 05634261001) and 3 µl of Hybridization Component A (tube 6, SeqCap EZ Hybridization and Wash Kit, Roche, Part # 05634261001) were added. The hybridization cocktail was vortexed for 10 sec and collected by centrifugation. Following

---

[2] https://arborbiosci.com/wp-content/uploads/2018/04/MYbaits-manual-v2.pdf

[3] http://sequencing.roche.com/products/nimblegen-seqcap-target-enrichment/seqcap-reagents.html (broken link), retrieved from https://dnatech.genomecenter.ucdavis.edu/wp-content/uploads/2013/06/NG_SeqCap_UsersGuide_EZ_Exome_SR_v1p2.pdf

denaturation in a heat block (95°C, 10 min) the sample was transferred to a 0.2 ml PCR tube containing 80 ng of biotinylated RNA probes (4.5 µL of myBaits Capture Probe).

10: The hybridization sample (15 µl) was incubated in a thermocycler (lid heated to 57°C) at 47°C for 64 h.

##### 3.2. First round of washing of the captured library

11: Streptavidin coupled magnetic beads were previously equilibrate as recommended by the Roche-Nimblegen protocol. Invitrogen Dynabeads MyOne™ Streptavidin C1 (Invitrogen, Thermo Fischer Scientific, Part # 65001) at 10 µg/µl were thoroughly vortexed, aliquoted (50 µl per hybridization) into 1.5-ml tubes and prepared for the affinity purification of captured DNA. The tubes were placed in a DynaMag-2 magnet (Invitrogen, Part # 123-21D) for 2 min. The clear liquid was discarded and 100 µl of 1 X Bead Wash Buffer (Tube 7, SeqCap EZ Hybridization and Wash Kit, Roche, Part # 05634261001) was added. Tubes were vortexed, placed back in the magnet, the clear liquid was removed, and the washing was repeated once. Dynabeads were resuspended in 50 µl 1 x Bead Wash Buffer, transferred into PCR plates and collected using a Agencourt SPRIPlate 96R (Agencourt, Part # A32782) . The clear supernatant was discarded.

12: The hybridization sample was added to the wet Dynabeads and mixed thoroughly by pipetting up and down. Using a thermocycler (lid heated to 57°C) at 47°C for 45 min the captured sample was bound to the Dynabeads. The sample was vortexed for 3 sec in 15-min intervals to ensure that the Dynabeads remain in suspension. Dynabeads plus bound DNA (15 µl) were washed by adding 100 µl 1 X Wash Buffer 1 (pre-heated to 47°C for 1 h) (Tube 1, SeqCap EZ Hybridization and Wash Kit, Roche, Part # 05634261001) and vortexing for 10 sec.

13: The suspension iwass transferred to a 1.5-ml tube and placed in a DynaMag-2 device, and the supernatant was discarded once clear. Washing was continued by adding 200 µl 1 X Stringent Wash Buffer (pre-heated to 47°C for 1 h) (Tube 4, SeqCap EZ Hybridization and Wash Kit, Roche, Part # 05634261001). The sample was mixed by pipetting, avoiding a major temperature drop, and incubated for 5 min at 47°C. The tube was placed in the DynaMag-2 magnet, the liquid was discarded and the washing at 47°C with 1 X Stringent Wash Buffer was repeated once.

14: 200 µl 1 X Wash Buffer 1 (pre-heated to room temperature) was added to the Dynabeads plus bound DNA. The sample was vortexed for 2 min and the liquid was collected to the tube's bottom. Following magnetic concentration the liquid was discarded, and the sample was washed at room temperature with 200 µl 1 X Wash Buffer 2 (vortexing for 1 min) (Tube 2, SeqCap EZ Hybridization and Wash Kit, Roche, Part # 05634261001), followed by a wash with 200 µl 1 X Wash Buffer 3 (vortexing for 30 sec) (Tube 3, SeqCap EZ Hybridization and Wash Kit, Roche, Part # 05634261001) as described for washing with Wash Buffer 1. The tube was removed from the magnet, the bead-bound captured library was resuspended in 25 µl PCR-grade water and the entire sample (beads + liquid) was transferred to a 0.2 ml PCR tube.

##### 3.3. Second round of hybridization of the barcoded libraries to biotinylated RNA probes

15: The captured sample (25 µl) was denatured by incubation in a thermocycler (lid heated to 105°C) at 95°C for 3 min. 21 µl of the denatured solution were quickly transferred on to a 1.5 ml microtube.

16: Next adapters and reagents were added to the captured sample:

- 1 µl (10 pmol/µl solution) of the P5 adapter blocking oligo
- 1 µl (10 pmol/µl solution) of the MP7 adapter blocking oligo,
- 1 µl of SeqCap EZ Developer Reagent.

The mixture was dried down in a SpeedVac at 43°C during 10 to 20 min.

17: 7.5 µl of 2 × Sequence Capture Hybridization Buffer (tube 5, SeqCap EZ Hybridization and Wash Kit) and 3 µl of Hybridization Component A (tube 6, SeqCap EZ Hybridization and Wash Kit) were added. The hybridization cocktail was vortexed for 10 sec and collected by centrifugation. Following denaturation in a heat block (95°C, 10 min) the sample was transferred to a 0.2 ml PCR tube containing 15 ng of biotinylated RNA probes (1 µL of myBaits Capture Probe) and 3,5 µl of UP water.

18: The hybridization sample (15 µl) was incubated in a thermocycler (lid heated to 57°C) at 47°C for 20 h.

##### 3.4. Second round of washing of the captured library

19: Streptavidin coupled magnetic beads were prepared as previously described (#11).

20: The hybridization sample was added to the wet Dynabeads and mixed thoroughly by pipetting up and down. Using a thermocycler (lid heated to 57°C) at 47°C for 45 min, the captured sample was bound to the Dynabeads. The sample was vortexed for 3 sec in 15-min intervals to ensure that the Dynabeads remained in suspension. Dynabeads plus bound DNA (15 µl) were washed by adding 100 µl 1 X Wash Buffer 1 (pre-heated to 47°C for 1 h) (Tube 1, SeqCap EZ Hybridization and Wash Kit) and vortexing for 10 sec.

21: The suspension was transferred to a 1.5-ml tube and placed in a DynaMag-2 device, and the supernatant was discarded once clear. Washing was continued by adding 200 µl 1 X Stringent Wash Buffer (pre-heated to 47°C for 1 h) (Tube 4, SeqCap EZ Hybridization and Wash Kit). The sample was mixed by pipetting, avoiding a major temperature drop, and incubated for 5 min at 47°C. The tube was placed in the DynaMag-2 magnet, the liquid was discarded and the washing at 47°C with 1 X Stringent Wash Buffer was repeated once.

22: 200 µl 1 X Wash Buffer 1 (pre-heated to room temperature) was added to the Dynabeads plus bound DNA. The sample was vortexed for 2 min and the liquid was collected to the tube's bottom. Following magnetic concentration, the liquid was discarded and the sample was washed at room temperature with 200 µl 1 X Wash Buffer 2 (vortexing for 1 min) (Tube 2, SeqCap EZ Hybridization and Wash Kit), followed by a wash with 200 µl 1 X Wash Buffer 3 (vortexing for 30 sec) (Tube 3, SeqCap EZ Hybridization and Wash Kit) as described for washing with Wash Buffer 1. The tube was removed from the magnet, the bead-bound captured library was resuspended in 22 µl PCR-grade water and the entire sample (beads + liquid) was transferred to a 0.2 ml PCR tube.

#### 4. PCR post-capture

23: An on-beads PCR amplification was undertaken to enrich library fragments, extend the adaptor sequence and incorporate an index to the P7 adaptor. The PCR reaction was realized using the KAPA® HiFi HotStart ReadyMix PCR Kit (KAPABiosystems, Boston, MA, Part # KR0370) in a final volume of 50 µl with:

- 15 pmol of SOL-PE-PCR_F primer (Rohland and Reich 2012), AATGATACGGCGACCACCGAGATCTACACTCTTTCCCTACACGACGCTCTTC
- 15 pmol of SOL-MPE-INDX_R indexed primers, CAAGCAGAAGACGGCATACGAGATXXXXXXGTGACTGGAGTTCAGACGTGT

This primer carried 6 bases of the official TruSeq Illumina Index.
Thermocycling parameters: 2 minutes at 98°C, followed by 18 cycles of 20 seconds at 98°C; 30 seconds at 62°C and 30 seconds at 72°C, with a final elongation of 5 minutes at 72°C.
Reaction volume = 50 µL.
A clean-up step was performed with 1.8x volume Agencourt AMPure XP magnetic beads.
Elution volume = 20 µL.

24: Indexed libraries were individually controlled (sizing and estimation of the concentration) by electrophoresis on an AATI Fragment Analyzer™ device with the DNF-474 High Sensitivity Fragment Analysis Kit.

#### 5. Sequencing

25: Six indexed libraries, corresponding to 288 captured barcoded DNA samples, were equally mixed. The final pooled library was quantify by qPCR with the KAPA Library Quantification Kit (Part # KK4824).

26: The final pooled library was sequenced following the Illumina paired-end protocol on Illumina sequencers. DNA samples of *Oenocarpus bataua* were individually sequenced with 3 runs of Illumina MiSeq sequencer using the paired-end protocol for 2 x 150 cycles. DNA samples of *Astrocaryum* spp. and *Oenocarpus bacaba*, were equally mixed and provided to the Get-PlaGe core facility (GenoToul platform, INRA Toulouse, France http://www.genotoul.fr).

The final pooled library was sequenced using the Illumina paired-end protocol on a single lane of a HiSeq3000 sequencer, for 2 x 150 cycles.

**Table S2.1.** Capture Adapters.

| Barcode | | Barcoded P5 adapter | | | | Barcoded P7 adapter | | | |
|---|---|---|---|---|---|---|---|---|---|
| | | Direct oligo | | Complementary oligo | | Direct oligo | | Complementary oligo | |
| **Id** | **Sequence** | **Id oligo** | **sequence** | **id oligo** | **sequence** | **Id oligo** | **sequence** | **id oligo** | **sequence** |
| Barcode1 | AGCGCA | Barcoded-P5-1 | CTTTCCCTACACGACGCTCTTCCGATCTAGCGCA | Barcoded-P5-comp-1 | TGCGCTAGATCGGAA | Barcoded-MP7-1 | TGACTGGAGTTCAGACGTGTGCTCTTCCGATCTAGCGCA | Barcoded-MP7-comp1 | TGCGCTAGATCGGAAGAGC |
| Barcode2 | CTCAGC | Barcoded-P5-2 | CTTTCCCTACACGACGCTCTTCCGATCTCTCAGC | Barcoded-P5-comp-2 | GCTGAGAGATCGGAA | Barcoded-MP7-2 | TGACTGGAGTTCAGACGTGTGCTCTTCCGATCTCTCAGC | Barcoded-MP7-comp2 | GCTGAGAGATCGGAAGAGC |
| Barcode3 | TAGATC | Barcoded-P5-3 | CTTTCCCTACACGACGCTCTTCCGATCTTAGATC | Barcoded-P5-comp-3 | GATCTAAGATCGGAA | Barcoded-MP7-3 | TGACTGGAGTTCAGACGTGTGCTCTTCCGATCTTAGATC | Barcoded-MP7-comp3 | GATCTAAGATCGGAAGAGC |
| Barcode4 | ACTGAT | Barcoded-P5-4 | CTTTCCCTACACGACGCTCTTCCGATCTACTGAT | Barcoded-P5-comp-4 | ATCAGTAGATCGGAA | Barcoded-MP7-4 | TGACTGGAGTTCAGACGTGTGCTCTTCCGATCTACTGAT | Barcoded-MP7-comp4 | ATCAGTAGATCGGAAGAGC |
| Barcode5 | ATACAC | Barcoded-P5-5 | CTTTCCCTACACGACGCTCTTCCGATCTATACAC | Barcoded-P5-comp-5 | GTGTATAGATCGGAA | Barcoded-MP7-5 | TGACTGGAGTTCAGACGTGTGCTCTTCCGATCTATACAC | Barcoded-MP7-comp5 | GTGTATAGATCGGAAGAGC |
| Barcode6 | ATGTGT | Barcoded-P5-6 | CTTTCCCTACACGACGCTCTTCCGATCTATGTGT | Barcoded-P5-comp-6 | ACACATAGATCGGAA | Barcoded-MP7-6 | TGACTGGAGTTCAGACGTGTGCTCTTCCGATCTATGTGT | Barcoded-MP7-comp6 | ACACATAGATCGGAAGAGC |
| Barcode7 | CACTTA | Barcoded-P5-7 | CTTTCCCTACACGACGCTCTTCCGATCTCACTTA | Barcoded-P5-comp-7 | TAAGTGAGATCGGAA | Barcoded-MP7-7 | TGACTGGAGTTCAGACGTGTGCTCTTCCGATCTCACTTA | Barcoded-MP7-comp7 | TAAGTGAGATCGGAAGAGC |
| Barcode8 | CCATAG | Barcoded-P5-8 | CTTTCCCTACACGACGCTCTTCCGATCTCCATAG | Barcoded-P5-comp-8 | CTATGGAGATCGGAA | Barcoded-MP7-8 | TGACTGGAGTTCAGACGTGTGCTCTTCCGATCTCCATAG | Barcoded-MP7-comp8 | CTATGGAGATCGGAAGAGC |
| Barcode9 | GAGGCA | Barcoded-P5-9 | CTTTCCCTACACGACGCTCTTCCGATCTGAGGCA | Barcoded-P5-comp-9 | TGCCTCAGATCGGAA | Barcoded-MP7-9 | TGACTGGAGTTCAGACGTGTGCTCTTCCGATCTGAGGCA | Barcoded-MP7-comp9 | TGCCTCAGATCGGAAGAGC |
| Barcode10 | GGTACG | Barcoded-P5-10 | CTTTCCCTACACGACGCTCTTCCGATCTGGTACG | Barcoded-P5-comp-10 | CGTACCAGATCGGAA | Barcoded-MP7-10 | TGACTGGAGTTCAGACGTGTGCTCTTCCGATCTGGTACG | Barcoded-MP7-comp10 | CGTACCAGATCGGAAGAGC |
| Barcode11 | GTGCCT | Barcoded-P5-11 | CTTTCCCTACACGACGCTCTTCCGATCTGTGCCT | Barcoded-P5-comp-11 | AGGCACAGATCGGAA | Barcoded-MP7-11 | TGACTGGAGTTCAGACGTGTGCTCTTCCGATCTGTGCCT | Barcoded-MP7-comp11 | AGGCACAGATCGGAAGAGC |
| Barcode12 | TGTGTG | Barcoded-P5-12 | CTTTCCCTACACGACGCTCTTCCGATCTTGTGTG | Barcoded-P5-comp-12 | CACACAAGATCGGAA | Barcoded-MP7-12 | TGACTGGAGTTCAGACGTGTGCTCTTCCGATCTTGTGTG | Barcoded-MP7-comp12 | CACACAAGATCGGAAGAGC |
| Barcode13 | AACCTT | Barcoded-P5-13 | CTTTCCCTACACGACGCTCTTCCGATCTAACCTT | Barcoded-P5-comp-13 | AAGGTTAGATCGGAA | Barcoded-MP7-13 | TGACTGGAGTTCAGACGTGTGCTCTTCCGATCTAACCTT | Barcoded-MP7-comp13 | AAGGTTAGATCGGAAGAGC |
| Barcode14 | AAGTTG | Barcoded-P5-14 | CTTTCCCTACACGACGCTCTTCCGATCTAAGTTG | Barcoded-P5-comp-14 | CAACTTAGATCGGAA | Barcoded-MP7-14 | TGACTGGAGTTCAGACGTGTGCTCTTCCGATCTAAGTTG | Barcoded-MP7-comp14 | CAACTTAGATCGGAAGAGC |

| | | | | | | | | | |
|---|---|---|---|---|---|---|---|---|---|
| Barcode15 | AATACA | Barcoded-P5-15 | CTTTCCCTACACGACGCTCTTCCGATCTAATACA | Barcoded-P5-comp-15 | TGTATTAGATCGGAA | Barcoded-MP7-15 | TGACTGGAGTTCAGACGTGTGCTCTTCCGATCTAATACA | Barcoded-MP7-comp15 | TGTATTAGATCGGAAGAGC |
| Barcode16 | ACAATA | Barcoded-P5-16 | CTTTCCCTACACGACGCTCTTCCGATCTACAATA | Barcoded-P5-comp-16 | TATTGTAGATCGGAA | Barcoded-MP7-16 | TGACTGGAGTTCAGACGTGTGCTCTTCCGATCTACAATA | Barcoded-MP7-comp16 | TATTGTAGATCGGAAGAGC |
| Barcode17 | ACCACA | Barcoded-P5-17 | CTTTCCCTACACGACGCTCTTCCGATCTACCACA | Barcoded-P5-comp-17 | TGTGGTAGATCGGAA | Barcoded-MP7-17 | TGACTGGAGTTCAGACGTGTGCTCTTCCGATCTACCACA | Barcoded-MP7-comp17 | TGTGGTAGATCGGAAGAGC |
| Barcode18 | ACTCAC | Barcoded-P5-18 | CTTTCCCTACACGACGCTCTTCCGATCTACTCAC | Barcoded-P5-comp-18 | GTGAGTAGATCGGAA | Barcoded-MP7-18 | TGACTGGAGTTCAGACGTGTGCTCTTCCGATCTACTCAC | Barcoded-MP7-comp18 | GTGAGTAGATCGGAAGAGC |
| Barcode19 | AGAAGG | Barcoded-P5-19 | CTTTCCCTACACGACGCTCTTCCGATCTAGAAGG | Barcoded-P5-comp-19 | CCTTCTAGATCGGAA | Barcoded-MP7-19 | TGACTGGAGTTCAGACGTGTGCTCTTCCGATCTAGAAGG | Barcoded-MP7-comp19 | CCTTCTAGATCGGAAGAGC |
| Barcode20 | AGGCGA | Barcoded-P5-20 | CTTTCCCTACACGACGCTCTTCCGATCTAGGCGA | Barcoded-P5-comp-20 | TCGCCTAGATCGGAA | Barcoded-MP7-20 | TGACTGGAGTTCAGACGTGTGCTCTTCCGATCTAGGCGA | Barcoded-MP7-comp20 | TCGCCTAGATCGGAAGAGC |
| Barcode21 | AGGTTC | Barcoded-P5-21 | CTTTCCCTACACGACGCTCTTCCGATCTAGGTTC | Barcoded-P5-comp-21 | GAACCTAGATCGGAA | Barcoded-MP7-21 | TGACTGGAGTTCAGACGTGTGCTCTTCCGATCTAGGTTC | Barcoded-MP7-comp21 | GAACCTAGATCGGAAGAGC |
| Barcode22 | AGTGGC | Barcoded-P5-22 | CTTTCCCTACACGACGCTCTTCCGATCTAGTGGC | Barcoded-P5-comp-22 | GCCACTAGATCGGAA | Barcoded-MP7-22 | TGACTGGAGTTCAGACGTGTGCTCTTCCGATCTAGTGGC | Barcoded-MP7-comp22 | GCCACTAGATCGGAAGAGC |
| Barcode23 | ATGTAG | Barcoded-P5-23 | CTTTCCCTACACGACGCTCTTCCGATCTATGTAG | Barcoded-P5-comp-23 | CTACATAGATCGGAA | Barcoded-MP7-23 | TGACTGGAGTTCAGACGTGTGCTCTTCCGATCTATGTAG | Barcoded-MP7-comp23 | CTACATAGATCGGAAGAGC |
| Barcode24 | CACCAC | Barcoded-P5-24 | CTTTCCCTACACGACGCTCTTCCGATCTCACCAC | Barcoded-P5-comp-24 | GTGGTGAGATCGGAA | Barcoded-MP7-24 | TGACTGGAGTTCAGACGTGTGCTCTTCCGATCTCACCAC | Barcoded-MP7-comp24 | GTGGTGAGATCGGAAGAGC |
| Barcode25 | CACTGC | Barcoded-P5-25 | CTTTCCCTACACGACGCTCTTCCGATCTCACTGC | Barcoded-P5-comp-25 | GCAGTGAGATCGGAA | Barcoded-MP7-25 | TGACTGGAGTTCAGACGTGTGCTCTTCCGATCTCACTGC | Barcoded-MP7-comp25 | GCAGTGAGATCGGAAGAGC |
| Barcode26 | CCGACG | Barcoded-P5-26 | CTTTCCCTACACGACGCTCTTCCGATCTCCGACG | Barcoded-P5-comp-26 | CGTCGGAGATCGGAA | Barcoded-MP7-26 | TGACTGGAGTTCAGACGTGTGCTCTTCCGATCTCCGACG | Barcoded-MP7-comp26 | CGTCGGAGATCGGAAGAGC |
| Barcode27 | CCTCCA | Barcoded-P5-27 | CTTTCCCTACACGACGCTCTTCCGATCTCCTCCA | Barcoded-P5-comp-27 | TGGAGGAGATCGGAA | Barcoded-MP7-27 | TGACTGGAGTTCAGACGTGTGCTCTTCCGATCTCCTCCA | Barcoded-MP7-comp27 | TGGAGGAGATCGGAAGAGC |
| Barcode28 | CCTGTA | Barcoded-P5-28 | CTTTCCCTACACGACGCTCTTCCGATCTCCTGTA | Barcoded-P5-comp-28 | TACAGGAGATCGGAA | Barcoded-MP7-28 | TGACTGGAGTTCAGACGTGTGCTCTTCCGATCTCCTGTA | Barcoded-MP7-comp28 | TACAGGAGATCGGAAGAGC |
| Barcode29 | CGACTC | Barcoded-P5-29 | CTTTCCCTACACGACGCTCTTCCGATCTCGACTC | Barcoded-P5-comp-29 | GAGTCGAGATCGGAA | Barcoded-MP7-29 | TGACTGGAGTTCAGACGTGTGCTCTTCCGATCTCGACTC | Barcoded-MP7-comp29 | GAGTCGAGATCGGAAGAGC |
| Barcode30 | CTAAGT | Barcoded-P5-30 | CTTTCCCTACACGACGCTCTTCCGATCTCTAAGT | Barcoded-P5-comp-30 | ACTTAGAGATCGGAA | Barcoded-MP7-30 | TGACTGGAGTTCAGACGTGTGCTCTTCCGATCTCTAAGT | Barcoded-MP7-comp30 | ACTTAGAGATCGGAAGAGC |
| Barcode31 | CTATGA | Barcoded-P5-31 | CTTTCCCTACACGACGCTCTTCCGATCTCTATGA | Barcoded-P5-comp-31 | TCATAGAGATCGGAA | Barcoded-MP7-31 | TGACTGGAGTTCAGACGTGTGCTCTTCCGATCTCTATGA | Barcoded-MP7-comp31 | TCATAGAGATCGGAAGAGC |
| Barcode32 | CTATTG | Barcoded-P5-32 | CTTTCCCTACACGACGCTCTTCCGATCTCTATTG | Barcoded-P5-comp-32 | CAATAGAGATCGGAA | Barcoded-MP7-32 | TGACTGGAGTTCAGACGTGTGCTCTTCCGATCTCTATTG | Barcoded-MP7-comp32 | CAATAGAGATCGGAAGAGC |

| | | | | | | | | | |
|---|---|---|---|---|---|---|---|---|---|
| Barcode33 | CTCTCG | Barcoded-P5-33 | CTTTCCCTACACGACGCTCTTCCGATCTCTCTCG | Barcoded-P5-comp-33 | CGAGAGAGATCGGAA | Barcoded-MP7-33 | TGACTGGAGTTCAGACGTGTGCTCTTCCGATCTCTCTCG | Barcoded-MP7-comp33 | CGAGAGAGATCGGAAGAGC |
| Barcode34 | CTGCTG | Barcoded-P5-34 | CTTTCCCTACACGACGCTCTTCCGATCTCTGCTG | Barcoded-P5-comp-34 | CAGCAGAGATCGGAA | Barcoded-MP7-34 | TGACTGGAGTTCAGACGTGTGCTCTTCCGATCTCTGCTG | Barcoded-MP7-comp34 | CAGCAGAGATCGGAAGAGC |
| Barcode35 | GAAGTC | Barcoded-P5-35 | CTTTCCCTACACGACGCTCTTCCGATCTGAAGTC | Barcoded-P5-comp-35 | GACTTCAGATCGGAA | Barcoded-MP7-35 | TGACTGGAGTTCAGACGTGTGCTCTTCCGATCTGAAGTC | Barcoded-MP7-comp35 | GACTTCAGATCGGAAGAGC |
| Barcode36 | GACAGA | Barcoded-P5-36 | CTTTCCCTACACGACGCTCTTCCGATCTGACAGA | Barcoded-P5-comp-36 | TCTGTCAGATCGGAA | Barcoded-MP7-36 | TGACTGGAGTTCAGACGTGTGCTCTTCCGATCTGACAGA | Barcoded-MP7-comp36 | TCTGTCAGATCGGAAGAGC |
| Barcode37 | GCATTC | Barcoded-P5-37 | CTTTCCCTACACGACGCTCTTCCGATCTGCATTC | Barcoded-P5-comp-37 | GAATGCAGATCGGAA | Barcoded-MP7-37 | TGACTGGAGTTCAGACGTGTGCTCTTCCGATCTGCATTC | Barcoded-MP7-comp37 | GAATGCAGATCGGAAGAGC |
| Barcode38 | GCCTTG | Barcoded-P5-38 | CTTTCCCTACACGACGCTCTTCCGATCTGCCTTG | Barcoded-P5-comp-38 | CAAGGCAGATCGGAA | Barcoded-MP7-38 | TGACTGGAGTTCAGACGTGTGCTCTTCCGATCTGCCTTG | Barcoded-MP7-comp38 | CAAGGCAGATCGGAAGAGC |
| Barcode39 | GCGCTA | Barcoded-P5-39 | CTTTCCCTACACGACGCTCTTCCGATCTGCGCTA | Barcoded-P5-comp-39 | TAGCGCAGATCGGAA | Barcoded-MP7-39 | TGACTGGAGTTCAGACGTGTGCTCTTCCGATCTGCGCTA | Barcoded-MP7-comp39 | TAGCGCAGATCGGAAGAGC |
| Barcode40 | GCTAAG | Barcoded-P5-40 | CTTTCCCTACACGACGCTCTTCCGATCTGCTAAG | Barcoded-P5-comp-40 | CTTAGCAGATCGGAA | Barcoded-MP7-40 | TGACTGGAGTTCAGACGTGTGCTCTTCCGATCTGCTAAG | Barcoded-MP7-comp40 | CTTAGCAGATCGGAAGAGC |
| Barcode41 | GCTGCC | Barcoded-P5-41 | CTTTCCCTACACGACGCTCTTCCGATCTGCTGCC | Barcoded-P5-comp-41 | GGCAGCAGATCGGAA | Barcoded-MP7-41 | TGACTGGAGTTCAGACGTGTGCTCTTCCGATCTGCTGCC | Barcoded-MP7-comp41 | GGCAGCAGATCGGAAGAGC |
| Barcode42 | GGCATC | Barcoded-P5-42 | CTTTCCCTACACGACGCTCTTCCGATCTGGCATC | Barcoded-P5-comp-42 | GATGCCAGATCGGAA | Barcoded-MP7-42 | TGACTGGAGTTCAGACGTGTGCTCTTCCGATCTGGCATC | Barcoded-MP7-comp42 | GATGCCAGATCGGAAGAGC |
| Barcode43 | GGCGAC | Barcoded-P5-43 | CTTTCCCTACACGACGCTCTTCCGATCTGGCGAC | Barcoded-P5-comp-43 | GTCGCCAGATCGGAA | Barcoded-MP7-43 | TGACTGGAGTTCAGACGTGTGCTCTTCCGATCTGGCGAC | Barcoded-MP7-comp43 | GTCGCCAGATCGGAAGAGC |
| Barcode44 | GGTCGT | Barcoded-P5-44 | CTTTCCCTACACGACGCTCTTCCGATCTGGTCGT | Barcoded-P5-comp-44 | ACGACCAGATCGGAA | Barcoded-MP7-44 | TGACTGGAGTTCAGACGTGTGCTCTTCCGATCTGGTCGT | Barcoded-MP7-comp44 | ACGACCAGATCGGAAGAGC |
| Barcode45 | GGTTGG | Barcoded-P5-45 | CTTTCCCTACACGACGCTCTTCCGATCTGGTTGG | Barcoded-P5-comp-45 | CCAACCAGATCGGAA | Barcoded-MP7-45 | TGACTGGAGTTCAGACGTGTGCTCTTCCGATCTGGTTGG | Barcoded-MP7-comp45 | CCAACCAGATCGGAAGAGC |
| Barcode46 | GTGTCC | Barcoded-P5-46 | CTTTCCCTACACGACGCTCTTCCGATCTGTGTCC | Barcoded-P5-comp-46 | GGACACAGATCGGAA | Barcoded-MP7-46 | TGACTGGAGTTCAGACGTGTGCTCTTCCGATCTGTGTCC | Barcoded-MP7-comp46 | GGACACAGATCGGAAGAGC |
| Barcode47 | TAAGGT | Barcoded-P5-47 | CTTTCCCTACACGACGCTCTTCCGATCTTAAGGT | Barcoded-P5-comp-47 | ACCTTAAGATCGGAA | Barcoded-MP7-47 | TGACTGGAGTTCAGACGTGTGCTCTTCCGATCTTAAGGT | Barcoded-MP7-comp47 | ACCTTAAGATCGGAAGAGC |
| Barcode48 | TACGAT | Barcoded-P5-48 | CTTTCCCTACACGACGCTCTTCCGATCTTACGAT | Barcoded-P5-comp-48 | ATCGTAAGATCGGAA | Barcoded-MP7-48 | TGACTGGAGTTCAGACGTGTGCTCTTCCGATCTTACGAT | Barcoded-MP7-comp48 | ATCGTAAGATCGGAAGAGC |
