## Supplementary material 3 for "Natural and human-mediated drivers of microevolution in Neotropical palms: a historical genomics approach"

**Supplementary Material 3 (S3)**

**Sequencing and bioinformatics (405 libraries)**

This supplementary Material provides supplementary figures, tables and information regarding the sequencing and bioinformatics treatment of the full dataset (405 libraries):

- Sequencing quality check: Figure S3.1
- Capture success: Table S3.1
- Mapping on-target coverage: Figure S3.2
- Sequencing and mapping statistics: Table S3.2
- Methodological consideration on the differences between *Astrocaryum spp.* and *Oenocarpus spp.*

**Figure S3.1.** Quality check: per-base sequence quality in one library (here sample "F2_AP02.1") following reads cleaning. Full fastQC results are provided in Supplementary Material 3.

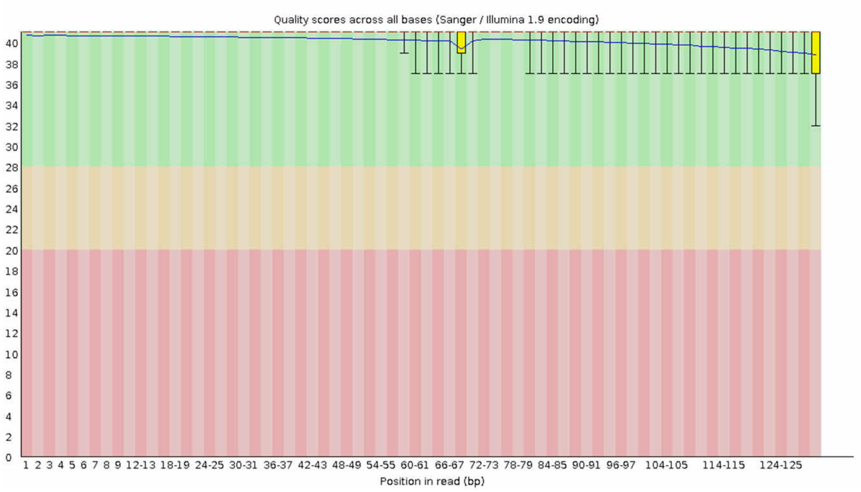


**Table S3.1.** Capture success across all *Astrocaryum spp.* and *Oenocarpus spp.* libraries, respectively.

| | ***Astrocaryum spp.*** | ***Oenocarpus spp.*** |
|---|---|---|
| N probes successfully mapped (out of 20,000 probes) | 19,899 | 19,835 |
| N probes unmapped | 101 | 165 |
| Mean coverage on target (sd) | 4812.6X (3994.1) | 1670.3X (1823.2) |

**Figure S3.2.** Mean on-target coverage across all *Astrocaryum spp.* (left) and *Oenocarpus spp.* (right) libraries.

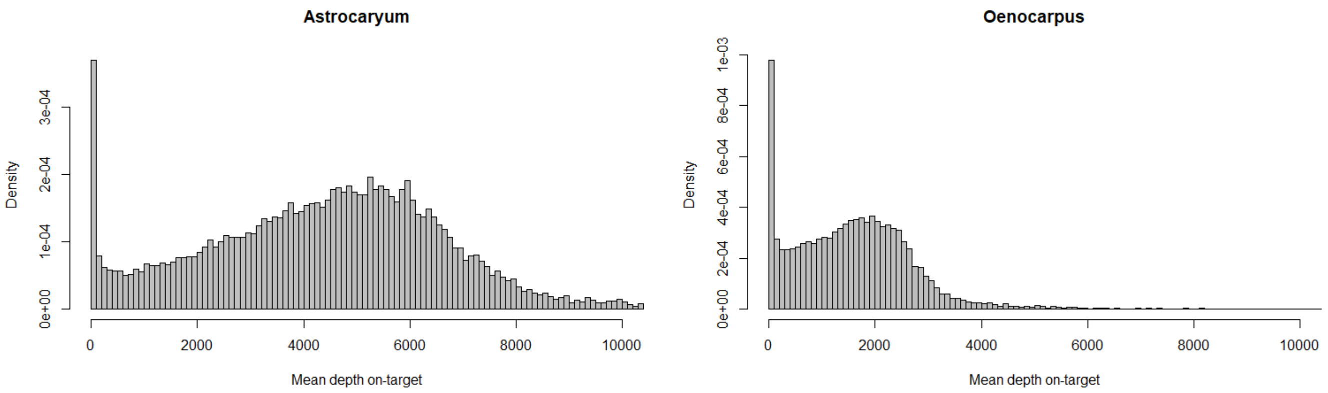

**Table S3.2.** Sequencing and mapping statistics.

| Library | N raw reads (R1) | N clean reads (R1) | N mapped reads | Mean Insert size | Mean depth | Mean depth on-target |
|---|---|---|---|---|---|---|
| **F10_AP01** | 638838 | 834176 | 834176 | 233.9 | 5.95326 | 13.099 |
| **F10_AP03** | 1164808 | 996212 | 996212 | 230.1 | 3.79262 | 9.41535 |
| **F10_AP05** | 1768114 | 1608949 | 1608949 | 224.5 | 4.76956 | 14.4439 |
| **F10_AP07** | 2407522 | 2322271 | 2322271 | 191.4 | 5.93711 | 20.9022 |
| **F10_AP09** | 2223711 | 2192470 | 2192470 | 218.9 | 5.68616 | 19.3389 |
| **F10_AP11** | 1395521 | 1256092 | 1256092 | 236.8 | 4.54549 | 12.3871 |
| **F10_AP13** | 1987519 | 1865704 | 1865704 | 217.4 | 5.40289 | 17.3685 |
| **F10_AP14** | 2257203 | 2163688 | 2163688 | 218.6 | 5.52898 | 19.0065 |
| **F10_AP16** | 1405372 | 1320719 | 1320719 | 221.5 | 4.92849 | 13.9149 |
| **F10_AP20** | 1497334 | 1342363 | 1342363 | 227.6 | 4.36091 | 12.3869 |
| **F10_AS01** | 2143605 | 2127129 | 2127129 | 214.8 | 6.05953 | 20.3062 |
| **F10_AS06** | 1614585 | 1605043 | 1605043 | 227.7 | 6.27002 | 17.5638 |
| **F10_AS07** | 1035830 | 955845 | 955845 | 222.1 | 4.44462 | 11.0565 |
| **F10_OP03** | 253468 | 232701 | 232701 | 199.8 | 2.73761 | 4.01665 |
| **F10_OP04** | 646137 | 480063 | 480063 | 218.5 | 3.6821 | 6.47213 |
| **F10_OP09** | 1015714 | 768743 | 768743 | 230.6 | 4.3897 | 9.58491 |
| **F10_OP10** | 383006 | 344247 | 344247 | 194.4 | 3.24327 | 5.40235 |
| **F10_OP11** | 543636 | 491671 | 491671 | 194.5 | 3.77503 | 7.44743 |
| **F10_OP12** | 392200 | 348393 | 348393 | 201.9 | 3.20514 | 5.38321 |
| **F10_OP13** | 615607 | 576022 | 576022 | 187.7 | 4.20861 | 8.7481 |
| **F11_AP04** | 1152431 | 997958 | 997958 | 218.1 | 4.40099 | 10.848 |
| **F11_AP12** | 1343886 | 1191265 | 1191265 | 229.8 | 4.16857 | 11.592 |
| **F11_AP14** | 1331725 | 1120301 | 1120301 | 220.6 | 4.14143 | 11.0817 |
| **F11_AP16** | 1604705 | 1544863 | 1544863 | 229.6 | 4.81465 | 14.9133 |
| **F11_OC01** | 786277 | 689700 | 689700 | 224.7 | 5.57188 | 10.98 |

| | | | | | | |
|---|---|---|---|---|---|---|
| **F11_OC03** | 1145664 | 1018651 | 1018651 | 223.1 | 6.63567 | 15.3448 |
| **F11_OC04** | 1034938 | 840067 | 840067 | 236.7 | 5.20151 | 11.7713 |
| **F11_OC08** | 1605261 | 1394384 | 1394384 | 228.9 | 6.25601 | 18.4113 |
| **F11_OC09** | 1257331 | 1072980 | 1072980 | 230.4 | 6.03891 | 14.955 |
| **F11_OC10** | 1045387 | 833814 | 833814 | 228.7 | 5.25311 | 11.9952 |
| **F11_OC13** | 967959 | 818617 | 818617 | 228.8 | 5.22622 | 11.7224 |
| **F11_OP03** | 546785 | 485344 | 485344 | 204.6 | 3.7759 | 7.24582 |
| **F11_OP07** | 119543 | 109369 | 109369 | 209.2 | 2.20202 | 2.61242 |
| **F11_OP09** | 245466 | 234519 | 234519 | 198.1 | 3.07831 | 4.45519 |
| **F11_OP11** | 307516 | 273882 | 273882 | 195.6 | 3.03977 | 4.78631 |
| **F11_OP17** | 382707 | 347015 | 347015 | 196.8 | 3.24801 | 5.60989 |
| **F11_OP19** | 388548 | 365916 | 365916 | 196 | 3.4378 | 5.97797 |
| **F12_AP05** | 895928 | 896202 | 896202 | 228.2 | 4.90474 | 11.3338 |
| **F12_AP09** | 808370 | 838348 | 838348 | 239.6 | 6.19048 | 12.3503 |
| **F12_AP13** | 780212 | 833686 | 833686 | 221.2 | 6.46383 | 12.6067 |
| **F12_AP15** | 669364 | 627858 | 627858 | 250.5 | 3.79906 | 7.81319 |
| **F12_AP19** | 911621 | 892657 | 892657 | 238.4 | 4.35768 | 10.4031 |
| **F12_AS01** | 2317012 | 2244216 | 2244216 | 230.9 | 6.03866 | 20.1332 |
| **F12_AS02** | 1998776 | 1985249 | 1985249 | 226.4 | 5.99627 | 19.1344 |
| **F12_AS05** | 1846234 | 1789798 | 1789798 | 224.4 | 5.89426 | 17.7225 |
| **F12_AS10** | 1704297 | 1533154 | 1533154 | 240.9 | 4.8728 | 14.5268 |
| **F12_AS13** | 1537818 | 1415971 | 1415971 | 230.4 | 4.86211 | 13.3371 |
| **F12_AS15** | 1240231 | 1064990 | 1064990 | 248 | 4.22908 | 10.5723 |
| **F12_AS16** | 1047090 | 852338 | 852338 | 241.3 | 3.88449 | 8.81178 |
| **F12_OP01** | 427682 | 393821 | 393821 | 137.8 | 4.33673 | 7.45088 |
| **F12_OP10** | 565537 | 513081 | 513081 | 144.9 | 4.74808 | 8.9205 |
| **F12_OP11** | 246839 | 224949 | 224949 | 142.3 | 3.41641 | 4.93564 |
| **F12_OP12** | 287889 | 247452 | 247452 | 137.9 | 3.28091 | 4.94026 |
| **F12_OP15** | 1071831 | 1014116 | 1014116 | 140.6 | 5.87116 | 15.5688 |
| **F2_AP02** | 797649 | 728988 | 728988 | 241.3 | 3.88407 | 8.48869 |

| | | | | | | |
|---|---|---|---|---|---|---|
| **F2_AP06** | 1527070 | 1419111 | 1419111 | 219.6 | 4.52214 | 13.3359 |
| **F2_AP14** | 1010177 | 940615 | 940615 | 222.1 | 4.13798 | 10.2922 |
| **F2_AP17** | 1337720 | 1173255 | 1173255 | 214.5 | 4.43431 | 12.0494 |
| **F2_OC02** | 594142 | 451646 | 451646 | 233.1 | 3.79913 | 6.87808 |
| **F2_OC03** | 736653 | 613765 | 613765 | 236.4 | 4.6253 | 9.44468 |
| **F2_OC05** | 1103860 | 956181 | 956181 | 239.1 | 5.76861 | 13.6264 |
| **F2_OC06** | 653381 | 509806 | 509806 | 231.9 | 4.06714 | 7.71249 |
| **F2_OP01** | 1661476 | 1252074 | 1252074 | 211.7 | 5.4824 | 14.7785 |
| **F2_OP06** | 1233883 | 948031 | 948031 | 217.9 | 4.92082 | 11.8946 |
| **F4_AP01** | 1598936 | 1507981 | 1507981 | 223.5 | 5.11343 | 14.8799 |
| **F4_AP03** | 1307525 | 1197236 | 1197236 | 224 | 4.24901 | 11.5461 |
| **F4_AP07** | 1119765 | 988547 | 988547 | 237.7 | 3.86974 | 9.73063 |
| **F4_AP09** | 734143 | 611594 | 611594 | 226.9 | 3.30154 | 6.61972 |
| **F4_AP10** | 1674973 | 1590493 | 1590493 | 218.1 | 4.74622 | 14.6223 |
| **F4_AP12** | 1273886 | 1249240 | 1249240 | 215.5 | 4.68994 | 12.6822 |
| **F4_AP13** | 1206985 | 1075906 | 1075906 | 233.3 | 3.96764 | 10.5632 |
| **F4_AP14** | 1481414 | 1460345 | 1460345 | 226.6 | 5.05579 | 14.8663 |
| **F4_AP17** | 1593312 | 1549696 | 1549696 | 221.1 | 5.17089 | 15.5678 |
| **F4_AP18** | 845684 | 808593 | 808593 | 221.9 | 4.20929 | 9.60196 |
| **F4_AP19** | 1356928 | 1263055 | 1263055 | 220.2 | 4.35722 | 12.3209 |
| **F4_AP20** | 924640 | 827059 | 827059 | 226.9 | 3.82147 | 8.91255 |
| **F5_AP03** | 1227575 | 1122622 | 1122622 | 232.2 | 4.24787 | 11.5168 |
| **F5_AP15** | 1975777 | 1898288 | 1898288 | 216.7 | 5.04549 | 16.8015 |
| **F5_AP17** | 2231676 | 2047448 | 2047448 | 224.6 | 4.95294 | 17.3971 |
| **F5_OC01** | 768122 | 674862 | 674862 | 230.8 | 5.24548 | 10.5755 |
| **F5_OC06** | 1273744 | 1146201 | 1146201 | 231.8 | 6.3897 | 16.3854 |
| **F5_OC07** | 616472 | 535015 | 535015 | 237.7 | 4.60855 | 8.53815 |
| **F5_OC09** | 569267 | 456198 | 456198 | 241.5 | 3.85112 | 6.9465 |
| **F5_OP01** | 1016574 | 756079 | 756079 | 225.7 | 4.12276 | 9.06249 |
| **F5_OP05** | 483786 | 328252 | 328252 | 249.8 | 2.83317 | 4.5154 |

| | | | | | | |
|---|---|---|---|---|---|---|
| **F8_AP01** | 1020091 | 937822 | 937822 | 223.1 | 4.13593 | 10.0518 |
| **F8_AP06** | 951444 | 879343 | 879343 | 237 | 4.58715 | 10.3806 |
| **F8_AP12** | 1192988 | 1132421 | 1132421 | 239.4 | 4.959 | 12.6009 |
| **F8_AP16** | 1143749 | 1042653 | 1042653 | 226.5 | 4.20929 | 10.7106 |
| **F8_AP19** | 1007124 | 913465 | 913465 | 227 | 4.11858 | 9.68066 |
| **F8_OC01** | 1320091 | 1073167 | 1073167 | 231.7 | 5.55462 | 14.3966 |
| **F8_OC03** | 1656983 | 1464534 | 1464534 | 228.3 | 6.93138 | 19.693 |
| **F8_OC04** | 1044303 | 828455 | 828455 | 242.3 | 4.93414 | 11.3462 |
| **F8_OC06** | 725973 | 563766 | 563766 | 243.3 | 4.2057 | 8.32205 |
| **F8_OP02** | 647177 | 471245 | 471245 | 237.2 | 3.66323 | 6.5111 |
| **F8_OP05** | 462055 | 313108 | 313108 | 239.5 | 2.90164 | 4.42013 |
| **F8_OP06** | 689420 | 517353 | 517353 | 230.8 | 3.70851 | 6.95601 |
| **F8_OP07** | 594827 | 432407 | 432407 | 237.1 | 3.40053 | 5.8851 |
| **F8_OP10** | 1937239 | 1458389 | 1458389 | 220.3 | 5.66124 | 16.8123 |
| **F8_OP15** | 871122 | 630926 | 630926 | 234.8 | 3.86122 | 7.91515 |
| **F8_OP17** | 1585473 | 1180164 | 1180164 | 214.4 | 5.20087 | 14.0113 |
| **F8_OP19** | 920555 | 691292 | 691292 | 218.8 | 4.4197 | 9.14231 |
| **F9_AP02** | 631406 | 810531 | 810531 | 228.9 | 5.50788 | 12.1656 |
| **F9_AP05** | 570589 | 713166 | 713166 | 240.7 | 5.14185 | 10.7592 |
| **F9_AP06** | 529428 | 666448 | 666448 | 254.4 | 4.87132 | 10.0883 |
| **F9_AP07** | 545079 | 709267 | 709267 | 231 | 5.39481 | 11.1342 |
| **F9_AP08** | 716137 | 955623 | 955623 | 234.3 | 6.6224 | 14.725 |
| **F9_AP09** | 732824 | 977432 | 977432 | 240.5 | 6.6808 | 14.8586 |
| **F9_AP10** | 824920 | 1092098 | 1092098 | 245 | 6.45727 | 15.6388 |
| **F9_AP11** | 653789 | 846391 | 846391 | 243.7 | 5.71772 | 12.5813 |
| **F9_AP12** | 1020413 | 1353085 | 1353085 | 245.6 | 7.06661 | 18.4948 |
| **F9_AP15** | 841358 | 1105137 | 1105137 | 245.4 | 6.43322 | 15.6851 |
| **F9_AS01** | 440292 | 535526 | 535526 | 253.4 | 4.464 | 8.23252 |
| **F9_AS02** | 720293 | 922734 | 922734 | 235.4 | 6.24634 | 13.6603 |
| **F9_AS03** | 781282 | 983233 | 983233 | 222.2 | 6.37482 | 14.8475 |

| | | | | | | |
|---|---|---|---|---|---|---|
| **F9_AS04** | 807729 | 911404 | 911404 | 236.7 | 5.75649 | 13.1109 |
| **F9_AS05** | 471671 | 576541 | 576541 | 226.3 | 4.70812 | 8.97898 |
| **F9_AS06** | 627299 | 771316 | 771316 | 236.1 | 5.41399 | 11.3428 |
| **F9_AS07** | 1839638 | 1861671 | 1861671 | 207.4 | 6.86526 | 20.7867 |
| **F9_AS08** | 1743161 | 1653855 | 1653855 | 217 | 5.48377 | 17.2337 |
| **F9_AS09** | 1510022 | 1514577 | 1514577 | 203.7 | 5.83601 | 16.8538 |
| **F9_AS10** | 2220100 | 2120070 | 2120070 | 197.2 | 6.1356 | 20.5483 |
| **F9_AS11** | 854917 | 1116370 | 1116370 | 237.2 | 6.88978 | 16.1886 |
| **F9_AS12** | 1848304 | 1891634 | 1891634 | 216 | 7.053 | 21.3172 |
| **F9_AS13** | 1116615 | 1043714 | 1043714 | 224.5 | 4.65406 | 11.6476 |
| **F9_AS14** | 1302188 | 1222916 | 1222916 | 237.9 | 4.52276 | 12.4661 |
| **F9_AS15** | 668122 | 598939 | 598939 | 218.4 | 3.76619 | 7.38161 |
| **F9_OC01** | 1820707 | 1568006 | 1568006 | 224.9 | 6.88212 | 20.0458 |
| **F9_OC03** | 365850 | 437074 | 437074 | 256.6 | 5.09308 | 8.63661 |
| **F9_OC04** | 405830 | 474991 | 474991 | 260.8 | 5.2657 | 9.28716 |
| **F9_OC05** | 416587 | 486279 | 486279 | 254 | 5.3445 | 9.40927 |
| **F9_OC06** | 355204 | 413739 | 413739 | 256.6 | 4.85406 | 8.30008 |
| **F9_OC07** | 369416 | 432361 | 432361 | 254 | 5.05752 | 8.70321 |
| **F9_OC08** | 440268 | 550240 | 550240 | 260.8 | 5.98548 | 10.6555 |
| **F9_OC09** | 701156 | 584711 | 584711 | 233.4 | 4.56536 | 8.76832 |
| **F9_OP03** | 169804 | 145356 | 145356 | 196.4 | 2.2703 | 2.95258 |
| **F9_OP08** | 238305 | 209686 | 209686 | 201.2 | 2.74611 | 3.79182 |
| **F9_OP11** | 544165 | 528029 | 528029 | 197.2 | 4.03955 | 8.03294 |
| **F9_OP12** | 657782 | 623824 | 623824 | 191.4 | 4.23682 | 9.31174 |
| **F9_OP14** | 635213 | 442358 | 442358 | 239.3 | 3.36092 | 5.69806 |
| **F9_OP15** | 321814 | 300047 | 300047 | 195.3 | 3.21252 | 5.19589 |
| **LAU_AP01** | 1315188 | 1169896 | 1169896 | 234.1 | 4.02276 | 11.0091 |
| **LAU_AP03** | 1696056 | 1502196 | 1502196 | 227.8 | 4.36188 | 13.0559 |
| **LAU_AP05** | 1909204 | 1783453 | 1783453 | 233.2 | 4.73436 | 15.3781 |
| **LAU_AP07** | 764332 | 604941 | 604941 | 228.5 | 3.30924 | 6.82457 |

| | | | | | | |
|---|---|---|---|---|---|---|
| **LAU_AP08** | 1141138 | 1007219 | 1007219 | 244.1 | 3.95872 | 10.0755 |
| **LAU_AP09** | 1380087 | 1102656 | 1102656 | 226 | 3.96747 | 10.4706 |
| **LAU_AP10** | 1264724 | 1028214 | 1028214 | 230.7 | 4.2972 | 10.1614 |
| **LAU_AP13** | 2352748 | 2185349 | 2185349 | 212.4 | 5.10594 | 17.6043 |
| **LAU_AP14** | 1384131 | 1154018 | 1154018 | 246.6 | 4.03766 | 10.8906 |
| **LAU_AP17** | 1111260 | 1049782 | 1049782 | 232.3 | 4.46751 | 11.3691 |
| **LAU_AP18** | 1014174 | 943193 | 943193 | 240.3 | 4.22628 | 10.0941 |
| **LAU_AP20** | 1843241 | 1701808 | 1701808 | 236.5 | 4.90165 | 15.6115 |
| **LAU_AS01** | 845301 | 840204 | 840204 | 243.1 | 5.51797 | 11.1198 |
| **LAU_AS02** | 1073714 | 1075540 | 1075540 | 232.8 | 6.17003 | 14.2005 |
| **LAU_AS03** | 512115 | 488455 | 488455 | 236.4 | 4.55182 | 7.29771 |
| **LAU_AS04** | 1133117 | 1080793 | 1080793 | 224.4 | 4.72882 | 12.2546 |
| **LAU_AS08** | 1018966 | 943847 | 943847 | 221.1 | 4.41774 | 10.6272 |
| **LAU_AS10** | 649944 | 579061 | 579061 | 250.9 | 3.78487 | 7.30676 |
| **LAU_AS11** | 775637 | 769813 | 769813 | 241.9 | 4.36982 | 9.78024 |
| **LAU_AS12** | 759807 | 706661 | 706661 | 225.5 | 4.14231 | 8.86456 |
| **LAU_AS13** | 921248 | 859408 | 859408 | 219.3 | 4.47659 | 10.54 |
| **LAU_AS14** | 822281 | 716174 | 716174 | 236 | 4.06073 | 8.63028 |
| **LAU_AS17** | 1155585 | 1093686 | 1093686 | 233.1 | 4.95866 | 12.6385 |
| **LAU_AS20** | 1204695 | 1009275 | 1009275 | 224.9 | 4.28207 | 10.8151 |
| **LAU_OC02** | 727253 | 569036 | 569036 | 232.3 | 4.29293 | 8.38579 |
| **LAU_OC03** | 692558 | 548350 | 548350 | 236.8 | 4.21685 | 8.12991 |
| **LAU_OC05** | 784745 | 651047 | 651047 | 218.6 | 4.88802 | 9.76077 |
| **LAU_OC06** | 1017638 | 848268 | 848268 | 229 | 5.13267 | 11.9471 |
| **LAU_OC07** | 845945 | 655964 | 655964 | 238 | 4.52192 | 9.18207 |
| **LAU_OC08** | 1143935 | 889266 | 889266 | 230.2 | 5.21844 | 12.1695 |
| **LAU_OC10** | 1346327 | 1136255 | 1136255 | 234.2 | 5.87247 | 15.4793 |
| **LAU_OC12** | 492551 | 320710 | 320710 | 254.6 | 3.14995 | 5.0914 |
| **LAU_OC15** | 832711 | 639510 | 639510 | 226.5 | 4.58007 | 9.39101 |
| **LAU_OC17** | 608501 | 424547 | 424547 | 235.4 | 3.77056 | 6.73629 |

| | | | | | | |
|---|---|---|---|---|---|---|
| **LAU_OC18** | 1122134 | 938459 | 938459 | 229.6 | 5.37663 | 12.9555 |
| **LAU_OC20** | 1460312 | 1224046 | 1224046 | 227.2 | 5.85151 | 16.0584 |
| **LAU_OP01** | 524445 | 473280 | 473280 | 188.1 | 3.81864 | 7.34905 |
| **LAU_OP04** | 153590 | 137791 | 137791 | 201.7 | 2.29731 | 2.92327 |
| **LAU_OP05** | 267009 | 226596 | 226596 | 192.1 | 2.78729 | 4.16247 |
| **LAU_OP08** | 258214 | 238289 | 238289 | 196.3 | 2.85703 | 4.24364 |
| **LAU_OP09** | 266576 | 225736 | 225736 | 211.2 | 2.51901 | 3.78441 |
| **LAU_OP10** | 330223 | 293274 | 293274 | 203.4 | 2.96144 | 4.80242 |
| **LAU_OP11** | 245037 | 209753 | 209753 | 192.2 | 2.69428 | 3.8743 |
| **LAU_OP12** | 652078 | 588015 | 588015 | 189.1 | 4.10743 | 8.73052 |
| **LAU_OP13** | 258394 | 242512 | 242512 | 199 | 2.88599 | 4.39975 |
| **LAU_OP14** | 242880 | 210424 | 210424 | 200.3 | 2.49475 | 3.59903 |
| **LAU_OP16** | 425586 | 407887 | 407887 | 188 | 3.86934 | 6.83415 |
| **LAU_OP18** | 177746 | 168346 | 168346 | 198.3 | 2.58145 | 3.46854 |
| **MC87_AP01** | 971440 | 906564 | 906564 | 248.1 | 4.26142 | 10.2011 |
| **MC87_AP02** | 1532960 | 1498603 | 1498603 | 228.8 | 5.4237 | 15.4751 |
| **MC87_AP04** | 2522457 | 2460369 | 2460369 | 234.8 | 5.93829 | 22.1341 |
| **MC87_AP06** | 2327778 | 2249112 | 2249112 | 223.5 | 5.6138 | 19.8517 |
| **MC87_AP09** | 2914619 | 2812743 | 2812743 | 220.1 | 5.75614 | 22.5306 |
| **MC87_AP11** | 2366341 | 2286908 | 2286908 | 221.9 | 5.60683 | 19.8967 |
| **MC87_AP14** | 1083962 | 972237 | 972237 | 209.2 | 4.69528 | 11.5853 |
| **MC87_AP15** | 836704 | 784546 | 784546 | 245.3 | 4.17432 | 9.19343 |
| **MC87_AP17** | 1718427 | 1629971 | 1629971 | 220 | 4.98517 | 15.442 |
| **MC87_AP18** | 597387 | 536287 | 536287 | 247.2 | 3.62532 | 7.26129 |
| **MC87_AP19** | 1427232 | 1329958 | 1329958 | 234.5 | 4.85155 | 13.6396 |
| **MC87_AP20** | 1740683 | 1704056 | 1704056 | 218.9 | 5.422 | 16.7587 |
| **MC87_AS01** | 791667 | 682017 | 682017 | 246.1 | 3.7966 | 8.13991 |
| **MC87_AS03** | 919831 | 744367 | 744367 | 236.2 | 3.934 | 8.78895 |
| **MC87_AS05** | 1385057 | 1331739 | 1331739 | 227 | 4.97257 | 14.157 |
| **MC87_AS06** | 1556113 | 1439574 | 1439574 | 240.9 | 4.78931 | 14.4895 |

| | | | | | | |
|---|---|---|---|---|---|---|
| **MC87_AS07** | 907090 | 798555 | 798555 | 242.7 | 4.06425 | 9.1122 |
| **MC87_AS08** | 660956 | 616337 | 616337 | 266.8 | 4.4105 | 8.18056 |
| **MC87_AS11** | 743526 | 723125 | 723125 | 245.4 | 5.05949 | 10.1608 |
| **MC87_AS13** | 2143544 | 1982833 | 1982833 | 232.5 | 5.05015 | 17.6655 |
| **MC87_AS15** | 786169 | 723001 | 723001 | 257.6 | 3.84128 | 8.35545 |
| **MC87_AS17** | 1537657 | 1471993 | 1471993 | 240.2 | 5.32979 | 15.5173 |
| **MC87_AS18** | 2041454 | 1946826 | 1946826 | 240 | 5.62544 | 18.7588 |
| **MC87_AS20** | 1958415 | 1839350 | 1839350 | 243.3 | 5.07791 | 16.6486 |
| **MC87_OB01** | 281137 | 268257 | 268257 | 198.6 | 3.11072 | 4.89588 |
| **MC87_OB02** | 216223 | 199214 | 199214 | 196.8 | 2.56957 | 3.73134 |
| **MC87_OB03** | 365876 | 319896 | 319896 | 192.4 | 3.19272 | 5.34471 |
| **MC87_OB06** | 169473 | 148847 | 148847 | 191.1 | 2.38333 | 3.17236 |
| **MC87_OB08** | 393766 | 357927 | 357927 | 193.4 | 3.40168 | 5.98709 |
| **MC87_OB10** | 266025 | 231642 | 231642 | 196.3 | 2.6623 | 4.08585 |
| **MC87_OB11** | 235375 | 210252 | 210252 | 189.9 | 2.69705 | 3.87622 |
| **MC87_OB14** | 549615 | 448319 | 448319 | 214.6 | 3.31443 | 6.17391 |
| **MC87_OB15** | 165837 | 147361 | 147361 | 201.2 | 2.29472 | 3.03319 |
| **MC87_OB16** | 403741 | 316723 | 316723 | 218.2 | 2.93782 | 4.84403 |
| **MC87_OB18** | 441912 | 360462 | 360462 | 217.7 | 3.04592 | 5.36217 |
| **MC87_OB20** | 369058 | 316599 | 316599 | 194.7 | 3.13877 | 5.22469 |
| **MCM_AP01** | 1535791 | 1479169 | 1479169 | 224 | 5.17256 | 15.0365 |
| **MCM_AP03** | 2124768 | 1992500 | 1992500 | 225.7 | 5.35785 | 16.8429 |
| **MCM_AP04** | 966013 | 822621 | 822621 | 231.1 | 5.33537 | 10.9707 |
| **MCM_AP06** | 920592 | 871232 | 871232 | 218.9 | 4.6461 | 10.4908 |
| **MCM_AP07** | 2013582 | 1905638 | 1905638 | 218.8 | 5.36277 | 17.6949 |
| **MCM_AP10** | 2246947 | 1956118 | 1956118 | 211.2 | 5.43905 | 18.4612 |
| **MCM_AP13** | 2346482 | 2181503 | 2181503 | 213.7 | 5.55875 | 19.3499 |
| **MCM_AP14** | 1273129 | 1114518 | 1114518 | 239.4 | 4.65686 | 11.7483 |
| **MCM_AP15** | 1407827 | 1371899 | 1371899 | 223.1 | 4.93686 | 14.1694 |
| **MCM_AP17** | 1238826 | 1124550 | 1124550 | 245.4 | 4.18162 | 10.885 |

| | | | | | | |
|---|---|---|---|---|---|---|
| **MCM_AP19** | 1274332 | 1170177 | 1170177 | 241 | 4.78404 | 12.3598 |
| **MCM_AP20** | 2057354 | 1953329 | 1953329 | 218.7 | 5.17925 | 17.5303 |
| **MCM_AS01** | 1866972 | 1737320 | 1737320 | 231.1 | 4.96566 | 16.0631 |
| **MCM_AS02** | 2141392 | 2083407 | 2083407 | 226.3 | 5.25926 | 18.3504 |
| **MCM_AS03** | 774588 | 707755 | 707755 | 218.8 | 4.23016 | 9.71814 |
| **MCM_AS04** | 1396103 | 1305379 | 1305379 | 239.9 | 4.86092 | 13.4507 |
| **MCM_AS05** | 1333655 | 1225986 | 1225986 | 239.4 | 4.55381 | 12.3779 |
| **MCM_AS08** | 1349067 | 1248595 | 1248595 | 236.6 | 4.5773 | 12.7067 |
| **MCM_AS10** | 1816104 | 1797828 | 1797828 | 240.3 | 4.94375 | 16.8649 |
| **MCM_AS12** | 1341985 | 1220425 | 1220425 | 247.2 | 4.54347 | 12.0716 |
| **MCM_AS14** | 628690 | 491773 | 491773 | 229.6 | 4.92173 | 7.63974 |
| **MCM_AS15** | 1786112 | 1732932 | 1732932 | 231.3 | 5.35328 | 16.7645 |
| **MCM_AS17** | 1579391 | 1515479 | 1515479 | 240.9 | 5.24524 | 15.2499 |
| **MCM_AS20** | 1888634 | 1935619 | 1935619 | 226.4 | 5.46318 | 18.6461 |
| **MCM_OP01** | 359940 | 320448 | 320448 | 184.2 | 3.12656 | 5.19526 |
| **MCM_OP02** | 769031 | 713028 | 713028 | 191.2 | 4.41174 | 10.4297 |
| **MCM_OP04** | 220480 | 191669 | 191669 | 199.2 | 2.50928 | 3.50281 |
| **MCM_OP06** | 486522 | 450401 | 450401 | 188.7 | 3.86985 | 7.28312 |
| **MCM_OP09** | 425627 | 374220 | 374220 | 182.5 | 3.74308 | 6.43207 |
| **MCM_OP10** | 579400 | 506744 | 506744 | 143.1 | 4.88486 | 9.2007 |
| **MCM_OP12** | 394081 | 378455 | 378455 | 146 | 4.20671 | 7.31227 |
| **MCM_OP15** | 400541 | 390091 | 390091 | 148.7 | 4.40156 | 7.58013 |
| **MCM_OP16** | 673213 | 627453 | 627453 | 147.2 | 5.31233 | 11.2445 |
| **MCM_OP17** | 366683 | 335729 | 335729 | 142.7 | 3.88665 | 6.38348 |
| **MCM_OP18** | 189010 | 171533 | 171533 | 144.8 | 3.01023 | 4.00422 |
| **MCM_OP20** | 489801 | 458386 | 458386 | 145.3 | 4.53203 | 8.33893 |
| **PAR_AP01** | 1715844 | 1620851 | 1620851 | 227.7 | 4.92987 | 15.3122 |
| **PAR_AP02** | 1868716 | 1793580 | 1793580 | 223.5 | 5.24814 | 17.1734 |
| **PAR_AP04** | 1583087 | 1427210 | 1427210 | 246.8 | 4.42548 | 12.9332 |
| **PAR_AP06** | 2488710 | 2397285 | 2397285 | 221.9 | 5.53552 | 20.4912 |

| | | | | | | |
|---|---|---|---|---|---|---|
| **PAR_AP10** | 1105557 | 1015243 | 1015243 | 233.9 | 4.41302 | 11.474 |
| **PAR_AP11** | 919851 | 870306 | 870306 | 221.5 | 4.25276 | 10.1533 |
| **PAR_AP12** | 483882 | 454514 | 454514 | 225.7 | 4.04178 | 7.25148 |
| **PAR_AP14** | 1855685 | 1763023 | 1763023 | 227.3 | 5.23511 | 16.9009 |
| **PAR_AP18** | 1347827 | 1239437 | 1239437 | 236 | 4.46099 | 12.3132 |
| **PAR_AP20** | 1065156 | 964838 | 964838 | 229 | 4.56142 | 10.8438 |
| **PAR_AP22** | 891498 | 819247 | 819247 | 256 | 3.94351 | 8.88877 |
| **PAR_AP25** | 769560 | 729704 | 729704 | 217.5 | 4.4181 | 9.75231 |
| **PAR_AS01** | 1239916 | 1189425 | 1189425 | 243.9 | 4.19441 | 11.8916 |
| **PAR_AS03** | 1163551 | 977877 | 977877 | 229.2 | 4.41099 | 11.1606 |
| **PAR_AS04** | 1030340 | 953058 | 953058 | 228.3 | 4.4465 | 11.3062 |
| **PAR_AS07** | 1177038 | 1111249 | 1111249 | 242.8 | 4.65944 | 12.0567 |
| **PAR_AS09** | 2023231 | 1918737 | 1918737 | 233.6 | 5.15151 | 16.9901 |
| **PAR_AS11** | 1243569 | 1191670 | 1191670 | 232.7 | 4.48469 | 12.3592 |
| **PAR_AS12** | 1224399 | 1170651 | 1170651 | 237.3 | 4.86444 | 12.918 |
| **PAR_AS17** | 1931383 | 1762011 | 1762011 | 241.1 | 4.91493 | 15.6445 |
| **PAR_AS19** | 1495296 | 1336365 | 1336365 | 247.1 | 4.55197 | 12.8225 |
| **PAR_AS21** | 838751 | 732487 | 732487 | 269.8 | 3.5037 | 7.60989 |
| **PAR_AS24** | 933408 | 838916 | 838916 | 249.3 | 3.96929 | 9.2924 |
| **PAR_AS25** | 1912776 | 1788859 | 1788859 | 231.7 | 5.15392 | 16.6855 |
| **PAR_OB03** | 337346 | 308597 | 308597 | 143.7 | 3.74937 | 6.03748 |
| **PAR_OB05** | 211110 | 198662 | 198662 | 146.2 | 3.22736 | 4.52372 |
| **PAR_OB06** | 508915 | 425911 | 425911 | 184.1 | 3.47024 | 6.12272 |
| **PAR_OB10** | 248426 | 224359 | 224359 | 139.9 | 3.33064 | 4.89093 |
| **PAR_OB12** | 307725 | 277682 | 277682 | 142.1 | 3.59073 | 5.48423 |
| **PAR_OB14** | 189680 | 168833 | 168833 | 143.1 | 3.05 | 4.09933 |
| **PAR_OB22** | 413179 | 401674 | 401674 | 144.2 | 4.49084 | 7.89877 |
| **PAR_OB23** | 241771 | 216467 | 216467 | 139.5 | 3.25076 | 4.72324 |
| **PAR_OB24** | 292063 | 271408 | 271408 | 142.2 | 3.92021 | 5.90336 |
| **PAR_OB25** | 224103 | 197438 | 197438 | 139.8 | 3.43967 | 4.97779 |

| | | | | | | |
|---|---|---|---|---|---|---|
| **PAR_OB26** | 202669 | 175803 | 175803 | 138.3 | 3.03265 | 4.30528 |
| **PAR_OB27** | 274879 | 252445 | 252445 | 142.6 | 3.5839 | 5.27548 |
| **PAR_OC01** | 875698 | 682038 | 682038 | 225.6 | 4.75289 | 10.1664 |
| **PAR_OC03** | 273852 | 237692 | 237692 | 211.6 | 3.75604 | 5.55365 |
| **PAR_OC04** | 116965 | 103646 | 103646 | 211.6 | 3.11978 | 3.93323 |
| **PAR_OC05** | 1110071 | 810253 | 810253 | 214 | 4.59525 | 10.1239 |
| **PAR_OC09** | 237979 | 171221 | 171221 | 214.5 | 3.0033 | 4.44144 |
| **PAR_OC12** | 516479 | 382104 | 382104 | 226 | 3.46604 | 5.71912 |
| **PAR_OC14** | 938851 | 726044 | 726044 | 222.7 | 4.42341 | 9.59759 |
| **PAR_OC17** | 684119 | 483039 | 483039 | 211.5 | 3.90733 | 6.84905 |
| **PAR_OC18** | 67169 | 47504 | 47504 | 217.7 | 2.30643 | 2.8666 |
| **PAR_OC20** | 1635954 | 1274034 | 1274034 | 205.5 | 5.56344 | 15.3418 |
| **PAR_OC21** | 1097347 | 821025 | 821025 | 212.8 | 4.60779 | 10.4263 |
| **PAR_OC22** | 1244719 | 926739 | 926739 | 219.4 | 4.89819 | 11.5645 |
| **SPA_AP01** | 1987640 | 1861906 | 1861906 | 240.1 | 5.11558 | 17.2547 |
| **SPA_AP02** | 1257745 | 1172694 | 1172694 | 241.3 | 4.49105 | 11.9356 |
| **SPA_AP04** | 864152 | 820037 | 820037 | 232.7 | 4.25483 | 9.85296 |
| **SPA_AP06** | 2024076 | 1970880 | 1970880 | 224.7 | 5.12963 | 17.5535 |
| **SPA_AP08** | 2689602 | 2874082 | 2874082 | 190.9 | 6.60164 | 26.362 |
| **SPA_AP09** | 98400 | 108237 | 108237 | 156.5 | 2.98102 | 3.38749 |
| **SPA_AP11** | 1030153 | 963728 | 963728 | 232.8 | 4.60314 | 11.3105 |
| **SPA_AP12** | 1147106 | 1094699 | 1094699 | 227.5 | 4.5564 | 11.7732 |
| **SPA_AP14** | 2223789 | 2151506 | 2151506 | 224.3 | 5.39569 | 18.7299 |
| **SPA_AP16** | 1130288 | 1008297 | 1008297 | 239.9 | 3.97926 | 9.85538 |
| **SPA_AP18** | 2248899 | 2116808 | 2116808 | 228.3 | 5.47184 | 19.075 |
| **SPA_AP20** | 1152183 | 1074688 | 1074688 | 226.1 | 4.40295 | 11.9117 |
| **SPA_AS01** | 2474192 | 2362520 | 2362520 | 229.6 | 5.39913 | 20.014 |
| **SPA_AS02** | 1154974 | 1076735 | 1076735 | 235.8 | 4.21087 | 11.4408 |
| **SPA_AS03** | 1346473 | 1326302 | 1326302 | 237.6 | 5.04891 | 14.1964 |
| **SPA_AS05** | 1650211 | 1548453 | 1548453 | 233.4 | 5.1739 | 15.9629 |

| | | | | | | |
|---|---|---|---|---|---|---|
| **SPA_AS07** | 1305897 | 1217807 | 1217807 | 227.6 | 4.66915 | 12.912 |
| **SPA_AS10** | 1559697 | 1467887 | 1467887 | 228.4 | 5.16867 | 15.2 |
| **SPA_AS11** | 1114123 | 1052926 | 1052926 | 236.6 | 4.7034 | 12.0628 |
| **SPA_AS13** | 1490580 | 1444554 | 1444554 | 239.6 | 5.13322 | 15.2727 |
| **SPA_AS16** | 3137802 | 2925315 | 2925315 | 231.4 | 5.88624 | 23.4431 |
| **SPA_AS18** | 2368186 | 2301792 | 2301792 | 230 | 5.56493 | 20.6181 |
| **SPA_AS19** | 825982 | 771891 | 771891 | 237 | 4.4163 | 9.93251 |
| **SPA_AS20** | 767269 | 620186 | 620186 | 286.5 | 3.03865 | 6.40793 |
| **SPA_OC01** | 986504 | 729447 | 729447 | 215.3 | 4.38676 | 9.33119 |
| **SPA_OC03** | 663063 | 473627 | 473627 | 228.7 | 3.58415 | 6.35305 |
| **SPA_OC04** | 1013420 | 766444 | 766444 | 214.3 | 4.55075 | 9.88576 |
| **SPA_OC06** | 1045226 | 785422 | 785422 | 224 | 4.52282 | 10.0043 |
| **SPA_OC08** | 1145023 | 888339 | 888339 | 214.1 | 4.89324 | 11.3113 |
| **SPA_OC10** | 830147 | 607602 | 607602 | 227 | 4.0803 | 7.87039 |
| **SPA_OC11** | 943640 | 706692 | 706692 | 219.5 | 4.51274 | 9.20624 |
| **SPA_OC12** | 971930 | 741042 | 741042 | 224.5 | 4.49017 | 9.56701 |
| **SPA_OC14** | 1089409 | 794897 | 794897 | 208.6 | 4.52885 | 9.83726 |
| **SPA_OC16** | 1005779 | 628656 | 628656 | 208.7 | 4.19353 | 8.38671 |
| **SPA_OC19** | 1465154 | 1083097 | 1083097 | 214.3 | 5.12288 | 12.8532 |
| **SPA_OC20** | 1439143 | 1124497 | 1124497 | 211.5 | 5.23203 | 13.7459 |
| **SPA_OP01** | 450524 | 406584 | 406584 | 144.5 | 4.06195 | 7.09407 |
| **SPA_OP02** | 411828 | 391194 | 391194 | 145.8 | 4.28653 | 7.53742 |
| **SPA_OP03** | 342303 | 320546 | 320546 | 142.7 | 4.00569 | 6.23845 |
| **SPA_OP07** | 535435 | 494724 | 494724 | 141.9 | 4.72275 | 8.78355 |
| **SPA_OP09** | 830378 | 787564 | 787564 | 142.9 | 5.76676 | 13.2647 |
| **SPA_OP10** | 688474 | 653471 | 653471 | 146.1 | 5.1602 | 10.9908 |
| **SPA_OP11** | 999231 | 928416 | 928416 | 143.6 | 5.86486 | 14.6085 |
| **SPA_OP13** | 318591 | 294204 | 294204 | 154.3 | 3.71161 | 5.85467 |
| **SPA_OP14** | 806465 | 762841 | 762841 | 144.6 | 5.3719 | 12.3693 |
| **SPA_OP17** | 339798 | 305541 | 305541 | 143.2 | 3.54339 | 5.6485 |

| | | | | | | |
|---|---|---|---|---|---|---|
| **SPA_OP18** | 359688 | 340842 | 340842 | 152.6 | 3.99764 | 6.5898 |
| **SPA_OP19** | 732318 | 654410 | 654410 | 147.4 | 5.16392 | 10.8054 |
| **SPB_AP02** | 1961391 | 1952135 | 1952135 | 231.7 | 5.38957 | 18.8808 |
| **SPB_AP03** | 1282504 | 1273877 | 1273877 | 239.2 | 5.85737 | 15.1429 |
| **SPB_AP06** | 911075 | 907012 | 907012 | 250.5 | 5.38219 | 11.9338 |
| **SPB_AP08** | 821989 | 790377 | 790377 | 249.5 | 4.87233 | 10.2459 |
| **SPB_AP09** | 1029866 | 1003463 | 1003463 | 227.7 | 5.81555 | 13.0988 |
| **SPB_AP12** | 1419989 | 1429749 | 1429749 | 233.6 | 6.02055 | 16.3098 |
| **SPB_AP13** | 1510999 | 1530018 | 1530018 | 233.4 | 6.18092 | 17.6223 |
| **SPB_AP15** | 753395 | 684744 | 684744 | 230 | 4.22615 | 9.22813 |
| **SPB_AP17** | 1140489 | 1083511 | 1083511 | 234.1 | 4.57949 | 11.9617 |
| **SPB_AP18** | 949471 | 858745 | 858745 | 244.1 | 4.05347 | 9.92294 |
| **SPB_AP19** | 1255812 | 1124749 | 1124749 | 222.2 | 4.76644 | 12.7506 |
| **SPB_AP20** | 1559643 | 1570661 | 1570661 | 236 | 5.92118 | 17.3155 |
| **SPB_AS01** | 448319 | 371575 | 371575 | 291.2 | 2.70275 | 4.53986 |
| **SPB_AS02** | 576169 | 465051 | 465051 | 279.6 | 2.78997 | 5.33518 |
| **SPB_AS05** | 428054 | 359654 | 359654 | 290.9 | 2.86832 | 4.98217 |
| **SPB_AS07** | 744830 | 636467 | 636467 | 277.7 | 3.40155 | 7.30949 |
| **SPB_AS08** | 494025 | 410706 | 410706 | 287.2 | 2.73038 | 4.97489 |
| **SPB_AS10** | 770854 | 673803 | 673803 | 248.9 | 3.36157 | 7.39769 |
| **SPB_AS14** | 178499 | 156189 | 156189 | 260.1 | 2.82652 | 4.2518 |
| **SPB_AS15** | 1018311 | 845123 | 845123 | 239.5 | 4.01815 | 8.90939 |
| **SPB_AS16** | 1800278 | 1660946 | 1660946 | 226.2 | 5.32796 | 16.3462 |
| **SPB_AS18** | 1752113 | 1651031 | 1651031 | 235 | 5.74309 | 16.4685 |
| **SPB_AS19** | 1125444 | 1025211 | 1025211 | 234.2 | 5.53331 | 12.3949 |
| **SPB_AS20** | 1861898 | 1766507 | 1766507 | 231.5 | 5.73234 | 17.5886 |
| **SPB_OC01** | 482038 | 339078 | 339078 | 236.1 | 3.10315 | 4.96283 |
| **SPB_OC03** | 1160824 | 824913 | 824913 | 214.2 | 4.68403 | 10.19 |
| **SPB_OC05** | 677153 | 519989 | 519989 | 213 | 4.23427 | 8.04691 |
| **SPB_OC06** | 886418 | 621642 | 621642 | 243.9 | 3.84687 | 7.7439 |

| | | | | | | |
|---|---|---|---|---|---|---|
| **SPB_OC07** | 846908 | 610045 | 610045 | 220.9 | 3.97123 | 7.91504 |
| **SPB_OC09** | 1649401 | 1259252 | 1259252 | 209.7 | 5.43062 | 14.9711 |
| **SPB_OC11** | 1157643 | 881237 | 881237 | 210.2 | 4.73629 | 11.0625 |
| **SPB_OC13** | 1455158 | 1114065 | 1114065 | 218.3 | 5.13713 | 13.4308 |
| **SPB_OC14** | 1123551 | 829510 | 829510 | 221.1 | 4.58502 | 10.4044 |
| **SPB_OC19** | 1493980 | 1123652 | 1123652 | 210.8 | 5.10676 | 13.4529 |
| **SPB_OC20** | 989801 | 725419 | 725419 | 213.5 | 4.619 | 9.48437 |
| **SPB_OC21** | 1298385 | 966016 | 966016 | 223.7 | 4.90069 | 12.0869 |
| **SPB_OP01** | 504324 | 470704 | 470704 | 140.7 | 4.52705 | 8.56033 |
| **SPB_OP02** | 463081 | 437653 | 437653 | 146.7 | 4.40896 | 7.99176 |
| **SPB_OP05** | 219719 | 196083 | 196083 | 147.5 | 3.15017 | 4.43925 |
| **SPB_OP06** | 586591 | 543145 | 543145 | 146.2 | 4.9192 | 9.67912 |
| **SPB_OP07** | 451070 | 435346 | 435346 | 150.9 | 4.42946 | 8.11499 |
| **SPB_OP08** | 515276 | 472503 | 472503 | 146.7 | 4.61126 | 8.49976 |
| **SPB_OP09** | 122000 | 110349 | 110349 | 141.7 | 2.50306 | 3.18483 |
| **SPB_OP12** | 143535 | 123472 | 123472 | 139.9 | 2.70459 | 3.46341 |
| **SPB_OP15** | 707757 | 652249 | 652249 | 144 | 5.27418 | 10.9067 |
| **SPB_OP16** | 384246 | 343890 | 343890 | 144.4 | 3.96539 | 6.29427 |
| **SPB_OP17** | 393510 | 364409 | 364409 | 144.4 | 4.05324 | 6.76719 |
| **SPB_OP20** | 329152 | 290471 | 290471 | 145.1 | 3.50626 | 5.44646 |

**Methodological consideration on the differences between *Astrocaryum spp.* and *Oenocarpus spp.*** Even if all species were submitted to a standardized experimental protocol and bioinformatics pipeline, the total number of reads, mapping coverage, and overall success rate of the capture experiment was higher in *Astrocaryum spp.* than in *Oenocarpus spp.* libraries, resulting in many SNPs detected in the former. The exact cause remains uncertain in the absence of marked differences in gDNA and library quantity and quality between the different species. This may alternatively be due to a greater phylogenetic proximity between *Astrocaryum* and *Elaeis* (both belonging to the *Cocoseae* tribe) from which the capture kit has been designed, than between *Oenocarpus* and *Elaeis* (the former belonging from the *Areceae* tribe). It is nonetheless important to notice that we cannot assert that *Astrocaryum sp.* would be more polymorphic than *Oenocarpus sp.* from this rough number of SNPs because of a potential recruitment bias. Nevertheless, each genus being analyzed independently from each other, these differences did not biased neither the bioinformatics treatment nor subsequent population genetics analyses.
