## Supplementary material 4 for "Natural and human-mediated drivers of microevolution in Neotropical palms: a historical genomics approach": outF2_AP02.1_pf_trimmed_masked_sickled_fastqc.html

outF2\_AP02.1\_pf\_trimmed\_masked\_sickled.fastq FastQC Report 

FastQC Report

mar. 25 juin 2019  
outF2\_AP02.1\_pf\_trimmed\_masked\_sickled.fastq

### Summary

- Basic Statistics
- Per base sequence quality
- Per tile sequence quality
- Per sequence quality scores
- Per base sequence content
- Per sequence GC content
- Per base N content
- Sequence Length Distribution
- Sequence Duplication Levels
- Overrepresented sequences
- Adapter Content
- Kmer Content

### Basic Statistics

| Measure | Value |
| --- | --- |
| Filename | outF2\_AP02.1\_pf\_trimmed\_masked\_sickled.fastq |
| File type | Conventional base calls |
| Encoding | Sanger / Illumina 1.9 |
| Total Sequences | 622793 |
| Sequences flagged as poor quality | 0 |
| Sequence length | 100-131 |
| %GC | 43 |

### Per base sequence quality

### Per tile sequence quality

### Per sequence quality scores

### Per base sequence content

### Per sequence GC content

### Per base N content

### Sequence Length Distribution

### Sequence Duplication Levels

### Overrepresented sequences

No overrepresented sequences

### Adapter Content

### Kmer Content

| Sequence | Count | PValue | Obs/Exp Max | Max Obs/Exp Position |
| --- | --- | --- | --- | --- |
| GGCCCGT | 75 | 0.0038374711 | 20.544844 | 46-47 |
| CACGTCT | 225 | 0.008309264 | 10.57777 | 122-123 |
| CGATCTA | 490 | 0.0033462918 | 9.8491335 | 5 |
| TGTAAGA | 330 | 0.003314267 | 8.425798 | 48-49 |
| TCGGAAG | 675 | 5.735281E-5 | 7.0518475 | 122-123 |
| TTAAAGA | 985 | 0.0028174315 | 6.716614 | 1 |
| CCTTCTA | 1170 | 4.7384674E-8 | 6.4224205 | 124-125 |
| CGGAAGA | 605 | 0.006401021 | 6.2101097 | 124-125 |
| CTTCTAG | 1090 | 3.2222582E-5 | 5.5460134 | 118-119 |
| TAGATCG | 1025 | 3.7855582E-4 | 5.3073134 | 122-123 |

Produced by FastQC (version 0.11.5)
