## Supplementary material 4 for "Natural and human-mediated drivers of microevolution in Neotropical palms: a historical genomics approach": outF2_AP02.2_pf_trimmed_masked_sickled_fastqc.html

outF2\_AP02.2\_pf\_trimmed\_masked\_sickled.fastq FastQC Report 

FastQC Report

mar. 25 juin 2019  
outF2\_AP02.2\_pf\_trimmed\_masked\_sickled.fastq

### Summary

- Basic Statistics
- Per base sequence quality
- Per tile sequence quality
- Per sequence quality scores
- Per base sequence content
- Per sequence GC content
- Per base N content
- Sequence Length Distribution
- Sequence Duplication Levels
- Overrepresented sequences
- Adapter Content
- Kmer Content

### Basic Statistics

| Measure | Value |
| --- | --- |
| Filename | outF2\_AP02.2\_pf\_trimmed\_masked\_sickled.fastq |
| File type | Conventional base calls |
| Encoding | Sanger / Illumina 1.9 |
| Total Sequences | 622793 |
| Sequences flagged as poor quality | 0 |
| Sequence length | 100-131 |
| %GC | 44 |

### Per base sequence quality

### Per tile sequence quality

### Per sequence quality scores

### Per base sequence content

### Per sequence GC content

### Per base N content

### Sequence Length Distribution

### Sequence Duplication Levels

### Overrepresented sequences

No overrepresented sequences

### Adapter Content

### Kmer Content

| Sequence | Count | PValue | Obs/Exp Max | Max Obs/Exp Position |
| --- | --- | --- | --- | --- |
| TAGAAGG | 565 | 7.1664545E-8 | 13.025982 | 6 |
| CGATCTA | 460 | 2.248762E-5 | 12.1568165 | 1 |
| TCTAGAA | 480 | 0.004839838 | 9.418617 | 4 |
| CCTTCTA | 990 | 3.8926373E-10 | 8.686729 | 124-125 |
| CTAGAAG | 525 | 0.009951859 | 8.626731 | 5 |
| ATCTAGA | 725 | 2.1229085E-4 | 8.558161 | 3 |
| TCGGAAG | 575 | 7.508061E-5 | 8.371366 | 122-123 |
| CGGAAGA | 570 | 0.0043679033 | 7.184513 | 124-125 |
| TAGATCG | 895 | 4.7191934E-6 | 7.1710014 | 122-123 |
| CTTCTAG | 930 | 1.0523973E-4 | 6.2249165 | 118-119 |
| TCTAGAT | 1175 | 2.9465014E-5 | 5.92493 | 124-125 |
| TTCTAGA | 995 | 0.0010213361 | 5.6440115 | 122-123 |
| ATCGGAA | 1095 | 4.3461274E-4 | 5.609825 | 124-125 |
| GCCTTCT | 950 | 0.002839572 | 5.60392 | 124-125 |
| GAAGAGC | 1015 | 0.006195641 | 5.2450485 | 124-125 |

Produced by FastQC (version 0.11.5)
