## Supplementary material 4 for "Natural and human-mediated drivers of microevolution in Neotropical palms: a historical genomics approach": outF2_AP06.1_pf_trimmed_masked_sickled_fastqc.html

outF2\_AP06.1\_pf\_trimmed\_masked\_sickled.fastq FastQC Report 

FastQC Report

mar. 25 juin 2019  
outF2\_AP06.1\_pf\_trimmed\_masked\_sickled.fastq

### Summary

- Basic Statistics
- Per base sequence quality
- Per tile sequence quality
- Per sequence quality scores
- Per base sequence content
- Per sequence GC content
- Per base N content
- Sequence Length Distribution
- Sequence Duplication Levels
- Overrepresented sequences
- Adapter Content
- Kmer Content

### Basic Statistics

| Measure | Value |
| --- | --- |
| Filename | outF2\_AP06.1\_pf\_trimmed\_masked\_sickled.fastq |
| File type | Conventional base calls |
| Encoding | Sanger / Illumina 1.9 |
| Total Sequences | 1223601 |
| Sequences flagged as poor quality | 0 |
| Sequence length | 100-131 |
| %GC | 44 |

### Per base sequence quality

### Per tile sequence quality

### Per sequence quality scores

### Per base sequence content

### Per sequence GC content

### Per base N content

### Sequence Length Distribution

### Sequence Duplication Levels

### Overrepresented sequences

No overrepresented sequences

### Adapter Content

### Kmer Content

| Sequence | Count | PValue | Obs/Exp Max | Max Obs/Exp Position |
| --- | --- | --- | --- | --- |
| CTAGGCG | 285 | 0.0 | 49.87364 | 5 |
| TAGGCGA | 360 | 0.0 | 39.484962 | 6 |
| TCTAGGC | 550 | 0.0 | 26.888735 | 4 |
| ATCTAGG | 1160 | 0.0 | 13.755582 | 3 |
| AGGCGAG | 520 | 3.1744876E-6 | 12.53416 | 7 |
| CGATCTA | 1190 | 0.0 | 12.35638 | 1 |
| GCACACG | 460 | 1.5995283E-7 | 10.903312 | 124-125 |
| CGCCTAG | 1515 | 0.0 | 9.846713 | 118-119 |
| TCGCCTA | 1720 | 0.0 | 9.789435 | 124-125 |
| CACACGT | 445 | 1.3941006E-5 | 9.66072 | 124-125 |
| ACACGTC | 395 | 6.261947E-5 | 9.576647 | 116-117 |
| GCCTAGA | 1585 | 0.0 | 9.279468 | 120-121 |
| GATCTAG | 1545 | 1.8189894E-12 | 9.170649 | 2 |
| CGGAAGA | 1090 | 1.8189894E-12 | 8.437369 | 122-123 |
| CGTCTGA | 500 | 5.171941E-4 | 7.8815365 | 124-125 |
| CCTAGAT | 1955 | 0.0 | 7.8815017 | 120-121 |
| TAGATCG | 2100 | 0.0 | 7.5797253 | 122-123 |
| GTCGCCT | 645 | 1.3230702E-4 | 7.220576 | 124-125 |
| TCGGAAG | 1210 | 8.913048E-10 | 7.0159435 | 122-123 |
| GTCTAAG | 440 | 0.0055895266 | 7.0057416 | 74-75 |

Produced by FastQC (version 0.11.5)
