## Supplementary material 4 for "Natural and human-mediated drivers of microevolution in Neotropical palms: a historical genomics approach": outF2_AP06.2_pf_trimmed_masked_sickled_fastqc.html

outF2\_AP06.2\_pf\_trimmed\_masked\_sickled.fastq FastQC Report 

FastQC Report

mar. 25 juin 2019  
outF2\_AP06.2\_pf\_trimmed\_masked\_sickled.fastq

### Summary

- Basic Statistics
- Per base sequence quality
- Per tile sequence quality
- Per sequence quality scores
- Per base sequence content
- Per sequence GC content
- Per base N content
- Sequence Length Distribution
- Sequence Duplication Levels
- Overrepresented sequences
- Adapter Content
- Kmer Content

### Basic Statistics

| Measure | Value |
| --- | --- |
| Filename | outF2\_AP06.2\_pf\_trimmed\_masked\_sickled.fastq |
| File type | Conventional base calls |
| Encoding | Sanger / Illumina 1.9 |
| Total Sequences | 1223601 |
| Sequences flagged as poor quality | 0 |
| Sequence length | 100-131 |
| %GC | 44 |

### Per base sequence quality

### Per tile sequence quality

### Per sequence quality scores

### Per base sequence content

### Per sequence GC content

### Per base N content

### Sequence Length Distribution

### Sequence Duplication Levels

### Overrepresented sequences

No overrepresented sequences

### Adapter Content

### Kmer Content

| Sequence | Count | PValue | Obs/Exp Max | Max Obs/Exp Position |
| --- | --- | --- | --- | --- |
| CTAGGCG | 215 | 0.0 | 39.142513 | 5 |
| TAGGCGA | 330 | 0.0 | 27.24334 | 6 |
| TCTAGGC | 390 | 1.8189894E-12 | 21.556171 | 4 |
| CGTGTAG | 225 | 0.0030252105 | 12.113923 | 118-119 |
| CGCCTAG | 1490 | 0.0 | 11.498355 | 118-119 |
| GAGCGTC | 520 | 8.132156E-8 | 11.44929 | 124-125 |
| TCGCCTA | 1685 | 0.0 | 11.104695 | 124-125 |
| ATCTAGG | 830 | 8.993084E-9 | 10.804523 | 3 |
| GTCGTGT | 255 | 0.009096941 | 10.449875 | 116-117 |
| AAGAGCG | 785 | 2.4374458E-10 | 10.292901 | 124-125 |
| AGAGCGT | 500 | 7.253804E-6 | 10.206224 | 124-125 |
| GCCTAGA | 1705 | 0.0 | 10.091348 | 120-121 |
| ATTCGCC | 470 | 4.1018142E-5 | 9.952878 | 124-125 |
| TTTCGCC | 345 | 0.0033176923 | 9.861087 | 124-125 |
| AATCGCC | 430 | 2.440357E-4 | 9.615309 | 122-123 |
| TAGATCG | 2055 | 0.0 | 9.4562235 | 122-123 |
| CGGAAGA | 1145 | 0.0 | 9.388572 | 122-123 |
| CGATCTA | 980 | 1.3503814E-7 | 9.069982 | 1 |
| CCTAGAT | 2090 | 0.0 | 8.806769 | 120-121 |
| AGCGTCG | 455 | 0.0033658082 | 8.411723 | 124-125 |

Produced by FastQC (version 0.11.5)
