## Supplementary material 4 for "Natural and human-mediated drivers of microevolution in Neotropical palms: a historical genomics approach": outF2_AP14.1_pf_trimmed_masked_sickled_fastqc.html

outF2\_AP14.1\_pf\_trimmed\_masked\_sickled.fastq FastQC Report 

FastQC Report

mar. 25 juin 2019  
outF2\_AP14.1\_pf\_trimmed\_masked\_sickled.fastq

### Summary

- Basic Statistics
- Per base sequence quality
- Per tile sequence quality
- Per sequence quality scores
- Per base sequence content
- Per sequence GC content
- Per base N content
- Sequence Length Distribution
- Sequence Duplication Levels
- Overrepresented sequences
- Adapter Content
- Kmer Content

### Basic Statistics

| Measure | Value |
| --- | --- |
| Filename | outF2\_AP14.1\_pf\_trimmed\_masked\_sickled.fastq |
| File type | Conventional base calls |
| Encoding | Sanger / Illumina 1.9 |
| Total Sequences | 814270 |
| Sequences flagged as poor quality | 0 |
| Sequence length | 100-131 |
| %GC | 44 |

### Per base sequence quality

### Per tile sequence quality

### Per sequence quality scores

### Per base sequence content

### Per sequence GC content

### Per base N content

### Sequence Length Distribution

### Sequence Duplication Levels

### Overrepresented sequences

No overrepresented sequences

### Adapter Content

### Kmer Content

| Sequence | Count | PValue | Obs/Exp Max | Max Obs/Exp Position |
| --- | --- | --- | --- | --- |
| CTAGGTT | 560 | 5.456968E-12 | 17.179525 | 5 |
| ATCTAGG | 650 | 3.6379788E-12 | 15.721935 | 3 |
| TCTAGGT | 635 | 4.3655746E-11 | 15.14381 | 4 |
| TAGGTTC | 660 | 7.8216544E-11 | 14.580221 | 6 |
| CGATCTA | 805 | 1.4370016E-10 | 12.657487 | 1 |
| ACGTCTG | 355 | 1.1078373E-7 | 12.602369 | 124-125 |
| ACACGTC | 360 | 2.0186435E-6 | 11.357007 | 122-123 |
| CGTCTGA | 385 | 4.0052673E-6 | 10.726492 | 124-125 |
| CACGTCT | 355 | 2.1423846E-5 | 10.557218 | 122-123 |
| GATCTAG | 940 | 2.1091182E-8 | 10.227567 | 2 |
| GCACACG | 440 | 2.253968E-6 | 9.981436 | 120-121 |
| CACACGT | 410 | 1.0574968E-5 | 9.887801 | 120-121 |
| TCGGAAG | 955 | 5.7953002E-9 | 7.567512 | 124-125 |
| TAGATCG | 1470 | 0.0 | 7.4915185 | 124-125 |
| CGGAAGA | 875 | 7.637027E-8 | 7.472789 | 124-125 |
| AACCTAG | 1520 | 0.0 | 7.471496 | 124-125 |
| CCTAGAT | 1525 | 0.0 | 6.9259124 | 122-123 |
| AGCACAC | 635 | 2.468866E-4 | 6.8629065 | 118-119 |
| GAACCTA | 1680 | 0.0 | 6.7599244 | 124-125 |
| ACCTAGA | 1415 | 2.910383E-11 | 6.741968 | 122-123 |

Produced by FastQC (version 0.11.5)
