## Supplementary material 4 for "Natural and human-mediated drivers of microevolution in Neotropical palms: a historical genomics approach": outF2_AP14.2_pf_trimmed_masked_sickled_fastqc.html

outF2\_AP14.2\_pf\_trimmed\_masked\_sickled.fastq FastQC Report 

FastQC Report

mar. 25 juin 2019  
outF2\_AP14.2\_pf\_trimmed\_masked\_sickled.fastq

### Summary

- Basic Statistics
- Per base sequence quality
- Per tile sequence quality
- Per sequence quality scores
- Per base sequence content
- Per sequence GC content
- Per base N content
- Sequence Length Distribution
- Sequence Duplication Levels
- Overrepresented sequences
- Adapter Content
- Kmer Content

### Basic Statistics

| Measure | Value |
| --- | --- |
| Filename | outF2\_AP14.2\_pf\_trimmed\_masked\_sickled.fastq |
| File type | Conventional base calls |
| Encoding | Sanger / Illumina 1.9 |
| Total Sequences | 814270 |
| Sequences flagged as poor quality | 0 |
| Sequence length | 100-131 |
| %GC | 44 |

### Per base sequence quality

### Per tile sequence quality

### Per sequence quality scores

### Per base sequence content

### Per sequence GC content

### Per base N content

### Sequence Length Distribution

### Sequence Duplication Levels

### Overrepresented sequences

No overrepresented sequences

### Adapter Content

### Kmer Content

| Sequence | Count | PValue | Obs/Exp Max | Max Obs/Exp Position |
| --- | --- | --- | --- | --- |
| TCTAGGT | 455 | 4.1600288E-9 | 16.171469 | 4 |
| CGTGTAG | 225 | 1.3262278E-4 | 14.467849 | 124-125 |
| ATCTAGG | 570 | 5.4442353E-9 | 13.892788 | 3 |
| CGATCTA | 640 | 2.9598596E-8 | 12.31029 | 1 |
| CTAGGTT | 510 | 4.1485146E-6 | 12.236417 | 5 |
| TAGGTTC | 540 | 7.824376E-6 | 11.55962 | 6 |
| CGTCGTG | 355 | 4.1120606E-5 | 11.462205 | 124-125 |
| CGGAAGA | 805 | 7.2759576E-12 | 10.614999 | 124-125 |
| GTCGTGT | 265 | 0.008702637 | 10.512004 | 122-123 |
| AAGAGCG | 600 | 9.751784E-8 | 10.172707 | 124-125 |
| TCGGAAG | 825 | 1.2914825E-10 | 9.864443 | 124-125 |
| GTGTAGG | 430 | 2.8631164E-4 | 9.462983 | 124-125 |
| TAGATCG | 1415 | 0.0 | 9.202167 | 124-125 |
| AGAGCGT | 425 | 0.002705146 | 8.61688 | 124-125 |
| GATCTAG | 920 | 4.0515733E-6 | 8.602474 | 2 |
| CCTAGAT | 1460 | 0.0 | 8.177147 | 122-123 |
| GAGCGTC | 470 | 0.0012683515 | 8.150025 | 118-119 |
| AACCTAG | 1470 | 0.0 | 7.676293 | 120-121 |
| ACCTAGA | 1575 | 0.0 | 7.4922795 | 124-125 |
| GGAACCT | 675 | 4.0763477E-4 | 7.233925 | 124-125 |

Produced by FastQC (version 0.11.5)
