## Supplementary material 4 for "Natural and human-mediated drivers of microevolution in Neotropical palms: a historical genomics approach": outF2_AP17.1_pf_trimmed_masked_sickled_fastqc.html

outF2\_AP17.1\_pf\_trimmed\_masked\_sickled.fastq FastQC Report 

FastQC Report

mar. 25 juin 2019  
outF2\_AP17.1\_pf\_trimmed\_masked\_sickled.fastq

### Summary

- Basic Statistics
- Per base sequence quality
- Per tile sequence quality
- Per sequence quality scores
- Per base sequence content
- Per sequence GC content
- Per base N content
- Sequence Length Distribution
- Sequence Duplication Levels
- Overrepresented sequences
- Adapter Content
- Kmer Content

### Basic Statistics

| Measure | Value |
| --- | --- |
| Filename | outF2\_AP17.1\_pf\_trimmed\_masked\_sickled.fastq |
| File type | Conventional base calls |
| Encoding | Sanger / Illumina 1.9 |
| Total Sequences | 1055624 |
| Sequences flagged as poor quality | 0 |
| Sequence length | 100-131 |
| %GC | 44 |

### Per base sequence quality

### Per tile sequence quality

### Per sequence quality scores

### Per base sequence content

### Per sequence GC content

### Per base N content

### Sequence Length Distribution

### Sequence Duplication Levels

### Overrepresented sequences

No overrepresented sequences

### Adapter Content

### Kmer Content

| Sequence | Count | PValue | Obs/Exp Max | Max Obs/Exp Position |
| --- | --- | --- | --- | --- |
| CTAGTGG | 730 | 0.0 | 21.6455 | 5 |
| TCTAGTG | 770 | 0.0 | 20.504002 | 4 |
| TAGTGGC | 790 | 0.0 | 20.006433 | 6 |
| ATCTAGT | 905 | 0.0 | 17.427502 | 3 |
| CGATCTA | 1075 | 0.0 | 15.677144 | 1 |
| GATCTAG | 1410 | 0.0 | 12.004344 | 2 |
| ACACGTC | 435 | 4.4874832E-7 | 11.315913 | 124-125 |
| AGTGGCG | 490 | 3.5426786E-4 | 10.759129 | 7 |
| GCACACG | 500 | 3.2154094E-7 | 10.363934 | 122-123 |
| CCACTAG | 1730 | 0.0 | 10.068094 | 124-125 |
| AGTGGCT | 610 | 2.4704312E-4 | 9.602865 | 7 |
| CACACGT | 445 | 8.040699E-5 | 9.359835 | 124-125 |
| CACGTCT | 455 | 1.0247102E-4 | 9.154124 | 124-125 |
| GCCACTA | 1885 | 0.0 | 8.838465 | 124-125 |
| CACTAGA | 1795 | 0.0 | 8.437836 | 124-125 |
| TCGGAAG | 1015 | 3.43789E-10 | 8.207146 | 124-125 |
| ACTAGAT | 1695 | 0.0 | 7.3719053 | 124-125 |
| AGCCACT | 1110 | 1.6707418E-8 | 7.163608 | 124-125 |
| ACCATAC | 435 | 0.0047991113 | 7.116435 | 60-61 |
| AGCACAC | 845 | 3.646859E-5 | 6.721559 | 124-125 |

Produced by FastQC (version 0.11.5)
