## Supplementary material 4 for "Natural and human-mediated drivers of microevolution in Neotropical palms: a historical genomics approach": outF2_AP17.2_pf_trimmed_masked_sickled_fastqc.html

outF2\_AP17.2\_pf\_trimmed\_masked\_sickled.fastq FastQC Report 

FastQC Report

mar. 25 juin 2019  
outF2\_AP17.2\_pf\_trimmed\_masked\_sickled.fastq

### Summary

- Basic Statistics
- Per base sequence quality
- Per tile sequence quality
- Per sequence quality scores
- Per base sequence content
- Per sequence GC content
- Per base N content
- Sequence Length Distribution
- Sequence Duplication Levels
- Overrepresented sequences
- Adapter Content
- Kmer Content

### Basic Statistics

| Measure | Value |
| --- | --- |
| Filename | outF2\_AP17.2\_pf\_trimmed\_masked\_sickled.fastq |
| File type | Conventional base calls |
| Encoding | Sanger / Illumina 1.9 |
| Total Sequences | 1055624 |
| Sequences flagged as poor quality | 0 |
| Sequence length | 100-131 |
| %GC | 45 |

### Per base sequence quality

### Per tile sequence quality

### Per sequence quality scores

### Per base sequence content

### Per sequence GC content

### Per base N content

### Sequence Length Distribution

### Sequence Duplication Levels

### Overrepresented sequences

No overrepresented sequences

### Adapter Content

### Kmer Content

| Sequence | Count | PValue | Obs/Exp Max | Max Obs/Exp Position |
| --- | --- | --- | --- | --- |
| CTAGTGG | 655 | 0.0 | 26.466991 | 5 |
| TAGTGGC | 660 | 0.0 | 25.487585 | 6 |
| TCTAGTG | 685 | 0.0 | 24.425919 | 4 |
| ATCTAGT | 795 | 0.0 | 23.07743 | 3 |
| CGATCTA | 885 | 0.0 | 19.310852 | 1 |
| AGTGGCG | 305 | 6.2457366E-6 | 16.543518 | 7 |
| GATCTAG | 1250 | 0.0 | 13.766923 | 2 |
| GGGTATT | 295 | 0.0015559832 | 13.234157 | 4 |
| CCACTAG | 1755 | 0.0 | 11.351804 | 124-125 |
| GCCACTA | 1820 | 0.0 | 11.184347 | 124-125 |
| AGTGGCC | 405 | 0.0012633309 | 11.074398 | 7 |
| CACTAGA | 1675 | 0.0 | 10.859721 | 124-125 |
| CTAGGCC | 255 | 0.0062395423 | 9.134279 | 30-31 |
| GGTTTTA | 550 | 0.0016520504 | 9.0974655 | 3 |
| AGTGGCT | 560 | 0.0018033417 | 9.010309 | 7 |
| CGCCACT | 475 | 5.482897E-4 | 8.868442 | 122-123 |
| GTTTTAG | 595 | 0.003276571 | 8.436178 | 4 |
| TCGGAAG | 1090 | 7.592462E-9 | 7.946716 | 124-125 |
| ACTAGAT | 1790 | 0.0 | 7.766079 | 122-123 |
| AGCCACT | 1030 | 1.984572E-7 | 7.568668 | 124-125 |

Produced by FastQC (version 0.11.5)
