## Supplementary material 4 for "Natural and human-mediated drivers of microevolution in Neotropical palms: a historical genomics approach": outF4_AP01.1_pf_trimmed_masked_sickled_fastqc.html

outF4\_AP01.1\_pf\_trimmed\_masked\_sickled.fastq FastQC Report 

FastQC Report

mar. 25 juin 2019  
outF4\_AP01.1\_pf\_trimmed\_masked\_sickled.fastq

### Summary

- Basic Statistics
- Per base sequence quality
- Per tile sequence quality
- Per sequence quality scores
- Per base sequence content
- Per sequence GC content
- Per base N content
- Sequence Length Distribution
- Sequence Duplication Levels
- Overrepresented sequences
- Adapter Content
- Kmer Content

### Basic Statistics

| Measure | Value |
| --- | --- |
| Filename | outF4\_AP01.1\_pf\_trimmed\_masked\_sickled.fastq |
| File type | Conventional base calls |
| Encoding | Sanger / Illumina 1.9 |
| Total Sequences | 1278774 |
| Sequences flagged as poor quality | 0 |
| Sequence length | 100-131 |
| %GC | 45 |

### Per base sequence quality

### Per tile sequence quality

### Per sequence quality scores

### Per base sequence content

### Per sequence GC content

### Per base N content

### Sequence Length Distribution

### Sequence Duplication Levels

### Overrepresented sequences

No overrepresented sequences

### Adapter Content

### Kmer Content

| Sequence | Count | PValue | Obs/Exp Max | Max Obs/Exp Position |
| --- | --- | --- | --- | --- |
| CTATGTA | 855 | 0.0 | 21.253096 | 5 |
| TATGTAG | 920 | 0.0 | 19.742884 | 6 |
| CTAACCG | 165 | 0.007362064 | 18.354946 | 5 |
| TCTATGT | 1130 | 0.0 | 16.61295 | 4 |
| CGATCTA | 1170 | 0.0 | 15.992859 | 1 |
| ATCTATG | 1460 | 0.0 | 12.442204 | 3 |
| ACGTCTG | 340 | 1.6951199E-4 | 9.970032 | 124-125 |
| GATCTAT | 1910 | 0.0 | 9.821577 | 2 |
| ACACGTC | 445 | 2.6808062E-5 | 9.141063 | 124-125 |
| GCACACG | 495 | 1.0335136E-5 | 8.85564 | 122-123 |
| ATGTAGT | 825 | 4.141005E-5 | 8.808623 | 7 |
| CACACGT | 490 | 8.2983606E-5 | 8.3015785 | 124-125 |
| CTACATA | 2060 | 0.0 | 8.184358 | 122-123 |
| TACATAG | 1985 | 0.0 | 7.8554807 | 124-125 |
| ATGTAGG | 715 | 0.0081537785 | 7.6228466 | 7 |
| CACGTCT | 405 | 0.009508732 | 7.493234 | 122-123 |
| TAGATCG | 1910 | 0.0 | 7.414771 | 122-123 |
| ACATAGA | 2205 | 0.0 | 7.37918 | 124-125 |
| TCGGAAG | 1145 | 6.7666406E-10 | 7.10528 | 124-125 |
| AGCACAC | 955 | 6.1168976E-8 | 7.0990806 | 124-125 |

Produced by FastQC (version 0.11.5)
