## Supplementary material 4 for "Natural and human-mediated drivers of microevolution in Neotropical palms: a historical genomics approach": outF4_AP01.2_pf_trimmed_masked_sickled_fastqc.html

outF4\_AP01.2\_pf\_trimmed\_masked\_sickled.fastq FastQC Report 

FastQC Report

mar. 25 juin 2019  
outF4\_AP01.2\_pf\_trimmed\_masked\_sickled.fastq

### Summary

- Basic Statistics
- Per base sequence quality
- Per tile sequence quality
- Per sequence quality scores
- Per base sequence content
- Per sequence GC content
- Per base N content
- Sequence Length Distribution
- Sequence Duplication Levels
- Overrepresented sequences
- Adapter Content
- Kmer Content

### Basic Statistics

| Measure | Value |
| --- | --- |
| Filename | outF4\_AP01.2\_pf\_trimmed\_masked\_sickled.fastq |
| File type | Conventional base calls |
| Encoding | Sanger / Illumina 1.9 |
| Total Sequences | 1278774 |
| Sequences flagged as poor quality | 0 |
| Sequence length | 100-131 |
| %GC | 45 |

### Per base sequence quality

### Per tile sequence quality

### Per sequence quality scores

### Per base sequence content

### Per sequence GC content

### Per base N content

### Sequence Length Distribution

### Sequence Duplication Levels

### Overrepresented sequences

No overrepresented sequences

### Adapter Content

### Kmer Content

| Sequence | Count | PValue | Obs/Exp Max | Max Obs/Exp Position |
| --- | --- | --- | --- | --- |
| CTATGTA | 730 | 0.0 | 20.311708 | 5 |
| TATGTAG | 805 | 0.0 | 18.439926 | 6 |
| TCTATGT | 935 | 0.0 | 17.666933 | 4 |
| CGATCTA | 1100 | 0.0 | 14.88333 | 1 |
| ATCTATG | 1405 | 0.0 | 12.543963 | 3 |
| AGAGCGT | 460 | 3.9171573E-7 | 11.435569 | 124-125 |
| GAGCGTC | 505 | 1.0793519E-7 | 11.217832 | 124-125 |
| AAGAGCG | 720 | 1.6370905E-11 | 11.010429 | 122-123 |
| GCGTCGT | 425 | 2.93946E-5 | 10.259129 | 122-123 |
| ATGTAGC | 645 | 5.2478397E-5 | 9.731521 | 7 |
| CTACATA | 2080 | 0.0 | 9.147126 | 122-123 |
| TACATAG | 2050 | 0.0 | 9.0798 | 124-125 |
| ACATAGA | 2260 | 0.0 | 8.773237 | 124-125 |
| TAGATCG | 1760 | 0.0 | 8.736614 | 124-125 |
| GATCTAT | 1695 | 0.0 | 8.705283 | 2 |
| CATAGAT | 2305 | 0.0 | 8.250859 | 124-125 |
| AGCGTCG | 460 | 0.0058112075 | 7.916932 | 124-125 |
| TCGGAAG | 1135 | 8.6583896E-10 | 7.8433046 | 124-125 |
| ATGTAGG | 655 | 0.0063362685 | 7.840594 | 7 |
| CGGAAGA | 1020 | 5.248785E-7 | 7.140763 | 124-125 |

Produced by FastQC (version 0.11.5)
