## Supplementary material 4 for "Natural and human-mediated drivers of microevolution in Neotropical palms: a historical genomics approach": outF4_AP03.1_pf_trimmed_masked_sickled_fastqc.html

outF4\_AP03.1\_pf\_trimmed\_masked\_sickled.fastq FastQC Report 

FastQC Report

mar. 25 juin 2019  
outF4\_AP03.1\_pf\_trimmed\_masked\_sickled.fastq

### Summary

- Basic Statistics
- Per base sequence quality
- Per tile sequence quality
- Per sequence quality scores
- Per base sequence content
- Per sequence GC content
- Per base N content
- Sequence Length Distribution
- Sequence Duplication Levels
- Overrepresented sequences
- Adapter Content
- Kmer Content

### Basic Statistics

| Measure | Value |
| --- | --- |
| Filename | outF4\_AP03.1\_pf\_trimmed\_masked\_sickled.fastq |
| File type | Conventional base calls |
| Encoding | Sanger / Illumina 1.9 |
| Total Sequences | 1040263 |
| Sequences flagged as poor quality | 0 |
| Sequence length | 100-131 |
| %GC | 43 |

### Per base sequence quality

### Per tile sequence quality

### Per sequence quality scores

### Per base sequence content

### Per sequence GC content

### Per base N content

### Sequence Length Distribution

### Sequence Duplication Levels

### Overrepresented sequences

No overrepresented sequences

### Adapter Content

### Kmer Content

| Sequence | Count | PValue | Obs/Exp Max | Max Obs/Exp Position |
| --- | --- | --- | --- | --- |
| CGATCTC | 1280 | 0.0 | 32.36773 | 1 |
| TCTCACC | 1315 | 0.0 | 32.041435 | 4 |
| TCACCAC | 1430 | 0.0 | 28.66438 | 6 |
| CTCACCA | 1455 | 0.0 | 27.769058 | 5 |
| ATCTCAC | 1925 | 0.0 | 21.589348 | 3 |
| GATCTCA | 2045 | 0.0 | 20.616884 | 2 |
| CACCACG | 595 | 0.0 | 19.956541 | 7 |
| CACCACT | 905 | 0.0 | 15.088688 | 7 |
| ACCACGA | 370 | 2.2230948E-5 | 14.468656 | 8 |
| AGTAGCG | 220 | 0.0014403851 | 10.90282 | 20-21 |
| CACCACA | 1045 | 1.1677912E-9 | 10.226533 | 7 |
| TCGGAAG | 935 | 0.0 | 9.549973 | 124-125 |
| ACCACTT | 900 | 1.5124606E-6 | 9.252794 | 8 |
| ACCACAA | 825 | 5.1083945E-5 | 8.651963 | 8 |
| GCACACG | 375 | 0.0028377166 | 8.572055 | 124-125 |
| CGGAAGA | 860 | 3.3305696E-9 | 8.306255 | 124-125 |
| CACACGT | 390 | 0.0040445453 | 8.242361 | 124-125 |
| GGTGAGA | 1830 | 0.0 | 7.2214494 | 124-125 |
| CGTGGTG | 685 | 4.2702355E-5 | 7.212231 | 122-123 |
| GTGGTGA | 1945 | 0.0 | 6.8943954 | 122-123 |

Produced by FastQC (version 0.11.5)
