## Supplementary material 4 for "Natural and human-mediated drivers of microevolution in Neotropical palms: a historical genomics approach": outF4_AP03.2_pf_trimmed_masked_sickled_fastqc.html

outF4\_AP03.2\_pf\_trimmed\_masked\_sickled.fastq FastQC Report 

FastQC Report

mar. 25 juin 2019  
outF4\_AP03.2\_pf\_trimmed\_masked\_sickled.fastq

### Summary

- Basic Statistics
- Per base sequence quality
- Per tile sequence quality
- Per sequence quality scores
- Per base sequence content
- Per sequence GC content
- Per base N content
- Sequence Length Distribution
- Sequence Duplication Levels
- Overrepresented sequences
- Adapter Content
- Kmer Content

### Basic Statistics

| Measure | Value |
| --- | --- |
| Filename | outF4\_AP03.2\_pf\_trimmed\_masked\_sickled.fastq |
| File type | Conventional base calls |
| Encoding | Sanger / Illumina 1.9 |
| Total Sequences | 1040263 |
| Sequences flagged as poor quality | 0 |
| Sequence length | 100-131 |
| %GC | 44 |

### Per base sequence quality

### Per tile sequence quality

### Per sequence quality scores

### Per base sequence content

### Per sequence GC content

### Per base N content

### Sequence Length Distribution

### Sequence Duplication Levels

### Overrepresented sequences

No overrepresented sequences

### Adapter Content

### Kmer Content

| Sequence | Count | PValue | Obs/Exp Max | Max Obs/Exp Position |
| --- | --- | --- | --- | --- |
| CGATCTC | 1270 | 0.0 | 31.288177 | 1 |
| TCTCACC | 1330 | 0.0 | 29.649534 | 4 |
| TCACCAC | 1375 | 0.0 | 28.733364 | 6 |
| CTCACCA | 1595 | 0.0 | 25.488274 | 5 |
| CACCACG | 420 | 0.0 | 24.159197 | 7 |
| GATCTCA | 2025 | 0.0 | 20.835815 | 2 |
| ATCTCAC | 2155 | 0.0 | 18.819687 | 3 |
| GAGCGTC | 330 | 1.4899031E-5 | 12.656945 | 124-125 |
| ACCACGA | 405 | 8.587433E-5 | 12.530187 | 8 |
| CACCACA | 1060 | 0.0 | 12.2315445 | 7 |
| AGCGTCG | 315 | 0.0018098347 | 10.607726 | 124-125 |
| AGAGCGT | 400 | 1.3786535E-4 | 10.175529 | 122-123 |
| AAGAGCG | 605 | 1.1889897E-6 | 9.418671 | 122-123 |
| CCACGAT | 505 | 0.0074687926 | 8.935601 | 9 |
| CACCACT | 1000 | 1.4186571E-6 | 8.455719 | 7 |
| CGGAAGA | 915 | 4.974936E-8 | 8.21664 | 124-125 |
| TCGGAAG | 915 | 4.974936E-8 | 8.21664 | 124-125 |
| GTGGTGA | 1700 | 0.0 | 7.370809 | 124-125 |
| TGGTGAG | 1775 | 1.8189894E-12 | 7.0593667 | 124-125 |
| GGTGAGA | 1740 | 3.6379788E-12 | 6.9613204 | 124-125 |

Produced by FastQC (version 0.11.5)
