## Supplementary material 4 for "Natural and human-mediated drivers of microevolution in Neotropical palms: a historical genomics approach": outF4_AP07.1_pf_trimmed_masked_sickled_fastqc.html

outF4\_AP07.1\_pf\_trimmed\_masked\_sickled.fastq FastQC Report 

FastQC Report

mar. 25 juin 2019  
outF4\_AP07.1\_pf\_trimmed\_masked\_sickled.fastq

### Summary

- Basic Statistics
- Per base sequence quality
- Per tile sequence quality
- Per sequence quality scores
- Per base sequence content
- Per sequence GC content
- Per base N content
- Sequence Length Distribution
- Sequence Duplication Levels
- Overrepresented sequences
- Adapter Content
- Kmer Content

### Basic Statistics

| Measure | Value |
| --- | --- |
| Filename | outF4\_AP07.1\_pf\_trimmed\_masked\_sickled.fastq |
| File type | Conventional base calls |
| Encoding | Sanger / Illumina 1.9 |
| Total Sequences | 880460 |
| Sequences flagged as poor quality | 0 |
| Sequence length | 100-131 |
| %GC | 42 |

### Per base sequence quality

### Per tile sequence quality

### Per sequence quality scores

### Per base sequence content

### Per sequence GC content

### Per base N content

### Sequence Length Distribution

### Sequence Duplication Levels

### Overrepresented sequences

No overrepresented sequences

### Adapter Content

### Kmer Content

| Sequence | Count | PValue | Obs/Exp Max | Max Obs/Exp Position |
| --- | --- | --- | --- | --- |
| CACGTCT | 325 | 6.002665E-11 | 16.710154 | 124-125 |
| CTCACTG | 855 | 0.0 | 16.636618 | 5 |
| TCACTGC | 875 | 0.0 | 15.576276 | 6 |
| CGATCTC | 880 | 0.0 | 15.422071 | 1 |
| ACGTCTG | 280 | 7.6640936E-7 | 14.223523 | 124-125 |
| TCTCACT | 1010 | 0.0 | 14.076078 | 4 |
| CACACGT | 395 | 1.9075742E-8 | 12.706317 | 122-123 |
| ACACGTC | 390 | 1.9466461E-7 | 12.068443 | 124-125 |
| CGTCTGA | 305 | 2.8767807E-5 | 11.870601 | 124-125 |
| GCACACG | 415 | 4.950198E-7 | 11.230109 | 122-123 |
| ATCTCAC | 1495 | 0.0 | 10.296651 | 3 |
| GATCTCA | 1525 | 1.6370905E-11 | 8.9283495 | 2 |
| AGCACAC | 675 | 4.5669367E-8 | 8.927697 | 120-121 |
| CACTGCA | 735 | 0.001401747 | 8.067433 | 7 |
| CGGTGGC | 405 | 0.0024912485 | 7.6104403 | 74-75 |
| TCGGAAG | 850 | 1.886765E-7 | 7.591757 | 122-123 |
| CGGAAGA | 770 | 2.1371197E-5 | 6.983759 | 122-123 |
| GCAGTGA | 1580 | 1.8189894E-12 | 6.87443 | 124-125 |
| AGAGCAC | 890 | 3.269378E-5 | 6.291227 | 118-119 |
| TGCAGTG | 945 | 4.7741414E-5 | 6.130003 | 124-125 |

Produced by FastQC (version 0.11.5)
