## Supplementary material 4 for "Natural and human-mediated drivers of microevolution in Neotropical palms: a historical genomics approach": outF4_AP07.2_pf_trimmed_masked_sickled_fastqc.html

outF4\_AP07.2\_pf\_trimmed\_masked\_sickled.fastq FastQC Report 

FastQC Report

mar. 25 juin 2019  
outF4\_AP07.2\_pf\_trimmed\_masked\_sickled.fastq

### Summary

- Basic Statistics
- Per base sequence quality
- Per tile sequence quality
- Per sequence quality scores
- Per base sequence content
- Per sequence GC content
- Per base N content
- Sequence Length Distribution
- Sequence Duplication Levels
- Overrepresented sequences
- Adapter Content
- Kmer Content

### Basic Statistics

| Measure | Value |
| --- | --- |
| Filename | outF4\_AP07.2\_pf\_trimmed\_masked\_sickled.fastq |
| File type | Conventional base calls |
| Encoding | Sanger / Illumina 1.9 |
| Total Sequences | 880460 |
| Sequences flagged as poor quality | 0 |
| Sequence length | 100-131 |
| %GC | 42 |

### Per base sequence quality

### Per tile sequence quality

### Per sequence quality scores

### Per base sequence content

### Per sequence GC content

### Per base N content

### Sequence Length Distribution

### Sequence Duplication Levels

### Overrepresented sequences

No overrepresented sequences

### Adapter Content

### Kmer Content

| Sequence | Count | PValue | Obs/Exp Max | Max Obs/Exp Position |
| --- | --- | --- | --- | --- |
| CGTGTAG | 210 | 2.7595033E-6 | 18.02479 | 124-125 |
| TCGTGTA | 220 | 4.3004966E-6 | 17.205482 | 124-125 |
| AGCGTCG | 335 | 1.16968295E-7 | 14.362277 | 120-121 |
| GAGCGTC | 340 | 1.4018588E-7 | 14.151066 | 120-121 |
| TCACTGC | 655 | 2.6338967E-9 | 12.988093 | 6 |
| CGTCGTG | 295 | 8.2497856E-5 | 12.584398 | 122-123 |
| GTCGTGT | 230 | 0.0023141655 | 12.554002 | 122-123 |
| CTCACTG | 710 | 8.89122E-9 | 11.968149 | 5 |
| GCGTCGT | 245 | 0.003715014 | 11.785388 | 122-123 |
| AGAGCGT | 450 | 3.315181E-7 | 11.58291 | 120-121 |
| TATACCC | 340 | 0.0043855845 | 11.52589 | 1 |
| CGATCTC | 860 | 1.7704224E-7 | 9.76446 | 1 |
| CGGAAGA | 740 | 1.2601959E-8 | 9.661938 | 124-125 |
| AAGAGCG | 580 | 4.670843E-6 | 9.426758 | 124-125 |
| TCTCACT | 910 | 3.531477E-7 | 9.311211 | 4 |
| TCGGAAG | 900 | 5.0331437E-9 | 8.708094 | 122-123 |
| CAGTGAG | 1425 | 0.0 | 8.263998 | 124-125 |
| GCAGTGA | 1650 | 0.0 | 8.1566725 | 124-125 |
| GAGATCG | 1255 | 3.0195224E-9 | 7.37269 | 124-125 |
| ATCTCAC | 1320 | 1.2319779E-6 | 7.265732 | 3 |

Produced by FastQC (version 0.11.5)
