## Supplementary material 4 for "Natural and human-mediated drivers of microevolution in Neotropical palms: a historical genomics approach": outF4_AP09.1_pf_trimmed_masked_sickled_fastqc.html

outF4\_AP09.1\_pf\_trimmed\_masked\_sickled.fastq FastQC Report 

FastQC Report

mar. 25 juin 2019  
outF4\_AP09.1\_pf\_trimmed\_masked\_sickled.fastq

### Summary

- Basic Statistics
- Per base sequence quality
- Per tile sequence quality
- Per sequence quality scores
- Per base sequence content
- Per sequence GC content
- Per base N content
- Sequence Length Distribution
- Sequence Duplication Levels
- Overrepresented sequences
- Adapter Content
- Kmer Content

### Basic Statistics

| Measure | Value |
| --- | --- |
| Filename | outF4\_AP09.1\_pf\_trimmed\_masked\_sickled.fastq |
| File type | Conventional base calls |
| Encoding | Sanger / Illumina 1.9 |
| Total Sequences | 573648 |
| Sequences flagged as poor quality | 0 |
| Sequence length | 100-131 |
| %GC | 42 |

### Per base sequence quality

### Per tile sequence quality

### Per sequence quality scores

### Per base sequence content

### Per sequence GC content

### Per base N content

### Sequence Length Distribution

### Sequence Duplication Levels

### Overrepresented sequences

No overrepresented sequences

### Adapter Content

### Kmer Content

| Sequence | Count | PValue | Obs/Exp Max | Max Obs/Exp Position |
| --- | --- | --- | --- | --- |
| CACACGT | 210 | 0.0024609582 | 12.451001 | 124-125 |
| ATCGTCG | 325 | 5.092704E-4 | 10.343908 | 124-125 |
| CGTCGGA | 645 | 9.255018E-9 | 9.844978 | 124-125 |
| AGAGCAC | 385 | 2.2268954E-4 | 9.70208 | 124-125 |
| TCGGAAG | 375 | 2.4534966E-4 | 9.6091385 | 120-121 |
| CGGAAGA | 380 | 0.0029501952 | 8.534432 | 120-121 |
| TCGTCGG | 400 | 0.0033892442 | 8.404426 | 124-125 |
| TGCGCAG | 415 | 3.3500706E-4 | 8.205611 | 76-77 |
| TATCGGA | 650 | 0.004869354 | 8.072312 | 4 |
| CGGAGAT | 590 | 4.063636E-5 | 7.939754 | 120-121 |
| GTCGGAG | 755 | 1.0710464E-6 | 7.9158683 | 124-125 |
| GAGATCG | 715 | 3.497984E-4 | 6.669226 | 122-123 |
| CGATTCG | 975 | 0.0035379059 | 6.5720854 | 2 |
| TTAATAT | 1045 | 0.007648671 | 6.103145 | 1 |
| CCACCTG | 735 | 6.977376E-4 | 5.8158813 | 46-47 |
| TCATTCG | 680 | 0.0021592758 | 5.735031 | 36-37 |
| ACAAACC | 805 | 0.0029633052 | 5.186952 | 26-27 |

Produced by FastQC (version 0.11.5)
