## Supplementary material 4 for "Natural and human-mediated drivers of microevolution in Neotropical palms: a historical genomics approach": outF4_AP09.2_pf_trimmed_masked_sickled_fastqc.html

outF4\_AP09.2\_pf\_trimmed\_masked\_sickled.fastq FastQC Report 

FastQC Report

mar. 25 juin 2019  
outF4\_AP09.2\_pf\_trimmed\_masked\_sickled.fastq

### Summary

- Basic Statistics
- Per base sequence quality
- Per tile sequence quality
- Per sequence quality scores
- Per base sequence content
- Per sequence GC content
- Per base N content
- Sequence Length Distribution
- Sequence Duplication Levels
- Overrepresented sequences
- Adapter Content
- Kmer Content

### Basic Statistics

| Measure | Value |
| --- | --- |
| Filename | outF4\_AP09.2\_pf\_trimmed\_masked\_sickled.fastq |
| File type | Conventional base calls |
| Encoding | Sanger / Illumina 1.9 |
| Total Sequences | 573648 |
| Sequences flagged as poor quality | 0 |
| Sequence length | 100-131 |
| %GC | 42 |

### Per base sequence quality

### Per tile sequence quality

### Per sequence quality scores

### Per base sequence content

### Per sequence GC content

### Per base N content

### Sequence Length Distribution

### Sequence Duplication Levels

### Overrepresented sequences

No overrepresented sequences

### Adapter Content

### Kmer Content

| Sequence | Count | PValue | Obs/Exp Max | Max Obs/Exp Position |
| --- | --- | --- | --- | --- |
| GAGCGTC | 175 | 2.347822E-4 | 16.979868 | 124-125 |
| AAGAGCG | 240 | 8.996491E-6 | 15.918624 | 124-125 |
| AGCGTCG | 275 | 1.9074978E-6 | 15.436242 | 124-125 |
| AGAGCGT | 195 | 6.3567143E-4 | 14.897027 | 122-123 |
| CGTCGGA | 615 | 5.0931703E-10 | 11.734054 | 124-125 |
| ATCCGGC | 445 | 0.0027861802 | 10.068963 | 4 |
| TCGTCGG | 340 | 0.0035954602 | 9.764439 | 122-123 |
| TCGGAAG | 420 | 0.0016526693 | 9.096357 | 124-125 |
| GTCGGAG | 625 | 1.0690326E-5 | 8.829531 | 124-125 |
| CGGAAGA | 435 | 0.0022738252 | 8.782689 | 124-125 |
| CGGAGAT | 545 | 2.098738E-4 | 8.5678215 | 124-125 |
| TATGGGT | 330 | 0.0041974955 | 8.207833 | 76-77 |
| GAGATCG | 690 | 7.073173E-5 | 7.593592 | 120-121 |
| ATCGGAA | 795 | 2.142188E-4 | 6.941455 | 124-125 |
| TCGGAGA | 665 | 0.0030481431 | 6.6669016 | 120-121 |
| GAAGAGC | 695 | 0.003563828 | 6.5681653 | 122-123 |
| TGGAAAG | 815 | 1.4702021E-4 | 6.075259 | 96-97 |

Produced by FastQC (version 0.11.5)
