## Supplementary material 4 for "Natural and human-mediated drivers of microevolution in Neotropical palms: a historical genomics approach": outF4_AP10.1_pf_trimmed_masked_sickled_fastqc.html

outF4\_AP10.1\_pf\_trimmed\_masked\_sickled.fastq FastQC Report 

FastQC Report

mar. 25 juin 2019  
outF4\_AP10.1\_pf\_trimmed\_masked\_sickled.fastq

### Summary

- Basic Statistics
- Per base sequence quality
- Per tile sequence quality
- Per sequence quality scores
- Per base sequence content
- Per sequence GC content
- Per base N content
- Sequence Length Distribution
- Sequence Duplication Levels
- Overrepresented sequences
- Adapter Content
- Kmer Content

### Basic Statistics

| Measure | Value |
| --- | --- |
| Filename | outF4\_AP10.1\_pf\_trimmed\_masked\_sickled.fastq |
| File type | Conventional base calls |
| Encoding | Sanger / Illumina 1.9 |
| Total Sequences | 1368093 |
| Sequences flagged as poor quality | 0 |
| Sequence length | 100-131 |
| %GC | 43 |

### Per base sequence quality

### Per tile sequence quality

### Per sequence quality scores

### Per base sequence content

### Per sequence GC content

### Per base N content

### Sequence Length Distribution

### Sequence Duplication Levels

### Overrepresented sequences

No overrepresented sequences

### Adapter Content

### Kmer Content

| Sequence | Count | PValue | Obs/Exp Max | Max Obs/Exp Position |
| --- | --- | --- | --- | --- |
| CGATCTC | 1445 | 0.0 | 14.053462 | 1 |
| GATCTCC | 2190 | 0.0 | 9.572511 | 2 |
| ACGTCTG | 435 | 1.6276992E-4 | 8.773211 | 124-125 |
| ATCTCCT | 2360 | 0.0 | 8.642708 | 3 |
| CACACGT | 525 | 1.5158483E-5 | 8.590909 | 124-125 |
| GCACACG | 520 | 1.3774293E-4 | 7.9479246 | 122-123 |
| ACACGTC | 480 | 5.094875E-4 | 7.8927307 | 122-123 |
| CGTCTGA | 550 | 2.4223E-4 | 7.5696125 | 124-125 |
| TCCTCCA | 3325 | 0.0 | 7.4039726 | 6 |
| TCGGAAG | 1320 | 3.6379788E-12 | 7.359346 | 124-125 |
| TCTCCTC | 3235 | 0.0 | 7.0491757 | 4 |
| CGGAAGA | 1260 | 3.252353E-9 | 6.6083922 | 124-125 |
| GAGATCG | 2290 | 0.0 | 6.3166914 | 122-123 |
| AGGAGAT | 3335 | 0.0 | 6.137779 | 124-125 |
| GGAGATC | 2970 | 0.0 | 6.0743804 | 124-125 |
| ATCGGAA | 2720 | 0.0 | 5.8673773 | 124-125 |
| CTCCTCC | 4020 | 0.0 | 5.6745663 | 5 |
| AGCACAC | 1045 | 6.9327296E-5 | 5.602843 | 122-123 |
| GAGGAGA | 3775 | 0.0 | 5.4223824 | 124-125 |
| AAGAGCA | 2145 | 1.09139364E-10 | 5.337547 | 124-125 |

Produced by FastQC (version 0.11.5)
