## Supplementary material 4 for "Natural and human-mediated drivers of microevolution in Neotropical palms: a historical genomics approach": outF4_AP10.2_pf_trimmed_masked_sickled_fastqc.html

outF4\_AP10.2\_pf\_trimmed\_masked\_sickled.fastq FastQC Report 

FastQC Report

mar. 25 juin 2019  
outF4\_AP10.2\_pf\_trimmed\_masked\_sickled.fastq

### Summary

- Basic Statistics
- Per base sequence quality
- Per tile sequence quality
- Per sequence quality scores
- Per base sequence content
- Per sequence GC content
- Per base N content
- Sequence Length Distribution
- Sequence Duplication Levels
- Overrepresented sequences
- Adapter Content
- Kmer Content

### Basic Statistics

| Measure | Value |
| --- | --- |
| Filename | outF4\_AP10.2\_pf\_trimmed\_masked\_sickled.fastq |
| File type | Conventional base calls |
| Encoding | Sanger / Illumina 1.9 |
| Total Sequences | 1368093 |
| Sequences flagged as poor quality | 0 |
| Sequence length | 100-131 |
| %GC | 43 |

### Per base sequence quality

### Per tile sequence quality

### Per sequence quality scores

### Per base sequence content

### Per sequence GC content

### Per base N content

### Sequence Length Distribution

### Sequence Duplication Levels

### Overrepresented sequences

No overrepresented sequences

### Adapter Content

### Kmer Content

| Sequence | Count | PValue | Obs/Exp Max | Max Obs/Exp Position |
| --- | --- | --- | --- | --- |
| AAGAGCG | 880 | 0.0 | 12.871787 | 124-125 |
| CTTCCGA | 690 | 1.0024014E-6 | 10.632011 | 1 |
| CGTCGTG | 495 | 1.0125932E-6 | 10.624331 | 124-125 |
| CGTGTAG | 325 | 4.3831652E-4 | 10.514972 | 118-119 |
| CGATCTC | 1315 | 0.0 | 10.299276 | 1 |
| AGCGTCG | 470 | 8.008352E-6 | 10.122105 | 122-123 |
| GCGTCGT | 345 | 0.0010632694 | 9.548355 | 114-115 |
| AGAGCGT | 605 | 1.2931341E-6 | 9.3612995 | 124-125 |
| TCGGAAG | 1375 | 0.0 | 8.532156 | 124-125 |
| GATCTCC | 1875 | 0.0 | 8.168903 | 2 |
| TTCCGAT | 1070 | 3.4128298E-6 | 7.9525933 | 2 |
| GGAGATC | 2475 | 0.0 | 7.8456607 | 124-125 |
| GAGATCG | 2105 | 0.0 | 7.3451376 | 122-123 |
| CGGAAGA | 1260 | 1.4133548E-9 | 7.2367687 | 122-123 |
| GAGCGTC | 565 | 0.0069157067 | 6.853381 | 120-121 |
| TGTAGGG | 575 | 0.00819365 | 6.734192 | 120-121 |
| ATCTCCT | 2170 | 9.265932E-9 | 6.2924953 | 3 |
| ATCGGAA | 2440 | 0.0 | 6.1344466 | 124-125 |
| GAGGAGA | 3580 | 0.0 | 5.979958 | 122-123 |
| AGGAGAT | 2990 | 0.0 | 5.9531245 | 124-125 |

Produced by FastQC (version 0.11.5)
