## Supplementary material 4 for "Natural and human-mediated drivers of microevolution in Neotropical palms: a historical genomics approach": outF4_AP12.1_pf_trimmed_masked_sickled_fastqc.html

outF4\_AP12.1\_pf\_trimmed\_masked\_sickled.fastq FastQC Report 

FastQC Report

mar. 25 juin 2019  
outF4\_AP12.1\_pf\_trimmed\_masked\_sickled.fastq

### Summary

- Basic Statistics
- Per base sequence quality
- Per tile sequence quality
- Per sequence quality scores
- Per base sequence content
- Per sequence GC content
- Per base N content
- Sequence Length Distribution
- Sequence Duplication Levels
- Overrepresented sequences
- Adapter Content
- Kmer Content

### Basic Statistics

| Measure | Value |
| --- | --- |
| Filename | outF4\_AP12.1\_pf\_trimmed\_masked\_sickled.fastq |
| File type | Conventional base calls |
| Encoding | Sanger / Illumina 1.9 |
| Total Sequences | 1043461 |
| Sequences flagged as poor quality | 0 |
| Sequence length | 100-131 |
| %GC | 44 |

### Per base sequence quality

### Per tile sequence quality

### Per sequence quality scores

### Per base sequence content

### Per sequence GC content

### Per base N content

### Sequence Length Distribution

### Sequence Duplication Levels

### Overrepresented sequences

No overrepresented sequences

### Adapter Content

### Kmer Content

| Sequence | Count | PValue | Obs/Exp Max | Max Obs/Exp Position |
| --- | --- | --- | --- | --- |
| TCCTGTA | 720 | 0.0 | 21.775208 | 6 |
| CTCCTGT | 835 | 0.0 | 18.061108 | 5 |
| CCTGTAC | 315 | 4.2252123E-6 | 17.237137 | 7 |
| CGATCTC | 1120 | 0.0 | 13.9521 | 1 |
| ACACGTC | 280 | 1.4021025E-6 | 13.481847 | 124-125 |
| TCTCCTG | 1245 | 0.0 | 12.590435 | 4 |
| GCACACG | 310 | 4.827998E-6 | 12.073651 | 122-123 |
| GATCTCC | 1760 | 0.0 | 9.581596 | 2 |
| CACACGT | 290 | 0.004648016 | 9.466877 | 124-125 |
| ATCTCCT | 1825 | 0.0 | 9.238535 | 3 |
| TACAGGA | 1425 | 0.0 | 9.151314 | 124-125 |
| ACAGGAG | 1460 | 0.0 | 8.461832 | 124-125 |
| GTACAGG | 530 | 1.7902395E-4 | 7.769984 | 124-125 |
| AGCACAC | 700 | 1.0357226E-5 | 7.3537345 | 124-125 |
| TCGGAAG | 775 | 5.6686986E-6 | 7.0848885 | 124-125 |
| CAGGAGA | 1640 | 0.0 | 6.905336 | 124-125 |
| CGGAAGA | 730 | 2.5952913E-5 | 6.8877687 | 118-119 |
| TTACAGG | 665 | 0.0023815646 | 6.1926184 | 124-125 |
| GAGATCG | 1370 | 1.1048614E-6 | 5.4639883 | 122-123 |
| AGAGCAC | 950 | 0.0045234966 | 5.01432 | 122-123 |

Produced by FastQC (version 0.11.5)
