## Supplementary material 4 for "Natural and human-mediated drivers of microevolution in Neotropical palms: a historical genomics approach": outF4_AP12.2_pf_trimmed_masked_sickled_fastqc.html

outF4\_AP12.2\_pf\_trimmed\_masked\_sickled.fastq FastQC Report 

FastQC Report

mar. 25 juin 2019  
outF4\_AP12.2\_pf\_trimmed\_masked\_sickled.fastq

### Summary

- Basic Statistics
- Per base sequence quality
- Per tile sequence quality
- Per sequence quality scores
- Per base sequence content
- Per sequence GC content
- Per base N content
- Sequence Length Distribution
- Sequence Duplication Levels
- Overrepresented sequences
- Adapter Content
- Kmer Content

### Basic Statistics

| Measure | Value |
| --- | --- |
| Filename | outF4\_AP12.2\_pf\_trimmed\_masked\_sickled.fastq |
| File type | Conventional base calls |
| Encoding | Sanger / Illumina 1.9 |
| Total Sequences | 1043461 |
| Sequences flagged as poor quality | 0 |
| Sequence length | 100-131 |
| %GC | 44 |

### Per base sequence quality

### Per tile sequence quality

### Per sequence quality scores

### Per base sequence content

### Per sequence GC content

### Per base N content

### Sequence Length Distribution

### Sequence Duplication Levels

### Overrepresented sequences

No overrepresented sequences

### Adapter Content

### Kmer Content

| Sequence | Count | PValue | Obs/Exp Max | Max Obs/Exp Position |
| --- | --- | --- | --- | --- |
| TCCTGTA | 730 | 0.0 | 22.09529 | 6 |
| CTCCTGT | 805 | 0.0 | 19.99421 | 5 |
| TCTCCTG | 1110 | 0.0 | 14.998474 | 4 |
| CGATCTC | 1105 | 0.0 | 14.9479475 | 1 |
| CTTCCGA | 575 | 3.5106495E-10 | 14.858304 | 1 |
| GTCGTGT | 210 | 0.0014692721 | 13.335918 | 124-125 |
| CCTGTAA | 405 | 7.0090224E-5 | 12.805128 | 7 |
| GATCTCC | 1625 | 0.0 | 11.2691345 | 2 |
| CGGGATA | 190 | 0.007849987 | 10.659849 | 16-17 |
| ATCTCCT | 1700 | 0.0 | 10.116015 | 3 |
| TACAGGA | 1415 | 0.0 | 9.948503 | 122-123 |
| ACAGGAG | 1375 | 0.0 | 9.601861 | 124-125 |
| AGCGTCG | 425 | 3.019064E-4 | 9.413589 | 124-125 |
| GTACAGG | 505 | 2.208519E-5 | 9.291836 | 122-123 |
| GAGCGTC | 345 | 0.005490357 | 9.27716 | 124-125 |
| AGAGCGT | 425 | 3.796971E-4 | 9.200739 | 122-123 |
| AAGAGCG | 495 | 1.804656E-4 | 8.689587 | 122-123 |
| CAGGAGA | 1595 | 0.0 | 8.528298 | 124-125 |
| TCGGAAG | 730 | 1.8797746E-5 | 7.672719 | 124-125 |
| TTCCGAT | 1225 | 1.5363522E-4 | 6.5401225 | 2 |

Produced by FastQC (version 0.11.5)
