## Supplementary material 4 for "Natural and human-mediated drivers of microevolution in Neotropical palms: a historical genomics approach": outF4_AP13.1_pf_trimmed_masked_sickled_fastqc.html

outF4\_AP13.1\_pf\_trimmed\_masked\_sickled.fastq FastQC Report 

FastQC Report

mar. 25 juin 2019  
outF4\_AP13.1\_pf\_trimmed\_masked\_sickled.fastq

### Summary

- Basic Statistics
- Per base sequence quality
- Per tile sequence quality
- Per sequence quality scores
- Per base sequence content
- Per sequence GC content
- Per base N content
- Sequence Length Distribution
- Sequence Duplication Levels
- Overrepresented sequences
- Adapter Content
- Kmer Content

### Basic Statistics

| Measure | Value |
| --- | --- |
| Filename | outF4\_AP13.1\_pf\_trimmed\_masked\_sickled.fastq |
| File type | Conventional base calls |
| Encoding | Sanger / Illumina 1.9 |
| Total Sequences | 943593 |
| Sequences flagged as poor quality | 0 |
| Sequence length | 100-131 |
| %GC | 43 |

### Per base sequence quality

### Per tile sequence quality

### Per sequence quality scores

### Per base sequence content

### Per sequence GC content

### Per base N content

### Sequence Length Distribution

### Sequence Duplication Levels

### Overrepresented sequences

No overrepresented sequences

### Adapter Content

### Kmer Content

| Sequence | Count | PValue | Obs/Exp Max | Max Obs/Exp Position |
| --- | --- | --- | --- | --- |
| CTCGACT | 610 | 0.0 | 29.487501 | 5 |
| TCTCGAC | 640 | 0.0 | 28.986612 | 4 |
| CGATCTC | 1015 | 0.0 | 17.626427 | 1 |
| TCGACTC | 1085 | 0.0 | 17.129818 | 6 |
| ATCTCGA | 1175 | 0.0 | 15.779001 | 3 |
| CGACTCT | 375 | 4.8810797E-4 | 12.403428 | 7 |
| GTGTACT | 210 | 0.0079045845 | 10.649742 | 92-93 |
| GATCTCG | 1910 | 0.0 | 10.310856 | 2 |
| GTCGAGA | 960 | 0.0 | 9.767542 | 124-125 |
| CGACTCC | 495 | 0.00493881 | 9.396536 | 7 |
| AGAGTCG | 470 | 0.0012601843 | 8.155634 | 122-123 |
| CGACTCA | 745 | 0.0019442122 | 7.804169 | 7 |
| AGTCGAG | 995 | 7.857852E-8 | 7.4606347 | 124-125 |
| GAGTCGA | 1405 | 5.529728E-10 | 6.820548 | 122-123 |
| CGGAAGA | 690 | 0.0010362084 | 6.666344 | 122-123 |
| GAGATCG | 1160 | 7.705141E-6 | 6.0626116 | 124-125 |
| TCGGAAG | 685 | 0.008613082 | 6.035015 | 120-121 |
| GAAGAGC | 1175 | 6.0512786E-5 | 5.6527047 | 124-125 |

Produced by FastQC (version 0.11.5)
