## Supplementary material 4 for "Natural and human-mediated drivers of microevolution in Neotropical palms: a historical genomics approach": outF4_AP13.2_pf_trimmed_masked_sickled_fastqc.html

outF4\_AP13.2\_pf\_trimmed\_masked\_sickled.fastq FastQC Report 

FastQC Report

mar. 25 juin 2019  
outF4\_AP13.2\_pf\_trimmed\_masked\_sickled.fastq

### Summary

- Basic Statistics
- Per base sequence quality
- Per tile sequence quality
- Per sequence quality scores
- Per base sequence content
- Per sequence GC content
- Per base N content
- Sequence Length Distribution
- Sequence Duplication Levels
- Overrepresented sequences
- Adapter Content
- Kmer Content

### Basic Statistics

| Measure | Value |
| --- | --- |
| Filename | outF4\_AP13.2\_pf\_trimmed\_masked\_sickled.fastq |
| File type | Conventional base calls |
| Encoding | Sanger / Illumina 1.9 |
| Total Sequences | 943593 |
| Sequences flagged as poor quality | 0 |
| Sequence length | 100-131 |
| %GC | 43 |

### Per base sequence quality

### Per tile sequence quality

### Per sequence quality scores

### Per base sequence content

### Per sequence GC content

### Per base N content

### Sequence Length Distribution

### Sequence Duplication Levels

### Overrepresented sequences

No overrepresented sequences

### Adapter Content

### Kmer Content

| Sequence | Count | PValue | Obs/Exp Max | Max Obs/Exp Position |
| --- | --- | --- | --- | --- |
| TCTCGAC | 545 | 0.0 | 22.818108 | 4 |
| CTCGACT | 610 | 0.0 | 18.659632 | 5 |
| CGATCTC | 860 | 0.0 | 14.943365 | 1 |
| TCGACTC | 820 | 0.0 | 14.702968 | 6 |
| ATCTCGA | 1015 | 0.0 | 12.786455 | 3 |
| CGACTCC | 440 | 1.452308E-4 | 11.843854 | 7 |
| GACGGAC | 180 | 0.0051703304 | 11.275395 | 32-33 |
| CGACTCT | 380 | 0.0078082727 | 10.666394 | 7 |
| AAGAGCG | 475 | 4.6624435E-5 | 9.836954 | 124-125 |
| CGGAAGA | 680 | 5.047259E-7 | 9.085456 | 122-123 |
| GTCGAGA | 965 | 1.1150405E-9 | 8.803679 | 124-125 |
| GATCTCG | 1555 | 7.2759576E-12 | 8.709019 | 2 |
| CTTTGGG | 610 | 0.0036333208 | 8.340001 | 4 |
| GAGATCG | 1095 | 1.378794E-9 | 8.146419 | 124-125 |
| AGTCGAG | 955 | 2.3859684E-8 | 7.981018 | 120-121 |
| GAGTCGA | 1265 | 2.9385046E-7 | 6.5118423 | 122-123 |
| TGAGTCG | 680 | 0.007166759 | 6.1430655 | 116-117 |
| TCGAGAT | 1355 | 3.8631297E-6 | 5.9562902 | 124-125 |
| GATGTAT | 865 | 0.0024911505 | 5.2597823 | 96-97 |
| GCCTGTA | 760 | 0.009827538 | 5.042595 | 10-11 |

Produced by FastQC (version 0.11.5)
