## Supplementary material 4 for "Natural and human-mediated drivers of microevolution in Neotropical palms: a historical genomics approach": outF4_AP14.1_pf_trimmed_masked_sickled_fastqc.html

outF4\_AP14.1\_pf\_trimmed\_masked\_sickled.fastq FastQC Report 

FastQC Report

mar. 25 juin 2019  
outF4\_AP14.1\_pf\_trimmed\_masked\_sickled.fastq

### Summary

- Basic Statistics
- Per base sequence quality
- Per tile sequence quality
- Per sequence quality scores
- Per base sequence content
- Per sequence GC content
- Per base N content
- Sequence Length Distribution
- Sequence Duplication Levels
- Overrepresented sequences
- Adapter Content
- Kmer Content

### Basic Statistics

| Measure | Value |
| --- | --- |
| Filename | outF4\_AP14.1\_pf\_trimmed\_masked\_sickled.fastq |
| File type | Conventional base calls |
| Encoding | Sanger / Illumina 1.9 |
| Total Sequences | 1206240 |
| Sequences flagged as poor quality | 0 |
| Sequence length | 100-131 |
| %GC | 44 |

### Per base sequence quality

### Per tile sequence quality

### Per sequence quality scores

### Per base sequence content

### Per sequence GC content

### Per base N content

### Sequence Length Distribution

### Sequence Duplication Levels

### Overrepresented sequences

No overrepresented sequences

### Adapter Content

### Kmer Content

| Sequence | Count | PValue | Obs/Exp Max | Max Obs/Exp Position |
| --- | --- | --- | --- | --- |
| CTCTAAG | 1020 | 0.0 | 17.249266 | 5 |
| TCTAAGT | 1040 | 0.0 | 16.33143 | 6 |
| CGATCTC | 1515 | 0.0 | 11.176143 | 1 |
| ATCTCTA | 1565 | 0.0 | 10.843233 | 3 |
| TCTCTAA | 1640 | 0.0 | 10.721871 | 4 |
| CACACGT | 400 | 8.499442E-5 | 9.312771 | 124-125 |
| CTAAGTA | 525 | 0.0056546074 | 9.243757 | 7 |
| ACACGTC | 400 | 8.707757E-4 | 8.466155 | 124-125 |
| GCACACG | 445 | 2.9159858E-4 | 8.312911 | 122-123 |
| CTAAGTC | 525 | 0.0050025424 | 8.088287 | 7 |
| GATCTCT | 2035 | 1.8189894E-12 | 8.03905 | 2 |
| TTCCGAT | 1555 | 1.1319571E-8 | 7.7776093 | 1 |
| CTTAGAG | 1925 | 0.0 | 7.036804 | 124-125 |
| CGTCTGA | 545 | 0.0023509795 | 6.8350616 | 124-125 |
| CGGAAGA | 1215 | 2.5011104E-9 | 6.689308 | 124-125 |
| TCGGAAG | 1165 | 5.9391823E-8 | 6.3506365 | 122-123 |
| ACTTAGA | 1985 | 0.0 | 6.312297 | 124-125 |
| GAGCACA | 1195 | 8.539246E-8 | 6.234491 | 124-125 |
| GAGATCG | 2010 | 0.0 | 6.1905065 | 122-123 |
| CAATTCG | 685 | 0.001540268 | 5.900446 | 56-57 |

Produced by FastQC (version 0.11.5)
