## Supplementary material 4 for "Natural and human-mediated drivers of microevolution in Neotropical palms: a historical genomics approach": outF4_AP14.2_pf_trimmed_masked_sickled_fastqc.html

outF4\_AP14.2\_pf\_trimmed\_masked\_sickled.fastq FastQC Report 

FastQC Report

mar. 25 juin 2019  
outF4\_AP14.2\_pf\_trimmed\_masked\_sickled.fastq

### Summary

- Basic Statistics
- Per base sequence quality
- Per tile sequence quality
- Per sequence quality scores
- Per base sequence content
- Per sequence GC content
- Per base N content
- Sequence Length Distribution
- Sequence Duplication Levels
- Overrepresented sequences
- Adapter Content
- Kmer Content

### Basic Statistics

| Measure | Value |
| --- | --- |
| Filename | outF4\_AP14.2\_pf\_trimmed\_masked\_sickled.fastq |
| File type | Conventional base calls |
| Encoding | Sanger / Illumina 1.9 |
| Total Sequences | 1206240 |
| Sequences flagged as poor quality | 0 |
| Sequence length | 100-131 |
| %GC | 44 |

### Per base sequence quality

### Per tile sequence quality

### Per sequence quality scores

### Per base sequence content

### Per sequence GC content

### Per base N content

### Sequence Length Distribution

### Sequence Duplication Levels

### Overrepresented sequences

No overrepresented sequences

### Adapter Content

### Kmer Content

| Sequence | Count | PValue | Obs/Exp Max | Max Obs/Exp Position |
| --- | --- | --- | --- | --- |
| TCTAAGT | 820 | 0.0 | 18.17279 | 6 |
| CTCTAAG | 935 | 0.0 | 16.523191 | 5 |
| CGATCTC | 1230 | 0.0 | 13.841806 | 1 |
| GCGGTGT | 230 | 0.003431136 | 11.911875 | 122-123 |
| ATCTCTA | 1545 | 0.0 | 10.727541 | 3 |
| GAGCGTC | 455 | 4.842037E-6 | 10.557972 | 124-125 |
| TCTCTAA | 1470 | 0.0 | 10.498202 | 4 |
| AGAGCGT | 465 | 7.075803E-5 | 9.470009 | 124-125 |
| TTCCGAT | 1345 | 2.7284841E-11 | 9.282758 | 1 |
| GATCTCT | 1755 | 0.0 | 9.097643 | 2 |
| CTAAGTA | 590 | 1.1345204E-5 | 8.740551 | 7 |
| AAGAGCG | 720 | 2.391167E-6 | 8.153962 | 122-123 |
| CGGAAGA | 1145 | 1.2860255E-9 | 7.691798 | 124-125 |
| CTTAGAG | 1680 | 0.0 | 7.3869147 | 124-125 |
| ACTTAGA | 1835 | 0.0 | 7.199271 | 124-125 |
| CACTTAG | 625 | 0.0021652274 | 6.888467 | 122-123 |
| TACTTAG | 890 | 6.55819E-5 | 6.443268 | 120-121 |
| TTAGAGA | 2080 | 5.456968E-12 | 6.1588173 | 124-125 |
| TAGAGAT | 2120 | 0.0 | 6.131235 | 120-121 |
| TCGGAAG | 1095 | 2.0075116E-5 | 6.076377 | 122-123 |

Produced by FastQC (version 0.11.5)
