## Supplementary material 4 for "Natural and human-mediated drivers of microevolution in Neotropical palms: a historical genomics approach": outF4_AP17.1_pf_trimmed_masked_sickled_fastqc.html

outF4\_AP17.1\_pf\_trimmed\_masked\_sickled.fastq FastQC Report 

FastQC Report

mar. 25 juin 2019  
outF4\_AP17.1\_pf\_trimmed\_masked\_sickled.fastq

### Summary

- Basic Statistics
- Per base sequence quality
- Per tile sequence quality
- Per sequence quality scores
- Per base sequence content
- Per sequence GC content
- Per base N content
- Sequence Length Distribution
- Sequence Duplication Levels
- Overrepresented sequences
- Adapter Content
- Kmer Content

### Basic Statistics

| Measure | Value |
| --- | --- |
| Filename | outF4\_AP17.1\_pf\_trimmed\_masked\_sickled.fastq |
| File type | Conventional base calls |
| Encoding | Sanger / Illumina 1.9 |
| Total Sequences | 1296735 |
| Sequences flagged as poor quality | 0 |
| Sequence length | 100-131 |
| %GC | 44 |

### Per base sequence quality

### Per tile sequence quality

### Per sequence quality scores

### Per base sequence content

### Per sequence GC content

### Per base N content

### Sequence Length Distribution

### Sequence Duplication Levels

### Overrepresented sequences

No overrepresented sequences

### Adapter Content

### Kmer Content

| Sequence | Count | PValue | Obs/Exp Max | Max Obs/Exp Position |
| --- | --- | --- | --- | --- |
| CTCTATG | 1130 | 0.0 | 12.288437 | 5 |
| GCACACG | 395 | 5.3445656E-6 | 10.471532 | 124-125 |
| TCTCTAT | 1710 | 0.0 | 9.178898 | 4 |
| CGATCTC | 1460 | 4.0745363E-10 | 8.668624 | 1 |
| TCTATGA | 1575 | 1.9463187E-10 | 8.43611 | 6 |
| GATCTCT | 1870 | 1.0186341E-10 | 7.7481833 | 2 |
| CCGAACT | 450 | 6.88726E-4 | 7.6736536 | 86-87 |
| ATCTCTA | 1735 | 1.1337761E-8 | 7.308899 | 3 |
| TCGGAAG | 1205 | 1.4279067E-9 | 6.8651533 | 124-125 |
| AGCACAC | 880 | 4.868527E-6 | 6.658744 | 124-125 |
| CGGAAGA | 1150 | 2.8247086E-8 | 6.594029 | 124-125 |
| CATAGAG | 2150 | 0.0 | 6.506735 | 122-123 |
| CTATGAA | 1115 | 0.0013703807 | 6.5037127 | 7 |
| GAGATCG | 2065 | 0.0 | 6.342925 | 124-125 |
| TCATAGA | 2880 | 0.0 | 6.279154 | 122-123 |
| CCGTCGC | 620 | 0.0034190943 | 5.995942 | 56-57 |
| ATAGAGA | 2880 | 0.0 | 5.9841647 | 124-125 |
| TAGAGAT | 2585 | 0.0 | 5.7336864 | 124-125 |
| CCTCTTA | 670 | 0.008080578 | 5.5484834 | 56-57 |
| AGAGCAC | 1095 | 1.386166E-4 | 5.3513193 | 124-125 |

Produced by FastQC (version 0.11.5)
