## Supplementary material 4 for "Natural and human-mediated drivers of microevolution in Neotropical palms: a historical genomics approach": outF4_AP17.2_pf_trimmed_masked_sickled_fastqc.html

outF4\_AP17.2\_pf\_trimmed\_masked\_sickled.fastq FastQC Report 

FastQC Report

mar. 25 juin 2019  
outF4\_AP17.2\_pf\_trimmed\_masked\_sickled.fastq

### Summary

- Basic Statistics
- Per base sequence quality
- Per tile sequence quality
- Per sequence quality scores
- Per base sequence content
- Per sequence GC content
- Per base N content
- Sequence Length Distribution
- Sequence Duplication Levels
- Overrepresented sequences
- Adapter Content
- Kmer Content

### Basic Statistics

| Measure | Value |
| --- | --- |
| Filename | outF4\_AP17.2\_pf\_trimmed\_masked\_sickled.fastq |
| File type | Conventional base calls |
| Encoding | Sanger / Illumina 1.9 |
| Total Sequences | 1296735 |
| Sequences flagged as poor quality | 0 |
| Sequence length | 100-131 |
| %GC | 44 |

### Per base sequence quality

### Per tile sequence quality

### Per sequence quality scores

### Per base sequence content

### Per sequence GC content

### Per base N content

### Sequence Length Distribution

### Sequence Duplication Levels

### Overrepresented sequences

No overrepresented sequences

### Adapter Content

### Kmer Content

| Sequence | Count | PValue | Obs/Exp Max | Max Obs/Exp Position |
| --- | --- | --- | --- | --- |
| CTCTATG | 1015 | 0.0 | 11.894157 | 5 |
| AGAGCGT | 435 | 3.5800702E-5 | 10.077046 | 124-125 |
| CGATCTC | 1275 | 9.094947E-11 | 9.356233 | 1 |
| TCGGAAG | 1025 | 1.8189894E-12 | 9.330764 | 124-125 |
| CGGAAGA | 1000 | 9.094947E-12 | 9.165532 | 124-125 |
| ATCTCTA | 1560 | 5.4569682E-11 | 8.442152 | 3 |
| CATAGAG | 2030 | 0.0 | 8.279587 | 122-123 |
| GATCTCT | 1745 | 9.094947E-12 | 8.182492 | 2 |
| AAGAGCG | 660 | 4.6975063E-5 | 7.8492694 | 124-125 |
| TCTCTAT | 1685 | 2.6739144E-10 | 7.8314514 | 4 |
| ATAGAGA | 2535 | 0.0 | 7.3883886 | 124-125 |
| GAGATCG | 1600 | 1.8189894E-12 | 7.222837 | 124-125 |
| TCTATGA | 1630 | 6.746086E-8 | 7.060123 | 6 |
| TAGAGAT | 2395 | 0.0 | 6.6555552 | 124-125 |
| TCATAGA | 2610 | 0.0 | 6.589439 | 122-123 |
| ATCGGAA | 2115 | 1.1616066E-8 | 5.2756677 | 124-125 |
| AGAGATC | 2295 | 4.616595E-9 | 5.109458 | 122-123 |

Produced by FastQC (version 0.11.5)
