## Supplementary material 4 for "Natural and human-mediated drivers of microevolution in Neotropical palms: a historical genomics approach": outF4_AP18.1_pf_trimmed_masked_sickled_fastqc.html

outF4\_AP18.1\_pf\_trimmed\_masked\_sickled.fastq FastQC Report 

FastQC Report

mar. 25 juin 2019  
outF4\_AP18.1\_pf\_trimmed\_masked\_sickled.fastq

### Summary

- Basic Statistics
- Per base sequence quality
- Per tile sequence quality
- Per sequence quality scores
- Per base sequence content
- Per sequence GC content
- Per base N content
- Sequence Length Distribution
- Sequence Duplication Levels
- Overrepresented sequences
- Adapter Content
- Kmer Content

### Basic Statistics

| Measure | Value |
| --- | --- |
| Filename | outF4\_AP18.1\_pf\_trimmed\_masked\_sickled.fastq |
| File type | Conventional base calls |
| Encoding | Sanger / Illumina 1.9 |
| Total Sequences | 687686 |
| Sequences flagged as poor quality | 0 |
| Sequence length | 100-131 |
| %GC | 43 |

### Per base sequence quality

### Per tile sequence quality

### Per sequence quality scores

### Per base sequence content

### Per sequence GC content

### Per base N content

### Sequence Length Distribution

### Sequence Duplication Levels

### Overrepresented sequences

No overrepresented sequences

### Adapter Content

### Kmer Content

| Sequence | Count | PValue | Obs/Exp Max | Max Obs/Exp Position |
| --- | --- | --- | --- | --- |
| CTTCCGA | 295 | 8.567297E-4 | 14.321416 | 1 |
| CTCTATT | 830 | 3.3482138E-8 | 10.9397 | 5 |
| CGATCTC | 720 | 7.277631E-7 | 10.8973465 | 1 |
| TCTATTG | 860 | 6.781745E-5 | 8.44397 | 6 |
| ATCTCTA | 965 | 2.9330158E-5 | 8.148682 | 3 |
| CGGAAGA | 650 | 4.382864E-6 | 7.8137856 | 122-123 |
| CAATAGA | 1565 | 0.0 | 7.632431 | 124-125 |
| AATAGAG | 1420 | 0.0 | 7.210113 | 124-125 |
| AGAGCAC | 590 | 6.540594E-4 | 6.941262 | 124-125 |
| TCAATAG | 845 | 3.4041386E-6 | 6.8120184 | 122-123 |
| GAGATCG | 1100 | 1.6585545E-8 | 6.771948 | 122-123 |
| TCGGAAG | 650 | 2.905447E-4 | 6.7719474 | 122-123 |
| TCTCTAT | 1085 | 9.94119E-4 | 6.6894555 | 4 |
| CTGGAGG | 540 | 0.0053821104 | 6.314784 | 72-73 |
| GATCTCT | 1240 | 0.004483545 | 5.8502483 | 2 |
| CCAAGTT | 700 | 0.002502224 | 5.6646833 | 24-25 |
| AAGAGCA | 1000 | 0.001483248 | 5.11918 | 124-125 |

Produced by FastQC (version 0.11.5)
