## Supplementary material 4 for "Natural and human-mediated drivers of microevolution in Neotropical palms: a historical genomics approach": outF4_AP18.2_pf_trimmed_masked_sickled_fastqc.html

outF4\_AP18.2\_pf\_trimmed\_masked\_sickled.fastq FastQC Report 

FastQC Report

mar. 25 juin 2019  
outF4\_AP18.2\_pf\_trimmed\_masked\_sickled.fastq

### Summary

- Basic Statistics
- Per base sequence quality
- Per tile sequence quality
- Per sequence quality scores
- Per base sequence content
- Per sequence GC content
- Per base N content
- Sequence Length Distribution
- Sequence Duplication Levels
- Overrepresented sequences
- Adapter Content
- Kmer Content

### Basic Statistics

| Measure | Value |
| --- | --- |
| Filename | outF4\_AP18.2\_pf\_trimmed\_masked\_sickled.fastq |
| File type | Conventional base calls |
| Encoding | Sanger / Illumina 1.9 |
| Total Sequences | 687686 |
| Sequences flagged as poor quality | 0 |
| Sequence length | 100-131 |
| %GC | 43 |

### Per base sequence quality

### Per tile sequence quality

### Per sequence quality scores

### Per base sequence content

### Per sequence GC content

### Per base N content

### Sequence Length Distribution

### Sequence Duplication Levels

### Overrepresented sequences

No overrepresented sequences

### Adapter Content

### Kmer Content

| Sequence | Count | PValue | Obs/Exp Max | Max Obs/Exp Position |
| --- | --- | --- | --- | --- |
| TAAGCGA | 95 | 0.008914074 | 17.755558 | 102-103 |
| GCAATAG | 560 | 6.4806954E-8 | 10.460268 | 122-123 |
| AAGAGCG | 430 | 3.4879238E-4 | 9.278277 | 124-125 |
| CGGAAGA | 550 | 6.100254E-6 | 9.230394 | 122-123 |
| CAATAGA | 1530 | 0.0 | 9.126671 | 124-125 |
| CGATCTC | 640 | 5.373662E-4 | 8.884858 | 1 |
| TCGGAAG | 565 | 6.517775E-5 | 8.473612 | 124-125 |
| AATAGAG | 1330 | 7.2759576E-12 | 7.799333 | 124-125 |
| GAGATCG | 1010 | 4.2517822E-6 | 6.7152677 | 124-125 |
| GGAAAGT | 635 | 0.0055481237 | 5.739977 | 74-75 |
| AGAGATC | 1410 | 1.2418332E-6 | 5.700536 | 120-121 |
| TAGAGAT | 1310 | 7.477956E-5 | 5.259164 | 120-121 |
| GGAAGAG | 1245 | 8.325402E-4 | 5.0186877 | 122-123 |

Produced by FastQC (version 0.11.5)
