## Supplementary material 4 for "Natural and human-mediated drivers of microevolution in Neotropical palms: a historical genomics approach": outF4_AP19.1_pf_trimmed_masked_sickled_fastqc.html

outF4\_AP19.1\_pf\_trimmed\_masked\_sickled.fastq FastQC Report 

FastQC Report

mar. 25 juin 2019  
outF4\_AP19.1\_pf\_trimmed\_masked\_sickled.fastq

### Summary

- Basic Statistics
- Per base sequence quality
- Per tile sequence quality
- Per sequence quality scores
- Per base sequence content
- Per sequence GC content
- Per base N content
- Sequence Length Distribution
- Sequence Duplication Levels
- Overrepresented sequences
- Adapter Content
- Kmer Content

### Basic Statistics

| Measure | Value |
| --- | --- |
| Filename | outF4\_AP19.1\_pf\_trimmed\_masked\_sickled.fastq |
| File type | Conventional base calls |
| Encoding | Sanger / Illumina 1.9 |
| Total Sequences | 1108159 |
| Sequences flagged as poor quality | 0 |
| Sequence length | 100-131 |
| %GC | 43 |

### Per base sequence quality

### Per tile sequence quality

### Per sequence quality scores

### Per base sequence content

### Per sequence GC content

### Per base N content

### Sequence Length Distribution

### Sequence Duplication Levels

### Overrepresented sequences

No overrepresented sequences

### Adapter Content

### Kmer Content

| Sequence | Count | PValue | Obs/Exp Max | Max Obs/Exp Position |
| --- | --- | --- | --- | --- |
| GACGTGT | 175 | 5.3039184E-7 | 27.425077 | 5 |
| ACGTGTG | 225 | 4.715912E-6 | 21.335543 | 6 |
| CGTGTGC | 205 | 5.567764E-5 | 20.487087 | 7 |
| CGACGTG | 255 | 1.4004354E-5 | 18.810707 | 4 |
| GCGACGT | 310 | 7.543415E-5 | 15.45834 | 3 |
| GAGCGAC | 285 | 7.155942E-4 | 14.667227 | 1 |
| AGCGACG | 355 | 2.3934155E-4 | 13.49572 | 2 |
| ACACGTC | 405 | 2.921297E-9 | 12.901332 | 124-125 |
| GCACACG | 425 | 8.8833986E-8 | 11.376971 | 122-123 |
| GTGTGCT | 560 | 9.6500444E-4 | 9.649204 | 8 |
| CGAGAGA | 2080 | 0.0 | 8.710267 | 120-121 |
| CACACGT | 445 | 1.993259E-4 | 8.610552 | 124-125 |
| GCGAGAG | 890 | 1.891749E-10 | 8.449902 | 120-121 |
| GAGCACA | 1090 | 1.8189894E-12 | 8.308931 | 124-125 |
| GAGATCG | 1975 | 0.0 | 8.289515 | 124-125 |
| AGAGATC | 2525 | 0.0 | 7.725471 | 124-125 |
| CGGAAGA | 1010 | 2.0154403E-9 | 7.5229626 | 122-123 |
| ACGAGAG | 620 | 1.6147595E-4 | 7.104729 | 118-119 |
| TCGGAAG | 1140 | 1.9254003E-8 | 6.722273 | 124-125 |
| AGAGCAC | 1050 | 2.0975858E-7 | 6.634971 | 124-125 |

Produced by FastQC (version 0.11.5)
