## Supplementary material 4 for "Natural and human-mediated drivers of microevolution in Neotropical palms: a historical genomics approach": outF4_AP19.2_pf_trimmed_masked_sickled_fastqc.html

outF4\_AP19.2\_pf\_trimmed\_masked\_sickled.fastq FastQC Report 

FastQC Report

mar. 25 juin 2019  
outF4\_AP19.2\_pf\_trimmed\_masked\_sickled.fastq

### Summary

- Basic Statistics
- Per base sequence quality
- Per tile sequence quality
- Per sequence quality scores
- Per base sequence content
- Per sequence GC content
- Per base N content
- Sequence Length Distribution
- Sequence Duplication Levels
- Overrepresented sequences
- Adapter Content
- Kmer Content

### Basic Statistics

| Measure | Value |
| --- | --- |
| Filename | outF4\_AP19.2\_pf\_trimmed\_masked\_sickled.fastq |
| File type | Conventional base calls |
| Encoding | Sanger / Illumina 1.9 |
| Total Sequences | 1108159 |
| Sequences flagged as poor quality | 0 |
| Sequence length | 100-131 |
| %GC | 43 |

### Per base sequence quality

### Per tile sequence quality

### Per sequence quality scores

### Per base sequence content

### Per sequence GC content

### Per base N content

### Sequence Length Distribution

### Sequence Duplication Levels

### Overrepresented sequences

No overrepresented sequences

### Adapter Content

### Kmer Content

| Sequence | Count | PValue | Obs/Exp Max | Max Obs/Exp Position |
| --- | --- | --- | --- | --- |
| GAGCGTC | 445 | 5.456968E-12 | 15.361419 | 124-125 |
| CTACGTT | 140 | 0.009999495 | 12.938688 | 72-73 |
| AGCGTCG | 405 | 8.111056E-8 | 12.907163 | 124-125 |
| AGAGCGT | 475 | 5.0351446E-8 | 11.851596 | 124-125 |
| GAGATCG | 1885 | 0.0 | 10.452667 | 124-125 |
| TCGGAAG | 1005 | 0.0 | 10.002681 | 124-125 |
| CGAGAGA | 1975 | 0.0 | 9.982808 | 122-123 |
| AAGAGCG | 695 | 1.2982127E-8 | 9.645259 | 122-123 |
| ACCGAGA | 345 | 0.0052665146 | 9.324238 | 124-125 |
| CCGAGAG | 670 | 1.4748421E-6 | 8.433937 | 118-119 |
| ACGAGAG | 575 | 9.183205E-5 | 8.229306 | 122-123 |
| CGGAAGA | 945 | 1.899025E-8 | 8.084705 | 124-125 |
| AGAGATC | 2485 | 0.0 | 7.928885 | 124-125 |
| AGACCGA | 510 | 0.00213272 | 7.7317824 | 122-123 |
| GAGAGAT | 2535 | 0.0 | 7.4552517 | 124-125 |
| TCGAGAG | 1240 | 5.862603E-9 | 7.1341696 | 124-125 |
| ATCGGAA | 1800 | 1.8371793E-10 | 6.255009 | 124-125 |
| GGAAGAG | 2075 | 1.6734703E-10 | 5.8136063 | 124-125 |
| ATCGAGA | 1220 | 2.2415663E-5 | 5.6739635 | 120-121 |
| TATATCA | 1380 | 0.00448698 | 5.3915224 | 1 |

Produced by FastQC (version 0.11.5)
