## Supplementary material 4 for "Natural and human-mediated drivers of microevolution in Neotropical palms: a historical genomics approach": outF4_AP20.1_pf_trimmed_masked_sickled_fastqc.html

outF4\_AP20.1\_pf\_trimmed\_masked\_sickled.fastq FastQC Report 

FastQC Report

mar. 25 juin 2019  
outF4\_AP20.1\_pf\_trimmed\_masked\_sickled.fastq

### Summary

- Basic Statistics
- Per base sequence quality
- Per tile sequence quality
- Per sequence quality scores
- Per base sequence content
- Per sequence GC content
- Per base N content
- Sequence Length Distribution
- Sequence Duplication Levels
- Overrepresented sequences
- Adapter Content
- Kmer Content

### Basic Statistics

| Measure | Value |
| --- | --- |
| Filename | outF4\_AP20.1\_pf\_trimmed\_masked\_sickled.fastq |
| File type | Conventional base calls |
| Encoding | Sanger / Illumina 1.9 |
| Total Sequences | 737670 |
| Sequences flagged as poor quality | 0 |
| Sequence length | 100-131 |
| %GC | 43 |

### Per base sequence quality

### Per tile sequence quality

### Per sequence quality scores

### Per base sequence content

### Per sequence GC content

### Per base N content

### Sequence Length Distribution

### Sequence Duplication Levels

### Overrepresented sequences

No overrepresented sequences

### Adapter Content

### Kmer Content

| Sequence | Count | PValue | Obs/Exp Max | Max Obs/Exp Position |
| --- | --- | --- | --- | --- |
| GCACACG | 185 | 7.672292E-5 | 15.427946 | 124-125 |
| CTTCCGA | 390 | 5.854355E-4 | 12.137722 | 1 |
| CACACGT | 215 | 0.004140179 | 11.6158085 | 124-125 |
| TCGGAAG | 560 | 2.4901929E-6 | 8.919282 | 124-125 |
| GCAGAGA | 1230 | 1.8189894E-12 | 7.8315635 | 124-125 |
| CAGAGAT | 1235 | 2.0008883E-11 | 7.5109735 | 124-125 |
| CAGCAGA | 1485 | 2.1100277E-10 | 6.4867506 | 124-125 |
| AGAGATC | 1355 | 1.0792064E-8 | 6.2475166 | 122-123 |
| AGCAGAG | 1390 | 1.8571882E-8 | 6.090205 | 122-123 |
| GAGATCG | 990 | 1.3645012E-4 | 5.700596 | 122-123 |
| CCAGCAG | 875 | 0.005482304 | 5.3006015 | 124-125 |

Produced by FastQC (version 0.11.5)
