## Supplementary material 4 for "Natural and human-mediated drivers of microevolution in Neotropical palms: a historical genomics approach": outF4_AP20.2_pf_trimmed_masked_sickled_fastqc.html

outF4\_AP20.2\_pf\_trimmed\_masked\_sickled.fastq FastQC Report 

FastQC Report

mar. 25 juin 2019  
outF4\_AP20.2\_pf\_trimmed\_masked\_sickled.fastq

### Summary

- Basic Statistics
- Per base sequence quality
- Per tile sequence quality
- Per sequence quality scores
- Per base sequence content
- Per sequence GC content
- Per base N content
- Sequence Length Distribution
- Sequence Duplication Levels
- Overrepresented sequences
- Adapter Content
- Kmer Content

### Basic Statistics

| Measure | Value |
| --- | --- |
| Filename | outF4\_AP20.2\_pf\_trimmed\_masked\_sickled.fastq |
| File type | Conventional base calls |
| Encoding | Sanger / Illumina 1.9 |
| Total Sequences | 737670 |
| Sequences flagged as poor quality | 0 |
| Sequence length | 100-131 |
| %GC | 43 |

### Per base sequence quality

### Per tile sequence quality

### Per sequence quality scores

### Per base sequence content

### Per sequence GC content

### Per base N content

### Sequence Length Distribution

### Sequence Duplication Levels

### Overrepresented sequences

No overrepresented sequences

### Adapter Content

### Kmer Content

| Sequence | Count | PValue | Obs/Exp Max | Max Obs/Exp Position |
| --- | --- | --- | --- | --- |
| TCGGAAG | 505 | 9.605374E-7 | 10.666407 | 124-125 |
| AAGAGCG | 320 | 0.0032651245 | 9.879513 | 120-121 |
| GCAGAGA | 1315 | 0.0 | 8.822637 | 124-125 |
| CAGCAGA | 1440 | 0.0 | 8.703423 | 122-123 |
| CGAATCT | 745 | 2.6439485E-4 | 8.386691 | 5 |
| GAGATCG | 965 | 1.6121703E-8 | 8.158164 | 124-125 |
| CAGAGAT | 1345 | 3.45608E-11 | 7.7016516 | 124-125 |
| TATGGAT | 775 | 0.003695128 | 7.3085313 | 4 |
| AGCAGAG | 1465 | 2.4556357E-10 | 7.0708 | 124-125 |
| ATCGGAA | 1060 | 5.008227E-6 | 6.645218 | 124-125 |
| AGAGATC | 1315 | 2.667548E-8 | 6.611385 | 120-121 |

Produced by FastQC (version 0.11.5)
