## Supplementary material 4 for "Natural and human-mediated drivers of microevolution in Neotropical palms: a historical genomics approach": outF5_AP03.1_pf_trimmed_masked_sickled_fastqc.html

outF5\_AP03.1\_pf\_trimmed\_masked\_sickled.fastq FastQC Report 

FastQC Report

mar. 25 juin 2019  
outF5\_AP03.1\_pf\_trimmed\_masked\_sickled.fastq

### Summary

- Basic Statistics
- Per base sequence quality
- Per tile sequence quality
- Per sequence quality scores
- Per base sequence content
- Per sequence GC content
- Per base N content
- Sequence Length Distribution
- Sequence Duplication Levels
- Overrepresented sequences
- Adapter Content
- Kmer Content

### Basic Statistics

| Measure | Value |
| --- | --- |
| Filename | outF5\_AP03.1\_pf\_trimmed\_masked\_sickled.fastq |
| File type | Conventional base calls |
| Encoding | Sanger / Illumina 1.9 |
| Total Sequences | 982342 |
| Sequences flagged as poor quality | 0 |
| Sequence length | 100-131 |
| %GC | 43 |

### Per base sequence quality

### Per tile sequence quality

### Per sequence quality scores

### Per base sequence content

### Per sequence GC content

### Per base N content

### Sequence Length Distribution

### Sequence Duplication Levels

### Overrepresented sequences

No overrepresented sequences

### Adapter Content

### Kmer Content

| Sequence | Count | PValue | Obs/Exp Max | Max Obs/Exp Position |
| --- | --- | --- | --- | --- |
| CTGAAGT | 955 | 0.0 | 19.530622 | 5 |
| TCGTTTA | 170 | 0.009086947 | 17.696152 | 5 |
| TGAAGTC | 1165 | 0.0 | 16.54288 | 6 |
| TCTGAAG | 1600 | 0.0 | 12.024005 | 4 |
| GAAGTCG | 525 | 4.0548148E-5 | 11.477674 | 7 |
| ATCTGAA | 1890 | 0.0 | 10.490606 | 3 |
| TTTCCCC | 530 | 5.781286E-4 | 10.202699 | 3 |
| CTTCCGA | 530 | 5.9548905E-4 | 10.169957 | 1 |
| CACGTCT | 370 | 3.5023046E-4 | 9.275445 | 124-125 |
| AAGTCGA | 525 | 0.005954781 | 9.185487 | 8 |
| CACACGT | 395 | 0.0069177924 | 7.7645106 | 122-123 |
| ACTTCAG | 1845 | 0.0 | 7.2033896 | 122-123 |
| CGATCTG | 3005 | 0.0 | 7.1748123 | 1 |
| TCGGAAG | 935 | 2.5626468E-7 | 6.973944 | 124-125 |
| GAAGTCA | 1010 | 0.0035978144 | 6.562729 | 7 |
| CGGAAGA | 865 | 3.1800828E-5 | 6.303366 | 122-123 |
| GACTTCA | 2045 | 0.0 | 6.165625 | 122-123 |
| GATCTGA | 4155 | 0.0 | 5.637177 | 2 |
| CTTCAGA | 2385 | 0.0 | 5.530087 | 120-121 |
| TTCAGAT | 2500 | 0.0 | 5.4910636 | 124-125 |

Produced by FastQC (version 0.11.5)
