## Supplementary material 4 for "Natural and human-mediated drivers of microevolution in Neotropical palms: a historical genomics approach": outF5_AP03.2_pf_trimmed_masked_sickled_fastqc.html

outF5\_AP03.2\_pf\_trimmed\_masked\_sickled.fastq FastQC Report 

FastQC Report

mar. 25 juin 2019  
outF5\_AP03.2\_pf\_trimmed\_masked\_sickled.fastq

### Summary

- Basic Statistics
- Per base sequence quality
- Per tile sequence quality
- Per sequence quality scores
- Per base sequence content
- Per sequence GC content
- Per base N content
- Sequence Length Distribution
- Sequence Duplication Levels
- Overrepresented sequences
- Adapter Content
- Kmer Content

### Basic Statistics

| Measure | Value |
| --- | --- |
| Filename | outF5\_AP03.2\_pf\_trimmed\_masked\_sickled.fastq |
| File type | Conventional base calls |
| Encoding | Sanger / Illumina 1.9 |
| Total Sequences | 982342 |
| Sequences flagged as poor quality | 0 |
| Sequence length | 100-131 |
| %GC | 44 |

### Per base sequence quality

### Per tile sequence quality

### Per sequence quality scores

### Per base sequence content

### Per sequence GC content

### Per base N content

### Sequence Length Distribution

### Sequence Duplication Levels

### Overrepresented sequences

No overrepresented sequences

### Adapter Content

### Kmer Content

| Sequence | Count | PValue | Obs/Exp Max | Max Obs/Exp Position |
| --- | --- | --- | --- | --- |
| CTGAAGT | 805 | 0.0 | 18.326134 | 5 |
| TGAAGTC | 1050 | 0.0 | 14.051553 | 6 |
| TCTGAAG | 1475 | 0.0 | 10.750777 | 4 |
| CTTCCGA | 450 | 0.00299833 | 9.981625 | 1 |
| ATCTGAA | 1775 | 0.0 | 9.554371 | 3 |
| CGTCGTG | 360 | 0.006461288 | 9.095134 | 124-125 |
| AGAGCGT | 480 | 0.0076805437 | 7.6740193 | 124-125 |
| ACTTCAG | 1830 | 0.0 | 7.662823 | 122-123 |
| GACTTCA | 2045 | 0.0 | 7.444957 | 122-123 |
| TCGGAAG | 920 | 5.2832347E-6 | 7.117931 | 124-125 |
| GAAGTCA | 810 | 0.005593649 | 7.004247 | 7 |
| AGACTTC | 905 | 3.2063406E-5 | 6.783663 | 124-125 |
| AAGAGCG | 605 | 0.00783653 | 6.7649755 | 124-125 |
| CTTCAGA | 2335 | 0.0 | 6.660676 | 124-125 |
| TTCAGAT | 2515 | 0.0 | 6.3467035 | 124-125 |
| CGGAAGA | 855 | 8.115533E-4 | 6.2229867 | 124-125 |
| CGATCTG | 2705 | 1.7826096E-10 | 6.0194182 | 1 |
| GGACTTC | 960 | 1.7840226E-4 | 5.9906487 | 118-119 |
| GATCTGA | 3745 | 0.0 | 5.574287 | 2 |

Produced by FastQC (version 0.11.5)
