## Supplementary material 4 for "Natural and human-mediated drivers of microevolution in Neotropical palms: a historical genomics approach": outF5_AP15.1_pf_trimmed_masked_sickled_fastqc.html

outF5\_AP15.1\_pf\_trimmed\_masked\_sickled.fastq FastQC Report 

FastQC Report

mar. 25 juin 2019  
outF5\_AP15.1\_pf\_trimmed\_masked\_sickled.fastq

### Summary

- Basic Statistics
- Per base sequence quality
- Per tile sequence quality
- Per sequence quality scores
- Per base sequence content
- Per sequence GC content
- Per base N content
- Sequence Length Distribution
- Sequence Duplication Levels
- Overrepresented sequences
- Adapter Content
- Kmer Content

### Basic Statistics

| Measure | Value |
| --- | --- |
| Filename | outF5\_AP15.1\_pf\_trimmed\_masked\_sickled.fastq |
| File type | Conventional base calls |
| Encoding | Sanger / Illumina 1.9 |
| Total Sequences | 1610212 |
| Sequences flagged as poor quality | 0 |
| Sequence length | 100-131 |
| %GC | 44 |

### Per base sequence quality

### Per tile sequence quality

### Per sequence quality scores

### Per base sequence content

### Per sequence GC content

### Per base N content

### Sequence Length Distribution

### Sequence Duplication Levels

### Overrepresented sequences

No overrepresented sequences

### Adapter Content

### Kmer Content

| Sequence | Count | PValue | Obs/Exp Max | Max Obs/Exp Position |
| --- | --- | --- | --- | --- |
| CTGACAG | 1160 | 0.0 | 16.095871 | 5 |
| TGACAGA | 1350 | 0.0 | 13.379717 | 6 |
| TCTGACA | 1440 | 0.0 | 12.549051 | 4 |
| ATCTGAC | 1670 | 0.0 | 11.902058 | 3 |
| CTTCCGA | 810 | 2.1082087E-9 | 11.853271 | 1 |
| CCCGCGA | 210 | 0.005872881 | 11.084556 | 116-117 |
| GCACACG | 555 | 3.697278E-8 | 9.867059 | 124-125 |
| ACACGTC | 455 | 0.0027940688 | 7.5222774 | 124-125 |
| TCGGAAG | 1415 | 0.0 | 7.498355 | 124-125 |
| CACACGT | 460 | 0.0031094768 | 7.440513 | 124-125 |
| GTCTGTC | 910 | 1.6768354E-7 | 7.1461635 | 124-125 |
| GACAGAT | 1185 | 5.12455E-5 | 7.1130414 | 7 |
| CTGTCAG | 2490 | 0.0 | 7.0102186 | 124-125 |
| TGTCAGA | 2485 | 0.0 | 6.2391324 | 120-121 |
| CGGAAGA | 1375 | 4.307367E-9 | 6.2229743 | 124-125 |
| TCTGTCA | 2900 | 0.0 | 6.1371403 | 124-125 |
| AGCACAC | 1095 | 4.0742325E-6 | 5.938821 | 124-125 |
| TTCCGAT | 1855 | 1.4210538E-5 | 5.8373623 | 2 |
| AGAGCAC | 1435 | 4.884314E-7 | 5.441361 | 122-123 |
| GTCAGAT | 2620 | 0.0 | 5.312686 | 122-123 |

Produced by FastQC (version 0.11.5)
