## Supplementary material 4 for "Natural and human-mediated drivers of microevolution in Neotropical palms: a historical genomics approach": outF5_AP15.2_pf_trimmed_masked_sickled_fastqc.html

outF5\_AP15.2\_pf\_trimmed\_masked\_sickled.fastq FastQC Report 

FastQC Report

mar. 25 juin 2019  
outF5\_AP15.2\_pf\_trimmed\_masked\_sickled.fastq

### Summary

- Basic Statistics
- Per base sequence quality
- Per tile sequence quality
- Per sequence quality scores
- Per base sequence content
- Per sequence GC content
- Per base N content
- Sequence Length Distribution
- Sequence Duplication Levels
- Overrepresented sequences
- Adapter Content
- Kmer Content

### Basic Statistics

| Measure | Value |
| --- | --- |
| Filename | outF5\_AP15.2\_pf\_trimmed\_masked\_sickled.fastq |
| File type | Conventional base calls |
| Encoding | Sanger / Illumina 1.9 |
| Total Sequences | 1610212 |
| Sequences flagged as poor quality | 0 |
| Sequence length | 100-131 |
| %GC | 44 |

### Per base sequence quality

### Per tile sequence quality

### Per sequence quality scores

### Per base sequence content

### Per sequence GC content

### Per base N content

### Sequence Length Distribution

### Sequence Duplication Levels

### Overrepresented sequences

No overrepresented sequences

### Adapter Content

### Kmer Content

| Sequence | Count | PValue | Obs/Exp Max | Max Obs/Exp Position |
| --- | --- | --- | --- | --- |
| CTGACAG | 1135 | 0.0 | 19.081844 | 5 |
| TCTGACA | 1415 | 0.0 | 16.080479 | 4 |
| ATCTGAC | 1535 | 0.0 | 15.203882 | 3 |
| TGACAGA | 1530 | 0.0 | 14.156412 | 6 |
| AGCGTCG | 550 | 1.4551915E-11 | 13.146602 | 124-125 |
| AGAGCGT | 615 | 1.03682396E-10 | 11.757124 | 124-125 |
| GAGCGTC | 560 | 3.5033736E-9 | 11.477193 | 124-125 |
| GACAGAC | 570 | 1.37390125E-5 | 10.9899435 | 7 |
| TCGGAAG | 1365 | 0.0 | 8.534323 | 124-125 |
| AAGAGCG | 880 | 4.9849405E-8 | 8.216627 | 124-125 |
| GCGTCGT | 440 | 0.004161169 | 8.216627 | 124-125 |
| GTCTGTC | 830 | 6.031951E-8 | 8.124663 | 118-119 |
| TGTCAGA | 2330 | 0.0 | 7.4133797 | 124-125 |
| TCTGTCA | 2570 | 0.0 | 7.346297 | 124-125 |
| CTGTCAG | 2435 | 0.0 | 7.2586756 | 124-125 |
| CGGAAGA | 1330 | 4.3837645E-10 | 7.248753 | 124-125 |
| GACAGAG | 1120 | 3.906007E-4 | 6.6100235 | 7 |
| CCTATGC | 660 | 0.0037447435 | 6.5385804 | 122-123 |
| GTCAGAT | 2540 | 0.0 | 6.3326416 | 122-123 |
| TTCACGA | 605 | 0.005816434 | 6.2683787 | 106-107 |

Produced by FastQC (version 0.11.5)
