## Supplementary material 4 for "Natural and human-mediated drivers of microevolution in Neotropical palms: a historical genomics approach": outF5_AP17.1_pf_trimmed_masked_sickled_fastqc.html

outF5\_AP17.1\_pf\_trimmed\_masked\_sickled.fastq FastQC Report 

FastQC Report

mar. 25 juin 2019  
outF5\_AP17.1\_pf\_trimmed\_masked\_sickled.fastq

### Summary

- Basic Statistics
- Per base sequence quality
- Per tile sequence quality
- Per sequence quality scores
- Per base sequence content
- Per sequence GC content
- Per base N content
- Sequence Length Distribution
- Sequence Duplication Levels
- Overrepresented sequences
- Adapter Content
- Kmer Content

### Basic Statistics

| Measure | Value |
| --- | --- |
| Filename | outF5\_AP17.1\_pf\_trimmed\_masked\_sickled.fastq |
| File type | Conventional base calls |
| Encoding | Sanger / Illumina 1.9 |
| Total Sequences | 1784633 |
| Sequences flagged as poor quality | 0 |
| Sequence length | 100-131 |
| %GC | 43 |

### Per base sequence quality

### Per tile sequence quality

### Per sequence quality scores

### Per base sequence content

### Per sequence GC content

### Per base N content

### Sequence Length Distribution

### Sequence Duplication Levels

### Overrepresented sequences

No overrepresented sequences

### Adapter Content

### Kmer Content

| Sequence | Count | PValue | Obs/Exp Max | Max Obs/Exp Position |
| --- | --- | --- | --- | --- |
| CTTCCGA | 945 | 2.687193E-8 | 10.071605 | 1 |
| CACACGT | 555 | 2.827801E-6 | 8.837236 | 124-125 |
| GATCTGC | 1940 | 2.928573E-10 | 7.3820343 | 2 |
| CTGCATT | 2005 | 5.7843863E-10 | 7.156679 | 5 |
| ACACGTC | 485 | 0.0048648594 | 7.106768 | 120-121 |
| GCACACG | 675 | 3.0436716E-4 | 6.747159 | 124-125 |
| TGCATTC | 2335 | 3.2923708E-10 | 6.658112 | 6 |
| ACGTCTG | 525 | 0.009673453 | 6.6194935 | 122-123 |
| CGGAAGA | 1470 | 5.638867E-11 | 6.5653005 | 120-121 |
| AGAGCAC | 1565 | 1.1514203E-9 | 6.044089 | 124-125 |
| ATCTGCA | 2600 | 4.090907E-9 | 5.968172 | 3 |
| AATGCAG | 3035 | 0.0 | 5.8869843 | 124-125 |
| GAATGCA | 3495 | 0.0 | 5.813828 | 124-125 |
| TCGGAAG | 1565 | 8.727511E-9 | 5.7735515 | 122-123 |
| TTCCGAT | 2020 | 3.5761323E-4 | 5.0218544 | 2 |

Produced by FastQC (version 0.11.5)
