## Supplementary material 4 for "Natural and human-mediated drivers of microevolution in Neotropical palms: a historical genomics approach": outF5_AP17.2_pf_trimmed_masked_sickled_fastqc.html

outF5\_AP17.2\_pf\_trimmed\_masked\_sickled.fastq FastQC Report 

FastQC Report

mar. 25 juin 2019  
outF5\_AP17.2\_pf\_trimmed\_masked\_sickled.fastq

### Summary

- Basic Statistics
- Per base sequence quality
- Per tile sequence quality
- Per sequence quality scores
- Per base sequence content
- Per sequence GC content
- Per base N content
- Sequence Length Distribution
- Sequence Duplication Levels
- Overrepresented sequences
- Adapter Content
- Kmer Content

### Basic Statistics

| Measure | Value |
| --- | --- |
| Filename | outF5\_AP17.2\_pf\_trimmed\_masked\_sickled.fastq |
| File type | Conventional base calls |
| Encoding | Sanger / Illumina 1.9 |
| Total Sequences | 1784633 |
| Sequences flagged as poor quality | 0 |
| Sequence length | 100-131 |
| %GC | 44 |

### Per base sequence quality

### Per tile sequence quality

### Per sequence quality scores

### Per base sequence content

### Per sequence GC content

### Per base N content

### Sequence Length Distribution

### Sequence Duplication Levels

### Overrepresented sequences

No overrepresented sequences

### Adapter Content

### Kmer Content

| Sequence | Count | PValue | Obs/Exp Max | Max Obs/Exp Position |
| --- | --- | --- | --- | --- |
| AGAGCGT | 585 | 2.1827873E-11 | 12.832886 | 124-125 |
| AGCGTCG | 595 | 9.593768E-7 | 9.569094 | 122-123 |
| TCGGAAG | 1385 | 5.0931703E-11 | 9.003246 | 7 |
| GATCTGC | 1650 | 0.0 | 8.883558 | 2 |
| GAGCGTC | 670 | 3.4127352E-6 | 8.7148695 | 124-125 |
| CGGAAGA | 1380 | 1.8371793E-10 | 8.616188 | 8 |
| ATCGGAA | 2730 | 0.0 | 8.315808 | 6 |
| AAGAGCG | 905 | 4.242611E-8 | 8.295291 | 124-125 |
| GCGTCGT | 485 | 0.007123127 | 7.7394204 | 124-125 |
| CTGCATT | 1795 | 1.3169483E-9 | 7.261049 | 5 |
| GAATGCA | 3335 | 0.0 | 6.5030212 | 124-125 |
| AATGCAG | 2985 | 0.0 | 6.42719 | 124-125 |
| GCAGATC | 3010 | 0.0 | 6.3681016 | 2 |
| CGAATGC | 855 | 3.1240442E-4 | 6.192635 | 116-117 |
| ATCTGCA | 2570 | 1.7516868E-9 | 5.937023 | 3 |
| TGCATTC | 2155 | 2.1774722E-6 | 5.530688 | 6 |
| TCTGCAT | 2675 | 2.6448106E-8 | 5.496141 | 4 |
| TGCAGAT | 3730 | 0.0 | 5.4114194 | 1 |

Produced by FastQC (version 0.11.5)
