## Supplementary material 4 for "Natural and human-mediated drivers of microevolution in Neotropical palms: a historical genomics approach": outF8_AP01.1_pf_trimmed_masked_sickled_fastqc.html

outF8\_AP01.1\_pf\_trimmed\_masked\_sickled.fastq FastQC Report 

FastQC Report

mar. 25 juin 2019  
outF8\_AP01.1\_pf\_trimmed\_masked\_sickled.fastq

### Summary

- Basic Statistics
- Per base sequence quality
- Per tile sequence quality
- Per sequence quality scores
- Per base sequence content
- Per sequence GC content
- Per base N content
- Sequence Length Distribution
- Sequence Duplication Levels
- Overrepresented sequences
- Adapter Content
- Kmer Content

### Basic Statistics

| Measure | Value |
| --- | --- |
| Filename | outF8\_AP01.1\_pf\_trimmed\_masked\_sickled.fastq |
| File type | Conventional base calls |
| Encoding | Sanger / Illumina 1.9 |
| Total Sequences | 823787 |
| Sequences flagged as poor quality | 0 |
| Sequence length | 100-131 |
| %GC | 43 |

### Per base sequence quality

### Per tile sequence quality

### Per sequence quality scores

### Per base sequence content

### Per sequence GC content

### Per base N content

### Sequence Length Distribution

### Sequence Duplication Levels

### Overrepresented sequences

No overrepresented sequences

### Adapter Content

### Kmer Content

| Sequence | Count | PValue | Obs/Exp Max | Max Obs/Exp Position |
| --- | --- | --- | --- | --- |
| ACGTCTG | 220 | 4.8445648E-4 | 12.41539 | 124-125 |
| CACGTCT | 270 | 2.106358E-4 | 11.3807745 | 124-125 |
| CACACGT | 330 | 1.0078222E-5 | 11.301104 | 122-123 |
| TCTAAGT | 520 | 4.580611E-4 | 10.463336 | 4 |
| TAAGTTG | 550 | 7.627167E-4 | 9.899337 | 6 |
| ACACGTC | 305 | 7.8940066E-4 | 9.863421 | 118-119 |
| GCACACG | 345 | 2.4372978E-4 | 9.616421 | 116-117 |
| CTAAGTT | 545 | 0.007845547 | 8.881787 | 5 |
| GAGGCGG | 535 | 2.2953379E-4 | 7.6044807 | 122-123 |
| AGCACAC | 555 | 3.2313677E-4 | 7.3821244 | 124-125 |
| CAACTTA | 1300 | 2.5465852E-11 | 7.0910983 | 124-125 |
| AACTTAG | 1290 | 1.5279511E-10 | 6.8813987 | 124-125 |
| GATCTAA | 1415 | 4.948617E-4 | 5.973644 | 2 |
| ACTTAGA | 1245 | 2.1879714E-7 | 5.9411197 | 120-121 |
| TCGGAAG | 750 | 0.0014851827 | 5.918003 | 124-125 |
| GCCAGAA | 650 | 0.005501134 | 5.7446337 | 68-69 |
| CTTAGAT | 1425 | 2.1926207E-6 | 5.2710958 | 124-125 |

Produced by FastQC (version 0.11.5)
