## Supplementary material 4 for "Natural and human-mediated drivers of microevolution in Neotropical palms: a historical genomics approach": outF8_AP01.2_pf_trimmed_masked_sickled_fastqc.html

outF8\_AP01.2\_pf\_trimmed\_masked\_sickled.fastq FastQC Report 

FastQC Report

mar. 25 juin 2019  
outF8\_AP01.2\_pf\_trimmed\_masked\_sickled.fastq

### Summary

- Basic Statistics
- Per base sequence quality
- Per tile sequence quality
- Per sequence quality scores
- Per base sequence content
- Per sequence GC content
- Per base N content
- Sequence Length Distribution
- Sequence Duplication Levels
- Overrepresented sequences
- Adapter Content
- Kmer Content

### Basic Statistics

| Measure | Value |
| --- | --- |
| Filename | outF8\_AP01.2\_pf\_trimmed\_masked\_sickled.fastq |
| File type | Conventional base calls |
| Encoding | Sanger / Illumina 1.9 |
| Total Sequences | 823787 |
| Sequences flagged as poor quality | 0 |
| Sequence length | 100-131 |
| %GC | 44 |

### Per base sequence quality

### Per tile sequence quality

### Per sequence quality scores

### Per base sequence content

### Per sequence GC content

### Per base N content

### Sequence Length Distribution

### Sequence Duplication Levels

### Overrepresented sequences

No overrepresented sequences

### Adapter Content

### Kmer Content

| Sequence | Count | PValue | Obs/Exp Max | Max Obs/Exp Position |
| --- | --- | --- | --- | --- |
| AGCGTCG | 290 | 7.589412E-5 | 12.697015 | 124-125 |
| AGAGCGT | 355 | 3.559171E-6 | 12.406506 | 122-123 |
| TAGGGAA | 305 | 0.0016431243 | 10.731174 | 124-125 |
| TCTAAGT | 535 | 9.0458656E-5 | 10.606702 | 4 |
| GAGCGTC | 340 | 9.030918E-4 | 9.719482 | 114-115 |
| CTAGCCA | 475 | 0.00418338 | 9.587248 | 6 |
| TAAGTTG | 615 | 3.5712804E-4 | 9.255981 | 6 |
| CTAAGTT | 625 | 4.2348905E-4 | 9.09971 | 5 |
| CTTAGAT | 1225 | 5.456968E-12 | 8.349511 | 124-125 |
| CAACTTA | 1225 | 4.7293724E-11 | 8.015531 | 124-125 |
| TCGGAAG | 630 | 1.7283499E-4 | 7.792877 | 124-125 |
| AAGAGCG | 545 | 0.0035162193 | 7.3466372 | 122-123 |
| ACTTAGA | 1115 | 3.3123797E-8 | 7.3385835 | 124-125 |
| AACTTAG | 1175 | 1.1230441E-8 | 7.31204 | 124-125 |
| CGATCTA | 810 | 0.005929256 | 6.962018 | 1 |
| CGGAAGA | 625 | 0.0027878752 | 6.724568 | 118-119 |
| GGCCACC | 880 | 0.0026105675 | 5.6443295 | 118-119 |
| GCAACTT | 955 | 0.0030593122 | 5.569255 | 124-125 |
| TAGATCG | 1120 | 5.976216E-4 | 5.4793663 | 124-125 |
| AGGGGAT | 785 | 0.0066468916 | 5.214368 | 86-87 |

Produced by FastQC (version 0.11.5)
