## Supplementary material 4 for "Natural and human-mediated drivers of microevolution in Neotropical palms: a historical genomics approach": outF8_AP06.1_pf_trimmed_masked_sickled_fastqc.html

outF8\_AP06.1\_pf\_trimmed\_masked\_sickled.fastq FastQC Report 

FastQC Report

mar. 25 juin 2019  
outF8\_AP06.1\_pf\_trimmed\_masked\_sickled.fastq

### Summary

- Basic Statistics
- Per base sequence quality
- Per tile sequence quality
- Per sequence quality scores
- Per base sequence content
- Per sequence GC content
- Per base N content
- Sequence Length Distribution
- Sequence Duplication Levels
- Overrepresented sequences
- Adapter Content
- Kmer Content

### Basic Statistics

| Measure | Value |
| --- | --- |
| Filename | outF8\_AP06.1\_pf\_trimmed\_masked\_sickled.fastq |
| File type | Conventional base calls |
| Encoding | Sanger / Illumina 1.9 |
| Total Sequences | 747145 |
| Sequences flagged as poor quality | 0 |
| Sequence length | 100-131 |
| %GC | 44 |

### Per base sequence quality

### Per tile sequence quality

### Per sequence quality scores

### Per base sequence content

### Per sequence GC content

### Per base N content

### Sequence Length Distribution

### Sequence Duplication Levels

### Overrepresented sequences

No overrepresented sequences

### Adapter Content

### Kmer Content

| Sequence | Count | PValue | Obs/Exp Max | Max Obs/Exp Position |
| --- | --- | --- | --- | --- |
| CTAATAC | 480 | 0.0 | 27.793783 | 5 |
| TAATACA | 605 | 0.0 | 23.050465 | 6 |
| CGATCTA | 800 | 0.0 | 16.619783 | 1 |
| TCTAATA | 890 | 0.0 | 16.334858 | 4 |
| ACACGTC | 210 | 3.692924E-4 | 12.821384 | 124-125 |
| CACGTCT | 215 | 4.5037334E-4 | 12.523213 | 124-125 |
| ATCTAAT | 1355 | 0.0 | 10.734271 | 3 |
| CTTCCGA | 395 | 0.0075731394 | 10.710101 | 1 |
| GATCTAA | 1260 | 1.8189894E-12 | 10.568711 | 2 |
| GCACACG | 260 | 0.002340571 | 10.284726 | 122-123 |
| CACACGT | 265 | 0.0025893168 | 10.160342 | 124-125 |
| GTATTAG | 915 | 7.2759576E-12 | 8.699509 | 120-121 |
| AGCACAC | 560 | 5.472104E-6 | 8.414033 | 124-125 |
| TGTATTA | 1035 | 1.1823431E-10 | 7.69087 | 120-121 |
| GAGCACA | 760 | 5.655622E-6 | 7.0855017 | 124-125 |
| TAGATCG | 1060 | 6.574919E-8 | 6.6677246 | 124-125 |
| TCGGAAG | 750 | 4.680709E-5 | 6.6017137 | 118-119 |
| TATTAGA | 1245 | 3.2772732E-8 | 6.2175994 | 124-125 |
| GGATTCT | 830 | 0.00215879 | 5.320072 | 90-91 |
| GGAAGAG | 1410 | 3.1092168E-6 | 5.174974 | 120-121 |

Produced by FastQC (version 0.11.5)
