## Supplementary material 4 for "Natural and human-mediated drivers of microevolution in Neotropical palms: a historical genomics approach": outF8_AP06.2_pf_trimmed_masked_sickled_fastqc.html

outF8\_AP06.2\_pf\_trimmed\_masked\_sickled.fastq FastQC Report 

FastQC Report

mar. 25 juin 2019  
outF8\_AP06.2\_pf\_trimmed\_masked\_sickled.fastq

### Summary

- Basic Statistics
- Per base sequence quality
- Per tile sequence quality
- Per sequence quality scores
- Per base sequence content
- Per sequence GC content
- Per base N content
- Sequence Length Distribution
- Sequence Duplication Levels
- Overrepresented sequences
- Adapter Content
- Kmer Content

### Basic Statistics

| Measure | Value |
| --- | --- |
| Filename | outF8\_AP06.2\_pf\_trimmed\_masked\_sickled.fastq |
| File type | Conventional base calls |
| Encoding | Sanger / Illumina 1.9 |
| Total Sequences | 747145 |
| Sequences flagged as poor quality | 0 |
| Sequence length | 100-131 |
| %GC | 45 |

### Per base sequence quality

### Per tile sequence quality

### Per sequence quality scores

### Per base sequence content

### Per sequence GC content

### Per base N content

### Sequence Length Distribution

### Sequence Duplication Levels

### Overrepresented sequences

No overrepresented sequences

### Adapter Content

### Kmer Content

| Sequence | Count | PValue | Obs/Exp Max | Max Obs/Exp Position |
| --- | --- | --- | --- | --- |
| CTAATAC | 480 | 0.0 | 29.384827 | 5 |
| TAATACA | 575 | 0.0 | 25.529142 | 6 |
| CGATCTA | 630 | 0.0 | 20.407393 | 1 |
| TCTAATA | 850 | 0.0 | 15.227621 | 4 |
| CTTCCGA | 450 | 7.406379E-8 | 14.90627 | 1 |
| AATACAC | 325 | 1.8761655E-4 | 13.88772 | 7 |
| AATACAG | 390 | 5.9935533E-5 | 13.019738 | 7 |
| GAGCGTC | 310 | 1.4579065E-4 | 11.839318 | 124-125 |
| GATCTAA | 1225 | 0.0 | 11.458298 | 2 |
| AGCGTCG | 290 | 0.0011079523 | 11.2496195 | 124-125 |
| AAGAGCG | 445 | 3.948595E-6 | 10.738457 | 122-123 |
| ATCTAAT | 1510 | 0.0 | 9.309359 | 3 |
| AGAGCGT | 345 | 0.0069960696 | 9.0071745 | 120-121 |
| GTATTAG | 910 | 1.9863364E-9 | 8.53702 | 120-121 |
| CTGTATT | 430 | 0.0045737806 | 8.130023 | 120-121 |
| TCGGAAG | 615 | 3.3730597E-5 | 8.059373 | 118-119 |
| TAGATCG | 985 | 3.0802767E-8 | 7.8661685 | 124-125 |
| TGTATTA | 1000 | 7.429062E-7 | 6.9918194 | 120-121 |
| TTCCGAT | 895 | 0.002121232 | 6.9005837 | 2 |
| CGGAAGA | 560 | 0.008805389 | 6.6835384 | 116-117 |

Produced by FastQC (version 0.11.5)
