## Supplementary material 4 for "Natural and human-mediated drivers of microevolution in Neotropical palms: a historical genomics approach": outF8_AP12.1_pf_trimmed_masked_sickled_fastqc.html

outF8\_AP12.1\_pf\_trimmed\_masked\_sickled.fastq FastQC Report 

FastQC Report

mar. 25 juin 2019  
outF8\_AP12.1\_pf\_trimmed\_masked\_sickled.fastq

### Summary

- Basic Statistics
- Per base sequence quality
- Per tile sequence quality
- Per sequence quality scores
- Per base sequence content
- Per sequence GC content
- Per base N content
- Sequence Length Distribution
- Sequence Duplication Levels
- Overrepresented sequences
- Adapter Content
- Kmer Content

### Basic Statistics

| Measure | Value |
| --- | --- |
| Filename | outF8\_AP12.1\_pf\_trimmed\_masked\_sickled.fastq |
| File type | Conventional base calls |
| Encoding | Sanger / Illumina 1.9 |
| Total Sequences | 944369 |
| Sequences flagged as poor quality | 0 |
| Sequence length | 100-131 |
| %GC | 44 |

### Per base sequence quality

### Per tile sequence quality

### Per sequence quality scores

### Per base sequence content

### Per sequence GC content

### Per base N content

### Sequence Length Distribution

### Sequence Duplication Levels

### Overrepresented sequences

No overrepresented sequences

### Adapter Content

### Kmer Content

| Sequence | Count | PValue | Obs/Exp Max | Max Obs/Exp Position |
| --- | --- | --- | --- | --- |
| GATCTAC | 875 | 0.0 | 24.994778 | 2 |
| CTACAAT | 935 | 0.0 | 21.450836 | 5 |
| CGATCTA | 975 | 0.0 | 21.134981 | 1 |
| TACAATA | 1005 | 0.0 | 20.560389 | 6 |
| TCTACAA | 1135 | 0.0 | 18.733849 | 4 |
| ATCTACA | 1110 | 0.0 | 18.61148 | 3 |
| CTTCCGA | 555 | 2.3250323E-8 | 14.19639 | 1 |
| ACACGTC | 235 | 6.556956E-5 | 12.89771 | 124-125 |
| ACAATAG | 615 | 1.5521819E-5 | 10.868408 | 7 |
| CACACGT | 325 | 1.1474159E-4 | 10.362263 | 124-125 |
| GCACACG | 305 | 7.7738916E-4 | 9.8800955 | 122-123 |
| ACAATAT | 1150 | 4.12565E-8 | 8.982546 | 7 |
| CTTTAGC | 550 | 0.008171987 | 8.837965 | 3 |
| ATTGTAG | 1400 | 0.0 | 7.457128 | 124-125 |
| GTAGATC | 1110 | 3.0213414E-9 | 6.97819 | 124-125 |
| TATTGTA | 1500 | 3.6379788E-12 | 6.7354712 | 124-125 |
| TTCCGAT | 1345 | 0.0016289885 | 5.8718624 | 2 |
| TTGTAGA | 1495 | 6.2809704E-9 | 5.856931 | 124-125 |
| TGTAGAT | 1340 | 1.4464422E-7 | 5.7804413 | 124-125 |
| CGGAAGA | 700 | 0.0052067162 | 5.7732606 | 124-125 |

Produced by FastQC (version 0.11.5)
