## Supplementary material 4 for "Natural and human-mediated drivers of microevolution in Neotropical palms: a historical genomics approach": outF8_AP12.2_pf_trimmed_masked_sickled_fastqc.html

outF8\_AP12.2\_pf\_trimmed\_masked\_sickled.fastq FastQC Report 

FastQC Report

mar. 25 juin 2019  
outF8\_AP12.2\_pf\_trimmed\_masked\_sickled.fastq

### Summary

- Basic Statistics
- Per base sequence quality
- Per tile sequence quality
- Per sequence quality scores
- Per base sequence content
- Per sequence GC content
- Per base N content
- Sequence Length Distribution
- Sequence Duplication Levels
- Overrepresented sequences
- Adapter Content
- Kmer Content

### Basic Statistics

| Measure | Value |
| --- | --- |
| Filename | outF8\_AP12.2\_pf\_trimmed\_masked\_sickled.fastq |
| File type | Conventional base calls |
| Encoding | Sanger / Illumina 1.9 |
| Total Sequences | 944369 |
| Sequences flagged as poor quality | 0 |
| Sequence length | 100-131 |
| %GC | 44 |

### Per base sequence quality

### Per tile sequence quality

### Per sequence quality scores

### Per base sequence content

### Per sequence GC content

### Per base N content

### Sequence Length Distribution

### Sequence Duplication Levels

### Overrepresented sequences

No overrepresented sequences

### Adapter Content

### Kmer Content

| Sequence | Count | PValue | Obs/Exp Max | Max Obs/Exp Position |
| --- | --- | --- | --- | --- |
| GATCTAC | 855 | 0.0 | 21.820301 | 2 |
| CGATCTA | 830 | 0.0 | 20.990177 | 1 |
| CTACAAT | 885 | 0.0 | 20.539057 | 5 |
| TACAATA | 920 | 0.0 | 20.381952 | 6 |
| ATCTACA | 1010 | 0.0 | 18.518053 | 3 |
| TCTACAA | 1110 | 0.0 | 15.835636 | 4 |
| AGCGTCG | 295 | 8.625406E-6 | 13.346782 | 122-123 |
| GCGTCGT | 230 | 0.003280755 | 11.983089 | 122-123 |
| GTCGTGT | 245 | 0.0044208537 | 11.514276 | 124-125 |
| ACAATAC | 450 | 2.1350695E-4 | 11.363209 | 7 |
| AGAGCGT | 320 | 3.3836393E-4 | 10.813195 | 120-121 |
| CGTCGTG | 310 | 0.0021334104 | 10.399991 | 124-125 |
| TATTGTA | 1360 | 1.8189894E-12 | 8.000728 | 124-125 |
| ACAATAA | 1135 | 7.695895E-6 | 7.5087285 | 7 |
| ATTGTAG | 1320 | 3.43789E-10 | 7.3272667 | 124-125 |
| GTAGATC | 1105 | 3.7083737E-8 | 7.2941113 | 124-125 |
| TCGGAAG | 745 | 2.4351885E-4 | 6.8704576 | 122-123 |
| GTGGAAA | 830 | 0.007123861 | 6.831542 | 4 |
| TGTAGAT | 1250 | 4.9141818E-8 | 6.7703943 | 124-125 |
| CGGAAGA | 730 | 0.0011123777 | 6.6246514 | 124-125 |

Produced by FastQC (version 0.11.5)
