## Supplementary material 4 for "Natural and human-mediated drivers of microevolution in Neotropical palms: a historical genomics approach": outF8_AP16.1_pf_trimmed_masked_sickled_fastqc.html

outF8\_AP16.1\_pf\_trimmed\_masked\_sickled.fastq FastQC Report 

FastQC Report

mar. 25 juin 2019  
outF8\_AP16.1\_pf\_trimmed\_masked\_sickled.fastq

### Summary

- Basic Statistics
- Per base sequence quality
- Per tile sequence quality
- Per sequence quality scores
- Per base sequence content
- Per sequence GC content
- Per base N content
- Sequence Length Distribution
- Sequence Duplication Levels
- Overrepresented sequences
- Adapter Content
- Kmer Content

### Basic Statistics

| Measure | Value |
| --- | --- |
| Filename | outF8\_AP16.1\_pf\_trimmed\_masked\_sickled.fastq |
| File type | Conventional base calls |
| Encoding | Sanger / Illumina 1.9 |
| Total Sequences | 910918 |
| Sequences flagged as poor quality | 0 |
| Sequence length | 100-131 |
| %GC | 43 |

### Per base sequence quality

### Per tile sequence quality

### Per sequence quality scores

### Per base sequence content

### Per sequence GC content

### Per base N content

### Sequence Length Distribution

### Sequence Duplication Levels

### Overrepresented sequences

No overrepresented sequences

### Adapter Content

### Kmer Content

| Sequence | Count | PValue | Obs/Exp Max | Max Obs/Exp Position |
| --- | --- | --- | --- | --- |
| TACCACA | 630 | 0.0 | 25.803263 | 6 |
| CTACCAC | 710 | 0.0 | 22.89842 | 5 |
| GATCTAC | 870 | 0.0 | 17.963848 | 2 |
| TCTACCA | 915 | 0.0 | 17.744274 | 4 |
| ATCTACC | 885 | 0.0 | 17.667294 | 3 |
| CGATCTA | 910 | 0.0 | 17.126257 | 1 |
| CTTCCGA | 465 | 1.2331679E-5 | 12.890732 | 1 |
| GTGGTAG | 1315 | 0.0 | 10.449957 | 124-125 |
| ACCACAA | 985 | 4.190224E-8 | 9.784858 | 7 |
| ACGTCTG | 285 | 0.003990595 | 9.643292 | 124-125 |
| TGTGGTA | 1445 | 0.0 | 9.272077 | 124-125 |
| TGGTAGA | 1580 | 0.0 | 8.697273 | 124-125 |
| GGTAGAT | 1385 | 0.0 | 7.689394 | 124-125 |
| TCGGAAG | 850 | 2.9783168E-6 | 6.8708463 | 124-125 |
| GTAGATC | 1300 | 7.839844E-9 | 6.342319 | 124-125 |
| TAGATCG | 1460 | 4.0745363E-10 | 6.3121543 | 122-123 |
| AGAGCAC | 835 | 1.187629E-4 | 6.171418 | 124-125 |
| CGGAAGA | 790 | 3.886686E-4 | 6.0880914 | 124-125 |
| TCAGCAC | 650 | 0.003387995 | 6.000545 | 106-107 |
| TTGTGGT | 1220 | 2.3522542E-5 | 5.350249 | 124-125 |

Produced by FastQC (version 0.11.5)
