## Supplementary material 4 for "Natural and human-mediated drivers of microevolution in Neotropical palms: a historical genomics approach": outF8_AP16.2_pf_trimmed_masked_sickled_fastqc.html

outF8\_AP16.2\_pf\_trimmed\_masked\_sickled.fastq FastQC Report 

FastQC Report

mar. 25 juin 2019  
outF8\_AP16.2\_pf\_trimmed\_masked\_sickled.fastq

### Summary

- Basic Statistics
- Per base sequence quality
- Per tile sequence quality
- Per sequence quality scores
- Per base sequence content
- Per sequence GC content
- Per base N content
- Sequence Length Distribution
- Sequence Duplication Levels
- Overrepresented sequences
- Adapter Content
- Kmer Content

### Basic Statistics

| Measure | Value |
| --- | --- |
| Filename | outF8\_AP16.2\_pf\_trimmed\_masked\_sickled.fastq |
| File type | Conventional base calls |
| Encoding | Sanger / Illumina 1.9 |
| Total Sequences | 910918 |
| Sequences flagged as poor quality | 0 |
| Sequence length | 100-131 |
| %GC | 43 |

### Per base sequence quality

### Per tile sequence quality

### Per sequence quality scores

### Per base sequence content

### Per sequence GC content

### Per base N content

### Sequence Length Distribution

### Sequence Duplication Levels

### Overrepresented sequences

No overrepresented sequences

### Adapter Content

### Kmer Content

| Sequence | Count | PValue | Obs/Exp Max | Max Obs/Exp Position |
| --- | --- | --- | --- | --- |
| CTACCAC | 510 | 0.0 | 19.982132 | 5 |
| TACCACA | 520 | 0.0 | 19.61152 | 6 |
| GATCTAC | 660 | 0.0 | 15.372917 | 2 |
| CGTGTAG | 175 | 0.0054882676 | 14.166362 | 124-125 |
| ATCTACC | 820 | 0.0 | 13.7688265 | 3 |
| CGATCTA | 865 | 0.0 | 12.966224 | 1 |
| TCTACCA | 930 | 3.6561687E-10 | 10.930659 | 4 |
| GTGGTAG | 1235 | 0.0 | 10.706022 | 124-125 |
| TCCGCAC | 275 | 0.008669934 | 10.51745 | 124-125 |
| GTGTGGT | 585 | 7.565177E-8 | 10.351127 | 122-123 |
| AGAGCGT | 335 | 0.0033000077 | 9.867117 | 124-125 |
| TGTGGTA | 1315 | 0.0 | 9.740495 | 124-125 |
| GGTAGAT | 1325 | 0.0 | 9.3551445 | 124-125 |
| TGGTAGA | 1460 | 0.0 | 8.20711 | 124-125 |
| AAGAGCG | 580 | 5.4964895E-4 | 7.836278 | 124-125 |
| TCGGAAG | 740 | 1.46580915E-5 | 7.8170238 | 124-125 |
| CGGAAGA | 765 | 5.164169E-6 | 7.724064 | 120-121 |
| TAGATCG | 1360 | 1.0913936E-11 | 7.7176776 | 122-123 |
| ATGTGGT | 860 | 1.3743958E-5 | 7.206725 | 124-125 |
| AGAGTGT | 500 | 0.004426471 | 7.175461 | 110-111 |

Produced by FastQC (version 0.11.5)
