## Supplementary material 4 for "Natural and human-mediated drivers of microevolution in Neotropical palms: a historical genomics approach": outF8_AP19.1_pf_trimmed_masked_sickled_fastqc.html

outF8\_AP19.1\_pf\_trimmed\_masked\_sickled.fastq FastQC Report 

FastQC Report

mar. 25 juin 2019  
outF8\_AP19.1\_pf\_trimmed\_masked\_sickled.fastq

### Summary

- Basic Statistics
- Per base sequence quality
- Per tile sequence quality
- Per sequence quality scores
- Per base sequence content
- Per sequence GC content
- Per base N content
- Sequence Length Distribution
- Sequence Duplication Levels
- Overrepresented sequences
- Adapter Content
- Kmer Content

### Basic Statistics

| Measure | Value |
| --- | --- |
| Filename | outF8\_AP19.1\_pf\_trimmed\_masked\_sickled.fastq |
| File type | Conventional base calls |
| Encoding | Sanger / Illumina 1.9 |
| Total Sequences | 800867 |
| Sequences flagged as poor quality | 0 |
| Sequence length | 100-131 |
| %GC | 44 |

### Per base sequence quality

### Per tile sequence quality

### Per sequence quality scores

### Per base sequence content

### Per sequence GC content

### Per base N content

### Sequence Length Distribution

### Sequence Duplication Levels

### Overrepresented sequences

No overrepresented sequences

### Adapter Content

### Kmer Content

| Sequence | Count | PValue | Obs/Exp Max | Max Obs/Exp Position |
| --- | --- | --- | --- | --- |
| TACTCAC | 500 | 0.0 | 50.517162 | 6 |
| CTACTCA | 615 | 0.0 | 42.04874 | 5 |
| TCTACTC | 765 | 0.0 | 34.581207 | 4 |
| ACTCACG | 235 | 0.0 | 33.291992 | 7 |
| GATCTAC | 835 | 0.0 | 32.35687 | 2 |
| CGATCTA | 945 | 0.0 | 29.155296 | 1 |
| ATCTACT | 965 | 0.0 | 26.143032 | 3 |
| ACTCACA | 435 | 1.0131771E-9 | 17.985329 | 7 |
| ACTCACT | 480 | 1.9826984E-10 | 17.55299 | 7 |
| CTCACGC | 255 | 0.0054435055 | 14.183014 | 8 |
| CTACGGA | 140 | 0.0070666075 | 13.636348 | 98-99 |
| CACACGT | 360 | 2.2180711E-5 | 10.524024 | 124-125 |
| CACGTCT | 290 | 0.0045097233 | 9.501314 | 124-125 |
| ACACGTC | 315 | 0.009425868 | 8.685723 | 122-123 |
| GCACACG | 360 | 0.0027241611 | 8.610565 | 124-125 |
| GTGAGTA | 975 | 1.202352E-9 | 7.7169313 | 122-123 |
| TGAGTAG | 1035 | 5.0386006E-10 | 7.600008 | 122-123 |
| TTGCGCA | 620 | 7.244669E-6 | 7.5420113 | 82-83 |
| AGAGCAC | 780 | 5.9045797E-6 | 7.065079 | 124-125 |
| GAGTAGA | 1200 | 1.3224053E-9 | 6.888453 | 124-125 |

Produced by FastQC (version 0.11.5)
