## Supplementary material 4 for "Natural and human-mediated drivers of microevolution in Neotropical palms: a historical genomics approach": outF8_AP19.2_pf_trimmed_masked_sickled_fastqc.html

outF8\_AP19.2\_pf\_trimmed\_masked\_sickled.fastq FastQC Report 

FastQC Report

mar. 25 juin 2019  
outF8\_AP19.2\_pf\_trimmed\_masked\_sickled.fastq

### Summary

- Basic Statistics
- Per base sequence quality
- Per tile sequence quality
- Per sequence quality scores
- Per base sequence content
- Per sequence GC content
- Per base N content
- Sequence Length Distribution
- Sequence Duplication Levels
- Overrepresented sequences
- Adapter Content
- Kmer Content

### Basic Statistics

| Measure | Value |
| --- | --- |
| Filename | outF8\_AP19.2\_pf\_trimmed\_masked\_sickled.fastq |
| File type | Conventional base calls |
| Encoding | Sanger / Illumina 1.9 |
| Total Sequences | 800867 |
| Sequences flagged as poor quality | 0 |
| Sequence length | 100-131 |
| %GC | 45 |

### Per base sequence quality

### Per tile sequence quality

### Per sequence quality scores

### Per base sequence content

### Per sequence GC content

### Per base N content

### Sequence Length Distribution

### Sequence Duplication Levels

### Overrepresented sequences

No overrepresented sequences

### Adapter Content

### Kmer Content

| Sequence | Count | PValue | Obs/Exp Max | Max Obs/Exp Position |
| --- | --- | --- | --- | --- |
| TACTCAC | 555 | 0.0 | 44.98408 | 6 |
| CTACTCA | 570 | 0.0 | 44.66821 | 5 |
| TCTACTC | 715 | 0.0 | 33.196075 | 4 |
| CGATCTA | 815 | 0.0 | 30.983885 | 1 |
| GATCTAC | 785 | 0.0 | 30.178036 | 2 |
| ATCTACT | 860 | 0.0 | 27.593588 | 3 |
| ACTCACG | 205 | 1.2658165E-7 | 24.834946 | 7 |
| ACTCACT | 355 | 0.0 | 23.902176 | 7 |
| ACTCACA | 615 | 3.6379788E-12 | 15.636819 | 7 |
| CTCACAT | 530 | 8.643736E-5 | 10.654255 | 8 |
| GGTTTTC | 565 | 1.6696475E-4 | 9.983042 | 2 |
| AGAGCGT | 450 | 3.7258392E-4 | 9.21755 | 124-125 |
| GCGGGGG | 265 | 0.0058390214 | 9.207978 | 54-55 |
| AAGAGCG | 635 | 1.7736184E-6 | 9.144972 | 124-125 |
| TGAGTAG | 1075 | 2.7284841E-11 | 8.700561 | 122-123 |
| GTGAGTA | 950 | 1.884473E-9 | 8.561193 | 122-123 |
| AGCGTCG | 415 | 0.0028737085 | 8.559568 | 120-121 |
| GAGCGTC | 455 | 0.004212484 | 8.204634 | 124-125 |
| GAGTAGA | 1070 | 1.3969839E-9 | 8.140735 | 124-125 |
| TGTGAGT | 615 | 1.1136901E-4 | 8.09346 | 124-125 |

Produced by FastQC (version 0.11.5)
