## Supplementary material 4 for "Natural and human-mediated drivers of microevolution in Neotropical palms: a historical genomics approach": outF9_AP2.1_pf_trimmed_masked_sickled_fastqc.html

outF9\_AP2.1\_pf\_trimmed\_masked\_sickled.fastq FastQC Report 

FastQC Report

mar. 25 juin 2019  
outF9\_AP2.1\_pf\_trimmed\_masked\_sickled.fastq

### Summary

- Basic Statistics
- Per base sequence quality
- Per tile sequence quality
- Per sequence quality scores
- Per base sequence content
- Per sequence GC content
- Per base N content
- Sequence Length Distribution
- Sequence Duplication Levels
- Overrepresented sequences
- Adapter Content
- Kmer Content

### Basic Statistics

| Measure | Value |
| --- | --- |
| Filename | outF9\_AP2.1\_pf\_trimmed\_masked\_sickled.fastq |
| File type | Conventional base calls |
| Encoding | Sanger / Illumina 1.9 |
| Total Sequences | 578960 |
| Sequences flagged as poor quality | 0 |
| Sequence length | 100-131 |
| %GC | 43 |

### Per base sequence quality

### Per tile sequence quality

### Per sequence quality scores

### Per base sequence content

### Per sequence GC content

### Per base N content

### Sequence Length Distribution

### Sequence Duplication Levels

### Overrepresented sequences

No overrepresented sequences

### Adapter Content

### Kmer Content

| Sequence | Count | PValue | Obs/Exp Max | Max Obs/Exp Position |
| --- | --- | --- | --- | --- |
| CAGCGCG | 115 | 0.0021838727 | 16.2698 | 94-95 |
| GAATTAT | 550 | 0.0065238387 | 9.082789 | 3 |
| GCCTTCA | 680 | 0.0014273718 | 5.9373245 | 74-75 |
| GCGCAGC | 815 | 0.0020084882 | 5.350402 | 82-83 |
| CCTGATG | 790 | 0.0075710034 | 5.1563444 | 30-31 |
| TTGCGCA | 800 | 0.009333279 | 5.064603 | 80-81 |

Produced by FastQC (version 0.11.5)
