## Supplementary material 4 for "Natural and human-mediated drivers of microevolution in Neotropical palms: a historical genomics approach": outF9_AP2.2_pf_trimmed_masked_sickled_fastqc.html

outF9\_AP2.2\_pf\_trimmed\_masked\_sickled.fastq FastQC Report 

FastQC Report

mar. 25 juin 2019  
outF9\_AP2.2\_pf\_trimmed\_masked\_sickled.fastq

### Summary

- Basic Statistics
- Per base sequence quality
- Per tile sequence quality
- Per sequence quality scores
- Per base sequence content
- Per sequence GC content
- Per base N content
- Sequence Length Distribution
- Sequence Duplication Levels
- Overrepresented sequences
- Adapter Content
- Kmer Content

### Basic Statistics

| Measure | Value |
| --- | --- |
| Filename | outF9\_AP2.2\_pf\_trimmed\_masked\_sickled.fastq |
| File type | Conventional base calls |
| Encoding | Sanger / Illumina 1.9 |
| Total Sequences | 578960 |
| Sequences flagged as poor quality | 0 |
| Sequence length | 100-131 |
| %GC | 43 |

### Per base sequence quality

### Per tile sequence quality

### Per sequence quality scores

### Per base sequence content

### Per sequence GC content

### Per base N content

### Sequence Length Distribution

### Sequence Duplication Levels

### Overrepresented sequences

No overrepresented sequences

### Adapter Content

### Kmer Content

| Sequence | Count | PValue | Obs/Exp Max | Max Obs/Exp Position |
| --- | --- | --- | --- | --- |
| CTAGCCG | 95 | 5.955258E-4 | 19.747234 | 80-81 |
| TGAGGCA | 585 | 7.2759576E-12 | 16.796793 | 6 |
| CTGAGGC | 480 | 2.8630893E-9 | 16.631235 | 5 |
| TCTGAGG | 545 | 1.537228E-8 | 14.646349 | 4 |
| CTATTGG | 340 | 0.002224358 | 12.618421 | 2 |
| CTATCCG | 410 | 0.009441929 | 10.395471 | 1 |
| ATCTGAG | 840 | 3.922014E-7 | 10.211169 | 3 |
| TATCCGA | 495 | 0.0031913035 | 9.90537 | 2 |
| CATTTGC | 680 | 4.6708074E-4 | 9.009854 | 3 |
| CTCACAG | 1010 | 4.6676814E-6 | 8.5111685 | 4 |
| CACAGCC | 1025 | 4.36064E-4 | 7.1898465 | 6 |
| TCACAGC | 1045 | 5.4463267E-4 | 7.051595 | 5 |
| ACAGCCT | 920 | 0.009044351 | 6.663731 | 7 |
| ATCTCAC | 1220 | 4.5052185E-4 | 6.530845 | 2 |
| CGATCTG | 1530 | 2.7264878E-5 | 6.3673472 | 1 |
| GATCTGA | 1945 | 1.3547574E-4 | 5.3569202 | 2 |
| GCCTGTA | 770 | 0.0072976355 | 5.172601 | 10-11 |
| TGTTTTT | 1555 | 0.008775002 | 5.090295 | 1 |

Produced by FastQC (version 0.11.5)
