## Supplementary material 4 for "Natural and human-mediated drivers of microevolution in Neotropical palms: a historical genomics approach": outF9_AP5.1_pf_trimmed_masked_sickled_fastqc.html

outF9\_AP5.1\_pf\_trimmed\_masked\_sickled.fastq FastQC Report 

FastQC Report

mar. 25 juin 2019  
outF9\_AP5.1\_pf\_trimmed\_masked\_sickled.fastq

### Summary

- Basic Statistics
- Per base sequence quality
- Per tile sequence quality
- Per sequence quality scores
- Per base sequence content
- Per sequence GC content
- Per base N content
- Sequence Length Distribution
- Sequence Duplication Levels
- Overrepresented sequences
- Adapter Content
- Kmer Content

### Basic Statistics

| Measure | Value |
| --- | --- |
| Filename | outF9\_AP5.1\_pf\_trimmed\_masked\_sickled.fastq |
| File type | Conventional base calls |
| Encoding | Sanger / Illumina 1.9 |
| Total Sequences | 520323 |
| Sequences flagged as poor quality | 0 |
| Sequence length | 100-131 |
| %GC | 43 |

### Per base sequence quality

### Per tile sequence quality

### Per sequence quality scores

### Per base sequence content

### Per sequence GC content

### Per base N content

### Sequence Length Distribution

### Sequence Duplication Levels

### Overrepresented sequences

No overrepresented sequences

### Adapter Content

### Kmer Content

| Sequence | Count | PValue | Obs/Exp Max | Max Obs/Exp Position |
| --- | --- | --- | --- | --- |
| TGGTACG | 250 | 0.0 | 75.04986 | 6 |
| GGTACGC | 85 | 6.2937033E-10 | 58.87455 | 7 |
| GTACGGT | 60 | 1.6142672E-5 | 52.21308 | 8 |
| GGTACGG | 100 | 2.6575435E-9 | 50.04337 | 7 |
| CTGGTAC | 395 | 0.0 | 49.10312 | 5 |
| GTACGCA | 75 | 6.0331797E-5 | 41.77047 | 8 |
| GGTACGT | 105 | 2.200104E-7 | 41.70281 | 7 |
| GTACGTT | 80 | 8.8258486E-5 | 39.15981 | 8 |
| GGTACGA | 115 | 4.4911576E-7 | 38.076477 | 7 |
| GTACGAG | 110 | 5.725433E-4 | 28.479864 | 8 |
| ATCTGGT | 700 | 0.0 | 27.680162 | 3 |
| TCTGGTA | 720 | 0.0 | 26.92216 | 4 |
| GATCTGG | 1460 | 0.0 | 13.608565 | 2 |
| CGATCTG | 1750 | 0.0 | 11.656689 | 1 |
| GTTGTGT | 255 | 0.0034141697 | 9.826353 | 102-103 |
| GCTTGCG | 450 | 0.0066146813 | 6.88426 | 68-69 |
| GATCGCA | 935 | 0.008498706 | 6.7065845 | 9 |
| TCATTGG | 1525 | 8.0010126E-4 | 5.7519984 | 8 |
| TTGCGCA | 790 | 0.0014121591 | 5.50184 | 70-71 |
| CATTGGA | 1735 | 6.864483E-4 | 5.421317 | 9 |

Produced by FastQC (version 0.11.5)
