## Supplementary material 4 for "Natural and human-mediated drivers of microevolution in Neotropical palms: a historical genomics approach": outF9_AP5.2_pf_trimmed_masked_sickled_fastqc.html

outF9\_AP5.2\_pf\_trimmed\_masked\_sickled.fastq FastQC Report 

FastQC Report

mar. 25 juin 2019  
outF9\_AP5.2\_pf\_trimmed\_masked\_sickled.fastq

### Summary

- Basic Statistics
- Per base sequence quality
- Per tile sequence quality
- Per sequence quality scores
- Per base sequence content
- Per sequence GC content
- Per base N content
- Sequence Length Distribution
- Sequence Duplication Levels
- Overrepresented sequences
- Adapter Content
- Kmer Content

### Basic Statistics

| Measure | Value |
| --- | --- |
| Filename | outF9\_AP5.2\_pf\_trimmed\_masked\_sickled.fastq |
| File type | Conventional base calls |
| Encoding | Sanger / Illumina 1.9 |
| Total Sequences | 520323 |
| Sequences flagged as poor quality | 0 |
| Sequence length | 100-131 |
| %GC | 43 |

### Per base sequence quality

### Per tile sequence quality

### Per sequence quality scores

### Per base sequence content

### Per sequence GC content

### Per base N content

### Sequence Length Distribution

### Sequence Duplication Levels

### Overrepresented sequences

No overrepresented sequences

### Adapter Content

### Kmer Content

| Sequence | Count | PValue | Obs/Exp Max | Max Obs/Exp Position |
| --- | --- | --- | --- | --- |
| TGGTACG | 305 | 0.0 | 68.25807 | 6 |
| GGTACGG | 100 | 0.0 | 67.12114 | 7 |
| CTGGTAC | 375 | 0.0 | 55.55636 | 5 |
| GGTACGC | 125 | 9.094947E-12 | 48.81538 | 7 |
| GTACGCT | 75 | 0.0034630424 | 32.527 | 8 |
| ATCTGGT | 685 | 0.0 | 32.15374 | 3 |
| TCTGGTA | 675 | 0.0 | 31.762669 | 4 |
| GGTACGT | 100 | 3.8259933E-4 | 30.50961 | 7 |
| GGTACGA | 170 | 3.5419725E-7 | 28.71493 | 7 |
| GTACGAT | 115 | 8.691764E-4 | 26.516575 | 8 |
| GATCTGG | 1360 | 0.0 | 16.64835 | 2 |
| CTTTTCG | 240 | 0.0032889443 | 15.298485 | 2 |
| CGATCTG | 1615 | 0.0 | 13.561705 | 1 |
| CAGCCTG | 940 | 1.740948E-4 | 7.7857184 | 8 |
| AGTCAGA | 410 | 0.0024721078 | 7.615706 | 60-61 |
| TCACAGC | 995 | 3.179137E-4 | 7.3899975 | 5 |
| ACAGCCT | 1010 | 3.9619725E-4 | 7.249809 | 7 |
| CTCACAG | 1040 | 5.299615E-4 | 7.068067 | 4 |
| CTTGCGC | 455 | 0.0051660757 | 7.0618496 | 112-113 |
| CACAGCC | 965 | 0.0018776458 | 6.9797573 | 6 |

Produced by FastQC (version 0.11.5)
