## Supplementary material 4 for "Natural and human-mediated drivers of microevolution in Neotropical palms: a historical genomics approach": outF9_AP6.2_pf_trimmed_masked_sickled_fastqc.html

outF9\_AP6.2\_pf\_trimmed\_masked\_sickled.fastq FastQC Report 

FastQC Report

mar. 25 juin 2019  
outF9\_AP6.2\_pf\_trimmed\_masked\_sickled.fastq

### Summary

- Basic Statistics
- Per base sequence quality
- Per tile sequence quality
- Per sequence quality scores
- Per base sequence content
- Per sequence GC content
- Per base N content
- Sequence Length Distribution
- Sequence Duplication Levels
- Overrepresented sequences
- Adapter Content
- Kmer Content

### Basic Statistics

| Measure | Value |
| --- | --- |
| Filename | outF9\_AP6.2\_pf\_trimmed\_masked\_sickled.fastq |
| File type | Conventional base calls |
| Encoding | Sanger / Illumina 1.9 |
| Total Sequences | 485039 |
| Sequences flagged as poor quality | 0 |
| Sequence length | 100-131 |
| %GC | 43 |

### Per base sequence quality

### Per tile sequence quality

### Per sequence quality scores

### Per base sequence content

### Per sequence GC content

### Per base N content

### Sequence Length Distribution

### Sequence Duplication Levels

### Overrepresented sequences

No overrepresented sequences

### Adapter Content

### Kmer Content

| Sequence | Count | PValue | Obs/Exp Max | Max Obs/Exp Position |
| --- | --- | --- | --- | --- |
| CGATCTT | 350 | 1.0793412E-5 | 15.613368 | 1 |
| CTTGTGT | 385 | 2.4880908E-5 | 14.293379 | 5 |
| CTTCCGA | 255 | 0.005182338 | 14.28674 | 1 |
| TTGTGTG | 405 | 6.0340896E-4 | 12.092365 | 6 |
| TCTTGTG | 615 | 1.7197034E-4 | 9.9519005 | 4 |
| ATCTTGT | 690 | 5.5632595E-4 | 8.852716 | 3 |
| GATCTTG | 775 | 1.8075081E-4 | 8.6851425 | 2 |
| CCTTCGT | 620 | 4.2551337E-4 | 6.5628643 | 58-59 |
| CGTTTTG | 590 | 0.0019139815 | 6.3133674 | 62-63 |
| TTCGTTT | 615 | 0.0030449007 | 6.0573997 | 60-61 |
| TTGCGCA | 745 | 5.597811E-4 | 5.916646 | 92-93 |
| GCGCAGC | 735 | 0.003111919 | 5.5609436 | 94-95 |

Produced by FastQC (version 0.11.5)
