## Supplementary material 4 for "Natural and human-mediated drivers of microevolution in Neotropical palms: a historical genomics approach": outF9_AP7.1_pf_trimmed_masked_sickled_fastqc.html

outF9\_AP7.1\_pf\_trimmed\_masked\_sickled.fastq FastQC Report 

FastQC Report

mar. 25 juin 2019  
outF9\_AP7.1\_pf\_trimmed\_masked\_sickled.fastq

### Summary

- Basic Statistics
- Per base sequence quality
- Per tile sequence quality
- Per sequence quality scores
- Per base sequence content
- Per sequence GC content
- Per base N content
- Sequence Length Distribution
- Sequence Duplication Levels
- Overrepresented sequences
- Adapter Content
- Kmer Content

### Basic Statistics

| Measure | Value |
| --- | --- |
| Filename | outF9\_AP7.1\_pf\_trimmed\_masked\_sickled.fastq |
| File type | Conventional base calls |
| Encoding | Sanger / Illumina 1.9 |
| Total Sequences | 507557 |
| Sequences flagged as poor quality | 0 |
| Sequence length | 100-131 |
| %GC | 43 |

### Per base sequence quality

### Per tile sequence quality

### Per sequence quality scores

### Per base sequence content

### Per sequence GC content

### Per base N content

### Sequence Length Distribution

### Sequence Duplication Levels

### Overrepresented sequences

No overrepresented sequences

### Adapter Content

### Kmer Content

| Sequence | Count | PValue | Obs/Exp Max | Max Obs/Exp Position |
| --- | --- | --- | --- | --- |
| ACGCTTA | 80 | 0.005117345 | 19.54758 | 36-37 |
| CCCCGCA | 80 | 0.005191206 | 19.499197 | 92-93 |
| TCGATAA | 245 | 0.003230689 | 15.33954 | 9 |
| CGACCGG | 175 | 0.0024401718 | 12.465344 | 78-79 |
| TGAGTGG | 275 | 0.006444679 | 9.09757 | 52-53 |
| TCATTAT | 690 | 0.0048904205 | 8.068068 | 1 |
| GGCGGCA | 520 | 4.0817173E-4 | 7.2328997 | 108-109 |
| TCCGGGC | 480 | 0.001382433 | 7.186394 | 104-105 |
| CTCACAG | 960 | 0.0014493832 | 7.152657 | 4 |
| CAGCCTG | 885 | 0.005121867 | 7.066591 | 8 |
| TCACAGC | 1010 | 0.0024587284 | 6.8034782 | 5 |
| CATCGAT | 465 | 0.008012376 | 6.749077 | 42-43 |
| AGCCTGT | 930 | 0.008156993 | 6.7351027 | 9 |
| ATCTCAC | 1155 | 0.0014764138 | 6.458869 | 2 |
| CATTTTT | 1260 | 5.9574476E-4 | 6.3818836 | 1 |
| AAGTTTT | 905 | 2.2039434E-4 | 5.5126677 | 86-87 |
| TTCGTTT | 740 | 0.003686517 | 5.4819865 | 98-99 |
| TATATAT | 1715 | 7.0580293E-4 | 5.410074 | 1 |
| AATCTCA | 1610 | 0.0018718126 | 5.378711 | 1 |

Produced by FastQC (version 0.11.5)
