## Supplementary material 4 for "Natural and human-mediated drivers of microevolution in Neotropical palms: a historical genomics approach": outF9_AP8.1_pf_trimmed_masked_sickled_fastqc.html

outF9\_AP8.1\_pf\_trimmed\_masked\_sickled.fastq FastQC Report 

FastQC Report

mar. 25 juin 2019  
outF9\_AP8.1\_pf\_trimmed\_masked\_sickled.fastq

### Summary

- Basic Statistics
- Per base sequence quality
- Per tile sequence quality
- Per sequence quality scores
- Per base sequence content
- Per sequence GC content
- Per base N content
- Sequence Length Distribution
- Sequence Duplication Levels
- Overrepresented sequences
- Adapter Content
- Kmer Content

### Basic Statistics

| Measure | Value |
| --- | --- |
| Filename | outF9\_AP8.1\_pf\_trimmed\_masked\_sickled.fastq |
| File type | Conventional base calls |
| Encoding | Sanger / Illumina 1.9 |
| Total Sequences | 657107 |
| Sequences flagged as poor quality | 0 |
| Sequence length | 100-131 |
| %GC | 44 |

### Per base sequence quality

### Per tile sequence quality

### Per sequence quality scores

### Per base sequence content

### Per sequence GC content

### Per base N content

### Sequence Length Distribution

### Sequence Duplication Levels

### Overrepresented sequences

No overrepresented sequences

### Adapter Content

### Kmer Content

| Sequence | Count | PValue | Obs/Exp Max | Max Obs/Exp Position |
| --- | --- | --- | --- | --- |
| CTAATAC | 495 | 0.0 | 30.277029 | 5 |
| TAATACA | 575 | 0.0 | 26.04166 | 6 |
| CGATCTA | 690 | 0.0 | 22.39444 | 1 |
| CTTCCGA | 415 | 1.6370905E-11 | 20.851114 | 1 |
| TCTAATA | 755 | 0.0 | 20.661083 | 4 |
| AATACAC | 315 | 6.0959293E-5 | 15.846723 | 7 |
| GATCTAA | 1090 | 0.0 | 14.246221 | 2 |
| GCTTAGA | 265 | 0.005756176 | 14.063438 | 2 |
| ATCTAAT | 1205 | 0.0 | 12.945325 | 3 |
| ATACACT | 345 | 0.0021639795 | 12.665214 | 8 |
| TTCCGAT | 850 | 1.9645086E-10 | 12.422705 | 2 |
| AATACAG | 425 | 7.713864E-4 | 11.745217 | 7 |
| CGCCTCA | 190 | 0.004536865 | 11.474371 | 94-95 |
| CCATTAC | 370 | 0.008658013 | 7.571816 | 48-49 |
| CCGATCT | 1915 | 4.860748E-4 | 5.213282 | 4 |

Produced by FastQC (version 0.11.5)
