## Supplementary material 4 for "Natural and human-mediated drivers of microevolution in Neotropical palms: a historical genomics approach": outF9_AP8.2_pf_trimmed_masked_sickled_fastqc.html

outF9\_AP8.2\_pf\_trimmed\_masked\_sickled.fastq FastQC Report 

FastQC Report

mar. 25 juin 2019  
outF9\_AP8.2\_pf\_trimmed\_masked\_sickled.fastq

### Summary

- Basic Statistics
- Per base sequence quality
- Per tile sequence quality
- Per sequence quality scores
- Per base sequence content
- Per sequence GC content
- Per base N content
- Sequence Length Distribution
- Sequence Duplication Levels
- Overrepresented sequences
- Adapter Content
- Kmer Content

### Basic Statistics

| Measure | Value |
| --- | --- |
| Filename | outF9\_AP8.2\_pf\_trimmed\_masked\_sickled.fastq |
| File type | Conventional base calls |
| Encoding | Sanger / Illumina 1.9 |
| Total Sequences | 657107 |
| Sequences flagged as poor quality | 0 |
| Sequence length | 100-131 |
| %GC | 44 |

### Per base sequence quality

### Per tile sequence quality

### Per sequence quality scores

### Per base sequence content

### Per sequence GC content

### Per base N content

### Sequence Length Distribution

### Sequence Duplication Levels

### Overrepresented sequences

No overrepresented sequences

### Adapter Content

### Kmer Content

| Sequence | Count | PValue | Obs/Exp Max | Max Obs/Exp Position |
| --- | --- | --- | --- | --- |
| CTAATAC | 505 | 0.0 | 42.34604 | 5 |
| TAATACA | 595 | 0.0 | 36.007015 | 6 |
| CGATCTA | 620 | 0.0 | 34.209625 | 1 |
| TCTAATA | 805 | 0.0 | 27.31735 | 4 |
| GATCTAA | 1050 | 0.0 | 19.738735 | 2 |
| AATACAC | 320 | 2.0347761E-7 | 19.114939 | 7 |
| ATCTAAT | 1275 | 0.0 | 16.261917 | 3 |
| AATACAG | 465 | 1.4567227E-4 | 11.838929 | 7 |
| AATACAA | 775 | 1.8177033E-4 | 8.681881 | 7 |

Produced by FastQC (version 0.11.5)
