## Supplementary material 4 for "Natural and human-mediated drivers of microevolution in Neotropical palms: a historical genomics approach": outF9_AP9.1_pf_trimmed_masked_sickled_fastqc.html

outF9\_AP9.1\_pf\_trimmed\_masked\_sickled.fastq FastQC Report 

FastQC Report

mar. 25 juin 2019  
outF9\_AP9.1\_pf\_trimmed\_masked\_sickled.fastq

### Summary

- Basic Statistics
- Per base sequence quality
- Per tile sequence quality
- Per sequence quality scores
- Per base sequence content
- Per sequence GC content
- Per base N content
- Sequence Length Distribution
- Sequence Duplication Levels
- Overrepresented sequences
- Adapter Content
- Kmer Content

### Basic Statistics

| Measure | Value |
| --- | --- |
| Filename | outF9\_AP9.1\_pf\_trimmed\_masked\_sickled.fastq |
| File type | Conventional base calls |
| Encoding | Sanger / Illumina 1.9 |
| Total Sequences | 675860 |
| Sequences flagged as poor quality | 0 |
| Sequence length | 100-131 |
| %GC | 44 |

### Per base sequence quality

### Per tile sequence quality

### Per sequence quality scores

### Per base sequence content

### Per sequence GC content

### Per base N content

### Sequence Length Distribution

### Sequence Duplication Levels

### Overrepresented sequences

No overrepresented sequences

### Adapter Content

### Kmer Content

| Sequence | Count | PValue | Obs/Exp Max | Max Obs/Exp Position |
| --- | --- | --- | --- | --- |
| CTTCCGA | 475 | 0.0 | 35.160065 | 1 |
| GATCTAC | 650 | 0.0 | 25.93501 | 6 |
| CGATCTA | 680 | 0.0 | 25.714981 | 5 |
| CTACAAT | 735 | 0.0 | 22.985636 | 9 |
| ATCTACA | 800 | 0.0 | 20.29489 | 7 |
| TTCCGAT | 955 | 0.0 | 18.876217 | 2 |
| TCTACAA | 990 | 0.0 | 17.04919 | 8 |
| TACAATA | 725 | 0.0 | 12.917821 | 6 |
| GCGACAT | 225 | 0.0012621413 | 11.076134 | 76-77 |
| TTTACAG | 455 | 0.0013599079 | 10.97608 | 4 |
| CCGATCT | 2005 | 0.0 | 9.029263 | 4 |
| ACAATAC | 515 | 4.5913024E-5 | 7.863213 | 10-11 |

Produced by FastQC (version 0.11.5)
