## Supplementary material 4 for "Natural and human-mediated drivers of microevolution in Neotropical palms: a historical genomics approach": outF9_AP9.2_pf_trimmed_masked_sickled_fastqc.html

outF9\_AP9.2\_pf\_trimmed\_masked\_sickled.fastq FastQC Report 

FastQC Report

mar. 25 juin 2019  
outF9\_AP9.2\_pf\_trimmed\_masked\_sickled.fastq

### Summary

- Basic Statistics
- Per base sequence quality
- Per tile sequence quality
- Per sequence quality scores
- Per base sequence content
- Per sequence GC content
- Per base N content
- Sequence Length Distribution
- Sequence Duplication Levels
- Overrepresented sequences
- Adapter Content
- Kmer Content

### Basic Statistics

| Measure | Value |
| --- | --- |
| Filename | outF9\_AP9.2\_pf\_trimmed\_masked\_sickled.fastq |
| File type | Conventional base calls |
| Encoding | Sanger / Illumina 1.9 |
| Total Sequences | 675860 |
| Sequences flagged as poor quality | 0 |
| Sequence length | 100-131 |
| %GC | 44 |

### Per base sequence quality

### Per tile sequence quality

### Per sequence quality scores

### Per base sequence content

### Per sequence GC content

### Per base N content

### Sequence Length Distribution

### Sequence Duplication Levels

### Overrepresented sequences

No overrepresented sequences

### Adapter Content

### Kmer Content

| Sequence | Count | PValue | Obs/Exp Max | Max Obs/Exp Position |
| --- | --- | --- | --- | --- |
| CTTCCGA | 400 | 0.0 | 33.46798 | 1 |
| GATCTAC | 565 | 0.0 | 21.672321 | 2 |
| CGATCTA | 645 | 0.0 | 19.811913 | 1 |
| CTACAAT | 650 | 0.0 | 18.848497 | 5 |
| ATCTACA | 785 | 0.0 | 17.166405 | 3 |
| TACAATA | 735 | 0.0 | 16.670034 | 6 |
| TTCCGAT | 820 | 0.0 | 15.679396 | 2 |
| TCTACAA | 795 | 0.0 | 15.415514 | 4 |
| TAGGCGA | 145 | 6.787389E-4 | 14.770249 | 10-11 |
| TAATTTG | 735 | 0.009449571 | 7.496854 | 2 |
| CCGATCT | 1950 | 1.0912117E-8 | 6.9132657 | 4 |
| TTTTGAC | 675 | 0.007985653 | 5.553903 | 54-55 |
| TCGTTTT | 850 | 5.736441E-4 | 5.49629 | 60-61 |
| CCTTCGT | 905 | 0.0012657539 | 5.180687 | 58-59 |

Produced by FastQC (version 0.11.5)
