## Supplementary material 4 for "Natural and human-mediated drivers of microevolution in Neotropical palms: a historical genomics approach": outF9_AP10.1_pf_trimmed_masked_sickled_fastqc.html

outF9\_AP10.1\_pf\_trimmed\_masked\_sickled.fastq FastQC Report 

FastQC Report

mar. 25 juin 2019  
outF9\_AP10.1\_pf\_trimmed\_masked\_sickled.fastq

### Summary

- Basic Statistics
- Per base sequence quality
- Per tile sequence quality
- Per sequence quality scores
- Per base sequence content
- Per sequence GC content
- Per base N content
- Sequence Length Distribution
- Sequence Duplication Levels
- Overrepresented sequences
- Adapter Content
- Kmer Content

### Basic Statistics

| Measure | Value |
| --- | --- |
| Filename | outF9\_AP10.1\_pf\_trimmed\_masked\_sickled.fastq |
| File type | Conventional base calls |
| Encoding | Sanger / Illumina 1.9 |
| Total Sequences | 763560 |
| Sequences flagged as poor quality | 0 |
| Sequence length | 100-131 |
| %GC | 44 |

### Per base sequence quality

### Per tile sequence quality

### Per sequence quality scores

### Per base sequence content

### Per sequence GC content

### Per base N content

### Sequence Length Distribution

### Sequence Duplication Levels

### Overrepresented sequences

No overrepresented sequences

### Adapter Content

### Kmer Content

| Sequence | Count | PValue | Obs/Exp Max | Max Obs/Exp Position |
| --- | --- | --- | --- | --- |
| CGATCTA | 705 | 0.0 | 15.819356 | 1 |
| TCTAGAA | 720 | 0.0 | 15.653936 | 4 |
| TAGAAGG | 740 | 1.8189894E-12 | 15.236098 | 6 |
| CTAGAAG | 740 | 1.6370905E-11 | 14.381729 | 5 |
| GATCTAG | 815 | 5.456968E-12 | 13.765777 | 2 |
| ATCTAGA | 920 | 3.6379788E-12 | 12.918179 | 3 |
| CCAACGT | 220 | 0.0010709629 | 11.296065 | 62-63 |
| TTCCGAT | 1025 | 4.6180972E-5 | 7.858233 | 1 |
| AGTATCC | 375 | 0.009189037 | 7.521653 | 108-109 |
| TCCAACG | 440 | 0.0049572606 | 7.0924864 | 60-61 |
| TTGCAGT | 770 | 9.303211E-4 | 5.686471 | 52-53 |

Produced by FastQC (version 0.11.5)
