## Supplementary material 4 for "Natural and human-mediated drivers of microevolution in Neotropical palms: a historical genomics approach": outF9_AP10.2_pf_trimmed_masked_sickled_fastqc.html

outF9\_AP10.2\_pf\_trimmed\_masked\_sickled.fastq FastQC Report 

FastQC Report

mar. 25 juin 2019  
outF9\_AP10.2\_pf\_trimmed\_masked\_sickled.fastq

### Summary

- Basic Statistics
- Per base sequence quality
- Per tile sequence quality
- Per sequence quality scores
- Per base sequence content
- Per sequence GC content
- Per base N content
- Sequence Length Distribution
- Sequence Duplication Levels
- Overrepresented sequences
- Adapter Content
- Kmer Content

### Basic Statistics

| Measure | Value |
| --- | --- |
| Filename | outF9\_AP10.2\_pf\_trimmed\_masked\_sickled.fastq |
| File type | Conventional base calls |
| Encoding | Sanger / Illumina 1.9 |
| Total Sequences | 763560 |
| Sequences flagged as poor quality | 0 |
| Sequence length | 100-131 |
| %GC | 44 |

### Per base sequence quality

### Per tile sequence quality

### Per sequence quality scores

### Per base sequence content

### Per sequence GC content

### Per base N content

### Sequence Length Distribution

### Sequence Duplication Levels

### Overrepresented sequences

No overrepresented sequences

### Adapter Content

### Kmer Content

| Sequence | Count | PValue | Obs/Exp Max | Max Obs/Exp Position |
| --- | --- | --- | --- | --- |
| CGATCTA | 590 | 7.1323666E-7 | 12.377517 | 1 |
| TCTAGAA | 605 | 8.715124E-7 | 12.17411 | 4 |
| TAGAAGG | 640 | 2.2010707E-5 | 10.529656 | 6 |
| CTAGAAG | 710 | 6.8517635E-5 | 9.496109 | 5 |
| GATCTAG | 730 | 9.436002E-5 | 9.221934 | 2 |
| ATCTAGA | 775 | 1.8138063E-4 | 8.684073 | 3 |
| ACCCATT | 565 | 0.008531219 | 6.0406027 | 64-65 |
| CCGACCC | 675 | 0.0063841506 | 5.6677837 | 108-109 |

Produced by FastQC (version 0.11.5)
