## Supplementary material 4 for "Natural and human-mediated drivers of microevolution in Neotropical palms: a historical genomics approach": outF9_AP11.1_pf_trimmed_masked_sickled_fastqc.html

outF9\_AP11.1\_pf\_trimmed\_masked\_sickled.fastq FastQC Report 

FastQC Report

mar. 25 juin 2019  
outF9\_AP11.1\_pf\_trimmed\_masked\_sickled.fastq

### Summary

- Basic Statistics
- Per base sequence quality
- Per tile sequence quality
- Per sequence quality scores
- Per base sequence content
- Per sequence GC content
- Per base N content
- Sequence Length Distribution
- Sequence Duplication Levels
- Overrepresented sequences
- Adapter Content
- Kmer Content

### Basic Statistics

| Measure | Value |
| --- | --- |
| Filename | outF9\_AP11.1\_pf\_trimmed\_masked\_sickled.fastq |
| File type | Conventional base calls |
| Encoding | Sanger / Illumina 1.9 |
| Total Sequences | 601115 |
| Sequences flagged as poor quality | 0 |
| Sequence length | 100-131 |
| %GC | 44 |

### Per base sequence quality

### Per tile sequence quality

### Per sequence quality scores

### Per base sequence content

### Per sequence GC content

### Per base N content

### Sequence Length Distribution

### Sequence Duplication Levels

### Overrepresented sequences

No overrepresented sequences

### Adapter Content

### Kmer Content

| Sequence | Count | PValue | Obs/Exp Max | Max Obs/Exp Position |
| --- | --- | --- | --- | --- |
| ACGTAAC | 35 | 0.0065127495 | 53.37029 | 2 |
| CTAGGTT | 475 | 0.0 | 28.98479 | 5 |
| TCTAGGT | 460 | 0.0 | 28.524647 | 4 |
| ATCTAGG | 485 | 0.0 | 28.35002 | 3 |
| TAGGTTC | 510 | 0.0 | 26.981497 | 6 |
| GATCTAG | 595 | 0.0 | 23.022478 | 2 |
| CGATCTA | 595 | 0.0 | 22.908989 | 1 |
| CTTCCGA | 395 | 8.5252395E-8 | 17.254238 | 1 |
| ACTCGGG | 225 | 0.0018610099 | 16.662014 | 4 |
| AGGTTCC | 515 | 3.1006083E-4 | 10.91357 | 7 |
| CTTGTCC | 415 | 0.008529012 | 10.539227 | 4 |
| TCGGAAG | 260 | 0.004068816 | 9.620253 | 84-85 |
| TTCCGAT | 900 | 7.500501E-4 | 7.610208 | 2 |
| CCGAGAA | 370 | 0.008724269 | 7.565251 | 78-79 |
| TCCGACC | 580 | 0.009701292 | 5.9656525 | 118-119 |

Produced by FastQC (version 0.11.5)
