## Supplementary material 4 for "Natural and human-mediated drivers of microevolution in Neotropical palms: a historical genomics approach": outF9_AP11.2_pf_trimmed_masked_sickled_fastqc.html

outF9\_AP11.2\_pf\_trimmed\_masked\_sickled.fastq FastQC Report 

FastQC Report

mar. 25 juin 2019  
outF9\_AP11.2\_pf\_trimmed\_masked\_sickled.fastq

### Summary

- Basic Statistics
- Per base sequence quality
- Per tile sequence quality
- Per sequence quality scores
- Per base sequence content
- Per sequence GC content
- Per base N content
- Sequence Length Distribution
- Sequence Duplication Levels
- Overrepresented sequences
- Adapter Content
- Kmer Content

### Basic Statistics

| Measure | Value |
| --- | --- |
| Filename | outF9\_AP11.2\_pf\_trimmed\_masked\_sickled.fastq |
| File type | Conventional base calls |
| Encoding | Sanger / Illumina 1.9 |
| Total Sequences | 601115 |
| Sequences flagged as poor quality | 0 |
| Sequence length | 100-131 |
| %GC | 44 |

### Per base sequence quality

### Per tile sequence quality

### Per sequence quality scores

### Per base sequence content

### Per sequence GC content

### Per base N content

### Sequence Length Distribution

### Sequence Duplication Levels

### Overrepresented sequences

No overrepresented sequences

### Adapter Content

### Kmer Content

| Sequence | Count | PValue | Obs/Exp Max | Max Obs/Exp Position |
| --- | --- | --- | --- | --- |
| CTAGGTT | 350 | 1.8189894E-12 | 24.53023 | 5 |
| TAGGTTC | 455 | 0.0 | 21.56693 | 6 |
| TCTAGGT | 455 | 7.094059E-11 | 18.857811 | 4 |
| CGATCTA | 490 | 1.0913936E-11 | 18.640491 | 1 |
| ATCTAGG | 490 | 2.0372681E-10 | 17.515436 | 3 |
| CTTCCGA | 385 | 7.755443E-8 | 17.397793 | 1 |
| GATCTAG | 605 | 2.5102054E-10 | 15.188676 | 2 |
| CGGAAGA | 245 | 0.0037397956 | 15.006642 | 8 |
| TATTCGT | 125 | 0.0039244764 | 14.900085 | 40-41 |
| AGTGCGA | 140 | 0.0089388145 | 13.159692 | 20-21 |
| TTCCGAT | 775 | 0.0017117319 | 7.9046445 | 2 |
| TGAGCAC | 410 | 0.00273803 | 7.5369954 | 42-43 |
| ACTTTCC | 690 | 0.0010897886 | 6.072314 | 114-115 |

Produced by FastQC (version 0.11.5)
