## Supplementary material 4 for "Natural and human-mediated drivers of microevolution in Neotropical palms: a historical genomics approach": outF9_AP12.2_pf_trimmed_masked_sickled_fastqc.html

outF9\_AP12.2\_pf\_trimmed\_masked\_sickled.fastq FastQC Report 

FastQC Report

mar. 25 juin 2019  
outF9\_AP12.2\_pf\_trimmed\_masked\_sickled.fastq

### Summary

- Basic Statistics
- Per base sequence quality
- Per tile sequence quality
- Per sequence quality scores
- Per base sequence content
- Per sequence GC content
- Per base N content
- Sequence Length Distribution
- Sequence Duplication Levels
- Overrepresented sequences
- Adapter Content
- Kmer Content

### Basic Statistics

| Measure | Value |
| --- | --- |
| Filename | outF9\_AP12.2\_pf\_trimmed\_masked\_sickled.fastq |
| File type | Conventional base calls |
| Encoding | Sanger / Illumina 1.9 |
| Total Sequences | 933166 |
| Sequences flagged as poor quality | 0 |
| Sequence length | 100-131 |
| %GC | 44 |

### Per base sequence quality

### Per tile sequence quality

### Per sequence quality scores

### Per base sequence content

### Per sequence GC content

### Per base N content

### Sequence Length Distribution

### Sequence Duplication Levels

### Overrepresented sequences

No overrepresented sequences

### Adapter Content

### Kmer Content

| Sequence | Count | PValue | Obs/Exp Max | Max Obs/Exp Position |
| --- | --- | --- | --- | --- |
| CTATGTA | 525 | 1.8189894E-12 | 18.709549 | 5 |
| CGATCTA | 570 | 0.0 | 18.175222 | 1 |
| TATGTAG | 620 | 1.8189894E-12 | 16.84634 | 6 |
| CGGTAGC | 165 | 0.0016899572 | 13.09072 | 34-35 |
| TCTATGT | 810 | 1.4497346E-9 | 12.136907 | 4 |
| ATCTATG | 940 | 1.03682396E-10 | 11.750993 | 3 |
| GACCCGG | 385 | 9.1734255E-5 | 9.247591 | 118-119 |
| CCCGGGA | 335 | 0.0023425904 | 8.755311 | 120-121 |
| GATCTAT | 1275 | 2.07001E-8 | 8.6507 | 2 |
| CCGGTCA | 1140 | 1.9116444E-4 | 7.0066676 | 4 |
| CCGACCC | 750 | 8.396055E-6 | 6.8999057 | 116-117 |
| CGGTCAT | 1620 | 8.111678E-6 | 6.4422345 | 5 |
| GGTCATT | 1840 | 1.3120534E-6 | 6.3443055 | 6 |
| CCGGGCG | 560 | 0.006470278 | 6.2041626 | 90-91 |
| CACAGCC | 1685 | 6.1189185E-4 | 5.4694 | 6 |
| CTCACAG | 1715 | 7.748786E-4 | 5.374031 | 4 |
| GTCATTG | 2065 | 5.746458E-5 | 5.347297 | 7 |
| GTTGCGC | 870 | 0.004310946 | 5.0337486 | 68-69 |

Produced by FastQC (version 0.11.5)
