## Supplementary material 4 for "Natural and human-mediated drivers of microevolution in Neotropical palms: a historical genomics approach": outF9_AP15.1_pf_trimmed_masked_sickled_fastqc.html

outF9\_AP15.1\_pf\_trimmed\_masked\_sickled.fastq FastQC Report 

FastQC Report

mar. 25 juin 2019  
outF9\_AP15.1\_pf\_trimmed\_masked\_sickled.fastq

### Summary

- Basic Statistics
- Per base sequence quality
- Per tile sequence quality
- Per sequence quality scores
- Per base sequence content
- Per sequence GC content
- Per base N content
- Sequence Length Distribution
- Sequence Duplication Levels
- Overrepresented sequences
- Adapter Content
- Kmer Content

### Basic Statistics

| Measure | Value |
| --- | --- |
| Filename | outF9\_AP15.1\_pf\_trimmed\_masked\_sickled.fastq |
| File type | Conventional base calls |
| Encoding | Sanger / Illumina 1.9 |
| Total Sequences | 774140 |
| Sequences flagged as poor quality | 0 |
| Sequence length | 100-131 |
| %GC | 44 |

### Per base sequence quality

### Per tile sequence quality

### Per sequence quality scores

### Per base sequence content

### Per sequence GC content

### Per base N content

### Sequence Length Distribution

### Sequence Duplication Levels

### Overrepresented sequences

No overrepresented sequences

### Adapter Content

### Kmer Content

| Sequence | Count | PValue | Obs/Exp Max | Max Obs/Exp Position |
| --- | --- | --- | --- | --- |
| TCCTGTA | 605 | 0.0 | 19.574348 | 6 |
| CTCCTGT | 740 | 0.0 | 19.364595 | 5 |
| TGTCCGC | 165 | 0.0063925907 | 18.80838 | 3 |
| CTTCGTA | 110 | 0.0016004636 | 17.044476 | 102-103 |
| TCTCCTG | 905 | 0.0 | 15.127123 | 4 |
| CCTGTAA | 295 | 6.7352573E-4 | 14.783877 | 7 |
| CGATCTC | 945 | 0.0 | 13.691021 | 1 |
| CCTGTAG | 340 | 0.001967455 | 12.827187 | 7 |
| CTTCCGA | 450 | 0.0013846485 | 10.952817 | 1 |
| GATCTCC | 1580 | 1.2732926E-11 | 9.012627 | 2 |
| ATCTCCT | 1485 | 3.1650416E-10 | 8.777245 | 3 |
| TGTCTCA | 445 | 0.0056468714 | 6.9980173 | 24-25 |
| ACACTTG | 550 | 8.793455E-4 | 6.763474 | 64-65 |
| GTTGCGC | 785 | 0.00661185 | 5.216757 | 112-113 |

Produced by FastQC (version 0.11.5)
