## Supplementary material 4 for "Natural and human-mediated drivers of microevolution in Neotropical palms: a historical genomics approach": outF9_AP15.2_pf_trimmed_masked_sickled_fastqc.html

outF9\_AP15.2\_pf\_trimmed\_masked\_sickled.fastq FastQC Report 

FastQC Report

mar. 25 juin 2019  
outF9\_AP15.2\_pf\_trimmed\_masked\_sickled.fastq

### Summary

- Basic Statistics
- Per base sequence quality
- Per tile sequence quality
- Per sequence quality scores
- Per base sequence content
- Per sequence GC content
- Per base N content
- Sequence Length Distribution
- Sequence Duplication Levels
- Overrepresented sequences
- Adapter Content
- Kmer Content

### Basic Statistics

| Measure | Value |
| --- | --- |
| Filename | outF9\_AP15.2\_pf\_trimmed\_masked\_sickled.fastq |
| File type | Conventional base calls |
| Encoding | Sanger / Illumina 1.9 |
| Total Sequences | 774140 |
| Sequences flagged as poor quality | 0 |
| Sequence length | 100-131 |
| %GC | 44 |

### Per base sequence quality

### Per tile sequence quality

### Per sequence quality scores

### Per base sequence content

### Per sequence GC content

### Per base N content

### Sequence Length Distribution

### Sequence Duplication Levels

### Overrepresented sequences

No overrepresented sequences

### Adapter Content

### Kmer Content

| Sequence | Count | PValue | Obs/Exp Max | Max Obs/Exp Position |
| --- | --- | --- | --- | --- |
| CTCCTGT | 635 | 0.0 | 23.16701 | 5 |
| TCCTGTA | 640 | 0.0 | 22.055336 | 6 |
| TCTCCTG | 850 | 0.0 | 16.572428 | 4 |
| CCTGTAG | 315 | 7.1409595E-5 | 15.556642 | 7 |
| CGATCTC | 930 | 0.0 | 15.052488 | 1 |
| CTTCCGA | 390 | 4.6169193E-4 | 12.485007 | 1 |
| GATCTCC | 1450 | 1.8189894E-12 | 9.701652 | 2 |
| ATCTCCT | 1495 | 5.0931703E-11 | 9.002965 | 3 |
| TTCGTTT | 870 | 8.0736267E-4 | 5.358314 | 74-75 |

Produced by FastQC (version 0.11.5)
