## Supplementary material 4 for "Natural and human-mediated drivers of microevolution in Neotropical palms: a historical genomics approach": outF9_AS1.1_pf_trimmed_masked_sickled_fastqc.html

outF9\_AS1.1\_pf\_trimmed\_masked\_sickled.fastq FastQC Report 

FastQC Report

mar. 25 juin 2019  
outF9\_AS1.1\_pf\_trimmed\_masked\_sickled.fastq

### Summary

- Basic Statistics
- Per base sequence quality
- Per tile sequence quality
- Per sequence quality scores
- Per base sequence content
- Per sequence GC content
- Per base N content
- Sequence Length Distribution
- Sequence Duplication Levels
- Overrepresented sequences
- Adapter Content
- Kmer Content

### Basic Statistics

| Measure | Value |
| --- | --- |
| Filename | outF9\_AS1.1\_pf\_trimmed\_masked\_sickled.fastq |
| File type | Conventional base calls |
| Encoding | Sanger / Illumina 1.9 |
| Total Sequences | 397571 |
| Sequences flagged as poor quality | 0 |
| Sequence length | 100-131 |
| %GC | 43 |

### Per base sequence quality

### Per tile sequence quality

### Per sequence quality scores

### Per base sequence content

### Per sequence GC content

### Per base N content

### Sequence Length Distribution

### Sequence Duplication Levels

### Overrepresented sequences

No overrepresented sequences

### Adapter Content

### Kmer Content

| Sequence | Count | PValue | Obs/Exp Max | Max Obs/Exp Position |
| --- | --- | --- | --- | --- |
| GGTACGC | 35 | 0.0063497317 | 53.709778 | 5 |
| AGGCGGG | 110 | 5.548133E-4 | 28.63116 | 116-117 |
| TGTGCCT | 195 | 9.4104143E-7 | 25.673054 | 6 |
| CTGTGCC | 295 | 3.3463715E-5 | 16.992924 | 5 |
| TCTGTGC | 300 | 7.432367E-4 | 14.589908 | 4 |
| ATCTGTG | 350 | 0.0023937572 | 12.494012 | 3 |
| GATCTGT | 405 | 5.299232E-4 | 12.279445 | 2 |
| GTTGCGC | 270 | 0.006070434 | 9.163577 | 68-69 |
| CCGACCC | 285 | 0.00822151 | 8.831041 | 116-117 |
| GCTCAAA | 460 | 0.0071951095 | 6.8241997 | 40-41 |
| CCTTCAA | 615 | 0.0026017814 | 6.142758 | 56-57 |

Produced by FastQC (version 0.11.5)
