## Supplementary material 4 for "Natural and human-mediated drivers of microevolution in Neotropical palms: a historical genomics approach": outF9_AS1.2_pf_trimmed_masked_sickled_fastqc.html

outF9\_AS1.2\_pf\_trimmed\_masked\_sickled.fastq FastQC Report 

FastQC Report

mar. 25 juin 2019  
outF9\_AS1.2\_pf\_trimmed\_masked\_sickled.fastq

### Summary

- Basic Statistics
- Per base sequence quality
- Per tile sequence quality
- Per sequence quality scores
- Per base sequence content
- Per sequence GC content
- Per base N content
- Sequence Length Distribution
- Sequence Duplication Levels
- Overrepresented sequences
- Adapter Content
- Kmer Content

### Basic Statistics

| Measure | Value |
| --- | --- |
| Filename | outF9\_AS1.2\_pf\_trimmed\_masked\_sickled.fastq |
| File type | Conventional base calls |
| Encoding | Sanger / Illumina 1.9 |
| Total Sequences | 397571 |
| Sequences flagged as poor quality | 0 |
| Sequence length | 100-131 |
| %GC | 43 |

### Per base sequence quality

### Per tile sequence quality

### Per sequence quality scores

### Per base sequence content

### Per sequence GC content

### Per base N content

### Sequence Length Distribution

### Sequence Duplication Levels

### Overrepresented sequences

No overrepresented sequences

### Adapter Content

### Kmer Content

| Sequence | Count | PValue | Obs/Exp Max | Max Obs/Exp Position |
| --- | --- | --- | --- | --- |
| TGTGCCT | 290 | 1.7751154E-6 | 18.869028 | 6 |
| CTGTGCC | 285 | 3.2315264E-5 | 17.062117 | 5 |
| TCTGTGC | 285 | 3.299563E-5 | 17.020754 | 4 |
| ATCTGTG | 300 | 5.108911E-5 | 16.17407 | 3 |
| GATCTGT | 535 | 5.6752504E-4 | 10.219791 | 2 |
| CCCATTG | 340 | 0.004135266 | 8.220828 | 42-43 |

Produced by FastQC (version 0.11.5)
