## Supplementary material 4 for "Natural and human-mediated drivers of microevolution in Neotropical palms: a historical genomics approach": outF9_AS2.1_pf_trimmed_masked_sickled_fastqc.html

outF9\_AS2.1\_pf\_trimmed\_masked\_sickled.fastq FastQC Report 

FastQC Report

mar. 25 juin 2019  
outF9\_AS2.1\_pf\_trimmed\_masked\_sickled.fastq

### Summary

- Basic Statistics
- Per base sequence quality
- Per tile sequence quality
- Per sequence quality scores
- Per base sequence content
- Per sequence GC content
- Per base N content
- Sequence Length Distribution
- Sequence Duplication Levels
- Overrepresented sequences
- Adapter Content
- Kmer Content

### Basic Statistics

| Measure | Value |
| --- | --- |
| Filename | outF9\_AS2.1\_pf\_trimmed\_masked\_sickled.fastq |
| File type | Conventional base calls |
| Encoding | Sanger / Illumina 1.9 |
| Total Sequences | 660327 |
| Sequences flagged as poor quality | 0 |
| Sequence length | 100-131 |
| %GC | 44 |

### Per base sequence quality

### Per tile sequence quality

### Per sequence quality scores

### Per base sequence content

### Per sequence GC content

### Per base N content

### Sequence Length Distribution

### Sequence Duplication Levels

### Overrepresented sequences

No overrepresented sequences

### Adapter Content

### Kmer Content

| Sequence | Count | PValue | Obs/Exp Max | Max Obs/Exp Position |
| --- | --- | --- | --- | --- |
| GGGGGGG | 75 | 0.0034917283 | 20.882456 | 60-61 |
| TCTAACC | 455 | 0.0013678854 | 10.9683 | 4 |
| CAGCCTG | 920 | 8.263669E-9 | 10.860354 | 8 |
| AGCCTGT | 870 | 4.249887E-8 | 10.7632885 | 9 |
| CTCACAG | 1040 | 5.185939E-9 | 10.197092 | 4 |
| CGATCTA | 500 | 0.0032669746 | 9.877816 | 1 |
| ACAGCCT | 970 | 2.0910738E-7 | 9.65213 | 7 |
| ATCTCAC | 1125 | 1.9947765E-7 | 8.839664 | 2 |
| TTTCCGG | 560 | 0.008307287 | 8.819479 | 1 |
| CACAGCC | 1005 | 3.4765617E-6 | 8.699785 | 6 |
| TCACAGC | 1015 | 3.999352E-6 | 8.609943 | 5 |
| AGGCCTG | 355 | 0.0056514596 | 7.941216 | 60-61 |
| CGGTCAT | 835 | 0.0029614454 | 7.475699 | 5 |
| AATCTCA | 1520 | 3.523819E-7 | 7.310883 | 1 |
| GATCTAA | 920 | 0.00792602 | 6.7558565 | 2 |
| TGTATCA | 1035 | 2.7162969E-8 | 6.6059895 | 12-13 |
| TTAGCTT | 525 | 0.0040422124 | 6.490444 | 64-65 |
| CATTGGA | 1780 | 3.9706156E-6 | 6.3128505 | 9 |
| TCATTGG | 1740 | 1.8754561E-5 | 6.101147 | 8 |
| GCCTGTA | 790 | 2.229807E-4 | 5.893417 | 10-11 |

Produced by FastQC (version 0.11.5)
