## Supplementary material 4 for "Natural and human-mediated drivers of microevolution in Neotropical palms: a historical genomics approach": outF9_AS2.2_pf_trimmed_masked_sickled_fastqc.html

outF9\_AS2.2\_pf\_trimmed\_masked\_sickled.fastq FastQC Report 

FastQC Report

mar. 25 juin 2019  
outF9\_AS2.2\_pf\_trimmed\_masked\_sickled.fastq

### Summary

- Basic Statistics
- Per base sequence quality
- Per tile sequence quality
- Per sequence quality scores
- Per base sequence content
- Per sequence GC content
- Per base N content
- Sequence Length Distribution
- Sequence Duplication Levels
- Overrepresented sequences
- Adapter Content
- Kmer Content

### Basic Statistics

| Measure | Value |
| --- | --- |
| Filename | outF9\_AS2.2\_pf\_trimmed\_masked\_sickled.fastq |
| File type | Conventional base calls |
| Encoding | Sanger / Illumina 1.9 |
| Total Sequences | 660327 |
| Sequences flagged as poor quality | 0 |
| Sequence length | 100-131 |
| %GC | 44 |

### Per base sequence quality

### Per tile sequence quality

### Per sequence quality scores

### Per base sequence content

### Per sequence GC content

### Per base N content

### Sequence Length Distribution

### Sequence Duplication Levels

### Overrepresented sequences

No overrepresented sequences

### Adapter Content

### Kmer Content

| Sequence | Count | PValue | Obs/Exp Max | Max Obs/Exp Position |
| --- | --- | --- | --- | --- |
| CTAACCT | 305 | 0.0010017658 | 14.028044 | 5 |
| TCTAACC | 430 | 7.0369424E-5 | 12.798134 | 4 |
| CGATCTA | 465 | 0.0020508536 | 10.447969 | 1 |
| TAACCTT | 470 | 0.00213005 | 10.40045 | 6 |
| TTCCGGT | 590 | 1.16450086E-4 | 10.344803 | 2 |
| ATCTAAC | 550 | 7.0058217E-4 | 9.990628 | 3 |
| CCGGTCA | 635 | 2.3994996E-4 | 9.629392 | 4 |
| TTTCCGG | 695 | 5.9981634E-5 | 9.611756 | 1 |
| CGACCCG | 430 | 3.1428196E-4 | 8.254614 | 118-119 |
| CGGTCAT | 975 | 2.9274908E-5 | 8.149625 | 5 |
| GGTCATT | 1090 | 1.3861327E-5 | 7.8480463 | 6 |
| TGCGCAG | 875 | 8.6401997E-10 | 7.8432436 | 70-71 |
| CATTGGA | 1990 | 3.6379788E-11 | 7.678667 | 9 |
| TTGCGCA | 920 | 2.384695E-9 | 7.4596066 | 70-71 |
| GTCATTG | 1325 | 3.1466625E-6 | 7.3672423 | 7 |
| TCATTGG | 1750 | 1.0682925E-8 | 7.3288026 | 8 |
| TCCGACC | 530 | 3.67792E-4 | 7.299191 | 116-117 |
| GCGCAGC | 900 | 1.2330929E-8 | 7.2769856 | 72-73 |
| TCGTTTT | 740 | 1.120173E-5 | 6.766344 | 56-57 |
| ATCCGAC | 625 | 3.4392378E-4 | 6.679044 | 114-115 |

Produced by FastQC (version 0.11.5)
