## Supplementary material 4 for "Natural and human-mediated drivers of microevolution in Neotropical palms: a historical genomics approach": outF9_AS3.1_pf_trimmed_masked_sickled_fastqc.html

outF9\_AS3.1\_pf\_trimmed\_masked\_sickled.fastq FastQC Report 

FastQC Report

mar. 25 juin 2019  
outF9\_AS3.1\_pf\_trimmed\_masked\_sickled.fastq

### Summary

- Basic Statistics
- Per base sequence quality
- Per tile sequence quality
- Per sequence quality scores
- Per base sequence content
- Per sequence GC content
- Per base N content
- Sequence Length Distribution
- Sequence Duplication Levels
- Overrepresented sequences
- Adapter Content
- Kmer Content

### Basic Statistics

| Measure | Value |
| --- | --- |
| Filename | outF9\_AS3.1\_pf\_trimmed\_masked\_sickled.fastq |
| File type | Conventional base calls |
| Encoding | Sanger / Illumina 1.9 |
| Total Sequences | 712264 |
| Sequences flagged as poor quality | 0 |
| Sequence length | 100-131 |
| %GC | 44 |

### Per base sequence quality

### Per tile sequence quality

### Per sequence quality scores

### Per base sequence content

### Per sequence GC content

### Per base N content

### Sequence Length Distribution

### Sequence Duplication Levels

### Overrepresented sequences

No overrepresented sequences

### Adapter Content

### Kmer Content

| Sequence | Count | PValue | Obs/Exp Max | Max Obs/Exp Position |
| --- | --- | --- | --- | --- |
| CTAGGCG | 1060 | 0.0 | 115.93858 | 9 |
| TCTAGGC | 1185 | 0.0 | 103.052505 | 8 |
| CTTCCGA | 1315 | 0.0 | 94.58272 | 1 |
| ATCTAGG | 1325 | 0.0 | 91.24375 | 7 |
| CGATCTA | 1540 | 0.0 | 80.88782 | 5 |
| GATCTAG | 1700 | 0.0 | 72.02023 | 6 |
| TTCCGAT | 1785 | 0.0 | 69.74724 | 2 |
| TAGGCGA | 1105 | 0.0 | 54.493156 | 10-11 |
| CCGATCT | 2560 | 0.0 | 47.93265 | 4 |
| AGGCGAC | 540 | 0.0 | 40.810043 | 10-11 |
| AGGCGAG | 520 | 0.0 | 32.23242 | 10-11 |
| AGGCGAA | 465 | 0.0 | 32.03987 | 10-11 |
| GGCGACT | 310 | 0.0 | 29.10805 | 12-13 |
| GGCGAAT | 250 | 0.0 | 28.626259 | 12-13 |
| GGCGAGT | 265 | 0.0 | 24.657564 | 12-13 |
| GGCGACA | 325 | 0.0 | 23.934998 | 12-13 |
| GCGAATC | 180 | 1.2478267E-9 | 20.743666 | 12-13 |
| GCGACTT | 170 | 1.4591933E-8 | 20.133558 | 12-13 |
| GCGACTA | 110 | 7.237954E-5 | 19.800774 | 12-13 |
| GGCGATA | 145 | 3.075441E-5 | 17.167173 | 12-13 |

Produced by FastQC (version 0.11.5)
