## Supplementary material 4 for "Natural and human-mediated drivers of microevolution in Neotropical palms: a historical genomics approach": outF9_AS3.2_pf_trimmed_masked_sickled_fastqc.html

outF9\_AS3.2\_pf\_trimmed\_masked\_sickled.fastq FastQC Report 

FastQC Report

mar. 25 juin 2019  
outF9\_AS3.2\_pf\_trimmed\_masked\_sickled.fastq

### Summary

- Basic Statistics
- Per base sequence quality
- Per tile sequence quality
- Per sequence quality scores
- Per base sequence content
- Per sequence GC content
- Per base N content
- Sequence Length Distribution
- Sequence Duplication Levels
- Overrepresented sequences
- Adapter Content
- Kmer Content

### Basic Statistics

| Measure | Value |
| --- | --- |
| Filename | outF9\_AS3.2\_pf\_trimmed\_masked\_sickled.fastq |
| File type | Conventional base calls |
| Encoding | Sanger / Illumina 1.9 |
| Total Sequences | 712264 |
| Sequences flagged as poor quality | 0 |
| Sequence length | 100-131 |
| %GC | 44 |

### Per base sequence quality

### Per tile sequence quality

### Per sequence quality scores

### Per base sequence content

### Per sequence GC content

### Per base N content

### Sequence Length Distribution

### Sequence Duplication Levels

### Overrepresented sequences

No overrepresented sequences

### Adapter Content

### Kmer Content

| Sequence | Count | PValue | Obs/Exp Max | Max Obs/Exp Position |
| --- | --- | --- | --- | --- |
| CTAGGCG | 445 | 0.0 | 83.786476 | 9 |
| TCTAGGC | 580 | 0.0 | 64.356476 | 8 |
| CTTCCGA | 715 | 0.0 | 58.44936 | 1 |
| ATCTAGG | 750 | 0.0 | 50.569786 | 7 |
| CGATCTA | 850 | 0.0 | 47.470776 | 5 |
| TTCCGAT | 1030 | 0.0 | 39.65306 | 2 |
| GATCTAG | 1025 | 0.0 | 37.635628 | 6 |
| TAGGCGA | 510 | 0.0 | 36.551174 | 10-11 |
| AGGCGAG | 380 | 0.0 | 24.92986 | 10-11 |
| CCGATCT | 1875 | 0.0 | 21.824974 | 4 |
| AGGCGAC | 270 | 1.8189894E-12 | 18.109144 | 10-11 |
| GGCGAGT | 220 | 5.712487E-6 | 13.890018 | 12-13 |
| GGCGACT | 205 | 4.530405E-5 | 13.415725 | 12-13 |
| AGGCGAT | 340 | 4.7971003E-4 | 8.987994 | 10-11 |
| AGGCGAA | 315 | 0.0024013326 | 8.731194 | 10-11 |
| GGCGAGC | 365 | 9.7228814E-4 | 8.372066 | 12-13 |
| CCGACCC | 530 | 3.487129E-4 | 7.3332076 | 116-117 |
| CATTGGA | 2185 | 4.1836756E-11 | 7.2732077 | 9 |
| AGCCTGT | 1120 | 1.6378178E-4 | 7.094625 | 9 |
| TCATTGG | 2020 | 3.092282E-9 | 6.9673376 | 8 |

Produced by FastQC (version 0.11.5)
