## Supplementary material 4 for "Natural and human-mediated drivers of microevolution in Neotropical palms: a historical genomics approach": outF9_AS4.1_pf_trimmed_masked_sickled_fastqc.html

outF9\_AS4.1\_pf\_trimmed\_masked\_sickled.fastq FastQC Report 

FastQC Report

mar. 25 juin 2019  
outF9\_AS4.1\_pf\_trimmed\_masked\_sickled.fastq

### Summary

- Basic Statistics
- Per base sequence quality
- Per tile sequence quality
- Per sequence quality scores
- Per base sequence content
- Per sequence GC content
- Per base N content
- Sequence Length Distribution
- Sequence Duplication Levels
- Overrepresented sequences
- Adapter Content
- Kmer Content

### Basic Statistics

| Measure | Value |
| --- | --- |
| Filename | outF9\_AS4.1\_pf\_trimmed\_masked\_sickled.fastq |
| File type | Conventional base calls |
| Encoding | Sanger / Illumina 1.9 |
| Total Sequences | 671600 |
| Sequences flagged as poor quality | 0 |
| Sequence length | 100-131 |
| %GC | 43 |

### Per base sequence quality

### Per tile sequence quality

### Per sequence quality scores

### Per base sequence content

### Per sequence GC content

### Per base N content

### Sequence Length Distribution

### Sequence Duplication Levels

### Overrepresented sequences

No overrepresented sequences

### Adapter Content

### Kmer Content

| Sequence | Count | PValue | Obs/Exp Max | Max Obs/Exp Position |
| --- | --- | --- | --- | --- |
| TAGTGGC | 470 | 0.0 | 37.19014 | 6 |
| CTAGTGG | 575 | 0.0 | 31.519785 | 5 |
| TCTAGTG | 610 | 0.0 | 29.670977 | 4 |
| AGTGGCG | 265 | 2.5465852E-11 | 28.304655 | 7 |
| ATCTAGT | 680 | 0.0 | 27.527836 | 3 |
| GATCTAG | 815 | 0.0 | 25.126522 | 2 |
| CGATCTA | 860 | 0.0 | 23.637377 | 1 |
| GTGGCGT | 200 | 3.3726952E-5 | 21.861418 | 8 |
| TTCCGAT | 880 | 3.092282E-11 | 12.60009 | 1 |
| TATATAC | 375 | 0.004461442 | 11.498749 | 1 |
| CGACCAG | 200 | 0.0057742116 | 11.108999 | 116-117 |
| ATCGGAA | 1075 | 7.495153E-5 | 7.557671 | 5 |
| CTCACCC | 480 | 0.0015624007 | 7.104257 | 74-75 |
| ATACAGG | 720 | 8.269861E-6 | 6.9064937 | 92-93 |
| TAACTCA | 900 | 6.8076406E-7 | 6.5915475 | 108-109 |
| CCGATCT | 1835 | 6.5677086E-6 | 6.1206307 | 3 |
| ACAGGCT | 785 | 2.019686E-4 | 5.936346 | 94-95 |
| TGCGCAG | 850 | 9.8361874E-5 | 5.831671 | 70-71 |
| ATTAACT | 910 | 3.784158E-5 | 5.8296323 | 106-107 |
| AATGTGC | 975 | 1.1428019E-4 | 5.419802 | 76-77 |

Produced by FastQC (version 0.11.5)
