## Supplementary material 4 for "Natural and human-mediated drivers of microevolution in Neotropical palms: a historical genomics approach": outF9_AS4.2_pf_trimmed_masked_sickled_fastqc.html

outF9\_AS4.2\_pf\_trimmed\_masked\_sickled.fastq FastQC Report 

FastQC Report

mar. 25 juin 2019  
outF9\_AS4.2\_pf\_trimmed\_masked\_sickled.fastq

### Summary

- Basic Statistics
- Per base sequence quality
- Per tile sequence quality
- Per sequence quality scores
- Per base sequence content
- Per sequence GC content
- Per base N content
- Sequence Length Distribution
- Sequence Duplication Levels
- Overrepresented sequences
- Adapter Content
- Kmer Content

### Basic Statistics

| Measure | Value |
| --- | --- |
| Filename | outF9\_AS4.2\_pf\_trimmed\_masked\_sickled.fastq |
| File type | Conventional base calls |
| Encoding | Sanger / Illumina 1.9 |
| Total Sequences | 671600 |
| Sequences flagged as poor quality | 0 |
| Sequence length | 100-131 |
| %GC | 43 |

### Per base sequence quality

### Per tile sequence quality

### Per sequence quality scores

### Per base sequence content

### Per sequence GC content

### Per base N content

### Sequence Length Distribution

### Sequence Duplication Levels

### Overrepresented sequences

No overrepresented sequences

### Adapter Content

### Kmer Content

| Sequence | Count | PValue | Obs/Exp Max | Max Obs/Exp Position |
| --- | --- | --- | --- | --- |
| CTAGTGG | 485 | 0.0 | 40.296894 | 5 |
| TAGTGGC | 500 | 0.0 | 39.02742 | 6 |
| TCTAGTG | 560 | 0.0 | 31.641039 | 4 |
| CGATCTA | 595 | 0.0 | 31.349428 | 1 |
| ATCTAGT | 705 | 0.0 | 26.792204 | 3 |
| GATCTAG | 940 | 0.0 | 20.705233 | 2 |
| AGTGGCG | 245 | 8.737767E-6 | 19.86497 | 7 |
| AGTGGCC | 320 | 0.0014909974 | 13.307977 | 7 |
| CGTTTGA | 185 | 0.003776108 | 11.7598 | 36-37 |
| AGTGGCA | 685 | 4.3140117E-6 | 10.657483 | 7 |
| AGGATCG | 460 | 0.0018469027 | 10.580255 | 7 |
| CCGACCC | 415 | 2.62729E-4 | 8.392818 | 98-99 |
| TCCGGGC | 395 | 0.0018470389 | 7.8453946 | 64-65 |
| CAAAGGG | 440 | 0.004916192 | 7.098331 | 58-59 |
| TTGCACA | 1440 | 5.777459E-4 | 5.901213 | 9 |
| CTTGCAC | 1455 | 6.387068E-4 | 5.8551097 | 8 |
| GCGCAGC | 920 | 0.001325814 | 5.162578 | 98-99 |
| TTGCGCA | 925 | 0.0014344845 | 5.132065 | 96-97 |

Produced by FastQC (version 0.11.5)
