## Supplementary material 4 for "Natural and human-mediated drivers of microevolution in Neotropical palms: a historical genomics approach": outF9_AS5.1_pf_trimmed_masked_sickled_fastqc.html

outF9\_AS5.1\_pf\_trimmed\_masked\_sickled.fastq FastQC Report 

FastQC Report

mar. 25 juin 2019  
outF9\_AS5.1\_pf\_trimmed\_masked\_sickled.fastq

### Summary

- Basic Statistics
- Per base sequence quality
- Per tile sequence quality
- Per sequence quality scores
- Per base sequence content
- Per sequence GC content
- Per base N content
- Sequence Length Distribution
- Sequence Duplication Levels
- Overrepresented sequences
- Adapter Content
- Kmer Content

### Basic Statistics

| Measure | Value |
| --- | --- |
| Filename | outF9\_AS5.1\_pf\_trimmed\_masked\_sickled.fastq |
| File type | Conventional base calls |
| Encoding | Sanger / Illumina 1.9 |
| Total Sequences | 433570 |
| Sequences flagged as poor quality | 0 |
| Sequence length | 100-131 |
| %GC | 42 |

### Per base sequence quality

### Per tile sequence quality

### Per sequence quality scores

### Per base sequence content

### Per sequence GC content

### Per base N content

### Sequence Length Distribution

### Sequence Duplication Levels

### Overrepresented sequences

No overrepresented sequences

### Adapter Content

### Kmer Content

| Sequence | Count | PValue | Obs/Exp Max | Max Obs/Exp Position |
| --- | --- | --- | --- | --- |
| TTTCCGG | 315 | 6.736127E-5 | 15.660247 | 1 |
| TTCCGGT | 360 | 1.9942092E-4 | 13.785601 | 2 |
| CCGGTCA | 405 | 5.0999416E-4 | 12.335949 | 4 |
| CGGTCAT | 560 | 3.9558E-6 | 12.285063 | 5 |
| GGTCATT | 730 | 6.777349E-6 | 10.259627 | 6 |
| TCATTGG | 1090 | 1.0641088E-8 | 9.756659 | 8 |
| GTCATTG | 840 | 3.3795186E-6 | 9.664994 | 7 |
| TCCGGTC | 540 | 0.005622871 | 9.247456 | 3 |
| CATTGGA | 1320 | 2.3677057E-7 | 8.055652 | 9 |
| GTTGCGC | 355 | 0.0063507655 | 7.838015 | 68-69 |
| TCGTTTT | 495 | 2.180698E-4 | 7.63722 | 56-57 |
| CGTTTTG | 505 | 2.7998007E-4 | 7.473591 | 58-59 |
| GGCAATC | 470 | 0.0011720994 | 7.2987313 | 88-89 |
| TTCGTTT | 485 | 0.0014687091 | 7.145131 | 56-57 |
| TTCGCAG | 820 | 1.0128188E-6 | 6.861452 | 34-35 |
| AATCTCA | 1025 | 0.003287306 | 6.6174088 | 1 |
| CATTCGC | 860 | 2.2235272E-6 | 6.5415177 | 32-33 |
| GCGCAGC | 660 | 0.0010182709 | 6.106203 | 72-73 |
| CGCAGCT | 610 | 0.002822258 | 6.0985036 | 72-73 |
| CTTCGTT | 570 | 0.007885626 | 6.085985 | 54-55 |

Produced by FastQC (version 0.11.5)
