## Supplementary material 4 for "Natural and human-mediated drivers of microevolution in Neotropical palms: a historical genomics approach": outF9_AS5.2_pf_trimmed_masked_sickled_fastqc.html

outF9\_AS5.2\_pf\_trimmed\_masked\_sickled.fastq FastQC Report 

FastQC Report

mar. 25 juin 2019  
outF9\_AS5.2\_pf\_trimmed\_masked\_sickled.fastq

### Summary

- Basic Statistics
- Per base sequence quality
- Per tile sequence quality
- Per sequence quality scores
- Per base sequence content
- Per sequence GC content
- Per base N content
- Sequence Length Distribution
- Sequence Duplication Levels
- Overrepresented sequences
- Adapter Content
- Kmer Content

### Basic Statistics

| Measure | Value |
| --- | --- |
| Filename | outF9\_AS5.2\_pf\_trimmed\_masked\_sickled.fastq |
| File type | Conventional base calls |
| Encoding | Sanger / Illumina 1.9 |
| Total Sequences | 433570 |
| Sequences flagged as poor quality | 0 |
| Sequence length | 100-131 |
| %GC | 42 |

### Per base sequence quality

### Per tile sequence quality

### Per sequence quality scores

### Per base sequence content

### Per sequence GC content

### Per base N content

### Sequence Length Distribution

### Sequence Duplication Levels

### Overrepresented sequences

No overrepresented sequences

### Adapter Content

### Kmer Content

| Sequence | Count | PValue | Obs/Exp Max | Max Obs/Exp Position |
| --- | --- | --- | --- | --- |
| CGACCCG | 190 | 1.3260524E-5 | 15.280592 | 118-119 |
| CCGGTCA | 465 | 1.0170614E-5 | 13.1301365 | 4 |
| CCGACCC | 270 | 1.9124727E-6 | 13.112179 | 116-117 |
| CTCTAAA | 330 | 0.0018278376 | 12.951089 | 5 |
| CGGTCAT | 695 | 2.9829607E-8 | 12.298876 | 5 |
| TTCCGGT | 400 | 5.7470164E-4 | 12.162124 | 2 |
| TCCGACC | 345 | 1.8229548E-7 | 12.12747 | 116-117 |
| TTTCCGG | 435 | 0.0012167426 | 11.121476 | 1 |
| CCGGGCG | 235 | 0.0017910086 | 10.620112 | 82-83 |
| GGTCATT | 865 | 5.255788E-8 | 10.605687 | 6 |
| GTCATTG | 935 | 1.4479156E-8 | 10.47346 | 7 |
| TCCGGGC | 305 | 1.3067917E-4 | 10.228388 | 82-83 |
| CGGGCGG | 250 | 0.0029376657 | 10.005956 | 84-85 |
| CTCACAG | 745 | 1.1215292E-5 | 9.834385 | 4 |
| CGGCAAT | 260 | 0.0039050342 | 9.667563 | 88-89 |
| CAGCCTG | 700 | 5.747255E-5 | 9.647276 | 8 |
| GCGGCAA | 260 | 0.0039764126 | 9.646394 | 86-87 |
| TCACAGC | 830 | 3.8696526E-6 | 9.562852 | 5 |
| ACAGCCT | 710 | 6.9413756E-5 | 9.482354 | 7 |
| ATCCGAC | 440 | 4.5897887E-6 | 9.438713 | 114-115 |

Produced by FastQC (version 0.11.5)
