## Supplementary material 4 for "Natural and human-mediated drivers of microevolution in Neotropical palms: a historical genomics approach": outF9_AS6.1_pf_trimmed_masked_sickled_fastqc.html

outF9\_AS6.1\_pf\_trimmed\_masked\_sickled.fastq FastQC Report 

FastQC Report

mar. 25 juin 2019  
outF9\_AS6.1\_pf\_trimmed\_masked\_sickled.fastq

### Summary

- Basic Statistics
- Per base sequence quality
- Per tile sequence quality
- Per sequence quality scores
- Per base sequence content
- Per sequence GC content
- Per base N content
- Sequence Length Distribution
- Sequence Duplication Levels
- Overrepresented sequences
- Adapter Content
- Kmer Content

### Basic Statistics

| Measure | Value |
| --- | --- |
| Filename | outF9\_AS6.1\_pf\_trimmed\_masked\_sickled.fastq |
| File type | Conventional base calls |
| Encoding | Sanger / Illumina 1.9 |
| Total Sequences | 568922 |
| Sequences flagged as poor quality | 0 |
| Sequence length | 100-131 |
| %GC | 43 |

### Per base sequence quality

### Per tile sequence quality

### Per sequence quality scores

### Per base sequence content

### Per sequence GC content

### Per base N content

### Sequence Length Distribution

### Sequence Duplication Levels

### Overrepresented sequences

No overrepresented sequences

### Adapter Content

### Kmer Content

| Sequence | Count | PValue | Obs/Exp Max | Max Obs/Exp Position |
| --- | --- | --- | --- | --- |
| TCTCGAC | 385 | 0.0 | 27.5587 | 4 |
| CTCGACT | 375 | 3.6379788E-12 | 23.335169 | 5 |
| CGACTCG | 250 | 1.880807E-4 | 17.478693 | 7 |
| TCGACTC | 590 | 1.2914825E-10 | 15.879304 | 6 |
| TCGCCCG | 140 | 0.008097416 | 13.358001 | 20-21 |
| ATCTCGA | 730 | 3.1850504E-9 | 12.820895 | 3 |
| CGATCTC | 690 | 2.2429958E-8 | 12.555301 | 1 |
| AATTGCT | 550 | 0.0065509183 | 9.07816 | 4 |
| CCCGAGA | 285 | 0.0077176527 | 8.900082 | 122-123 |
| GTTTTTC | 945 | 0.0012329863 | 7.2629128 | 3 |
| GATCTCG | 1705 | 2.080511E-6 | 6.5664988 | 2 |
| TTCGAGA | 565 | 0.007975382 | 6.0797586 | 32-33 |

Produced by FastQC (version 0.11.5)
