## Supplementary material 4 for "Natural and human-mediated drivers of microevolution in Neotropical palms: a historical genomics approach": outF9_AS6.2_pf_trimmed_masked_sickled_fastqc.html

outF9\_AS6.2\_pf\_trimmed\_masked\_sickled.fastq FastQC Report 

FastQC Report

mar. 25 juin 2019  
outF9\_AS6.2\_pf\_trimmed\_masked\_sickled.fastq

### Summary

- Basic Statistics
- Per base sequence quality
- Per tile sequence quality
- Per sequence quality scores
- Per base sequence content
- Per sequence GC content
- Per base N content
- Sequence Length Distribution
- Sequence Duplication Levels
- Overrepresented sequences
- Adapter Content
- Kmer Content

### Basic Statistics

| Measure | Value |
| --- | --- |
| Filename | outF9\_AS6.2\_pf\_trimmed\_masked\_sickled.fastq |
| File type | Conventional base calls |
| Encoding | Sanger / Illumina 1.9 |
| Total Sequences | 568922 |
| Sequences flagged as poor quality | 0 |
| Sequence length | 100-131 |
| %GC | 44 |

### Per base sequence quality

### Per tile sequence quality

### Per sequence quality scores

### Per base sequence content

### Per sequence GC content

### Per base N content

### Sequence Length Distribution

### Sequence Duplication Levels

### Overrepresented sequences

No overrepresented sequences

### Adapter Content

### Kmer Content

| Sequence | Count | PValue | Obs/Exp Max | Max Obs/Exp Position |
| --- | --- | --- | --- | --- |
| CTCGACT | 345 | 0.0 | 40.721188 | 5 |
| TCTCGAC | 435 | 0.0 | 33.67203 | 4 |
| TCGACTC | 580 | 0.0 | 25.348495 | 6 |
| CGACTCG | 265 | 7.122053E-7 | 20.758307 | 7 |
| CGATCTC | 735 | 0.0 | 19.821234 | 1 |
| ATCTCGA | 715 | 0.0 | 19.637697 | 3 |
| CGACTCC | 335 | 0.0020313791 | 12.771693 | 7 |
| GCTCGGT | 190 | 0.0035066523 | 11.876521 | 118-119 |
| GATCTCG | 1420 | 0.0 | 11.178777 | 2 |
| GACTCCT | 405 | 0.008223841 | 10.590922 | 8 |
| GGAGTCG | 285 | 4.948733E-4 | 10.376476 | 124-125 |
| GCTCCAG | 465 | 0.0090045 | 6.6680408 | 62-63 |

Produced by FastQC (version 0.11.5)
