## Supplementary material 4 for "Natural and human-mediated drivers of microevolution in Neotropical palms: a historical genomics approach": outF9_AS07.1_pf_trimmed_masked_sickled_fastqc.html

outF9\_AS07.1\_pf\_trimmed\_masked\_sickled.fastq FastQC Report 

FastQC Report

mar. 25 juin 2019  
outF9\_AS07.1\_pf\_trimmed\_masked\_sickled.fastq

### Summary

- Basic Statistics
- Per base sequence quality
- Per tile sequence quality
- Per sequence quality scores
- Per base sequence content
- Per sequence GC content
- Per base N content
- Sequence Length Distribution
- Sequence Duplication Levels
- Overrepresented sequences
- Adapter Content
- Kmer Content

### Basic Statistics

| Measure | Value |
| --- | --- |
| Filename | outF9\_AS07.1\_pf\_trimmed\_masked\_sickled.fastq |
| File type | Conventional base calls |
| Encoding | Sanger / Illumina 1.9 |
| Total Sequences | 1526184 |
| Sequences flagged as poor quality | 0 |
| Sequence length | 100-131 |
| %GC | 45 |

### Per base sequence quality

### Per tile sequence quality

### Per sequence quality scores

### Per base sequence content

### Per sequence GC content

### Per base N content

### Sequence Length Distribution

### Sequence Duplication Levels

### Overrepresented sequences

No overrepresented sequences

### Adapter Content

### Kmer Content

| Sequence | Count | PValue | Obs/Exp Max | Max Obs/Exp Position |
| --- | --- | --- | --- | --- |
| CTAGAAG | 1520 | 0.0 | 11.152722 | 5 |
| TAGAAGG | 1345 | 0.0 | 10.800387 | 6 |
| TCTAGAA | 1465 | 0.0 | 10.740231 | 4 |
| CGATCTA | 1745 | 0.0 | 10.015799 | 1 |
| GATCTAG | 1860 | 0.0 | 9.418785 | 2 |
| ATCTAGA | 1680 | 0.0 | 9.353554 | 3 |
| CGTTGTA | 340 | 4.2084686E-4 | 9.107313 | 50-51 |
| ACGTCTG | 1770 | 0.0 | 8.998898 | 124-125 |
| GCACACG | 2415 | 0.0 | 8.98106 | 124-125 |
| CACACGT | 2200 | 0.0 | 8.934496 | 124-125 |
| CGTCTGA | 1800 | 0.0 | 8.848917 | 124-125 |
| GGCTACC | 560 | 0.009787859 | 8.645872 | 8 |
| CACGTCT | 1980 | 0.0 | 8.163109 | 122-123 |
| ACACGTC | 2050 | 0.0 | 7.720111 | 122-123 |
| AGCACAC | 3135 | 0.0 | 7.567025 | 124-125 |
| GAGCACA | 3565 | 0.0 | 6.9895906 | 122-123 |
| CGGAAGA | 4030 | 0.0 | 6.9797163 | 124-125 |
| TCGGAAG | 4420 | 0.0 | 6.8238974 | 124-125 |
| AGAGCAC | 3595 | 0.0 | 6.7439313 | 122-123 |
| ACGATCG | 465 | 0.008615264 | 6.699525 | 78-79 |

Produced by FastQC (version 0.11.5)
