## Supplementary material 4 for "Natural and human-mediated drivers of microevolution in Neotropical palms: a historical genomics approach": outF9_AS07.2_pf_trimmed_masked_sickled_fastqc.html

outF9\_AS07.2\_pf\_trimmed\_masked\_sickled.fastq FastQC Report 

FastQC Report

mar. 25 juin 2019  
outF9\_AS07.2\_pf\_trimmed\_masked\_sickled.fastq

### Summary

- Basic Statistics
- Per base sequence quality
- Per tile sequence quality
- Per sequence quality scores
- Per base sequence content
- Per sequence GC content
- Per base N content
- Sequence Length Distribution
- Sequence Duplication Levels
- Overrepresented sequences
- Adapter Content
- Kmer Content

### Basic Statistics

| Measure | Value |
| --- | --- |
| Filename | outF9\_AS07.2\_pf\_trimmed\_masked\_sickled.fastq |
| File type | Conventional base calls |
| Encoding | Sanger / Illumina 1.9 |
| Total Sequences | 1526184 |
| Sequences flagged as poor quality | 0 |
| Sequence length | 100-131 |
| %GC | 45 |

### Per base sequence quality

### Per tile sequence quality

### Per sequence quality scores

### Per base sequence content

### Per sequence GC content

### Per base N content

### Sequence Length Distribution

### Sequence Duplication Levels

### Overrepresented sequences

No overrepresented sequences

### Adapter Content

### Kmer Content

| Sequence | Count | PValue | Obs/Exp Max | Max Obs/Exp Position |
| --- | --- | --- | --- | --- |
| TAGAAGG | 1205 | 0.0 | 12.313258 | 6 |
| CGATCTA | 1485 | 0.0 | 11.437706 | 1 |
| CTAGAAG | 1510 | 0.0 | 11.327755 | 5 |
| TCTAGAA | 1385 | 0.0 | 11.101787 | 4 |
| GATCTAG | 1715 | 0.0 | 9.6185055 | 2 |
| ATCTAGA | 1765 | 0.0 | 9.036108 | 3 |
| AAGAGCG | 3105 | 0.0 | 8.777663 | 122-123 |
| CGTCGTG | 1515 | 0.0 | 8.691678 | 124-125 |
| AGCGTCG | 1845 | 0.0 | 8.650992 | 124-125 |
| GCGTCGT | 1450 | 0.0 | 8.324079 | 122-123 |
| TCGGAAG | 4185 | 0.0 | 8.19984 | 124-125 |
| CGGAAGA | 3915 | 0.0 | 8.051887 | 124-125 |
| CCTTCTA | 6820 | 0.0 | 7.9571376 | 124-125 |
| GTGTAGG | 1415 | 0.0 | 7.8959413 | 124-125 |
| GTCGTGT | 1315 | 5.456968E-12 | 7.8895073 | 124-125 |
| GAGCGTC | 2075 | 0.0 | 7.8843894 | 124-125 |
| CGTGTAG | 1230 | 9.094947E-11 | 7.785893 | 124-125 |
| CCGGTCA | 890 | 2.0526914E-4 | 7.678389 | 4 |
| TCGTGTA | 1245 | 2.0372681E-10 | 7.505582 | 122-123 |
| AGAGCGT | 2255 | 0.0 | 7.4244523 | 122-123 |

Produced by FastQC (version 0.11.5)
