## Supplementary material 4 for "Natural and human-mediated drivers of microevolution in Neotropical palms: a historical genomics approach": outF9_AS08.1_pf_trimmed_masked_sickled_fastqc.html

outF9\_AS08.1\_pf\_trimmed\_masked\_sickled.fastq FastQC Report 

FastQC Report

mar. 25 juin 2019  
outF9\_AS08.1\_pf\_trimmed\_masked\_sickled.fastq

### Summary

- Basic Statistics
- Per base sequence quality
- Per tile sequence quality
- Per sequence quality scores
- Per base sequence content
- Per sequence GC content
- Per base N content
- Sequence Length Distribution
- Sequence Duplication Levels
- Overrepresented sequences
- Adapter Content
- Kmer Content

### Basic Statistics

| Measure | Value |
| --- | --- |
| Filename | outF9\_AS08.1\_pf\_trimmed\_masked\_sickled.fastq |
| File type | Conventional base calls |
| Encoding | Sanger / Illumina 1.9 |
| Total Sequences | 1408541 |
| Sequences flagged as poor quality | 0 |
| Sequence length | 100-131 |
| %GC | 44 |

### Per base sequence quality

### Per tile sequence quality

### Per sequence quality scores

### Per base sequence content

### Per sequence GC content

### Per base N content

### Sequence Length Distribution

### Sequence Duplication Levels

### Overrepresented sequences

No overrepresented sequences

### Adapter Content

### Kmer Content

| Sequence | Count | PValue | Obs/Exp Max | Max Obs/Exp Position |
| --- | --- | --- | --- | --- |
| CTAGGCG | 335 | 0.0 | 49.484344 | 5 |
| TAGGCGA | 505 | 0.0 | 32.839466 | 6 |
| TCTAGGC | 680 | 0.0 | 24.350698 | 4 |
| ATCTAGG | 1040 | 0.0 | 16.481205 | 3 |
| AGGCGAC | 405 | 3.3311517E-6 | 14.632311 | 7 |
| CGATCTA | 1380 | 0.0 | 12.7867985 | 1 |
| CGCCTAG | 1735 | 0.0 | 10.29002 | 124-125 |
| TCGCCTA | 1950 | 0.0 | 9.155479 | 124-125 |
| GATCTAG | 1900 | 0.0 | 9.0127325 | 2 |
| ACGTCTG | 575 | 4.9097453E-6 | 8.482957 | 120-121 |
| AGGCGAG | 630 | 0.0031758244 | 8.4658375 | 7 |
| GCCTAGA | 1920 | 0.0 | 7.8107686 | 124-125 |
| CCTAGAT | 2335 | 0.0 | 7.5623026 | 122-123 |
| TAGATCG | 2420 | 0.0 | 7.3773494 | 124-125 |
| CACGTCT | 620 | 1.14592935E-4 | 7.305311 | 120-121 |
| CTCGCCT | 1210 | 1.200533E-10 | 7.2966847 | 122-123 |
| GTCGCCT | 735 | 1.3753175E-5 | 7.207338 | 122-123 |
| ATTCGCC | 540 | 0.0013690963 | 7.1939907 | 122-123 |
| TTCGCCT | 865 | 2.5442532E-6 | 6.9407077 | 122-123 |
| CACACGT | 735 | 9.23827E-5 | 6.8012137 | 124-125 |

Produced by FastQC (version 0.11.5)
