## Supplementary material 4 for "Natural and human-mediated drivers of microevolution in Neotropical palms: a historical genomics approach": outF9_AS08.2_pf_trimmed_masked_sickled_fastqc.html

outF9\_AS08.2\_pf\_trimmed\_masked\_sickled.fastq FastQC Report 

FastQC Report

mar. 25 juin 2019  
outF9\_AS08.2\_pf\_trimmed\_masked\_sickled.fastq

### Summary

- Basic Statistics
- Per base sequence quality
- Per tile sequence quality
- Per sequence quality scores
- Per base sequence content
- Per sequence GC content
- Per base N content
- Sequence Length Distribution
- Sequence Duplication Levels
- Overrepresented sequences
- Adapter Content
- Kmer Content

### Basic Statistics

| Measure | Value |
| --- | --- |
| Filename | outF9\_AS08.2\_pf\_trimmed\_masked\_sickled.fastq |
| File type | Conventional base calls |
| Encoding | Sanger / Illumina 1.9 |
| Total Sequences | 1408541 |
| Sequences flagged as poor quality | 0 |
| Sequence length | 100-131 |
| %GC | 44 |

### Per base sequence quality

### Per tile sequence quality

### Per sequence quality scores

### Per base sequence content

### Per sequence GC content

### Per base N content

### Sequence Length Distribution

### Sequence Duplication Levels

### Overrepresented sequences

No overrepresented sequences

### Adapter Content

### Kmer Content

| Sequence | Count | PValue | Obs/Exp Max | Max Obs/Exp Position |
| --- | --- | --- | --- | --- |
| CTAGGCG | 265 | 0.0 | 48.764603 | 5 |
| TAGGCGA | 355 | 0.0 | 38.024426 | 6 |
| TCTAGGC | 480 | 0.0 | 28.050505 | 4 |
| CACGCTA | 125 | 0.004324441 | 14.68511 | 78-79 |
| ATCTAGG | 960 | 0.0 | 14.032094 | 3 |
| ATACGCA | 160 | 0.008985374 | 13.150092 | 106-107 |
| CGATCTA | 1295 | 0.0 | 12.463122 | 1 |
| CGCCTAG | 1700 | 0.0 | 11.925166 | 124-125 |
| GTCGCCT | 810 | 0.0 | 11.170006 | 122-123 |
| TCGCCTA | 1950 | 0.0 | 10.396298 | 124-125 |
| AGGCGAG | 545 | 1.2187424E-4 | 10.300331 | 7 |
| GCCTATC | 470 | 0.004309332 | 9.553767 | 3 |
| GATCTAG | 1720 | 0.0 | 9.110511 | 2 |
| GCCTAGA | 1880 | 0.0 | 7.6382384 | 124-125 |
| GAGCGTC | 750 | 1.11595145E-4 | 7.320727 | 124-125 |
| GGCGTTC | 420 | 0.0044222395 | 7.176679 | 46-47 |
| TAGATCG | 2390 | 0.0 | 7.0686135 | 124-125 |
| CCTAGAT | 2095 | 0.0 | 7.066986 | 122-123 |
| CTCGCCT | 1270 | 2.7224814E-8 | 6.983734 | 124-125 |
| TCGGAAG | 1480 | 7.437848E-8 | 6.27817 | 124-125 |

Produced by FastQC (version 0.11.5)
