## Supplementary material 4 for "Natural and human-mediated drivers of microevolution in Neotropical palms: a historical genomics approach": outF9_AS09.1_pf_trimmed_masked_sickled_fastqc.html

outF9\_AS09.1\_pf\_trimmed\_masked\_sickled.fastq FastQC Report 

FastQC Report

mar. 25 juin 2019  
outF9\_AS09.1\_pf\_trimmed\_masked\_sickled.fastq

### Summary

- Basic Statistics
- Per base sequence quality
- Per tile sequence quality
- Per sequence quality scores
- Per base sequence content
- Per sequence GC content
- Per base N content
- Sequence Length Distribution
- Sequence Duplication Levels
- Overrepresented sequences
- Adapter Content
- Kmer Content

### Basic Statistics

| Measure | Value |
| --- | --- |
| Filename | outF9\_AS09.1\_pf\_trimmed\_masked\_sickled.fastq |
| File type | Conventional base calls |
| Encoding | Sanger / Illumina 1.9 |
| Total Sequences | 1254650 |
| Sequences flagged as poor quality | 0 |
| Sequence length | 100-131 |
| %GC | 44 |

### Per base sequence quality

### Per tile sequence quality

### Per sequence quality scores

### Per base sequence content

### Per sequence GC content

### Per base N content

### Sequence Length Distribution

### Sequence Duplication Levels

### Overrepresented sequences

No overrepresented sequences

### Adapter Content

### Kmer Content

| Sequence | Count | PValue | Obs/Exp Max | Max Obs/Exp Position |
| --- | --- | --- | --- | --- |
| CTAGGTT | 975 | 0.0 | 29.066399 | 5 |
| ATCTAGG | 1110 | 0.0 | 26.55379 | 3 |
| TCTAGGT | 1160 | 0.0 | 24.396954 | 4 |
| TAGGTTC | 1260 | 0.0 | 22.970404 | 6 |
| CGATCTA | 1460 | 0.0 | 20.106289 | 1 |
| GATCTAG | 1965 | 0.0 | 15.60256 | 2 |
| AGGTTCG | 675 | 1.09139364E-10 | 14.295613 | 7 |
| GGTTCGT | 445 | 7.382334E-6 | 13.549405 | 8 |
| CACACGT | 1895 | 0.0 | 9.95382 | 124-125 |
| GCACACG | 2050 | 0.0 | 9.870394 | 124-125 |
| AGGTTCC | 815 | 3.6086367E-6 | 9.61994 | 7 |
| ACACGTC | 1710 | 0.0 | 9.42623 | 124-125 |
| ACGTCTG | 1495 | 0.0 | 8.717235 | 124-125 |
| CTTCCGA | 895 | 1.2877794E-5 | 8.701798 | 1 |
| GTCACAC | 925 | 4.3655746E-11 | 8.5275135 | 124-125 |
| CACGTCT | 1620 | 0.0 | 7.8329086 | 124-125 |
| CGTCTGA | 1540 | 0.0 | 7.794418 | 124-125 |
| AGTCACA | 1175 | 1.8189894E-11 | 7.5435963 | 122-123 |
| TCGGAAG | 3550 | 0.0 | 7.490473 | 122-123 |
| AGCACAC | 2630 | 0.0 | 7.258974 | 122-123 |

Produced by FastQC (version 0.11.5)
