## Supplementary material 4 for "Natural and human-mediated drivers of microevolution in Neotropical palms: a historical genomics approach": outF9_AS09.2_pf_trimmed_masked_sickled_fastqc.html

outF9\_AS09.2\_pf\_trimmed\_masked\_sickled.fastq FastQC Report 

FastQC Report

mar. 25 juin 2019  
outF9\_AS09.2\_pf\_trimmed\_masked\_sickled.fastq

### Summary

- Basic Statistics
- Per base sequence quality
- Per tile sequence quality
- Per sequence quality scores
- Per base sequence content
- Per sequence GC content
- Per base N content
- Sequence Length Distribution
- Sequence Duplication Levels
- Overrepresented sequences
- Adapter Content
- Kmer Content

### Basic Statistics

| Measure | Value |
| --- | --- |
| Filename | outF9\_AS09.2\_pf\_trimmed\_masked\_sickled.fastq |
| File type | Conventional base calls |
| Encoding | Sanger / Illumina 1.9 |
| Total Sequences | 1254650 |
| Sequences flagged as poor quality | 0 |
| Sequence length | 100-131 |
| %GC | 44 |

### Per base sequence quality

### Per tile sequence quality

### Per sequence quality scores

### Per base sequence content

### Per sequence GC content

### Per base N content

### Sequence Length Distribution

### Sequence Duplication Levels

### Overrepresented sequences

No overrepresented sequences

### Adapter Content

### Kmer Content

| Sequence | Count | PValue | Obs/Exp Max | Max Obs/Exp Position |
| --- | --- | --- | --- | --- |
| CTAGGTT | 835 | 0.0 | 28.01346 | 5 |
| TCTAGGT | 925 | 0.0 | 24.652475 | 4 |
| ATCTAGG | 1000 | 0.0 | 23.36874 | 3 |
| TAGGTTC | 995 | 0.0 | 21.790451 | 6 |
| CGATCTA | 1155 | 0.0 | 19.6341 | 1 |
| GATCTAG | 1570 | 0.0 | 15.602793 | 2 |
| AGGTTCG | 560 | 2.3101165E-10 | 15.277255 | 7 |
| CTTCCGA | 590 | 5.5479177E-10 | 14.413592 | 1 |
| AGCGTGA | 330 | 0.0030469333 | 12.101897 | 5 |
| CGTCGTG | 1315 | 0.0 | 9.577272 | 124-125 |
| GCGTCGT | 1260 | 0.0 | 9.350469 | 124-125 |
| CGTGTAG | 1000 | 5.456968E-12 | 9.344019 | 124-125 |
| GTTCTAC | 550 | 0.0013089385 | 9.3330145 | 8 |
| AGCGTCG | 1620 | 0.0 | 9.330739 | 122-123 |
| GAGCGTC | 1625 | 0.0 | 9.302029 | 122-123 |
| AGTGTAG | 825 | 2.0263542E-9 | 9.161089 | 122-123 |
| GTCGTGT | 1110 | 0.0 | 9.150039 | 124-125 |
| GAGTGTA | 755 | 4.335743E-8 | 8.956729 | 122-123 |
| AAGAGCG | 2535 | 0.0 | 8.814357 | 124-125 |
| TCGTGTA | 1035 | 1.200533E-10 | 8.635514 | 124-125 |

Produced by FastQC (version 0.11.5)
