## Supplementary material 4 for "Natural and human-mediated drivers of microevolution in Neotropical palms: a historical genomics approach": outF9_AS10.1_pf_trimmed_masked_sickled_fastqc.html

outF9\_AS10.1\_pf\_trimmed\_masked\_sickled.fastq FastQC Report 

FastQC Report

mar. 25 juin 2019  
outF9\_AS10.1\_pf\_trimmed\_masked\_sickled.fastq

### Summary

- Basic Statistics
- Per base sequence quality
- Per tile sequence quality
- Per sequence quality scores
- Per base sequence content
- Per sequence GC content
- Per base N content
- Sequence Length Distribution
- Sequence Duplication Levels
- Overrepresented sequences
- Adapter Content
- Kmer Content

### Basic Statistics

| Measure | Value |
| --- | --- |
| Filename | outF9\_AS10.1\_pf\_trimmed\_masked\_sickled.fastq |
| File type | Conventional base calls |
| Encoding | Sanger / Illumina 1.9 |
| Total Sequences | 1812771 |
| Sequences flagged as poor quality | 0 |
| Sequence length | 100-131 |
| %GC | 44 |

### Per base sequence quality

### Per tile sequence quality

### Per sequence quality scores

### Per base sequence content

### Per sequence GC content

### Per base N content

### Sequence Length Distribution

### Sequence Duplication Levels

### Overrepresented sequences

No overrepresented sequences

### Adapter Content

### Kmer Content

| Sequence | Count | PValue | Obs/Exp Max | Max Obs/Exp Position |
| --- | --- | --- | --- | --- |
| CTAGTGG | 1580 | 0.0 | 32.45105 | 5 |
| TAGTGGC | 1620 | 0.0 | 31.289568 | 6 |
| TCTAGTG | 1655 | 0.0 | 30.583387 | 4 |
| ATCTAGT | 1855 | 0.0 | 26.62036 | 3 |
| AGTGGCG | 740 | 0.0 | 23.886753 | 7 |
| CGATCTA | 2275 | 0.0 | 21.591635 | 1 |
| GATCTAG | 2845 | 0.0 | 17.552685 | 2 |
| AGTGGCC | 930 | 0.0 | 14.571775 | 7 |
| GTGGCGT | 500 | 1.4047328E-7 | 14.149829 | 8 |
| CACGTCT | 1950 | 0.0 | 10.899962 | 124-125 |
| ACGTCTG | 1745 | 0.0 | 9.870383 | 124-125 |
| ACACGTC | 1985 | 0.0 | 9.600071 | 124-125 |
| CGTCTGA | 1730 | 0.0 | 9.532307 | 124-125 |
| CACACGT | 2165 | 0.0 | 9.482518 | 122-123 |
| CGGAAGA | 4085 | 0.0 | 9.434056 | 122-123 |
| GCACACG | 2310 | 0.0 | 8.731378 | 122-123 |
| ACTCCAG | 1705 | 0.0 | 8.597401 | 124-125 |
| GTGGCCT | 970 | 4.70588E-6 | 8.509348 | 8 |
| TCGGAAG | 4535 | 0.0 | 8.418511 | 122-123 |
| AGTGGCA | 1690 | 1.6189006E-10 | 8.018787 | 7 |

Produced by FastQC (version 0.11.5)
