## Supplementary material 4 for "Natural and human-mediated drivers of microevolution in Neotropical palms: a historical genomics approach": outF9_AS10.2_pf_trimmed_masked_sickled_fastqc.html

outF9\_AS10.2\_pf\_trimmed\_masked\_sickled.fastq FastQC Report 

FastQC Report

mar. 25 juin 2019  
outF9\_AS10.2\_pf\_trimmed\_masked\_sickled.fastq

### Summary

- Basic Statistics
- Per base sequence quality
- Per tile sequence quality
- Per sequence quality scores
- Per base sequence content
- Per sequence GC content
- Per base N content
- Sequence Length Distribution
- Sequence Duplication Levels
- Overrepresented sequences
- Adapter Content
- Kmer Content

### Basic Statistics

| Measure | Value |
| --- | --- |
| Filename | outF9\_AS10.2\_pf\_trimmed\_masked\_sickled.fastq |
| File type | Conventional base calls |
| Encoding | Sanger / Illumina 1.9 |
| Total Sequences | 1812771 |
| Sequences flagged as poor quality | 0 |
| Sequence length | 100-131 |
| %GC | 44 |

### Per base sequence quality

### Per tile sequence quality

### Per sequence quality scores

### Per base sequence content

### Per sequence GC content

### Per base N content

### Sequence Length Distribution

### Sequence Duplication Levels

### Overrepresented sequences

No overrepresented sequences

### Adapter Content

### Kmer Content

| Sequence | Count | PValue | Obs/Exp Max | Max Obs/Exp Position |
| --- | --- | --- | --- | --- |
| TAGTGGC | 1200 | 0.0 | 40.06678 | 6 |
| CTAGTGG | 1345 | 0.0 | 35.28864 | 5 |
| TCTAGTG | 1575 | 0.0 | 30.771235 | 4 |
| ATCTAGT | 1820 | 0.0 | 26.001358 | 3 |
| AGTGGCG | 715 | 0.0 | 23.722427 | 7 |
| CGATCTA | 2045 | 0.0 | 22.686348 | 1 |
| GATCTAG | 2595 | 0.0 | 18.417774 | 2 |
| GTGGCGT | 415 | 6.957571E-6 | 13.6280985 | 8 |
| AGTGGCC | 965 | 5.456968E-12 | 11.717814 | 7 |
| CGTGTAG | 1165 | 0.0 | 10.859402 | 124-125 |
| AAGAGCG | 3160 | 0.0 | 10.275766 | 124-125 |
| CGGAAGA | 4010 | 0.0 | 10.262005 | 122-123 |
| GTTGCGC | 895 | 0.0 | 9.828465 | 50-51 |
| AGTGGCT | 1095 | 5.875336E-10 | 9.810326 | 7 |
| TCGGAAG | 4190 | 0.0 | 9.428309 | 122-123 |
| GTGTAGG | 1420 | 0.0 | 9.206274 | 124-125 |
| TGTAGGG | 1415 | 0.0 | 8.940779 | 124-125 |
| AGTGGCA | 1455 | 1.6370905E-11 | 8.93735 | 7 |
| ACAAACC | 2840 | 0.0 | 8.926522 | 3 |
| TGCGCAG | 1380 | 0.0 | 8.833841 | 52-53 |

Produced by FastQC (version 0.11.5)
