## Supplementary material 4 for "Natural and human-mediated drivers of microevolution in Neotropical palms: a historical genomics approach": outF9_AS11.2_pf_trimmed_masked_sickled_fastqc.html

outF9\_AS11.2\_pf\_trimmed\_masked\_sickled.fastq FastQC Report 

FastQC Report

mar. 25 juin 2019  
outF9\_AS11.2\_pf\_trimmed\_masked\_sickled.fastq

### Summary

- Basic Statistics
- Per base sequence quality
- Per tile sequence quality
- Per sequence quality scores
- Per base sequence content
- Per sequence GC content
- Per base N content
- Sequence Length Distribution
- Sequence Duplication Levels
- Overrepresented sequences
- Adapter Content
- Kmer Content

### Basic Statistics

| Measure | Value |
| --- | --- |
| Filename | outF9\_AS11.2\_pf\_trimmed\_masked\_sickled.fastq |
| File type | Conventional base calls |
| Encoding | Sanger / Illumina 1.9 |
| Total Sequences | 786627 |
| Sequences flagged as poor quality | 0 |
| Sequence length | 100-131 |
| %GC | 44 |

### Per base sequence quality

### Per tile sequence quality

### Per sequence quality scores

### Per base sequence content

### Per sequence GC content

### Per base N content

### Sequence Length Distribution

### Sequence Duplication Levels

### Overrepresented sequences

No overrepresented sequences

### Adapter Content

### Kmer Content

| Sequence | Count | PValue | Obs/Exp Max | Max Obs/Exp Position |
| --- | --- | --- | --- | --- |
| CTTCCGA | 455 | 0.0016348378 | 10.736886 | 1 |
| CTCTATG | 680 | 4.4652322E-4 | 9.05151 | 5 |
| CGATCTC | 870 | 6.433634E-4 | 7.7209935 | 1 |
| ATCTCTA | 915 | 0.008451138 | 6.7114706 | 3 |
| TAATCAT | 920 | 0.008781331 | 6.6848555 | 4 |
| TGCTTGA | 1245 | 6.03438E-4 | 6.376374 | 1 |
| TTGCACA | 1585 | 0.0016975695 | 5.4216785 | 7 |

Produced by FastQC (version 0.11.5)
