## Supplementary material 4 for "Natural and human-mediated drivers of microevolution in Neotropical palms: a historical genomics approach": outF9_AS12.1_pf_trimmed_masked_sickled_fastqc.html

outF9\_AS12.1\_pf\_trimmed\_masked\_sickled.fastq FastQC Report 

FastQC Report

mar. 25 juin 2019  
outF9\_AS12.1\_pf\_trimmed\_masked\_sickled.fastq

### Summary

- Basic Statistics
- Per base sequence quality
- Per tile sequence quality
- Per sequence quality scores
- Per base sequence content
- Per sequence GC content
- Per base N content
- Sequence Length Distribution
- Sequence Duplication Levels
- Overrepresented sequences
- Adapter Content
- Kmer Content

### Basic Statistics

| Measure | Value |
| --- | --- |
| Filename | outF9\_AS12.1\_pf\_trimmed\_masked\_sickled.fastq |
| File type | Conventional base calls |
| Encoding | Sanger / Illumina 1.9 |
| Total Sequences | 1518746 |
| Sequences flagged as poor quality | 0 |
| Sequence length | 100-131 |
| %GC | 45 |

### Per base sequence quality

### Per tile sequence quality

### Per sequence quality scores

### Per base sequence content

### Per sequence GC content

### Per base N content

### Sequence Length Distribution

### Sequence Duplication Levels

### Overrepresented sequences

No overrepresented sequences

### Adapter Content

### Kmer Content

| Sequence | Count | PValue | Obs/Exp Max | Max Obs/Exp Position |
| --- | --- | --- | --- | --- |
| CTATGTA | 1120 | 0.0 | 29.238977 | 5 |
| TATGTAG | 1245 | 0.0 | 26.810183 | 6 |
| TCTATGT | 1425 | 0.0 | 22.5492 | 4 |
| CGATCTA | 1625 | 0.0 | 20.075113 | 1 |
| ATCTATG | 2080 | 0.0 | 15.1523285 | 3 |
| GATCTAT | 2590 | 0.0 | 13.0968275 | 2 |
| ATGTAGC | 965 | 1.6370905E-11 | 11.961455 | 7 |
| GCGTGTC | 190 | 0.004261675 | 11.571805 | 86-87 |
| CGTCTGA | 915 | 0.0 | 9.515703 | 122-123 |
| ACACGTC | 995 | 0.0 | 9.087184 | 122-123 |
| ACGTCTG | 935 | 1.8189894E-12 | 8.889577 | 120-121 |
| CACGTCT | 1020 | 7.2759576E-12 | 8.257292 | 124-125 |
| ATGTAGT | 1115 | 2.312585E-6 | 8.172861 | 7 |
| TCGGAAG | 2355 | 0.0 | 8.154207 | 124-125 |
| AGCACAC | 1590 | 0.0 | 8.051639 | 124-125 |
| ATGTAGG | 990 | 3.836154E-5 | 7.9774833 | 7 |
| CACACGT | 1080 | 5.456968E-12 | 7.9523883 | 118-119 |
| GAGCACA | 2070 | 0.0 | 7.3238587 | 124-125 |
| GCACACG | 1115 | 5.802576E-10 | 7.156309 | 120-121 |
| AGAGCAC | 2075 | 0.0 | 7.1010604 | 122-123 |

Produced by FastQC (version 0.11.5)
