## Supplementary material 4 for "Natural and human-mediated drivers of microevolution in Neotropical palms: a historical genomics approach": outF9_AS12.2_pf_trimmed_masked_sickled_fastqc.html

outF9\_AS12.2\_pf\_trimmed\_masked\_sickled.fastq FastQC Report 

FastQC Report

mar. 25 juin 2019  
outF9\_AS12.2\_pf\_trimmed\_masked\_sickled.fastq

### Summary

- Basic Statistics
- Per base sequence quality
- Per tile sequence quality
- Per sequence quality scores
- Per base sequence content
- Per sequence GC content
- Per base N content
- Sequence Length Distribution
- Sequence Duplication Levels
- Overrepresented sequences
- Adapter Content
- Kmer Content

### Basic Statistics

| Measure | Value |
| --- | --- |
| Filename | outF9\_AS12.2\_pf\_trimmed\_masked\_sickled.fastq |
| File type | Conventional base calls |
| Encoding | Sanger / Illumina 1.9 |
| Total Sequences | 1518746 |
| Sequences flagged as poor quality | 0 |
| Sequence length | 100-131 |
| %GC | 45 |

### Per base sequence quality

### Per tile sequence quality

### Per sequence quality scores

### Per base sequence content

### Per sequence GC content

### Per base N content

### Sequence Length Distribution

### Sequence Duplication Levels

### Overrepresented sequences

No overrepresented sequences

### Adapter Content

### Kmer Content

| Sequence | Count | PValue | Obs/Exp Max | Max Obs/Exp Position |
| --- | --- | --- | --- | --- |
| TATGTAG | 1130 | 0.0 | 25.500927 | 6 |
| CTATGTA | 1075 | 0.0 | 25.172733 | 5 |
| TCTATGT | 1325 | 0.0 | 20.40895 | 4 |
| CGATCTA | 1455 | 0.0 | 18.820524 | 1 |
| ATCTATG | 1830 | 0.0 | 16.311546 | 3 |
| ATGTAGG | 985 | 0.0 | 12.281911 | 7 |
| AGAGCGT | 1040 | 0.0 | 11.1576 | 124-125 |
| GATCTAT | 2475 | 0.0 | 11.121723 | 2 |
| GAGCGTC | 1040 | 0.0 | 10.388111 | 124-125 |
| ATGTAGC | 805 | 5.1156167E-7 | 10.018785 | 7 |
| AAGAGCG | 1525 | 0.0 | 9.74705 | 122-123 |
| TGCCCCG | 310 | 0.0010508613 | 9.5608 | 98-99 |
| TCGGAAG | 2235 | 0.0 | 9.48865 | 124-125 |
| AGCGTCG | 940 | 2.1827873E-11 | 9.364853 | 124-125 |
| CGTGTAG | 585 | 1.2937307E-6 | 9.361202 | 122-123 |
| GTCGTGT | 580 | 2.0955085E-6 | 9.034691 | 118-119 |
| CGGAAGA | 2200 | 0.0 | 8.548331 | 124-125 |
| TACATAG | 3645 | 0.0 | 8.370599 | 122-123 |
| GTGTAGG | 850 | 4.345202E-8 | 8.283484 | 122-123 |
| ACATAGA | 3735 | 0.0 | 8.168898 | 122-123 |

Produced by FastQC (version 0.11.5)
