## Supplementary material 4 for "Natural and human-mediated drivers of microevolution in Neotropical palms: a historical genomics approach": outF9_AS13.1_pf_trimmed_masked_sickled_fastqc.html

outF9\_AS13.1\_pf\_trimmed\_masked\_sickled.fastq FastQC Report 

FastQC Report

mar. 25 juin 2019  
outF9\_AS13.1\_pf\_trimmed\_masked\_sickled.fastq

### Summary

- Basic Statistics
- Per base sequence quality
- Per tile sequence quality
- Per sequence quality scores
- Per base sequence content
- Per sequence GC content
- Per base N content
- Sequence Length Distribution
- Sequence Duplication Levels
- Overrepresented sequences
- Adapter Content
- Kmer Content

### Basic Statistics

| Measure | Value |
| --- | --- |
| Filename | outF9\_AS13.1\_pf\_trimmed\_masked\_sickled.fastq |
| File type | Conventional base calls |
| Encoding | Sanger / Illumina 1.9 |
| Total Sequences | 889823 |
| Sequences flagged as poor quality | 0 |
| Sequence length | 100-131 |
| %GC | 44 |

### Per base sequence quality

### Per tile sequence quality

### Per sequence quality scores

### Per base sequence content

### Per sequence GC content

### Per base N content

### Sequence Length Distribution

### Sequence Duplication Levels

### Overrepresented sequences

No overrepresented sequences

### Adapter Content

### Kmer Content

| Sequence | Count | PValue | Obs/Exp Max | Max Obs/Exp Position |
| --- | --- | --- | --- | --- |
| TCACCAC | 1030 | 0.0 | 19.07844 | 6 |
| CGATCTC | 1095 | 0.0 | 17.797546 | 1 |
| TCTCACC | 1170 | 0.0 | 16.766426 | 4 |
| CTCACCA | 1270 | 0.0 | 14.540354 | 5 |
| CACCACG | 490 | 1.526927E-6 | 13.3779545 | 7 |
| ACGTCTG | 480 | 1.2732926E-11 | 13.258101 | 124-125 |
| ATCTCAC | 1540 | 0.0 | 13.108223 | 3 |
| GATCTCA | 1565 | 0.0 | 12.505012 | 2 |
| CACACGT | 645 | 1.9645086E-10 | 10.414634 | 124-125 |
| ACACGTC | 625 | 1.4224497E-9 | 10.111317 | 122-123 |
| GCACACG | 655 | 4.08545E-9 | 9.512154 | 120-121 |
| CACGTCT | 510 | 8.274408E-6 | 9.012043 | 124-125 |
| TCGCCCA | 365 | 0.0039403974 | 8.266145 | 114-115 |
| CGTCTGA | 530 | 1.2667854E-4 | 8.004891 | 124-125 |
| CGGAAGA | 1220 | 1.6552804E-10 | 7.1944137 | 122-123 |
| AGCACAC | 1015 | 7.170238E-7 | 6.572079 | 122-123 |
| GGTGAGA | 2135 | 0.0 | 6.4133058 | 122-123 |
| GTGGTGA | 2180 | 0.0 | 6.1923532 | 120-121 |
| AGAGCAC | 1210 | 3.1597847E-7 | 6.1359806 | 124-125 |
| TCGGAAG | 1245 | 1.2399141E-7 | 6.1164713 | 120-121 |

Produced by FastQC (version 0.11.5)
