## Supplementary material 4 for "Natural and human-mediated drivers of microevolution in Neotropical palms: a historical genomics approach": outF9_AS13.2_pf_trimmed_masked_sickled_fastqc.html

outF9\_AS13.2\_pf\_trimmed\_masked\_sickled.fastq FastQC Report 

FastQC Report

mar. 25 juin 2019  
outF9\_AS13.2\_pf\_trimmed\_masked\_sickled.fastq

### Summary

- Basic Statistics
- Per base sequence quality
- Per tile sequence quality
- Per sequence quality scores
- Per base sequence content
- Per sequence GC content
- Per base N content
- Sequence Length Distribution
- Sequence Duplication Levels
- Overrepresented sequences
- Adapter Content
- Kmer Content

### Basic Statistics

| Measure | Value |
| --- | --- |
| Filename | outF9\_AS13.2\_pf\_trimmed\_masked\_sickled.fastq |
| File type | Conventional base calls |
| Encoding | Sanger / Illumina 1.9 |
| Total Sequences | 889823 |
| Sequences flagged as poor quality | 0 |
| Sequence length | 100-131 |
| %GC | 44 |

### Per base sequence quality

### Per tile sequence quality

### Per sequence quality scores

### Per base sequence content

### Per sequence GC content

### Per base N content

### Sequence Length Distribution

### Sequence Duplication Levels

### Overrepresented sequences

No overrepresented sequences

### Adapter Content

### Kmer Content

| Sequence | Count | PValue | Obs/Exp Max | Max Obs/Exp Position |
| --- | --- | --- | --- | --- |
| CGATCTC | 970 | 0.0 | 16.702074 | 1 |
| TCACCAC | 1020 | 0.0 | 15.999461 | 6 |
| TCTCACC | 935 | 0.0 | 15.034898 | 4 |
| GATCTCA | 1280 | 0.0 | 13.604593 | 2 |
| CTCACCA | 1140 | 0.0 | 13.321711 | 5 |
| CACCACG | 370 | 5.8764464E-4 | 12.1327305 | 7 |
| TCGTCAG | 345 | 0.004721467 | 11.412479 | 5 |
| ATCTCAC | 1440 | 0.0 | 10.935659 | 3 |
| AGCGTCG | 495 | 9.1143374E-7 | 10.710589 | 122-123 |
| CGTCGTG | 500 | 8.954177E-6 | 10.026664 | 124-125 |
| CGTGTAG | 375 | 6.7868613E-4 | 10.026664 | 124-125 |
| ACGTGGT | 340 | 0.0034041873 | 9.830063 | 124-125 |
| AGAGCGT | 580 | 9.494306E-7 | 9.575173 | 120-121 |
| GTCGTGT | 395 | 0.0013622472 | 9.292245 | 122-123 |
| GAGCGTC | 605 | 8.686095E-6 | 8.97704 | 124-125 |
| GCGTCGT | 445 | 5.2070984E-4 | 8.914286 | 120-121 |
| GTGTAGG | 465 | 0.0048034955 | 8.0860195 | 124-125 |
| AAGAGCG | 905 | 2.0674997E-7 | 7.8477573 | 124-125 |
| GGTGAGA | 1955 | 0.0 | 7.5098443 | 122-123 |
| GAGATCG | 1750 | 0.0 | 7.400633 | 124-125 |

Produced by FastQC (version 0.11.5)
