## Supplementary material 4 for "Natural and human-mediated drivers of microevolution in Neotropical palms: a historical genomics approach": outF9_AS14.1_pf_trimmed_masked_sickled_fastqc.html

outF9\_AS14.1\_pf\_trimmed\_masked\_sickled.fastq FastQC Report 

FastQC Report

mar. 25 juin 2019  
outF9\_AS14.1\_pf\_trimmed\_masked\_sickled.fastq

### Summary

- Basic Statistics
- Per base sequence quality
- Per tile sequence quality
- Per sequence quality scores
- Per base sequence content
- Per sequence GC content
- Per base N content
- Sequence Length Distribution
- Sequence Duplication Levels
- Overrepresented sequences
- Adapter Content
- Kmer Content

### Basic Statistics

| Measure | Value |
| --- | --- |
| Filename | outF9\_AS14.1\_pf\_trimmed\_masked\_sickled.fastq |
| File type | Conventional base calls |
| Encoding | Sanger / Illumina 1.9 |
| Total Sequences | 1024600 |
| Sequences flagged as poor quality | 0 |
| Sequence length | 100-131 |
| %GC | 43 |

### Per base sequence quality

### Per tile sequence quality

### Per sequence quality scores

### Per base sequence content

### Per sequence GC content

### Per base N content

### Sequence Length Distribution

### Sequence Duplication Levels

### Overrepresented sequences

No overrepresented sequences

### Adapter Content

### Kmer Content

| Sequence | Count | PValue | Obs/Exp Max | Max Obs/Exp Position |
| --- | --- | --- | --- | --- |
| CTCACTG | 830 | 1.8189894E-12 | 13.580887 | 5 |
| TCACTGC | 955 | 2.8721843E-9 | 10.572497 | 6 |
| TCTCACT | 1095 | 2.4920155E-10 | 10.281298 | 4 |
| ACGCAGT | 285 | 0.0028351753 | 10.050085 | 124-125 |
| CGATCTC | 1070 | 1.9754225E-9 | 9.920052 | 1 |
| CACACGT | 500 | 0.004517419 | 7.160685 | 124-125 |
| TCGGAAG | 985 | 4.029389E-7 | 6.794784 | 122-123 |
| GATCTCA | 1770 | 1.8111677E-7 | 6.6875205 | 2 |
| GCAGTGA | 1930 | 0.0 | 6.6783595 | 124-125 |
| CGGAAGA | 1005 | 5.7085526E-7 | 6.6595645 | 122-123 |
| ATCTCAC | 1670 | 2.2590575E-5 | 6.0289984 | 3 |
| AGCACAC | 885 | 1.8693876E-4 | 5.9704385 | 122-123 |
| CAGTGAG | 1745 | 1.4861143E-9 | 5.7449627 | 124-125 |
| GAGATCG | 1600 | 2.6087764E-7 | 5.3705134 | 124-125 |

Produced by FastQC (version 0.11.5)
