## Supplementary material 4 for "Natural and human-mediated drivers of microevolution in Neotropical palms: a historical genomics approach": outF9_AS14.2_pf_trimmed_masked_sickled_fastqc.html

outF9\_AS14.2\_pf\_trimmed\_masked\_sickled.fastq FastQC Report 

FastQC Report

mar. 25 juin 2019  
outF9\_AS14.2\_pf\_trimmed\_masked\_sickled.fastq

### Summary

- Basic Statistics
- Per base sequence quality
- Per tile sequence quality
- Per sequence quality scores
- Per base sequence content
- Per sequence GC content
- Per base N content
- Sequence Length Distribution
- Sequence Duplication Levels
- Overrepresented sequences
- Adapter Content
- Kmer Content

### Basic Statistics

| Measure | Value |
| --- | --- |
| Filename | outF9\_AS14.2\_pf\_trimmed\_masked\_sickled.fastq |
| File type | Conventional base calls |
| Encoding | Sanger / Illumina 1.9 |
| Total Sequences | 1024600 |
| Sequences flagged as poor quality | 0 |
| Sequence length | 100-131 |
| %GC | 43 |

### Per base sequence quality

### Per tile sequence quality

### Per sequence quality scores

### Per base sequence content

### Per sequence GC content

### Per base N content

### Sequence Length Distribution

### Sequence Duplication Levels

### Overrepresented sequences

No overrepresented sequences

### Adapter Content

### Kmer Content

| Sequence | Count | PValue | Obs/Exp Max | Max Obs/Exp Position |
| --- | --- | --- | --- | --- |
| GTATGCG | 150 | 2.485994E-4 | 22.47468 | 2 |
| CTCACTG | 685 | 1.8189894E-12 | 14.817891 | 5 |
| CGTCTAC | 115 | 0.004236533 | 14.730341 | 10-11 |
| CACTGCG | 235 | 0.004926878 | 14.398269 | 7 |
| TCGCAGT | 330 | 1.7702798E-5 | 12.4463005 | 122-123 |
| TCACTGC | 930 | 3.092282E-11 | 11.527761 | 6 |
| CGATCTC | 935 | 4.8748916E-10 | 10.754009 | 1 |
| TCTCACT | 1010 | 1.4370016E-10 | 10.59327 | 4 |
| CGTCGTG | 365 | 0.005629797 | 9.248812 | 124-125 |
| AGCGTCG | 425 | 0.002486303 | 8.697767 | 122-123 |
| GATCTCA | 1500 | 3.45608E-11 | 8.6152935 | 2 |
| AGAGCGT | 475 | 9.135502E-4 | 8.424853 | 120-121 |
| AAGAGCG | 755 | 1.6057256E-6 | 8.38365 | 124-125 |
| GAGCGTC | 465 | 0.004390731 | 8.167297 | 124-125 |
| CGCAGTG | 620 | 1.0025667E-4 | 8.167297 | 124-125 |
| GCAGTGA | 2000 | 0.0 | 7.8065753 | 124-125 |
| TCGGAAG | 1005 | 3.8809958E-8 | 7.765005 | 122-123 |
| ATCTCAC | 1655 | 2.3137545E-9 | 7.4700828 | 3 |
| CAGTGAG | 1665 | 7.2759576E-12 | 7.0963106 | 124-125 |
| GGCAGTG | 1100 | 9.146315E-7 | 6.905079 | 124-125 |

Produced by FastQC (version 0.11.5)
