## Supplementary material 4 for "Natural and human-mediated drivers of microevolution in Neotropical palms: a historical genomics approach": outF9_AS15.1_pf_trimmed_masked_sickled_fastqc.html

outF9\_AS15.1\_pf\_trimmed\_masked\_sickled.fastq FastQC Report 

FastQC Report

mar. 25 juin 2019  
outF9\_AS15.1\_pf\_trimmed\_masked\_sickled.fastq

### Summary

- Basic Statistics
- Per base sequence quality
- Per tile sequence quality
- Per sequence quality scores
- Per base sequence content
- Per sequence GC content
- Per base N content
- Sequence Length Distribution
- Sequence Duplication Levels
- Overrepresented sequences
- Adapter Content
- Kmer Content

### Basic Statistics

| Measure | Value |
| --- | --- |
| Filename | outF9\_AS15.1\_pf\_trimmed\_masked\_sickled.fastq |
| File type | Conventional base calls |
| Encoding | Sanger / Illumina 1.9 |
| Total Sequences | 534487 |
| Sequences flagged as poor quality | 0 |
| Sequence length | 100-131 |
| %GC | 43 |

### Per base sequence quality

### Per tile sequence quality

### Per sequence quality scores

### Per base sequence content

### Per sequence GC content

### Per base N content

### Sequence Length Distribution

### Sequence Duplication Levels

### Overrepresented sequences

No overrepresented sequences

### Adapter Content

### Kmer Content

| Sequence | Count | PValue | Obs/Exp Max | Max Obs/Exp Position |
| --- | --- | --- | --- | --- |
| CGGTCAT | 380 | 0.007074229 | 10.807825 | 5 |
| ACGTCTG | 310 | 5.3037395E-4 | 10.29839 | 120-121 |
| CGGAAGA | 735 | 1.4006218E-10 | 9.821992 | 122-123 |
| GCACACG | 425 | 7.004969E-5 | 9.47732 | 124-125 |
| CGTCTGA | 385 | 3.1407888E-4 | 9.375538 | 122-123 |
| CACGTCT | 385 | 3.740292E-4 | 9.213568 | 120-121 |
| CACACGT | 420 | 9.8065706E-5 | 9.189905 | 118-119 |
| ACACGTC | 380 | 0.0025508332 | 8.672427 | 124-125 |
| TTGCGCA | 330 | 0.003560712 | 8.3588705 | 70-71 |
| TCGGAGA | 1155 | 1.8189894E-12 | 8.242769 | 124-125 |
| CGTCGGA | 1165 | 3.6379788E-12 | 8.055719 | 122-123 |
| CGGAGAT | 1070 | 2.382876E-10 | 7.8709264 | 124-125 |
| GTATCCA | 355 | 0.0065460517 | 7.8118634 | 56-57 |
| GTCGGAG | 1165 | 2.4920155E-10 | 7.436049 | 122-123 |
| AGCACAC | 630 | 6.185099E-4 | 6.974651 | 124-125 |
| TCGGAAG | 820 | 7.993929E-6 | 6.9214115 | 120-121 |
| AAGAGCA | 1090 | 1.6616286E-7 | 6.7187004 | 124-125 |
| ATTCGCA | 500 | 0.0033275874 | 6.611901 | 32-33 |
| AGAGCAC | 720 | 4.6302547E-4 | 6.517301 | 122-123 |
| GAAGAGC | 1045 | 4.038442E-6 | 6.3072205 | 124-125 |

Produced by FastQC (version 0.11.5)
