## Supplementary material 4 for "Natural and human-mediated drivers of microevolution in Neotropical palms: a historical genomics approach": outF9_AS15.2_pf_trimmed_masked_sickled_fastqc.html

outF9\_AS15.2\_pf\_trimmed\_masked\_sickled.fastq FastQC Report 

FastQC Report

mar. 25 juin 2019  
outF9\_AS15.2\_pf\_trimmed\_masked\_sickled.fastq

### Summary

- Basic Statistics
- Per base sequence quality
- Per tile sequence quality
- Per sequence quality scores
- Per base sequence content
- Per sequence GC content
- Per base N content
- Sequence Length Distribution
- Sequence Duplication Levels
- Overrepresented sequences
- Adapter Content
- Kmer Content

### Basic Statistics

| Measure | Value |
| --- | --- |
| Filename | outF9\_AS15.2\_pf\_trimmed\_masked\_sickled.fastq |
| File type | Conventional base calls |
| Encoding | Sanger / Illumina 1.9 |
| Total Sequences | 534487 |
| Sequences flagged as poor quality | 0 |
| Sequence length | 100-131 |
| %GC | 43 |

### Per base sequence quality

### Per tile sequence quality

### Per sequence quality scores

### Per base sequence content

### Per sequence GC content

### Per base N content

### Sequence Length Distribution

### Sequence Duplication Levels

### Overrepresented sequences

No overrepresented sequences

### Adapter Content

### Kmer Content

| Sequence | Count | PValue | Obs/Exp Max | Max Obs/Exp Position |
| --- | --- | --- | --- | --- |
| TTTGCCC | 250 | 0.0075086365 | 13.509418 | 4 |
| ACCGTCG | 295 | 1.19996286E-4 | 12.088324 | 120-121 |
| CGGAAGA | 615 | 8.367351E-11 | 11.888753 | 122-123 |
| AAGAGCG | 525 | 1.278313E-7 | 11.078371 | 124-125 |
| TCGGAAG | 670 | 7.021845E-8 | 9.46217 | 120-121 |
| TCGGAGA | 1035 | 7.2759576E-12 | 9.231976 | 124-125 |
| AGTAGGA | 285 | 0.0070565357 | 8.997489 | 94-95 |
| CGGAGAT | 985 | 4.129106E-10 | 8.660081 | 122-123 |
| CGTCGGA | 1225 | 0.0 | 8.621367 | 122-123 |
| AGAGCGT | 430 | 0.003052438 | 8.501841 | 122-123 |
| CATTCGC | 510 | 9.53185E-6 | 8.072461 | 32-33 |
| GCGTCGG | 445 | 0.005203561 | 8.013607 | 120-121 |
| CGCAGCC | 610 | 1.4216712E-6 | 7.768996 | 36-37 |
| GCGCAGC | 435 | 7.5391884E-4 | 7.6077824 | 72-73 |
| GTCGGAG | 1130 | 6.040864E-9 | 7.5488315 | 122-123 |
| TTGCGCA | 400 | 0.0027531912 | 7.5324664 | 70-71 |
| TGCGCAG | 405 | 0.0031091399 | 7.4394727 | 70-71 |
| AGCGTCG | 510 | 0.0031350655 | 7.4327188 | 116-117 |
| TTCGCAG | 565 | 3.6468537E-5 | 7.297945 | 34-35 |
| TCATTGG | 800 | 0.0052153803 | 7.053575 | 8 |

Produced by FastQC (version 0.11.5)
