## Supplementary material 4 for "Natural and human-mediated drivers of microevolution in Neotropical palms: a historical genomics approach": outF10_AP1.1_pf_trimmed_masked_sickled_fastqc.html

outF10\_AP1.1\_pf\_trimmed\_masked\_sickled.fastq FastQC Report 

FastQC Report

mar. 25 juin 2019  
outF10\_AP1.1\_pf\_trimmed\_masked\_sickled.fastq

### Summary

- Basic Statistics
- Per base sequence quality
- Per tile sequence quality
- Per sequence quality scores
- Per base sequence content
- Per sequence GC content
- Per base N content
- Sequence Length Distribution
- Sequence Duplication Levels
- Overrepresented sequences
- Adapter Content
- Kmer Content

### Basic Statistics

| Measure | Value |
| --- | --- |
| Filename | outF10\_AP1.1\_pf\_trimmed\_masked\_sickled.fastq |
| File type | Conventional base calls |
| Encoding | Sanger / Illumina 1.9 |
| Total Sequences | 583584 |
| Sequences flagged as poor quality | 0 |
| Sequence length | 100-131 |
| %GC | 44 |

### Per base sequence quality

### Per tile sequence quality

### Per sequence quality scores

### Per base sequence content

### Per sequence GC content

### Per base N content

### Sequence Length Distribution

### Sequence Duplication Levels

### Overrepresented sequences

No overrepresented sequences

### Adapter Content

### Kmer Content

| Sequence | Count | PValue | Obs/Exp Max | Max Obs/Exp Position |
| --- | --- | --- | --- | --- |
| GCGTAAT | 30 | 0.0035296776 | 62.302216 | 4 |
| TAGTTAC | 80 | 0.005210554 | 19.486958 | 92-93 |
| CTCTATT | 595 | 8.043913E-6 | 11.529456 | 8 |
| TCTATTG | 625 | 1.3944418E-5 | 10.972093 | 9 |
| CGATCTC | 600 | 1.1205753E-4 | 10.383703 | 4 |
| TCTCTAT | 725 | 6.376862E-6 | 10.313946 | 7 |
| ATCTCTA | 740 | 8.2180595E-6 | 10.09671 | 6 |
| TTCCGAT | 725 | 8.114379E-5 | 9.349391 | 1 |
| GATCTCT | 945 | 0.0012504008 | 7.2534137 | 5 |
| CGGTCAT | 960 | 0.001477644 | 7.1400795 | 5 |
| GGTCATT | 1180 | 0.0018484648 | 6.3318353 | 6 |
| GTCATTG | 1235 | 0.003053976 | 6.054745 | 7 |
| TCATTGG | 1780 | 0.0010416822 | 5.255383 | 8 |
| CATTGGA | 1930 | 5.4899894E-4 | 5.168203 | 9 |
| CCGATCT | 1715 | 0.0038058674 | 5.082698 | 3 |

Produced by FastQC (version 0.11.5)
