## Supplementary material 4 for "Natural and human-mediated drivers of microevolution in Neotropical palms: a historical genomics approach": outF10_AP1.2_pf_trimmed_masked_sickled_fastqc.html

outF10\_AP1.2\_pf\_trimmed\_masked\_sickled.fastq FastQC Report 

FastQC Report

mar. 25 juin 2019  
outF10\_AP1.2\_pf\_trimmed\_masked\_sickled.fastq

### Summary

- Basic Statistics
- Per base sequence quality
- Per tile sequence quality
- Per sequence quality scores
- Per base sequence content
- Per sequence GC content
- Per base N content
- Sequence Length Distribution
- Sequence Duplication Levels
- Overrepresented sequences
- Adapter Content
- Kmer Content

### Basic Statistics

| Measure | Value |
| --- | --- |
| Filename | outF10\_AP1.2\_pf\_trimmed\_masked\_sickled.fastq |
| File type | Conventional base calls |
| Encoding | Sanger / Illumina 1.9 |
| Total Sequences | 583584 |
| Sequences flagged as poor quality | 0 |
| Sequence length | 100-131 |
| %GC | 44 |

### Per base sequence quality

### Per tile sequence quality

### Per sequence quality scores

### Per base sequence content

### Per sequence GC content

### Per base N content

### Sequence Length Distribution

### Sequence Duplication Levels

### Overrepresented sequences

No overrepresented sequences

### Adapter Content

### Kmer Content

| Sequence | Count | PValue | Obs/Exp Max | Max Obs/Exp Position |
| --- | --- | --- | --- | --- |
| TGCCGCT | 220 | 0.0010167792 | 11.365867 | 84-85 |
| CTTAAAA | 695 | 0.0057290294 | 7.927606 | 4 |
| TCCGGGC | 465 | 0.00104529 | 7.3783007 | 82-83 |
| CGGCAAT | 555 | 1.0180397E-4 | 7.374555 | 88-89 |
| CGGTCAT | 1065 | 6.9910317E-4 | 6.8991294 | 5 |
| GTTGCGC | 645 | 9.1492606E-5 | 6.8052664 | 68-69 |
| CCGACCC | 580 | 0.0010823311 | 6.6404157 | 116-117 |
| CATTGGA | 1940 | 5.32571E-7 | 6.298149 | 9 |
| GGTCATT | 1285 | 8.586686E-4 | 6.192219 | 6 |
| TCGTTTT | 770 | 1.442307E-4 | 6.084096 | 56-57 |
| TTCGTTT | 835 | 6.761033E-5 | 5.984516 | 56-57 |
| CGTTTTG | 855 | 9.340703E-5 | 5.852334 | 58-59 |
| CCTTCGT | 805 | 2.6147047E-4 | 5.824661 | 54-55 |
| TCATTGG | 1895 | 1.50767755E-5 | 5.8134003 | 8 |
| CTTCGTT | 865 | 1.1120527E-4 | 5.782014 | 54-55 |
| GTCATTG | 1415 | 0.0027498389 | 5.6182575 | 7 |
| TGCGCAG | 1005 | 2.6416496E-5 | 5.615433 | 70-71 |
| GCGCAGC | 1005 | 2.7692418E-5 | 5.598887 | 72-73 |
| GGCAATC | 855 | 5.364976E-4 | 5.5234523 | 88-89 |
| TTCGCAG | 1265 | 6.5779313E-6 | 5.2055397 | 34-35 |

Produced by FastQC (version 0.11.5)
