## Supplementary material 4 for "Natural and human-mediated drivers of microevolution in Neotropical palms: a historical genomics approach": outF10_AP03.1_pf_trimmed_masked_sickled_fastqc.html

outF10\_AP03.1\_pf\_trimmed\_masked\_sickled.fastq FastQC Report 

FastQC Report

mar. 25 juin 2019  
outF10\_AP03.1\_pf\_trimmed\_masked\_sickled.fastq

### Summary

- Basic Statistics
- Per base sequence quality
- Per tile sequence quality
- Per sequence quality scores
- Per base sequence content
- Per sequence GC content
- Per base N content
- Sequence Length Distribution
- Sequence Duplication Levels
- Overrepresented sequences
- Adapter Content
- Kmer Content

### Basic Statistics

| Measure | Value |
| --- | --- |
| Filename | outF10\_AP03.1\_pf\_trimmed\_masked\_sickled.fastq |
| File type | Conventional base calls |
| Encoding | Sanger / Illumina 1.9 |
| Total Sequences | 896255 |
| Sequences flagged as poor quality | 0 |
| Sequence length | 100-131 |
| %GC | 42 |

### Per base sequence quality

### Per tile sequence quality

### Per sequence quality scores

### Per base sequence content

### Per sequence GC content

### Per base N content

### Sequence Length Distribution

### Sequence Duplication Levels

### Overrepresented sequences

No overrepresented sequences

### Adapter Content

### Kmer Content

| Sequence | Count | PValue | Obs/Exp Max | Max Obs/Exp Position |
| --- | --- | --- | --- | --- |
| TAGCGCA | 90 | 1.1301745E-7 | 45.399284 | 6 |
| CTAGCGC | 120 | 2.4607289E-8 | 38.90691 | 5 |
| TCTAGCG | 175 | 6.887294E-7 | 26.615778 | 4 |
| AGCGCAG | 160 | 0.007629892 | 18.240784 | 7 |
| GCACACG | 755 | 1.8189894E-12 | 10.818197 | 124-125 |
| GACGAGC | 320 | 4.6732192E-4 | 10.441712 | 124-125 |
| ACACGTC | 585 | 2.3654138E-8 | 10.154143 | 124-125 |
| CACGTTG | 240 | 0.0031026665 | 9.94134 | 26-27 |
| CACACGT | 640 | 9.025825E-9 | 9.861617 | 124-125 |
| GCGCTAG | 1775 | 0.0 | 9.269064 | 122-123 |
| CTGCGCT | 830 | 1.34823495E-8 | 8.241684 | 120-121 |
| TCGGAAG | 1310 | 0.0 | 8.093723 | 122-123 |
| TGTGCGC | 495 | 5.6302355E-4 | 7.818992 | 116-117 |
| CGCTAGA | 1795 | 0.0 | 7.7292833 | 118-119 |
| GCTAGAT | 2315 | 0.0 | 7.6978493 | 124-125 |
| CACGTCT | 580 | 2.0435151E-4 | 7.681259 | 124-125 |
| GTGCGCT | 775 | 2.7677597E-6 | 7.432907 | 120-121 |
| AGCACAC | 1055 | 9.167707E-9 | 7.3900266 | 124-125 |
| GAGCACA | 1220 | 3.6925485E-10 | 7.3034925 | 124-125 |
| TGCGCTA | 1915 | 0.0 | 7.1731863 | 124-125 |

Produced by FastQC (version 0.11.5)
