## Supplementary material 4 for "Natural and human-mediated drivers of microevolution in Neotropical palms: a historical genomics approach": outF10_AP03.2_pf_trimmed_masked_sickled_fastqc.html

outF10\_AP03.2\_pf\_trimmed\_masked\_sickled.fastq FastQC Report 

FastQC Report

mar. 25 juin 2019  
outF10\_AP03.2\_pf\_trimmed\_masked\_sickled.fastq

### Summary

- Basic Statistics
- Per base sequence quality
- Per tile sequence quality
- Per sequence quality scores
- Per base sequence content
- Per sequence GC content
- Per base N content
- Sequence Length Distribution
- Sequence Duplication Levels
- Overrepresented sequences
- Adapter Content
- Kmer Content

### Basic Statistics

| Measure | Value |
| --- | --- |
| Filename | outF10\_AP03.2\_pf\_trimmed\_masked\_sickled.fastq |
| File type | Conventional base calls |
| Encoding | Sanger / Illumina 1.9 |
| Total Sequences | 896255 |
| Sequences flagged as poor quality | 0 |
| Sequence length | 100-131 |
| %GC | 42 |

### Per base sequence quality

### Per tile sequence quality

### Per sequence quality scores

### Per base sequence content

### Per sequence GC content

### Per base N content

### Sequence Length Distribution

### Sequence Duplication Levels

### Overrepresented sequences

No overrepresented sequences

### Adapter Content

### Kmer Content

| Sequence | Count | PValue | Obs/Exp Max | Max Obs/Exp Position |
| --- | --- | --- | --- | --- |
| CTAGCGC | 180 | 0.0 | 42.892067 | 5 |
| TAGCGCA | 185 | 0.0 | 38.812195 | 6 |
| TCTAGCG | 225 | 0.0 | 34.262478 | 4 |
| AGCGCAC | 160 | 4.3161397E-4 | 20.712276 | 7 |
| GTACGAT | 130 | 0.0051878723 | 14.287438 | 60-61 |
| CTGGGGG | 155 | 0.0013795368 | 13.447999 | 44-45 |
| ATCTAGC | 615 | 2.2835593E-8 | 12.541786 | 3 |
| AGTGCGC | 285 | 4.567406E-4 | 12.501886 | 124-125 |
| AGCGTCG | 540 | 7.219569E-8 | 11.546883 | 124-125 |
| GGCACTC | 220 | 0.00121974 | 11.121749 | 56-57 |
| CGATCTA | 665 | 9.555515E-7 | 10.67032 | 1 |
| TCTGCGC | 410 | 2.8038127E-4 | 9.482921 | 114-115 |
| TCGGAAG | 1410 | 0.0 | 9.461512 | 122-123 |
| CTGCGCT | 880 | 3.074092E-10 | 9.424681 | 120-121 |
| AGAGCGT | 685 | 2.9713374E-7 | 9.423638 | 122-123 |
| CGCTAGA | 1785 | 0.0 | 9.231961 | 124-125 |
| ATCTACC | 655 | 1.00815996E-4 | 9.1665745 | 1 |
| GCGCTAG | 1815 | 0.0 | 9.010002 | 122-123 |
| CGGAAGA | 1365 | 0.0 | 8.827614 | 122-123 |
| GAGCGTC | 635 | 1.1009051E-5 | 8.810236 | 122-123 |

Produced by FastQC (version 0.11.5)
