## Supplementary material 4 for "Natural and human-mediated drivers of microevolution in Neotropical palms: a historical genomics approach": outF10_AP05.1_pf_trimmed_masked_sickled_fastqc.html

outF10\_AP05.1\_pf\_trimmed\_masked\_sickled.fastq FastQC Report 

FastQC Report

mar. 25 juin 2019  
outF10\_AP05.1\_pf\_trimmed\_masked\_sickled.fastq

### Summary

- Basic Statistics
- Per base sequence quality
- Per tile sequence quality
- Per sequence quality scores
- Per base sequence content
- Per sequence GC content
- Per base N content
- Sequence Length Distribution
- Sequence Duplication Levels
- Overrepresented sequences
- Adapter Content
- Kmer Content

### Basic Statistics

| Measure | Value |
| --- | --- |
| Filename | outF10\_AP05.1\_pf\_trimmed\_masked\_sickled.fastq |
| File type | Conventional base calls |
| Encoding | Sanger / Illumina 1.9 |
| Total Sequences | 1425941 |
| Sequences flagged as poor quality | 0 |
| Sequence length | 100-131 |
| %GC | 44 |

### Per base sequence quality

### Per tile sequence quality

### Per sequence quality scores

### Per base sequence content

### Per sequence GC content

### Per base N content

### Sequence Length Distribution

### Sequence Duplication Levels

### Overrepresented sequences

No overrepresented sequences

### Adapter Content

### Kmer Content

| Sequence | Count | PValue | Obs/Exp Max | Max Obs/Exp Position |
| --- | --- | --- | --- | --- |
| TCTCAGC | 1515 | 0.0 | 13.375175 | 6 |
| CGATCTC | 1675 | 0.0 | 12.017007 | 1 |
| CTCTCAG | 1700 | 0.0 | 11.913202 | 5 |
| CTTCCGA | 855 | 4.8567017E-10 | 11.771044 | 1 |
| ACACGTC | 650 | 1.8189894E-12 | 11.523071 | 124-125 |
| GCACACG | 765 | 0.0 | 10.616394 | 122-123 |
| AAGCGAC | 240 | 0.0018644023 | 10.570657 | 92-93 |
| CACGTCT | 575 | 3.5488483E-9 | 10.439788 | 122-123 |
| CTCAGCG | 475 | 0.0028597862 | 10.039402 | 7 |
| ACGTCTG | 485 | 4.4452154E-6 | 9.464817 | 122-123 |
| CACACGT | 725 | 1.4115358E-9 | 9.347122 | 124-125 |
| CGGAAGA | 1570 | 0.0 | 9.31422 | 124-125 |
| GATCTCT | 2245 | 0.0 | 9.26777 | 2 |
| ATCTCTC | 2350 | 0.0 | 9.111552 | 3 |
| TCTCTCA | 2460 | 0.0 | 8.9520235 | 4 |
| TCGGAAG | 1745 | 0.0 | 8.175736 | 124-125 |
| CGTCTGA | 700 | 6.8867485E-7 | 8.152376 | 124-125 |
| TTCCGAT | 1840 | 3.6454549E-7 | 6.434938 | 1 |
| GCTGAGA | 2800 | 0.0 | 6.3690443 | 124-125 |
| AGCACAC | 1195 | 2.1241976E-7 | 6.267779 | 124-125 |

Produced by FastQC (version 0.11.5)
