## Supplementary material 4 for "Natural and human-mediated drivers of microevolution in Neotropical palms: a historical genomics approach": outF10_AP05.2_pf_trimmed_masked_sickled_fastqc.html

outF10\_AP05.2\_pf\_trimmed\_masked\_sickled.fastq FastQC Report 

FastQC Report

mar. 25 juin 2019  
outF10\_AP05.2\_pf\_trimmed\_masked\_sickled.fastq

### Summary

- Basic Statistics
- Per base sequence quality
- Per tile sequence quality
- Per sequence quality scores
- Per base sequence content
- Per sequence GC content
- Per base N content
- Sequence Length Distribution
- Sequence Duplication Levels
- Overrepresented sequences
- Adapter Content
- Kmer Content

### Basic Statistics

| Measure | Value |
| --- | --- |
| Filename | outF10\_AP05.2\_pf\_trimmed\_masked\_sickled.fastq |
| File type | Conventional base calls |
| Encoding | Sanger / Illumina 1.9 |
| Total Sequences | 1425941 |
| Sequences flagged as poor quality | 0 |
| Sequence length | 100-131 |
| %GC | 44 |

### Per base sequence quality

### Per tile sequence quality

### Per sequence quality scores

### Per base sequence content

### Per sequence GC content

### Per base N content

### Sequence Length Distribution

### Sequence Duplication Levels

### Overrepresented sequences

No overrepresented sequences

### Adapter Content

### Kmer Content

| Sequence | Count | PValue | Obs/Exp Max | Max Obs/Exp Position |
| --- | --- | --- | --- | --- |
| TCTCAGC | 1430 | 0.0 | 18.345818 | 6 |
| CTCTCAG | 1510 | 0.0 | 17.33503 | 5 |
| CTCAGCG | 520 | 1.3496901E-9 | 15.339764 | 7 |
| CGATCTC | 1665 | 0.0 | 14.190362 | 1 |
| GATCTCT | 2135 | 0.0 | 12.984524 | 2 |
| CCGCGCG | 225 | 0.0019127667 | 12.877063 | 124-125 |
| ATCTCTC | 2160 | 0.0 | 12.069027 | 3 |
| TCTCTCA | 2235 | 0.0 | 10.920746 | 4 |
| CTTCCGA | 690 | 1.0438016E-6 | 10.5987015 | 1 |
| AGCGTCG | 580 | 6.571281E-8 | 10.452176 | 122-123 |
| TCGGAAG | 1525 | 0.0 | 9.770886 | 124-125 |
| AAGAGCG | 1110 | 0.0 | 9.695086 | 124-125 |
| CGGAAGA | 1490 | 0.0 | 9.444825 | 124-125 |
| CGCTGAG | 700 | 1.020926E-7 | 9.237732 | 122-123 |
| AGAGCGT | 685 | 1.0966687E-7 | 9.194861 | 120-121 |
| GCGTCGT | 425 | 0.0023193127 | 8.765059 | 124-125 |
| GAGCGTC | 675 | 5.7619454E-6 | 8.382387 | 122-123 |
| TCTTCCG | 660 | 0.0077073975 | 7.6710887 | 1 |
| CTCAGCA | 1220 | 2.5565641E-6 | 7.472297 | 7 |
| CGTCGTG | 540 | 0.0037956657 | 7.2899075 | 120-121 |

Produced by FastQC (version 0.11.5)
