## Supplementary material 4 for "Natural and human-mediated drivers of microevolution in Neotropical palms: a historical genomics approach": outF10_AP07.1_pf_trimmed_masked_sickled_fastqc.html

outF10\_AP07.1\_pf\_trimmed\_masked\_sickled.fastq FastQC Report 

FastQC Report

mar. 25 juin 2019  
outF10\_AP07.1\_pf\_trimmed\_masked\_sickled.fastq

### Summary

- Basic Statistics
- Per base sequence quality
- Per tile sequence quality
- Per sequence quality scores
- Per base sequence content
- Per sequence GC content
- Per base N content
- Sequence Length Distribution
- Sequence Duplication Levels
- Overrepresented sequences
- Adapter Content
- Kmer Content

### Basic Statistics

| Measure | Value |
| --- | --- |
| Filename | outF10\_AP07.1\_pf\_trimmed\_masked\_sickled.fastq |
| File type | Conventional base calls |
| Encoding | Sanger / Illumina 1.9 |
| Total Sequences | 2017157 |
| Sequences flagged as poor quality | 0 |
| Sequence length | 100-131 |
| %GC | 44 |

### Per base sequence quality

### Per tile sequence quality

### Per sequence quality scores

### Per base sequence content

### Per sequence GC content

### Per base N content

### Sequence Length Distribution

### Sequence Duplication Levels

### Overrepresented sequences

No overrepresented sequences

### Adapter Content

### Kmer Content

| Sequence | Count | PValue | Obs/Exp Max | Max Obs/Exp Position |
| --- | --- | --- | --- | --- |
| CGATCTT | 1715 | 0.0 | 13.359828 | 1 |
| ATCTTAG | 1970 | 0.0 | 12.590791 | 3 |
| CTTAGAT | 2155 | 0.0 | 12.368628 | 5 |
| GATCTTA | 2200 | 0.0 | 10.993117 | 2 |
| CGTCTGA | 1955 | 0.0 | 10.878247 | 124-125 |
| TCTTAGA | 2185 | 0.0 | 10.804396 | 4 |
| ACGTCTG | 1960 | 0.0 | 10.678265 | 124-125 |
| ACACGTC | 2275 | 0.0 | 10.238416 | 124-125 |
| CACGTCT | 2100 | 0.0 | 10.21801 | 122-123 |
| GCACACG | 2725 | 0.0 | 9.662582 | 124-125 |
| CACACGT | 2480 | 0.0 | 9.255979 | 124-125 |
| AGAGCAC | 4285 | 0.0 | 9.217223 | 124-125 |
| AGCACAC | 3435 | 0.0 | 9.139476 | 124-125 |
| TCGGAAG | 5270 | 0.0 | 8.455288 | 124-125 |
| CGGAAGA | 5075 | 0.0 | 8.314556 | 124-125 |
| CTTCCGA | 990 | 4.2175583E-5 | 7.9175267 | 1 |
| GAGCACA | 4085 | 0.0 | 7.685214 | 124-125 |
| GTTGCGC | 770 | 4.9942173E-8 | 7.656513 | 50-51 |
| CGCGGAT | 450 | 7.49182E-4 | 7.6137233 | 60-61 |
| TTAGATC | 4015 | 0.0 | 7.5462675 | 6 |

Produced by FastQC (version 0.11.5)
