## Supplementary material 4 for "Natural and human-mediated drivers of microevolution in Neotropical palms: a historical genomics approach": outF10_AP07.2_pf_trimmed_masked_sickled_fastqc.html

outF10\_AP07.2\_pf\_trimmed\_masked\_sickled.fastq FastQC Report 

FastQC Report

mar. 25 juin 2019  
outF10\_AP07.2\_pf\_trimmed\_masked\_sickled.fastq

### Summary

- Basic Statistics
- Per base sequence quality
- Per tile sequence quality
- Per sequence quality scores
- Per base sequence content
- Per sequence GC content
- Per base N content
- Sequence Length Distribution
- Sequence Duplication Levels
- Overrepresented sequences
- Adapter Content
- Kmer Content

### Basic Statistics

| Measure | Value |
| --- | --- |
| Filename | outF10\_AP07.2\_pf\_trimmed\_masked\_sickled.fastq |
| File type | Conventional base calls |
| Encoding | Sanger / Illumina 1.9 |
| Total Sequences | 2017157 |
| Sequences flagged as poor quality | 0 |
| Sequence length | 100-131 |
| %GC | 44 |

### Per base sequence quality

### Per tile sequence quality

### Per sequence quality scores

### Per base sequence content

### Per sequence GC content

### Per base N content

### Sequence Length Distribution

### Sequence Duplication Levels

### Overrepresented sequences

No overrepresented sequences

### Adapter Content

### Kmer Content

| Sequence | Count | PValue | Obs/Exp Max | Max Obs/Exp Position |
| --- | --- | --- | --- | --- |
| CGATCTT | 1660 | 0.0 | 15.548059 | 1 |
| ATCTTAG | 1675 | 0.0 | 15.160094 | 3 |
| CTTAGAT | 1900 | 0.0 | 14.000244 | 5 |
| CCCGGGA | 640 | 0.0 | 13.569186 | 102-103 |
| GATCTTA | 1890 | 0.0 | 13.416415 | 2 |
| ACAAACC | 3115 | 0.0 | 12.968925 | 3 |
| GTTGCGC | 1110 | 0.0 | 12.420477 | 50-51 |
| TCTTAGA | 2170 | 0.0 | 12.242386 | 4 |
| CACAAAC | 3420 | 0.0 | 12.132546 | 2 |
| AACCATA | 3410 | 0.0 | 11.7092905 | 6 |
| TGCGCAG | 1775 | 0.0 | 11.58182 | 52-53 |
| ACCATAT | 3580 | 0.0 | 11.462848 | 7 |
| CCATATC | 3590 | 0.0 | 11.417585 | 8 |
| AGAGCGT | 2775 | 0.0 | 11.313648 | 124-125 |
| CGTTTTG | 1240 | 0.0 | 11.170427 | 40-41 |
| TCGTGTA | 1325 | 0.0 | 11.041724 | 122-123 |
| CGTGTAG | 1385 | 0.0 | 10.841368 | 122-123 |
| CAAACCA | 3835 | 0.0 | 10.69205 | 4 |
| CATATCA | 3725 | 0.0 | 10.68884 | 9 |
| TCGTTTT | 1185 | 0.0 | 10.653305 | 38-39 |

Produced by FastQC (version 0.11.5)
