## Supplementary material 4 for "Natural and human-mediated drivers of microevolution in Neotropical palms: a historical genomics approach": outF10_AP09.1_pf_trimmed_masked_sickled_fastqc.html

outF10\_AP09.1\_pf\_trimmed\_masked\_sickled.fastq FastQC Report 

FastQC Report

mar. 25 juin 2019  
outF10\_AP09.1\_pf\_trimmed\_masked\_sickled.fastq

### Summary

- Basic Statistics
- Per base sequence quality
- Per tile sequence quality
- Per sequence quality scores
- Per base sequence content
- Per sequence GC content
- Per base N content
- Sequence Length Distribution
- Sequence Duplication Levels
- Overrepresented sequences
- Adapter Content
- Kmer Content

### Basic Statistics

| Measure | Value |
| --- | --- |
| Filename | outF10\_AP09.1\_pf\_trimmed\_masked\_sickled.fastq |
| File type | Conventional base calls |
| Encoding | Sanger / Illumina 1.9 |
| Total Sequences | 1825810 |
| Sequences flagged as poor quality | 0 |
| Sequence length | 100-131 |
| %GC | 44 |

### Per base sequence quality

### Per tile sequence quality

### Per sequence quality scores

### Per base sequence content

### Per sequence GC content

### Per base N content

### Sequence Length Distribution

### Sequence Duplication Levels

### Overrepresented sequences

No overrepresented sequences

### Adapter Content

### Kmer Content

| Sequence | Count | PValue | Obs/Exp Max | Max Obs/Exp Position |
| --- | --- | --- | --- | --- |
| CTACTGA | 2000 | 0.0 | 42.698044 | 5 |
| TCTACTG | 2035 | 0.0 | 41.343117 | 4 |
| GATCTAC | 2235 | 0.0 | 37.35294 | 2 |
| CGATCTA | 2230 | 0.0 | 36.82038 | 1 |
| TACTGAT | 2320 | 0.0 | 36.548622 | 6 |
| ATCTACT | 2430 | 0.0 | 34.120766 | 3 |
| ACTGATA | 1490 | 0.0 | 13.016951 | 7 |
| ACTGATG | 2370 | 0.0 | 12.531218 | 7 |
| CACACGT | 1085 | 0.0 | 12.125795 | 124-125 |
| ACTGATT | 1645 | 0.0 | 11.790429 | 7 |
| ACTGATC | 1650 | 0.0 | 11.754701 | 7 |
| ACACGTC | 1030 | 0.0 | 10.480648 | 124-125 |
| CGGAAGA | 2420 | 0.0 | 9.618538 | 124-125 |
| GCACACG | 1180 | 0.0 | 8.819652 | 122-123 |
| TCGGAAG | 2510 | 0.0 | 8.467246 | 124-125 |
| ACGTCTG | 820 | 3.4635377E-8 | 7.8165503 | 124-125 |
| CTTCCGA | 1035 | 7.21326E-5 | 7.5832825 | 1 |
| AGCACAC | 1830 | 0.0 | 7.3380504 | 122-123 |
| CAGTAGA | 3780 | 0.0 | 7.3180842 | 124-125 |
| TCAGTAG | 3875 | 0.0 | 7.0516167 | 124-125 |

Produced by FastQC (version 0.11.5)
