## Supplementary material 4 for "Natural and human-mediated drivers of microevolution in Neotropical palms: a historical genomics approach": outF10_AP09.2_pf_trimmed_masked_sickled_fastqc.html

outF10\_AP09.2\_pf\_trimmed\_masked\_sickled.fastq FastQC Report 

FastQC Report

mar. 25 juin 2019  
outF10\_AP09.2\_pf\_trimmed\_masked\_sickled.fastq

### Summary

- Basic Statistics
- Per base sequence quality
- Per tile sequence quality
- Per sequence quality scores
- Per base sequence content
- Per sequence GC content
- Per base N content
- Sequence Length Distribution
- Sequence Duplication Levels
- Overrepresented sequences
- Adapter Content
- Kmer Content

### Basic Statistics

| Measure | Value |
| --- | --- |
| Filename | outF10\_AP09.2\_pf\_trimmed\_masked\_sickled.fastq |
| File type | Conventional base calls |
| Encoding | Sanger / Illumina 1.9 |
| Total Sequences | 1825810 |
| Sequences flagged as poor quality | 0 |
| Sequence length | 100-131 |
| %GC | 44 |

### Per base sequence quality

### Per tile sequence quality

### Per sequence quality scores

### Per base sequence content

### Per sequence GC content

### Per base N content

### Sequence Length Distribution

### Sequence Duplication Levels

### Overrepresented sequences

No overrepresented sequences

### Adapter Content

### Kmer Content

| Sequence | Count | PValue | Obs/Exp Max | Max Obs/Exp Position |
| --- | --- | --- | --- | --- |
| CTACTGA | 1865 | 0.0 | 44.246277 | 5 |
| TCTACTG | 1970 | 0.0 | 40.691566 | 4 |
| CGATCTA | 2220 | 0.0 | 35.542004 | 1 |
| ATCTACT | 2335 | 0.0 | 35.51902 | 3 |
| GATCTAC | 2235 | 0.0 | 35.49916 | 2 |
| TACTGAT | 2275 | 0.0 | 35.055225 | 6 |
| ACTGATA | 1380 | 0.0 | 17.4447 | 7 |
| ACTGATC | 1410 | 0.0 | 13.821434 | 7 |
| ACTGATG | 2100 | 0.0 | 11.463659 | 7 |
| CGTCGTG | 925 | 0.0 | 10.808961 | 124-125 |
| AGCGTCG | 985 | 0.0 | 10.345667 | 122-123 |
| GTCGTGT | 670 | 5.660695E-9 | 10.147518 | 124-125 |
| AGAGCGT | 1155 | 0.0 | 9.629769 | 120-121 |
| ACTGATT | 1740 | 0.0 | 9.55305 | 7 |
| CGGAAGA | 2220 | 0.0 | 9.547916 | 124-125 |
| GCGTCGT | 850 | 4.656613E-10 | 9.222157 | 122-123 |
| CCGCATA | 280 | 0.008332672 | 8.81818 | 80-81 |
| AAGAGCG | 1685 | 0.0 | 8.78188 | 124-125 |
| GAGCGTC | 1060 | 7.2759576E-12 | 8.683709 | 120-121 |
| CAGTAGA | 3710 | 0.0 | 8.623861 | 124-125 |

Produced by FastQC (version 0.11.5)
