## Supplementary material 4 for "Natural and human-mediated drivers of microevolution in Neotropical palms: a historical genomics approach": outF10_AP11.1_pf_trimmed_masked_sickled_fastqc.html

outF10\_AP11.1\_pf\_trimmed\_masked\_sickled.fastq FastQC Report 

FastQC Report

mar. 25 juin 2019  
outF10\_AP11.1\_pf\_trimmed\_masked\_sickled.fastq

### Summary

- Basic Statistics
- Per base sequence quality
- Per tile sequence quality
- Per sequence quality scores
- Per base sequence content
- Per sequence GC content
- Per base N content
- Sequence Length Distribution
- Sequence Duplication Levels
- Overrepresented sequences
- Adapter Content
- Kmer Content

### Basic Statistics

| Measure | Value |
| --- | --- |
| Filename | outF10\_AP11.1\_pf\_trimmed\_masked\_sickled.fastq |
| File type | Conventional base calls |
| Encoding | Sanger / Illumina 1.9 |
| Total Sequences | 1096205 |
| Sequences flagged as poor quality | 0 |
| Sequence length | 100-131 |
| %GC | 44 |

### Per base sequence quality

### Per tile sequence quality

### Per sequence quality scores

### Per base sequence content

### Per sequence GC content

### Per base N content

### Sequence Length Distribution

### Sequence Duplication Levels

### Overrepresented sequences

No overrepresented sequences

### Adapter Content

### Kmer Content

| Sequence | Count | PValue | Obs/Exp Max | Max Obs/Exp Position |
| --- | --- | --- | --- | --- |
| CTATACA | 675 | 0.0 | 29.550337 | 5 |
| TCTATAC | 700 | 0.0 | 28.48043 | 4 |
| TATACAC | 705 | 0.0 | 28.29025 | 6 |
| ATACACG | 200 | 4.3295444E-5 | 21.167143 | 7 |
| ATCTATA | 1415 | 0.0 | 15.357991 | 3 |
| CGATCTA | 1275 | 0.0 | 14.638416 | 1 |
| CACACGT | 460 | 2.7801434E-8 | 11.082319 | 124-125 |
| CTTCCGA | 625 | 2.0554313E-5 | 10.596324 | 1 |
| ATACACC | 700 | 6.011129E-6 | 10.36758 | 7 |
| GCACACG | 460 | 3.3051947E-7 | 10.343498 | 124-125 |
| GATCTAT | 1945 | 0.0 | 10.242426 | 2 |
| ACACGTC | 420 | 1.3907498E-4 | 8.901037 | 124-125 |
| ATACACA | 675 | 0.0049241837 | 8.063673 | 7 |
| GTATAGA | 1605 | 0.0 | 8.046477 | 124-125 |
| GTGTATA | 1565 | 0.0 | 7.5388865 | 122-123 |
| TGTATAG | 1635 | 0.0 | 7.422296 | 122-123 |
| CCCCGCC | 490 | 0.008585156 | 6.701728 | 114-115 |
| TCGGAAG | 1135 | 2.3285065E-7 | 6.2370176 | 122-123 |
| AGCACAC | 830 | 1.41902E-4 | 6.092096 | 122-123 |
| CGGAAGA | 1150 | 1.6756694E-6 | 5.91057 | 124-125 |

Produced by FastQC (version 0.11.5)
