## Supplementary material 4 for "Natural and human-mediated drivers of microevolution in Neotropical palms: a historical genomics approach": outF10_AP11.2_pf_trimmed_masked_sickled_fastqc.html

outF10\_AP11.2\_pf\_trimmed\_masked\_sickled.fastq FastQC Report 

FastQC Report

mar. 25 juin 2019  
outF10\_AP11.2\_pf\_trimmed\_masked\_sickled.fastq

### Summary

- Basic Statistics
- Per base sequence quality
- Per tile sequence quality
- Per sequence quality scores
- Per base sequence content
- Per sequence GC content
- Per base N content
- Sequence Length Distribution
- Sequence Duplication Levels
- Overrepresented sequences
- Adapter Content
- Kmer Content

### Basic Statistics

| Measure | Value |
| --- | --- |
| Filename | outF10\_AP11.2\_pf\_trimmed\_masked\_sickled.fastq |
| File type | Conventional base calls |
| Encoding | Sanger / Illumina 1.9 |
| Total Sequences | 1096205 |
| Sequences flagged as poor quality | 0 |
| Sequence length | 100-131 |
| %GC | 44 |

### Per base sequence quality

### Per tile sequence quality

### Per sequence quality scores

### Per base sequence content

### Per sequence GC content

### Per base N content

### Sequence Length Distribution

### Sequence Duplication Levels

### Overrepresented sequences

No overrepresented sequences

### Adapter Content

### Kmer Content

| Sequence | Count | PValue | Obs/Exp Max | Max Obs/Exp Position |
| --- | --- | --- | --- | --- |
| ATACACG | 230 | 0.0 | 36.88873 | 7 |
| TCTATAC | 645 | 0.0 | 34.185863 | 4 |
| TATACAC | 690 | 0.0 | 30.404106 | 6 |
| CTATACA | 740 | 0.0 | 29.114578 | 5 |
| TACACGA | 205 | 3.5222783E-6 | 22.065802 | 8 |
| CGATCTA | 1050 | 0.0 | 19.212114 | 1 |
| ATCTATA | 1420 | 0.0 | 15.505645 | 3 |
| GATCTAT | 1785 | 0.0 | 11.684445 | 2 |
| ATACACA | 695 | 1.2238061E-5 | 9.766225 | 7 |
| GCGTCGT | 370 | 8.746764E-4 | 9.753725 | 122-123 |
| GTATAGA | 1470 | 0.0 | 9.237527 | 124-125 |
| GTCGTGT | 410 | 0.001764065 | 9.032703 | 124-125 |
| AGCGTCG | 460 | 6.510061E-4 | 8.717097 | 122-123 |
| AAGAGCG | 665 | 7.496772E-6 | 8.219363 | 120-121 |
| TACACAT | 710 | 0.0015929957 | 7.9638896 | 8 |
| GTGTATA | 1485 | 0.0 | 7.758731 | 124-125 |
| TCGTGTA | 485 | 0.008031395 | 7.635894 | 124-125 |
| TCGGAAG | 1095 | 3.4428012E-8 | 7.3239536 | 122-123 |
| GAGCGTC | 535 | 0.00375579 | 7.297566 | 120-121 |
| TGTATAG | 1440 | 4.5474735E-11 | 7.2400336 | 122-123 |

Produced by FastQC (version 0.11.5)
