## Supplementary material 4 for "Natural and human-mediated drivers of microevolution in Neotropical palms: a historical genomics approach": outF10_AP13.1_pf_trimmed_masked_sickled_fastqc.html

outF10\_AP13.1\_pf\_trimmed\_masked\_sickled.fastq FastQC Report 

FastQC Report

mar. 25 juin 2019  
outF10\_AP13.1\_pf\_trimmed\_masked\_sickled.fastq

### Summary

- Basic Statistics
- Per base sequence quality
- Per tile sequence quality
- Per sequence quality scores
- Per base sequence content
- Per sequence GC content
- Per base N content
- Sequence Length Distribution
- Sequence Duplication Levels
- Overrepresented sequences
- Adapter Content
- Kmer Content

### Basic Statistics

| Measure | Value |
| --- | --- |
| Filename | outF10\_AP13.1\_pf\_trimmed\_masked\_sickled.fastq |
| File type | Conventional base calls |
| Encoding | Sanger / Illumina 1.9 |
| Total Sequences | 1622386 |
| Sequences flagged as poor quality | 0 |
| Sequence length | 100-131 |
| %GC | 44 |

### Per base sequence quality

### Per tile sequence quality

### Per sequence quality scores

### Per base sequence content

### Per sequence GC content

### Per base N content

### Sequence Length Distribution

### Sequence Duplication Levels

### Overrepresented sequences

No overrepresented sequences

### Adapter Content

### Kmer Content

| Sequence | Count | PValue | Obs/Exp Max | Max Obs/Exp Position |
| --- | --- | --- | --- | --- |
| CTATGTG | 1400 | 0.0 | 30.239025 | 5 |
| TATGTGT | 1720 | 0.0 | 24.620117 | 6 |
| TCTATGT | 1895 | 0.0 | 21.689625 | 4 |
| CGATCTA | 1945 | 0.0 | 21.052761 | 1 |
| ATCTATG | 2645 | 0.0 | 15.97944 | 3 |
| GATCTAT | 3070 | 0.0 | 13.958288 | 2 |
| TTCCGAT | 2005 | 0.0 | 11.412716 | 1 |
| ATGTGTC | 1055 | 8.54925E-11 | 10.902738 | 7 |
| ATGTGTA | 1055 | 9.840733E-10 | 10.328909 | 7 |
| CACACGT | 935 | 0.0 | 9.445712 | 124-125 |
| ATGTGTT | 1385 | 1.3096724E-10 | 9.179181 | 7 |
| ATGTGTG | 1190 | 7.640301E-8 | 8.648412 | 7 |
| ACACGTC | 870 | 1.3460522E-10 | 8.589668 | 124-125 |
| TACGATT | 335 | 0.0038204703 | 8.295059 | 70-71 |
| CACGTCT | 795 | 1.7686034E-8 | 8.118195 | 124-125 |
| CACATAG | 3040 | 0.0 | 7.8216324 | 124-125 |
| TCGGAAG | 2080 | 0.0 | 7.675514 | 124-125 |
| CTGTTTA | 730 | 0.009935034 | 7.456198 | 5 |
| GCACACG | 1040 | 7.6761353E-10 | 7.450703 | 122-123 |
| ACATAGA | 3300 | 0.0 | 7.102449 | 124-125 |

Produced by FastQC (version 0.11.5)
