## Supplementary material 4 for "Natural and human-mediated drivers of microevolution in Neotropical palms: a historical genomics approach": outF10_AP13.2_pf_trimmed_masked_sickled_fastqc.html

outF10\_AP13.2\_pf\_trimmed\_masked\_sickled.fastq FastQC Report 

FastQC Report

mar. 25 juin 2019  
outF10\_AP13.2\_pf\_trimmed\_masked\_sickled.fastq

### Summary

- Basic Statistics
- Per base sequence quality
- Per tile sequence quality
- Per sequence quality scores
- Per base sequence content
- Per sequence GC content
- Per base N content
- Sequence Length Distribution
- Sequence Duplication Levels
- Overrepresented sequences
- Adapter Content
- Kmer Content

### Basic Statistics

| Measure | Value |
| --- | --- |
| Filename | outF10\_AP13.2\_pf\_trimmed\_masked\_sickled.fastq |
| File type | Conventional base calls |
| Encoding | Sanger / Illumina 1.9 |
| Total Sequences | 1622386 |
| Sequences flagged as poor quality | 0 |
| Sequence length | 100-131 |
| %GC | 44 |

### Per base sequence quality

### Per tile sequence quality

### Per sequence quality scores

### Per base sequence content

### Per sequence GC content

### Per base N content

### Sequence Length Distribution

### Sequence Duplication Levels

### Overrepresented sequences

No overrepresented sequences

### Adapter Content

### Kmer Content

| Sequence | Count | PValue | Obs/Exp Max | Max Obs/Exp Position |
| --- | --- | --- | --- | --- |
| CTATGTG | 1405 | 0.0 | 31.09996 | 5 |
| TATGTGT | 1865 | 0.0 | 24.071772 | 6 |
| CGATCTA | 1860 | 0.0 | 23.56201 | 1 |
| TCTATGT | 1855 | 0.0 | 23.506216 | 4 |
| ATCTATG | 2460 | 0.0 | 17.95668 | 3 |
| CTTCCGA | 770 | 0.0 | 16.261723 | 1 |
| GATCTAT | 3155 | 0.0 | 13.781867 | 2 |
| TTCCGAT | 1860 | 0.0 | 12.852005 | 1 |
| ATGTGTG | 1265 | 0.0 | 12.267028 | 7 |
| ATGTGTC | 1005 | 0.0 | 12.009345 | 7 |
| AGAGCGT | 1015 | 9.094947E-12 | 9.186515 | 124-125 |
| CGCGTGG | 300 | 0.0070152073 | 9.0051155 | 102-103 |
| ATGTGTT | 1540 | 3.8198777E-11 | 8.5836735 | 7 |
| AAGAGCG | 1480 | 0.0 | 8.49159 | 124-125 |
| CACATAG | 3185 | 0.0 | 8.146284 | 124-125 |
| TCTTCCG | 710 | 0.0014941412 | 8.016341 | 1 |
| ATGTGTA | 1005 | 1.06591E-5 | 8.00623 | 7 |
| GAGTGTA | 520 | 0.0019663859 | 7.7962484 | 124-125 |
| TGTGTGA | 1115 | 5.161328E-6 | 7.7247505 | 8 |
| TCGGAAG | 1950 | 0.0 | 7.6922984 | 124-125 |

Produced by FastQC (version 0.11.5)
