## Supplementary material 4 for "Natural and human-mediated drivers of microevolution in Neotropical palms: a historical genomics approach": outF10_AP14.1_pf_trimmed_masked_sickled_fastqc.html

outF10\_AP14.1\_pf\_trimmed\_masked\_sickled.fastq FastQC Report 

FastQC Report

mar. 25 juin 2019  
outF10\_AP14.1\_pf\_trimmed\_masked\_sickled.fastq

### Summary

- Basic Statistics
- Per base sequence quality
- Per tile sequence quality
- Per sequence quality scores
- Per base sequence content
- Per sequence GC content
- Per base N content
- Sequence Length Distribution
- Sequence Duplication Levels
- Overrepresented sequences
- Adapter Content
- Kmer Content

### Basic Statistics

| Measure | Value |
| --- | --- |
| Filename | outF10\_AP14.1\_pf\_trimmed\_masked\_sickled.fastq |
| File type | Conventional base calls |
| Encoding | Sanger / Illumina 1.9 |
| Total Sequences | 1857218 |
| Sequences flagged as poor quality | 0 |
| Sequence length | 100-131 |
| %GC | 44 |

### Per base sequence quality

### Per tile sequence quality

### Per sequence quality scores

### Per base sequence content

### Per sequence GC content

### Per base N content

### Sequence Length Distribution

### Sequence Duplication Levels

### Overrepresented sequences

No overrepresented sequences

### Adapter Content

### Kmer Content

| Sequence | Count | PValue | Obs/Exp Max | Max Obs/Exp Position |
| --- | --- | --- | --- | --- |
| TCACTTA | 1225 | 0.0 | 16.827166 | 6 |
| CTCACTT | 1630 | 0.0 | 12.267846 | 5 |
| CACACGT | 1015 | 0.0 | 9.684693 | 124-125 |
| TCTCACT | 2120 | 0.0 | 9.430028 | 4 |
| CGATCTC | 1975 | 0.0 | 8.863608 | 1 |
| CACTTAC | 830 | 4.380774E-5 | 8.767072 | 7 |
| ACACGTC | 1020 | 0.0 | 8.640265 | 124-125 |
| GCACACG | 1160 | 0.0 | 8.116723 | 122-123 |
| GCGTCAA | 420 | 0.0019085066 | 7.8202453 | 116-117 |
| CGGAAGA | 2485 | 0.0 | 7.5022273 | 124-125 |
| ACGTCTG | 955 | 9.100404E-9 | 7.394292 | 122-123 |
| TCGGAAG | 2670 | 0.0 | 7.236316 | 124-125 |
| AAGTGAG | 4380 | 0.0 | 7.1971865 | 124-125 |
| CACGTCT | 960 | 9.116775E-8 | 6.9463716 | 120-121 |
| CACTTAA | 1070 | 8.265393E-4 | 6.8006263 | 7 |
| AGAGCAC | 2430 | 0.0 | 6.5560994 | 124-125 |
| TAAGTGA | 4460 | 0.0 | 6.5360827 | 124-125 |
| ATCTCAC | 2975 | 1.8189894E-12 | 6.513578 | 3 |
| CGTCTGA | 1090 | 1.13943315E-7 | 6.478485 | 122-123 |
| GATCTCA | 3195 | 0.0 | 6.439199 | 2 |

Produced by FastQC (version 0.11.5)
