## Supplementary material 4 for "Natural and human-mediated drivers of microevolution in Neotropical palms: a historical genomics approach": outF10_AP14.2_pf_trimmed_masked_sickled_fastqc.html

outF10\_AP14.2\_pf\_trimmed\_masked\_sickled.fastq FastQC Report 

FastQC Report

mar. 25 juin 2019  
outF10\_AP14.2\_pf\_trimmed\_masked\_sickled.fastq

### Summary

- Basic Statistics
- Per base sequence quality
- Per tile sequence quality
- Per sequence quality scores
- Per base sequence content
- Per sequence GC content
- Per base N content
- Sequence Length Distribution
- Sequence Duplication Levels
- Overrepresented sequences
- Adapter Content
- Kmer Content

### Basic Statistics

| Measure | Value |
| --- | --- |
| Filename | outF10\_AP14.2\_pf\_trimmed\_masked\_sickled.fastq |
| File type | Conventional base calls |
| Encoding | Sanger / Illumina 1.9 |
| Total Sequences | 1857218 |
| Sequences flagged as poor quality | 0 |
| Sequence length | 100-131 |
| %GC | 44 |

### Per base sequence quality

### Per tile sequence quality

### Per sequence quality scores

### Per base sequence content

### Per sequence GC content

### Per base N content

### Sequence Length Distribution

### Sequence Duplication Levels

### Overrepresented sequences

No overrepresented sequences

### Adapter Content

### Kmer Content

| Sequence | Count | PValue | Obs/Exp Max | Max Obs/Exp Position |
| --- | --- | --- | --- | --- |
| TCACTTA | 1375 | 0.0 | 16.29378 | 6 |
| CTCACTT | 1475 | 0.0 | 14.019932 | 5 |
| CGATCTC | 1675 | 0.0 | 12.605196 | 1 |
| TCTCACT | 1985 | 0.0 | 11.267847 | 4 |
| AAGAGCG | 1745 | 0.0 | 11.118875 | 124-125 |
| CACTTAG | 730 | 1.3835052E-7 | 11.018289 | 7 |
| GAGCGTC | 1120 | 0.0 | 10.252746 | 124-125 |
| AGAGCGT | 1245 | 0.0 | 9.859446 | 124-125 |
| ACGTCAG | 300 | 0.0011757518 | 9.444291 | 88-89 |
| AGCGTCG | 1010 | 1.2732926E-11 | 9.017095 | 124-125 |
| ACCGTTT | 320 | 0.008708257 | 8.770777 | 110-111 |
| TCGGAAG | 2580 | 0.0 | 8.420892 | 122-123 |
| CGGAAGA | 2550 | 0.0 | 8.367819 | 122-123 |
| GTAGGGA | 910 | 1.27820385E-8 | 8.267466 | 124-125 |
| CGTGTAG | 625 | 2.5757952E-5 | 8.236136 | 124-125 |
| GATCTCA | 2970 | 0.0 | 7.917065 | 2 |
| ATCTCAC | 3200 | 0.0 | 7.7086406 | 3 |
| TCGTGTA | 675 | 6.73061E-5 | 7.626052 | 124-125 |
| TAAGTGA | 4250 | 0.0 | 7.576679 | 122-123 |
| AGTGAGA | 4675 | 0.0 | 7.368818 | 124-125 |

Produced by FastQC (version 0.11.5)
