## Supplementary material 4 for "Natural and human-mediated drivers of microevolution in Neotropical palms: a historical genomics approach": outF10_AP16.1_pf_trimmed_masked_sickled_fastqc.html

outF10\_AP16.1\_pf\_trimmed\_masked\_sickled.fastq FastQC Report 

FastQC Report

mar. 25 juin 2019  
outF10\_AP16.1\_pf\_trimmed\_masked\_sickled.fastq

### Summary

- Basic Statistics
- Per base sequence quality
- Per tile sequence quality
- Per sequence quality scores
- Per base sequence content
- Per sequence GC content
- Per base N content
- Sequence Length Distribution
- Sequence Duplication Levels
- Overrepresented sequences
- Adapter Content
- Kmer Content

### Basic Statistics

| Measure | Value |
| --- | --- |
| Filename | outF10\_AP16.1\_pf\_trimmed\_masked\_sickled.fastq |
| File type | Conventional base calls |
| Encoding | Sanger / Illumina 1.9 |
| Total Sequences | 1145109 |
| Sequences flagged as poor quality | 0 |
| Sequence length | 100-131 |
| %GC | 44 |

### Per base sequence quality

### Per tile sequence quality

### Per sequence quality scores

### Per base sequence content

### Per sequence GC content

### Per base N content

### Sequence Length Distribution

### Sequence Duplication Levels

### Overrepresented sequences

No overrepresented sequences

### Adapter Content

### Kmer Content

| Sequence | Count | PValue | Obs/Exp Max | Max Obs/Exp Position |
| --- | --- | --- | --- | --- |
| TCCATAG | 1375 | 0.0 | 37.343258 | 6 |
| CGATCTC | 1680 | 0.0 | 30.074533 | 1 |
| CTCCATA | 1730 | 0.0 | 29.673746 | 5 |
| GATCTCC | 2550 | 0.0 | 20.332327 | 2 |
| ATCTCCA | 2545 | 0.0 | 19.902908 | 3 |
| CCATAGT | 720 | 0.0 | 19.303926 | 7 |
| TCTCCAT | 2830 | 0.0 | 18.12288 | 4 |
| CCATAGC | 875 | 0.0 | 17.95625 | 7 |
| CCATAGG | 710 | 2.3464963E-10 | 13.617957 | 7 |
| CTTCCGA | 585 | 5.12955E-8 | 13.366459 | 1 |
| CCATAGA | 1025 | 3.6379788E-12 | 11.791159 | 7 |
| CATAGTG | 545 | 7.048713E-4 | 9.985426 | 8 |
| TCGGAAG | 1155 | 0.0 | 8.5491705 | 124-125 |
| CGGAAGA | 1175 | 0.0 | 8.403653 | 124-125 |
| CTATGGA | 2025 | 0.0 | 8.070574 | 118-119 |
| CATAGCT | 860 | 6.3243665E-4 | 7.7341895 | 8 |
| GAGATCG | 1910 | 0.0 | 7.6655455 | 124-125 |
| GGAGATC | 2305 | 0.0 | 6.8953304 | 122-123 |
| TATGGAG | 2325 | 0.0 | 6.3573117 | 120-121 |
| ACTATGG | 790 | 5.6845706E-4 | 5.9106374 | 118-119 |

Produced by FastQC (version 0.11.5)
