## Supplementary material 4 for "Natural and human-mediated drivers of microevolution in Neotropical palms: a historical genomics approach": outF10_AP16.2_pf_trimmed_masked_sickled_fastqc.html

outF10\_AP16.2\_pf\_trimmed\_masked\_sickled.fastq FastQC Report 

FastQC Report

mar. 25 juin 2019  
outF10\_AP16.2\_pf\_trimmed\_masked\_sickled.fastq

### Summary

- Basic Statistics
- Per base sequence quality
- Per tile sequence quality
- Per sequence quality scores
- Per base sequence content
- Per sequence GC content
- Per base N content
- Sequence Length Distribution
- Sequence Duplication Levels
- Overrepresented sequences
- Adapter Content
- Kmer Content

### Basic Statistics

| Measure | Value |
| --- | --- |
| Filename | outF10\_AP16.2\_pf\_trimmed\_masked\_sickled.fastq |
| File type | Conventional base calls |
| Encoding | Sanger / Illumina 1.9 |
| Total Sequences | 1145109 |
| Sequences flagged as poor quality | 0 |
| Sequence length | 100-131 |
| %GC | 44 |

### Per base sequence quality

### Per tile sequence quality

### Per sequence quality scores

### Per base sequence content

### Per sequence GC content

### Per base N content

### Sequence Length Distribution

### Sequence Duplication Levels

### Overrepresented sequences

No overrepresented sequences

### Adapter Content

### Kmer Content

| Sequence | Count | PValue | Obs/Exp Max | Max Obs/Exp Position |
| --- | --- | --- | --- | --- |
| TCCATAG | 1350 | 0.0 | 37.81591 | 6 |
| CGATCTC | 1575 | 0.0 | 32.01752 | 1 |
| CTCCATA | 1700 | 0.0 | 28.640938 | 5 |
| GATCTCC | 2200 | 0.0 | 23.301369 | 2 |
| ATCTCCA | 2345 | 0.0 | 21.631548 | 3 |
| CTTCCGA | 535 | 0.0 | 21.181387 | 1 |
| CCATAGT | 570 | 0.0 | 21.12235 | 7 |
| TCTCCAT | 2520 | 0.0 | 19.931526 | 4 |
| CCATAGA | 1010 | 0.0 | 16.46169 | 7 |
| CCATAGG | 720 | 0.0 | 15.129302 | 7 |
| CCATAGC | 870 | 0.0 | 14.497771 | 7 |
| CATAGTG | 430 | 1.2990914E-4 | 11.9870205 | 8 |
| CGGAAGA | 945 | 0.0 | 11.841926 | 124-125 |
| TTCCGAT | 1215 | 0.0 | 10.782344 | 2 |
| TCGGAAG | 1075 | 0.0 | 9.666317 | 124-125 |
| CATAGCA | 700 | 1.2322175E-4 | 8.999779 | 8 |
| CTATGGA | 2165 | 0.0 | 8.725524 | 118-119 |
| AAGAGCG | 650 | 4.026502E-6 | 8.60817 | 124-125 |
| GAGATCG | 1755 | 0.0 | 8.425985 | 124-125 |
| GGAGATC | 2260 | 0.0 | 8.161977 | 122-123 |

Produced by FastQC (version 0.11.5)
