## Supplementary material 4 for "Natural and human-mediated drivers of microevolution in Neotropical palms: a historical genomics approach": outF10_AP20.1_pf_trimmed_masked_sickled_fastqc.html

outF10\_AP20.1\_pf\_trimmed\_masked\_sickled.fastq FastQC Report 

FastQC Report

mar. 25 juin 2019  
outF10\_AP20.1\_pf\_trimmed\_masked\_sickled.fastq

### Summary

- Basic Statistics
- Per base sequence quality
- Per tile sequence quality
- Per sequence quality scores
- Per base sequence content
- Per sequence GC content
- Per base N content
- Sequence Length Distribution
- Sequence Duplication Levels
- Overrepresented sequences
- Adapter Content
- Kmer Content

### Basic Statistics

| Measure | Value |
| --- | --- |
| Filename | outF10\_AP20.1\_pf\_trimmed\_masked\_sickled.fastq |
| File type | Conventional base calls |
| Encoding | Sanger / Illumina 1.9 |
| Total Sequences | 1172641 |
| Sequences flagged as poor quality | 0 |
| Sequence length | 100-131 |
| %GC | 43 |

### Per base sequence quality

### Per tile sequence quality

### Per sequence quality scores

### Per base sequence content

### Per sequence GC content

### Per base N content

### Sequence Length Distribution

### Sequence Duplication Levels

### Overrepresented sequences

No overrepresented sequences

### Adapter Content

### Kmer Content

| Sequence | Count | PValue | Obs/Exp Max | Max Obs/Exp Position |
| --- | --- | --- | --- | --- |
| CTGAGGC | 890 | 0.0 | 27.946527 | 5 |
| TGAGGCA | 1330 | 0.0 | 20.474913 | 6 |
| TCTGAGG | 1300 | 0.0 | 18.63776 | 4 |
| CACACGT | 605 | 0.0 | 14.741616 | 124-125 |
| GCACACG | 705 | 0.0 | 14.16868 | 124-125 |
| ACACGTC | 590 | 0.0 | 13.302435 | 124-125 |
| ATCTGAG | 2115 | 0.0 | 12.00465 | 3 |
| AGCACAC | 1115 | 0.0 | 9.9185295 | 124-125 |
| ACGTCTG | 540 | 1.4325087E-7 | 9.909642 | 124-125 |
| CACGTCT | 535 | 1.4736725E-7 | 9.890428 | 122-123 |
| CGTCTGA | 690 | 7.225026E-9 | 9.202397 | 122-123 |
| CGGAAGA | 1545 | 0.0 | 9.005266 | 124-125 |
| GAGGCAC | 620 | 0.002780742 | 8.590791 | 7 |
| GAGGCAA | 920 | 2.1239623E-5 | 8.362533 | 7 |
| GCCTCAG | 2755 | 0.0 | 8.287411 | 124-125 |
| TCGGAAG | 1640 | 0.0 | 8.048563 | 124-125 |
| CCTCAGA | 2840 | 0.0 | 7.536911 | 124-125 |
| GAGGCAG | 1045 | 1.03949584E-4 | 7.36223 | 7 |
| AGAGCAC | 1460 | 0.0 | 7.248463 | 122-123 |
| CGATCTG | 3940 | 0.0 | 6.865127 | 1 |

Produced by FastQC (version 0.11.5)
