## Supplementary material 4 for "Natural and human-mediated drivers of microevolution in Neotropical palms: a historical genomics approach": outF10_AP20.2_pf_trimmed_masked_sickled_fastqc.html

outF10\_AP20.2\_pf\_trimmed\_masked\_sickled.fastq FastQC Report 

FastQC Report

mar. 25 juin 2019  
outF10\_AP20.2\_pf\_trimmed\_masked\_sickled.fastq

### Summary

- Basic Statistics
- Per base sequence quality
- Per tile sequence quality
- Per sequence quality scores
- Per base sequence content
- Per sequence GC content
- Per base N content
- Sequence Length Distribution
- Sequence Duplication Levels
- Overrepresented sequences
- Adapter Content
- Kmer Content

### Basic Statistics

| Measure | Value |
| --- | --- |
| Filename | outF10\_AP20.2\_pf\_trimmed\_masked\_sickled.fastq |
| File type | Conventional base calls |
| Encoding | Sanger / Illumina 1.9 |
| Total Sequences | 1172641 |
| Sequences flagged as poor quality | 0 |
| Sequence length | 100-131 |
| %GC | 44 |

### Per base sequence quality

### Per tile sequence quality

### Per sequence quality scores

### Per base sequence content

### Per sequence GC content

### Per base N content

### Sequence Length Distribution

### Sequence Duplication Levels

### Overrepresented sequences

No overrepresented sequences

### Adapter Content

### Kmer Content

| Sequence | Count | PValue | Obs/Exp Max | Max Obs/Exp Position |
| --- | --- | --- | --- | --- |
| CTGAGGC | 965 | 0.0 | 22.643326 | 5 |
| TCTGAGG | 1090 | 0.0 | 21.553402 | 4 |
| TGAGGCA | 1205 | 0.0 | 18.179714 | 6 |
| GCGTCGT | 420 | 4.432877E-9 | 14.099834 | 124-125 |
| AGCGTCG | 490 | 1.8735591E-10 | 13.812083 | 124-125 |
| CGTCGTG | 435 | 7.2614057E-9 | 13.613633 | 124-125 |
| GTCGTGT | 295 | 6.521509E-5 | 12.904934 | 124-125 |
| CGTGTAG | 300 | 7.633102E-5 | 12.689852 | 124-125 |
| ATCTGAG | 1875 | 0.0 | 11.9146805 | 3 |
| TCGTGTA | 280 | 7.599024E-4 | 11.768232 | 122-123 |
| CAACCGG | 340 | 0.0043329466 | 11.545087 | 8 |
| GAGCGTC | 675 | 2.1282176E-10 | 11.279868 | 124-125 |
| CGGAAGA | 1475 | 0.0 | 11.184277 | 124-125 |
| GAGGCAC | 570 | 1.6278913E-5 | 10.822203 | 7 |
| AAGAGCG | 1020 | 0.0 | 10.587084 | 120-121 |
| AGAGCGT | 730 | 1.9826984E-10 | 10.409827 | 120-121 |
| TCGGAAG | 1450 | 0.0 | 9.918505 | 124-125 |
| GCCTCAG | 2610 | 0.0 | 9.886091 | 124-125 |
| GTGTAGG | 615 | 8.87269E-5 | 8.253562 | 124-125 |
| CCTCAGA | 2770 | 0.0 | 8.246113 | 124-125 |

Produced by FastQC (version 0.11.5)
