## Supplementary material 4 for "Natural and human-mediated drivers of microevolution in Neotropical palms: a historical genomics approach": outF10_AS01.1_pf_trimmed_masked_sickled_fastqc.html

outF10\_AS01.1\_pf\_trimmed\_masked\_sickled.fastq FastQC Report 

FastQC Report

mar. 25 juin 2019  
outF10\_AS01.1\_pf\_trimmed\_masked\_sickled.fastq

### Summary

- Basic Statistics
- Per base sequence quality
- Per tile sequence quality
- Per sequence quality scores
- Per base sequence content
- Per sequence GC content
- Per base N content
- Sequence Length Distribution
- Sequence Duplication Levels
- Overrepresented sequences
- Adapter Content
- Kmer Content

### Basic Statistics

| Measure | Value |
| --- | --- |
| Filename | outF10\_AS01.1\_pf\_trimmed\_masked\_sickled.fastq |
| File type | Conventional base calls |
| Encoding | Sanger / Illumina 1.9 |
| Total Sequences | 1760189 |
| Sequences flagged as poor quality | 0 |
| Sequence length | 100-131 |
| %GC | 44 |

### Per base sequence quality

### Per tile sequence quality

### Per sequence quality scores

### Per base sequence content

### Per sequence GC content

### Per base N content

### Sequence Length Distribution

### Sequence Duplication Levels

### Overrepresented sequences

No overrepresented sequences

### Adapter Content

### Kmer Content

| Sequence | Count | PValue | Obs/Exp Max | Max Obs/Exp Position |
| --- | --- | --- | --- | --- |
| CGATCTC | 2080 | 0.0 | 11.483281 | 1 |
| ATCTCCT | 2800 | 0.0 | 8.991023 | 3 |
| CTTCCGA | 955 | 3.2116986E-6 | 8.753747 | 1 |
| GCACACG | 1365 | 0.0 | 8.6427 | 124-125 |
| GATCTCC | 3000 | 0.0 | 8.384806 | 2 |
| CACACGT | 1205 | 0.0 | 7.9726796 | 122-123 |
| TCGGAAG | 2795 | 0.0 | 7.9451365 | 124-125 |
| CGCCCGA | 365 | 0.008484607 | 7.5897274 | 70-71 |
| CGGAAGA | 2530 | 0.0 | 7.543022 | 124-125 |
| ACACGTC | 1120 | 2.5102054E-10 | 7.435264 | 124-125 |
| CACGTCT | 1140 | 3.7107384E-10 | 7.304821 | 124-125 |
| TCCTCCA | 4165 | 0.0 | 7.06532 | 6 |
| CGTCTGA | 1150 | 4.296453E-9 | 6.862199 | 122-123 |
| ACGTCTG | 1050 | 4.760841E-8 | 6.783069 | 120-121 |
| AGCACAC | 2000 | 0.0 | 6.76609 | 124-125 |
| TCTCCTC | 3880 | 0.0 | 6.3373775 | 4 |
| GAAGAGC | 3490 | 0.0 | 6.064676 | 124-125 |
| CCTCCAG | 2010 | 1.3610588E-6 | 5.9798183 | 7 |
| AGAGCAC | 2220 | 1.8189894E-12 | 5.6266866 | 124-125 |
| GAGATCG | 3985 | 0.0 | 5.6170964 | 120-121 |

Produced by FastQC (version 0.11.5)
