## Supplementary material 4 for "Natural and human-mediated drivers of microevolution in Neotropical palms: a historical genomics approach": outF10_AS01.2_pf_trimmed_masked_sickled_fastqc.html

outF10\_AS01.2\_pf\_trimmed\_masked\_sickled.fastq FastQC Report 

FastQC Report

mar. 25 juin 2019  
outF10\_AS01.2\_pf\_trimmed\_masked\_sickled.fastq

### Summary

- Basic Statistics
- Per base sequence quality
- Per tile sequence quality
- Per sequence quality scores
- Per base sequence content
- Per sequence GC content
- Per base N content
- Sequence Length Distribution
- Sequence Duplication Levels
- Overrepresented sequences
- Adapter Content
- Kmer Content

### Basic Statistics

| Measure | Value |
| --- | --- |
| Filename | outF10\_AS01.2\_pf\_trimmed\_masked\_sickled.fastq |
| File type | Conventional base calls |
| Encoding | Sanger / Illumina 1.9 |
| Total Sequences | 1760189 |
| Sequences flagged as poor quality | 0 |
| Sequence length | 100-131 |
| %GC | 44 |

### Per base sequence quality

### Per tile sequence quality

### Per sequence quality scores

### Per base sequence content

### Per sequence GC content

### Per base N content

### Sequence Length Distribution

### Sequence Duplication Levels

### Overrepresented sequences

No overrepresented sequences

### Adapter Content

### Kmer Content

| Sequence | Count | PValue | Obs/Exp Max | Max Obs/Exp Position |
| --- | --- | --- | --- | --- |
| CGATCTC | 1760 | 0.0 | 12.198493 | 1 |
| AAGAGCG | 1765 | 0.0 | 10.082195 | 124-125 |
| TCGGAAG | 2615 | 0.0 | 10.05284 | 124-125 |
| CGGAAGA | 2575 | 0.0 | 8.779997 | 122-123 |
| AGCGTCG | 1030 | 1.200533E-10 | 8.638386 | 124-125 |
| GAGCGTC | 1175 | 2.5465852E-11 | 8.260769 | 124-125 |
| GCGTCGT | 800 | 7.7562436E-7 | 8.08867 | 124-125 |
| AGAGCGT | 1255 | 1.2732926E-11 | 8.056444 | 124-125 |
| GATCTCC | 2615 | 0.0 | 8.035989 | 2 |
| GTCGTGT | 720 | 1.355588E-5 | 7.8639846 | 124-125 |
| CGTGTAG | 635 | 2.1674967E-4 | 7.642837 | 124-125 |
| ATCTCCT | 2905 | 0.0 | 7.4405994 | 3 |
| GAGATCG | 3830 | 0.0 | 6.7581577 | 124-125 |
| GTAGGGA | 965 | 1.2822733E-5 | 6.705633 | 124-125 |
| TCGTGTA | 725 | 3.6366674E-4 | 6.649374 | 116-117 |
| TCTCCTC | 3440 | 0.0 | 6.454358 | 4 |
| GAAGAGC | 3490 | 0.0 | 6.373594 | 124-125 |
| TCCTCCA | 3805 | 0.0 | 6.2978244 | 6 |
| CGTCGTG | 855 | 9.380334E-4 | 6.149281 | 124-125 |
| GAGGAGA | 5855 | 0.0 | 6.1476655 | 124-125 |

Produced by FastQC (version 0.11.5)
