## Supplementary material 4 for "Natural and human-mediated drivers of microevolution in Neotropical palms: a historical genomics approach": outF10_AS06.1_pf_trimmed_masked_sickled_fastqc.html

outF10\_AS06.1\_pf\_trimmed\_masked\_sickled.fastq FastQC Report 

FastQC Report

mar. 25 juin 2019  
outF10\_AS06.1\_pf\_trimmed\_masked\_sickled.fastq

### Summary

- Basic Statistics
- Per base sequence quality
- Per tile sequence quality
- Per sequence quality scores
- Per base sequence content
- Per sequence GC content
- Per base N content
- Sequence Length Distribution
- Sequence Duplication Levels
- Overrepresented sequences
- Adapter Content
- Kmer Content

### Basic Statistics

| Measure | Value |
| --- | --- |
| Filename | outF10\_AS06.1\_pf\_trimmed\_masked\_sickled.fastq |
| File type | Conventional base calls |
| Encoding | Sanger / Illumina 1.9 |
| Total Sequences | 1308851 |
| Sequences flagged as poor quality | 0 |
| Sequence length | 100-131 |
| %GC | 45 |

### Per base sequence quality

### Per tile sequence quality

### Per sequence quality scores

### Per base sequence content

### Per sequence GC content

### Per base N content

### Sequence Length Distribution

### Sequence Duplication Levels

### Overrepresented sequences

No overrepresented sequences

### Adapter Content

### Kmer Content

| Sequence | Count | PValue | Obs/Exp Max | Max Obs/Exp Position |
| --- | --- | --- | --- | --- |
| CTCCTGT | 1265 | 0.0 | 26.778063 | 5 |
| TCCTGTA | 1270 | 0.0 | 26.217787 | 6 |
| TCTCCTG | 1550 | 0.0 | 22.613733 | 4 |
| CGATCTC | 1520 | 0.0 | 22.568512 | 1 |
| CCTGTAC | 475 | 1.5825208E-10 | 17.843147 | 7 |
| CCTGTAG | 795 | 0.0 | 17.5145 | 7 |
| ATCTCCT | 2465 | 0.0 | 13.958673 | 3 |
| GATCTCC | 2465 | 0.0 | 13.462616 | 2 |
| CTTCCGA | 755 | 9.036739E-9 | 11.956826 | 1 |
| CACACGT | 1040 | 0.0 | 10.800805 | 124-125 |
| ACGTCTG | 765 | 0.0 | 10.678871 | 124-125 |
| CGTCTGA | 855 | 0.0 | 10.35101 | 124-125 |
| CCTGTAA | 610 | 1.7739182E-4 | 9.924467 | 7 |
| CACGTCT | 945 | 0.0 | 9.666507 | 122-123 |
| GCACACG | 1105 | 0.0 | 9.549375 | 124-125 |
| TACAGGA | 3690 | 0.0 | 8.343584 | 122-123 |
| ACACGTC | 995 | 2.7284841E-11 | 8.210388 | 124-125 |
| AGCACAC | 1530 | 0.0 | 7.9606524 | 122-123 |
| TCGGAAG | 2320 | 0.0 | 7.9228473 | 124-125 |
| CAGGAGA | 3825 | 0.0 | 7.920162 | 124-125 |

Produced by FastQC (version 0.11.5)
