## Supplementary material 4 for "Natural and human-mediated drivers of microevolution in Neotropical palms: a historical genomics approach": outF10_AS06.2_pf_trimmed_masked_sickled_fastqc.html

outF10\_AS06.2\_pf\_trimmed\_masked\_sickled.fastq FastQC Report 

FastQC Report

mar. 25 juin 2019  
outF10\_AS06.2\_pf\_trimmed\_masked\_sickled.fastq

### Summary

- Basic Statistics
- Per base sequence quality
- Per tile sequence quality
- Per sequence quality scores
- Per base sequence content
- Per sequence GC content
- Per base N content
- Sequence Length Distribution
- Sequence Duplication Levels
- Overrepresented sequences
- Adapter Content
- Kmer Content

### Basic Statistics

| Measure | Value |
| --- | --- |
| Filename | outF10\_AS06.2\_pf\_trimmed\_masked\_sickled.fastq |
| File type | Conventional base calls |
| Encoding | Sanger / Illumina 1.9 |
| Total Sequences | 1308851 |
| Sequences flagged as poor quality | 0 |
| Sequence length | 100-131 |
| %GC | 45 |

### Per base sequence quality

### Per tile sequence quality

### Per sequence quality scores

### Per base sequence content

### Per sequence GC content

### Per base N content

### Sequence Length Distribution

### Sequence Duplication Levels

### Overrepresented sequences

No overrepresented sequences

### Adapter Content

### Kmer Content

| Sequence | Count | PValue | Obs/Exp Max | Max Obs/Exp Position |
| --- | --- | --- | --- | --- |
| TCCTGTA | 965 | 0.0 | 23.70401 | 6 |
| CTCCTGT | 1160 | 0.0 | 19.693087 | 5 |
| CGATCTC | 1560 | 0.0 | 17.023895 | 1 |
| TCTCCTG | 1470 | 0.0 | 16.292858 | 4 |
| CCTGTAG | 570 | 7.0222086E-8 | 13.049768 | 7 |
| CCTGTAA | 615 | 1.8913124E-7 | 12.094907 | 7 |
| GATCTCC | 2305 | 0.0 | 11.341508 | 2 |
| ATCTCCT | 2410 | 0.0 | 11.118377 | 3 |
| TTTCCGG | 680 | 8.167608E-7 | 10.802396 | 1 |
| CGTGTAG | 600 | 9.955329E-9 | 10.7356415 | 124-125 |
| AGCGTCG | 905 | 0.0 | 10.452948 | 122-123 |
| CTTCCGA | 615 | 3.4645513E-5 | 10.106556 | 1 |
| TCGTGTA | 560 | 4.80788E-7 | 10.064664 | 124-125 |
| CGTCGTG | 880 | 5.456968E-12 | 10.064664 | 124-125 |
| CCGGTCA | 775 | 3.8795442E-6 | 9.565484 | 4 |
| AGAGCGT | 1060 | 0.0 | 9.460742 | 120-121 |
| GTCGTGT | 650 | 3.6589336E-7 | 9.29046 | 124-125 |
| GCGTCGT | 770 | 7.064955E-9 | 9.214205 | 122-123 |
| GTGTAGG | 745 | 2.8740033E-8 | 9.186539 | 124-125 |
| CGGTCAT | 935 | 4.470894E-7 | 9.162024 | 5 |

Produced by FastQC (version 0.11.5)
