## Supplementary material 4 for "Natural and human-mediated drivers of microevolution in Neotropical palms: a historical genomics approach": outF10_AS07.1_pf_trimmed_masked_sickled_fastqc.html

outF10\_AS07.1\_pf\_trimmed\_masked\_sickled.fastq FastQC Report 

FastQC Report

mar. 25 juin 2019  
outF10\_AS07.1\_pf\_trimmed\_masked\_sickled.fastq

### Summary

- Basic Statistics
- Per base sequence quality
- Per tile sequence quality
- Per sequence quality scores
- Per base sequence content
- Per sequence GC content
- Per base N content
- Sequence Length Distribution
- Sequence Duplication Levels
- Overrepresented sequences
- Adapter Content
- Kmer Content

### Basic Statistics

| Measure | Value |
| --- | --- |
| Filename | outF10\_AS07.1\_pf\_trimmed\_masked\_sickled.fastq |
| File type | Conventional base calls |
| Encoding | Sanger / Illumina 1.9 |
| Total Sequences | 826637 |
| Sequences flagged as poor quality | 0 |
| Sequence length | 100-131 |
| %GC | 44 |

### Per base sequence quality

### Per tile sequence quality

### Per sequence quality scores

### Per base sequence content

### Per sequence GC content

### Per base N content

### Sequence Length Distribution

### Sequence Duplication Levels

### Overrepresented sequences

No overrepresented sequences

### Adapter Content

### Kmer Content

| Sequence | Count | PValue | Obs/Exp Max | Max Obs/Exp Position |
| --- | --- | --- | --- | --- |
| CTCGACT | 470 | 0.0 | 31.2339 | 5 |
| TCTCGAC | 610 | 0.0 | 23.08703 | 4 |
| CGACTCC | 385 | 5.857146E-9 | 18.31366 | 7 |
| CGATCTC | 905 | 0.0 | 15.459445 | 1 |
| TCGACTC | 1085 | 0.0 | 15.160089 | 6 |
| CACGACT | 120 | 0.0042062066 | 14.746052 | 10-11 |
| ATCTCGA | 1075 | 0.0 | 13.068018 | 3 |
| GCACACG | 485 | 1.1641532E-9 | 12.307782 | 124-125 |
| CACACGT | 430 | 3.5739504E-8 | 12.146779 | 124-125 |
| AGTCGAG | 1530 | 0.0 | 9.7537155 | 124-125 |
| ACGTCTG | 325 | 0.0011226477 | 9.491358 | 112-113 |
| CGTCTGA | 370 | 0.0019955256 | 8.910964 | 122-123 |
| GATCTCG | 1840 | 0.0 | 8.907322 | 2 |
| CGAGTGC | 270 | 0.008133241 | 8.843789 | 28-29 |
| GGAGTCG | 810 | 6.3919288E-9 | 8.593151 | 122-123 |
| GTCGAGA | 1385 | 0.0 | 7.9351535 | 122-123 |
| CGAGTCG | 615 | 4.4266308E-5 | 7.8862367 | 124-125 |
| CGGAAGA | 985 | 2.5853296E-8 | 7.438368 | 122-123 |
| TCGGAAG | 915 | 3.3201468E-7 | 7.3392725 | 124-125 |
| GAGATCG | 1420 | 1.8007995E-10 | 6.831036 | 124-125 |

Produced by FastQC (version 0.11.5)
