## Supplementary material 4 for "Natural and human-mediated drivers of microevolution in Neotropical palms: a historical genomics approach": outF10_AS07.2_pf_trimmed_masked_sickled_fastqc.html

outF10\_AS07.2\_pf\_trimmed\_masked\_sickled.fastq FastQC Report 

FastQC Report

mar. 25 juin 2019  
outF10\_AS07.2\_pf\_trimmed\_masked\_sickled.fastq

### Summary

- Basic Statistics
- Per base sequence quality
- Per tile sequence quality
- Per sequence quality scores
- Per base sequence content
- Per sequence GC content
- Per base N content
- Sequence Length Distribution
- Sequence Duplication Levels
- Overrepresented sequences
- Adapter Content
- Kmer Content

### Basic Statistics

| Measure | Value |
| --- | --- |
| Filename | outF10\_AS07.2\_pf\_trimmed\_masked\_sickled.fastq |
| File type | Conventional base calls |
| Encoding | Sanger / Illumina 1.9 |
| Total Sequences | 826637 |
| Sequences flagged as poor quality | 0 |
| Sequence length | 100-131 |
| %GC | 44 |

### Per base sequence quality

### Per tile sequence quality

### Per sequence quality scores

### Per base sequence content

### Per sequence GC content

### Per base N content

### Sequence Length Distribution

### Sequence Duplication Levels

### Overrepresented sequences

No overrepresented sequences

### Adapter Content

### Kmer Content

| Sequence | Count | PValue | Obs/Exp Max | Max Obs/Exp Position |
| --- | --- | --- | --- | --- |
| CTCGACT | 465 | 0.0 | 21.92419 | 5 |
| TCTCGAC | 475 | 0.0 | 21.381256 | 4 |
| CGACTCT | 280 | 4.712385E-5 | 16.33173 | 7 |
| CGATCTC | 750 | 9.094947E-12 | 13.425001 | 1 |
| ATCTCGA | 865 | 0.0 | 13.019789 | 3 |
| GCGTCGT | 280 | 5.995983E-4 | 12.104202 | 124-125 |
| TCGACTC | 870 | 1.0186341E-10 | 11.760583 | 6 |
| AAGAGCG | 725 | 5.456968E-12 | 11.686817 | 124-125 |
| GAGCGTC | 410 | 1.5789396E-4 | 10.03932 | 122-123 |
| CGAGTCG | 620 | 1.3309364E-7 | 9.958358 | 122-123 |
| AGTCGAG | 1470 | 0.0 | 9.510446 | 124-125 |
| AGCGTCG | 360 | 0.004863107 | 9.41438 | 124-125 |
| GGAGTCG | 620 | 2.871473E-6 | 8.82571 | 118-119 |
| GATCTCG | 1435 | 1.1823431E-10 | 8.64074 | 2 |
| TCGGAAG | 950 | 8.210918E-9 | 8.472942 | 124-125 |
| GTCGAGA | 1395 | 0.0 | 8.199622 | 124-125 |
| CGGAAGA | 880 | 2.941506E-7 | 7.9515977 | 122-123 |
| TGTAGGG | 450 | 0.009216989 | 7.518848 | 114-115 |
| GAGATCG | 1525 | 1.2732926E-11 | 7.287559 | 122-123 |
| GAGTCGA | 1915 | 0.0 | 6.8579946 | 124-125 |

Produced by FastQC (version 0.11.5)
