## Supplementary material 4 for "Natural and human-mediated drivers of microevolution in Neotropical palms: a historical genomics approach": outF11_AP04.1_pf_trimmed_masked_sickled_fastqc.html

outF11\_AP04.1\_pf\_trimmed\_masked\_sickled.fastq FastQC Report 

FastQC Report

mar. 25 juin 2019  
outF11\_AP04.1\_pf\_trimmed\_masked\_sickled.fastq

### Summary

- Basic Statistics
- Per base sequence quality
- Per tile sequence quality
- Per sequence quality scores
- Per base sequence content
- Per sequence GC content
- Per base N content
- Sequence Length Distribution
- Sequence Duplication Levels
- Overrepresented sequences
- Adapter Content
- Kmer Content

### Basic Statistics

| Measure | Value |
| --- | --- |
| Filename | outF11\_AP04.1\_pf\_trimmed\_masked\_sickled.fastq |
| File type | Conventional base calls |
| Encoding | Sanger / Illumina 1.9 |
| Total Sequences | 919858 |
| Sequences flagged as poor quality | 0 |
| Sequence length | 100-131 |
| %GC | 45 |

### Per base sequence quality

### Per tile sequence quality

### Per sequence quality scores

### Per base sequence content

### Per sequence GC content

### Per base N content

### Sequence Length Distribution

### Sequence Duplication Levels

### Overrepresented sequences

No overrepresented sequences

### Adapter Content

### Kmer Content

| Sequence | Count | PValue | Obs/Exp Max | Max Obs/Exp Position |
| --- | --- | --- | --- | --- |
| GGTACGG | 145 | 0.003972833 | 20.42156 | 7 |
| TGGTACG | 350 | 3.58541E-8 | 18.614868 | 6 |
| CACGTCT | 285 | 2.9576768E-9 | 16.596031 | 124-125 |
| CGTTTTA | 125 | 0.004218962 | 14.739431 | 52-53 |
| GGTACGT | 205 | 5.9677557E-5 | 13.028035 | 10-11 |
| CTGGTAC | 540 | 4.867261E-6 | 12.063175 | 5 |
| ACACGTC | 335 | 5.429596E-6 | 11.946856 | 124-125 |
| GCGTACC | 360 | 1.1070915E-6 | 11.9377 | 122-123 |
| CTTCCGA | 445 | 1.3642994E-4 | 11.923732 | 1 |
| TCGGAAG | 825 | 0.0 | 11.466349 | 124-125 |
| CGTACCA | 1240 | 0.0 | 11.443232 | 124-125 |
| ACGTCTG | 260 | 0.0011546063 | 11.194956 | 124-125 |
| CACACGT | 375 | 2.2627532E-5 | 10.505177 | 122-123 |
| GTACCAG | 1505 | 0.0 | 9.911813 | 124-125 |
| ATCTGGT | 1075 | 2.1369488E-8 | 9.353426 | 3 |
| CCGTACC | 445 | 0.0014372552 | 8.047888 | 122-123 |
| ACGTACC | 410 | 0.005370282 | 7.986645 | 124-125 |
| TCTGGTA | 975 | 3.7878114E-4 | 7.279995 | 4 |
| TCGTACC | 570 | 0.0020905843 | 6.9113 | 122-123 |
| TTCCGAT | 995 | 0.003732859 | 6.539555 | 2 |

Produced by FastQC (version 0.11.5)
