## Supplementary material 4 for "Natural and human-mediated drivers of microevolution in Neotropical palms: a historical genomics approach": outF11_AP04.2_pf_trimmed_masked_sickled_fastqc.html

outF11\_AP04.2\_pf\_trimmed\_masked\_sickled.fastq FastQC Report 

FastQC Report

mar. 25 juin 2019  
outF11\_AP04.2\_pf\_trimmed\_masked\_sickled.fastq

### Summary

- Basic Statistics
- Per base sequence quality
- Per tile sequence quality
- Per sequence quality scores
- Per base sequence content
- Per sequence GC content
- Per base N content
- Sequence Length Distribution
- Sequence Duplication Levels
- Overrepresented sequences
- Adapter Content
- Kmer Content

### Basic Statistics

| Measure | Value |
| --- | --- |
| Filename | outF11\_AP04.2\_pf\_trimmed\_masked\_sickled.fastq |
| File type | Conventional base calls |
| Encoding | Sanger / Illumina 1.9 |
| Total Sequences | 919858 |
| Sequences flagged as poor quality | 0 |
| Sequence length | 100-131 |
| %GC | 45 |

### Per base sequence quality

### Per tile sequence quality

### Per sequence quality scores

### Per base sequence content

### Per sequence GC content

### Per base N content

### Sequence Length Distribution

### Sequence Duplication Levels

### Overrepresented sequences

No overrepresented sequences

### Adapter Content

### Kmer Content

| Sequence | Count | PValue | Obs/Exp Max | Max Obs/Exp Position |
| --- | --- | --- | --- | --- |
| TGGTACG | 275 | 0.0 | 30.763083 | 6 |
| GGTACGG | 145 | 6.150718E-6 | 27.235004 | 7 |
| CTGGTAC | 560 | 2.8740033E-10 | 15.056159 | 5 |
| TCGGAAG | 770 | 0.0 | 13.22623 | 124-125 |
| CTTCCGA | 380 | 7.9261325E-4 | 11.708249 | 1 |
| GTACCTC | 345 | 0.004617645 | 11.446595 | 7 |
| GGTACGT | 180 | 0.0063802735 | 10.961377 | 10-11 |
| GCGCCCA | 250 | 0.00924723 | 10.426492 | 112-113 |
| GTACCAG | 1460 | 0.0 | 9.881927 | 124-125 |
| CGTACCA | 1270 | 0.0 | 9.806578 | 122-123 |
| GAGCGTC | 355 | 0.004275712 | 9.562627 | 124-125 |
| TCGTACC | 505 | 3.249078E-5 | 8.992283 | 114-115 |
| CCGTACC | 425 | 0.0018485872 | 8.986057 | 124-125 |
| ATCTGGT | 940 | 6.3239713E-7 | 8.943745 | 3 |
| GCGTACC | 380 | 0.007484768 | 8.933506 | 124-125 |
| CGGAAGA | 720 | 7.4875607E-7 | 8.840448 | 124-125 |
| TCTGGTA | 960 | 8.53086E-7 | 8.759944 | 4 |
| AAGAGCG | 575 | 5.0286057E-5 | 8.663898 | 122-123 |
| GTCCCAT | 390 | 2.7695575E-4 | 8.3524065 | 38-39 |
| GTACAAA | 610 | 0.0038223683 | 8.293228 | 5 |

Produced by FastQC (version 0.11.5)
