## Supplementary material 4 for "Natural and human-mediated drivers of microevolution in Neotropical palms: a historical genomics approach": outF11_AP12.1_pf_trimmed_masked_sickled_fastqc.html

outF11\_AP12.1\_pf\_trimmed\_masked\_sickled.fastq FastQC Report 

FastQC Report

mar. 25 juin 2019  
outF11\_AP12.1\_pf\_trimmed\_masked\_sickled.fastq

### Summary

- Basic Statistics
- Per base sequence quality
- Per tile sequence quality
- Per sequence quality scores
- Per base sequence content
- Per sequence GC content
- Per base N content
- Sequence Length Distribution
- Sequence Duplication Levels
- Overrepresented sequences
- Adapter Content
- Kmer Content

### Basic Statistics

| Measure | Value |
| --- | --- |
| Filename | outF11\_AP12.1\_pf\_trimmed\_masked\_sickled.fastq |
| File type | Conventional base calls |
| Encoding | Sanger / Illumina 1.9 |
| Total Sequences | 1030151 |
| Sequences flagged as poor quality | 0 |
| Sequence length | 100-131 |
| %GC | 44 |

### Per base sequence quality

### Per tile sequence quality

### Per sequence quality scores

### Per base sequence content

### Per sequence GC content

### Per base N content

### Sequence Length Distribution

### Sequence Duplication Levels

### Overrepresented sequences

No overrepresented sequences

### Adapter Content

### Kmer Content

| Sequence | Count | PValue | Obs/Exp Max | Max Obs/Exp Position |
| --- | --- | --- | --- | --- |
| TGTGCCT | 775 | 0.0 | 33.841198 | 6 |
| TCTGTGC | 935 | 0.0 | 29.234743 | 4 |
| CTGTGCC | 925 | 0.0 | 27.698135 | 5 |
| GATCTGT | 1125 | 0.0 | 23.665653 | 2 |
| ATCTGTG | 1135 | 0.0 | 22.481321 | 3 |
| GTGCCTG | 530 | 0.0 | 22.012165 | 7 |
| GTGCCTA | 350 | 1.9626896E-9 | 19.999624 | 7 |
| CTAAGCG | 80 | 0.007367169 | 18.353361 | 16-17 |
| GCACACG | 440 | 4.4507033E-8 | 11.957328 | 124-125 |
| CGATCTG | 2795 | 0.0 | 10.087462 | 1 |
| AGGCACA | 1740 | 0.0 | 9.71901 | 124-125 |
| CGGAAGA | 925 | 5.456968E-12 | 9.344258 | 124-125 |
| ACACGTC | 325 | 0.0056223394 | 9.250504 | 124-125 |
| TCGGAAG | 935 | 7.2759576E-12 | 9.24432 | 124-125 |
| AGCACAC | 805 | 3.623427E-9 | 8.869854 | 124-125 |
| CACGTCT | 340 | 0.008143539 | 8.842393 | 124-125 |
| GTGCCTT | 535 | 0.009101184 | 8.722577 | 7 |
| CAGGCAC | 660 | 1.2306446E-6 | 8.540948 | 124-125 |
| GGCTAGG | 435 | 3.2170015E-4 | 8.237322 | 100-101 |
| TAGGCAC | 655 | 1.0203899E-5 | 8.032403 | 124-125 |

Produced by FastQC (version 0.11.5)
