## Supplementary material 4 for "Natural and human-mediated drivers of microevolution in Neotropical palms: a historical genomics approach": outF11_AP12.2_pf_trimmed_masked_sickled_fastqc.html

outF11\_AP12.2\_pf\_trimmed\_masked\_sickled.fastq FastQC Report 

FastQC Report

mar. 25 juin 2019  
outF11\_AP12.2\_pf\_trimmed\_masked\_sickled.fastq

### Summary

- Basic Statistics
- Per base sequence quality
- Per tile sequence quality
- Per sequence quality scores
- Per base sequence content
- Per sequence GC content
- Per base N content
- Sequence Length Distribution
- Sequence Duplication Levels
- Overrepresented sequences
- Adapter Content
- Kmer Content

### Basic Statistics

| Measure | Value |
| --- | --- |
| Filename | outF11\_AP12.2\_pf\_trimmed\_masked\_sickled.fastq |
| File type | Conventional base calls |
| Encoding | Sanger / Illumina 1.9 |
| Total Sequences | 1030151 |
| Sequences flagged as poor quality | 0 |
| Sequence length | 100-131 |
| %GC | 44 |

### Per base sequence quality

### Per tile sequence quality

### Per sequence quality scores

### Per base sequence content

### Per sequence GC content

### Per base N content

### Sequence Length Distribution

### Sequence Duplication Levels

### Overrepresented sequences

No overrepresented sequences

### Adapter Content

### Kmer Content

| Sequence | Count | PValue | Obs/Exp Max | Max Obs/Exp Position |
| --- | --- | --- | --- | --- |
| TGTGCCT | 945 | 0.0 | 36.822712 | 6 |
| GTGCCTA | 305 | 0.0 | 36.11864 | 7 |
| CTGTGCC | 955 | 0.0 | 34.093258 | 5 |
| TCTGTGC | 1050 | 0.0 | 32.559746 | 4 |
| ATCTGTG | 1110 | 0.0 | 28.761467 | 3 |
| GATCTGT | 1170 | 0.0 | 27.654333 | 2 |
| GTGCCTT | 580 | 1.8189894E-12 | 16.144411 | 7 |
| TGCCTAA | 390 | 2.882498E-7 | 15.505634 | 8 |
| GTGCCTC | 615 | 7.2759576E-12 | 15.225621 | 7 |
| TGCCTAG | 295 | 1.0272167E-4 | 14.908345 | 8 |
| CGGAAGA | 775 | 0.0 | 12.27031 | 124-125 |
| AGAGCGT | 360 | 2.0890357E-5 | 12.246869 | 122-123 |
| AGCGTCG | 370 | 2.1032218E-5 | 12.238726 | 124-125 |
| TAGGCAC | 750 | 1.8189894E-12 | 12.075543 | 124-125 |
| CGATCTG | 2700 | 0.0 | 11.901105 | 1 |
| GAGCGTC | 420 | 5.887501E-6 | 11.859909 | 124-125 |
| TCGGAAG | 880 | 0.0 | 11.320822 | 124-125 |
| AGGCACA | 1700 | 0.0 | 10.921264 | 124-125 |
| GTGCCTG | 525 | 1.0109198E-4 | 10.491605 | 7 |
| GCACAGA | 1570 | 0.0 | 9.806572 | 124-125 |

Produced by FastQC (version 0.11.5)
