## Supplementary material 4 for "Natural and human-mediated drivers of microevolution in Neotropical palms: a historical genomics approach": outF11_AP14.1_pf_trimmed_masked_sickled_fastqc.html

outF11\_AP14.1\_pf\_trimmed\_masked\_sickled.fastq FastQC Report 

FastQC Report

mar. 25 juin 2019  
outF11\_AP14.1\_pf\_trimmed\_masked\_sickled.fastq

### Summary

- Basic Statistics
- Per base sequence quality
- Per tile sequence quality
- Per sequence quality scores
- Per base sequence content
- Per sequence GC content
- Per base N content
- Sequence Length Distribution
- Sequence Duplication Levels
- Overrepresented sequences
- Adapter Content
- Kmer Content

### Basic Statistics

| Measure | Value |
| --- | --- |
| Filename | outF11\_AP14.1\_pf\_trimmed\_masked\_sickled.fastq |
| File type | Conventional base calls |
| Encoding | Sanger / Illumina 1.9 |
| Total Sequences | 976111 |
| Sequences flagged as poor quality | 0 |
| Sequence length | 100-131 |
| %GC | 43 |

### Per base sequence quality

### Per tile sequence quality

### Per sequence quality scores

### Per base sequence content

### Per sequence GC content

### Per base N content

### Sequence Length Distribution

### Sequence Duplication Levels

### Overrepresented sequences

No overrepresented sequences

### Adapter Content

### Kmer Content

| Sequence | Count | PValue | Obs/Exp Max | Max Obs/Exp Position |
| --- | --- | --- | --- | --- |
| GCACACG | 365 | 2.1778942E-7 | 13.649585 | 122-123 |
| CTTGTGT | 660 | 2.401066E-10 | 13.595257 | 5 |
| CGATCTT | 730 | 9.094947E-11 | 13.004224 | 1 |
| ACACGTC | 340 | 1.7225779E-5 | 12.479352 | 124-125 |
| CACACGT | 355 | 2.6818361E-5 | 11.952055 | 124-125 |
| CGGAAGA | 935 | 0.0 | 10.89107 | 124-125 |
| TTGTGTG | 870 | 1.8371793E-8 | 10.319561 | 6 |
| TCGGAAG | 915 | 3.6379788E-12 | 10.201699 | 124-125 |
| TCTTGTG | 965 | 8.776624E-9 | 9.877394 | 4 |
| GACACAC | 365 | 0.0064346283 | 9.099723 | 122-123 |
| CGTCTGA | 370 | 0.00719511 | 8.976754 | 122-123 |
| ATCTTGT | 1015 | 2.0307925E-7 | 8.831055 | 3 |
| ACACAAG | 1890 | 0.0 | 8.78677 | 122-123 |
| CACAAGA | 2095 | 0.0 | 8.506211 | 124-125 |
| CACACAA | 2180 | 0.0 | 8.379676 | 122-123 |
| AGAGCAC | 830 | 1.2758537E-6 | 7.82574 | 120-121 |
| TCACACA | 1205 | 1.1114025E-9 | 7.7465196 | 124-125 |
| ACAAGAT | 2145 | 0.0 | 7.5167007 | 124-125 |
| AGCACAC | 830 | 4.7235335E-5 | 7.156833 | 124-125 |
| TTCACAC | 700 | 4.9134926E-4 | 7.117284 | 122-123 |

Produced by FastQC (version 0.11.5)
