## Supplementary material 4 for "Natural and human-mediated drivers of microevolution in Neotropical palms: a historical genomics approach": outF11_AP14.2_pf_trimmed_masked_sickled_fastqc.html

outF11\_AP14.2\_pf\_trimmed\_masked\_sickled.fastq FastQC Report 

FastQC Report

mar. 25 juin 2019  
outF11\_AP14.2\_pf\_trimmed\_masked\_sickled.fastq

### Summary

- Basic Statistics
- Per base sequence quality
- Per tile sequence quality
- Per sequence quality scores
- Per base sequence content
- Per sequence GC content
- Per base N content
- Sequence Length Distribution
- Sequence Duplication Levels
- Overrepresented sequences
- Adapter Content
- Kmer Content

### Basic Statistics

| Measure | Value |
| --- | --- |
| Filename | outF11\_AP14.2\_pf\_trimmed\_masked\_sickled.fastq |
| File type | Conventional base calls |
| Encoding | Sanger / Illumina 1.9 |
| Total Sequences | 976111 |
| Sequences flagged as poor quality | 0 |
| Sequence length | 100-131 |
| %GC | 43 |

### Per base sequence quality

### Per tile sequence quality

### Per sequence quality scores

### Per base sequence content

### Per sequence GC content

### Per base N content

### Sequence Length Distribution

### Sequence Duplication Levels

### Overrepresented sequences

No overrepresented sequences

### Adapter Content

### Kmer Content

| Sequence | Count | PValue | Obs/Exp Max | Max Obs/Exp Position |
| --- | --- | --- | --- | --- |
| CGATCTT | 800 | 0.0 | 21.089575 | 1 |
| CTTGTGT | 770 | 0.0 | 17.892231 | 5 |
| TTGTGTG | 1030 | 0.0 | 13.473016 | 6 |
| TCTTGTG | 1135 | 0.0 | 13.099336 | 4 |
| ATCTTGT | 1215 | 0.0 | 11.719214 | 3 |
| CGGAAGA | 935 | 0.0 | 11.002357 | 124-125 |
| GATCTTG | 1425 | 0.0 | 10.363525 | 2 |
| TGTGTGG | 535 | 8.4468233E-4 | 9.790056 | 7 |
| AGCGTCG | 390 | 0.0011176373 | 9.49588 | 124-125 |
| AGAGCGT | 425 | 2.9370622E-4 | 9.439314 | 122-123 |
| TCGGAAG | 935 | 1.9099389E-10 | 9.010255 | 122-123 |
| ACACAAG | 1915 | 0.0 | 9.008014 | 122-123 |
| AAGATCG | 1450 | 0.0 | 8.5767565 | 122-123 |
| CACACAA | 2230 | 0.0 | 8.303573 | 124-125 |
| CACAAGA | 2085 | 0.0 | 8.28897 | 124-125 |
| GAGCGTC | 465 | 0.005506228 | 7.9642878 | 124-125 |
| CCACACA | 1015 | 4.4732587E-8 | 7.702734 | 124-125 |
| AAGAGCG | 650 | 2.3219378E-4 | 7.596704 | 124-125 |
| ACAAGAT | 2215 | 0.0 | 7.430938 | 124-125 |
| GTCCACA | 600 | 0.00877486 | 6.6861806 | 122-123 |

Produced by FastQC (version 0.11.5)
