## Supplementary material 4 for "Natural and human-mediated drivers of microevolution in Neotropical palms: a historical genomics approach": outF11_AP16.1_pf_trimmed_masked_sickled_fastqc.html

outF11\_AP16.1\_pf\_trimmed\_masked\_sickled.fastq FastQC Report 

FastQC Report

mar. 25 juin 2019  
outF11\_AP16.1\_pf\_trimmed\_masked\_sickled.fastq

### Summary

- Basic Statistics
- Per base sequence quality
- Per tile sequence quality
- Per sequence quality scores
- Per base sequence content
- Per sequence GC content
- Per base N content
- Sequence Length Distribution
- Sequence Duplication Levels
- Overrepresented sequences
- Adapter Content
- Kmer Content

### Basic Statistics

| Measure | Value |
| --- | --- |
| Filename | outF11\_AP16.1\_pf\_trimmed\_masked\_sickled.fastq |
| File type | Conventional base calls |
| Encoding | Sanger / Illumina 1.9 |
| Total Sequences | 1283694 |
| Sequences flagged as poor quality | 0 |
| Sequence length | 100-131 |
| %GC | 44 |

### Per base sequence quality

### Per tile sequence quality

### Per sequence quality scores

### Per base sequence content

### Per sequence GC content

### Per base N content

### Sequence Length Distribution

### Sequence Duplication Levels

### Overrepresented sequences

No overrepresented sequences

### Adapter Content

### Kmer Content

| Sequence | Count | PValue | Obs/Exp Max | Max Obs/Exp Position |
| --- | --- | --- | --- | --- |
| TCTAACC | 1005 | 0.0 | 23.518007 | 4 |
| TAACCTT | 1090 | 0.0 | 22.811434 | 6 |
| CTAACCT | 1065 | 0.0 | 21.072514 | 5 |
| ATCTAAC | 1145 | 0.0 | 20.0964 | 3 |
| CGATCTA | 1235 | 0.0 | 18.564878 | 1 |
| CCGGTAA | 250 | 0.002862139 | 10.038965 | 62-63 |
| GATCTAA | 2340 | 0.0 | 9.8292055 | 2 |
| GCACACG | 500 | 1.0939993E-5 | 8.815866 | 124-125 |
| AACCTTG | 1275 | 2.4643668E-8 | 8.563692 | 7 |
| ACGTCTG | 395 | 8.215169E-4 | 8.516008 | 122-123 |
| CACGTCT | 480 | 7.780237E-5 | 8.347577 | 120-121 |
| GGTTAGA | 1865 | 0.0 | 7.2723174 | 124-125 |
| CACACGT | 515 | 0.0012752353 | 7.2423096 | 124-125 |
| AGGTTAG | 1785 | 0.0 | 7.028379 | 124-125 |
| CGTCTGA | 485 | 0.006161467 | 6.9357185 | 122-123 |
| AAGGTTA | 2055 | 0.0 | 6.3838987 | 122-123 |
| TAGATCG | 1800 | 1.8189894E-12 | 6.16701 | 122-123 |
| GGTTCCG | 555 | 0.0072941366 | 6.133139 | 46-47 |
| AACCTTA | 1090 | 0.0074309707 | 6.121599 | 7 |
| TCGGAAG | 1345 | 2.143679E-8 | 6.0503516 | 124-125 |

Produced by FastQC (version 0.11.5)
