## Supplementary material 4 for "Natural and human-mediated drivers of microevolution in Neotropical palms: a historical genomics approach": outF11_AP16.2_pf_trimmed_masked_sickled_fastqc.html

outF11\_AP16.2\_pf\_trimmed\_masked\_sickled.fastq FastQC Report 

FastQC Report

mar. 25 juin 2019  
outF11\_AP16.2\_pf\_trimmed\_masked\_sickled.fastq

### Summary

- Basic Statistics
- Per base sequence quality
- Per tile sequence quality
- Per sequence quality scores
- Per base sequence content
- Per sequence GC content
- Per base N content
- Sequence Length Distribution
- Sequence Duplication Levels
- Overrepresented sequences
- Adapter Content
- Kmer Content

### Basic Statistics

| Measure | Value |
| --- | --- |
| Filename | outF11\_AP16.2\_pf\_trimmed\_masked\_sickled.fastq |
| File type | Conventional base calls |
| Encoding | Sanger / Illumina 1.9 |
| Total Sequences | 1283694 |
| Sequences flagged as poor quality | 0 |
| Sequence length | 100-131 |
| %GC | 44 |

### Per base sequence quality

### Per tile sequence quality

### Per sequence quality scores

### Per base sequence content

### Per sequence GC content

### Per base N content

### Sequence Length Distribution

### Sequence Duplication Levels

### Overrepresented sequences

No overrepresented sequences

### Adapter Content

### Kmer Content

| Sequence | Count | PValue | Obs/Exp Max | Max Obs/Exp Position |
| --- | --- | --- | --- | --- |
| TCTAACC | 875 | 0.0 | 26.736822 | 4 |
| CTAACCT | 885 | 0.0 | 25.831467 | 5 |
| CGATCTA | 1080 | 0.0 | 23.078497 | 1 |
| TAACCTT | 1070 | 0.0 | 20.888033 | 6 |
| ATCTAAC | 1330 | 0.0 | 18.010138 | 3 |
| GATCTAA | 2205 | 0.0 | 11.880241 | 2 |
| AACCTTA | 925 | 2.1827873E-11 | 11.748566 | 7 |
| GGGGCGC | 190 | 0.006232208 | 10.996298 | 38-39 |
| TCGTGTA | 340 | 3.4997432E-4 | 10.77419 | 124-125 |
| GGTTAGA | 1695 | 0.0 | 10.085574 | 124-125 |
| AGGTTAG | 1700 | 0.0 | 9.577058 | 124-125 |
| AAGGTTA | 1885 | 0.0 | 8.231552 | 122-123 |
| GAGCGTC | 580 | 9.156402E-5 | 8.231552 | 122-123 |
| AGAGCGT | 500 | 0.0012839057 | 8.140498 | 124-125 |
| CTTCCGA | 720 | 0.0017962431 | 7.867669 | 1 |
| AAGAGCG | 825 | 2.4069777E-6 | 7.502886 | 120-121 |
| AGCGTCG | 520 | 0.0031112465 | 7.43976 | 120-121 |
| TCGGAAG | 1120 | 3.9646693E-8 | 7.268303 | 124-125 |
| GTTAGAT | 1755 | 1.8189894E-12 | 6.957691 | 124-125 |
| AACCTTC | 1290 | 4.2214E-5 | 6.65081 | 7 |

Produced by FastQC (version 0.11.5)
