## Supplementary material 4 for "Natural and human-mediated drivers of microevolution in Neotropical palms: a historical genomics approach": outF12_AP5.1_pf_trimmed_masked_sickled_fastqc.html

outF12\_AP5.1\_pf\_trimmed\_masked\_sickled.fastq FastQC Report 

FastQC Report

mar. 25 juin 2019  
outF12\_AP5.1\_pf\_trimmed\_masked\_sickled.fastq

### Summary

- Basic Statistics
- Per base sequence quality
- Per tile sequence quality
- Per sequence quality scores
- Per base sequence content
- Per sequence GC content
- Per base N content
- Sequence Length Distribution
- Sequence Duplication Levels
- Overrepresented sequences
- Adapter Content
- Kmer Content

### Basic Statistics

| Measure | Value |
| --- | --- |
| Filename | outF12\_AP5.1\_pf\_trimmed\_masked\_sickled.fastq |
| File type | Conventional base calls |
| Encoding | Sanger / Illumina 1.9 |
| Total Sequences | 715584 |
| Sequences flagged as poor quality | 0 |
| Sequence length | 100-131 |
| %GC | 44 |

### Per base sequence quality

### Per tile sequence quality

### Per sequence quality scores

### Per base sequence content

### Per sequence GC content

### Per base N content

### Sequence Length Distribution

### Sequence Duplication Levels

### Overrepresented sequences

No overrepresented sequences

### Adapter Content

### Kmer Content

| Sequence | Count | PValue | Obs/Exp Max | Max Obs/Exp Position |
| --- | --- | --- | --- | --- |
| ACACGTC | 270 | 1.0737749E-6 | 13.804269 | 124-125 |
| CACACGT | 310 | 5.056121E-6 | 12.023072 | 124-125 |
| ACGTCTG | 270 | 2.2602986E-4 | 11.294401 | 124-125 |
| GCACACG | 395 | 7.357481E-5 | 9.435829 | 124-125 |
| CAACTTA | 1440 | 0.0 | 8.7061 | 124-125 |
| AGCACAC | 555 | 4.429192E-6 | 8.547114 | 124-125 |
| AACTTAG | 1300 | 0.0 | 8.34048 | 124-125 |
| AGAGCAC | 730 | 2.4520277E-5 | 6.9152684 | 122-123 |
| TAGATCG | 1180 | 1.3096724E-9 | 6.891499 | 124-125 |
| ACTTAGA | 1170 | 7.934432E-9 | 6.660801 | 124-125 |
| CGGAAGA | 810 | 1.7624423E-5 | 6.5617023 | 118-119 |
| TCGGAAG | 770 | 5.1528717E-5 | 6.556034 | 122-123 |
| GAGCACA | 740 | 1.9949829E-4 | 6.410336 | 124-125 |
| CTTAGAT | 1305 | 1.3180397E-8 | 6.189298 | 122-123 |
| TGCATCC | 765 | 0.0032210457 | 5.5453105 | 112-113 |

Produced by FastQC (version 0.11.5)
