## Supplementary material 4 for "Natural and human-mediated drivers of microevolution in Neotropical palms: a historical genomics approach": outF12_AP5.2_pf_trimmed_masked_sickled_fastqc.html

outF12\_AP5.2\_pf\_trimmed\_masked\_sickled.fastq FastQC Report 

FastQC Report

mar. 25 juin 2019  
outF12\_AP5.2\_pf\_trimmed\_masked\_sickled.fastq

### Summary

- Basic Statistics
- Per base sequence quality
- Per tile sequence quality
- Per sequence quality scores
- Per base sequence content
- Per sequence GC content
- Per base N content
- Sequence Length Distribution
- Sequence Duplication Levels
- Overrepresented sequences
- Adapter Content
- Kmer Content

### Basic Statistics

| Measure | Value |
| --- | --- |
| Filename | outF12\_AP5.2\_pf\_trimmed\_masked\_sickled.fastq |
| File type | Conventional base calls |
| Encoding | Sanger / Illumina 1.9 |
| Total Sequences | 715584 |
| Sequences flagged as poor quality | 0 |
| Sequence length | 100-131 |
| %GC | 45 |

### Per base sequence quality

### Per tile sequence quality

### Per sequence quality scores

### Per base sequence content

### Per sequence GC content

### Per base N content

### Sequence Length Distribution

### Sequence Duplication Levels

### Overrepresented sequences

No overrepresented sequences

### Adapter Content

### Kmer Content

| Sequence | Count | PValue | Obs/Exp Max | Max Obs/Exp Position |
| --- | --- | --- | --- | --- |
| AGCGTCG | 305 | 8.904384E-6 | 13.304812 | 124-125 |
| GTCGTGT | 230 | 0.002615822 | 12.350338 | 124-125 |
| GCGTCGT | 260 | 0.006534974 | 10.925299 | 124-125 |
| AGAGCGT | 405 | 1.6989286E-5 | 10.780191 | 122-123 |
| CAACTTA | 1345 | 0.0 | 10.559768 | 124-125 |
| AACTTAG | 1285 | 0.0 | 10.421241 | 124-125 |
| GAGCGTC | 395 | 0.0017440342 | 9.043452 | 122-123 |
| TAGATCG | 1230 | 0.0 | 8.907734 | 124-125 |
| AAGAGCG | 520 | 2.614939E-4 | 8.3961115 | 122-123 |
| GCAACTT | 985 | 3.3600372E-8 | 7.8275523 | 124-125 |
| CGGGCGG | 435 | 6.7676604E-4 | 7.6855392 | 72-73 |
| ACTTAGA | 1185 | 1.9481377E-9 | 7.5337806 | 124-125 |
| CTTAGAT | 1250 | 1.1605152E-9 | 7.30309 | 122-123 |
| TCCGGGC | 460 | 0.0012575019 | 7.2512856 | 70-71 |
| AATGTGC | 800 | 0.004931939 | 7.0950274 | 5 |
| TCGGAAG | 735 | 1.8678048E-4 | 7.020125 | 122-123 |
| CGGAAGA | 745 | 2.2048443E-4 | 6.925895 | 122-123 |
| TCCGACC | 710 | 0.005097676 | 5.783869 | 104-105 |

Produced by FastQC (version 0.11.5)
