## Supplementary material 4 for "Natural and human-mediated drivers of microevolution in Neotropical palms: a historical genomics approach": outF12_AP9.1_pf_trimmed_masked_sickled_fastqc.html

outF12\_AP9.1\_pf\_trimmed\_masked\_sickled.fastq FastQC Report 

FastQC Report

mar. 25 juin 2019  
outF12\_AP9.1\_pf\_trimmed\_masked\_sickled.fastq

### Summary

- Basic Statistics
- Per base sequence quality
- Per tile sequence quality
- Per sequence quality scores
- Per base sequence content
- Per sequence GC content
- Per base N content
- Sequence Length Distribution
- Sequence Duplication Levels
- Overrepresented sequences
- Adapter Content
- Kmer Content

### Basic Statistics

| Measure | Value |
| --- | --- |
| Filename | outF12\_AP9.1\_pf\_trimmed\_masked\_sickled.fastq |
| File type | Conventional base calls |
| Encoding | Sanger / Illumina 1.9 |
| Total Sequences | 636436 |
| Sequences flagged as poor quality | 0 |
| Sequence length | 100-131 |
| %GC | 45 |

### Per base sequence quality

### Per tile sequence quality

### Per sequence quality scores

### Per base sequence content

### Per sequence GC content

### Per base N content

### Sequence Length Distribution

### Sequence Duplication Levels

### Overrepresented sequences

No overrepresented sequences

### Adapter Content

### Kmer Content

| Sequence | Count | PValue | Obs/Exp Max | Max Obs/Exp Position |
| --- | --- | --- | --- | --- |
| CTAATAC | 315 | 0.0 | 34.75404 | 5 |
| TAATACA | 460 | 0.0 | 25.119118 | 6 |
| TTACACG | 80 | 0.004926194 | 19.677177 | 92-93 |
| CGATCTA | 645 | 0.0 | 18.779194 | 1 |
| TCTAATA | 655 | 0.0 | 16.691084 | 4 |
| GATCTAA | 1090 | 0.0 | 12.247109 | 2 |
| ATCTAAT | 1385 | 1.2732926E-11 | 9.642365 | 3 |
| TGTATTA | 800 | 2.7284841E-11 | 9.26915 | 124-125 |
| TAGATCG | 785 | 1.7789716E-9 | 8.587516 | 124-125 |
| GTATTAG | 685 | 1.1091222E-5 | 7.3173194 | 120-121 |
| TATTAGA | 985 | 1.6603735E-8 | 7.165094 | 122-123 |
| CGGAAGA | 615 | 1.5575752E-4 | 7.124844 | 124-125 |
| TCGGAAG | 565 | 0.003602052 | 6.5622287 | 124-125 |
| TTAGATC | 1450 | 3.8868457E-6 | 5.114013 | 124-125 |

Produced by FastQC (version 0.11.5)
