## Supplementary material 4 for "Natural and human-mediated drivers of microevolution in Neotropical palms: a historical genomics approach": outF12_AP9.2_pf_trimmed_masked_sickled_fastqc.html

outF12\_AP9.2\_pf\_trimmed\_masked\_sickled.fastq FastQC Report 

FastQC Report

mar. 25 juin 2019  
outF12\_AP9.2\_pf\_trimmed\_masked\_sickled.fastq

### Summary

- Basic Statistics
- Per base sequence quality
- Per tile sequence quality
- Per sequence quality scores
- Per base sequence content
- Per sequence GC content
- Per base N content
- Sequence Length Distribution
- Sequence Duplication Levels
- Overrepresented sequences
- Adapter Content
- Kmer Content

### Basic Statistics

| Measure | Value |
| --- | --- |
| Filename | outF12\_AP9.2\_pf\_trimmed\_masked\_sickled.fastq |
| File type | Conventional base calls |
| Encoding | Sanger / Illumina 1.9 |
| Total Sequences | 636436 |
| Sequences flagged as poor quality | 0 |
| Sequence length | 100-131 |
| %GC | 46 |

### Per base sequence quality

### Per tile sequence quality

### Per sequence quality scores

### Per base sequence content

### Per sequence GC content

### Per base N content

### Sequence Length Distribution

### Sequence Duplication Levels

### Overrepresented sequences

No overrepresented sequences

### Adapter Content

### Kmer Content

| Sequence | Count | PValue | Obs/Exp Max | Max Obs/Exp Position |
| --- | --- | --- | --- | --- |
| CTAATAC | 375 | 0.0 | 33.08291 | 5 |
| TAATACA | 485 | 0.0 | 27.891117 | 6 |
| CGATCTA | 575 | 0.0 | 22.339941 | 1 |
| ACCCCCG | 80 | 0.0059384634 | 19.050953 | 50-51 |
| TCTAATA | 680 | 0.0 | 19.046705 | 4 |
| GATCTAA | 965 | 0.0 | 13.397124 | 2 |
| ATCTAAT | 1125 | 0.0 | 12.017204 | 3 |
| CGGAAGA | 480 | 6.396356E-6 | 10.312944 | 124-125 |
| AATACAT | 570 | 1.8521718E-4 | 9.880116 | 7 |
| TGTATTA | 740 | 1.7249477E-8 | 9.476758 | 124-125 |
| GTATTAG | 665 | 3.5662742E-7 | 9.304911 | 124-125 |
| AATACAA | 625 | 4.6684456E-4 | 9.010666 | 7 |
| ATACAAG | 515 | 0.008858863 | 8.750391 | 8 |
| TCGGAAG | 575 | 5.411032E-5 | 8.609065 | 124-125 |
| TAGATCG | 870 | 2.60261E-8 | 8.53485 | 124-125 |
| ACTGGGA | 360 | 0.0011974645 | 8.197582 | 28-29 |
| TATTAGA | 945 | 1.2822657E-6 | 7.2474227 | 122-123 |
| TTAGATC | 1375 | 1.2746896E-6 | 6.000258 | 124-125 |
| ATTAGAT | 1460 | 9.194817E-6 | 5.365373 | 120-121 |

Produced by FastQC (version 0.11.5)
