## Supplementary material 4 for "Natural and human-mediated drivers of microevolution in Neotropical palms: a historical genomics approach": outF12_AP13.1_pf_trimmed_masked_sickled_fastqc.html

outF12\_AP13.1\_pf\_trimmed\_masked\_sickled.fastq FastQC Report 

FastQC Report

mar. 25 juin 2019  
outF12\_AP13.1\_pf\_trimmed\_masked\_sickled.fastq

### Summary

- Basic Statistics
- Per base sequence quality
- Per tile sequence quality
- Per sequence quality scores
- Per base sequence content
- Per sequence GC content
- Per base N content
- Sequence Length Distribution
- Sequence Duplication Levels
- Overrepresented sequences
- Adapter Content
- Kmer Content

### Basic Statistics

| Measure | Value |
| --- | --- |
| Filename | outF12\_AP13.1\_pf\_trimmed\_masked\_sickled.fastq |
| File type | Conventional base calls |
| Encoding | Sanger / Illumina 1.9 |
| Total Sequences | 632041 |
| Sequences flagged as poor quality | 0 |
| Sequence length | 100-131 |
| %GC | 46 |

### Per base sequence quality

### Per tile sequence quality

### Per sequence quality scores

### Per base sequence content

### Per sequence GC content

### Per base N content

### Sequence Length Distribution

### Sequence Duplication Levels

### Overrepresented sequences

No overrepresented sequences

### Adapter Content

### Kmer Content

| Sequence | Count | PValue | Obs/Exp Max | Max Obs/Exp Position |
| --- | --- | --- | --- | --- |
| CTACAAT | 590 | 0.0 | 27.837656 | 5 |
| GATCTAC | 535 | 0.0 | 27.277437 | 2 |
| TACAATA | 660 | 0.0 | 26.739288 | 6 |
| CGATCTA | 600 | 0.0 | 25.268703 | 1 |
| ATCTACA | 685 | 0.0 | 23.961525 | 3 |
| TCTACAA | 780 | 0.0 | 23.39632 | 4 |
| TCTATAC | 295 | 0.0073903804 | 8.94734 | 118-119 |
| ACAATAA | 695 | 6.2191807E-4 | 8.755401 | 7 |
| TATTGTA | 1190 | 0.0 | 8.744983 | 124-125 |
| GTATTGT | 525 | 2.4375575E-5 | 8.271274 | 122-123 |
| ATTGTAG | 1125 | 1.8189894E-12 | 8.056667 | 124-125 |
| GTAGATC | 890 | 2.384695E-9 | 7.920881 | 124-125 |
| ACAATAT | 695 | 0.006050279 | 7.8798604 | 7 |
| GCCGTGG | 360 | 0.008281105 | 7.609625 | 10-11 |
| TGTAGAT | 1080 | 2.3283064E-10 | 7.4598775 | 124-125 |
| GGAGAGC | 465 | 0.0049359994 | 7.0953374 | 118-119 |
| TTGTAGA | 1160 | 8.047209E-9 | 6.656011 | 124-125 |
| TCGGAAG | 610 | 0.0011524702 | 6.6038256 | 124-125 |
| TAGATCG | 1030 | 1.990822E-6 | 6.1924224 | 124-125 |

Produced by FastQC (version 0.11.5)
