## Supplementary material 4 for "Natural and human-mediated drivers of microevolution in Neotropical palms: a historical genomics approach": outF12_AP13.2_pf_trimmed_masked_sickled_fastqc.html

outF12\_AP13.2\_pf\_trimmed\_masked\_sickled.fastq FastQC Report 

FastQC Report

mar. 25 juin 2019  
outF12\_AP13.2\_pf\_trimmed\_masked\_sickled.fastq

### Summary

- Basic Statistics
- Per base sequence quality
- Per tile sequence quality
- Per sequence quality scores
- Per base sequence content
- Per sequence GC content
- Per base N content
- Sequence Length Distribution
- Sequence Duplication Levels
- Overrepresented sequences
- Adapter Content
- Kmer Content

### Basic Statistics

| Measure | Value |
| --- | --- |
| Filename | outF12\_AP13.2\_pf\_trimmed\_masked\_sickled.fastq |
| File type | Conventional base calls |
| Encoding | Sanger / Illumina 1.9 |
| Total Sequences | 632041 |
| Sequences flagged as poor quality | 0 |
| Sequence length | 100-131 |
| %GC | 46 |

### Per base sequence quality

### Per tile sequence quality

### Per sequence quality scores

### Per base sequence content

### Per sequence GC content

### Per base N content

### Sequence Length Distribution

### Sequence Duplication Levels

### Overrepresented sequences

No overrepresented sequences

### Adapter Content

### Kmer Content

| Sequence | Count | PValue | Obs/Exp Max | Max Obs/Exp Position |
| --- | --- | --- | --- | --- |
| GATCTAC | 520 | 0.0 | 24.010088 | 2 |
| CTACAAT | 615 | 0.0 | 22.205503 | 5 |
| TACAATA | 600 | 0.0 | 21.814087 | 6 |
| TCTACAA | 735 | 0.0 | 20.095154 | 4 |
| CGATCTA | 690 | 0.0 | 19.655052 | 1 |
| ATCTACA | 695 | 0.0 | 18.805813 | 3 |
| ACAATAC | 270 | 6.8472006E-4 | 14.751045 | 7 |
| CGTATTG | 170 | 0.0061869407 | 13.911982 | 122-123 |
| AGCGTCG | 220 | 0.00233029 | 12.5418625 | 122-123 |
| GTAGATC | 865 | 3.459354E-8 | 9.841234 | 2 |
| TAGATCG | 940 | 4.9940354E-8 | 9.672555 | 3 |
| TATTGTA | 1090 | 0.0 | 9.250946 | 124-125 |
| ATTGTAG | 975 | 4.0017767E-11 | 9.101033 | 124-125 |
| TGTAGAT | 1095 | 4.4201443E-10 | 8.10366 | 124-125 |
| AAGAGCG | 430 | 0.0049229264 | 8.063594 | 120-121 |
| CGGAAGA | 615 | 1.5405852E-4 | 7.870074 | 124-125 |
| TTGTAGA | 1055 | 1.5481419E-8 | 7.64628 | 124-125 |
| AGTGGGA | 445 | 0.009419282 | 6.637076 | 28-29 |
| ATCGGAA | 1385 | 1.706976E-5 | 6.5740147 | 6 |
| TCAAGAG | 635 | 0.005240123 | 5.769514 | 74-75 |

Produced by FastQC (version 0.11.5)
