## Supplementary material 4 for "Natural and human-mediated drivers of microevolution in Neotropical palms: a historical genomics approach": outF12_AP15.1_pf_trimmed_masked_sickled_fastqc.html

outF12\_AP15.1\_pf\_trimmed\_masked\_sickled.fastq FastQC Report 

FastQC Report

mar. 25 juin 2019  
outF12\_AP15.1\_pf\_trimmed\_masked\_sickled.fastq

### Summary

- Basic Statistics
- Per base sequence quality
- Per tile sequence quality
- Per sequence quality scores
- Per base sequence content
- Per sequence GC content
- Per base N content
- Sequence Length Distribution
- Sequence Duplication Levels
- Overrepresented sequences
- Adapter Content
- Kmer Content

### Basic Statistics

| Measure | Value |
| --- | --- |
| Filename | outF12\_AP15.1\_pf\_trimmed\_masked\_sickled.fastq |
| File type | Conventional base calls |
| Encoding | Sanger / Illumina 1.9 |
| Total Sequences | 505234 |
| Sequences flagged as poor quality | 0 |
| Sequence length | 100-131 |
| %GC | 44 |

### Per base sequence quality

### Per tile sequence quality

### Per sequence quality scores

### Per base sequence content

### Per sequence GC content

### Per base N content

### Sequence Length Distribution

### Sequence Duplication Levels

### Overrepresented sequences

No overrepresented sequences

### Adapter Content

### Kmer Content

| Sequence | Count | PValue | Obs/Exp Max | Max Obs/Exp Position |
| --- | --- | --- | --- | --- |
| CGATCTA | 375 | 3.434252E-9 | 19.114378 | 1 |
| TACCACA | 315 | 4.2929933E-6 | 17.204544 | 6 |
| GATCTAC | 415 | 2.2064887E-7 | 15.870921 | 2 |
| CTACCAC | 345 | 1.0172913E-5 | 15.711677 | 5 |
| TCGGAAG | 490 | 3.8198777E-11 | 12.447925 | 120-121 |
| TCTACCA | 515 | 3.42119E-5 | 11.667619 | 4 |
| ATCTACC | 465 | 1.7570643E-4 | 11.601871 | 3 |
| CGGAAGA | 505 | 8.1672624E-10 | 11.407173 | 120-121 |
| TCGGACA | 205 | 0.009123139 | 10.445091 | 40-41 |
| AGAGCAC | 580 | 6.840855E-8 | 9.47901 | 124-125 |
| GTGGTAG | 800 | 1.6016202E-8 | 8.160835 | 124-125 |
| GTAGATC | 775 | 1.0269105E-7 | 7.8703012 | 120-121 |
| CTGTGGT | 485 | 5.82535E-4 | 7.7933097 | 124-125 |
| AAGGTGC | 365 | 0.007660275 | 7.6758842 | 58-59 |
| ATAATAG | 370 | 0.008966931 | 7.5417824 | 76-77 |
| TGTGGTA | 805 | 1.2575201E-6 | 7.256447 | 124-125 |
| TAGATCG | 800 | 1.8177416E-6 | 7.0889797 | 116-117 |
| TCCACTG | 500 | 0.008349492 | 6.720195 | 118-119 |
| GGTAGAT | 855 | 4.228818E-5 | 6.1804333 | 114-115 |
| GAAGAGC | 885 | 4.2810167E-5 | 6.175155 | 122-123 |

Produced by FastQC (version 0.11.5)
