## Supplementary material 4 for "Natural and human-mediated drivers of microevolution in Neotropical palms: a historical genomics approach": outF12_AP15.2_pf_trimmed_masked_sickled_fastqc.html

outF12\_AP15.2\_pf\_trimmed\_masked\_sickled.fastq FastQC Report 

FastQC Report

mar. 25 juin 2019  
outF12\_AP15.2\_pf\_trimmed\_masked\_sickled.fastq

### Summary

- Basic Statistics
- Per base sequence quality
- Per tile sequence quality
- Per sequence quality scores
- Per base sequence content
- Per sequence GC content
- Per base N content
- Sequence Length Distribution
- Sequence Duplication Levels
- Overrepresented sequences
- Adapter Content
- Kmer Content

### Basic Statistics

| Measure | Value |
| --- | --- |
| Filename | outF12\_AP15.2\_pf\_trimmed\_masked\_sickled.fastq |
| File type | Conventional base calls |
| Encoding | Sanger / Illumina 1.9 |
| Total Sequences | 505234 |
| Sequences flagged as poor quality | 0 |
| Sequence length | 100-131 |
| %GC | 44 |

### Per base sequence quality

### Per tile sequence quality

### Per sequence quality scores

### Per base sequence content

### Per sequence GC content

### Per base N content

### Sequence Length Distribution

### Sequence Duplication Levels

### Overrepresented sequences

No overrepresented sequences

### Adapter Content

### Kmer Content

| Sequence | Count | PValue | Obs/Exp Max | Max Obs/Exp Position |
| --- | --- | --- | --- | --- |
| CACTACG | 65 | 0.0025169072 | 34.715073 | 8 |
| AGACACG | 80 | 0.0074398327 | 27.817598 | 1 |
| GCGACTA | 45 | 0.008942312 | 26.790693 | 70-71 |
| AGAGCGT | 275 | 2.1883734E-6 | 15.233225 | 124-125 |
| AAGAGCG | 385 | 1.4842954E-9 | 15.233224 | 124-125 |
| AGGAGTG | 295 | 9.576966E-5 | 12.383553 | 122-123 |
| TCGGAAG | 485 | 6.9630914E-9 | 12.165808 | 120-121 |
| CGGAAGA | 445 | 3.6650454E-7 | 11.491448 | 120-121 |
| TGTGGTA | 735 | 1.1985321E-8 | 9.68916 | 124-125 |
| GAAGAGC | 780 | 4.970043E-7 | 8.326262 | 122-123 |
| GTGGTAG | 755 | 1.7756374E-6 | 8.322788 | 124-125 |
| TAGATCG | 730 | 5.641559E-7 | 8.257406 | 116-117 |
| GGTAGAT | 730 | 6.914288E-7 | 8.147476 | 114-115 |
| GTAGATC | 730 | 6.033908E-6 | 7.6382585 | 114-115 |
| GGAAATC | 365 | 0.00991237 | 7.4573736 | 72-73 |
| TGGTAGA | 865 | 5.3269323E-6 | 7.112372 | 118-119 |
| ATCGGAA | 910 | 0.0012954354 | 5.984481 | 124-125 |
| GGAAGAG | 1095 | 4.8923964E-4 | 5.5603466 | 122-123 |

Produced by FastQC (version 0.11.5)
