## Supplementary material 4 for "Natural and human-mediated drivers of microevolution in Neotropical palms: a historical genomics approach": outF12_AP19.1_pf_trimmed_masked_sickled_fastqc.html

outF12\_AP19.1\_pf\_trimmed\_masked\_sickled.fastq FastQC Report 

FastQC Report

mar. 25 juin 2019  
outF12\_AP19.1\_pf\_trimmed\_masked\_sickled.fastq

### Summary

- Basic Statistics
- Per base sequence quality
- Per tile sequence quality
- Per sequence quality scores
- Per base sequence content
- Per sequence GC content
- Per base N content
- Sequence Length Distribution
- Sequence Duplication Levels
- Overrepresented sequences
- Adapter Content
- Kmer Content

### Basic Statistics

| Measure | Value |
| --- | --- |
| Filename | outF12\_AP19.1\_pf\_trimmed\_masked\_sickled.fastq |
| File type | Conventional base calls |
| Encoding | Sanger / Illumina 1.9 |
| Total Sequences | 708862 |
| Sequences flagged as poor quality | 0 |
| Sequence length | 100-131 |
| %GC | 44 |

### Per base sequence quality

### Per tile sequence quality

### Per sequence quality scores

### Per base sequence content

### Per sequence GC content

### Per base N content

### Sequence Length Distribution

### Sequence Duplication Levels

### Overrepresented sequences

No overrepresented sequences

### Adapter Content

### Kmer Content

| Sequence | Count | PValue | Obs/Exp Max | Max Obs/Exp Position |
| --- | --- | --- | --- | --- |
| TACTCAC | 455 | 0.0 | 45.018066 | 6 |
| CTACTCA | 590 | 0.0 | 34.73483 | 5 |
| TCTACTC | 675 | 0.0 | 29.465725 | 4 |
| GATCTAC | 710 | 0.0 | 28.820549 | 2 |
| CGATCTA | 775 | 0.0 | 27.858282 | 1 |
| ATCTACT | 780 | 0.0 | 25.502861 | 3 |
| ACTCACG | 145 | 1.2236618E-4 | 24.950428 | 7 |
| ACTCACA | 455 | 3.6379788E-12 | 19.878088 | 7 |
| GCACACG | 245 | 5.2612013E-6 | 14.000703 | 124-125 |
| CACACGT | 240 | 6.720964E-5 | 12.863147 | 124-125 |
| ACTCACC | 395 | 5.560878E-4 | 12.212023 | 7 |
| CAAGCGA | 225 | 0.008775865 | 10.500301 | 120-121 |
| ACTCACT | 405 | 0.009271569 | 10.42168 | 7 |
| TGAGTAG | 945 | 9.404175E-10 | 7.810733 | 118-119 |
| CGGAAGA | 620 | 1.8804545E-5 | 7.672256 | 122-123 |
| GTGAGTA | 890 | 1.5945261E-8 | 7.635311 | 122-123 |
| GCCATTA | 495 | 2.2853362E-4 | 7.607244 | 88-89 |
| GAGTAGA | 1010 | 2.3082976E-9 | 7.471662 | 124-125 |
| TAGATCG | 1140 | 7.4578566E-11 | 7.4511266 | 122-123 |
| TCGGAAG | 705 | 1.3157898E-5 | 7.229178 | 122-123 |

Produced by FastQC (version 0.11.5)
