## Supplementary material 4 for "Natural and human-mediated drivers of microevolution in Neotropical palms: a historical genomics approach": outF12_AP19.2_pf_trimmed_masked_sickled_fastqc.html

outF12\_AP19.2\_pf\_trimmed\_masked\_sickled.fastq FastQC Report 

FastQC Report

mar. 25 juin 2019  
outF12\_AP19.2\_pf\_trimmed\_masked\_sickled.fastq

### Summary

- Basic Statistics
- Per base sequence quality
- Per tile sequence quality
- Per sequence quality scores
- Per base sequence content
- Per sequence GC content
- Per base N content
- Sequence Length Distribution
- Sequence Duplication Levels
- Overrepresented sequences
- Adapter Content
- Kmer Content

### Basic Statistics

| Measure | Value |
| --- | --- |
| Filename | outF12\_AP19.2\_pf\_trimmed\_masked\_sickled.fastq |
| File type | Conventional base calls |
| Encoding | Sanger / Illumina 1.9 |
| Total Sequences | 708862 |
| Sequences flagged as poor quality | 0 |
| Sequence length | 100-131 |
| %GC | 44 |

### Per base sequence quality

### Per tile sequence quality

### Per sequence quality scores

### Per base sequence content

### Per sequence GC content

### Per base N content

### Sequence Length Distribution

### Sequence Duplication Levels

### Overrepresented sequences

No overrepresented sequences

### Adapter Content

### Kmer Content

| Sequence | Count | PValue | Obs/Exp Max | Max Obs/Exp Position |
| --- | --- | --- | --- | --- |
| TACTCAC | 415 | 0.0 | 51.744957 | 6 |
| CTACTCA | 575 | 0.0 | 38.22029 | 5 |
| TCTACTC | 555 | 0.0 | 36.426376 | 4 |
| GATCTAC | 630 | 0.0 | 33.77455 | 2 |
| ACTCACG | 180 | 1.0786607E-9 | 31.303453 | 7 |
| ATCTACT | 700 | 0.0 | 29.634592 | 3 |
| CGATCTA | 790 | 0.0 | 29.589088 | 1 |
| ACTCACA | 375 | 7.0976967E-9 | 18.030788 | 7 |
| GCGTCGT | 205 | 5.0196548E-5 | 16.212263 | 124-125 |
| ACTCACT | 290 | 7.190047E-5 | 15.543782 | 7 |
| GAGCGTC | 265 | 2.1324613E-6 | 15.272949 | 122-123 |
| ACTCACC | 390 | 3.792953E-6 | 14.447746 | 7 |
| AAGAGCG | 395 | 5.63854E-7 | 12.620938 | 124-125 |
| AGCGTCG | 240 | 0.0030176842 | 12.116977 | 124-125 |
| AGAGCGT | 305 | 0.0017973271 | 10.6159525 | 122-123 |
| CGGAAGA | 585 | 8.107236E-7 | 9.685922 | 122-123 |
| GTGAGTA | 840 | 2.0008883E-10 | 9.636504 | 122-123 |
| AGTTGTC | 350 | 0.0010312144 | 9.579397 | 114-115 |
| GGTGAGT | 385 | 0.0014301857 | 9.242465 | 120-121 |
| TCGGAAG | 670 | 5.239999E-7 | 9.061191 | 122-123 |

Produced by FastQC (version 0.11.5)
