## Supplementary material 4 for "Natural and human-mediated drivers of microevolution in Neotropical palms: a historical genomics approach": outF12_AS1.1_pf_trimmed_masked_sickled_fastqc.html

outF12\_AS1.1\_pf\_trimmed\_masked\_sickled.fastq FastQC Report 

FastQC Report

mar. 25 juin 2019  
outF12\_AS1.1\_pf\_trimmed\_masked\_sickled.fastq

### Summary

- Basic Statistics
- Per base sequence quality
- Per tile sequence quality
- Per sequence quality scores
- Per base sequence content
- Per sequence GC content
- Per base N content
- Sequence Length Distribution
- Sequence Duplication Levels
- Overrepresented sequences
- Adapter Content
- Kmer Content

### Basic Statistics

| Measure | Value |
| --- | --- |
| Filename | outF12\_AS1.1\_pf\_trimmed\_masked\_sickled.fastq |
| File type | Conventional base calls |
| Encoding | Sanger / Illumina 1.9 |
| Total Sequences | 1840844 |
| Sequences flagged as poor quality | 0 |
| Sequence length | 100-131 |
| %GC | 45 |

### Per base sequence quality

### Per tile sequence quality

### Per sequence quality scores

### Per base sequence content

### Per sequence GC content

### Per base N content

### Sequence Length Distribution

### Sequence Duplication Levels

### Overrepresented sequences

No overrepresented sequences

### Adapter Content

### Kmer Content

| Sequence | Count | PValue | Obs/Exp Max | Max Obs/Exp Position |
| --- | --- | --- | --- | --- |
| CTATGTG | 1420 | 0.0 | 26.786062 | 5 |
| TATGTGT | 1640 | 0.0 | 22.457771 | 6 |
| CGATCTA | 2065 | 0.0 | 18.919884 | 1 |
| TCTATGT | 2060 | 0.0 | 18.161541 | 4 |
| ATCTATG | 2730 | 0.0 | 14.138172 | 3 |
| ATGTGTA | 1150 | 0.0 | 13.130802 | 7 |
| GATCTAT | 3170 | 0.0 | 12.927467 | 2 |
| TTCCGAT | 2360 | 0.0 | 9.168866 | 1 |
| ATGTGTC | 895 | 1.1622811E-5 | 8.773429 | 7 |
| CACATAG | 2680 | 0.0 | 8.280673 | 124-125 |
| GCACACG | 720 | 2.3713073E-7 | 8.061271 | 124-125 |
| ACATAGA | 2905 | 0.0 | 7.874369 | 124-125 |
| ACACATA | 3530 | 0.0 | 7.8342495 | 124-125 |
| CACACGT | 755 | 5.0599374E-7 | 7.6875696 | 124-125 |
| TGTGTAA | 905 | 0.0011015378 | 7.3426585 | 8 |
| GACGGGC | 470 | 0.0057163476 | 6.9897676 | 112-113 |
| ACACGTC | 590 | 6.5200357E-4 | 6.9441056 | 124-125 |
| TCGGAAG | 1860 | 0.0 | 6.920179 | 122-123 |
| TAGATCG | 2765 | 0.0 | 6.173933 | 124-125 |
| CGGAAGA | 1680 | 3.074092E-10 | 5.8935337 | 124-125 |

Produced by FastQC (version 0.11.5)
