## Supplementary material 4 for "Natural and human-mediated drivers of microevolution in Neotropical palms: a historical genomics approach": outF12_AS1.2_pf_trimmed_masked_sickled_fastqc.html

outF12\_AS1.2\_pf\_trimmed\_masked\_sickled.fastq FastQC Report 

FastQC Report

mar. 25 juin 2019  
outF12\_AS1.2\_pf\_trimmed\_masked\_sickled.fastq

### Summary

- Basic Statistics
- Per base sequence quality
- Per tile sequence quality
- Per sequence quality scores
- Per base sequence content
- Per sequence GC content
- Per base N content
- Sequence Length Distribution
- Sequence Duplication Levels
- Overrepresented sequences
- Adapter Content
- Kmer Content

### Basic Statistics

| Measure | Value |
| --- | --- |
| Filename | outF12\_AS1.2\_pf\_trimmed\_masked\_sickled.fastq |
| File type | Conventional base calls |
| Encoding | Sanger / Illumina 1.9 |
| Total Sequences | 1840844 |
| Sequences flagged as poor quality | 0 |
| Sequence length | 100-131 |
| %GC | 45 |

### Per base sequence quality

### Per tile sequence quality

### Per sequence quality scores

### Per base sequence content

### Per sequence GC content

### Per base N content

### Sequence Length Distribution

### Sequence Duplication Levels

### Overrepresented sequences

No overrepresented sequences

### Adapter Content

### Kmer Content

| Sequence | Count | PValue | Obs/Exp Max | Max Obs/Exp Position |
| --- | --- | --- | --- | --- |
| CTATGTG | 1465 | 0.0 | 27.364452 | 5 |
| TCTATGT | 1755 | 0.0 | 22.477934 | 4 |
| TATGTGT | 1920 | 0.0 | 21.174858 | 6 |
| CGATCTA | 2085 | 0.0 | 19.030428 | 1 |
| CTTCCGA | 1070 | 0.0 | 17.481823 | 1 |
| ATCTATG | 2520 | 0.0 | 16.332062 | 3 |
| GATCTAT | 3045 | 0.0 | 14.429771 | 2 |
| AGAGCGT | 810 | 0.0 | 11.60324 | 124-125 |
| TGGTCGC | 520 | 6.281432E-5 | 10.996233 | 7 |
| ATGTGTA | 1270 | 0.0 | 10.805746 | 7 |
| AGCGTCG | 630 | 2.6026464E-7 | 9.511712 | 122-123 |
| ACACATA | 3485 | 0.0 | 9.380451 | 124-125 |
| CACATAG | 2770 | 0.0 | 9.293884 | 124-125 |
| ATGTGTT | 1545 | 0.0 | 9.252493 | 7 |
| GCCGTAT | 285 | 0.001814418 | 9.005557 | 12-13 |
| GGTCGCC | 585 | 0.0022550987 | 8.792153 | 8 |
| GAGCGTC | 800 | 7.165727E-8 | 8.683512 | 124-125 |
| GCGTCGT | 530 | 2.3489945E-4 | 8.481122 | 124-125 |
| TTCCGAT | 2155 | 0.0 | 8.47342 | 2 |
| ATGTGTC | 950 | 5.367372E-6 | 8.426586 | 7 |

Produced by FastQC (version 0.11.5)
