## Supplementary material 4 for "Natural and human-mediated drivers of microevolution in Neotropical palms: a historical genomics approach": outF12_AS2.1_pf_trimmed_masked_sickled_fastqc.html

outF12\_AS2.1\_pf\_trimmed\_masked\_sickled.fastq FastQC Report 

FastQC Report

mar. 25 juin 2019  
outF12\_AS2.1\_pf\_trimmed\_masked\_sickled.fastq

### Summary

- Basic Statistics
- Per base sequence quality
- Per tile sequence quality
- Per sequence quality scores
- Per base sequence content
- Per sequence GC content
- Per base N content
- Sequence Length Distribution
- Sequence Duplication Levels
- Overrepresented sequences
- Adapter Content
- Kmer Content

### Basic Statistics

| Measure | Value |
| --- | --- |
| Filename | outF12\_AS2.1\_pf\_trimmed\_masked\_sickled.fastq |
| File type | Conventional base calls |
| Encoding | Sanger / Illumina 1.9 |
| Total Sequences | 1625860 |
| Sequences flagged as poor quality | 0 |
| Sequence length | 100-131 |
| %GC | 44 |

### Per base sequence quality

### Per tile sequence quality

### Per sequence quality scores

### Per base sequence content

### Per sequence GC content

### Per base N content

### Sequence Length Distribution

### Sequence Duplication Levels

### Overrepresented sequences

No overrepresented sequences

### Adapter Content

### Kmer Content

| Sequence | Count | PValue | Obs/Exp Max | Max Obs/Exp Position |
| --- | --- | --- | --- | --- |
| TCACTTA | 1125 | 0.0 | 15.079595 | 6 |
| CTCACTT | 1180 | 0.0 | 14.381682 | 5 |
| TCTCACT | 1770 | 0.0 | 11.635032 | 4 |
| CGATCTC | 1835 | 0.0 | 9.875982 | 1 |
| GAGATCG | 2420 | 0.0 | 7.8256674 | 124-125 |
| TAAGTGA | 2180 | 0.0 | 7.7564373 | 124-125 |
| AAGTGAG | 2405 | 0.0 | 7.481655 | 120-121 |
| GCACACG | 570 | 4.901393E-4 | 7.1195927 | 124-125 |
| ATCTCAC | 3290 | 0.0 | 6.992278 | 3 |
| GTAAGTG | 825 | 2.3903194E-6 | 6.968571 | 124-125 |
| CACTTAT | 1010 | 0.003404669 | 6.5980363 | 7 |
| CTAAGTG | 975 | 5.357195E-6 | 6.200774 | 122-123 |
| AGTGAGA | 2675 | 0.0 | 6.194711 | 124-125 |
| GATCTCA | 3035 | 7.8216544E-11 | 5.983092 | 2 |
| CGGAAGA | 1240 | 7.812792E-6 | 5.417343 | 122-123 |
| GTGAGAT | 3605 | 0.0 | 5.2174797 | 122-123 |
| TCGGAAG | 1310 | 2.0325035E-5 | 5.1278667 | 122-123 |

Produced by FastQC (version 0.11.5)
