## Supplementary material 4 for "Natural and human-mediated drivers of microevolution in Neotropical palms: a historical genomics approach": outF12_AS2.2_pf_trimmed_masked_sickled_fastqc.html

outF12\_AS2.2\_pf\_trimmed\_masked\_sickled.fastq FastQC Report 

FastQC Report

mar. 25 juin 2019  
outF12\_AS2.2\_pf\_trimmed\_masked\_sickled.fastq

### Summary

- Basic Statistics
- Per base sequence quality
- Per tile sequence quality
- Per sequence quality scores
- Per base sequence content
- Per sequence GC content
- Per base N content
- Sequence Length Distribution
- Sequence Duplication Levels
- Overrepresented sequences
- Adapter Content
- Kmer Content

### Basic Statistics

| Measure | Value |
| --- | --- |
| Filename | outF12\_AS2.2\_pf\_trimmed\_masked\_sickled.fastq |
| File type | Conventional base calls |
| Encoding | Sanger / Illumina 1.9 |
| Total Sequences | 1625860 |
| Sequences flagged as poor quality | 0 |
| Sequence length | 100-131 |
| %GC | 45 |

### Per base sequence quality

### Per tile sequence quality

### Per sequence quality scores

### Per base sequence content

### Per sequence GC content

### Per base N content

### Sequence Length Distribution

### Sequence Duplication Levels

### Overrepresented sequences

No overrepresented sequences

### Adapter Content

### Kmer Content

| Sequence | Count | PValue | Obs/Exp Max | Max Obs/Exp Position |
| --- | --- | --- | --- | --- |
| TCACTTA | 830 | 0.0 | 17.941336 | 6 |
| CTCACTT | 1130 | 0.0 | 13.186223 | 5 |
| TAACGTC | 170 | 0.0026906976 | 12.305359 | 38-39 |
| CGGGGTA | 195 | 0.0064314865 | 10.950011 | 50-51 |
| TCTCACT | 1650 | 0.0 | 9.3785 | 4 |
| GTCGTGT | 395 | 0.0016852723 | 9.078614 | 124-125 |
| CGATCTC | 1630 | 1.8189894E-12 | 8.737368 | 1 |
| GAGATCG | 2400 | 0.0 | 7.969006 | 124-125 |
| AAGTGAG | 2415 | 0.0 | 7.75452 | 124-125 |
| TAAGTGA | 2315 | 0.0 | 7.5731373 | 124-125 |
| CTAAGTG | 845 | 2.2160002E-6 | 7.5446215 | 124-125 |
| AGAGCGT | 595 | 0.0010647965 | 7.366308 | 124-125 |
| CACTTAT | 895 | 0.0017313071 | 7.035893 | 7 |
| AAGAGCG | 850 | 1.945027E-5 | 7.031476 | 124-125 |
| CTTCCGA | 820 | 0.0060603307 | 6.947273 | 1 |
| GAGCGTC | 635 | 0.002120734 | 6.9022884 | 124-125 |
| GATCTCA | 2880 | 0.0 | 6.7637243 | 2 |
| CGGAAGA | 1340 | 4.9545633E-8 | 6.4093633 | 122-123 |
| AGTGAGA | 2630 | 0.0 | 6.3638697 | 120-121 |
| ATCTCAC | 3145 | 1.364242E-10 | 5.654405 | 3 |

Produced by FastQC (version 0.11.5)
