## Supplementary material 4 for "Natural and human-mediated drivers of microevolution in Neotropical palms: a historical genomics approach": outF12_AS5.1_pf_trimmed_masked_sickled_fastqc.html

outF12\_AS5.1\_pf\_trimmed\_masked\_sickled.fastq FastQC Report 

FastQC Report

mar. 25 juin 2019  
outF12\_AS5.1\_pf\_trimmed\_masked\_sickled.fastq

### Summary

- Basic Statistics
- Per base sequence quality
- Per tile sequence quality
- Per sequence quality scores
- Per base sequence content
- Per sequence GC content
- Per base N content
- Sequence Length Distribution
- Sequence Duplication Levels
- Overrepresented sequences
- Adapter Content
- Kmer Content

### Basic Statistics

| Measure | Value |
| --- | --- |
| Filename | outF12\_AS5.1\_pf\_trimmed\_masked\_sickled.fastq |
| File type | Conventional base calls |
| Encoding | Sanger / Illumina 1.9 |
| Total Sequences | 1491754 |
| Sequences flagged as poor quality | 0 |
| Sequence length | 100-131 |
| %GC | 45 |

### Per base sequence quality

### Per tile sequence quality

### Per sequence quality scores

### Per base sequence content

### Per sequence GC content

### Per base N content

### Sequence Length Distribution

### Sequence Duplication Levels

### Overrepresented sequences

No overrepresented sequences

### Adapter Content

### Kmer Content

| Sequence | Count | PValue | Obs/Exp Max | Max Obs/Exp Position |
| --- | --- | --- | --- | --- |
| TCCATAG | 1085 | 0.0 | 14.502642 | 6 |
| CTCCATA | 1495 | 0.0 | 11.325675 | 5 |
| CGATCTC | 1785 | 0.0 | 9.442422 | 1 |
| ACGTCTG | 325 | 0.0012151151 | 9.41018 | 124-125 |
| GCACACG | 470 | 5.405524E-4 | 7.8493724 | 120-121 |
| CACGTCT | 480 | 5.884821E-4 | 7.787359 | 124-125 |
| CTATGGA | 2295 | 0.0 | 7.403312 | 124-125 |
| CGGAAGA | 1195 | 1.8444553E-9 | 6.78456 | 122-123 |
| CCATAGG | 1010 | 0.0034134847 | 6.5963182 | 7 |
| GATCTCC | 2845 | 1.03682396E-10 | 6.151186 | 2 |
| ATCTCCA | 3090 | 2.1827873E-11 | 6.059219 | 3 |
| TCTCCAT | 3005 | 3.947207E-10 | 5.8342104 | 4 |
| TATGGAG | 2595 | 0.0 | 5.630797 | 124-125 |
| GGGACTT | 685 | 0.009965856 | 5.443746 | 78-79 |
| TCGGAAG | 1240 | 8.014449E-6 | 5.4093914 | 120-121 |
| ACTATGG | 1020 | 4.2980324E-4 | 5.2608986 | 120-121 |
| GAGATCG | 2145 | 2.3101165E-10 | 5.1971474 | 122-123 |

Produced by FastQC (version 0.11.5)
