## Supplementary material 4 for "Natural and human-mediated drivers of microevolution in Neotropical palms: a historical genomics approach": outF12_AS5.2_pf_trimmed_masked_sickled_fastqc.html

outF12\_AS5.2\_pf\_trimmed\_masked\_sickled.fastq FastQC Report 

FastQC Report

mar. 25 juin 2019  
outF12\_AS5.2\_pf\_trimmed\_masked\_sickled.fastq

### Summary

- Basic Statistics
- Per base sequence quality
- Per tile sequence quality
- Per sequence quality scores
- Per base sequence content
- Per sequence GC content
- Per base N content
- Sequence Length Distribution
- Sequence Duplication Levels
- Overrepresented sequences
- Adapter Content
- Kmer Content

### Basic Statistics

| Measure | Value |
| --- | --- |
| Filename | outF12\_AS5.2\_pf\_trimmed\_masked\_sickled.fastq |
| File type | Conventional base calls |
| Encoding | Sanger / Illumina 1.9 |
| Total Sequences | 1491754 |
| Sequences flagged as poor quality | 0 |
| Sequence length | 100-131 |
| %GC | 45 |

### Per base sequence quality

### Per tile sequence quality

### Per sequence quality scores

### Per base sequence content

### Per sequence GC content

### Per base N content

### Sequence Length Distribution

### Sequence Duplication Levels

### Overrepresented sequences

No overrepresented sequences

### Adapter Content

### Kmer Content

| Sequence | Count | PValue | Obs/Exp Max | Max Obs/Exp Position |
| --- | --- | --- | --- | --- |
| ATACGCG | 75 | 0.0033210383 | 21.064829 | 90-91 |
| TCCATAG | 1215 | 1.8189894E-12 | 10.368815 | 6 |
| AAGAGCG | 765 | 5.1562893E-8 | 8.861742 | 124-125 |
| CTATGGA | 2215 | 0.0 | 8.821734 | 124-125 |
| CTCCATA | 1435 | 8.240022E-10 | 8.364154 | 5 |
| TATGGAG | 2470 | 0.0 | 8.233886 | 124-125 |
| AGAGCGT | 535 | 3.3825994E-4 | 8.1991825 | 124-125 |
| CGGAAGA | 1105 | 6.330447E-8 | 7.0856814 | 122-123 |
| CGATCTC | 1520 | 1.9706931E-7 | 7.0794463 | 1 |
| GGAGATC | 2635 | 0.0 | 6.507579 | 124-125 |
| GAGATCG | 2040 | 2.1827873E-11 | 6.0598674 | 124-125 |
| ACTATGG | 1040 | 3.1093523E-4 | 5.7516117 | 124-125 |
| TCGGAAG | 1165 | 5.1622043E-5 | 5.712641 | 122-123 |
| GCTATGG | 1175 | 5.8781265E-5 | 5.6640224 | 122-123 |
| TGGAGAT | 3365 | 0.0 | 5.214339 | 124-125 |
| TCTCCAT | 2965 | 3.98868E-8 | 5.192585 | 4 |
| TCTATGG | 1375 | 1.1270043E-4 | 5.12488 | 122-123 |
| ATGGAGA | 3640 | 0.0 | 5.054875 | 122-123 |

Produced by FastQC (version 0.11.5)
