## Supplementary material 4 for "Natural and human-mediated drivers of microevolution in Neotropical palms: a historical genomics approach": outF12_AS10.1_pf_trimmed_masked_sickled_fastqc.html

outF12\_AS10.1\_pf\_trimmed\_masked\_sickled.fastq FastQC Report 

FastQC Report

mar. 25 juin 2019  
outF12\_AS10.1\_pf\_trimmed\_masked\_sickled.fastq

### Summary

- Basic Statistics
- Per base sequence quality
- Per tile sequence quality
- Per sequence quality scores
- Per base sequence content
- Per sequence GC content
- Per base N content
- Sequence Length Distribution
- Sequence Duplication Levels
- Overrepresented sequences
- Adapter Content
- Kmer Content

### Basic Statistics

| Measure | Value |
| --- | --- |
| Filename | outF12\_AS10.1\_pf\_trimmed\_masked\_sickled.fastq |
| File type | Conventional base calls |
| Encoding | Sanger / Illumina 1.9 |
| Total Sequences | 1319928 |
| Sequences flagged as poor quality | 0 |
| Sequence length | 100-131 |
| %GC | 43 |

### Per base sequence quality

### Per tile sequence quality

### Per sequence quality scores

### Per base sequence content

### Per sequence GC content

### Per base N content

### Sequence Length Distribution

### Sequence Duplication Levels

### Overrepresented sequences

No overrepresented sequences

### Adapter Content

### Kmer Content

| Sequence | Count | PValue | Obs/Exp Max | Max Obs/Exp Position |
| --- | --- | --- | --- | --- |
| CTGAGGC | 960 | 0.0 | 12.94504 | 5 |
| TCTGAGG | 1085 | 0.0 | 11.987384 | 4 |
| TGAGGCA | 1205 | 0.0 | 11.296139 | 6 |
| GCACACG | 555 | 1.8873834E-8 | 10.303841 | 124-125 |
| TCGGAAG | 1140 | 0.0 | 8.46508 | 124-125 |
| GCCTCAG | 2160 | 0.0 | 8.346815 | 122-123 |
| CGTCTGA | 565 | 2.6242986E-5 | 8.223697 | 124-125 |
| CCTCAGA | 2280 | 0.0 | 7.994798 | 124-125 |
| CACACGT | 550 | 1.7172439E-4 | 7.7981343 | 124-125 |
| TGCCTCA | 2750 | 0.0 | 7.668165 | 124-125 |
| AATGTAG | 695 | 0.00797813 | 7.641465 | 3 |
| CACGTCT | 575 | 3.2562544E-4 | 7.377645 | 122-123 |
| ATCTGAG | 1990 | 6.511982E-10 | 7.116675 | 3 |
| AGAGCAC | 1315 | 5.4424163E-9 | 6.451933 | 122-123 |
| CTCAGAT | 3270 | 0.0 | 6.4487634 | 124-125 |
| GAGGCAG | 1300 | 2.130629E-4 | 6.3779244 | 7 |
| CGGAAGA | 1175 | 1.0016302E-6 | 6.083651 | 124-125 |
| AGCACAC | 1105 | 1.4834892E-5 | 5.8221364 | 124-125 |
| GAGCACA | 1200 | 9.2039845E-6 | 5.659063 | 124-125 |
| TTGCCTC | 1635 | 6.6454413E-7 | 5.133006 | 120-121 |

Produced by FastQC (version 0.11.5)
