## Supplementary material 4 for "Natural and human-mediated drivers of microevolution in Neotropical palms: a historical genomics approach": outF12_AS10.2_pf_trimmed_masked_sickled_fastqc.html

outF12\_AS10.2\_pf\_trimmed\_masked\_sickled.fastq FastQC Report 

FastQC Report

mar. 25 juin 2019  
outF12\_AS10.2\_pf\_trimmed\_masked\_sickled.fastq

### Summary

- Basic Statistics
- Per base sequence quality
- Per tile sequence quality
- Per sequence quality scores
- Per base sequence content
- Per sequence GC content
- Per base N content
- Sequence Length Distribution
- Sequence Duplication Levels
- Overrepresented sequences
- Adapter Content
- Kmer Content

### Basic Statistics

| Measure | Value |
| --- | --- |
| Filename | outF12\_AS10.2\_pf\_trimmed\_masked\_sickled.fastq |
| File type | Conventional base calls |
| Encoding | Sanger / Illumina 1.9 |
| Total Sequences | 1319928 |
| Sequences flagged as poor quality | 0 |
| Sequence length | 100-131 |
| %GC | 44 |

### Per base sequence quality

### Per tile sequence quality

### Per sequence quality scores

### Per base sequence content

### Per sequence GC content

### Per base N content

### Sequence Length Distribution

### Sequence Duplication Levels

### Overrepresented sequences

No overrepresented sequences

### Adapter Content

### Kmer Content

| Sequence | Count | PValue | Obs/Exp Max | Max Obs/Exp Position |
| --- | --- | --- | --- | --- |
| CTGAGGC | 800 | 0.0 | 16.074877 | 5 |
| TCTGAGG | 1010 | 0.0 | 13.811267 | 4 |
| TCGTGTA | 320 | 2.6820105E-4 | 11.08848 | 118-119 |
| CGTGTAG | 300 | 0.0016088631 | 10.759 | 120-121 |
| TGAGGCA | 1260 | 0.0 | 10.210401 | 6 |
| GCCTCAG | 2175 | 0.0 | 9.808311 | 124-125 |
| CCTCAGA | 2305 | 0.0 | 9.625338 | 124-125 |
| GTGTAGG | 590 | 9.838373E-6 | 8.889852 | 120-121 |
| AGAGCGT | 535 | 1.628371E-4 | 8.772481 | 124-125 |
| TGCCTCA | 2630 | 0.0 | 8.556476 | 122-123 |
| AAGAGCG | 775 | 1.9948093E-6 | 8.257966 | 124-125 |
| AGCGTCG | 460 | 0.0011773299 | 8.212224 | 114-115 |
| TCGGAAG | 1255 | 2.764864E-10 | 7.8192954 | 124-125 |
| CGTCGTG | 445 | 0.006873875 | 7.7700844 | 116-117 |
| GTCAGGG | 385 | 0.002182295 | 7.714103 | 40-41 |
| CTCAGAT | 3075 | 0.0 | 7.353842 | 124-125 |
| ATCTGAG | 1870 | 5.9117156E-10 | 7.1484833 | 3 |
| CGGAAGA | 1220 | 8.017014E-8 | 6.9944515 | 124-125 |
| CTGCGCA | 930 | 0.0034578464 | 6.588006 | 3 |
| ATGCCTC | 1580 | 8.172719E-9 | 6.3301067 | 122-123 |

Produced by FastQC (version 0.11.5)
