## Supplementary material 4 for "Natural and human-mediated drivers of microevolution in Neotropical palms: a historical genomics approach": outF12_AS13.1_pf_trimmed_masked_sickled_fastqc.html

outF12\_AS13.1\_pf\_trimmed\_masked\_sickled.fastq FastQC Report 

FastQC Report

mar. 25 juin 2019  
outF12\_AS13.1\_pf\_trimmed\_masked\_sickled.fastq

### Summary

- Basic Statistics
- Per base sequence quality
- Per tile sequence quality
- Per sequence quality scores
- Per base sequence content
- Per sequence GC content
- Per base N content
- Sequence Length Distribution
- Sequence Duplication Levels
- Overrepresented sequences
- Adapter Content
- Kmer Content

### Basic Statistics

| Measure | Value |
| --- | --- |
| Filename | outF12\_AS13.1\_pf\_trimmed\_masked\_sickled.fastq |
| File type | Conventional base calls |
| Encoding | Sanger / Illumina 1.9 |
| Total Sequences | 1208561 |
| Sequences flagged as poor quality | 0 |
| Sequence length | 100-131 |
| %GC | 44 |

### Per base sequence quality

### Per tile sequence quality

### Per sequence quality scores

### Per base sequence content

### Per sequence GC content

### Per base N content

### Sequence Length Distribution

### Sequence Duplication Levels

### Overrepresented sequences

No overrepresented sequences

### Adapter Content

### Kmer Content

| Sequence | Count | PValue | Obs/Exp Max | Max Obs/Exp Position |
| --- | --- | --- | --- | --- |
| GGTACGC | 195 | 3.9916653E-5 | 21.391706 | 7 |
| TGGTACG | 380 | 1.9099389E-10 | 20.376043 | 6 |
| CTGGTAC | 640 | 1.8841456E-7 | 12.098276 | 5 |
| CGTTATT | 180 | 0.0033368526 | 11.956617 | 70-71 |
| ACACGTC | 490 | 4.080175E-7 | 10.186045 | 124-125 |
| ACGTCTG | 350 | 1.5591587E-4 | 10.052851 | 122-123 |
| CGTACCA | 1630 | 0.0 | 9.066068 | 122-123 |
| CACGTCT | 435 | 1.2107495E-4 | 9.015236 | 124-125 |
| TCGTACC | 715 | 1.05701474E-7 | 8.4765 | 124-125 |
| GCACACG | 600 | 7.6195465E-6 | 8.209827 | 122-123 |
| ACGTACC | 520 | 1.0741978E-4 | 8.11961 | 122-123 |
| ATCGTAC | 455 | 0.0024277384 | 7.6306877 | 120-121 |
| GTACCAG | 1870 | 0.0 | 7.426658 | 120-121 |
| GAGTCGG | 595 | 0.0050444994 | 6.3545785 | 118-119 |
| TCGGAAG | 1300 | 2.4476321E-8 | 6.3075128 | 124-125 |
| TACCAGA | 2460 | 0.0 | 6.2317066 | 124-125 |
| ATCTGGT | 1375 | 4.192209E-4 | 6.0520005 | 3 |
| TCTGGTA | 1295 | 0.0013206941 | 5.976018 | 4 |
| CGGAAGA | 1155 | 5.036449E-6 | 5.8646927 | 124-125 |
| ACCAGAT | 2425 | 0.0 | 5.803708 | 122-123 |

Produced by FastQC (version 0.11.5)
