## Supplementary material 4 for "Natural and human-mediated drivers of microevolution in Neotropical palms: a historical genomics approach": outF12_AS13.2_pf_trimmed_masked_sickled_fastqc.html

outF12\_AS13.2\_pf\_trimmed\_masked\_sickled.fastq FastQC Report 

FastQC Report

mar. 25 juin 2019  
outF12\_AS13.2\_pf\_trimmed\_masked\_sickled.fastq

### Summary

- Basic Statistics
- Per base sequence quality
- Per tile sequence quality
- Per sequence quality scores
- Per base sequence content
- Per sequence GC content
- Per base N content
- Sequence Length Distribution
- Sequence Duplication Levels
- Overrepresented sequences
- Adapter Content
- Kmer Content

### Basic Statistics

| Measure | Value |
| --- | --- |
| Filename | outF12\_AS13.2\_pf\_trimmed\_masked\_sickled.fastq |
| File type | Conventional base calls |
| Encoding | Sanger / Illumina 1.9 |
| Total Sequences | 1208561 |
| Sequences flagged as poor quality | 0 |
| Sequence length | 100-131 |
| %GC | 44 |

### Per base sequence quality

### Per tile sequence quality

### Per sequence quality scores

### Per base sequence content

### Per sequence GC content

### Per base N content

### Sequence Length Distribution

### Sequence Duplication Levels

### Overrepresented sequences

No overrepresented sequences

### Adapter Content

### Kmer Content

| Sequence | Count | PValue | Obs/Exp Max | Max Obs/Exp Position |
| --- | --- | --- | --- | --- |
| CGCGACT | 100 | 6.289163E-4 | 28.030136 | 2 |
| TGGTACG | 325 | 0.0 | 27.781353 | 6 |
| GGTACGG | 140 | 0.0042648893 | 20.173443 | 7 |
| GTACGGA | 145 | 0.0052488176 | 19.46235 | 8 |
| CTGGTAC | 670 | 2.8740033E-10 | 13.442262 | 5 |
| GGCGTAC | 235 | 0.003347304 | 11.951263 | 120-121 |
| TCGTACC | 685 | 5.820766E-11 | 11.1287565 | 120-121 |
| CGTACCA | 1635 | 0.0 | 10.323881 | 122-123 |
| ACGTACC | 570 | 3.4086042E-7 | 10.319888 | 124-125 |
| GCGTACC | 430 | 2.54737E-4 | 9.574331 | 122-123 |
| GAGCGTC | 425 | 2.934329E-4 | 9.440494 | 120-121 |
| GTACCAG | 1845 | 0.0 | 9.371947 | 122-123 |
| AGCGTCG | 340 | 0.008131399 | 8.844106 | 114-115 |
| AAGAGCG | 625 | 2.9419334E-6 | 8.810656 | 118-119 |
| ATCGTAC | 405 | 0.002994177 | 8.521368 | 116-117 |
| CGGAAGA | 960 | 1.1497832E-8 | 8.315802 | 124-125 |
| TACCAGA | 2620 | 0.0 | 8.018452 | 124-125 |
| TCGGAAG | 1150 | 4.438334E-9 | 7.672612 | 124-125 |
| ACCAGAT | 2365 | 0.0 | 7.6393976 | 124-125 |
| ATCTGGT | 1190 | 2.2278746E-6 | 7.540817 | 3 |

Produced by FastQC (version 0.11.5)
