## Supplementary material 4 for "Natural and human-mediated drivers of microevolution in Neotropical palms: a historical genomics approach": outF12_AS15.1_pf_trimmed_masked_sickled_fastqc.html

outF12\_AS15.1\_pf\_trimmed\_masked\_sickled.fastq FastQC Report 

FastQC Report

mar. 25 juin 2019  
outF12\_AS15.1\_pf\_trimmed\_masked\_sickled.fastq

### Summary

- Basic Statistics
- Per base sequence quality
- Per tile sequence quality
- Per sequence quality scores
- Per base sequence content
- Per sequence GC content
- Per base N content
- Sequence Length Distribution
- Sequence Duplication Levels
- Overrepresented sequences
- Adapter Content
- Kmer Content

### Basic Statistics

| Measure | Value |
| --- | --- |
| Filename | outF12\_AS15.1\_pf\_trimmed\_masked\_sickled.fastq |
| File type | Conventional base calls |
| Encoding | Sanger / Illumina 1.9 |
| Total Sequences | 922250 |
| Sequences flagged as poor quality | 0 |
| Sequence length | 100-131 |
| %GC | 44 |

### Per base sequence quality

### Per tile sequence quality

### Per sequence quality scores

### Per base sequence content

### Per sequence GC content

### Per base N content

### Sequence Length Distribution

### Sequence Duplication Levels

### Overrepresented sequences

No overrepresented sequences

### Adapter Content

### Kmer Content

| Sequence | Count | PValue | Obs/Exp Max | Max Obs/Exp Position |
| --- | --- | --- | --- | --- |
| TGTGCCT | 720 | 0.0 | 20.195162 | 6 |
| CTGTGCC | 790 | 0.0 | 17.656528 | 5 |
| TCTGTGC | 840 | 0.0 | 17.258495 | 4 |
| ATCTGTG | 940 | 0.0 | 16.644106 | 3 |
| GATCTGT | 1135 | 0.0 | 14.784831 | 2 |
| GTGCCTA | 365 | 2.4016554E-5 | 14.350243 | 7 |
| ACGTCTG | 320 | 1.5703154E-7 | 14.020702 | 124-125 |
| CACACGT | 440 | 2.401066E-10 | 13.595834 | 124-125 |
| GTGCCTT | 450 | 1.1923292E-5 | 12.932936 | 7 |
| GCACACG | 435 | 4.0749E-8 | 12.033094 | 124-125 |
| TCGTACT | 210 | 0.0033649988 | 11.942714 | 118-119 |
| ACACGTC | 380 | 1.2642759E-6 | 11.806908 | 124-125 |
| CACGTCT | 345 | 8.36791E-5 | 10.68983 | 122-123 |
| CGTCTGA | 350 | 9.980294E-4 | 9.614197 | 124-125 |
| CGGAAGA | 880 | 2.3101165E-10 | 8.922266 | 124-125 |
| CTTCCGA | 520 | 0.008039751 | 8.8555155 | 1 |
| AGCACAC | 730 | 8.543611E-8 | 8.588471 | 122-123 |
| CCTAGTC | 410 | 0.0054145027 | 7.979327 | 120-121 |
| TCGGAAG | 960 | 1.1019438E-8 | 7.78928 | 124-125 |
| AGAGCAC | 1000 | 4.8985385E-9 | 7.6335554 | 120-121 |

Produced by FastQC (version 0.11.5)
