## Supplementary material 4 for "Natural and human-mediated drivers of microevolution in Neotropical palms: a historical genomics approach": outF12_AS15.2_pf_trimmed_masked_sickled_fastqc.html

outF12\_AS15.2\_pf\_trimmed\_masked\_sickled.fastq FastQC Report 

FastQC Report

mar. 25 juin 2019  
outF12\_AS15.2\_pf\_trimmed\_masked\_sickled.fastq

### Summary

- Basic Statistics
- Per base sequence quality
- Per tile sequence quality
- Per sequence quality scores
- Per base sequence content
- Per sequence GC content
- Per base N content
- Sequence Length Distribution
- Sequence Duplication Levels
- Overrepresented sequences
- Adapter Content
- Kmer Content

### Basic Statistics

| Measure | Value |
| --- | --- |
| Filename | outF12\_AS15.2\_pf\_trimmed\_masked\_sickled.fastq |
| File type | Conventional base calls |
| Encoding | Sanger / Illumina 1.9 |
| Total Sequences | 922250 |
| Sequences flagged as poor quality | 0 |
| Sequence length | 100-131 |
| %GC | 44 |

### Per base sequence quality

### Per tile sequence quality

### Per sequence quality scores

### Per base sequence content

### Per sequence GC content

### Per base N content

### Sequence Length Distribution

### Sequence Duplication Levels

### Overrepresented sequences

No overrepresented sequences

### Adapter Content

### Kmer Content

| Sequence | Count | PValue | Obs/Exp Max | Max Obs/Exp Position |
| --- | --- | --- | --- | --- |
| TGTGCCT | 650 | 0.0 | 31.356117 | 6 |
| CTGTGCC | 750 | 0.0 | 27.138502 | 5 |
| TCTGTGC | 720 | 0.0 | 26.642384 | 4 |
| GTGCCTA | 345 | 0.0 | 23.90634 | 7 |
| ATCTGTG | 855 | 0.0 | 22.384485 | 3 |
| GATCTGT | 1090 | 0.0 | 17.516441 | 2 |
| AGCGTCG | 310 | 1.3496901E-8 | 17.120375 | 122-123 |
| CGTGTAG | 170 | 0.0023347985 | 16.107635 | 124-125 |
| AGAGCGT | 440 | 2.924935E-9 | 14.521278 | 124-125 |
| AAGAGCG | 650 | 3.8380676E-10 | 11.936172 | 124-125 |
| GAGCGTC | 350 | 2.1190717E-4 | 11.37282 | 122-123 |
| GTGCCTT | 440 | 2.3488418E-4 | 11.246846 | 7 |
| CGTCGTG | 285 | 0.0053989314 | 11.209409 | 124-125 |
| TCGGAAG | 885 | 5.456968E-12 | 10.829428 | 124-125 |
| GTGCCTC | 460 | 3.5452424E-4 | 10.757853 | 7 |
| CGGAAGA | 850 | 2.8740033E-10 | 10.201503 | 124-125 |
| TGCCTAA | 460 | 0.0042824415 | 9.5602865 | 8 |
| TAGGCAC | 610 | 4.6776167E-6 | 9.425561 | 122-123 |
| CGATCTG | 2400 | 0.0 | 9.238259 | 1 |
| ACAGATC | 1600 | 0.0 | 8.292681 | 122-123 |

Produced by FastQC (version 0.11.5)
