## Supplementary material 4 for "Natural and human-mediated drivers of microevolution in Neotropical palms: a historical genomics approach": outF12_AS16.1_pf_trimmed_masked_sickled_fastqc.html

outF12\_AS16.1\_pf\_trimmed\_masked\_sickled.fastq FastQC Report 

FastQC Report

mar. 25 juin 2019  
outF12\_AS16.1\_pf\_trimmed\_masked\_sickled.fastq

### Summary

- Basic Statistics
- Per base sequence quality
- Per tile sequence quality
- Per sequence quality scores
- Per base sequence content
- Per sequence GC content
- Per base N content
- Sequence Length Distribution
- Sequence Duplication Levels
- Overrepresented sequences
- Adapter Content
- Kmer Content

### Basic Statistics

| Measure | Value |
| --- | --- |
| Filename | outF12\_AS16.1\_pf\_trimmed\_masked\_sickled.fastq |
| File type | Conventional base calls |
| Encoding | Sanger / Illumina 1.9 |
| Total Sequences | 749604 |
| Sequences flagged as poor quality | 0 |
| Sequence length | 100-131 |
| %GC | 43 |

### Per base sequence quality

### Per tile sequence quality

### Per sequence quality scores

### Per base sequence content

### Per sequence GC content

### Per base N content

### Sequence Length Distribution

### Sequence Duplication Levels

### Overrepresented sequences

No overrepresented sequences

### Adapter Content

### Kmer Content

| Sequence | Count | PValue | Obs/Exp Max | Max Obs/Exp Position |
| --- | --- | --- | --- | --- |
| CGATCTT | 495 | 0.0 | 24.225914 | 1 |
| CTTGTGT | 575 | 0.0 | 19.916254 | 5 |
| TTGTGTG | 735 | 0.0 | 16.362005 | 6 |
| TCTTGTG | 750 | 0.0 | 16.027138 | 4 |
| CGTCAGG | 135 | 0.008521859 | 13.255522 | 48-49 |
| ATCTTGT | 940 | 0.0 | 12.792825 | 3 |
| GCACACG | 225 | 0.0044108313 | 11.51762 | 118-119 |
| GATCTTG | 1135 | 7.2759576E-12 | 10.592058 | 2 |
| TCACAGG | 380 | 0.0043187193 | 8.18223 | 106-107 |
| ACACAAG | 1065 | 4.12183E-9 | 7.701031 | 124-125 |
| AAGATCG | 870 | 7.279523E-7 | 7.512 | 122-123 |
| CCACACA | 610 | 3.1027902E-4 | 7.407842 | 120-121 |
| CACAAGA | 1250 | 1.6298145E-9 | 7.186162 | 124-125 |
| AGCACAC | 590 | 0.0021187358 | 6.902206 | 118-119 |
| CACACAA | 1285 | 4.3948603E-8 | 6.447033 | 120-121 |
| ACAAGAT | 1345 | 5.264883E-8 | 6.388216 | 124-125 |
| CTAAAAA | 1185 | 0.0049139373 | 5.802347 | 8 |
| CAAGATC | 1685 | 5.529135E-6 | 5.019364 | 122-123 |

Produced by FastQC (version 0.11.5)
