## Supplementary material 4 for "Natural and human-mediated drivers of microevolution in Neotropical palms: a historical genomics approach": outF12_AS16.2_pf_trimmed_masked_sickled_fastqc.html

outF12\_AS16.2\_pf\_trimmed\_masked\_sickled.fastq FastQC Report 

FastQC Report

mar. 25 juin 2019  
outF12\_AS16.2\_pf\_trimmed\_masked\_sickled.fastq

### Summary

- Basic Statistics
- Per base sequence quality
- Per tile sequence quality
- Per sequence quality scores
- Per base sequence content
- Per sequence GC content
- Per base N content
- Sequence Length Distribution
- Sequence Duplication Levels
- Overrepresented sequences
- Adapter Content
- Kmer Content

### Basic Statistics

| Measure | Value |
| --- | --- |
| Filename | outF12\_AS16.2\_pf\_trimmed\_masked\_sickled.fastq |
| File type | Conventional base calls |
| Encoding | Sanger / Illumina 1.9 |
| Total Sequences | 749604 |
| Sequences flagged as poor quality | 0 |
| Sequence length | 100-131 |
| %GC | 43 |

### Per base sequence quality

### Per tile sequence quality

### Per sequence quality scores

### Per base sequence content

### Per sequence GC content

### Per base N content

### Sequence Length Distribution

### Sequence Duplication Levels

### Overrepresented sequences

No overrepresented sequences

### Adapter Content

### Kmer Content

| Sequence | Count | PValue | Obs/Exp Max | Max Obs/Exp Position |
| --- | --- | --- | --- | --- |
| CGATCTT | 520 | 0.0 | 18.341625 | 1 |
| CTTGTGT | 545 | 3.9535735E-8 | 13.632356 | 5 |
| AGCGTCG | 185 | 0.008810023 | 13.188369 | 122-123 |
| ATCTTGT | 795 | 4.5487468E-8 | 10.714224 | 3 |
| AGAGCGT | 275 | 0.007953778 | 10.64007 | 124-125 |
| TCTTGTG | 865 | 1.3327735E-8 | 10.533186 | 4 |
| GATCTTG | 1070 | 3.6561687E-10 | 10.063853 | 2 |
| TTGTGTG | 740 | 2.1103322E-5 | 9.325759 | 6 |
| GTGGACT | 260 | 0.007516665 | 8.929095 | 26-27 |
| AAGAGCG | 460 | 0.0043355497 | 8.178313 | 124-125 |
| AAGATCG | 965 | 1.2028795E-7 | 7.7969413 | 124-125 |
| CGGAAGA | 530 | 0.0022984275 | 7.672478 | 122-123 |
| ACACAAG | 1150 | 4.8894435E-9 | 7.6330924 | 124-125 |
| CACACAA | 1300 | 1.5897967E-9 | 7.1944237 | 122-123 |
| CACAAGA | 1300 | 2.7193892E-9 | 7.0121355 | 120-121 |
| ACAAGAT | 1385 | 1.1670449E-5 | 5.5784736 | 122-123 |
| CAAGATC | 1615 | 2.7664542E-5 | 5.035806 | 122-123 |

Produced by FastQC (version 0.11.5)
