## Supplementary material 4 for "Natural and human-mediated drivers of microevolution in Neotropical palms: a historical genomics approach": outLAU_AP01.1_pf_trimmed_masked_sickled_fastqc.html

outLAU\_AP01.1\_pf\_trimmed\_masked\_sickled.fastq FastQC Report 

FastQC Report

mar. 25 juin 2019  
outLAU\_AP01.1\_pf\_trimmed\_masked\_sickled.fastq

### Summary

- Basic Statistics
- Per base sequence quality
- Per tile sequence quality
- Per sequence quality scores
- Per base sequence content
- Per sequence GC content
- Per base N content
- Sequence Length Distribution
- Sequence Duplication Levels
- Overrepresented sequences
- Adapter Content
- Kmer Content

### Basic Statistics

| Measure | Value |
| --- | --- |
| Filename | outLAU\_AP01.1\_pf\_trimmed\_masked\_sickled.fastq |
| File type | Conventional base calls |
| Encoding | Sanger / Illumina 1.9 |
| Total Sequences | 1019968 |
| Sequences flagged as poor quality | 0 |
| Sequence length | 100-131 |
| %GC | 43 |

### Per base sequence quality

### Per tile sequence quality

### Per sequence quality scores

### Per base sequence content

### Per sequence GC content

### Per base N content

### Sequence Length Distribution

### Sequence Duplication Levels

### Overrepresented sequences

No overrepresented sequences

### Adapter Content

### Kmer Content

| Sequence | Count | PValue | Obs/Exp Max | Max Obs/Exp Position |
| --- | --- | --- | --- | --- |
| ACGTCTG | 315 | 2.8923278E-6 | 12.639501 | 124-125 |
| CACGTCT | 330 | 5.5967266E-6 | 11.914646 | 122-123 |
| ACACGTC | 350 | 1.077169E-5 | 11.233809 | 122-123 |
| CACACGT | 360 | 1.7388627E-5 | 10.7587185 | 120-121 |
| CGTCTGA | 340 | 8.729626E-5 | 10.645569 | 124-125 |
| GCACACG | 410 | 4.650583E-6 | 10.593639 | 124-125 |
| CTTCCGA | 490 | 0.004185906 | 9.587023 | 1 |
| AGCACAC | 765 | 2.4529982E-7 | 8.043319 | 124-125 |
| ATCTGCC | 745 | 0.0016583035 | 7.9314127 | 7 |
| CGGAAGA | 830 | 1.0773874E-7 | 7.849504 | 124-125 |
| TCGGAAG | 895 | 4.6775313E-8 | 7.683841 | 124-125 |
| AGAGCAC | 945 | 1.5164551E-7 | 7.1866117 | 122-123 |
| GCAGATC | 1845 | 0.0 | 6.393225 | 122-123 |
| GGCAGAT | 1495 | 1.6079866E-9 | 6.2163377 | 122-123 |
| AGGCAGA | 1515 | 3.079549E-9 | 6.042701 | 120-121 |
| CAAGGCA | 1965 | 1.6370905E-11 | 5.8943405 | 124-125 |
| GAGCACA | 1020 | 1.4473767E-4 | 5.677637 | 124-125 |
| CCCATTC | 765 | 0.0016837886 | 5.426475 | 10-11 |
| GAAGAGC | 1535 | 9.01553E-7 | 5.275818 | 120-121 |
| AAGGCAG | 1835 | 1.6359081E-7 | 5.064536 | 122-123 |

Produced by FastQC (version 0.11.5)
