## Supplementary material 4 for "Natural and human-mediated drivers of microevolution in Neotropical palms: a historical genomics approach": outLAU_AP01.2_pf_trimmed_masked_sickled_fastqc.html

outLAU\_AP01.2\_pf\_trimmed\_masked\_sickled.fastq FastQC Report 

FastQC Report

mar. 25 juin 2019  
outLAU\_AP01.2\_pf\_trimmed\_masked\_sickled.fastq

### Summary

- Basic Statistics
- Per base sequence quality
- Per tile sequence quality
- Per sequence quality scores
- Per base sequence content
- Per sequence GC content
- Per base N content
- Sequence Length Distribution
- Sequence Duplication Levels
- Overrepresented sequences
- Adapter Content
- Kmer Content

### Basic Statistics

| Measure | Value |
| --- | --- |
| Filename | outLAU\_AP01.2\_pf\_trimmed\_masked\_sickled.fastq |
| File type | Conventional base calls |
| Encoding | Sanger / Illumina 1.9 |
| Total Sequences | 1019968 |
| Sequences flagged as poor quality | 0 |
| Sequence length | 100-131 |
| %GC | 43 |

### Per base sequence quality

### Per tile sequence quality

### Per sequence quality scores

### Per base sequence content

### Per sequence GC content

### Per base N content

### Sequence Length Distribution

### Sequence Duplication Levels

### Overrepresented sequences

No overrepresented sequences

### Adapter Content

### Kmer Content

| Sequence | Count | PValue | Obs/Exp Max | Max Obs/Exp Position |
| --- | --- | --- | --- | --- |
| AGCGTCG | 355 | 2.8376235E-10 | 17.117392 | 124-125 |
| GTACGGA | 115 | 0.0027048318 | 15.757281 | 76-77 |
| AGAGCGT | 370 | 1.31331035E-8 | 14.827073 | 122-123 |
| GCGTCGT | 255 | 2.218811E-4 | 13.617197 | 124-125 |
| GAGCGTC | 365 | 1.9670788E-6 | 13.080903 | 124-125 |
| AAGAGCG | 570 | 2.1282176E-9 | 11.845651 | 122-123 |
| CGTCGTG | 265 | 0.007029658 | 10.819042 | 120-121 |
| CCCGTCG | 190 | 0.009985764 | 10.319255 | 16-17 |
| CGGAAGA | 860 | 3.810783E-9 | 8.844602 | 118-119 |
| TCGGAAG | 865 | 9.52532E-8 | 8.530427 | 124-125 |
| CTGCCTT | 855 | 1.6086534E-4 | 7.841036 | 9 |
| AGGCAGA | 1435 | 3.6379788E-12 | 7.7063465 | 120-121 |
| CAAGGCA | 1785 | 0.0 | 7.5380917 | 124-125 |
| GGCAGAT | 1460 | 2.0008883E-11 | 7.515092 | 122-123 |
| AAGGCAG | 1570 | 3.6379788E-12 | 7.3045764 | 120-121 |
| TCTGCCT | 845 | 0.0012329994 | 7.264288 | 8 |
| GATCTGC | 1040 | 0.0016249465 | 6.405886 | 2 |
| GAAGAGC | 1490 | 2.3264874E-8 | 6.3223457 | 120-121 |
| GCAGATC | 1795 | 9.304131E-8 | 5.642358 | 122-123 |

Produced by FastQC (version 0.11.5)
