## Supplementary material 4 for "Natural and human-mediated drivers of microevolution in Neotropical palms: a historical genomics approach": outLAU_AP03.1_pf_trimmed_masked_sickled_fastqc.html

outLAU\_AP03.1\_pf\_trimmed\_masked\_sickled.fastq FastQC Report 

FastQC Report

mar. 25 juin 2019  
outLAU\_AP03.1\_pf\_trimmed\_masked\_sickled.fastq

### Summary

- Basic Statistics
- Per base sequence quality
- Per tile sequence quality
- Per sequence quality scores
- Per base sequence content
- Per sequence GC content
- Per base N content
- Sequence Length Distribution
- Sequence Duplication Levels
- Overrepresented sequences
- Adapter Content
- Kmer Content

### Basic Statistics

| Measure | Value |
| --- | --- |
| Filename | outLAU\_AP03.1\_pf\_trimmed\_masked\_sickled.fastq |
| File type | Conventional base calls |
| Encoding | Sanger / Illumina 1.9 |
| Total Sequences | 1315885 |
| Sequences flagged as poor quality | 0 |
| Sequence length | 100-131 |
| %GC | 43 |

### Per base sequence quality

### Per tile sequence quality

### Per sequence quality scores

### Per base sequence content

### Per sequence GC content

### Per base N content

### Sequence Length Distribution

### Sequence Duplication Levels

### Overrepresented sequences

No overrepresented sequences

### Adapter Content

### Kmer Content

| Sequence | Count | PValue | Obs/Exp Max | Max Obs/Exp Position |
| --- | --- | --- | --- | --- |
| TGCGCTA | 505 | 0.0 | 53.299553 | 6 |
| GCGCTAC | 150 | 0.0 | 46.8276 | 7 |
| CTGCGCT | 615 | 0.0 | 43.74205 | 5 |
| TCTGCGC | 630 | 0.0 | 41.71618 | 4 |
| GCGCTAA | 170 | 9.094947E-12 | 37.875263 | 7 |
| ATCTGCG | 765 | 0.0 | 34.31924 | 3 |
| GCGCTAT | 210 | 3.6379788E-12 | 33.448284 | 7 |
| CGCTAAG | 150 | 1.7179445E-7 | 31.194925 | 8 |
| GCGCTAG | 240 | 5.275069E-10 | 26.828312 | 7 |
| CGCTATT | 240 | 2.2669522E-4 | 17.059723 | 8 |
| GATCTGC | 1870 | 0.0 | 14.6330595 | 2 |
| CTTCCGA | 675 | 3.0304363E-9 | 12.868526 | 1 |
| GCGCAGA | 1780 | 0.0 | 12.36036 | 124-125 |
| ACGTCTG | 465 | 8.274583E-9 | 12.029219 | 124-125 |
| AGCGCAG | 1960 | 0.0 | 11.795999 | 124-125 |
| CACACGT | 615 | 3.092282E-11 | 11.520666 | 124-125 |
| GCACACG | 650 | 7.2759576E-12 | 11.474024 | 124-125 |
| TAGCGCA | 1950 | 0.0 | 11.117291 | 122-123 |
| ACACGTC | 545 | 7.3378033E-9 | 10.947693 | 124-125 |
| GTAGCGC | 490 | 2.695415E-7 | 10.498168 | 122-123 |

Produced by FastQC (version 0.11.5)
