## Supplementary material 4 for "Natural and human-mediated drivers of microevolution in Neotropical palms: a historical genomics approach": outLAU_AP03.2_pf_trimmed_masked_sickled_fastqc.html

outLAU\_AP03.2\_pf\_trimmed\_masked\_sickled.fastq FastQC Report 

FastQC Report

mar. 25 juin 2019  
outLAU\_AP03.2\_pf\_trimmed\_masked\_sickled.fastq

### Summary

- Basic Statistics
- Per base sequence quality
- Per tile sequence quality
- Per sequence quality scores
- Per base sequence content
- Per sequence GC content
- Per base N content
- Sequence Length Distribution
- Sequence Duplication Levels
- Overrepresented sequences
- Adapter Content
- Kmer Content

### Basic Statistics

| Measure | Value |
| --- | --- |
| Filename | outLAU\_AP03.2\_pf\_trimmed\_masked\_sickled.fastq |
| File type | Conventional base calls |
| Encoding | Sanger / Illumina 1.9 |
| Total Sequences | 1315885 |
| Sequences flagged as poor quality | 0 |
| Sequence length | 100-131 |
| %GC | 44 |

### Per base sequence quality

### Per tile sequence quality

### Per sequence quality scores

### Per base sequence content

### Per sequence GC content

### Per base N content

### Sequence Length Distribution

### Sequence Duplication Levels

### Overrepresented sequences

No overrepresented sequences

### Adapter Content

### Kmer Content

| Sequence | Count | PValue | Obs/Exp Max | Max Obs/Exp Position |
| --- | --- | --- | --- | --- |
| TGCGCTA | 375 | 0.0 | 50.235245 | 6 |
| CTGCGCT | 460 | 0.0 | 40.87424 | 5 |
| TCTGCGC | 500 | 0.0 | 38.62849 | 4 |
| ATCTGCG | 510 | 0.0 | 37.804996 | 3 |
| GCGCTAT | 180 | 3.6379788E-11 | 33.79747 | 7 |
| GCGCTAC | 115 | 1.1795655E-6 | 33.663883 | 7 |
| GCGCTAA | 230 | 6.220944E-10 | 26.450193 | 7 |
| GCGCTAG | 120 | 0.0019712066 | 23.04373 | 7 |
| TTAGCGC | 730 | 0.0 | 15.422876 | 122-123 |
| GCGCAGA | 1765 | 0.0 | 15.056143 | 124-125 |
| GATCTGC | 1450 | 0.0 | 14.389374 | 2 |
| AGCGCAG | 1855 | 0.0 | 14.086894 | 124-125 |
| CTAGCGC | 480 | 1.5279511E-10 | 13.994124 | 120-121 |
| TAGCGCA | 1770 | 0.0 | 13.944934 | 122-123 |
| GTAGCGC | 435 | 7.838935E-8 | 12.941033 | 122-123 |
| CTAGTAC | 315 | 0.0028655336 | 12.201515 | 2 |
| AGAGCGT | 590 | 1.713488E-9 | 12.010888 | 124-125 |
| CGTGTAG | 310 | 9.705873E-4 | 11.429717 | 124-125 |
| GCGTCGT | 390 | 4.5228247E-5 | 11.356449 | 124-125 |
| AGCGTCG | 445 | 1.8394941E-5 | 10.704038 | 122-123 |

Produced by FastQC (version 0.11.5)
