## Supplementary material 4 for "Natural and human-mediated drivers of microevolution in Neotropical palms: a historical genomics approach": outLAU_AP05.1_pf_trimmed_masked_sickled_fastqc.html

outLAU\_AP05.1\_pf\_trimmed\_masked\_sickled.fastq FastQC Report 

FastQC Report

mar. 25 juin 2019  
outLAU\_AP05.1\_pf\_trimmed\_masked\_sickled.fastq

### Summary

- Basic Statistics
- Per base sequence quality
- Per tile sequence quality
- Per sequence quality scores
- Per base sequence content
- Per sequence GC content
- Per base N content
- Sequence Length Distribution
- Sequence Duplication Levels
- Overrepresented sequences
- Adapter Content
- Kmer Content

### Basic Statistics

| Measure | Value |
| --- | --- |
| Filename | outLAU\_AP05.1\_pf\_trimmed\_masked\_sickled.fastq |
| File type | Conventional base calls |
| Encoding | Sanger / Illumina 1.9 |
| Total Sequences | 1522560 |
| Sequences flagged as poor quality | 0 |
| Sequence length | 100-131 |
| %GC | 43 |

### Per base sequence quality

### Per tile sequence quality

### Per sequence quality scores

### Per base sequence content

### Per sequence GC content

### Per base N content

### Sequence Length Distribution

### Sequence Duplication Levels

### Overrepresented sequences

No overrepresented sequences

### Adapter Content

### Kmer Content

| Sequence | Count | PValue | Obs/Exp Max | Max Obs/Exp Position |
| --- | --- | --- | --- | --- |
| CTTCCGA | 855 | 2.910383E-11 | 12.625696 | 1 |
| ACACGAC | 400 | 0.008507194 | 10.544663 | 6 |
| TCGGAAG | 1260 | 0.0 | 9.5288105 | 124-125 |
| CTGCTAA | 1015 | 6.5935274E-7 | 8.919347 | 9 |
| GCACACG | 505 | 1.0713136E-5 | 8.83066 | 124-125 |
| CGGAAGA | 1170 | 1.4551915E-11 | 7.623049 | 124-125 |
| TCTGCTA | 1290 | 2.524559E-6 | 7.478774 | 8 |
| TTAGCAG | 2115 | 0.0 | 7.298663 | 124-125 |
| CACACGT | 485 | 0.005095322 | 7.072932 | 124-125 |
| TGCTAAG | 1115 | 4.4930915E-5 | 7.025272 | 6 |
| CTTAGCA | 2220 | 0.0 | 6.7563014 | 122-123 |
| TAGCAGA | 2265 | 0.0 | 6.209503 | 124-125 |
| GCAGATC | 2600 | 0.0 | 5.97279 | 120-121 |
| ACTTAGC | 825 | 7.488001E-4 | 5.7847347 | 122-123 |
| GATCTGC | 1690 | 1.351671E-4 | 5.7046356 | 6 |
| TCTAGTC | 715 | 0.007869839 | 5.5618525 | 114-115 |
| ATCTGCT | 1765 | 2.5159895E-4 | 5.4624124 | 7 |
| TCTTAGC | 1330 | 3.2005464E-6 | 5.4163766 | 124-125 |
| AGAGCAC | 1395 | 1.3387125E-6 | 5.4099054 | 124-125 |
| TTCCGAT | 1910 | 1.3947886E-4 | 5.348701 | 2 |

Produced by FastQC (version 0.11.5)
