## Supplementary material 4 for "Natural and human-mediated drivers of microevolution in Neotropical palms: a historical genomics approach": outLAU_AP05.2_pf_trimmed_masked_sickled_fastqc.html

outLAU\_AP05.2\_pf\_trimmed\_masked\_sickled.fastq FastQC Report 

FastQC Report

mar. 25 juin 2019  
outLAU\_AP05.2\_pf\_trimmed\_masked\_sickled.fastq

### Summary

- Basic Statistics
- Per base sequence quality
- Per tile sequence quality
- Per sequence quality scores
- Per base sequence content
- Per sequence GC content
- Per base N content
- Sequence Length Distribution
- Sequence Duplication Levels
- Overrepresented sequences
- Adapter Content
- Kmer Content

### Basic Statistics

| Measure | Value |
| --- | --- |
| Filename | outLAU\_AP05.2\_pf\_trimmed\_masked\_sickled.fastq |
| File type | Conventional base calls |
| Encoding | Sanger / Illumina 1.9 |
| Total Sequences | 1522560 |
| Sequences flagged as poor quality | 0 |
| Sequence length | 100-131 |
| %GC | 43 |

### Per base sequence quality

### Per tile sequence quality

### Per sequence quality scores

### Per base sequence content

### Per sequence GC content

### Per base N content

### Sequence Length Distribution

### Sequence Duplication Levels

### Overrepresented sequences

No overrepresented sequences

### Adapter Content

### Kmer Content

| Sequence | Count | PValue | Obs/Exp Max | Max Obs/Exp Position |
| --- | --- | --- | --- | --- |
| TCGGAAG | 1200 | 0.0 | 13.305621 | 124-125 |
| CTGCTAA | 895 | 1.2732926E-11 | 12.047176 | 5 |
| CTTCCGA | 745 | 2.4227847E-7 | 10.579184 | 1 |
| CGGAAGA | 1110 | 0.0 | 9.220805 | 124-125 |
| GCTAAGC | 560 | 0.0016383297 | 9.10637 | 7 |
| TGCTAAG | 1095 | 5.617767E-8 | 8.8142605 | 6 |
| GCGAACC | 300 | 0.009171888 | 8.715276 | 96-97 |
| TCTGCTA | 1195 | 2.634988E-8 | 8.530397 | 4 |
| TTAGCAG | 2045 | 0.0 | 7.8076997 | 124-125 |
| TAGCAGA | 2065 | 0.0 | 7.73208 | 124-125 |
| CTTAGCA | 2100 | 0.0 | 7.6032124 | 124-125 |
| AGAGCGT | 570 | 0.0043864343 | 7.182522 | 124-125 |
| AAGAGCG | 740 | 1.8503536E-4 | 7.0265446 | 122-123 |
| CCTTAGC | 1010 | 4.1009625E-6 | 6.7322116 | 122-123 |
| TTCCGAT | 1385 | 1.3133459E-4 | 6.126241 | 2 |
| ACTTAGC | 875 | 0.001414594 | 5.9424496 | 122-123 |
| TCTTAGC | 1055 | 2.6424078E-4 | 5.820906 | 124-125 |
| GATCTGC | 1790 | 2.152945E-5 | 5.6881638 | 2 |
| GCAGATC | 2685 | 1.8189894E-12 | 5.3735843 | 120-121 |
| ATCGGAA | 2670 | 1.6370905E-11 | 5.366716 | 124-125 |

Produced by FastQC (version 0.11.5)
