## Supplementary material 4 for "Natural and human-mediated drivers of microevolution in Neotropical palms: a historical genomics approach": outLAU_AP07.1_pf_trimmed_masked_sickled_fastqc.html

outLAU\_AP07.1\_pf\_trimmed\_masked\_sickled.fastq FastQC Report 

FastQC Report

mar. 25 juin 2019  
outLAU\_AP07.1\_pf\_trimmed\_masked\_sickled.fastq

### Summary

- Basic Statistics
- Per base sequence quality
- Per tile sequence quality
- Per sequence quality scores
- Per base sequence content
- Per sequence GC content
- Per base N content
- Sequence Length Distribution
- Sequence Duplication Levels
- Overrepresented sequences
- Adapter Content
- Kmer Content

### Basic Statistics

| Measure | Value |
| --- | --- |
| Filename | outLAU\_AP07.1\_pf\_trimmed\_masked\_sickled.fastq |
| File type | Conventional base calls |
| Encoding | Sanger / Illumina 1.9 |
| Total Sequences | 540703 |
| Sequences flagged as poor quality | 0 |
| Sequence length | 100-131 |
| %GC | 43 |

### Per base sequence quality

### Per tile sequence quality

### Per sequence quality scores

### Per base sequence content

### Per sequence GC content

### Per base N content

### Sequence Length Distribution

### Sequence Duplication Levels

### Overrepresented sequences

No overrepresented sequences

### Adapter Content

### Kmer Content

| Sequence | Count | PValue | Obs/Exp Max | Max Obs/Exp Position |
| --- | --- | --- | --- | --- |
| ACGTCTG | 175 | 1.7162396E-5 | 18.37023 | 124-125 |
| ACACGTC | 185 | 3.2606196E-5 | 17.048588 | 122-123 |
| TGCTGCC | 455 | 7.4884156E-8 | 14.890551 | 6 |
| CACACGT | 235 | 1.800947E-5 | 14.793835 | 120-121 |
| GCACACG | 250 | 2.224368E-5 | 14.466557 | 124-125 |
| ATCTGCT | 585 | 5.366019E-10 | 14.440214 | 3 |
| ACGGGAG | 125 | 0.006351332 | 13.857304 | 24-25 |
| GATCTGC | 545 | 4.908361E-8 | 13.407221 | 2 |
| CGTCTGA | 215 | 0.0016953469 | 13.083448 | 124-125 |
| TCGGAAG | 530 | 1.4570105E-9 | 12.131283 | 124-125 |
| CACGTCT | 230 | 0.003246295 | 11.998871 | 122-123 |
| GGGCAGC | 415 | 2.3937177E-5 | 10.449963 | 122-123 |
| TCTGCTG | 670 | 8.226065E-6 | 10.095459 | 4 |
| CTGCTGC | 855 | 1.521615E-5 | 8.584562 | 5 |
| AGGTATT | 330 | 0.0034252563 | 8.394755 | 68-69 |
| AGCACAC | 440 | 0.004917612 | 8.064176 | 122-123 |
| GCAGATC | 960 | 5.977199E-9 | 8.047572 | 120-121 |
| CAGCAGA | 1140 | 3.0340743E-9 | 7.3693047 | 118-119 |
| ATCGGAA | 1150 | 8.0897735E-8 | 6.9886746 | 124-125 |
| AGCAGAT | 1025 | 1.4776142E-6 | 6.7059035 | 118-119 |

Produced by FastQC (version 0.11.5)
