## Supplementary material 4 for "Natural and human-mediated drivers of microevolution in Neotropical palms: a historical genomics approach": outLAU_AP07.2_pf_trimmed_masked_sickled_fastqc.html

outLAU\_AP07.2\_pf\_trimmed\_masked\_sickled.fastq FastQC Report 

FastQC Report

mar. 25 juin 2019  
outLAU\_AP07.2\_pf\_trimmed\_masked\_sickled.fastq

### Summary

- Basic Statistics
- Per base sequence quality
- Per tile sequence quality
- Per sequence quality scores
- Per base sequence content
- Per sequence GC content
- Per base N content
- Sequence Length Distribution
- Sequence Duplication Levels
- Overrepresented sequences
- Adapter Content
- Kmer Content

### Basic Statistics

| Measure | Value |
| --- | --- |
| Filename | outLAU\_AP07.2\_pf\_trimmed\_masked\_sickled.fastq |
| File type | Conventional base calls |
| Encoding | Sanger / Illumina 1.9 |
| Total Sequences | 540703 |
| Sequences flagged as poor quality | 0 |
| Sequence length | 100-131 |
| %GC | 43 |

### Per base sequence quality

### Per tile sequence quality

### Per sequence quality scores

### Per base sequence content

### Per sequence GC content

### Per base N content

### Sequence Length Distribution

### Sequence Duplication Levels

### Overrepresented sequences

No overrepresented sequences

### Adapter Content

### Kmer Content

| Sequence | Count | PValue | Obs/Exp Max | Max Obs/Exp Position |
| --- | --- | --- | --- | --- |
| CGTGTTA | 45 | 0.009112532 | 26.687223 | 74-75 |
| GAGCGTC | 190 | 7.6440665E-6 | 20.173922 | 122-123 |
| ATCTGCT | 520 | 1.4551915E-11 | 16.182882 | 3 |
| TCGGAAG | 550 | 5.456968E-12 | 15.414541 | 124-125 |
| AGAGCGT | 210 | 4.9198064E-4 | 15.407422 | 120-121 |
| TCTGCTG | 625 | 1.8189894E-12 | 15.187115 | 4 |
| TGCTGCC | 605 | 9.094947E-12 | 14.92806 | 6 |
| ACGAAAC | 120 | 0.008621863 | 13.231565 | 16-17 |
| GATCTGC | 595 | 2.053639E-9 | 13.202997 | 2 |
| AAGAGCG | 320 | 6.0750106E-5 | 13.000012 | 120-121 |
| TCGGCAG | 260 | 0.005310146 | 11.233261 | 114-115 |
| TCGACAA | 250 | 0.0025517778 | 10.17738 | 64-65 |
| CGGAAGA | 485 | 1.8535256E-4 | 9.878983 | 122-123 |
| CTGCTGC | 1110 | 2.593879E-9 | 9.030955 | 5 |
| ATCGGAA | 1120 | 7.5153366E-8 | 8.0149145 | 124-125 |
| CAGCAGA | 1030 | 1.2290548E-7 | 7.7849636 | 118-119 |
| GAAGAGC | 805 | 2.9697266E-4 | 7.4341235 | 124-125 |
| GGCAGCA | 1015 | 1.1935737E-5 | 7.0807486 | 122-123 |
| GCAGATC | 1045 | 5.7263987E-6 | 7.0770936 | 120-121 |
| GGAAGAG | 900 | 5.3976936E-4 | 6.4346085 | 118-119 |

Produced by FastQC (version 0.11.5)
