## Supplementary material 4 for "Natural and human-mediated drivers of microevolution in Neotropical palms: a historical genomics approach": outLAU_AP08.1_pf_trimmed_masked_sickled_fastqc.html

outLAU\_AP08.1\_pf\_trimmed\_masked\_sickled.fastq FastQC Report 

FastQC Report

mar. 25 juin 2019  
outLAU\_AP08.1\_pf\_trimmed\_masked\_sickled.fastq

### Summary

- Basic Statistics
- Per base sequence quality
- Per tile sequence quality
- Per sequence quality scores
- Per base sequence content
- Per sequence GC content
- Per base N content
- Sequence Length Distribution
- Sequence Duplication Levels
- Overrepresented sequences
- Adapter Content
- Kmer Content

### Basic Statistics

| Measure | Value |
| --- | --- |
| Filename | outLAU\_AP08.1\_pf\_trimmed\_masked\_sickled.fastq |
| File type | Conventional base calls |
| Encoding | Sanger / Illumina 1.9 |
| Total Sequences | 872484 |
| Sequences flagged as poor quality | 0 |
| Sequence length | 100-131 |
| %GC | 43 |

### Per base sequence quality

### Per tile sequence quality

### Per sequence quality scores

### Per base sequence content

### Per sequence GC content

### Per base N content

### Sequence Length Distribution

### Sequence Duplication Levels

### Overrepresented sequences

No overrepresented sequences

### Adapter Content

### Kmer Content

| Sequence | Count | PValue | Obs/Exp Max | Max Obs/Exp Position |
| --- | --- | --- | --- | --- |
| ACGACGC | 140 | 1.16809184E-4 | 25.122675 | 8 |
| CTACACG | 185 | 3.0663876E-5 | 22.134281 | 4 |
| CGACGCT | 165 | 3.542063E-4 | 21.327631 | 9 |
| ACACGAC | 165 | 3.5994433E-4 | 21.27696 | 6 |
| ATCTGGC | 1070 | 0.0 | 20.198206 | 3 |
| CTGGCAT | 1055 | 0.0 | 19.980299 | 5 |
| TACACGA | 180 | 6.4412935E-4 | 19.517792 | 5 |
| TCTGGCA | 1115 | 0.0 | 19.411806 | 4 |
| GTGTTCG | 210 | 0.001823863 | 16.71364 | 4 |
| CACGACG | 230 | 0.0033323069 | 15.270255 | 7 |
| TGGCATC | 1500 | 0.0 | 14.432873 | 6 |
| CCCTACA | 265 | 0.008675383 | 13.218868 | 2 |
| TCGGAAA | 360 | 0.0048165973 | 11.381995 | 7 |
| GATCTGG | 2225 | 0.0 | 10.758278 | 2 |
| CGATCTG | 2470 | 0.0 | 9.881848 | 1 |
| GGCATCA | 1280 | 5.2241376E-9 | 8.688934 | 7 |
| TCTTCCG | 420 | 6.5459876E-4 | 7.7098646 | 14-15 |
| TCGGAAG | 680 | 2.0280264E-5 | 7.6289306 | 124-125 |
| CGTCTTC | 375 | 0.008340475 | 7.603785 | 88-89 |
| GTGGTTC | 555 | 0.0015964097 | 6.4166727 | 20-21 |

Produced by FastQC (version 0.11.5)
