## Supplementary material 4 for "Natural and human-mediated drivers of microevolution in Neotropical palms: a historical genomics approach": outLAU_AP08.2_pf_trimmed_masked_sickled_fastqc.html

outLAU\_AP08.2\_pf\_trimmed\_masked\_sickled.fastq FastQC Report 

FastQC Report

mar. 25 juin 2019  
outLAU\_AP08.2\_pf\_trimmed\_masked\_sickled.fastq

### Summary

- Basic Statistics
- Per base sequence quality
- Per tile sequence quality
- Per sequence quality scores
- Per base sequence content
- Per sequence GC content
- Per base N content
- Sequence Length Distribution
- Sequence Duplication Levels
- Overrepresented sequences
- Adapter Content
- Kmer Content

### Basic Statistics

| Measure | Value |
| --- | --- |
| Filename | outLAU\_AP08.2\_pf\_trimmed\_masked\_sickled.fastq |
| File type | Conventional base calls |
| Encoding | Sanger / Illumina 1.9 |
| Total Sequences | 872484 |
| Sequences flagged as poor quality | 0 |
| Sequence length | 100-131 |
| %GC | 43 |

### Per base sequence quality

### Per tile sequence quality

### Per sequence quality scores

### Per base sequence content

### Per sequence GC content

### Per base N content

### Sequence Length Distribution

### Sequence Duplication Levels

### Overrepresented sequences

No overrepresented sequences

### Adapter Content

### Kmer Content

| Sequence | Count | PValue | Obs/Exp Max | Max Obs/Exp Position |
| --- | --- | --- | --- | --- |
| TCTGGCA | 795 | 0.0 | 15.246824 | 4 |
| CTGGCAT | 830 | 0.0 | 14.656023 | 5 |
| GTCGTGT | 175 | 2.5612351E-5 | 14.254219 | 16-17 |
| ATCTGGC | 865 | 0.0 | 13.991523 | 3 |
| GCGTCGT | 220 | 1.5895694E-5 | 12.577858 | 14-15 |
| CGTTGTT | 175 | 0.003660472 | 11.809156 | 38-39 |
| AAGAGCG | 425 | 8.828578E-6 | 11.435707 | 124-125 |
| CGTGTAG | 175 | 0.0055897785 | 11.15788 | 18-19 |
| TGGCATC | 1205 | 0.0 | 11.041975 | 6 |
| AGAGCGT | 290 | 0.007817729 | 10.664962 | 124-125 |
| TCGGAAG | 700 | 1.5188816E-6 | 10.292173 | 7 |
| CGGAAGA | 650 | 7.0163587E-6 | 10.233167 | 8 |
| GAGCGTC | 275 | 1.5488743E-4 | 10.059187 | 12-13 |
| CGTCGTG | 265 | 0.0012110815 | 9.413164 | 16-17 |
| GCCAGAT | 1150 | 2.6830094E-9 | 9.017171 | 1 |
| ATCGGAA | 1335 | 2.9127477E-8 | 7.8903255 | 6 |
| GATCTGG | 1860 | 3.7449354E-8 | 6.498453 | 2 |
| TGCCAGA | 1230 | 2.6945945E-6 | 6.465865 | 124-125 |
| CGATCTG | 2215 | 7.9671736E-10 | 6.406402 | 1 |
| ATGCCAG | 1285 | 1.2864257E-6 | 6.352058 | 122-123 |

Produced by FastQC (version 0.11.5)
