## Supplementary material 4 for "Natural and human-mediated drivers of microevolution in Neotropical palms: a historical genomics approach": outLAU_AP09.1_pf_trimmed_masked_sickled_fastqc.html

outLAU\_AP09.1\_pf\_trimmed\_masked\_sickled.fastq FastQC Report 

FastQC Report

mar. 25 juin 2019  
outLAU\_AP09.1\_pf\_trimmed\_masked\_sickled.fastq

### Summary

- Basic Statistics
- Per base sequence quality
- Per tile sequence quality
- Per sequence quality scores
- Per base sequence content
- Per sequence GC content
- Per base N content
- Sequence Length Distribution
- Sequence Duplication Levels
- Overrepresented sequences
- Adapter Content
- Kmer Content

### Basic Statistics

| Measure | Value |
| --- | --- |
| Filename | outLAU\_AP09.1\_pf\_trimmed\_masked\_sickled.fastq |
| File type | Conventional base calls |
| Encoding | Sanger / Illumina 1.9 |
| Total Sequences | 994178 |
| Sequences flagged as poor quality | 0 |
| Sequence length | 100-131 |
| %GC | 43 |

### Per base sequence quality

### Per tile sequence quality

### Per sequence quality scores

### Per base sequence content

### Per sequence GC content

### Per base N content

### Sequence Length Distribution

### Sequence Duplication Levels

### Overrepresented sequences

No overrepresented sequences

### Adapter Content

### Kmer Content

| Sequence | Count | PValue | Obs/Exp Max | Max Obs/Exp Position |
| --- | --- | --- | --- | --- |
| TCTGGCG | 450 | 0.0 | 32.521835 | 4 |
| TGGCGAC | 480 | 0.0 | 28.184471 | 6 |
| CTGGCGA | 515 | 0.0 | 27.363564 | 5 |
| GGCGACT | 245 | 3.510962E-4 | 16.10799 | 7 |
| ACACGTC | 305 | 3.4169716E-8 | 15.87845 | 124-125 |
| CACACGT | 390 | 1.8189894E-10 | 15.522204 | 124-125 |
| ATCTGGC | 1015 | 0.0 | 15.48638 | 3 |
| ACGTCTG | 295 | 4.0497798E-7 | 15.048645 | 124-125 |
| CACGTCT | 310 | 1.6069936E-4 | 11.716761 | 124-125 |
| GGCGACC | 355 | 0.0057424344 | 11.116783 | 7 |
| GCACACG | 420 | 2.1138845E-5 | 10.569881 | 124-125 |
| CTTCCGA | 495 | 6.308011E-4 | 10.106433 | 1 |
| AGTCGCC | 360 | 8.111233E-4 | 9.834162 | 122-123 |
| TCGCCAG | 1190 | 0.0 | 9.495936 | 124-125 |
| CGGAAGA | 735 | 2.2018321E-7 | 8.7853565 | 124-125 |
| CGCCAGA | 945 | 8.731149E-11 | 8.77423 | 118-119 |
| TCGGAAG | 810 | 1.0682197E-7 | 8.47014 | 124-125 |
| GTCGCCA | 1205 | 3.783498E-10 | 7.7031355 | 124-125 |
| TGTCGCC | 550 | 8.327118E-4 | 7.537862 | 118-119 |
| GATCTGG | 2390 | 0.0 | 7.2683735 | 2 |

Produced by FastQC (version 0.11.5)
