## Supplementary material 4 for "Natural and human-mediated drivers of microevolution in Neotropical palms: a historical genomics approach": outLAU_AP09.2_pf_trimmed_masked_sickled_fastqc.html

outLAU\_AP09.2\_pf\_trimmed\_masked\_sickled.fastq FastQC Report 

FastQC Report

mar. 25 juin 2019  
outLAU\_AP09.2\_pf\_trimmed\_masked\_sickled.fastq

### Summary

- Basic Statistics
- Per base sequence quality
- Per tile sequence quality
- Per sequence quality scores
- Per base sequence content
- Per sequence GC content
- Per base N content
- Sequence Length Distribution
- Sequence Duplication Levels
- Overrepresented sequences
- Adapter Content
- Kmer Content

### Basic Statistics

| Measure | Value |
| --- | --- |
| Filename | outLAU\_AP09.2\_pf\_trimmed\_masked\_sickled.fastq |
| File type | Conventional base calls |
| Encoding | Sanger / Illumina 1.9 |
| Total Sequences | 994178 |
| Sequences flagged as poor quality | 0 |
| Sequence length | 100-131 |
| %GC | 43 |

### Per base sequence quality

### Per tile sequence quality

### Per sequence quality scores

### Per base sequence content

### Per sequence GC content

### Per base N content

### Sequence Length Distribution

### Sequence Duplication Levels

### Overrepresented sequences

No overrepresented sequences

### Adapter Content

### Kmer Content

| Sequence | Count | PValue | Obs/Exp Max | Max Obs/Exp Position |
| --- | --- | --- | --- | --- |
| TCTGGCG | 300 | 0.0 | 36.823704 | 4 |
| TGGCGAC | 395 | 0.0 | 30.8496 | 6 |
| CTGGCGA | 430 | 0.0 | 26.98496 | 5 |
| CGTGTAG | 125 | 1.7208555E-4 | 23.728983 | 124-125 |
| GCGTCGT | 235 | 7.299561E-5 | 15.517887 | 120-121 |
| GTCGTGT | 185 | 0.0031191637 | 15.423005 | 122-123 |
| GCGACCC | 210 | 0.003664667 | 15.053886 | 8 |
| GAGCGTC | 315 | 5.43422E-6 | 13.9566965 | 118-119 |
| GGCGACC | 270 | 0.001200779 | 13.696661 | 7 |
| ATCTGGC | 845 | 0.0 | 13.689333 | 3 |
| CGTCGTG | 270 | 0.0019805254 | 12.816583 | 124-125 |
| AGCGTCG | 285 | 3.7554713E-4 | 12.795451 | 120-121 |
| CGGAAGA | 760 | 9.094947E-12 | 12.358847 | 124-125 |
| AGAGCGT | 275 | 0.0041734837 | 11.6031475 | 120-121 |
| CGCCAGA | 900 | 3.8380676E-10 | 10.039235 | 122-123 |
| TCGGAAG | 760 | 2.9420335E-8 | 10.011425 | 122-123 |
| TCGCCAG | 1035 | 2.3646862E-10 | 9.5527315 | 124-125 |
| GTCGCCA | 1135 | 1.3587851E-9 | 8.7110815 | 124-125 |
| AAGAGCG | 520 | 8.8187604E-4 | 8.556125 | 124-125 |
| GATCTGG | 2000 | 2.7284841E-11 | 7.0555487 | 2 |

Produced by FastQC (version 0.11.5)
