## Supplementary material 4 for "Natural and human-mediated drivers of microevolution in Neotropical palms: a historical genomics approach": outLAU_AP10.1_pf_trimmed_masked_sickled_fastqc.html

outLAU\_AP10.1\_pf\_trimmed\_masked\_sickled.fastq FastQC Report 

FastQC Report

mar. 25 juin 2019  
outLAU\_AP10.1\_pf\_trimmed\_masked\_sickled.fastq

### Summary

- Basic Statistics
- Per base sequence quality
- Per tile sequence quality
- Per sequence quality scores
- Per base sequence content
- Per sequence GC content
- Per base N content
- Sequence Length Distribution
- Sequence Duplication Levels
- Overrepresented sequences
- Adapter Content
- Kmer Content

### Basic Statistics

| Measure | Value |
| --- | --- |
| Filename | outLAU\_AP10.1\_pf\_trimmed\_masked\_sickled.fastq |
| File type | Conventional base calls |
| Encoding | Sanger / Illumina 1.9 |
| Total Sequences | 980266 |
| Sequences flagged as poor quality | 0 |
| Sequence length | 100-131 |
| %GC | 45 |

### Per base sequence quality

### Per tile sequence quality

### Per sequence quality scores

### Per base sequence content

### Per sequence GC content

### Per base N content

### Sequence Length Distribution

### Sequence Duplication Levels

### Overrepresented sequences

No overrepresented sequences

### Adapter Content

### Kmer Content

| Sequence | Count | PValue | Obs/Exp Max | Max Obs/Exp Position |
| --- | --- | --- | --- | --- |
| TGGTCGT | 510 | 0.0 | 20.667189 | 6 |
| GGTCGTA | 225 | 0.0028466382 | 15.635899 | 7 |
| CTGGTCG | 725 | 3.6379788E-11 | 13.730614 | 5 |
| CGTCTGA | 355 | 5.3178155E-8 | 13.330607 | 122-123 |
| TCTGGTC | 925 | 1.8189894E-12 | 12.649598 | 4 |
| ATCTGGT | 1015 | 7.2759576E-12 | 11.513347 | 3 |
| CACGTCT | 420 | 5.8560545E-7 | 11.085162 | 120-121 |
| CACACGT | 430 | 9.2361915E-7 | 10.700244 | 118-119 |
| AAACGAC | 370 | 2.2827028E-4 | 9.679353 | 120-121 |
| ACGACCA | 1625 | 0.0 | 9.408729 | 122-123 |
| GCACACG | 440 | 8.77388E-5 | 9.285241 | 124-125 |
| GACCAGA | 1845 | 0.0 | 9.058772 | 124-125 |
| CGGAAGA | 995 | 3.8198777E-11 | 8.585349 | 124-125 |
| AACGACC | 680 | 3.7525388E-6 | 7.9000587 | 120-121 |
| CGACCAG | 1755 | 0.0 | 7.674668 | 122-123 |
| ACCAGAT | 1905 | 0.0 | 7.603663 | 124-125 |
| TCGGAAG | 1035 | 9.26957E-9 | 7.386079 | 122-123 |
| AGCACAC | 850 | 5.1746396E-5 | 6.5542884 | 124-125 |
| GACGACC | 810 | 0.0021044728 | 5.7478623 | 120-121 |
| CACGACC | 750 | 0.0056567155 | 5.7301764 | 120-121 |

Produced by FastQC (version 0.11.5)
