## Supplementary material 4 for "Natural and human-mediated drivers of microevolution in Neotropical palms: a historical genomics approach": outLAU_AP10.2_pf_trimmed_masked_sickled_fastqc.html

outLAU\_AP10.2\_pf\_trimmed\_masked\_sickled.fastq FastQC Report 

FastQC Report

mar. 25 juin 2019  
outLAU\_AP10.2\_pf\_trimmed\_masked\_sickled.fastq

### Summary

- Basic Statistics
- Per base sequence quality
- Per tile sequence quality
- Per sequence quality scores
- Per base sequence content
- Per sequence GC content
- Per base N content
- Sequence Length Distribution
- Sequence Duplication Levels
- Overrepresented sequences
- Adapter Content
- Kmer Content

### Basic Statistics

| Measure | Value |
| --- | --- |
| Filename | outLAU\_AP10.2\_pf\_trimmed\_masked\_sickled.fastq |
| File type | Conventional base calls |
| Encoding | Sanger / Illumina 1.9 |
| Total Sequences | 980266 |
| Sequences flagged as poor quality | 0 |
| Sequence length | 100-131 |
| %GC | 45 |

### Per base sequence quality

### Per tile sequence quality

### Per sequence quality scores

### Per base sequence content

### Per sequence GC content

### Per base N content

### Sequence Length Distribution

### Sequence Duplication Levels

### Overrepresented sequences

No overrepresented sequences

### Adapter Content

### Kmer Content

| Sequence | Count | PValue | Obs/Exp Max | Max Obs/Exp Position |
| --- | --- | --- | --- | --- |
| TGGTCGT | 555 | 0.0 | 21.985836 | 6 |
| CTGGTCG | 665 | 0.0 | 19.987629 | 5 |
| TCTGGTC | 720 | 0.0 | 17.655876 | 4 |
| GGTCGTA | 195 | 0.0015811952 | 17.074345 | 7 |
| ATCTGGT | 970 | 0.0 | 13.103249 | 3 |
| TCGTGTA | 275 | 6.2559E-4 | 12.04359 | 120-121 |
| GTACGAC | 260 | 0.005632564 | 11.146111 | 120-121 |
| GTAGGGA | 320 | 0.0012969754 | 11.039897 | 124-125 |
| ACGACCA | 1615 | 0.0 | 10.820651 | 122-123 |
| CGGAAGA | 970 | 0.0 | 10.47083 | 124-125 |
| CGACCAG | 1675 | 0.0 | 10.433046 | 122-123 |
| AAGAGCG | 635 | 6.7542715E-8 | 10.431399 | 124-125 |
| TTCCGAT | 1080 | 7.6397555E-11 | 10.134642 | 1 |
| AGAGCGT | 505 | 7.943419E-6 | 10.128193 | 122-123 |
| GACCAGA | 1900 | 0.0 | 9.994012 | 124-125 |
| TCGGAAG | 1015 | 1.4551915E-11 | 9.571537 | 124-125 |
| GTGTAGG | 410 | 0.0012795704 | 9.356227 | 122-123 |
| CGTCGTG | 480 | 9.0985806E-5 | 9.254087 | 118-119 |
| AGCGTCG | 485 | 4.213875E-4 | 9.105069 | 124-125 |
| TACGACC | 555 | 3.4280114E-5 | 8.951317 | 120-121 |

Produced by FastQC (version 0.11.5)
