## Supplementary material 4 for "Natural and human-mediated drivers of microevolution in Neotropical palms: a historical genomics approach": outLAU_AP13.1_pf_trimmed_masked_sickled_fastqc.html

outLAU\_AP13.1\_pf\_trimmed\_masked\_sickled.fastq FastQC Report 

FastQC Report

mar. 25 juin 2019  
outLAU\_AP13.1\_pf\_trimmed\_masked\_sickled.fastq

### Summary

- Basic Statistics
- Per base sequence quality
- Per tile sequence quality
- Per sequence quality scores
- Per base sequence content
- Per sequence GC content
- Per base N content
- Sequence Length Distribution
- Sequence Duplication Levels
- Overrepresented sequences
- Adapter Content
- Kmer Content

### Basic Statistics

| Measure | Value |
| --- | --- |
| Filename | outLAU\_AP13.1\_pf\_trimmed\_masked\_sickled.fastq |
| File type | Conventional base calls |
| Encoding | Sanger / Illumina 1.9 |
| Total Sequences | 1886371 |
| Sequences flagged as poor quality | 0 |
| Sequence length | 100-131 |
| %GC | 44 |

### Per base sequence quality

### Per tile sequence quality

### Per sequence quality scores

### Per base sequence content

### Per sequence GC content

### Per base N content

### Sequence Length Distribution

### Sequence Duplication Levels

### Overrepresented sequences

No overrepresented sequences

### Adapter Content

### Kmer Content

| Sequence | Count | PValue | Obs/Exp Max | Max Obs/Exp Position |
| --- | --- | --- | --- | --- |
| CTGGTTG | 1695 | 0.0 | 16.344131 | 5 |
| ATCTGGT | 2330 | 0.0 | 12.635345 | 3 |
| TGGTTGG | 2670 | 0.0 | 11.48864 | 6 |
| TCTGGTT | 2430 | 0.0 | 11.158579 | 4 |
| GCACACG | 980 | 0.0 | 10.083293 | 124-125 |
| CGGAAGA | 2175 | 0.0 | 9.927901 | 124-125 |
| CGTAAGG | 290 | 0.0045872345 | 9.482403 | 114-115 |
| ACGTCTG | 810 | 8.913048E-11 | 9.363395 | 122-123 |
| ACACGTC | 815 | 1.3096724E-10 | 9.182704 | 120-121 |
| CACACGT | 945 | 1.6370905E-11 | 8.9076 | 124-125 |
| TCGGAAG | 2275 | 0.0 | 8.848018 | 124-125 |
| TTCCGAT | 2510 | 0.0 | 8.645082 | 1 |
| CAACCAG | 4300 | 0.0 | 8.426194 | 124-125 |
| CACGTCT | 820 | 9.5041287E-7 | 7.388291 | 120-121 |
| CGTCTGA | 995 | 4.062349E-8 | 7.2594876 | 122-123 |
| CCAACCA | 5415 | 0.0 | 7.164273 | 124-125 |
| GGTTGGA | 1980 | 2.1786218E-7 | 6.259045 | 7 |
| GATCTGG | 5385 | 0.0 | 6.2275896 | 2 |
| AGCACAC | 1515 | 1.7262209E-9 | 6.1981173 | 122-123 |
| ACCAGAT | 4310 | 0.0 | 5.9440913 | 124-125 |

Produced by FastQC (version 0.11.5)
