## Supplementary material 4 for "Natural and human-mediated drivers of microevolution in Neotropical palms: a historical genomics approach": outLAU_AP13.2_pf_trimmed_masked_sickled_fastqc.html

outLAU\_AP13.2\_pf\_trimmed\_masked\_sickled.fastq FastQC Report 

FastQC Report

mar. 25 juin 2019  
outLAU\_AP13.2\_pf\_trimmed\_masked\_sickled.fastq

### Summary

- Basic Statistics
- Per base sequence quality
- Per tile sequence quality
- Per sequence quality scores
- Per base sequence content
- Per sequence GC content
- Per base N content
- Sequence Length Distribution
- Sequence Duplication Levels
- Overrepresented sequences
- Adapter Content
- Kmer Content

### Basic Statistics

| Measure | Value |
| --- | --- |
| Filename | outLAU\_AP13.2\_pf\_trimmed\_masked\_sickled.fastq |
| File type | Conventional base calls |
| Encoding | Sanger / Illumina 1.9 |
| Total Sequences | 1886371 |
| Sequences flagged as poor quality | 0 |
| Sequence length | 100-131 |
| %GC | 45 |

### Per base sequence quality

### Per tile sequence quality

### Per sequence quality scores

### Per base sequence content

### Per sequence GC content

### Per base N content

### Sequence Length Distribution

### Sequence Duplication Levels

### Overrepresented sequences

No overrepresented sequences

### Adapter Content

### Kmer Content

| Sequence | Count | PValue | Obs/Exp Max | Max Obs/Exp Position |
| --- | --- | --- | --- | --- |
| CTGGTTG | 1360 | 0.0 | 18.611828 | 5 |
| ATCTGGT | 1970 | 0.0 | 13.6462 | 3 |
| TGGTTGG | 2135 | 0.0 | 12.425375 | 6 |
| CGGAAGA | 2110 | 0.0 | 12.186666 | 124-125 |
| TCGGAAG | 2180 | 0.0 | 10.441787 | 124-125 |
| CGTGTAG | 475 | 5.933991E-6 | 10.380264 | 122-123 |
| AGCGTCG | 835 | 1.2732926E-11 | 10.333646 | 122-123 |
| CAACCAG | 4125 | 0.0 | 10.014739 | 124-125 |
| CGTCGTG | 800 | 4.0563464E-10 | 10.011547 | 124-125 |
| TCTGGTT | 2540 | 0.0 | 9.943555 | 4 |
| GAGCGTC | 915 | 7.2759576E-12 | 9.879213 | 122-123 |
| GCGTCGT | 765 | 2.0940206E-8 | 9.367529 | 124-125 |
| AGAGCGT | 925 | 1.4004399E-8 | 8.225174 | 120-121 |
| GGTTGGT | 1345 | 2.4558176E-8 | 7.9688478 | 7 |
| CCAACCA | 5295 | 0.0 | 7.72224 | 124-125 |
| GTCGTGT | 610 | 7.6180906E-4 | 7.601519 | 124-125 |
| TCGTGTA | 535 | 0.0029338838 | 7.4847865 | 120-121 |
| AAGAGCG | 1375 | 1.2732926E-10 | 7.3577676 | 124-125 |
| AACCAGA | 5080 | 0.0 | 7.0533066 | 124-125 |
| GTAGGGA | 910 | 2.2971379E-5 | 6.9484415 | 124-125 |

Produced by FastQC (version 0.11.5)
