## Supplementary material 4 for "Natural and human-mediated drivers of microevolution in Neotropical palms: a historical genomics approach": outLAU_AP14.1_pf_trimmed_masked_sickled_fastqc.html

outLAU\_AP14.1\_pf\_trimmed\_masked\_sickled.fastq FastQC Report 

FastQC Report

mar. 25 juin 2019  
outLAU\_AP14.1\_pf\_trimmed\_masked\_sickled.fastq

### Summary

- Basic Statistics
- Per base sequence quality
- Per tile sequence quality
- Per sequence quality scores
- Per base sequence content
- Per sequence GC content
- Per base N content
- Sequence Length Distribution
- Sequence Duplication Levels
- Overrepresented sequences
- Adapter Content
- Kmer Content

### Basic Statistics

| Measure | Value |
| --- | --- |
| Filename | outLAU\_AP14.1\_pf\_trimmed\_masked\_sickled.fastq |
| File type | Conventional base calls |
| Encoding | Sanger / Illumina 1.9 |
| Total Sequences | 1037205 |
| Sequences flagged as poor quality | 0 |
| Sequence length | 100-131 |
| %GC | 44 |

### Per base sequence quality

### Per tile sequence quality

### Per sequence quality scores

### Per base sequence content

### Per sequence GC content

### Per base N content

### Sequence Length Distribution

### Sequence Duplication Levels

### Overrepresented sequences

No overrepresented sequences

### Adapter Content

### Kmer Content

| Sequence | Count | PValue | Obs/Exp Max | Max Obs/Exp Position |
| --- | --- | --- | --- | --- |
| TGTGTCC | 565 | 0.0 | 42.5505 | 6 |
| CTGTGTC | 670 | 0.0 | 33.256615 | 5 |
| TCTGTGT | 825 | 0.0 | 27.685944 | 4 |
| ATCTGTG | 1005 | 0.0 | 23.28909 | 3 |
| GTGTCCG | 325 | 7.239578E-10 | 21.665619 | 7 |
| GATCTGT | 1155 | 0.0 | 19.23404 | 2 |
| GTGTCCA | 495 | 7.885319E-9 | 15.41031 | 7 |
| GTGTCCT | 455 | 7.8906487E-7 | 14.1858225 | 7 |
| GTGTCCC | 350 | 2.5225285E-4 | 13.41205 | 7 |
| CTTCCGA | 535 | 7.018451E-5 | 10.8758745 | 1 |
| GGACACA | 1160 | 0.0 | 8.775982 | 122-123 |
| GACACAG | 1100 | 0.0 | 8.593624 | 122-123 |
| ACACAGA | 1140 | 1.8189894E-12 | 8.414914 | 124-125 |
| CGATCTG | 3480 | 0.0 | 7.18964 | 1 |
| TTCCGAT | 1155 | 5.3781212E-5 | 7.086225 | 2 |
| CACAGAT | 1455 | 4.129106E-10 | 6.593128 | 124-125 |
| CGGAAGA | 755 | 6.3238613E-4 | 6.3529816 | 124-125 |
| TCGGAAG | 735 | 0.0038226123 | 5.9359407 | 122-123 |
| TGAGGGT | 650 | 0.005013621 | 5.793047 | 84-85 |
| TGCATGA | 1415 | 7.9447345E-4 | 5.7569118 | 1 |

Produced by FastQC (version 0.11.5)
