## Supplementary material 4 for "Natural and human-mediated drivers of microevolution in Neotropical palms: a historical genomics approach": outLAU_AP14.2_pf_trimmed_masked_sickled_fastqc.html

outLAU\_AP14.2\_pf\_trimmed\_masked\_sickled.fastq FastQC Report 

FastQC Report

mar. 25 juin 2019  
outLAU\_AP14.2\_pf\_trimmed\_masked\_sickled.fastq

### Summary

- Basic Statistics
- Per base sequence quality
- Per tile sequence quality
- Per sequence quality scores
- Per base sequence content
- Per sequence GC content
- Per base N content
- Sequence Length Distribution
- Sequence Duplication Levels
- Overrepresented sequences
- Adapter Content
- Kmer Content

### Basic Statistics

| Measure | Value |
| --- | --- |
| Filename | outLAU\_AP14.2\_pf\_trimmed\_masked\_sickled.fastq |
| File type | Conventional base calls |
| Encoding | Sanger / Illumina 1.9 |
| Total Sequences | 1037205 |
| Sequences flagged as poor quality | 0 |
| Sequence length | 100-131 |
| %GC | 44 |

### Per base sequence quality

### Per tile sequence quality

### Per sequence quality scores

### Per base sequence content

### Per sequence GC content

### Per base N content

### Sequence Length Distribution

### Sequence Duplication Levels

### Overrepresented sequences

No overrepresented sequences

### Adapter Content

### Kmer Content

| Sequence | Count | PValue | Obs/Exp Max | Max Obs/Exp Position |
| --- | --- | --- | --- | --- |
| TGTGTCC | 635 | 0.0 | 38.084038 | 6 |
| CTGTGTC | 700 | 0.0 | 33.758972 | 5 |
| TCTGTGT | 725 | 0.0 | 30.988527 | 4 |
| GTGTCCG | 255 | 8.0035534E-11 | 25.836325 | 7 |
| ATCTGTG | 950 | 0.0 | 24.175772 | 3 |
| GATCTGT | 1245 | 0.0 | 18.383602 | 2 |
| GTGTCCT | 420 | 1.09139364E-10 | 18.30073 | 7 |
| GTGTCCA | 455 | 3.43789E-10 | 16.89298 | 7 |
| GTGTCCC | 295 | 1.0388107E-4 | 14.888729 | 7 |
| TGTCCGA | 355 | 5.0384476E-4 | 12.356908 | 8 |
| CACAGAT | 1260 | 0.0 | 10.068581 | 124-125 |
| CGGGGAA | 255 | 0.00300079 | 9.981473 | 88-89 |
| ACACAGA | 1080 | 1.8189894E-12 | 9.466302 | 120-121 |
| GACACAG | 1120 | 1.8189894E-12 | 9.427879 | 122-123 |
| TGGTGTA | 485 | 0.0066317176 | 9.065928 | 6 |
| GGACACA | 1155 | 8.731149E-11 | 8.380338 | 122-123 |
| ACAGATC | 1445 | 5.456968E-12 | 7.916374 | 122-123 |
| ACTGTTA | 435 | 7.6375477E-4 | 7.5994363 | 46-47 |
| CGATCTG | 3145 | 0.0 | 7.39956 | 1 |

Produced by FastQC (version 0.11.5)
