## Supplementary material 4 for "Natural and human-mediated drivers of microevolution in Neotropical palms: a historical genomics approach": outLAU_AP17.1_pf_trimmed_masked_sickled_fastqc.html

outLAU\_AP17.1\_pf\_trimmed\_masked\_sickled.fastq FastQC Report 

FastQC Report

mar. 25 juin 2019  
outLAU\_AP17.1\_pf\_trimmed\_masked\_sickled.fastq

### Summary

- Basic Statistics
- Per base sequence quality
- Per tile sequence quality
- Per sequence quality scores
- Per base sequence content
- Per sequence GC content
- Per base N content
- Sequence Length Distribution
- Sequence Duplication Levels
- Overrepresented sequences
- Adapter Content
- Kmer Content

### Basic Statistics

| Measure | Value |
| --- | --- |
| Filename | outLAU\_AP17.1\_pf\_trimmed\_masked\_sickled.fastq |
| File type | Conventional base calls |
| Encoding | Sanger / Illumina 1.9 |
| Total Sequences | 889755 |
| Sequences flagged as poor quality | 0 |
| Sequence length | 100-131 |
| %GC | 44 |

### Per base sequence quality

### Per tile sequence quality

### Per sequence quality scores

### Per base sequence content

### Per sequence GC content

### Per base N content

### Sequence Length Distribution

### Sequence Duplication Levels

### Overrepresented sequences

No overrepresented sequences

### Adapter Content

### Kmer Content

| Sequence | Count | PValue | Obs/Exp Max | Max Obs/Exp Position |
| --- | --- | --- | --- | --- |
| TTAAGGT | 575 | 0.0 | 22.142324 | 6 |
| CTTAAGG | 615 | 0.0 | 18.727322 | 5 |
| TCTTAAG | 715 | 0.0 | 16.941362 | 4 |
| CGATCTT | 810 | 0.0 | 16.372133 | 1 |
| CTAGCCG | 135 | 0.0072547356 | 13.582492 | 32-33 |
| TAAGGTC | 335 | 0.0021581738 | 12.67067 | 7 |
| GATCTTA | 1075 | 1.0913936E-11 | 11.245465 | 2 |
| ATCTTAA | 1300 | 3.8198777E-11 | 9.774135 | 3 |
| TTCCGAT | 965 | 3.2254866E-7 | 9.369827 | 1 |
| GGTACCT | 350 | 0.0032250234 | 8.451642 | 116-117 |
| TAAGGTT | 695 | 0.006247653 | 7.8524294 | 7 |
| CCGAAAT | 370 | 0.007949614 | 7.6448035 | 86-87 |
| CCTTAAG | 1140 | 9.094947E-11 | 7.3847356 | 122-123 |
| GGGGTTC | 430 | 0.0039704223 | 7.2562203 | 60-61 |
| ACCTTAA | 1140 | 3.3574906E-8 | 6.536189 | 124-125 |
| AAGATCG | 995 | 6.4935866E-6 | 6.1271176 | 124-125 |
| CTTAAGA | 1120 | 1.231565E-6 | 6.0132847 | 122-123 |
| CGGAAGA | 710 | 0.0067190207 | 5.641751 | 120-121 |
| TTAAGAT | 1400 | 1.82449E-6 | 5.322325 | 124-125 |
| TAAGATC | 1280 | 6.5853295E-5 | 5.0274806 | 124-125 |

Produced by FastQC (version 0.11.5)
