## Supplementary material 4 for "Natural and human-mediated drivers of microevolution in Neotropical palms: a historical genomics approach": outLAU_AP17.2_pf_trimmed_masked_sickled_fastqc.html

outLAU\_AP17.2\_pf\_trimmed\_masked\_sickled.fastq FastQC Report 

FastQC Report

mar. 25 juin 2019  
outLAU\_AP17.2\_pf\_trimmed\_masked\_sickled.fastq

### Summary

- Basic Statistics
- Per base sequence quality
- Per tile sequence quality
- Per sequence quality scores
- Per base sequence content
- Per sequence GC content
- Per base N content
- Sequence Length Distribution
- Sequence Duplication Levels
- Overrepresented sequences
- Adapter Content
- Kmer Content

### Basic Statistics

| Measure | Value |
| --- | --- |
| Filename | outLAU\_AP17.2\_pf\_trimmed\_masked\_sickled.fastq |
| File type | Conventional base calls |
| Encoding | Sanger / Illumina 1.9 |
| Total Sequences | 889755 |
| Sequences flagged as poor quality | 0 |
| Sequence length | 100-131 |
| %GC | 44 |

### Per base sequence quality

### Per tile sequence quality

### Per sequence quality scores

### Per base sequence content

### Per sequence GC content

### Per base N content

### Sequence Length Distribution

### Sequence Duplication Levels

### Overrepresented sequences

No overrepresented sequences

### Adapter Content

### Kmer Content

| Sequence | Count | PValue | Obs/Exp Max | Max Obs/Exp Position |
| --- | --- | --- | --- | --- |
| CTTCCGA | 430 | 3.686364E-8 | 15.778922 | 1 |
| CTTAAGG | 545 | 8.128691E-6 | 11.520367 | 5 |
| TTAAGGT | 545 | 8.177147E-6 | 11.5142145 | 6 |
| CGATCTT | 630 | 4.4787237E-5 | 9.872262 | 1 |
| AGAGCGT | 330 | 0.0034707547 | 9.807151 | 124-125 |
| TCTTAAG | 655 | 6.283405E-5 | 9.572011 | 4 |
| TTAAAGC | 485 | 0.005252697 | 9.326372 | 1 |
| CCTTAAG | 1060 | 3.4197E-10 | 8.208337 | 122-123 |
| TAAGATC | 1230 | 4.4055923E-9 | 7.2357635 | 124-125 |
| CTTAAGA | 1100 | 4.8192305E-8 | 7.190774 | 122-123 |
| ACCTTAA | 1085 | 1.945773E-7 | 7.0841975 | 124-125 |
| GATCTTA | 890 | 0.001758641 | 7.024582 | 2 |
| AACCTAG | 480 | 0.0019017449 | 6.9736557 | 48-49 |
| AAGAGCG | 585 | 0.007885425 | 6.760557 | 122-123 |
| ATCTTAA | 1115 | 3.6614804E-4 | 6.644206 | 3 |
| AAGATCG | 1045 | 4.10124E-5 | 6.1939898 | 124-125 |
| TCGGAAG | 685 | 0.0076947706 | 6.1010222 | 118-119 |
| TTAAGAT | 1590 | 1.12539055E-7 | 5.851908 | 124-125 |

Produced by FastQC (version 0.11.5)
