## Supplementary material 4 for "Natural and human-mediated drivers of microevolution in Neotropical palms: a historical genomics approach": outLAU_AP18.1_pf_trimmed_masked_sickled_fastqc.html

outLAU\_AP18.1\_pf\_trimmed\_masked\_sickled.fastq FastQC Report 

FastQC Report

mar. 25 juin 2019  
outLAU\_AP18.1\_pf\_trimmed\_masked\_sickled.fastq

### Summary

- Basic Statistics
- Per base sequence quality
- Per tile sequence quality
- Per sequence quality scores
- Per base sequence content
- Per sequence GC content
- Per base N content
- Sequence Length Distribution
- Sequence Duplication Levels
- Overrepresented sequences
- Adapter Content
- Kmer Content

### Basic Statistics

| Measure | Value |
| --- | --- |
| Filename | outLAU\_AP18.1\_pf\_trimmed\_masked\_sickled.fastq |
| File type | Conventional base calls |
| Encoding | Sanger / Illumina 1.9 |
| Total Sequences | 797634 |
| Sequences flagged as poor quality | 0 |
| Sequence length | 100-131 |
| %GC | 43 |

### Per base sequence quality

### Per tile sequence quality

### Per sequence quality scores

### Per base sequence content

### Per sequence GC content

### Per base N content

### Sequence Length Distribution

### Sequence Duplication Levels

### Overrepresented sequences

No overrepresented sequences

### Adapter Content

### Kmer Content

| Sequence | Count | PValue | Obs/Exp Max | Max Obs/Exp Position |
| --- | --- | --- | --- | --- |
| CTTACGA | 275 | 0.0 | 52.729774 | 5 |
| TCTTACG | 290 | 0.0 | 52.069176 | 4 |
| TTACGAT | 320 | 0.0 | 45.31465 | 6 |
| ATCTTAC | 570 | 0.0 | 28.568676 | 3 |
| TACGATA | 165 | 9.872843E-6 | 25.624346 | 7 |
| CGATCTT | 670 | 0.0 | 24.253654 | 1 |
| TACGATT | 190 | 7.519524E-4 | 19.073761 | 7 |
| GATCTTA | 850 | 0.0 | 17.03243 | 2 |
| GCACACG | 250 | 4.189751E-7 | 15.003738 | 124-125 |
| TACGATG | 300 | 9.675621E-4 | 14.09339 | 7 |
| ACTGTCT | 380 | 0.005704321 | 11.12636 | 4 |
| TCGTAAG | 870 | 0.0 | 10.190627 | 124-125 |
| CGTAAGA | 765 | 9.094947E-12 | 9.806365 | 124-125 |
| CACACGT | 285 | 0.004243102 | 9.571763 | 124-125 |
| AATCGTA | 395 | 8.200931E-5 | 9.342766 | 120-121 |
| TATCGTA | 335 | 0.0016681927 | 9.088289 | 122-123 |
| ATCGTAA | 1005 | 0.0 | 8.821736 | 124-125 |
| AAGATCG | 1070 | 1.8189894E-12 | 8.285837 | 124-125 |
| GATCGTA | 560 | 4.2248914E-5 | 7.9159336 | 124-125 |
| GTAAGAT | 1030 | 6.91216E-11 | 7.882399 | 122-123 |

Produced by FastQC (version 0.11.5)
