## Supplementary material 4 for "Natural and human-mediated drivers of microevolution in Neotropical palms: a historical genomics approach": outLAU_AP18.2_pf_trimmed_masked_sickled_fastqc.html

outLAU\_AP18.2\_pf\_trimmed\_masked\_sickled.fastq FastQC Report 

FastQC Report

mar. 25 juin 2019  
outLAU\_AP18.2\_pf\_trimmed\_masked\_sickled.fastq

### Summary

- Basic Statistics
- Per base sequence quality
- Per tile sequence quality
- Per sequence quality scores
- Per base sequence content
- Per sequence GC content
- Per base N content
- Sequence Length Distribution
- Sequence Duplication Levels
- Overrepresented sequences
- Adapter Content
- Kmer Content

### Basic Statistics

| Measure | Value |
| --- | --- |
| Filename | outLAU\_AP18.2\_pf\_trimmed\_masked\_sickled.fastq |
| File type | Conventional base calls |
| Encoding | Sanger / Illumina 1.9 |
| Total Sequences | 797634 |
| Sequences flagged as poor quality | 0 |
| Sequence length | 100-131 |
| %GC | 43 |

### Per base sequence quality

### Per tile sequence quality

### Per sequence quality scores

### Per base sequence content

### Per sequence GC content

### Per base N content

### Sequence Length Distribution

### Sequence Duplication Levels

### Overrepresented sequences

No overrepresented sequences

### Adapter Content

### Kmer Content

| Sequence | Count | PValue | Obs/Exp Max | Max Obs/Exp Position |
| --- | --- | --- | --- | --- |
| TCTTACG | 205 | 0.0 | 47.037117 | 4 |
| CTTACGA | 245 | 0.0 | 44.087376 | 5 |
| TTACGAT | 310 | 0.0 | 34.8734 | 6 |
| TACGATA | 180 | 3.0986885E-5 | 22.103731 | 7 |
| CGATCTT | 535 | 0.0 | 18.926765 | 1 |
| ATCTTAC | 510 | 0.0 | 18.897043 | 3 |
| AGAGCGT | 335 | 9.6250005E-8 | 14.592585 | 124-125 |
| CTTGCGT | 150 | 0.008510915 | 13.257977 | 98-99 |
| GAGCGTC | 280 | 5.6863642E-5 | 13.094238 | 124-125 |
| TCGTAAG | 775 | 0.0 | 11.56423 | 124-125 |
| GATCTTA | 900 | 1.9826984E-10 | 11.319456 | 2 |
| TATCGTA | 335 | 3.619909E-4 | 10.734155 | 122-123 |
| ATCGTAA | 930 | 0.0 | 10.512937 | 124-125 |
| CGTAAGA | 720 | 1.7553248E-9 | 9.988728 | 122-123 |
| AAGAGCG | 450 | 6.662937E-5 | 9.521497 | 120-121 |
| AATCGTA | 375 | 0.0015604923 | 9.154785 | 118-119 |
| AAGATCG | 980 | 3.6379788E-11 | 9.145183 | 124-125 |
| GGTTCCG | 330 | 0.0033670247 | 8.411278 | 56-57 |
| GTAAGAT | 905 | 9.826181E-9 | 8.388324 | 122-123 |
| TCGGAAG | 675 | 4.717236E-5 | 7.845766 | 124-125 |

Produced by FastQC (version 0.11.5)
