## Supplementary material 4 for "Natural and human-mediated drivers of microevolution in Neotropical palms: a historical genomics approach": outLAU_AP20.1_pf_trimmed_masked_sickled_fastqc.html

outLAU\_AP20.1\_pf\_trimmed\_masked\_sickled.fastq FastQC Report 

FastQC Report

mar. 25 juin 2019  
outLAU\_AP20.1\_pf\_trimmed\_masked\_sickled.fastq

### Summary

- Basic Statistics
- Per base sequence quality
- Per tile sequence quality
- Per sequence quality scores
- Per base sequence content
- Per sequence GC content
- Per base N content
- Sequence Length Distribution
- Sequence Duplication Levels
- Overrepresented sequences
- Adapter Content
- Kmer Content

### Basic Statistics

| Measure | Value |
| --- | --- |
| Filename | outLAU\_AP20.1\_pf\_trimmed\_masked\_sickled.fastq |
| File type | Conventional base calls |
| Encoding | Sanger / Illumina 1.9 |
| Total Sequences | 1444166 |
| Sequences flagged as poor quality | 0 |
| Sequence length | 100-131 |
| %GC | 44 |

### Per base sequence quality

### Per tile sequence quality

### Per sequence quality scores

### Per base sequence content

### Per sequence GC content

### Per base N content

### Sequence Length Distribution

### Sequence Duplication Levels

### Overrepresented sequences

No overrepresented sequences

### Adapter Content

### Kmer Content

| Sequence | Count | PValue | Obs/Exp Max | Max Obs/Exp Position |
| --- | --- | --- | --- | --- |
| CTAACAC | 750 | 0.0 | 28.80896 | 5 |
| TAACACC | 840 | 0.0 | 27.157064 | 6 |
| ATCTAAC | 1485 | 0.0 | 16.54507 | 3 |
| CGATCTA | 1430 | 0.0 | 15.874366 | 1 |
| TCTAACA | 1585 | 0.0 | 14.764869 | 4 |
| ACACGTC | 540 | 0.0 | 13.376265 | 124-125 |
| CACACGT | 635 | 0.0 | 11.916762 | 124-125 |
| GCACACG | 740 | 0.0 | 10.625444 | 122-123 |
| GATCTAA | 2590 | 0.0 | 9.716267 | 2 |
| AACACCT | 1010 | 6.395203E-8 | 9.522048 | 7 |
| CGGAAGA | 1540 | 0.0 | 9.157406 | 124-125 |
| ACCTCGG | 530 | 0.006759948 | 9.045349 | 3 |
| CACGTCT | 530 | 1.903906E-5 | 8.436781 | 124-125 |
| CTTCCGA | 710 | 9.2562573E-4 | 8.413767 | 1 |
| AACACCG | 675 | 0.0052052774 | 8.014391 | 7 |
| AACACCC | 715 | 0.008718646 | 7.566033 | 7 |
| TGGTCGT | 420 | 0.003365147 | 7.3806005 | 74-75 |
| GTGTTAG | 2060 | 0.0 | 7.3019094 | 122-123 |
| GTTAGAT | 2615 | 0.0 | 6.9712954 | 124-125 |
| TCGGAAG | 1630 | 0.0 | 6.711404 | 122-123 |

Produced by FastQC (version 0.11.5)
