## Supplementary material 4 for "Natural and human-mediated drivers of microevolution in Neotropical palms: a historical genomics approach": outLAU_AP20.2_pf_trimmed_masked_sickled_fastqc.html

outLAU\_AP20.2\_pf\_trimmed\_masked\_sickled.fastq FastQC Report 

FastQC Report

mar. 25 juin 2019  
outLAU\_AP20.2\_pf\_trimmed\_masked\_sickled.fastq

### Summary

- Basic Statistics
- Per base sequence quality
- Per tile sequence quality
- Per sequence quality scores
- Per base sequence content
- Per sequence GC content
- Per base N content
- Sequence Length Distribution
- Sequence Duplication Levels
- Overrepresented sequences
- Adapter Content
- Kmer Content

### Basic Statistics

| Measure | Value |
| --- | --- |
| Filename | outLAU\_AP20.2\_pf\_trimmed\_masked\_sickled.fastq |
| File type | Conventional base calls |
| Encoding | Sanger / Illumina 1.9 |
| Total Sequences | 1444166 |
| Sequences flagged as poor quality | 0 |
| Sequence length | 100-131 |
| %GC | 44 |

### Per base sequence quality

### Per tile sequence quality

### Per sequence quality scores

### Per base sequence content

### Per sequence GC content

### Per base N content

### Sequence Length Distribution

### Sequence Duplication Levels

### Overrepresented sequences

No overrepresented sequences

### Adapter Content

### Kmer Content

| Sequence | Count | PValue | Obs/Exp Max | Max Obs/Exp Position |
| --- | --- | --- | --- | --- |
| CTAACAC | 710 | 0.0 | 35.64729 | 5 |
| TAACACC | 835 | 0.0 | 30.338678 | 6 |
| CGATCTA | 1345 | 0.0 | 18.982347 | 1 |
| ATCTAAC | 1400 | 0.0 | 18.411152 | 3 |
| TCTAACA | 1510 | 0.0 | 17.472301 | 4 |
| GTCGTGT | 490 | 1.2732926E-11 | 14.642841 | 124-125 |
| GCGTCGT | 480 | 1.2732926E-11 | 14.56791 | 122-123 |
| TCGTGTA | 435 | 1.0961594E-7 | 12.613238 | 124-125 |
| AGAGCGT | 750 | 1.0913936E-11 | 11.25489 | 124-125 |
| AAGAGCG | 1120 | 0.0 | 10.928298 | 124-125 |
| GATCTAA | 2460 | 0.0 | 10.682258 | 2 |
| GAGCGTC | 705 | 1.1586963E-9 | 10.232572 | 120-121 |
| AACACCG | 615 | 3.599641E-5 | 10.0716305 | 7 |
| AGCGTCG | 575 | 5.145903E-7 | 10.014973 | 122-123 |
| AACACCC | 795 | 5.88916E-7 | 9.916151 | 7 |
| CGGAAGA | 1470 | 0.0 | 9.47478 | 124-125 |
| GTGTTAG | 1780 | 0.0 | 9.247347 | 124-125 |
| CGTCGTG | 610 | 8.483121E-6 | 8.994686 | 124-125 |
| TCGGAAG | 1530 | 0.0 | 8.602964 | 122-123 |
| AACACCT | 850 | 1.3728146E-4 | 7.949586 | 7 |

Produced by FastQC (version 0.11.5)
