## Supplementary material 4 for "Natural and human-mediated drivers of microevolution in Neotropical palms: a historical genomics approach": outLAU_AS01.1_pf_trimmed_masked_sickled_fastqc.html

outLAU\_AS01.1\_pf\_trimmed\_masked\_sickled.fastq FastQC Report 

FastQC Report

mar. 25 juin 2019  
outLAU\_AS01.1\_pf\_trimmed\_masked\_sickled.fastq

### Summary

- Basic Statistics
- Per base sequence quality
- Per tile sequence quality
- Per sequence quality scores
- Per base sequence content
- Per sequence GC content
- Per base N content
- Sequence Length Distribution
- Sequence Duplication Levels
- Overrepresented sequences
- Adapter Content
- Kmer Content

### Basic Statistics

| Measure | Value |
| --- | --- |
| Filename | outLAU\_AS01.1\_pf\_trimmed\_masked\_sickled.fastq |
| File type | Conventional base calls |
| Encoding | Sanger / Illumina 1.9 |
| Total Sequences | 665601 |
| Sequences flagged as poor quality | 0 |
| Sequence length | 100-131 |
| %GC | 45 |

### Per base sequence quality

### Per tile sequence quality

### Per sequence quality scores

### Per base sequence content

### Per sequence GC content

### Per base N content

### Sequence Length Distribution

### Sequence Duplication Levels

### Overrepresented sequences

No overrepresented sequences

### Adapter Content

### Kmer Content

| Sequence | Count | PValue | Obs/Exp Max | Max Obs/Exp Position |
| --- | --- | --- | --- | --- |
| CTCTAAG | 530 | 7.3123374E-10 | 16.014746 | 5 |
| TCTAAGT | 555 | 1.373337E-9 | 15.316737 | 6 |
| CCGTAGC | 155 | 9.2751655E-4 | 14.173508 | 88-89 |
| ATCTCTA | 800 | 1.4551915E-9 | 12.131931 | 3 |
| AACGGTT | 200 | 0.006004444 | 11.050824 | 94-95 |
| TCTCTAA | 795 | 2.1391679E-7 | 10.672424 | 4 |
| CGATCTC | 875 | 7.3951014E-8 | 10.364769 | 1 |
| GATCTCT | 1130 | 2.888808E-6 | 8.044183 | 2 |
| TACTTAG | 555 | 5.497752E-5 | 7.7501817 | 118-119 |
| AACTTAG | 580 | 7.442737E-5 | 7.5635433 | 124-125 |
| TGAAGGC | 725 | 0.009164298 | 7.522697 | 2 |
| ACTTAGA | 1240 | 1.0913936E-11 | 7.34771 | 124-125 |
| CGGAAGA | 665 | 3.9999987E-4 | 6.596775 | 124-125 |
| TCGGAAG | 720 | 1.4732504E-4 | 6.5615354 | 124-125 |
| TTAGAGA | 1290 | 9.557516E-8 | 5.899288 | 118-119 |
| TAGAGAT | 1390 | 5.5610144E-8 | 5.782273 | 122-123 |
| CTTAGAG | 1125 | 8.15169E-6 | 5.6991625 | 124-125 |

Produced by FastQC (version 0.11.5)
