## Supplementary material 4 for "Natural and human-mediated drivers of microevolution in Neotropical palms: a historical genomics approach": outLAU_AS01.2_pf_trimmed_masked_sickled_fastqc.html

outLAU\_AS01.2\_pf\_trimmed\_masked\_sickled.fastq FastQC Report 

FastQC Report

mar. 25 juin 2019  
outLAU\_AS01.2\_pf\_trimmed\_masked\_sickled.fastq

### Summary

- Basic Statistics
- Per base sequence quality
- Per tile sequence quality
- Per sequence quality scores
- Per base sequence content
- Per sequence GC content
- Per base N content
- Sequence Length Distribution
- Sequence Duplication Levels
- Overrepresented sequences
- Adapter Content
- Kmer Content

### Basic Statistics

| Measure | Value |
| --- | --- |
| Filename | outLAU\_AS01.2\_pf\_trimmed\_masked\_sickled.fastq |
| File type | Conventional base calls |
| Encoding | Sanger / Illumina 1.9 |
| Total Sequences | 665601 |
| Sequences flagged as poor quality | 0 |
| Sequence length | 100-131 |
| %GC | 45 |

### Per base sequence quality

### Per tile sequence quality

### Per sequence quality scores

### Per base sequence content

### Per sequence GC content

### Per base N content

### Sequence Length Distribution

### Sequence Duplication Levels

### Overrepresented sequences

No overrepresented sequences

### Adapter Content

### Kmer Content

| Sequence | Count | PValue | Obs/Exp Max | Max Obs/Exp Position |
| --- | --- | --- | --- | --- |
| CTCTAAG | 400 | 7.106893E-5 | 12.784653 | 5 |
| CGATCTC | 590 | 1.8172959E-6 | 11.453937 | 1 |
| ACTTAGA | 1110 | 0.0 | 9.946588 | 124-125 |
| CGGAAGA | 640 | 2.323286E-7 | 9.583952 | 124-125 |
| TCTAAGT | 560 | 0.0015806741 | 9.140607 | 6 |
| CAAGGGG | 335 | 0.0029961732 | 8.520252 | 90-91 |
| TACTTAG | 490 | 0.0010028835 | 8.345209 | 124-125 |
| TCTCTAA | 760 | 0.0029795414 | 7.4710636 | 4 |
| GATCTCT | 915 | 2.9969585E-4 | 7.428878 | 2 |
| ATCTCTA | 780 | 0.0038941312 | 7.2691126 | 3 |
| CTTAGAG | 1040 | 1.5777186E-7 | 7.1689415 | 120-121 |
| TCGGAAG | 720 | 8.0517714E-4 | 6.8152547 | 124-125 |
| TTAGAGA | 1300 | 7.980598E-7 | 6.159663 | 122-123 |
| GAGATCG | 1160 | 3.420966E-5 | 5.86761 | 122-123 |
| AGAGATC | 1350 | 2.2400873E-6 | 5.813411 | 120-121 |
| ATCGGAA | 1205 | 4.4308465E-5 | 5.768929 | 124-125 |
| TAGAGAT | 1255 | 1.843954E-5 | 5.7424746 | 122-123 |

Produced by FastQC (version 0.11.5)
