## Supplementary material 4 for "Natural and human-mediated drivers of microevolution in Neotropical palms: a historical genomics approach": outLAU_AS02.1_pf_trimmed_masked_sickled_fastqc.html

outLAU\_AS02.1\_pf\_trimmed\_masked\_sickled.fastq FastQC Report 

FastQC Report

mar. 25 juin 2019  
outLAU\_AS02.1\_pf\_trimmed\_masked\_sickled.fastq

### Summary

- Basic Statistics
- Per base sequence quality
- Per tile sequence quality
- Per sequence quality scores
- Per base sequence content
- Per sequence GC content
- Per base N content
- Sequence Length Distribution
- Sequence Duplication Levels
- Overrepresented sequences
- Adapter Content
- Kmer Content

### Basic Statistics

| Measure | Value |
| --- | --- |
| Filename | outLAU\_AS02.1\_pf\_trimmed\_masked\_sickled.fastq |
| File type | Conventional base calls |
| Encoding | Sanger / Illumina 1.9 |
| Total Sequences | 862278 |
| Sequences flagged as poor quality | 0 |
| Sequence length | 100-131 |
| %GC | 45 |

### Per base sequence quality

### Per tile sequence quality

### Per sequence quality scores

### Per base sequence content

### Per sequence GC content

### Per base N content

### Sequence Length Distribution

### Sequence Duplication Levels

### Overrepresented sequences

No overrepresented sequences

### Adapter Content

### Kmer Content

| Sequence | Count | PValue | Obs/Exp Max | Max Obs/Exp Position |
| --- | --- | --- | --- | --- |
| CTCTATG | 660 | 3.6379788E-12 | 15.623115 | 5 |
| TCTATGA | 930 | 0.0 | 13.048581 | 6 |
| CGATCTC | 975 | 1.8189894E-12 | 12.390809 | 1 |
| ATCTCTA | 1065 | 9.094947E-12 | 11.382454 | 3 |
| TCTCTAT | 1105 | 1.9645086E-10 | 10.41699 | 4 |
| GATCTCT | 1290 | 3.274181E-10 | 9.390505 | 2 |
| CCGACGA | 455 | 0.004661256 | 7.1376166 | 110-111 |
| GAGATCG | 1155 | 1.3328587E-5 | 5.5352755 | 122-123 |

Produced by FastQC (version 0.11.5)
