## Supplementary material 4 for "Natural and human-mediated drivers of microevolution in Neotropical palms: a historical genomics approach": outLAU_AS02.2_pf_trimmed_masked_sickled_fastqc.html

outLAU\_AS02.2\_pf\_trimmed\_masked\_sickled.fastq FastQC Report 

FastQC Report

mar. 25 juin 2019  
outLAU\_AS02.2\_pf\_trimmed\_masked\_sickled.fastq

### Summary

- Basic Statistics
- Per base sequence quality
- Per tile sequence quality
- Per sequence quality scores
- Per base sequence content
- Per sequence GC content
- Per base N content
- Sequence Length Distribution
- Sequence Duplication Levels
- Overrepresented sequences
- Adapter Content
- Kmer Content

### Basic Statistics

| Measure | Value |
| --- | --- |
| Filename | outLAU\_AS02.2\_pf\_trimmed\_masked\_sickled.fastq |
| File type | Conventional base calls |
| Encoding | Sanger / Illumina 1.9 |
| Total Sequences | 862278 |
| Sequences flagged as poor quality | 0 |
| Sequence length | 100-131 |
| %GC | 45 |

### Per base sequence quality

### Per tile sequence quality

### Per sequence quality scores

### Per base sequence content

### Per sequence GC content

### Per base N content

### Sequence Length Distribution

### Sequence Duplication Levels

### Overrepresented sequences

No overrepresented sequences

### Adapter Content

### Kmer Content

| Sequence | Count | PValue | Obs/Exp Max | Max Obs/Exp Position |
| --- | --- | --- | --- | --- |
| CTCTATG | 725 | 0.0 | 15.727795 | 5 |
| ACGACAC | 140 | 0.008782083 | 13.195386 | 68-69 |
| ATCTCTA | 1120 | 0.0 | 13.194009 | 3 |
| CGATCTC | 935 | 0.0 | 12.682394 | 1 |
| TCTCTAT | 1080 | 0.0 | 12.648697 | 4 |
| TCTATGA | 1065 | 0.0 | 12.331604 | 6 |
| GATCTCT | 1180 | 0.0 | 11.555576 | 2 |
| CTTCCGA | 510 | 5.8493264E-5 | 11.071932 | 1 |
| AAGAGCG | 405 | 1.7342552E-4 | 9.946555 | 124-125 |
| CTATGAC | 475 | 0.004026832 | 9.631727 | 7 |
| AACACGA | 265 | 0.00953651 | 8.673503 | 12-13 |
| CTATGAA | 700 | 0.0012374263 | 8.169769 | 7 |
| GGCGAGA | 345 | 0.004892538 | 8.069875 | 82-83 |
| TAGAGAT | 1350 | 1.7315451E-6 | 7.11064 | 1 |
| TTCCGAT | 975 | 0.00460055 | 6.4098883 | 2 |
| CATAGAG | 1080 | 3.8410762E-6 | 6.327573 | 118-119 |
| TCATAGA | 1350 | 6.839946E-8 | 6.3041997 | 120-121 |
| GAGATCG | 1185 | 2.6715308E-4 | 6.2351446 | 3 |
| AGAGATC | 1575 | 1.1394763E-4 | 5.7716746 | 2 |
| ATCGGAA | 1595 | 1.2712329E-4 | 5.7279687 | 6 |

Produced by FastQC (version 0.11.5)
