## Supplementary material 4 for "Natural and human-mediated drivers of microevolution in Neotropical palms: a historical genomics approach": outLAU_AS03.2_pf_trimmed_masked_sickled_fastqc.html

outLAU\_AS03.2\_pf\_trimmed\_masked\_sickled.fastq FastQC Report 

FastQC Report

mar. 25 juin 2019  
outLAU\_AS03.2\_pf\_trimmed\_masked\_sickled.fastq

### Summary

- Basic Statistics
- Per base sequence quality
- Per tile sequence quality
- Per sequence quality scores
- Per base sequence content
- Per sequence GC content
- Per base N content
- Sequence Length Distribution
- Sequence Duplication Levels
- Overrepresented sequences
- Adapter Content
- Kmer Content

### Basic Statistics

| Measure | Value |
| --- | --- |
| Filename | outLAU\_AS03.2\_pf\_trimmed\_masked\_sickled.fastq |
| File type | Conventional base calls |
| Encoding | Sanger / Illumina 1.9 |
| Total Sequences | 408297 |
| Sequences flagged as poor quality | 0 |
| Sequence length | 100-131 |
| %GC | 45 |

### Per base sequence quality

### Per tile sequence quality

### Per sequence quality scores

### Per base sequence content

### Per sequence GC content

### Per base N content

### Sequence Length Distribution

### Sequence Duplication Levels

### Overrepresented sequences

No overrepresented sequences

### Adapter Content

### Kmer Content

| Sequence | Count | PValue | Obs/Exp Max | Max Obs/Exp Position |
| --- | --- | --- | --- | --- |
| AGAGCGT | 140 | 0.001343335 | 17.493162 | 124-125 |
| AAGAGCG | 245 | 0.0040140864 | 11.662107 | 124-125 |
| CGGAAGA | 315 | 2.6266667E-4 | 11.110738 | 120-121 |
| TCTATTG | 470 | 0.0039049126 | 9.665308 | 6 |
| TATGTTT | 635 | 0.0054409807 | 7.972097 | 1 |
| GAGATCG | 490 | 0.00944225 | 7.497069 | 124-125 |

Produced by FastQC (version 0.11.5)
