## Supplementary material 4 for "Natural and human-mediated drivers of microevolution in Neotropical palms: a historical genomics approach": outLAU_AS04.1_pf_trimmed_masked_sickled_fastqc.html

outLAU\_AS04.1\_pf\_trimmed\_masked\_sickled.fastq FastQC Report 

FastQC Report

mar. 25 juin 2019  
outLAU\_AS04.1\_pf\_trimmed\_masked\_sickled.fastq

### Summary

- Basic Statistics
- Per base sequence quality
- Per tile sequence quality
- Per sequence quality scores
- Per base sequence content
- Per sequence GC content
- Per base N content
- Sequence Length Distribution
- Sequence Duplication Levels
- Overrepresented sequences
- Adapter Content
- Kmer Content

### Basic Statistics

| Measure | Value |
| --- | --- |
| Filename | outLAU\_AS04.1\_pf\_trimmed\_masked\_sickled.fastq |
| File type | Conventional base calls |
| Encoding | Sanger / Illumina 1.9 |
| Total Sequences | 910560 |
| Sequences flagged as poor quality | 0 |
| Sequence length | 100-131 |
| %GC | 43 |

### Per base sequence quality

### Per tile sequence quality

### Per sequence quality scores

### Per base sequence content

### Per sequence GC content

### Per base N content

### Sequence Length Distribution

### Sequence Duplication Levels

### Overrepresented sequences

No overrepresented sequences

### Adapter Content

### Kmer Content

| Sequence | Count | PValue | Obs/Exp Max | Max Obs/Exp Position |
| --- | --- | --- | --- | --- |
| CACGTCT | 265 | 1.1712365E-5 | 12.956832 | 122-123 |
| ACGTCTG | 315 | 4.621643E-6 | 12.120675 | 124-125 |
| GCACACG | 370 | 2.8714094E-6 | 11.028854 | 120-121 |
| ACACGTC | 320 | 8.057637E-5 | 10.7298765 | 122-123 |
| CGTCTGA | 395 | 4.9154805E-6 | 10.544568 | 124-125 |
| CACACGT | 375 | 4.00324E-5 | 9.974985 | 120-121 |
| CGAGAGA | 1255 | 0.0 | 9.679858 | 124-125 |
| CCGAGAG | 550 | 3.2798798E-6 | 8.739972 | 122-123 |
| TCGGAAG | 850 | 4.5608977E-7 | 7.201193 | 120-121 |
| TCGAGAG | 820 | 1.1060896E-5 | 6.772527 | 124-125 |
| CGGAAGA | 840 | 1.8512656E-5 | 6.540115 | 122-123 |
| AGCACAC | 620 | 0.001304726 | 6.5322623 | 118-119 |
| AGAGCAC | 850 | 8.6954434E-4 | 5.71681 | 124-125 |

Produced by FastQC (version 0.11.5)
