## Supplementary material 4 for "Natural and human-mediated drivers of microevolution in Neotropical palms: a historical genomics approach": outLAU_AS04.2_pf_trimmed_masked_sickled_fastqc.html

outLAU\_AS04.2\_pf\_trimmed\_masked\_sickled.fastq FastQC Report 

FastQC Report

mar. 25 juin 2019  
outLAU\_AS04.2\_pf\_trimmed\_masked\_sickled.fastq

### Summary

- Basic Statistics
- Per base sequence quality
- Per tile sequence quality
- Per sequence quality scores
- Per base sequence content
- Per sequence GC content
- Per base N content
- Sequence Length Distribution
- Sequence Duplication Levels
- Overrepresented sequences
- Adapter Content
- Kmer Content

### Basic Statistics

| Measure | Value |
| --- | --- |
| Filename | outLAU\_AS04.2\_pf\_trimmed\_masked\_sickled.fastq |
| File type | Conventional base calls |
| Encoding | Sanger / Illumina 1.9 |
| Total Sequences | 910560 |
| Sequences flagged as poor quality | 0 |
| Sequence length | 100-131 |
| %GC | 44 |

### Per base sequence quality

### Per tile sequence quality

### Per sequence quality scores

### Per base sequence content

### Per sequence GC content

### Per base N content

### Sequence Length Distribution

### Sequence Duplication Levels

### Overrepresented sequences

No overrepresented sequences

### Adapter Content

### Kmer Content

| Sequence | Count | PValue | Obs/Exp Max | Max Obs/Exp Position |
| --- | --- | --- | --- | --- |
| AAGAGCG | 520 | 1.2065357E-8 | 11.725597 | 124-125 |
| AGCGTCG | 255 | 0.0065170773 | 10.929743 | 122-123 |
| CGAGAGA | 1205 | 0.0 | 10.457351 | 124-125 |
| CCGAGAG | 480 | 8.4657695E-5 | 9.315334 | 124-125 |
| AGAGCGT | 355 | 0.006092751 | 9.160278 | 124-125 |
| TCGAGAG | 760 | 4.0430295E-6 | 7.8583207 | 122-123 |
| TCGGAAG | 815 | 1.9278286E-6 | 7.61401 | 120-121 |
| GCGAGAG | 535 | 0.0025306782 | 7.597894 | 124-125 |
| CGGAAGA | 820 | 7.078211E-5 | 6.9400277 | 124-125 |
| CATTTAT | 915 | 0.002390153 | 6.8232374 | 1 |
| GAAGAGC | 1335 | 9.813502E-7 | 6.089698 | 124-125 |
| ATCAGGG | 1285 | 0.001886044 | 5.800122 | 4 |
| GAGAGAT | 1615 | 2.7884214E-5 | 5.0338993 | 124-125 |
| GAGATCG | 1330 | 1.5768257E-4 | 5.0156574 | 116-117 |

Produced by FastQC (version 0.11.5)
