## Supplementary material 4 for "Natural and human-mediated drivers of microevolution in Neotropical palms: a historical genomics approach": outLAU_AS08.1_pf_trimmed_masked_sickled_fastqc.html

outLAU\_AS08.1\_pf\_trimmed\_masked\_sickled.fastq FastQC Report 

FastQC Report

mar. 25 juin 2019  
outLAU\_AS08.1\_pf\_trimmed\_masked\_sickled.fastq

### Summary

- Basic Statistics
- Per base sequence quality
- Per tile sequence quality
- Per sequence quality scores
- Per base sequence content
- Per sequence GC content
- Per base N content
- Sequence Length Distribution
- Sequence Duplication Levels
- Overrepresented sequences
- Adapter Content
- Kmer Content

### Basic Statistics

| Measure | Value |
| --- | --- |
| Filename | outLAU\_AS08.1\_pf\_trimmed\_masked\_sickled.fastq |
| File type | Conventional base calls |
| Encoding | Sanger / Illumina 1.9 |
| Total Sequences | 815930 |
| Sequences flagged as poor quality | 0 |
| Sequence length | 100-131 |
| %GC | 43 |

### Per base sequence quality

### Per tile sequence quality

### Per sequence quality scores

### Per base sequence content

### Per sequence GC content

### Per base N content

### Sequence Length Distribution

### Sequence Duplication Levels

### Overrepresented sequences

No overrepresented sequences

### Adapter Content

### Kmer Content

| Sequence | Count | PValue | Obs/Exp Max | Max Obs/Exp Position |
| --- | --- | --- | --- | --- |
| TCGTTTG | 230 | 0.0029205575 | 15.575439 | 6 |
| GCCTTAC | 205 | 0.003194609 | 12.025825 | 124-125 |
| GCACTCG | 205 | 0.00812493 | 10.610256 | 56-57 |
| TCGGAAG | 760 | 4.2200554E-10 | 9.268023 | 124-125 |
| CGGAAGA | 680 | 3.9769882E-5 | 7.2508645 | 124-125 |
| GCAGAGA | 1515 | 0.0 | 7.2064233 | 124-125 |
| AGCAGAG | 1585 | 5.456968E-12 | 6.5902605 | 122-123 |
| CAGCAGA | 1875 | 0.0 | 6.386286 | 124-125 |
| CAGAGAT | 1665 | 1.6370905E-11 | 6.3456736 | 124-125 |
| TATAGTT | 595 | 0.002741681 | 6.115176 | 34-35 |
| GAGATCG | 1245 | 4.4437165E-6 | 5.5933404 | 122-123 |
| CCAGCAG | 945 | 0.0027549022 | 5.2175536 | 124-125 |

Produced by FastQC (version 0.11.5)
