## Supplementary material 4 for "Natural and human-mediated drivers of microevolution in Neotropical palms: a historical genomics approach": outLAU_AS08.2_pf_trimmed_masked_sickled_fastqc.html

outLAU\_AS08.2\_pf\_trimmed\_masked\_sickled.fastq FastQC Report 

FastQC Report

mar. 25 juin 2019  
outLAU\_AS08.2\_pf\_trimmed\_masked\_sickled.fastq

### Summary

- Basic Statistics
- Per base sequence quality
- Per tile sequence quality
- Per sequence quality scores
- Per base sequence content
- Per sequence GC content
- Per base N content
- Sequence Length Distribution
- Sequence Duplication Levels
- Overrepresented sequences
- Adapter Content
- Kmer Content

### Basic Statistics

| Measure | Value |
| --- | --- |
| Filename | outLAU\_AS08.2\_pf\_trimmed\_masked\_sickled.fastq |
| File type | Conventional base calls |
| Encoding | Sanger / Illumina 1.9 |
| Total Sequences | 815930 |
| Sequences flagged as poor quality | 0 |
| Sequence length | 100-131 |
| %GC | 43 |

### Per base sequence quality

### Per tile sequence quality

### Per sequence quality scores

### Per base sequence content

### Per sequence GC content

### Per base N content

### Sequence Length Distribution

### Sequence Duplication Levels

### Overrepresented sequences

No overrepresented sequences

### Adapter Content

### Kmer Content

| Sequence | Count | PValue | Obs/Exp Max | Max Obs/Exp Position |
| --- | --- | --- | --- | --- |
| GTCGTGT | 205 | 8.7157096E-4 | 14.290152 | 124-125 |
| TCGGAAG | 740 | 9.094947E-12 | 11.310739 | 124-125 |
| TCTCTGC | 775 | 3.7797327E-7 | 10.240427 | 4 |
| CGGAAGA | 645 | 1.8592073E-7 | 9.732495 | 124-125 |
| CTCTGCT | 815 | 7.701159E-6 | 9.061665 | 5 |
| CAGCAGA | 1845 | 0.0 | 8.619457 | 124-125 |
| GCAGAGA | 1455 | 0.0 | 8.341184 | 124-125 |
| CAGAGAT | 1415 | 0.0 | 8.281219 | 124-125 |
| AGCAGAG | 1595 | 0.0 | 7.899313 | 122-123 |
| CGATCTC | 860 | 1.6249072E-4 | 7.833756 | 1 |
| TCTGCTG | 960 | 5.917283E-5 | 7.70349 | 6 |
| ATCTCTG | 1060 | 0.0016309114 | 6.4033985 | 3 |
| GAGATCG | 1210 | 1.8064329E-6 | 6.2276516 | 120-121 |
| AGAGATC | 1420 | 2.6823182E-7 | 6.189045 | 124-125 |
| GATCTCT | 1105 | 0.0026407337 | 6.134687 | 2 |
| TCAGCAG | 1165 | 1.2116915E-4 | 5.7476025 | 124-125 |
| ATCGGAA | 1440 | 8.136623E-5 | 5.2312164 | 124-125 |
| GCAGCAG | 1590 | 7.170014E-5 | 5.000911 | 124-125 |

Produced by FastQC (version 0.11.5)
