## Supplementary material 4 for "Natural and human-mediated drivers of microevolution in Neotropical palms: a historical genomics approach": outLAU_AS10.1_pf_trimmed_masked_sickled_fastqc.html

outLAU\_AS10.1\_pf\_trimmed\_masked\_sickled.fastq FastQC Report 

FastQC Report

mar. 25 juin 2019  
outLAU\_AS10.1\_pf\_trimmed\_masked\_sickled.fastq

### Summary

- Basic Statistics
- Per base sequence quality
- Per tile sequence quality
- Per sequence quality scores
- Per base sequence content
- Per sequence GC content
- Per base N content
- Sequence Length Distribution
- Sequence Duplication Levels
- Overrepresented sequences
- Adapter Content
- Kmer Content

### Basic Statistics

| Measure | Value |
| --- | --- |
| Filename | outLAU\_AS10.1\_pf\_trimmed\_masked\_sickled.fastq |
| File type | Conventional base calls |
| Encoding | Sanger / Illumina 1.9 |
| Total Sequences | 499464 |
| Sequences flagged as poor quality | 0 |
| Sequence length | 100-131 |
| %GC | 44 |

### Per base sequence quality

### Per tile sequence quality

### Per sequence quality scores

### Per base sequence content

### Per sequence GC content

### Per base N content

### Sequence Length Distribution

### Sequence Duplication Levels

### Overrepresented sequences

No overrepresented sequences

### Adapter Content

### Kmer Content

| Sequence | Count | PValue | Obs/Exp Max | Max Obs/Exp Position |
| --- | --- | --- | --- | --- |
| GCGTCCA | 80 | 0.004258721 | 20.177965 | 102-103 |
| CTGAAGT | 450 | 8.367351E-11 | 18.651209 | 5 |
| TGAAGTC | 520 | 3.45608E-11 | 17.289827 | 6 |
| CGGCCCT | 130 | 0.0031327403 | 15.412624 | 116-117 |
| TCTGAAG | 990 | 4.8854417E-8 | 9.684983 | 4 |
| CGGAAGA | 615 | 1.5253372E-5 | 7.792958 | 122-123 |
| ATCTGAA | 1115 | 2.420297E-5 | 7.5250883 | 3 |
| TCGGAAG | 655 | 3.2086387E-5 | 7.3686166 | 124-125 |
| GGACTTC | 545 | 3.5017915E-4 | 7.3298783 | 114-115 |
| GAAGAGC | 830 | 1.9336348E-6 | 7.061063 | 124-125 |
| ACTTCAG | 1150 | 3.8580765E-9 | 6.89492 | 124-125 |
| TGACTTC | 610 | 9.2210737E-4 | 6.73443 | 122-123 |
| AAGAGCA | 805 | 4.7223188E-4 | 5.9955826 | 124-125 |
| CTTCAGA | 1340 | 6.3871084E-7 | 5.620401 | 122-123 |
| GACTTCA | 1130 | 4.8709117E-5 | 5.4033146 | 120-121 |

Produced by FastQC (version 0.11.5)
