## Supplementary material 4 for "Natural and human-mediated drivers of microevolution in Neotropical palms: a historical genomics approach": outLAU_AS10.2_pf_trimmed_masked_sickled_fastqc.html

outLAU\_AS10.2\_pf\_trimmed\_masked\_sickled.fastq FastQC Report 

FastQC Report

mar. 25 juin 2019  
outLAU\_AS10.2\_pf\_trimmed\_masked\_sickled.fastq

### Summary

- Basic Statistics
- Per base sequence quality
- Per tile sequence quality
- Per sequence quality scores
- Per base sequence content
- Per sequence GC content
- Per base N content
- Sequence Length Distribution
- Sequence Duplication Levels
- Overrepresented sequences
- Adapter Content
- Kmer Content

### Basic Statistics

| Measure | Value |
| --- | --- |
| Filename | outLAU\_AS10.2\_pf\_trimmed\_masked\_sickled.fastq |
| File type | Conventional base calls |
| Encoding | Sanger / Illumina 1.9 |
| Total Sequences | 499464 |
| Sequences flagged as poor quality | 0 |
| Sequence length | 100-131 |
| %GC | 44 |

### Per base sequence quality

### Per tile sequence quality

### Per sequence quality scores

### Per base sequence content

### Per sequence GC content

### Per base N content

### Sequence Length Distribution

### Sequence Duplication Levels

### Overrepresented sequences

No overrepresented sequences

### Adapter Content

### Kmer Content

| Sequence | Count | PValue | Obs/Exp Max | Max Obs/Exp Position |
| --- | --- | --- | --- | --- |
| CTGAAGT | 400 | 1.8189894E-12 | 21.044401 | 5 |
| TGAAGTC | 400 | 4.0017767E-11 | 19.633087 | 6 |
| TAGCGGA | 80 | 0.0057961205 | 19.130533 | 56-57 |
| GTCGTGC | 100 | 0.006851782 | 18.583563 | 112-113 |
| ACGGCGA | 170 | 0.0017408531 | 13.0378895 | 90-91 |
| CGGAAGA | 505 | 6.168193E-9 | 12.265129 | 122-123 |
| TCTGAAG | 705 | 9.3532435E-9 | 11.923595 | 4 |
| GTATGTG | 215 | 0.0010260125 | 11.35311 | 50-51 |
| ATCTGAA | 760 | 2.9376679E-8 | 11.034913 | 3 |
| AAGAGCG | 395 | 8.661601E-5 | 10.651427 | 124-125 |
| TCGGAAG | 540 | 1.8623541E-6 | 10.128718 | 124-125 |
| CACATGG | 315 | 0.001664176 | 9.089802 | 62-63 |
| ACTTCAG | 1005 | 3.074092E-10 | 8.791403 | 124-125 |
| CTTCAGA | 1310 | 1.3460522E-9 | 7.249847 | 122-123 |
| CGATCTG | 1430 | 7.6926517E-7 | 6.974082 | 1 |
| TTCGCAG | 425 | 0.005862829 | 6.970332 | 38-39 |
| CATTCGC | 425 | 0.006120006 | 6.939545 | 36-37 |
| TGACTTC | 620 | 0.009097018 | 6.6600976 | 122-123 |
| GACTTCA | 1130 | 3.1108666E-6 | 6.408039 | 120-121 |
| ATCGGAA | 955 | 8.554346E-5 | 6.318573 | 120-121 |

Produced by FastQC (version 0.11.5)
