## Supplementary material 4 for "Natural and human-mediated drivers of microevolution in Neotropical palms: a historical genomics approach": outLAU_AS11.1_pf_trimmed_masked_sickled_fastqc.html

outLAU\_AS11.1\_pf\_trimmed\_masked\_sickled.fastq FastQC Report 

FastQC Report

mar. 25 juin 2019  
outLAU\_AS11.1\_pf\_trimmed\_masked\_sickled.fastq

### Summary

- Basic Statistics
- Per base sequence quality
- Per tile sequence quality
- Per sequence quality scores
- Per base sequence content
- Per sequence GC content
- Per base N content
- Sequence Length Distribution
- Sequence Duplication Levels
- Overrepresented sequences
- Adapter Content
- Kmer Content

### Basic Statistics

| Measure | Value |
| --- | --- |
| Filename | outLAU\_AS11.1\_pf\_trimmed\_masked\_sickled.fastq |
| File type | Conventional base calls |
| Encoding | Sanger / Illumina 1.9 |
| Total Sequences | 606263 |
| Sequences flagged as poor quality | 0 |
| Sequence length | 100-131 |
| %GC | 44 |

### Per base sequence quality

### Per tile sequence quality

### Per sequence quality scores

### Per base sequence content

### Per sequence GC content

### Per base N content

### Sequence Length Distribution

### Sequence Duplication Levels

### Overrepresented sequences

No overrepresented sequences

### Adapter Content

### Kmer Content

| Sequence | Count | PValue | Obs/Exp Max | Max Obs/Exp Position |
| --- | --- | --- | --- | --- |
| CGTATTC | 130 | 0.0030661353 | 15.462859 | 118-119 |
| CTGACAG | 455 | 0.0018699679 | 10.564389 | 5 |
| TGACAGA | 505 | 0.0044033625 | 9.527218 | 6 |
| TCTGACA | 585 | 0.0014276877 | 9.243064 | 4 |
| GTCAGAT | 1035 | 0.0 | 8.97109 | 124-125 |
| CCCATTA | 305 | 0.0081091765 | 8.8465 | 120-121 |
| CTGTCAG | 1060 | 0.0 | 8.658718 | 122-123 |
| TGTCAGA | 1175 | 0.0 | 8.487542 | 124-125 |
| TCTGTCA | 1175 | 1.8189894E-12 | 7.811269 | 122-123 |
| CGGAAGA | 550 | 2.677637E-4 | 7.503093 | 124-125 |
| AGTGACA | 560 | 1.09163644E-4 | 7.333132 | 90-91 |
| AGAGCAC | 535 | 0.0016420444 | 7.0706716 | 124-125 |
| TCGGAAG | 620 | 0.007687924 | 6.1013055 | 124-125 |

Produced by FastQC (version 0.11.5)
