## Supplementary material 4 for "Natural and human-mediated drivers of microevolution in Neotropical palms: a historical genomics approach": outLAU_AS11.2_pf_trimmed_masked_sickled_fastqc.html

outLAU\_AS11.2\_pf\_trimmed\_masked\_sickled.fastq FastQC Report 

FastQC Report

mar. 25 juin 2019  
outLAU\_AS11.2\_pf\_trimmed\_masked\_sickled.fastq

### Summary

- Basic Statistics
- Per base sequence quality
- Per tile sequence quality
- Per sequence quality scores
- Per base sequence content
- Per sequence GC content
- Per base N content
- Sequence Length Distribution
- Sequence Duplication Levels
- Overrepresented sequences
- Adapter Content
- Kmer Content

### Basic Statistics

| Measure | Value |
| --- | --- |
| Filename | outLAU\_AS11.2\_pf\_trimmed\_masked\_sickled.fastq |
| File type | Conventional base calls |
| Encoding | Sanger / Illumina 1.9 |
| Total Sequences | 606263 |
| Sequences flagged as poor quality | 0 |
| Sequence length | 100-131 |
| %GC | 44 |

### Per base sequence quality

### Per tile sequence quality

### Per sequence quality scores

### Per base sequence content

### Per sequence GC content

### Per base N content

### Sequence Length Distribution

### Sequence Duplication Levels

### Overrepresented sequences

No overrepresented sequences

### Adapter Content

### Kmer Content

| Sequence | Count | PValue | Obs/Exp Max | Max Obs/Exp Position |
| --- | --- | --- | --- | --- |
| AGCGTCG | 255 | 6.0992606E-4 | 12.079135 | 118-119 |
| CTGACAG | 380 | 6.9714343E-4 | 11.887322 | 5 |
| TCTGACA | 450 | 2.2588659E-4 | 11.292956 | 4 |
| TGTCAGA | 990 | 0.0 | 9.962363 | 124-125 |
| GTCTGTC | 435 | 4.4893368E-5 | 9.870048 | 120-121 |
| CTGTCAG | 985 | 0.0 | 9.754806 | 122-123 |
| TGACAGA | 525 | 9.4743184E-4 | 9.666973 | 6 |
| AAGAGCG | 385 | 0.0010042918 | 9.606565 | 124-125 |
| TCTGTCA | 1045 | 0.0 | 9.577834 | 122-123 |
| ATCTGAC | 595 | 2.809498E-4 | 9.479083 | 3 |
| GTCAGAT | 1035 | 9.094947E-12 | 9.132167 | 124-125 |
| CGGAAGA | 560 | 0.003552398 | 7.338349 | 124-125 |
| TCGGAAG | 560 | 0.003552398 | 7.338349 | 124-125 |
| TTCTGTC | 625 | 0.0017021234 | 7.046221 | 122-123 |
| TGCGCAG | 560 | 0.0062918128 | 6.2203255 | 62-63 |
| GAAGAGC | 985 | 0.004179222 | 5.423673 | 124-125 |

Produced by FastQC (version 0.11.5)
