## Supplementary material 4 for "Natural and human-mediated drivers of microevolution in Neotropical palms: a historical genomics approach": outLAU_AS12.1_pf_trimmed_masked_sickled_fastqc.html

outLAU\_AS12.1\_pf\_trimmed\_masked\_sickled.fastq FastQC Report 

FastQC Report

mar. 25 juin 2019  
outLAU\_AS12.1\_pf\_trimmed\_masked\_sickled.fastq

### Summary

- Basic Statistics
- Per base sequence quality
- Per tile sequence quality
- Per sequence quality scores
- Per base sequence content
- Per sequence GC content
- Per base N content
- Sequence Length Distribution
- Sequence Duplication Levels
- Overrepresented sequences
- Adapter Content
- Kmer Content

### Basic Statistics

| Measure | Value |
| --- | --- |
| Filename | outLAU\_AS12.1\_pf\_trimmed\_masked\_sickled.fastq |
| File type | Conventional base calls |
| Encoding | Sanger / Illumina 1.9 |
| Total Sequences | 599988 |
| Sequences flagged as poor quality | 0 |
| Sequence length | 100-131 |
| %GC | 44 |

### Per base sequence quality

### Per tile sequence quality

### Per sequence quality scores

### Per base sequence content

### Per sequence GC content

### Per base N content

### Sequence Length Distribution

### Sequence Duplication Levels

### Overrepresented sequences

No overrepresented sequences

### Adapter Content

### Kmer Content

| Sequence | Count | PValue | Obs/Exp Max | Max Obs/Exp Position |
| --- | --- | --- | --- | --- |
| ACGTTTA | 80 | 0.005007635 | 19.621296 | 64-65 |
| GCACACG | 235 | 6.4501166E-4 | 11.999437 | 124-125 |
| ACACGTC | 245 | 9.1503817E-4 | 11.509665 | 124-125 |
| CACACGT | 250 | 0.0010836762 | 11.27947 | 124-125 |
| GATCTGC | 565 | 0.0011321428 | 9.48075 | 2 |
| CTGCATT | 670 | 5.3080404E-4 | 8.8954315 | 5 |
| GGTACAT | 300 | 0.0095617715 | 8.670215 | 104-105 |
| ATCTAGA | 640 | 0.003471766 | 8.381165 | 5 |
| TCGGAAG | 635 | 1.7693837E-6 | 8.326381 | 124-125 |
| CCTTTAA | 370 | 0.007889313 | 7.6508555 | 62-63 |
| TCTGCAT | 930 | 0.0017120618 | 7.04157 | 4 |
| GCAGATC | 1110 | 4.6085188E-7 | 6.3510537 | 124-125 |
| AATGCAG | 1155 | 1.0549084E-6 | 6.064696 | 122-123 |
| CGGAAGA | 640 | 0.008270749 | 6.0583096 | 124-125 |
| GAATGCA | 1445 | 4.2755346E-8 | 5.854397 | 124-125 |
| ATGCAGA | 1550 | 3.583591E-8 | 5.648971 | 122-123 |

Produced by FastQC (version 0.11.5)
