## Supplementary material 4 for "Natural and human-mediated drivers of microevolution in Neotropical palms: a historical genomics approach": outLAU_AS12.2_pf_trimmed_masked_sickled_fastqc.html

outLAU\_AS12.2\_pf\_trimmed\_masked\_sickled.fastq FastQC Report 

FastQC Report

mar. 25 juin 2019  
outLAU\_AS12.2\_pf\_trimmed\_masked\_sickled.fastq

### Summary

- Basic Statistics
- Per base sequence quality
- Per tile sequence quality
- Per sequence quality scores
- Per base sequence content
- Per sequence GC content
- Per base N content
- Sequence Length Distribution
- Sequence Duplication Levels
- Overrepresented sequences
- Adapter Content
- Kmer Content

### Basic Statistics

| Measure | Value |
| --- | --- |
| Filename | outLAU\_AS12.2\_pf\_trimmed\_masked\_sickled.fastq |
| File type | Conventional base calls |
| Encoding | Sanger / Illumina 1.9 |
| Total Sequences | 599988 |
| Sequences flagged as poor quality | 0 |
| Sequence length | 100-131 |
| %GC | 44 |

### Per base sequence quality

### Per tile sequence quality

### Per sequence quality scores

### Per base sequence content

### Per sequence GC content

### Per base N content

### Sequence Length Distribution

### Sequence Duplication Levels

### Overrepresented sequences

No overrepresented sequences

### Adapter Content

### Kmer Content

| Sequence | Count | PValue | Obs/Exp Max | Max Obs/Exp Position |
| --- | --- | --- | --- | --- |
| TACGGCT | 80 | 0.0054043904 | 19.36405 | 68-69 |
| GGTTTAT | 260 | 0.009729629 | 12.990186 | 2 |
| TCGGAAG | 590 | 3.3105607E-10 | 12.043739 | 124-125 |
| TGCATTC | 610 | 3.167557E-5 | 10.186729 | 6 |
| CTGCATT | 530 | 0.0010233906 | 9.585863 | 5 |
| CGGAAGA | 565 | 3.7721187E-5 | 8.877634 | 124-125 |
| AAGAGCG | 365 | 0.009407248 | 8.687029 | 120-121 |
| GCAGATC | 1010 | 3.5361154E-9 | 8.277001 | 124-125 |
| GAATGCA | 1190 | 9.455107E-9 | 7.37627 | 124-125 |
| AATGCAG | 1170 | 4.0809027E-7 | 6.7878494 | 124-125 |
| ATGCAGA | 1445 | 1.8135324E-9 | 6.786842 | 122-123 |
| TGCAGAT | 1435 | 1.2832152E-6 | 5.689964 | 118-119 |
| GAAGAGC | 865 | 0.008907747 | 5.498438 | 120-121 |

Produced by FastQC (version 0.11.5)
