## Supplementary material 4 for "Natural and human-mediated drivers of microevolution in Neotropical palms: a historical genomics approach": outLAU_AS13.1_pf_trimmed_masked_sickled_fastqc.html

outLAU\_AS13.1\_pf\_trimmed\_masked\_sickled.fastq FastQC Report 

FastQC Report

mar. 25 juin 2019  
outLAU\_AS13.1\_pf\_trimmed\_masked\_sickled.fastq

### Summary

- Basic Statistics
- Per base sequence quality
- Per tile sequence quality
- Per sequence quality scores
- Per base sequence content
- Per sequence GC content
- Per base N content
- Sequence Length Distribution
- Sequence Duplication Levels
- Overrepresented sequences
- Adapter Content
- Kmer Content

### Basic Statistics

| Measure | Value |
| --- | --- |
| Filename | outLAU\_AS13.1\_pf\_trimmed\_masked\_sickled.fastq |
| File type | Conventional base calls |
| Encoding | Sanger / Illumina 1.9 |
| Total Sequences | 738667 |
| Sequences flagged as poor quality | 0 |
| Sequence length | 100-131 |
| %GC | 44 |

### Per base sequence quality

### Per tile sequence quality

### Per sequence quality scores

### Per base sequence content

### Per sequence GC content

### Per base N content

### Sequence Length Distribution

### Sequence Duplication Levels

### Overrepresented sequences

No overrepresented sequences

### Adapter Content

### Kmer Content

| Sequence | Count | PValue | Obs/Exp Max | Max Obs/Exp Position |
| --- | --- | --- | --- | --- |
| ACACGTC | 275 | 0.0022389127 | 10.339683 | 122-123 |
| GCACACG | 310 | 5.9681287E-4 | 10.167935 | 120-121 |
| ATCTGCC | 565 | 0.001231349 | 9.394618 | 3 |
| AGCACAC | 545 | 1.5720434E-6 | 9.22693 | 124-125 |
| AACGGAT | 270 | 0.006347104 | 9.114979 | 48-49 |
| GATCTGC | 590 | 0.0018308322 | 8.994663 | 2 |
| GAGCACA | 810 | 9.695214E-10 | 8.868918 | 124-125 |
| GGTACCA | 285 | 0.008779404 | 8.761456 | 82-83 |
| TGCCTTG | 680 | 6.6023576E-4 | 8.703408 | 6 |
| TCGGAAG | 785 | 9.924406E-9 | 8.384056 | 118-119 |
| CTGTCGA | 355 | 0.0039480026 | 8.264186 | 102-103 |
| CTGCCTT | 645 | 0.003984827 | 8.25466 | 5 |
| AGAGCAC | 790 | 5.3174517E-8 | 8.184104 | 124-125 |
| TTAGCCT | 340 | 0.005147346 | 8.024364 | 36-37 |
| TCTGCCT | 700 | 0.008421759 | 7.5944214 | 4 |
| AGGCAGA | 1140 | 1.36788E-9 | 7.24684 | 124-125 |
| CGGAAGA | 700 | 6.267916E-5 | 7.0045776 | 120-121 |
| GGCAGAT | 1010 | 5.3178337E-7 | 6.686242 | 122-123 |
| AAGAGCA | 1165 | 1.3997669E-7 | 6.406831 | 122-123 |
| TCAAGGC | 770 | 2.174592E-4 | 6.3677974 | 120-121 |

Produced by FastQC (version 0.11.5)
