## Supplementary material 4 for "Natural and human-mediated drivers of microevolution in Neotropical palms: a historical genomics approach": outLAU_AS13.2_pf_trimmed_masked_sickled_fastqc.html

outLAU\_AS13.2\_pf\_trimmed\_masked\_sickled.fastq FastQC Report 

FastQC Report

mar. 25 juin 2019  
outLAU\_AS13.2\_pf\_trimmed\_masked\_sickled.fastq

### Summary

- Basic Statistics
- Per base sequence quality
- Per tile sequence quality
- Per sequence quality scores
- Per base sequence content
- Per sequence GC content
- Per base N content
- Sequence Length Distribution
- Sequence Duplication Levels
- Overrepresented sequences
- Adapter Content
- Kmer Content

### Basic Statistics

| Measure | Value |
| --- | --- |
| Filename | outLAU\_AS13.2\_pf\_trimmed\_masked\_sickled.fastq |
| File type | Conventional base calls |
| Encoding | Sanger / Illumina 1.9 |
| Total Sequences | 738667 |
| Sequences flagged as poor quality | 0 |
| Sequence length | 100-131 |
| %GC | 44 |

### Per base sequence quality

### Per tile sequence quality

### Per sequence quality scores

### Per base sequence content

### Per sequence GC content

### Per base N content

### Sequence Length Distribution

### Sequence Duplication Levels

### Overrepresented sequences

No overrepresented sequences

### Adapter Content

### Kmer Content

| Sequence | Count | PValue | Obs/Exp Max | Max Obs/Exp Position |
| --- | --- | --- | --- | --- |
| AAGAGCG | 530 | 3.6379788E-12 | 14.081683 | 122-123 |
| GAGCGTC | 380 | 3.9018487E-6 | 12.30437 | 124-125 |
| CGGAAGA | 585 | 8.767529E-9 | 10.821641 | 118-119 |
| TCGGAAG | 670 | 5.966285E-10 | 10.629839 | 118-119 |
| AGAGCGT | 375 | 0.006619155 | 9.067947 | 124-125 |
| GCAAGGC | 545 | 0.0019562899 | 7.7992673 | 124-125 |
| AGGCAGA | 1065 | 4.730955E-7 | 7.184113 | 124-125 |
| AAGGCAG | 1120 | 1.0901222E-6 | 6.831322 | 124-125 |
| GGCAGAT | 1005 | 1.7872735E-6 | 6.6297727 | 112-113 |
| GCAGATC | 1205 | 3.6245892E-6 | 6.349445 | 124-125 |
| CAAGGCA | 1250 | 9.848343E-6 | 5.970634 | 122-123 |
| GGAAGAG | 1270 | 1.993416E-5 | 5.7148333 | 120-121 |

Produced by FastQC (version 0.11.5)
