## Supplementary material 4 for "Natural and human-mediated drivers of microevolution in Neotropical palms: a historical genomics approach": outLAU_AS14.1_pf_trimmed_masked_sickled_fastqc.html

outLAU\_AS14.1\_pf\_trimmed\_masked\_sickled.fastq FastQC Report 

FastQC Report

mar. 25 juin 2019  
outLAU\_AS14.1\_pf\_trimmed\_masked\_sickled.fastq

### Summary

- Basic Statistics
- Per base sequence quality
- Per tile sequence quality
- Per sequence quality scores
- Per base sequence content
- Per sequence GC content
- Per base N content
- Sequence Length Distribution
- Sequence Duplication Levels
- Overrepresented sequences
- Adapter Content
- Kmer Content

### Basic Statistics

| Measure | Value |
| --- | --- |
| Filename | outLAU\_AS14.1\_pf\_trimmed\_masked\_sickled.fastq |
| File type | Conventional base calls |
| Encoding | Sanger / Illumina 1.9 |
| Total Sequences | 635776 |
| Sequences flagged as poor quality | 0 |
| Sequence length | 100-131 |
| %GC | 44 |

### Per base sequence quality

### Per tile sequence quality

### Per sequence quality scores

### Per base sequence content

### Per sequence GC content

### Per base N content

### Sequence Length Distribution

### Sequence Duplication Levels

### Overrepresented sequences

No overrepresented sequences

### Adapter Content

### Kmer Content

| Sequence | Count | PValue | Obs/Exp Max | Max Obs/Exp Position |
| --- | --- | --- | --- | --- |
| TGCGCTA | 170 | 0.0 | 41.211777 | 6 |
| TCTGCGC | 205 | 0.0 | 36.956867 | 4 |
| CTGCGCT | 195 | 1.8189894E-12 | 35.910534 | 5 |
| GCGCTAG | 90 | 2.6498982E-4 | 32.483177 | 7 |
| ATCTGCG | 280 | 1.2369128E-10 | 24.945692 | 3 |
| ATAGCGC | 180 | 9.230046E-4 | 14.182105 | 122-123 |
| GCACACG | 190 | 0.0012126187 | 13.678983 | 124-125 |
| GATCTGC | 665 | 3.2448952E-8 | 12.228014 | 2 |
| AGCGCAG | 625 | 5.366019E-10 | 10.693056 | 124-125 |
| TTAGCGC | 280 | 0.0018083896 | 10.608191 | 124-125 |
| GCGCAGA | 590 | 2.8981594E-7 | 9.439491 | 124-125 |
| TAGCGCA | 580 | 2.9345028E-7 | 9.43145 | 122-123 |
| CGCAGAT | 565 | 2.3871326E-5 | 8.284763 | 120-121 |
| GAAAAGC | 500 | 0.0042872005 | 6.4539456 | 12-13 |
| GCAGATC | 865 | 0.0029888938 | 5.580031 | 124-125 |
| CGATCTG | 1685 | 2.3817086E-4 | 5.481742 | 1 |

Produced by FastQC (version 0.11.5)
