## Supplementary material 4 for "Natural and human-mediated drivers of microevolution in Neotropical palms: a historical genomics approach": outLAU_AS14.2_pf_trimmed_masked_sickled_fastqc.html

outLAU\_AS14.2\_pf\_trimmed\_masked\_sickled.fastq FastQC Report 

FastQC Report

mar. 25 juin 2019  
outLAU\_AS14.2\_pf\_trimmed\_masked\_sickled.fastq

### Summary

- Basic Statistics
- Per base sequence quality
- Per tile sequence quality
- Per sequence quality scores
- Per base sequence content
- Per sequence GC content
- Per base N content
- Sequence Length Distribution
- Sequence Duplication Levels
- Overrepresented sequences
- Adapter Content
- Kmer Content

### Basic Statistics

| Measure | Value |
| --- | --- |
| Filename | outLAU\_AS14.2\_pf\_trimmed\_masked\_sickled.fastq |
| File type | Conventional base calls |
| Encoding | Sanger / Illumina 1.9 |
| Total Sequences | 635776 |
| Sequences flagged as poor quality | 0 |
| Sequence length | 100-131 |
| %GC | 44 |

### Per base sequence quality

### Per tile sequence quality

### Per sequence quality scores

### Per base sequence content

### Per sequence GC content

### Per base N content

### Sequence Length Distribution

### Sequence Duplication Levels

### Overrepresented sequences

No overrepresented sequences

### Adapter Content

### Kmer Content

| Sequence | Count | PValue | Obs/Exp Max | Max Obs/Exp Position |
| --- | --- | --- | --- | --- |
| TGCGCTA | 135 | 6.002665E-11 | 40.954597 | 6 |
| CTGCGCT | 190 | 2.415618E-9 | 29.031029 | 5 |
| TCTGCGC | 205 | 5.5661076E-9 | 26.846054 | 4 |
| ATCTGCG | 240 | 3.0735464E-8 | 22.860271 | 3 |
| ATAGCGC | 210 | 8.924976E-4 | 14.244621 | 122-123 |
| TTAGCGC | 195 | 0.008984012 | 13.148881 | 122-123 |
| AGCGCAG | 520 | 4.703179E-8 | 11.907336 | 124-125 |
| TAGCGCA | 525 | 8.653842E-8 | 11.395697 | 122-123 |
| AAGAGCG | 340 | 2.2259106E-4 | 11.311905 | 122-123 |
| GAGCGTC | 275 | 0.00522597 | 11.257845 | 124-125 |
| GATCTGC | 500 | 6.656506E-5 | 10.93101 | 2 |
| GCGCAGA | 475 | 3.700532E-6 | 10.795923 | 122-123 |
| AGAGCGT | 280 | 0.007714931 | 10.683466 | 122-123 |
| CGCAGAT | 515 | 1.3486988E-6 | 10.387863 | 120-121 |
| CGGAAGA | 465 | 6.9138815E-4 | 8.663517 | 118-119 |
| TCGGAAG | 475 | 0.0034787715 | 8.3799 | 124-125 |
| GCAGATC | 870 | 1.193117E-6 | 7.859102 | 122-123 |
| ATGCACA | 515 | 0.009783899 | 7.4680543 | 122-123 |

Produced by FastQC (version 0.11.5)
