## Supplementary material 6 for "Natural and human-mediated drivers of microevolution in Neotropical palms: a historical genomics approach"

**Supplementary Material 6 (S6)**

**Population genetics – genome-wide processes**

This supplementary material provides additional information regarding the genome-wide genetic structure across populations:

- Genetic differentiation ($F_{ST}$, Weir and Cockerham 1984): Tables S6.1 & S6.2
- Individual ancestry: Figures S6.1 & S6.2
- Isolation-by-distance: Figure S6.3
- $f_3$ admixture tests: Figure S6.4
- Intra-population inbreeding coefficients ($F_{IS}$): Figure S6.5
- BayPass sanity check: Figure S6.6 to S6.9
- Genome-wide structure across populations estimated by BayPass: Figure S6.10
- Detection of climate adaptation and domestication SNPs: Figure S6.11
- Shared outlier SNPs: Tables S6.3 to S6.6

**Table S6.1.** Weir & Cockerham's $F_{ST}$ in *Astrocaryum spp*. “AP” refers to *A. paramaca* and “AS” to *A. sciophilum*.

| | **AP SPA -** | **AP SPA +** | **AP LAU -** | **AP PAR -** | **AP NOU_N -** | **AP NOU_N +** | **AP NOU_S -** | **AP NOU_S +** | **AP MC87 -** | **AP MC87 +** | **AS SPA -** | **AS SPA +** | **AS LAU -** | **AS PAR -** | **AS NOU_N -** | **AS NOU_N +** | **AS MC87 -** | **AS MC87 +** |
|---|---|---|---|---|---|---|---|---|---|---|---|---|---|---|---|---|---|---|
| **AP SPA-** | 0 | 0.007 | 0.015 | 0.018 | 0.035 | 0.046 | 0.048 | 0.043 | 0.403 | 0.086 | 0.534 | 0.541 | 0.544 | 0.543 | 0.54 | 0.548 | 0.39 | 0.486 |
| **AP_SPA+** | 0.007 | 0 | 0.021 | 0.024 | 0.038 | 0.044 | 0.044 | 0.042 | 0.397 | 0.079 | 0.525 | 0.533 | 0.537 | 0.535 | 0.531 | 0.54 | 0.381 | 0.477 |
| **AP LAU-** | 0.015 | 0.021 | 0 | 0.018 | 0.034 | 0.05 | 0.047 | 0.042 | 0.398 | 0.083 | 0.53 | 0.538 | 0.542 | 0.54 | 0.536 | 0.545 | 0.383 | 0.482 |
| **AP_PAR-** | 0.018 | 0.024 | 0.018 | 0 | 0.025 | 0.038 | 0.037 | 0.035 | 0.407 | 0.074 | 0.537 | 0.544 | 0.548 | 0.546 | 0.543 | 0.551 | 0.388 | 0.487 |
| **AP_NOU_N-** | 0.035 | 0.038 | 0.034 | 0.025 | 0 | 0.005 | 0.004 | 0.006 | 0.387 | 0.043 | 0.511 | 0.521 | 0.523 | 0.523 | 0.519 | 0.528 | 0.38 | 0.47 |
| **AP NOU_N+** | 0.046 | 0.044 | 0.05 | 0.038 | 0.005 | 0 | 0.008 | 0.01 | 0.394 | 0.039 | 0.513 | 0.525 | 0.527 | 0.527 | 0.523 | 0.532 | 0.382 | 0.474 |
| **AP_NOU_S-** | 0.048 | 0.044 | 0.047 | 0.037 | 0.004 | 0.008 | 0 | 0.003 | 0.396 | 0.042 | 0.524 | 0.534 | 0.537 | 0.536 | 0.532 | 0.541 | 0.383 | 0.479 |
| **AP NOU_S+** | 0.043 | 0.042 | 0.042 | 0.035 | 0.006 | 0.01 | 0.003 | 0 | 0.394 | 0.038 | 0.525 | 0.532 | 0.537 | 0.535 | 0.531 | 0.54 | 0.385 | 0.479 |
| **AP_MC87-** | 0.403 | 0.397 | 0.398 | 0.407 | 0.387 | 0.394 | 0.396 | 0.394 | 0 | 0.271 | 0.541 | 0.548 | 0.55 | 0.551 | 0.547 | 0.553 | 0.296 | 0.467 |
| **AP_MC87+** | 0.086 | 0.079 | 0.083 | 0.074 | 0.043 | 0.039 | 0.042 | 0.038 | 0.271 | 0 | 0.471 | 0.491 | 0.491 | 0.494 | 0.489 | 0.499 | 0.313 | 0.431 |
| **AS_SPA-** | 0.534 | 0.525 | 0.53 | 0.537 | 0.511 | 0.513 | 0.524 | 0.525 | 0.541 | 0.471 | 0 | 0.011 | 0.052 | 0.039 | 0.039 | 0.051 | 0.075 | 0.048 |
| **AS SPA+** | 0.541 | 0.533 | 0.538 | 0.544 | 0.521 | 0.525 | 0.534 | 0.532 | 0.548 | 0.491 | 0.011 | 0 | 0.051 | 0.047 | 0.039 | 0.055 | 0.103 | 0.057 |
| **AS_LAU-** | 0.544 | 0.537 | 0.542 | 0.548 | 0.523 | 0.527 | 0.537 | 0.537 | 0.55 | 0.491 | 0.052 | 0.051 | 0 | 0.026 | 0.05 | 0.055 | 0.099 | 0.059 |
| **AS_PAR-** | 0.543 | 0.535 | 0.54 | 0.546 | 0.523 | 0.527 | 0.536 | 0.535 | 0.551 | 0.494 | 0.039 | 0.047 | 0.026 | 0 | 0.038 | 0.05 | 0.107 | 0.059 |
| **AS NOU_N-** | 0.54 | 0.531 | 0.536 | 0.543 | 0.519 | 0.523 | 0.532 | 0.531 | 0.547 | 0.489 | 0.039 | 0.039 | 0.05 | 0.038 | 0 | 0.011 | 0.074 | 0.017 |
| **AS_NOU_N+** | 0.548 | 0.54 | 0.545 | 0.551 | 0.528 | 0.532 | 0.541 | 0.54 | 0.553 | 0.499 | 0.051 | 0.055 | 0.055 | 0.05 | 0.011 | 0 | 0.083 | 0.018 |
| **AS MC87-** | 0.39 | 0.381 | 0.383 | 0.388 | 0.38 | 0.382 | 0.383 | 0.385 | 0.296 | 0.313 | 0.075 | 0.103 | 0.099 | 0.107 | 0.074 | 0.083 | 0 | 0.014 |
| **AS_MC87+** | 0.486 | 0.477 | 0.482 | 0.487 | 0.47 | 0.474 | 0.479 | 0.479 | 0.467 | 0.431 | 0.048 | 0.057 | 0.059 | 0.059 | 0.017 | 0.018 | 0.014 | 0 |

**Table S6.2.** Weir & Cockerham's $F_{ST}$ in *Oenocarpus spp*. “OBc” refers to *O. bacaba* and “OBt” to *O. bataua*.

| | OBc SPA - | OBc SPA + | OBc LAU - | OBc PAR - | OBc NOU_N + | OBc NOU_S + | OBt SPA - | OBt SPA + | OBt LAU - | OBt PAR - | OBt NOU_N - | OBt NOU_N + | OBt NOU_S + | OBt MC87 - | OBt MC87 + |
|---|---|---|---|---|---|---|---|---|---|---|---|---|---|---|---|
| **OBc SPA-** | 0 | 0.005 | 0.061 | 0.047 | 0.014 | 0.015 | 0.295 | 0.296 | 0.298 | 0.294 | 0.291 | 0.285 | 0.282 | 0.292 | 0.288 |
| **OBc SPA+** | 0.005 | 0 | 0.042 | 0.028 | 0.019 | 0.017 | 0.241 | 0.244 | 0.238 | 0.235 | 0.237 | 0.231 | 0.228 | 0.238 | 0.233 |
| **OBc LAU-** | 0.061 | 0.042 | 0 | -0.005 | 0.064 | 0.06 | 0.254 | 0.257 | 0.25 | 0.248 | 0.249 | 0.242 | 0.24 | 0.25 | 0.243 |
| **OBc PAR-** | 0.047 | 0.028 | -0.005 | 0 | 0.052 | 0.048 | 0.248 | 0.253 | 0.241 | 0.238 | 0.243 | 0.238 | 0.233 | 0.245 | 0.237 |
| **OBc NOU_N+** | 0.014 | 0.019 | 0.064 | 0.052 | 0 | 0.002 | 0.294 | 0.295 | 0.297 | 0.294 | 0.29 | 0.284 | 0.283 | 0.29 | 0.287 |
| **OBc NOU_S+** | 0.015 | 0.017 | 0.06 | 0.048 | 0.002 | 0 | 0.293 | 0.294 | 0.295 | 0.293 | 0.288 | 0.283 | 0.281 | 0.288 | 0.285 |
| **OBt SPA-** | 0.295 | 0.241 | 0.254 | 0.248 | 0.294 | 0.293 | 0 | 0 | 0.009 | 0.002 | 0.006 | 0.009 | 0.011 | 0.008 | 0.011 |
| **OBt SPA+** | 0.296 | 0.244 | 0.257 | 0.253 | 0.295 | 0.294 | 0 | 0 | 0.01 | 0.005 | 0.008 | 0.011 | 0.014 | 0.01 | 0.013 |
| **OBt LAU-** | 0.298 | 0.238 | 0.25 | 0.241 | 0.297 | 0.295 | 0.009 | 0.01 | 0 | 0.003 | 0.011 | 0.008 | 0.013 | 0.013 | 0.011 |
| **OBt PAR-** | 0.294 | 0.235 | 0.248 | 0.238 | 0.294 | 0.293 | 0.002 | 0.005 | 0.003 | 0 | 0.008 | 0.006 | 0.011 | 0.01 | 0.011 |
| **OBt NOU_N-** | 0.291 | 0.237 | 0.249 | 0.243 | 0.29 | 0.288 | 0.006 | 0.008 | 0.011 | 0.008 | 0 | -0.001 | 0.005 | 0.003 | 0.002 |
| **OBt NOU_N+** | 0.285 | 0.231 | 0.242 | 0.238 | 0.284 | 0.283 | 0.009 | 0.011 | 0.008 | 0.006 | -0.001 | 0 | 0.004 | 0.005 | 0.003 |
| **OBt NOU_S+** | 0.282 | 0.228 | 0.24 | 0.233 | 0.283 | 0.281 | 0.011 | 0.014 | 0.013 | 0.011 | 0.005 | 0.004 | 0 | 0.009 | 0.007 |
| **OBt MC87-** | 0.292 | 0.238 | 0.25 | 0.245 | 0.29 | 0.288 | 0.008 | 0.01 | 0.013 | 0.01 | 0.003 | 0.005 | 0.009 | 0 | 0 |
| **OBt MC87+** | 0.288 | 0.233 | 0.243 | 0.237 | 0.287 | 0.285 | 0.011 | 0.013 | 0.011 | 0.011 | 0.002 | 0.003 | 0.007 | 0 | 0 |

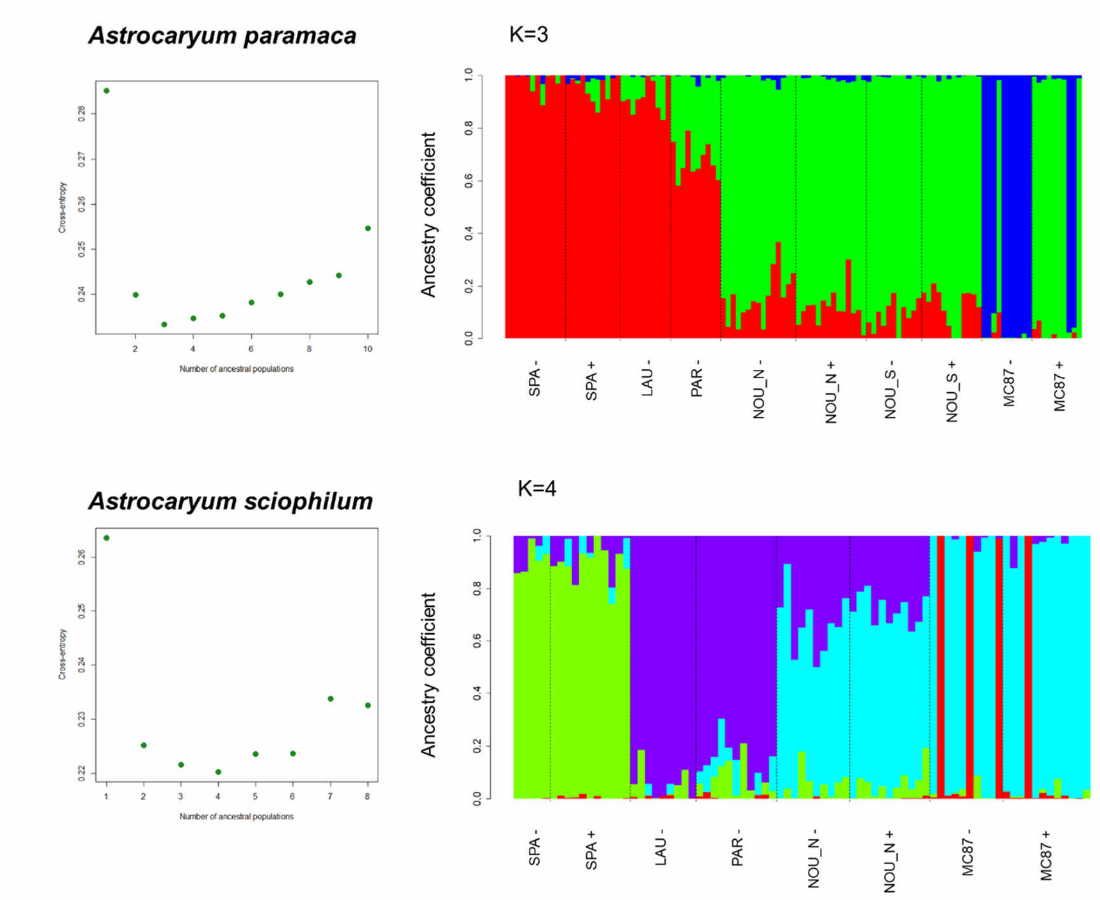


**Figure S6.1.** Value of cross-entropy criterion and individual ancestry coefficients computed with sNMF for *Astrocaryum paramaca* (upper panel, K=3) and *Astrocaryum sciophilum* (lower panel, K=4), respectively.

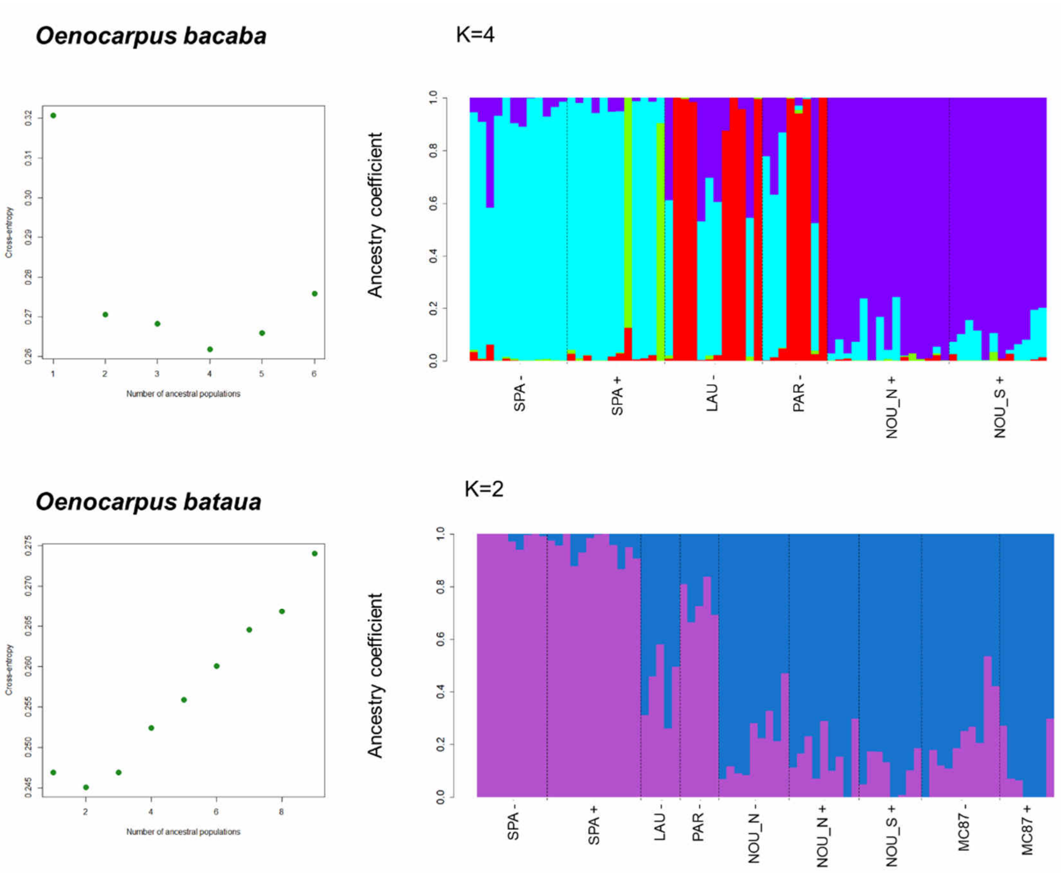


**Figure S6.2.** Value of cross-entropy criterion and individual ancestry coefficients computed with sNMF for *Oenocarpus bacaba* (upper panel, K=4) and *Astrocaryum bataua* (lower panel, K=2), respectively.

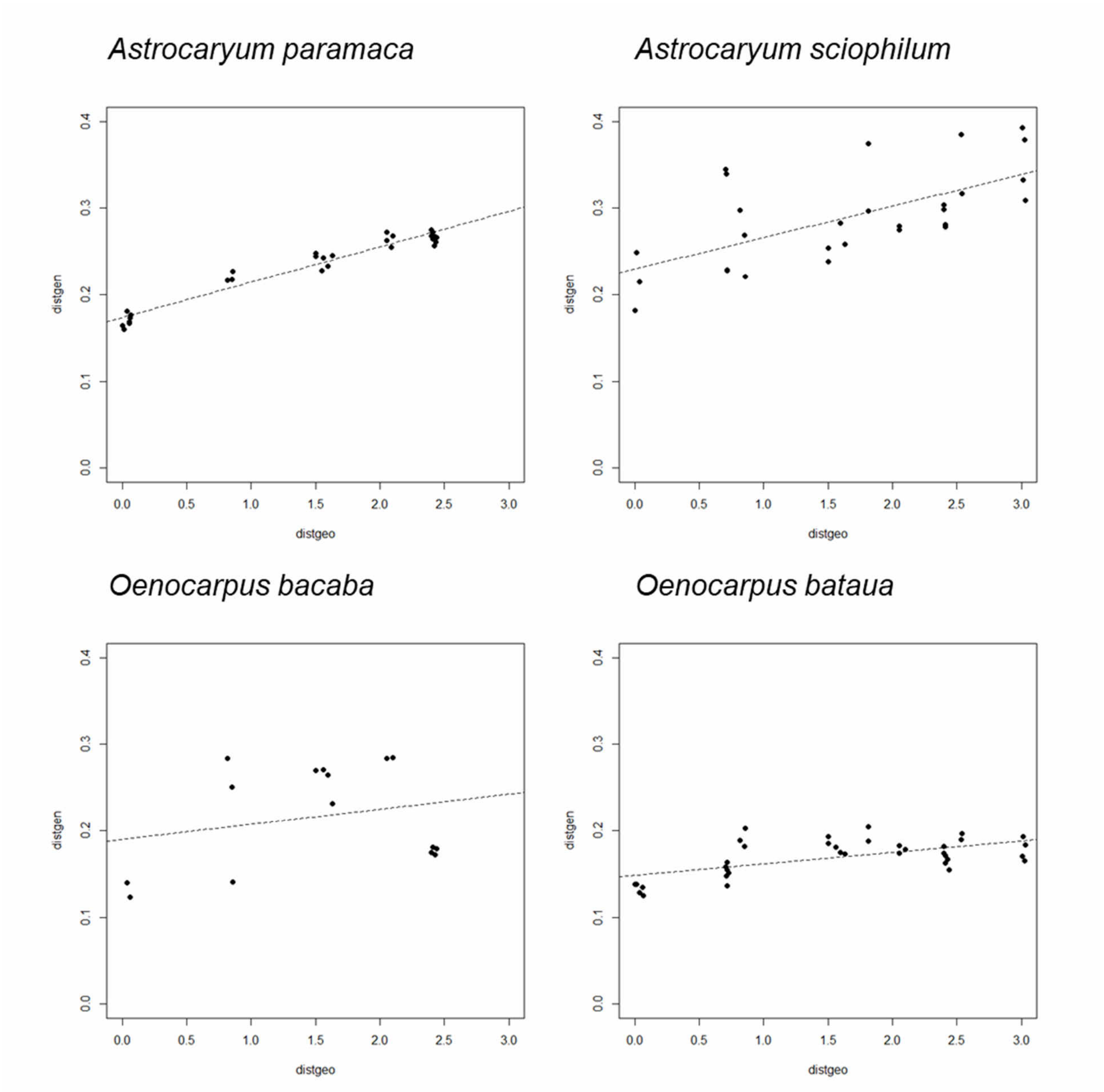


**Figure S6.3.** Relationship between geographic and genetic distances illustrating patterns of isolation-by-distance in *A. paramaca* (excluding populations from MC87), *A. sciophylum* and *O. bataua*. Mantel test was not significant in *O. bacaba*.

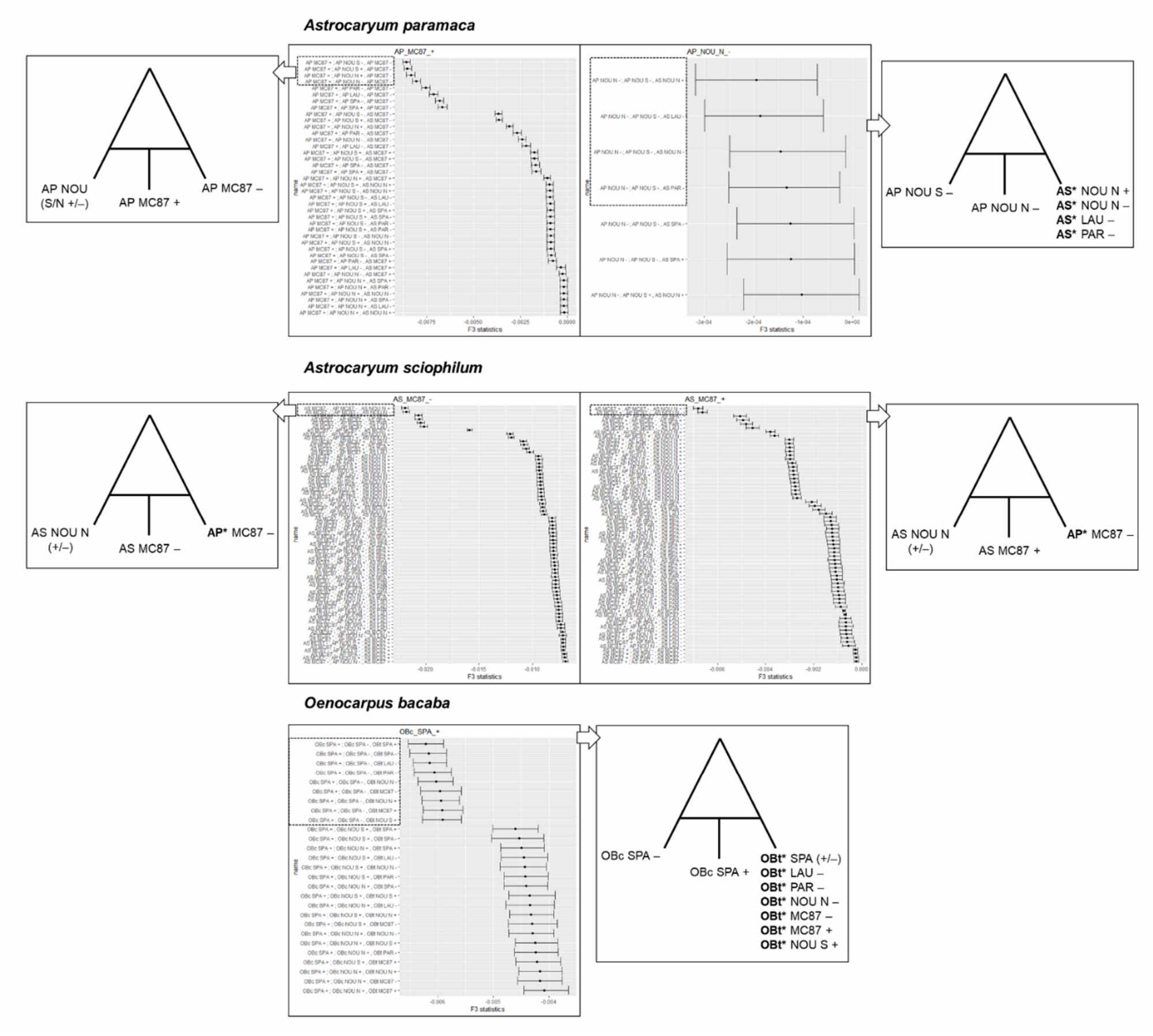


**Figure S6.4.** Significantly negative $f_3$ statistics (p-value <0.05) with 1% and 99% CI. "AP" refers to *A. paramaca*, "AS" to *A. sciophilum*. "OBc" to *O. bacaba* and "OBt" to *O. bataua*.

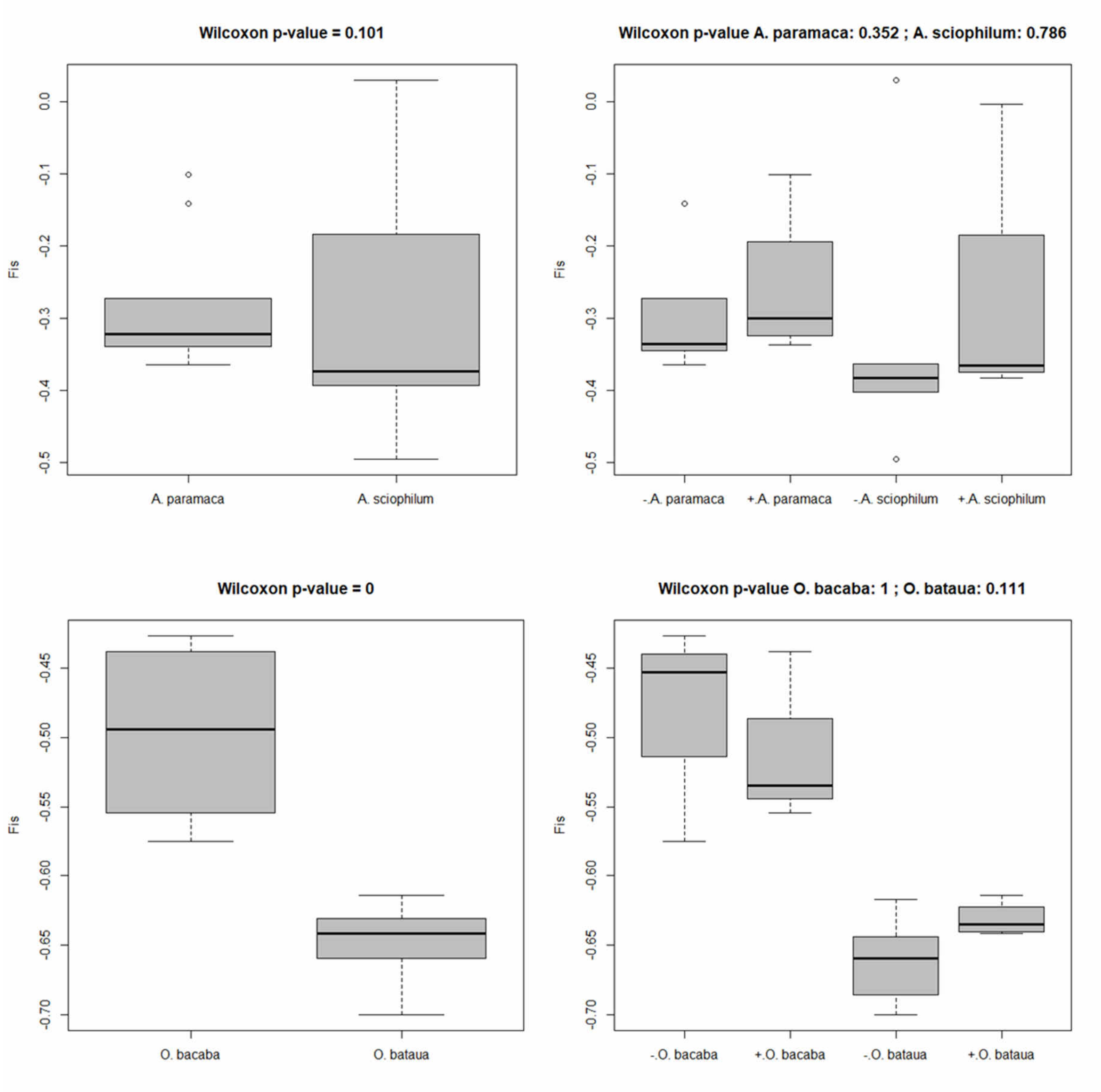


**Figure S6.5.** Average inbreeding coefficients (*Fis*) across markers and populations in the different species (left) and local conditions of pre-Columbian occupation (right).

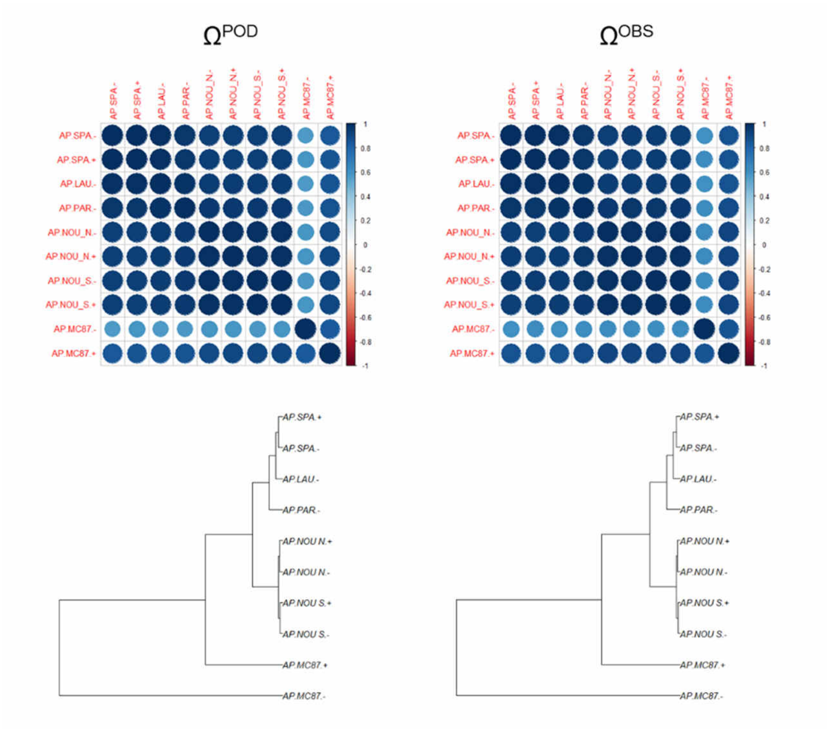


**Figure S6.6.** BayPass sanity check in *A. paramaca*. Comparison of estimated population covariance between simulated ($\Omega^{POD}$, left) and observed data ($\Omega^{OBS}$, right). $\overline{FMD}$ = 1.06. Correlogram and trees were realized with the functions `corrplot` and `hclust`.

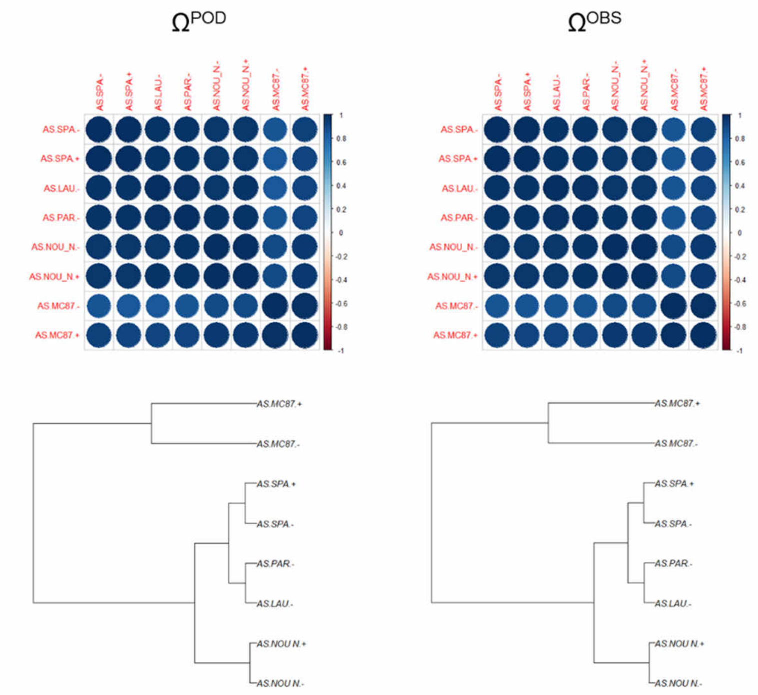


**Figure S6.7.** BayPass sanity check in *A. sciophilum*. $\overline{FMD}$ = 1.25.

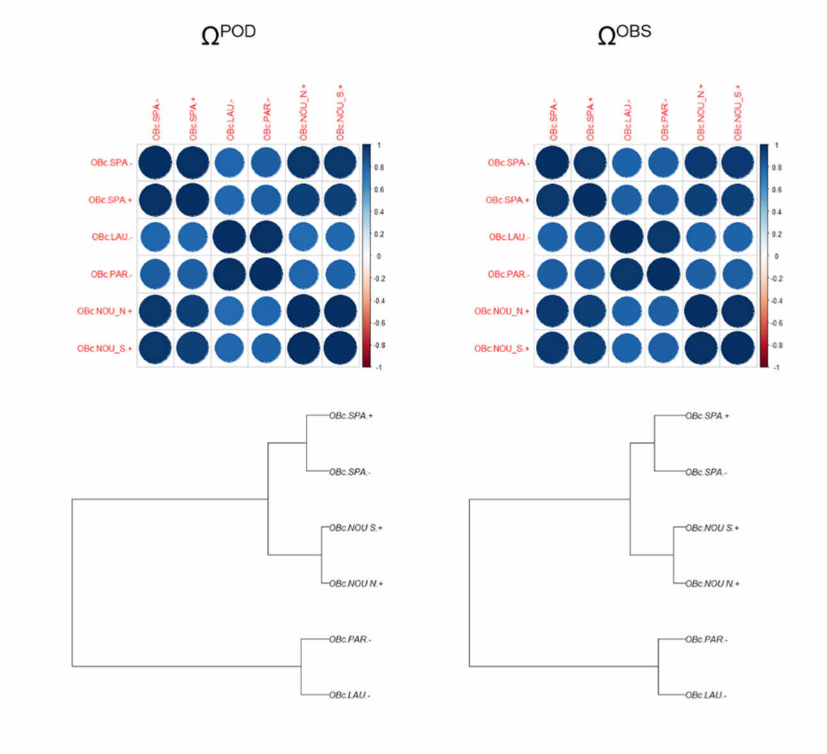


**Figure S6.8.** BayPass sanity check in *O. bacaba*. $\overline{FMD}$ = 2.07.

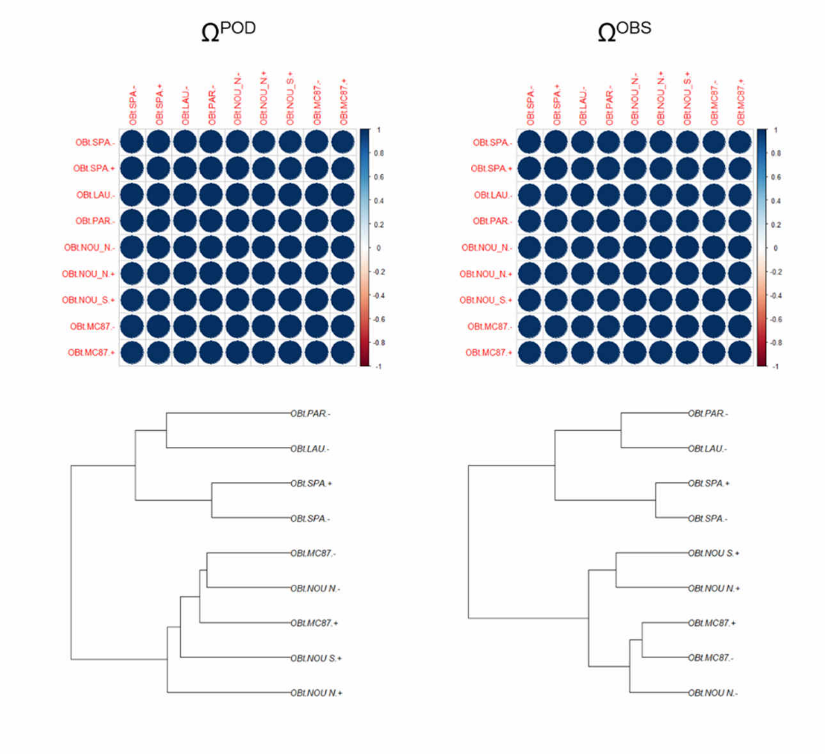


**Figure S6.9.** BayPass sanity check in *O. bataua*. $\overline{FMD}$ = 2.90.

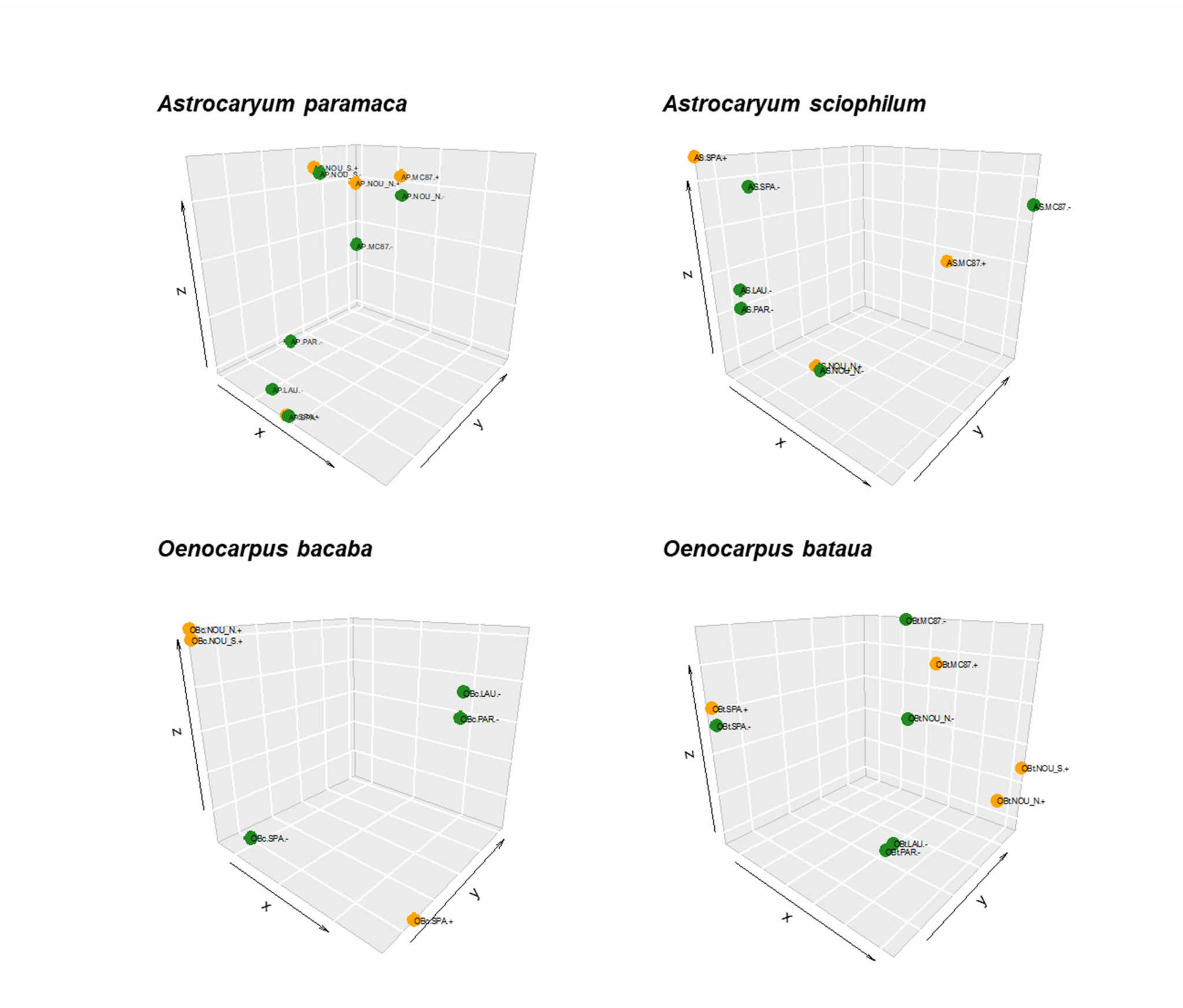


**Figure S6.10.** Genome-wide structure across populations (from covariance matrices, $\Omega^{OBS}$, estimated by BayPass). Yellow: population on ring ditches (+), green: populations in natural forest areas (–).

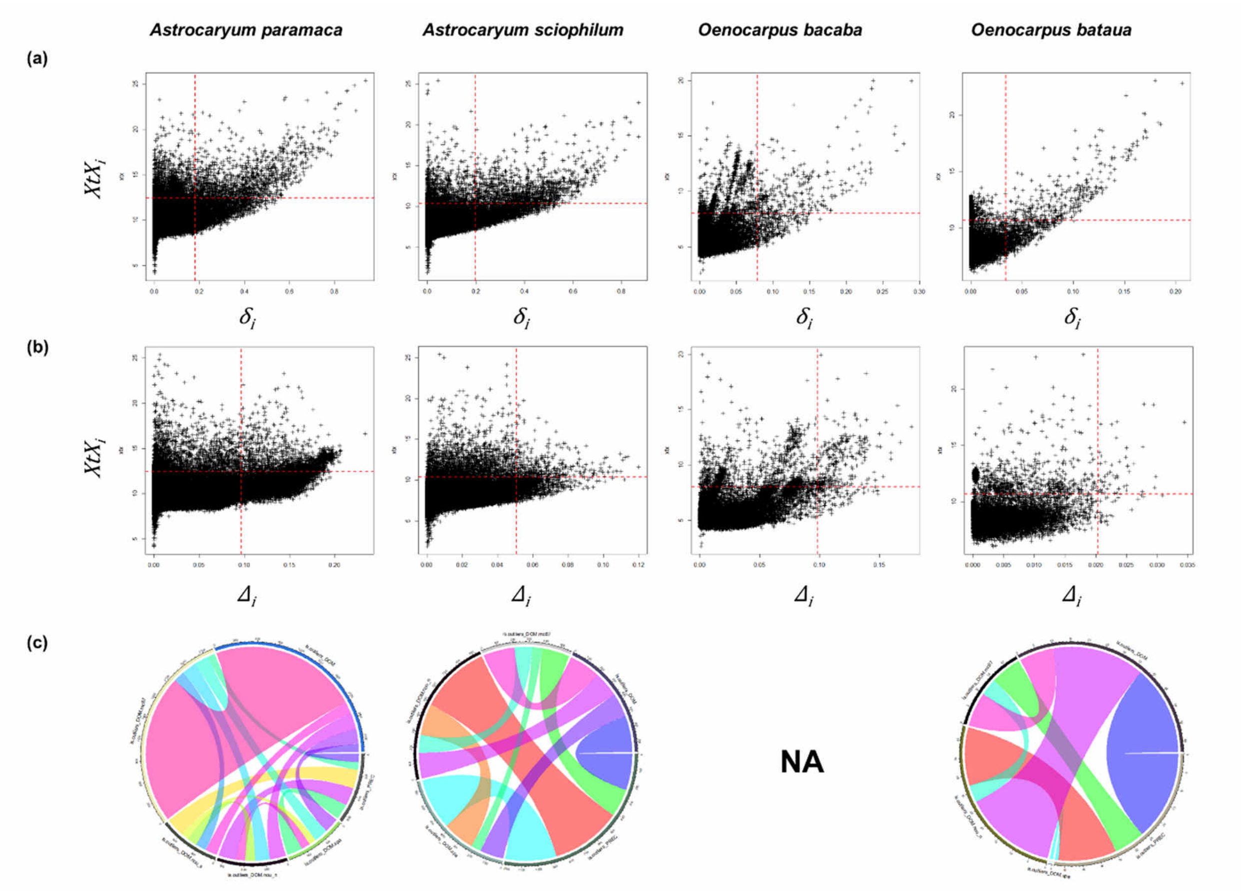


**Figure S6.11.** (a) Detection of climate adaptation SNPs (across all populations and study sites) in the four species based on the joint distribution of *XtX* and *δ*. (b) Detection of and domestication SNPs (across all populations and study sites) based on the joint distribution of *XtX* and *Δ*. (c) Proportion of shared outliers (see also Tables S5.3 to S5.6).

**Table S6.3.** Number of shared outliers (out of 178,358 SNPs) in *A. paramaca*.

| | PREC | DOM (SPA) | DOM (NOU_N) | DOM (NOU_S) | DOM (MC87) | DOM (multi-site) |
|---|---|---|---|---|---|---|
| PREC | | 223 | 262 | 273 | 108 | 120 |
| DOM (SPA) | | | 128 | 103 | 205 | 213 |
| DOM (NOU_N) | | | | 144 | 213 | 235 |
| DOM (NOU_S) | | | | | 160 | 173 |
| DOM (MC87) | | | | | | 2158 |
| DOM (multi-sites) | | | | | | |

**Table S6.4.** Number of shared outliers (out of 178,358 SNPs) in *A. sciophilum*.

| | PREC | DOM (SPA) | DOM (NOU_N) | DOM (MC87) | DOM (multi-sites) |
|---|---|---|---|---|---|
| PREC | | 334 | 459 | 181 | 236 |

| | | | | | |
|---|---|---|---|---|---|
| DOM (SPA) | | | 205 | 66 | 155 |
| DOM (NOU_N) | | | | 116 | 167 |
| DOM (MC87) | | | | | 205 |
| DOM (multi-sites) | | | | | |

**Table S6.5.** Number of shared outliers (out of 23,252 SNPs) in *O. bacaba*.

| | PREC | DOM (SPA) | DOM (NOU_N) | DOM (MC87) | DOM (multi-sites) |
|---|---|---|---|---|---|
| PREC | | 46 | NA | NA | 46 |
| DOM (SPA) | | | NA | NA | 284 |
| DOM (NOU_N) | | | | NA | NA |
| DOM (MC87) | | | | | NA |
| DOM (multi-sites) | | | | | |

**Table S6.6.** Number of shared outliers (out of 23,252 SNPs) in *O. bacaba*.

| | PREC | DOM (SPA) | DOM (NOU_N) | DOM (MC87) | DOM (multi-sites) |
|---|---|---|---|---|---|
| PREC | | 1 | 17 | 9 | 26 |
| DOM (SPA) | | | 0 | 0 | 0 |
| DOM (NOU_N) | | | | 5 | 27 |
| DOM (MC87) | | | | | 10 |
| DOM (multi-sites) | | | | | |
